# Multiplexed Label-Free Biomarker Detection by Targeted Disassembly of Variable-Length DNA Payload Chains

**DOI:** 10.1101/2022.03.25.485867

**Authors:** Matthew Aquilina, Katherine E. Dunn

## Abstract

Simultaneously studying different types of biomarkers (DNA, RNA, proteins, metabolites) has the potential to significantly improve understanding and diagnosis for many complex diseases. However, extracting biomarkers of different types involves using several technically complex or expensive methodologies, often requiring specialized laboratories and personnel. Streamlining detection through the use of a single multiplexed assay would greatly facilitate the process of accessing and interpreting patient biomarker data. In this work, we present a method for multiplexed biomarker detection based on variable-length DNA payload chains, which are systematically disassembled in the presence of specific biomolecular targets, leading to fragments of different sizes that yield characteristic band patterns in gel electrophoresis. This strategy has enabled us to detect with high sensitivity and specificity DNA sequences including BRCA1, an RNA sequence (miR-141) and the steroid aldosterone. We show that our assay can be multiplexed, enabling simultaneous detection of different types of biomarker. Furthermore, we show that our method suffers no loss of sensitivity when conducted in fetal bovine serum and can be applied using capillary electrophoresis, which may be more amenable to automation and integration in healthcare settings.

**ToC Graphic:** 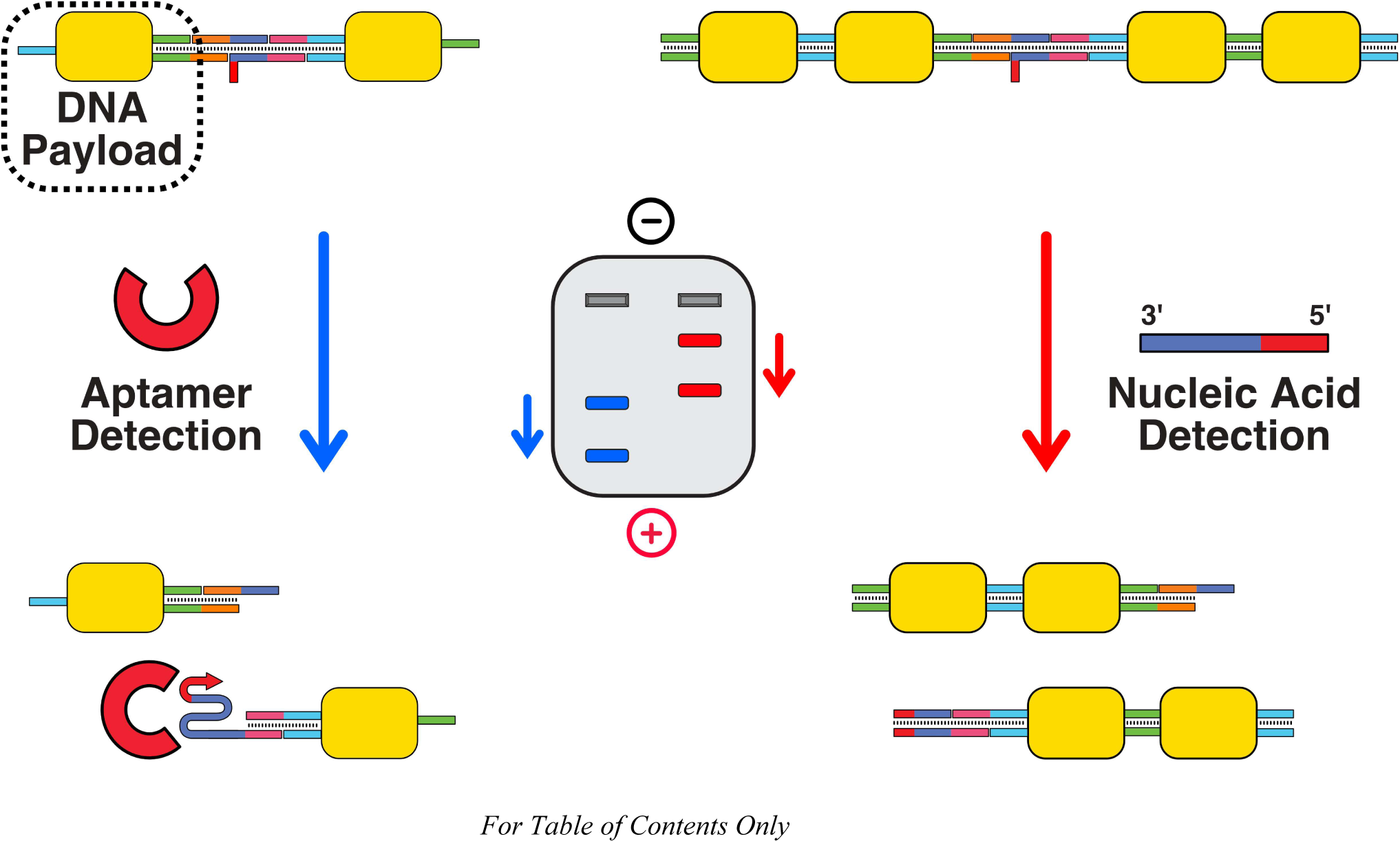

## Introduction

Significant advances in bioinformatics have led to major breakthroughs in our understanding and treatment of many complex diseases^1–4^. A large variety of molecular biomarkers from across the-ome spectrum (genomics, proteomics, metabolomics, *etc*.), extracted from blood, urine, *etc*., can now be used to help inform disease diagnosis, monitoring and medication^5–7^. As a direct result of this, healthcare systems have begun the transition to ‘precision medicine’, where patients are provided with personalized treatments selected for maximum efficacy against their unique multi-omic biomarker profile^8–11^. However, precision medicine can only be achieved if the methods for detecting and quantifying biomarkers are inexpensive and efficient enough to be applied at scale in clinical settings rather than in limited-throughput laboratories^12,13^.

At present, building up a multi-omic biomarker profile involves a large number of different techniques, each with their own set of limitations and complications^14^. Profiling the proteome often relies on expensive enzyme-linked immunosorbent assay (ELISA) tests (or other immunoassays)^15^, the metabolome depends almost exclusively on the technically complex method of mass spectrometry^16^ and the genome/transcriptome require techniques such as next-generation sequencing and PCR^17^. Building a multi-omic biomarker profile using a combination of these techniques is a technical and logistical challenge, and is currently extremely difficult to achieve in clinical settings^18,19^.

DNA nanotechnology-based diagnostic methods are versatile alternatives that could help address many of the challenges of biomarker detection. DNA nanostructures have been shown to be capable of detecting a variety of different biomarkers, by incorporating or conjugating with affinity reagents such as aptamers and antibodies^20–23^. Furthermore, DNA nanotechnology-based assays are compatible with a large range of readouts, including electrochemistry^24^, pH^25^, conformational changes detected via gel electrophoresis^26–28^, fluorescence imaging^29,30^ and naked-eye colorimetric changes^31^. This flexibility in both target analyte and readout could allow a single DNA nanotechnology-based assay to detect multiple classes of biomarkers simultaneously. Such assays would remove the requirement for several separate protocols and machines to detect different classes of biomarkers, significantly lowering the barrier for deployment in healthcare settings. While a number of such multiplexed detection methods have been outlined in the literature^24,26,32^, limitations such as scalability, regulations and cost^33^ have prevented their widespread use outside research laboratories.

In this work, we present a new technique for multiplexed biomarker detection, with a number of notable advantages. In brief, our detection principle revolves around the manipulation of the length of DNA nanostructure chains. These chains are assembled from DNA payload subunits, each of which consists of a double-stranded DNA (dsDNA) core flanked by two single-stranded DNA (ssDNA) sticky ends. The number of payloads within a chain dictates its length, which can be probed easily using techniques such as electrophoresis. To detect biomarkers, we use strand displacement or aptamer-binding events to disassemble long payload chains whenever a target biomarker is present. This action releases shorter payload chains that are designed to possess a unique length for each individual biomarker detected. This results in a characteristic pattern of bands on a gel electrophoresis scan, which can be interpreted to confirm the presence and quantity of each biomarker, in a single run. Just four strands of unlabeled DNA are required per payload unit, which makes them significantly less expensive than DNA origami-based detection systems^34^, which typically require 500+ unique strands per structure.

We show that our DNA payload assay can detect different classes of biomarkers with high sensitivity (limit of detection (LOD) of 2.9nM for DNA, and 222nM for a steroid target), high specificity (resistant to even single point mutations in target nucleic acids) and in a clinically-relevant medium (fetal bovine serum). We demonstrate the technique’s multiplexing capabilities by detecting multiple biomarkers simultaneously. Furthermore, we show that our assay can be performed with capillary electrophoresis, which is amenable to automation and integration in high-throughput clinical testing.

## Results and Discussion

### Formation of Payloads and Principle of Detection

Gel electrophoresis separates biomolecules such as DNA based on shape, size and charge^35^. Thus, for any detection technique using gel electrophoresis as the primary readout mechanism, some form of structural or conformational change needs to be induced in a nanostructure to cause an easily distinguishable shift in the corresponding band on a gel scan. Typically, electrophoresis-based biomarker detection assays use structural changes in relatively massive DNA origami scaffolds^26^ or nanostructures with complex shapes^28^ to produce easily-distinguishable gel bands. In contrast, we focused on keeping the nanostructures as small and as simple as possible, to keep costs low and reduce the number of steps in the assembly process. Our detection method is based on a 4-strand DNA payload unit (Figure 1A left dotted box). The payload unit is made up of two components: a 30-base pair (bp) dsDNA core and two uniquely addressable 15-nucleotide (nt) sticky ends. Due to their simplicity, payloads can be assembled efficiently and in high concentrations using a short ~1.5 hour annealing protocol (Methods). In isolation, these payloads produce a single well-defined band on a gel electrophoresis scan with an intercalating fluorescent dye due to their dsDNA core. When two payloads are bound together via a combination of adapter and linker strands, a dimer structure (‘Target-Specific Detection Unit’, Figure 1A right dotted box) is formed and this moves more slowly through the gel than the original single unit payloads.

**Figure 1:**
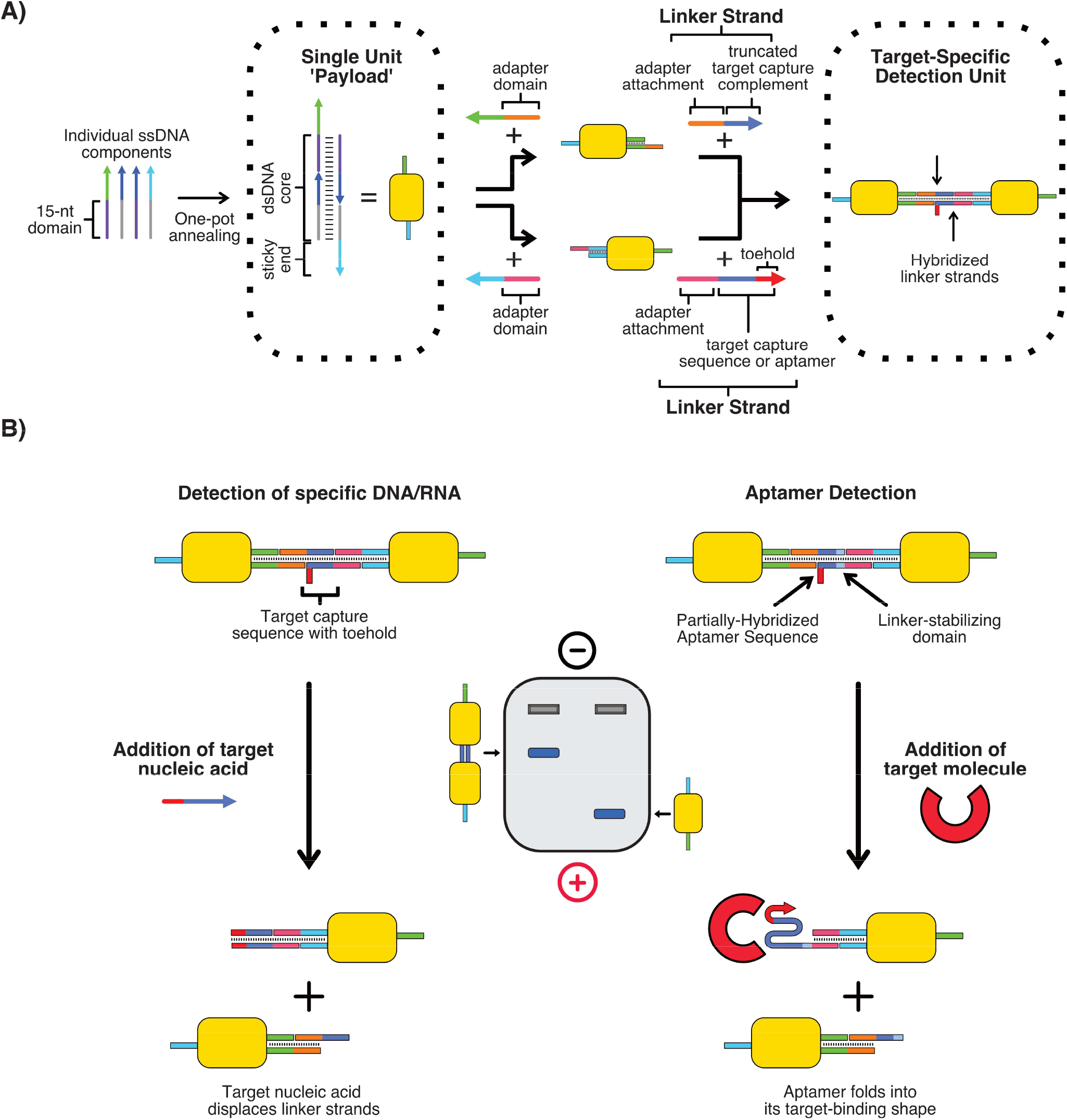
Assembly of payload structures and detection principle. A) Payload structure consists of a 30-bp dsDNA core with two unique 15-nt sticky ends. Adapter strands can be used to standardize attachment points. Payloads can then be attached together using partially hybridized linker strands. B) In the presence of the target biomarker, the connection between the two payloads in the dimer is broken, leaving individual single payloads. This causes a clear gel band shift, allowing for target detection. When targeting DNA/RNA, the capture strands are unraveled through toehold-mediated strand displacement. For targeting other types of biomarkers, the capture strands consist of a partially-hybridized aptamer. In the presence of the aptamer’s target, the aptamer should de-hybridize and fold into its target-binding shape, breaking the payload connection.

To detect a nucleic acid (DNA/RNA), the target capture domain in the ‘Target-Specific Detection Unit’ is set to contain the sequence complementary to the target nucleic acid sequence. On the matching linker, the complementary domain is truncated such that the final few bases of the target capture domain are unpaired, providing a single-stranded toehold. With no target present, the complementary linker strands hybridize, binding the two payloads together into a dimer. However, when the target is added, it binds to the toehold, invades the duplex and unravels the payload linkage via toehold-mediated strand displacement^36^. This breaks apart the target-specific detection unit, releasing the two payloads and shifting the corresponding gel band to the single-payload location, enabling detection of the target (Figure 1B, left).

For detection of targets other than nucleic acids, the target capture domains of the linker are designed to incorporate a partially hybridized aptamer sequence. In the absence of the target, the linkers keep the payloads attached in the dimer configuration, leaving a portion of the aptamer unhybridized (Figure 1B, right). However, the presence of the target biomarker causes the aptamer to unravel its hybridized structure and preferentially fold into its specific binding conformation. The number of unhybridized aptamer bases required to allow for preferential binding to the target will depend on the dissociation constant and structure of the aptamer selected. Additionally, an extra domain can be added to the linker pair to stabilize the linker hybridization if the aptamer duplex is too short to support hybridization under normal conditions. Once the aptamer starts to fold to bind to its target, this will cause the payload linkers to de-hybridize, which releases the individual payloads and causes a corresponding gel band shift (Figure 1B, right). Together, the aptamer and strand displacement mechanisms can detect a large variety of biomarkers, as aptamers have been developed for a wide range of targets^20^.

### Biomarker Detection: Sensitivity and Specificity

To test and characterize our methodology, we designed target-specific detection units to target different classes of biomarkers. We selected a panel consisting of DNA (a 25-nt fragment of the BRCA1 gene, identical to that tested by Chandrasekaran et al.^26^), microRNA (miR-141, also identical to that tested by Chandrasekaran et al.^26^), and aldosterone, a steroid with a well-defined aptamer^37^. Each of the targets selected can act as a clinically relevant biomarker for different diseases: BRCA1 mutations are highly associated with breast cancer^38^, miR-141 is expressed abnormally in many cancer tumors^39^ while aldosterone is used for the evaluation of high blood pressure and diagnosing/monitoring adrenal gland tumors and other disorders^40^. In particular, we deliberately selected a steroid biomarker for our analysis as steroids are often very challenging to detect due to their low circulating concentration in blood (pM-nM) ^41–43^. Successfully detecting a steroid at a clinically relevant concentration indicates that detection of most other plasma analytes, which circulate at much higher concentrations^43,44^, should be easily achievable with our assay. We set the target capture domain according to the type of biomarker – the complementary sequence for nucleic acids or the appropriate aptamer sequence for other biomolecules. We prepared dimer payloads as described in the Methods. We combined each dimer with its target biomarker and observed the formation or increase in intensity of a single fast gel band for each dimer/target pair tested (Figure 2).

**Figure 2:**
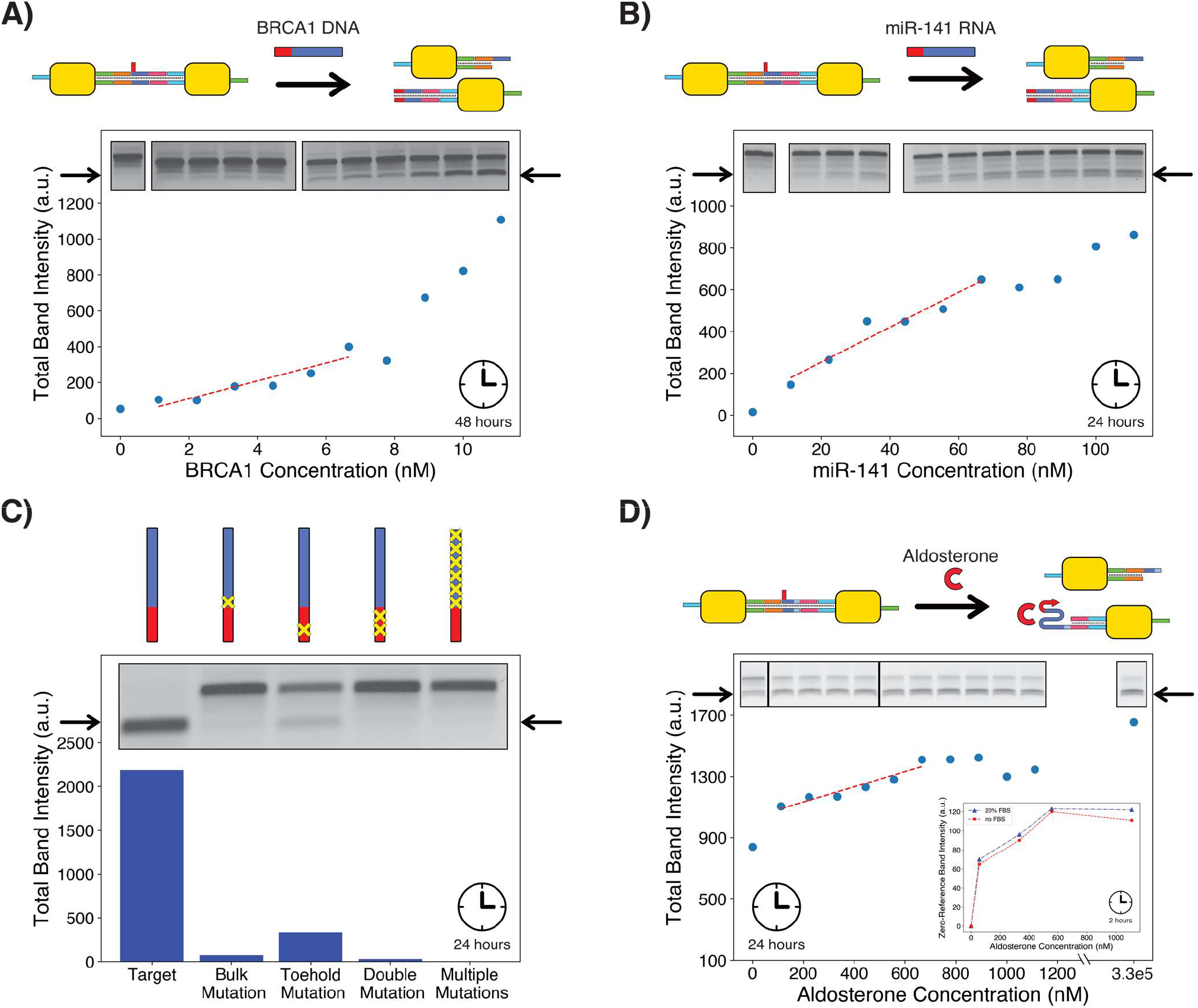
Detection of various biomarkers using purified dimer payloads; each panel has a different y-axis range due to variations in band size and image brightness. All gels are TBE-based, except for panel C, which is TAE-based (TBE version of this result available in Figure S2). Full gel images for panels A, B and D are available in Figures S3-S5. None of the gel images are computer-adjusted and only vary in exposure time (between 10-25s). a.u. = arbitrary units. A) Detection of single-stranded BRCA1 DNA fragment, graph points correspond to the gel bands embedded within the figure. LOD calculated to be 2.9nM, using the linear trendline y=49.81x + 8.80 shown on the graph. B) Detection of miR-141 RNA fragment. Results similar to those for BRCA1, but LOD is higher (18.7nM, with linear trendline y=8.33x + 86.67 shown on graph). Output quantification encompasses both visible single-payload bands. C) Specificity analysis on BRCA1 DNA fragment. A clear reduction in output signal is observed against all incorrect targets, including those with just one mismatch. All targets were introduced at a concentration of 111nM and incubated for 24 hours. D) Detection of aldosterone using aptamer payloads. Due to the aptamer partially dissociating without any target present, the output band quickly saturates, preventing accurate quantification of aldosterone at higher concentrations, but still allowing for detection (LOD 222nM, with linear trendline y=0.50x + 1033.93 shown on graph). Output quantification encompasses both visible single-payload bands. The inset shows detection of aldosterone in 20% FBS produces results which are very close to those in standard buffer. ‘Zero-Reference Band Intensity’ refers to the pixel intensity above that of the baseline output band (no target present). The actual bands corresponding to these results are provided in Figure S6. All samples for FBS comparison were incubated with their targets for 2 hours at 21°C with constant agitation prior to gel loading. Only the top output band was considered for quantification.

We analyzed the sensitivity of our assay by measuring the intensity of the output band formed after incubation with a range of target concentrations. Figures 2A and S1 show sensitivity profiles for quantification of the BRCA1 fragment after 48- and 24-hour incubation respectively, while Figure 2B shows the sensitivity profile for miR-141 RNA quantification after 24-hour incubation. We found that the assay produced a visible output gel signal in the presence of the target nucleic acid at concentrations as low as ~1nM (9μl sample containing 77.5pg of DNA) for BRCA1 after 48-hour incubation, with the LOD calculated to be 2.9nM (based on linear trendline in Figure 2A, details on calculation in the Methods). The concentration at which the output band is visible drops to ~3-5nM after a 24-hour incubation (Figure S1). For miR-141, 24-hour incubation (Figure 2B) resulted in a visible output band at a target concentration of ~10nM (9μl sample containing 698.0pg of RNA) with a LOD of 18.7nM (calculated as before). The reduction in LOD for the miR-141 RNA can be attributed to the fact that each target nucleic acid will display different rates of strand displacement according to its sequence^45^, and whether RNA or DNA is invading the linker DNA duplex^46^. We also analyzed the specificity of our detection method by challenging our assay with various off-target variants of the BRCA1 fragment, including single mutations on both the toehold and bulk areas of the strand. The results show that our system displayed a high degree of sequence specificity (Figure 2C and S2). When exposed to targets at a concentration of 111nM for 24 hours, the signal was dramatically lower for mutated sequences than for the perfectly matched target. The intensity of the output band was 85% lower for a single mismatch on the toehold, and decreased by nearly 100% in the cases of a double mutation in the toehold, a single mutation in the displacement domain or many mutations in the displacement domain. Our technique could therefore be used to distinguish accurately between strands with a range of single nucleotide polymorphisms.

Figure 2D shows the sensitivity profile for aldosterone detection after 24-hour incubation with dimer payloads equipped with an aldosterone aptamer^37^. In contrast to the DNA/RNA payloads, the aldosterone dimer payloads appeared to be in equilibrium with the detached single payload structures (as evidenced by the presence of single-payload gel bands), even with no target present. This caused issues with accurate quantification of the target biomarker at higher concentrations, since the extra single payloads add an additional bias to the output band signal and cause it to saturate earlier. Despite this, aldosterone could still be detected at lower concentrations, with an LOD of 222nM. We expect that the signal profile for each individual target biomarker will depend on the stability and dissociation constant of the complementary aptamer. Furthermore, we demonstrated that this LOD is also maintained in a biologically relevant medium by testing aldosterone detection in 20% fetal bovine serum (FBS). As shown in the inset of Figure 2D and Figure S6, detection in FBS results in weaker output bands, but the output signal ratio to the no-target output band remains constant, allowing for quantification with no degradation in LOD.

We also attempted to detect another biomarker, the enzyme thrombin, using a dimer payload equipped with a thrombin aptamer^47^ as the linker sequence. While we were able to detect a band shift in the presence of thrombin, this resulted in a very slow band rather than the faster single-payload band we were expecting (Figure S7A). This indicates that the aptamer caused some unknown aggregation to occur after breaking its hybridized state. This experiment indicates that not every single aptamer will work with our method immediately, and might require further optimization to conditions and/or sequence to achieve the desired effect. Despite this unexpected result, the thrombin dimer payloads could still be used to detect a target DNA strand containing the sequence complementary to that of the thrombin aptamer (Figure S7B). We continued to use this dimer payload and target sequence pair in subsequent experiments.

### Multiplexed Detection

Using just payload dimers allows our technique to act as a single-target detection system. However, the nature of our payload attachment system also enables us to perform one-pot multiplexed detection without the need for any additional components or labels. The more payloads we conjugate in a chain, the slower the bands produced in the resulting gel (Figure 3, A and B). Chain elongation can be controlled by capping one of the sticky ends of the seed payload, after which elongation can only proceed in one direction (Figure 3A). With this elongation technique, the number of payloads on either side of a detection linker can be predetermined (Figure 4A) and thus allow for precise control of the position of the output band following detection.

**Figure 3:**
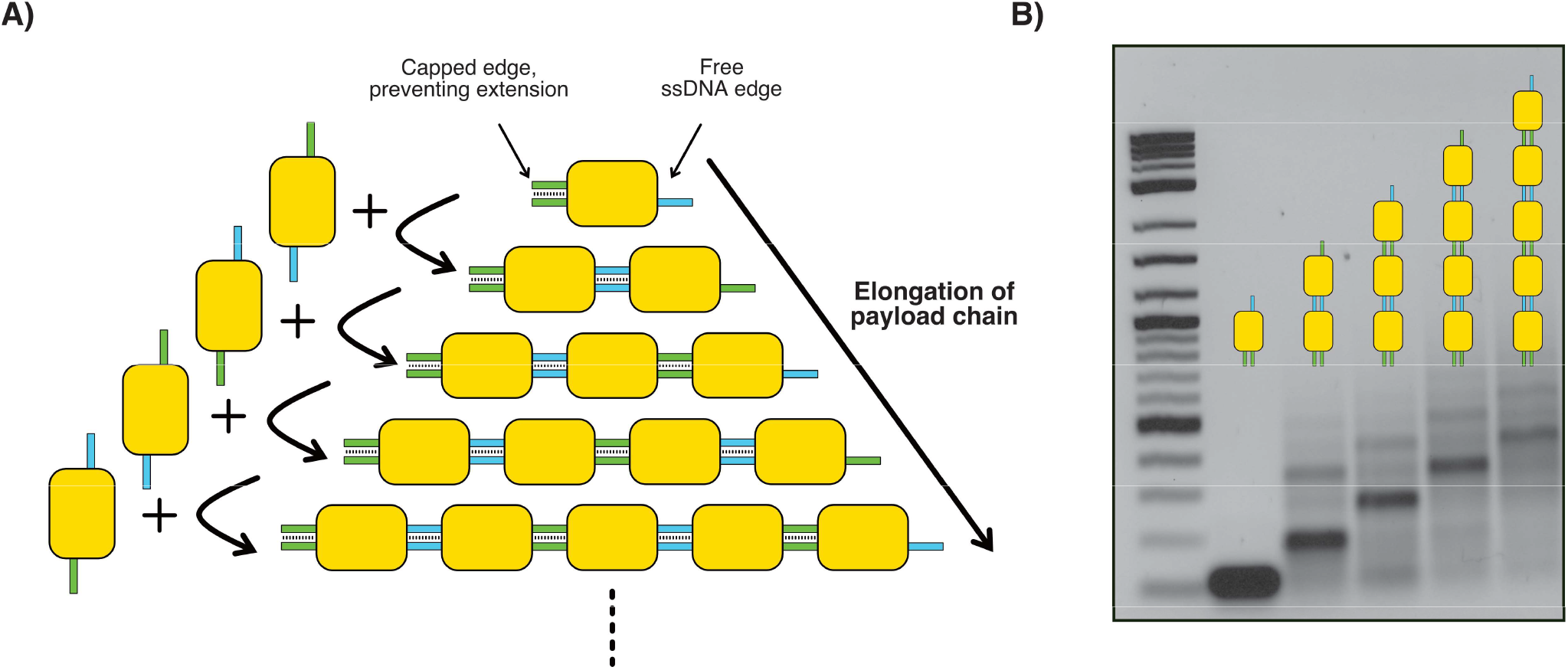
Multi-payload formation. A) Payloads can be attached together to form longer chains. ssDNA caps can be used to control the direction of elongation by blocking off one of the sticky ends. B) Longer payload chains produce slower bands for each additional payload added to the chain. The payload chains shown here were produced by sequentially adding equimolar quantities of payloads with the correct sticky ends to elongate the current chain, allowing incubation at 30°C for 30 minutes between each step. The final resulting payloads shown here were not purified and thus show some off-pathway products.

**Figure 4:**
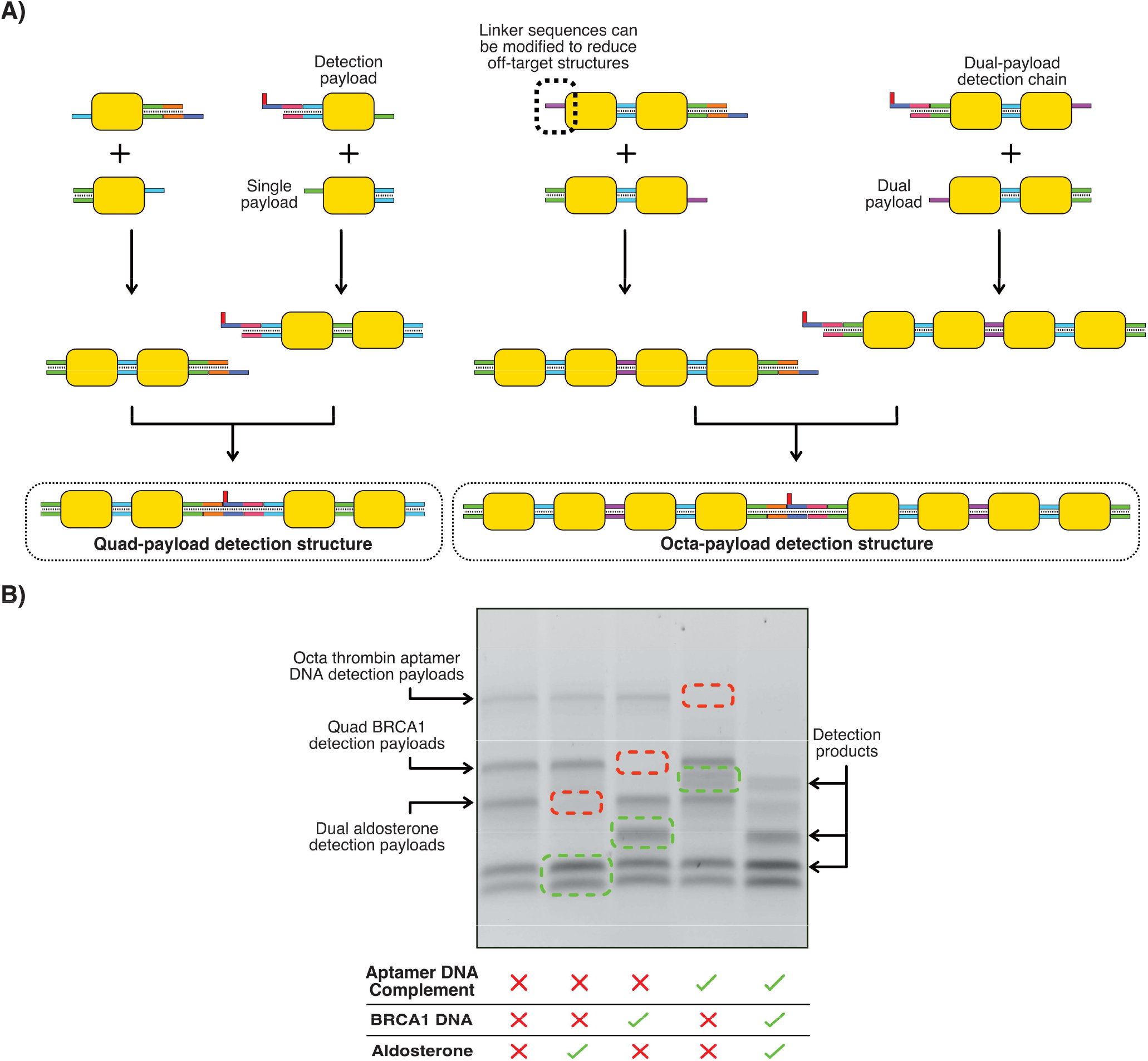
Multiplexed biomarker detection. A) Payload detection structures of variable size can be produced by controlling the length of the payload chain. Different sticky end sequences can be used in each payload to improve yield or modify payload chain assembly process. All detection structures were assembled via stepwise addition of the correct payload to elongate the chain with a 30°C incubation for 30 minutes between each step (exact details in methods section). Final products are purified before use in a detection assay. B) Multiplexed detection assay for thrombin aptamer complement, BRCA1 and aldosterone. Detection can be carried out for each target individually, or all three in one pot or some combination of targets. All targets were added in excess and were left to incubate for 30 minutes at 30°C before gel electrophoresis.

Combining payload detectors with different output bands results in a characteristic band pattern that allows for simultaneous detection of multiple biomarkers, as each target biomarker will produce a unique gel band independent of any other structure in the mixture. We subsequently refer to individual payload chains according to the number of payloads within the chain i.e. ‘dual’ refers to a chain with two payloads, ‘quad’ refers a chain with four payloads, ‘octa’ refers to a chain with eight payloads, *etc*.

To demonstrate this principle, we created three different detection payload chains for two DNA targets (the previously tested BRCA1 fragment and the DNA complement to the thrombin aptamer) and aldosterone. We designed each detection chain to contain a different number of payloads: 4 on each side (octa) for the thrombin aptamer complement, 2 on each side (quad) for the BRCA1 fragment and 1 on each side (dual) for the aldosterone target. After assembly and purification, we combined each of the detection chains in one mixture, and exposed them to different combinations of the targets (Figure 4B). The results clearly show that each of the detection structures disassociated only in the presence of their target, producing a unique, faster gel band corresponding to the payload chain generated after splitting apart the longer detection chain. The simultaneous detection of multiple biomarkers produced a gel band pattern which could be interpreted to enable detection of all targeted biomarkers. An additional gel image showing detection of RNA with a payload chain containing 6 payloads (hexa-chain) is available in the supplementary information (Figure S8).

### Capillary Electrophoresis

The results of Figure 2 and Figure 4 have shown that our technique is capable of targeted individual and multiplexed detection of various biomarkers. However, conventional gel electrophoresis requires many steps, even when using precast gel systems^48^. In contrast, capillary electrophoresis systems, as used in Sanger sequencing^49^, present an opportunity for automating the entire process of detection and quantification. Capillary electrophoresis machines separate nanostructures by forcing them through capillary tubes filled with a gel/stain mixture, while automatically detecting fluorescence via a built-in detector at the end of the capillary tubes (Figure 5A). To test whether our structures were compatible with such a system, we evaluated the detection of our biomarker panel (BRCA1 DNA, miR-141 RNA and aldosterone) with an Agilent capillary electrophoresis system. We used unpurified dual detection payloads for each of the targets, due to the machine’s relatively high DNA LOD (0.5ng/μl) and the challenges inherent in producing large amounts of purified payloads. The plots in Figures 5B-D show the results obtained after mixing each payload detector with different concentrations of each target. Each plot shows that the principle of detection works properly with capillary electrophoresis, for both nucleic acid and aptamer targets. Dual detection payloads are left intact when no target is present and dissociate into single payloads when their target is added, producing in each case an output band with an intensity that depends on the target concentration.

**Figure 5:**
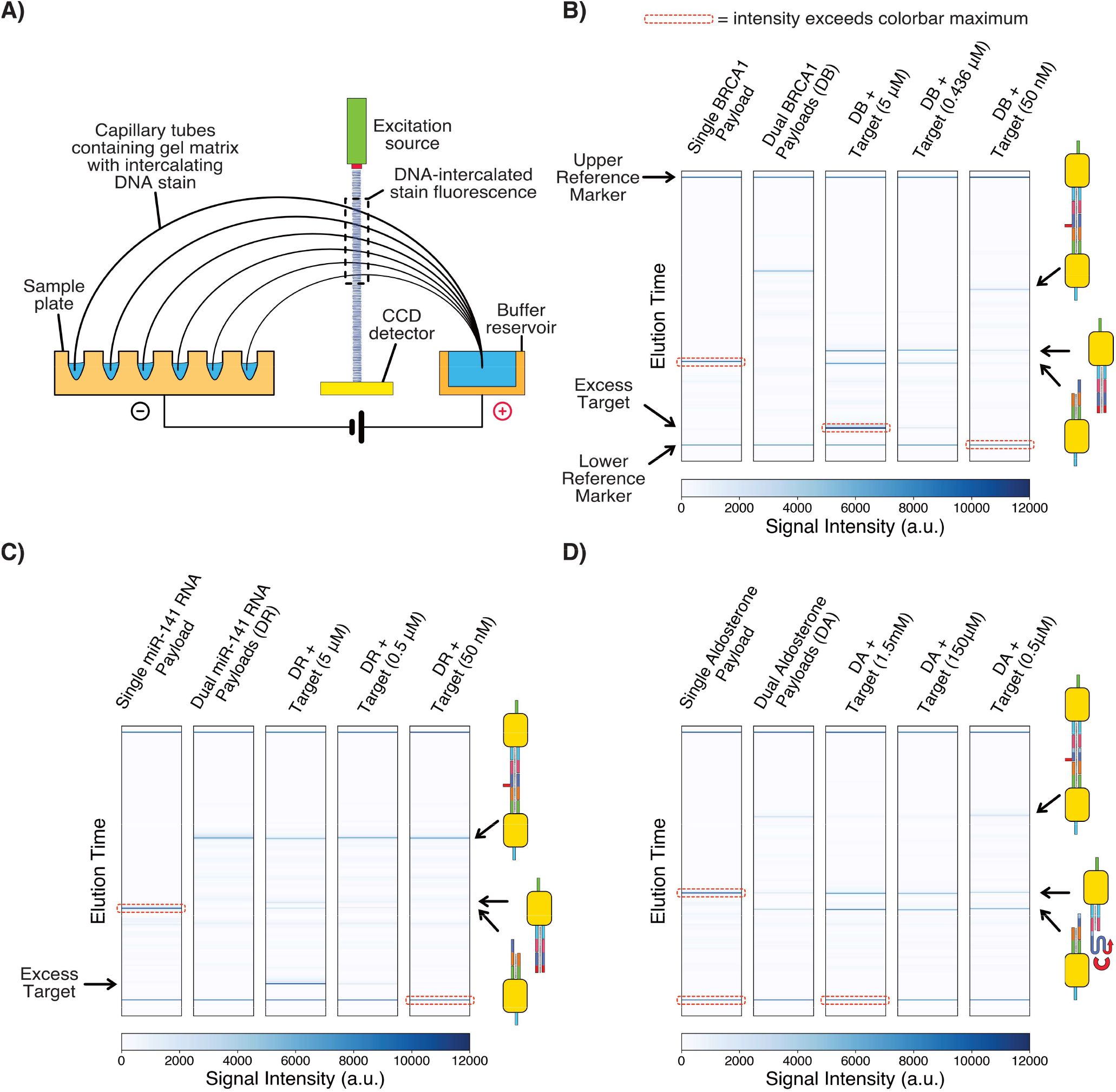
Capillary electrophoresis results. All target and payload detector mixtures were allowed to incubate at room temperature for ~75 minutes prior to capillary electrophoresis. All detection payloads were used without purification and thus some impurities are visible around the main payload bands. a.u. = arbitrary units. A) Principle of capillary electrophoresis detection. B) Detection of BRCA1 DNA. C). Detection of miR-141 RNA. D) Detection of aldosterone. As before, dimer payloads are in equilibrium with single payloads, even when no target is present.

The capillary results also displayed several unique advantages over those of standard gel electrophoresis. As can be seen in all three panels of Figure 5, the bands produced for each structure were very thin and specific, with much less variation than in standard gel electrophoresis. This means that different dual payloads will produce slightly different bands due to variations in the sequences of the target capture domains. This makes it simpler to produce a multiplexed readout, as multiple dual payloads could be mixed together, with each occupying different spaces on the gel scan (an example is shown in Figure S9). Additionally, working with unpurified structures ensures that the concentration of the detection chains is significantly higher than the purified chains tested in Figure 2. This higher concentration indirectly had the effect of allowing the detection reaction to occur significantly faster. In fact, we achieved detection of 50nM of miR-141 RNA (Figure 5C) after incubation for just 75 minutes, as opposed to the 24 hours it took for standard gel electrophoresis. The combination of a more automation-friendly procedure, rapid detection and straightforward interpretability makes capillary electrophoresis more suited for clinical applications.

## Conclusions

We have presented a technique for multiplexed biomarker detection, with a number of key features that give it great potential as a diagnostic method. We have shown that our approach can detect specific nucleic acids (genome/transcriptome) with a LOD of up to 2.9nM (22.5ng/ml) with a standard gel electrophoresis setup, with 100% mutation specificity. Enzyme-free DNA/RNA amplification techniques^50^ could be coupled with our assay to improve the LOD further. Detection of non-nucleic acid biomarkers can be taken care of via aptamers, and we have demonstrated this by successfully detecting the steroid aldosterone (part of the metabolome) with a LOD of 222nM (80.0ng/ml) for this particular aptamer. This is 2 to 3 orders of magnitude above the physiological range for plasma aldosterone^41^, but in the physiological normal range for the more abundant and more widely measured steroid hormone cortisol^42^. We thus establish that the method we describe, even without further refinement to enhance the LOD further, is already capable of detecting steroid hormones at levels that are physiologically relevant for some. Detecting other small molecule targets can be achieved by replacing the linkers with the correct aptamer for the required target. A large number of aptamers targeting many different types of molecules^20^ already exist, which significantly increases the range of targets our detection method could be applied on. Furthermore, we have shown that detection in a more complex medium such as FBS does not have an impact on our method’s ability to quantitate a target biomarker. While our assay results are reproducible under the conditions we have tested, further testing in biological conditions/media with a large number of repeated measurements would be required to properly quantify the precision and reliability of our technique in a clinical setting.

Our DNA payload chaining mechanism allows our method to perform multiplexed detection without any changes to the detection assay format. We have demonstrated this principle with the simultaneous detection of a panel of two nucleic acids and aldosterone in one pot. The entire technique requires just 6-10 unique DNA strands for the bulk payload chains, and 2 extra strands per target to allow for specific detection. To properly assemble our higher order payload chains and prove our detection principle, we used low-throughput gel purification (details in methods). However, nanostructure purification could be performed using higher-throughput methods such as size-exclusion chromatography which would significantly improve nanostructure yield and production scalability^51^.

Importantly, capillary electrophoresis can help to automate some of the more laborious parts of electrophoresis, facilitating application for diagnostic use. Furthermore, capillary electrophoresis machines can be produced for $500^52^ and the running costs are lower than using standard gel electrophoresis^53,54^. Coupling our technique’s low production requirements, multiplexing capability and easy automation, the barriers to introduction of targeted biomarker testing to the clinic are significantly lower than the current alternative of combining multiple conventional techniques to achieve the same outcome.

## Methods

All nanostructures used in this paper were composed of synthetic DNA strands bought from Integrated DNA Technologies (IDT) with standard desalting. For sequences longer than 60 bases, the Ultramer option was selected, in order to ensure maximum sequence purity. Upon receipt, all strands were resuspended in ultra-pure water at a concentration of 100μM (most oligonucleotides) or 50μM (Ultramers only) and stored at −20°C. The full list of sequences used is provided in the supplementary information (nucleic_acid_sequences.xlsx).

### Preparation of individual DNA Payloads

To prepare a single payload block composed of 4 individual strands (or 4 individual strands + sticky-end caps), the strands were combined in a 1xTAE-Mg buffer (1xTAE at pH 8.3 + 12.5mM magnesium acetate) at a constant volume of 200μl. The final concentration of each strand was either 1μM, 2μM or 5μM according to the final payload construct desired (details in supplementary material). Once combined, the strand mixture was annealed in a thermocycler using the following protocol: heat to 95°C for 5 min, then slowly cool at 1°C/min down to 27°C, then remove mixture from thermocycler and allow to cool to room temperature. All single payload structures were constructed using this method, after which the nanostructures were used unpurified for subsequent reactions.

### Preparation of Dual-Payload Detection Structures

For detection structures that only contain two payloads (‘dual’), two unpurified single payloads along with their detection linkers and adapters were combined in equimolar quantities and allowed to anneal at 30°C for 30 minutes. For detection assays performed using capillary electrophoresis, these structures were used without further purification. For assays performed with ‘normal’ gel electrophoresis, purification was carried out (as described in the purification section).

### Preparation of Multiple-Payload Detection Structures

For preparation of multiple-payload structures, each individual payload was first assembled/annealed separately. Subsequently, payloads were combined in sequence (exact steps differ for each unique structure, full details and steps available in the supplementary information) in equimolar proportions to produce the final linked structure, heating to 30°C for 30 minutes at each mixing step. All final nanostructures were purified (as detailed in purification section) to remove off-target structures.

### Gel Electrophoresis

#### Loading Dye

For all gels, a non-EDTA based loading dye (LD) was prepared at 6x concentration with the following formulation: 25mg (to give final concentration ~0.25% w/v) of Bromophenol Blue (Fisher Scientific - m32712) was combined with 1.5g (to give final concentration ~15% w/v) of Ficoll-400 (Merck - F2637) and topped up with ultra-pure water to a final volume of 10ml in a centrifuge tube. This was mixed vigorously and centrifuged for 10 minutes at 5000rpm to ensure all sediment settled at the bottom of the tube.

#### TAE Gels

For TAE gels, samples were combined with GelRed (Merck – SCT123) and 6xLD to a final concentration of 1xGelRed and 1xLD. For 12-lane gels, this consisted of combining 9μl of sample with 1μl of 12xGelRed and 2μl of 6xLD. For 8-lane gels, 18ul of sample was combined with 2μl of 12xGelRed and 4μl of 6xLD. The final sample mixtures were loaded into wells of unstained 1xTAE agarose gels. For detection assays, 2% agarose gels were used, made using high-resolution agarose (Cambridge Bioscience – MB-133-0025). For purification and non-quantitative tests, multi-purpose agarose was used (Fisher Scientific – 10766834). Gels for detection assays were run at 100V for 1-2 hours in a 1xTAE running buffer.

#### TBE Gels

For TBE gels, only 8-lane gel casts were used. Gel samples were prepared by combining 9μl of the unmodified sample with 9μl of 0.5xTBE (to a final concentration of 0.25xTBE). The final mixture was combined with 2μl of 12xGelRed and 4μl of 6xLD to a final concentration of 1xGelRed and 1xLD. Samples were then loaded into unstained 0.5xTBE 2% agarose gels, and run at 100V for 1-2 hours in a 0.5xTBE running buffer. The same tanks, agarose type and power supplies were used as for TAE gels.

#### Gel Imaging

All gels were scanned using a UVP BioDoc-It imager, which produces 8-bit grayscale TIFF images. Image exposure was varied between 4 to 25s, according to the brightness of the bands tested.

#### Quantitative Gel Analysis

All gel detection assays were quantified using a custom Python analysis system (tested using Python 3.8). In brief, this involved the following steps: 1) Relevant lanes were selected using a manual rectangular selection tool. 2) All lanes were forced to have the same length and width, but could have different orientations. 3) A trace profile of the selected lane was built by calculating the mean pixel intensity throughout the lane selected, resulting in a 1D lane profile signal. 4) The background signal of each lane was estimated using the baseline correction with asymmetric least squares smoothing^55^ algorithm, after which this was subtracted from each lane profile. 5) Bands of interest were selected manually using the same selection tool, and their positions were transferred to the 1D profile. 6) Finally, the total area of the relevant band was estimated by summing the value of the clean profile signal (raw signal minus background) throughout the defined band width. A comparison of the results of our approach versus that of the established GelAnalyzer^56^ software tool are provided in Figure S10.

#### LOD Quantification

We quantified the LOD for each sensitivity profile using the following steps: 1) We fitted a linear trendline on the data with the least squares method, using only the 6 lowest target concentration data points (excluding the blank value). 2). We calculated the standard deviation of the trendline (using Excel’s *linest* function) at the y-intercept. 3) Finally, we calculated the LOD through the equation: LOD = 3.3* (standard deviation at y-intercept/trendline gradient) as recommended by the ICH Harmonised Tripartite Guidelines^57^. The LOD calculations and raw data for the profiles in Figure 2 are provided in LOD_analysis.xlsx. The linear trendlines calculated are displayed in Figure 2. Images were not modified in any way prior to analysis or display (apart from inversion to make the bands dark and the background light). The scripts used for image analysis are provided as supplementary information.

### Nanostructure Purification

Purification of specific payload-structures was achieved via a combination of gel purification and centrifugal concentration. Hexa-payload and octa-payload structures were purified via separation in a 1.5% agarose 1xTAE gel with a 1xTAE running buffer, pre-stained with 1xSYBR Safe (ThermoFisher – S33102). 20μl of each sample was added to each well, mixed with 4μl of 70% glycerol (Merck – G5516). The bands containing the target structures were located using a blue light transilluminator and extracted with a scalpel. The target nanostructures were extracted from the excised bands using Freeze N’ Squeeze spin columns (Bio-Rad) by following the manufacturer’s protocol. 50μl of 1xTAE + 125mM magnesium acetate was added to the resulting solution, after which this was topped up to 500μl with 1xTAE. Finally, the resulting solution was concentrated to 15-25μl using either 100K (Merck – UFC5100) or 10K (Merck – UFC5010) MWCO Amicon centrifugal filters. Final concentration of the structures was estimated using a nanodrop by measuring the absorbance of the final solution at 260nm and converting this value to the target concentration using the structure’s extinction coefficient. The extinction coefficient of each payload chain was approximated by adding up the extinction coefficient of all ssDNA and dsDNA components in the nanostructure using the base composition method^58^.

For dual- and quad-payload structures, gel purification was carried out in an unstained gel, and 1μl of 100xSYBR Safe was added to each unpurified sample instead. This was done because the dual/quad payloads generally had a higher concentration yield than hexa/octa payloads. Running these high-concentration samples in a pre-stained gel accumulated excessive SYBR Safe in the final excised band, which interfered with nanodrop readings at 260nm. Mixing SYBR Safe with the samples instead of the entire gel significantly reduced the amount of SYBR Safe that got extracted with the purified structures, but still allowed visual identification of the target bands using a transilluminator.

### Preparation of Aldosterone for Testing

The aldosterone used for testing was bought from Merck (A9477), dissolved in 100% ethanol to a concentration of 30mM and stored at −20°C (following the protocol defined in the paper that presented the associated aptamer^37^). All subsequent dilutions of the original aldosterone stock were also prepared in 100% ethanol.

### Preparation of RNA for Testing

The miR-141 RNA was bought lyophilized with standard desalting from IDT. This was resuspended in nuclease-free ultra-pure water (ThermoFisher – AM9930) to a concentration of 100μM and was stored at −80°C. Fresh dilutions of RNA were prepared for each quantitative test from the original stock, also in nuclease-free ultra-pure water. When preparing dilutions, an aliquot of the stock was first extracted and its concentration verified via Nanodrop (measuring absorbance at 260nm and converting to concentration using extinction coefficient provided by IDT) before any subsequent operations.

### Preparation of DNA for Testing

All target DNA was bought lyophilized with standard desalting from IDT, resuspended to 100μM in ultra-pure water and stored at −20°C. When preparing dilutions, an aliquot of the stock was first extracted and its concentration verified via Nanodrop (measuring absorbance at 260nm) before any subsequent operations. All DNA target dilutions were prepared in 1xTAE-Mg and ~0.5μM of off-target blocking oligos (sequence defined in supplementary file), in order to prevent any of the DNA target from adhering to the container walls^26^.

### Biomarker Testing

To perform a detection assay, purified chains of payload units were first diluted fivefold in 1xTAE-Mg with 0.5μM off-target blocking oligos. Subsequently, 1μl of a diluted purified chain of payload units was added to the target biomarker at the desired concentration, topped up to a volume of either 9μl (24-hour BRCA1 detection and 24-hour aldosterone detection) or 18μl (mutations analysis, 24-hour miR-141 RNA detection and 48-hour BRCA1 detection) in 1xTAE-Mg with 0.5μM of off-target blocking oligos. The biomarker concentration indicated in all figures is that of the final biomarker concentration in the mixture. After mixing, samples were left on the bench at room temperature or incubated in a thermocycler at 20°C for the desired detection timespan (marked on the graphs of Figure 2, and indicated in the supplementary figure captions). After incubation, samples were added to gels using the defined gel preparation protocol.

For detecting aldosterone samples in 12-lane gels, an additional 2μl of 6xLD was added to each sample for extra weighting, as the ethanol can cause the samples to drift out of wells prior to electrophoresis.

### Aldosterone Testing with FBS

When performing detection in FBS, the same purified and diluted payload units were used as in non-FBS tests. All assays were prepared in a 96-well plate. To perform a detection assay, 1μl of a purified chain of payload units were added to aldosterone at the desired concentration along with 1.8μl of 100% FBS (Merck – F7524) and topped up to 9μl in 1xTAE-Mg with 0.5μM of off-target blocking oligos (final concentration of 20% FBS). Non-FBS assays were also prepared with the same aldosterone target concentrations in the same plate for comparison. All samples were then incubated at 21°C and agitated at 400rpm for 2 hours. After incubation, samples were added to gels using the defined gel preparation protocol.

### Multiplexed Biomarker Testing

For multiplexed biomarker testing, the protocol is identical to that of single biomarker testing, other than that all detection structures (chains with different numbers of payloads) are combined in one solution (9μl) when detecting different targets simultaneously. For the results in Figure 4, all targets were added in excess (484nM for BRCA1, 5.6μM for the thrombin aptamer complement and 1.6mM for aldosterone), and were left to incubate for 30 minutes at 30°C before gel electrophoresis in a TBE gel. The purified octa-thrombin payloads were not diluted prior to use in the detection assay, while both the quad and dual purified detection payloads were diluted tenfold (to ensure all output bands have similar intensities).

### Capillary Electrophoresis Detection

For capillary electrophoresis tests, final samples (payload chains or payload chains with target biomarkers) were prepared to a volume of 20μl in a 96-well plate. No dyes or stains were added to samples prior to testing. DNA payload structures were used unpurified for capillary testing. When target detection testing was being performed, these were allowed to incubate for 1-2 hours prior to the capillary electrophoresis run. Samples were run using the Agilent 5400 Fragment Analyzer with the DNF-905 kit.

Final results were exported in raw CSV format (intensity signal with associated elution time), and results interpreted using a custom Python script (linked in the supplementary information).

### Preparation of Thrombin for Testing

The thrombin used for testing was bought from Merck in lyophilized form (T1063), centrifuged, dissolved in ultra-pure water to a concentration of 20μM, split into aliquots in plastic PCR tubes and stored at −20°C. All subsequent dilutions were prepared in ultra-pure water. Details for specific tests conducted using thrombin are provided in the supplementary methods file (Figure S7).

## Supporting information

Nucleic Acid Sequences

LOD analysis

Underlying data - gel images etc

## ASSOCIATED CONTENT

### Supporting Information

The following files are provided:

- supplementary_methods_figures.pdf (PDF)
- LOD_analysis.xlsx (Excel workbook)
- nucleic_acid_sequences.xlsx (Excel workbook)
- gel_data.zip (Zip file containing all raw gel images and capillary electrophoresis data)

These files contain all the data and DNA sequences used in this paper. All code used to interpret and display gel data is provided in our GitHub repository here: https://github.com/mattaq31/Gel-Analysis-Scripts

### Author Contributions

Both MA and KED were involved in the conceptualization and design of the assay methodology. MA carried out all experimental/software work. KED acquired the project funding and supervised the project. MA prepared all figures and wrote the first manuscript draft. Both authors reviewed, commented on and approved the final submitted manuscript.

## ACKNOWLEDGMENTS

This work was supported by the Medical Research Council [grant number MR/N013166/1], as part of the Precision Medicine Doctoral Training Programme. Capillary electrophoresis was carried out by the Edinburgh Genome Foundry, a synthetic biology research facility specializing in the assembly of large DNA fragments at the University of Edinburgh. We thank Robert K. Semple (Centre for Cardiovascular Sciences, University of Edinburgh) for his advice on the clinical utility and medical relevance of the biomarkers we have analyzed, as well as for reviewing the final manuscript. We thank Arun R. Chandrasekaran (The RNA Institute, SUNY Albany) and Aurélie Lacroix (Sixfold Bioscience) for their invaluable advice which helped us greatly improve and optimize our gel electrophoresis assays for our DNA payloads. We thank Katalin Kis (School of Engineering, University of Edinburgh) for her indispensable assistance and support in the laboratory, especially during the restrictions and uncertainty caused by the COVID-19 pandemic. For the purpose of open access, the author has applied a Creative Commons Attribution (CC BY) licence to any Author Accepted Manuscript version arising from this submission

