## Supplementary Methods and Figures for "Multiplexed Label-Free Biomarker Detection by Targeted Disassembly of Variable-Length DNA Payload Chains"

### Nomenclature

With reference to the sequence names in nucleic_acid_sequences.xlsx, the following list contains the individual payloads created to produce the multi-payload detection structures. Each of the individual payloads was annealed individually in a 1xTAE-Mg solution using the protocol defined in the methods section of the main paper.

**U1cap**: bulk_1 + bulk_2 + lhs_green + rhs_blue + lhs_green_cap*

**U2cap**: bulk_1 + bulk_2 + lhs_blue + rhs_green + lhs_blue_cap

**U3**: bulk_1 + bulk_2 + lhs_blue + rhs_purple

**U4**: bulk_1 + bulk_2 + lhs_purple + rhs_blue

**U1-BRCA**: bulk_1 + bulk_2 + lhs_green + rhs_blue + rhs_blue_adapter + BRCA_target_capture

**U2-BRCAco**: bulk_1 + bulk_2 + lhs_blue + rhs_green + rhs_green_adapter_complement + BRCA_target_capture_complement

**U5-RNA**: bulk_1 + bulk_2 + lhs_purple + rhs_green + rhs_green_adapter + miR_target_capture

**U5-RNAco**: bulk_1 + bulk_2 + lhs_purple + rhs_green + rhs_green_adapter_complement + miR_target_capture_complement

**U2-Thr**: bulk_1 + bulk_2 + lhs_blue + rhs_green + rhs_green_adapter + thrombin_target_capture_complement

**U2-Thrco**: bulk_1 + bulk_2 + lhs_blue + rhs_green + rhs_green_adapter_complement + thrombin_target_capture_complement

*'lhs_green_cap' and 'lhs_green_cap_extended' were used interchangeably.

**Preparation of Quad Detection Payloads**

Quad detection payloads contain two payloads on either side of the detection linkers (Figure 4A). These were prepared as follows:

1) 45µl of U1cap (5µM) were combined with 45µl of U2-BRCAco (5µM). Separately, 45µl of U2cap (5µM) were combined with 45 µl of U1-BRCA (5µM). These were allowed to anneal for 30 minutes at 30ºC. These form two individual dual-payload chains.
2) The two dual-payload solutions were combined to form the final quad-payload structure. These were again allowed to incubate at 30ºC for 30 minutes.
3) The final structures were purified before use.

**Preparation of Hexa Detection Payloads**

Hexa detection payloads contain three payloads on either side of the detection linkers. These were prepared as follows:

1) 16µl of U1cap (5µM) were combined with 80µl of U3 (1µM) and 64µl of 1xTAE-Mg. The mixture was allowed to anneal for 30 minutes at 30ºC. This forms a dual-payload chain (U1U3).
2) 80µl of U1U3 were combined with 20µl of U5-RNA (2µM). Separately, another 80µl of U1U3 were combined with 20µl of U5-RNAco (2µM). These were again allowed to incubate at 30ºC for 30 minutes. These form individual triple-payload chains.
3) The two triple-payload solutions were combined to form the final hexa-payload structure. These were again allowed to incubate at 30ºC for 30 minutes.
4) The final structures were purified before use.

**Preparation of Octa Detection Payloads**

Octa detection payloads contain four payloads on either side of the detection linkers (Figure 4A). These were prepared as follows:

1) 8µl of U1cap (5µM) were combined with 40µl of U3 (1µM) and 32µl of 1xTAE-Mg. The mixture was allowed to anneal for 30 minutes at 30ºC. This forms a dual-payload chain (U1U3).
2) 40µl of U4 (1µM) were combined with 20µl of U2-Thr (2µM). Separately, another 40µl of U4 (1µM) were combined with 20µl of U2-Thrco (2µM). The mixtures were allowed to anneal for 30 minutes at 30ºC. These form the U4U2-Thr and U4U2-Thrco dual payload chains respectively.
3) 40µl of U1U3 were combined with 30µl of U4U2-Thr. Separately, another 40µl of U1U3 were combined with 30µl of U4U2-Thrco. These were again allowed to incubate at 30ºC for 30 minutes. These form individual quad-payload chains.
4) The two quad-payload solutions were combined to form the final octa-payload structure. These were again allowed to incubate at 30ºC for 30 minutes.
5) The final structures were purified before use.

### Supplementary Figures

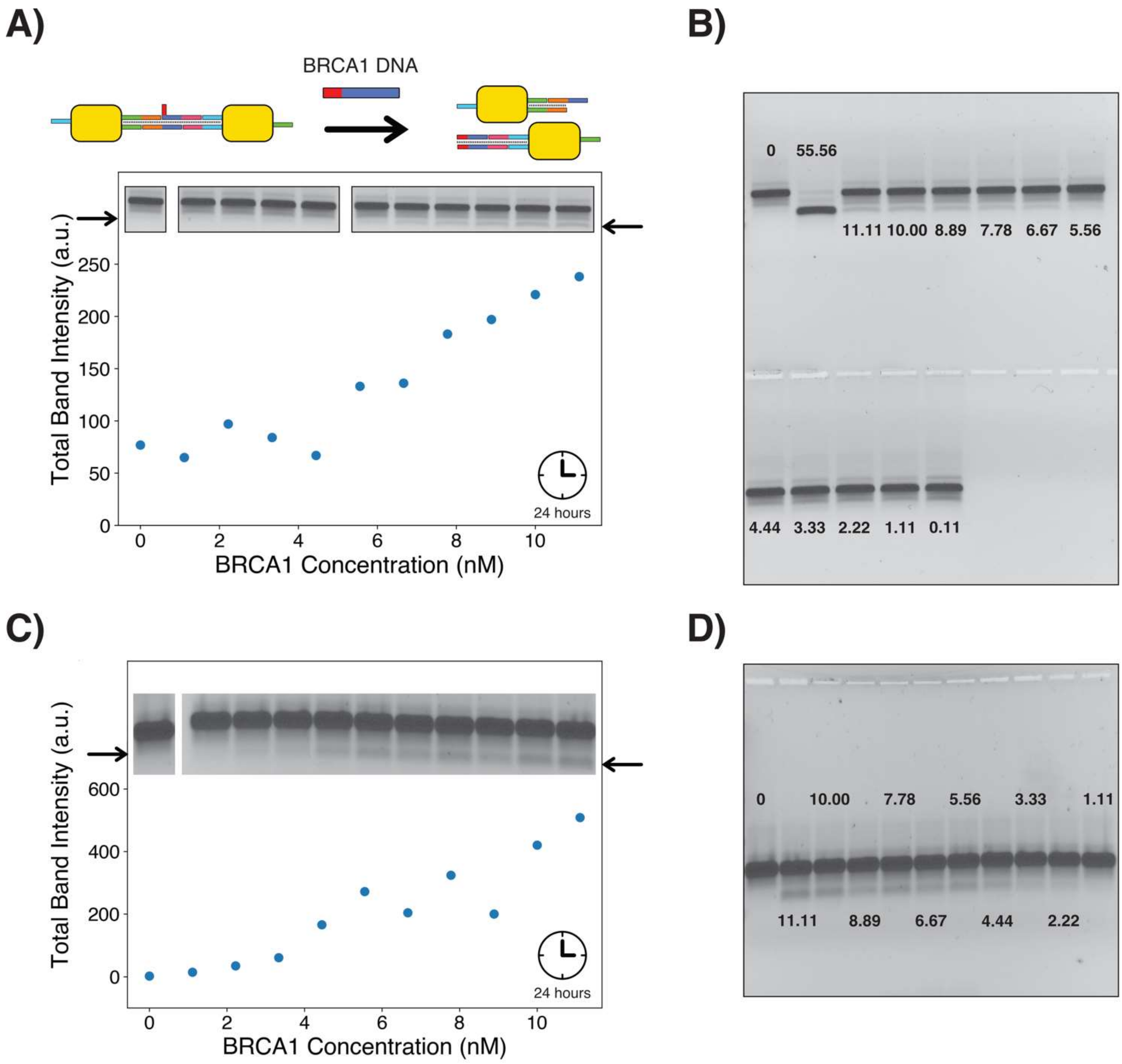


*Figure S1: Detection of single stranded BRCA1 DNA after 24-hour incubation with dual detection payload chains (in contrast to 48-hour incubation for main paper). A) Detection of BRCA1 in a TBE-based gel. Lowest visible output band is at 5.56nM BRCA1 concentration. B) Corresponding complete gel for profile in panel A. The numbers inserted in the gel indicate the concentration of BRCA1 added to each sample in nM. C) Detection of BRCA1 in a TAE-based gel. Lowest visible output band is at 3.33nM BRCA1 concentration. D) Corresponding complete gel for detection in panel C. The numbers inserted in the gel indicate the concentration of BRCA1 added to each sample in nM.*

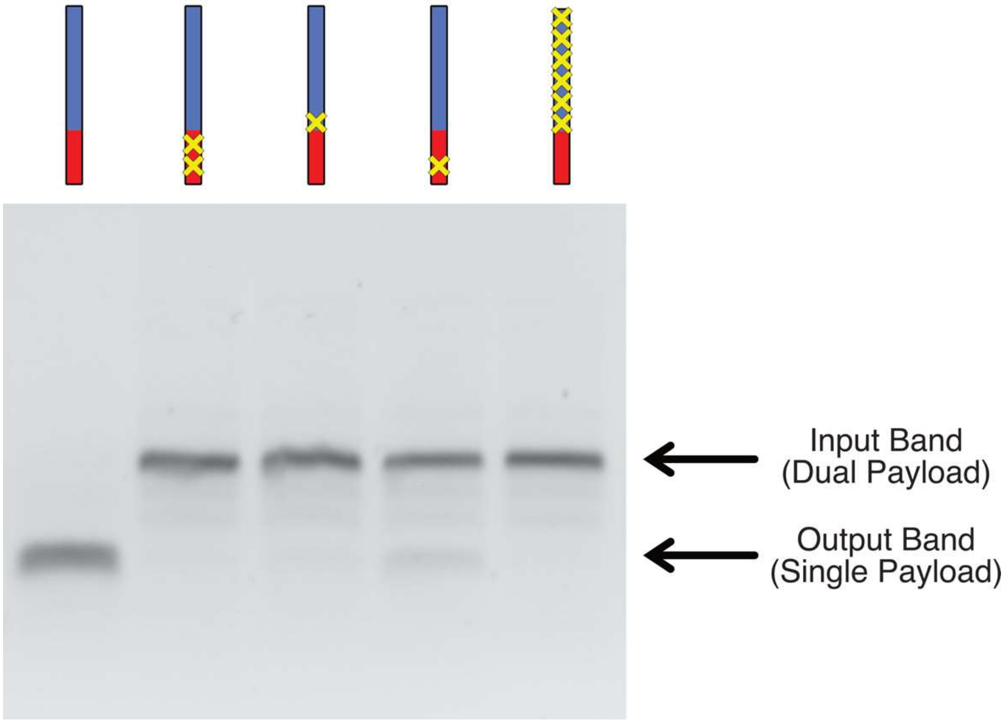


*Figure S2: Specificity analysis on BRCA1 DNA fragment in a TBE-based gel. A clear reduction in output signal is observed against all incorrect targets, including those with just one mismatch. All targets were introduced at a concentration of 111nM and incubated for 24 hours. Mutation position is indicated in DNA strands at the top of the gel.*

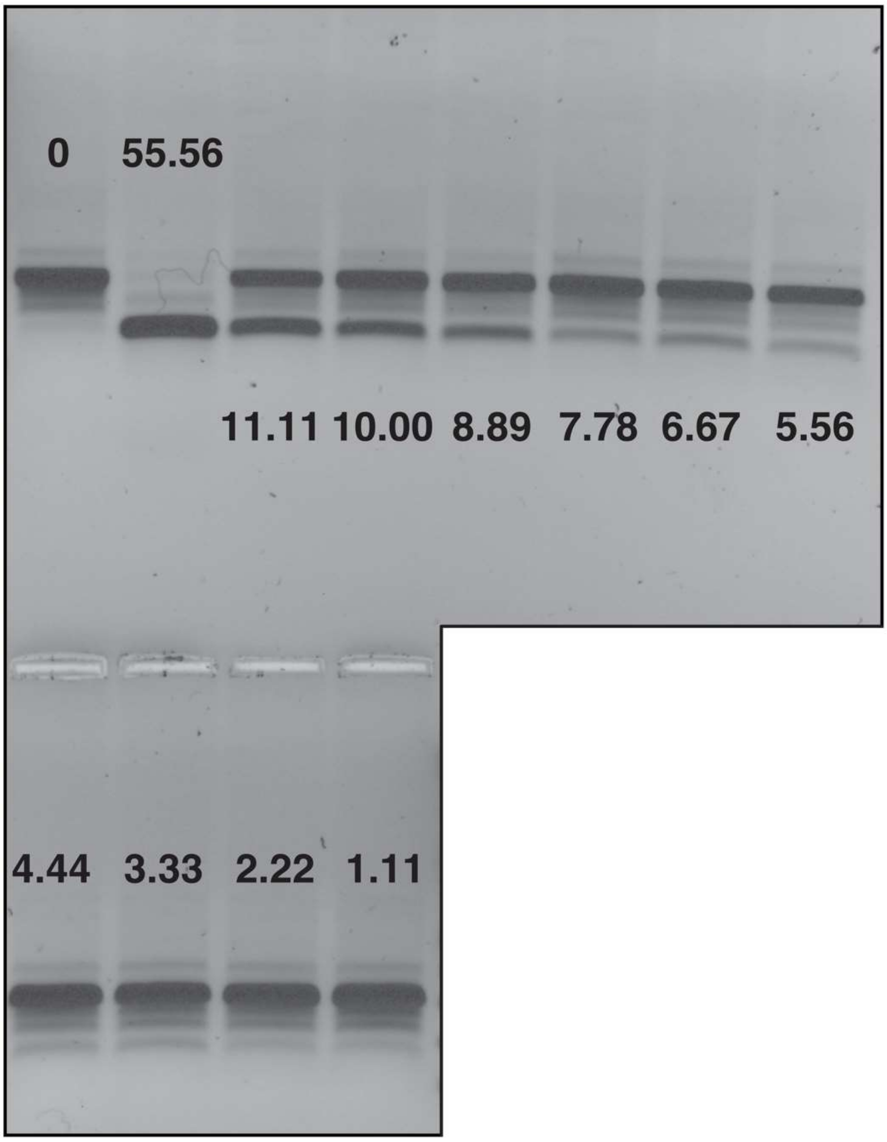


*Figure S3: Raw gel image corresponding to sensitivity profile in Figure 2A. The numbers inserted in the gel indicate the concentration of BRCA1 added to each sample in nM.*

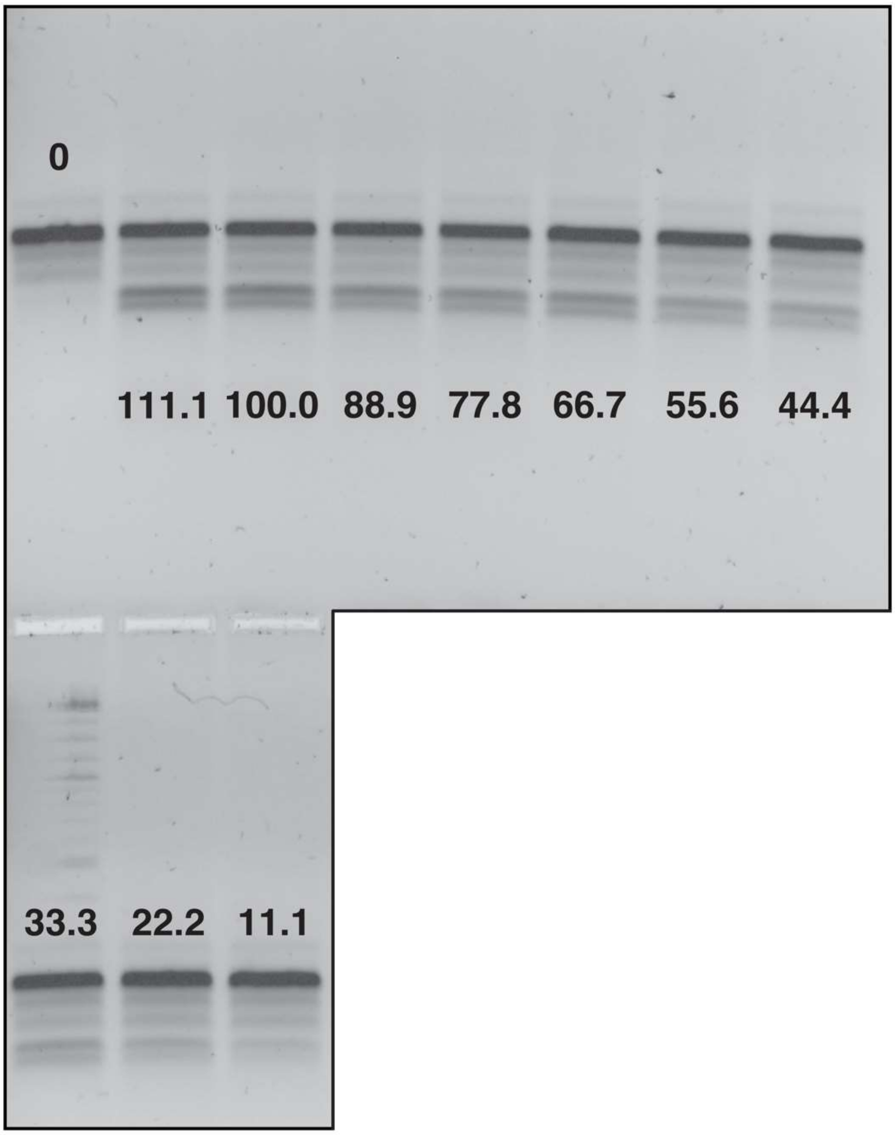


*Figure S4: Raw gel image corresponding to sensitivity profile in Figure 2B. The numbers inserted in the gel indicate the concentration of miR-141 added to each sample in nM.*

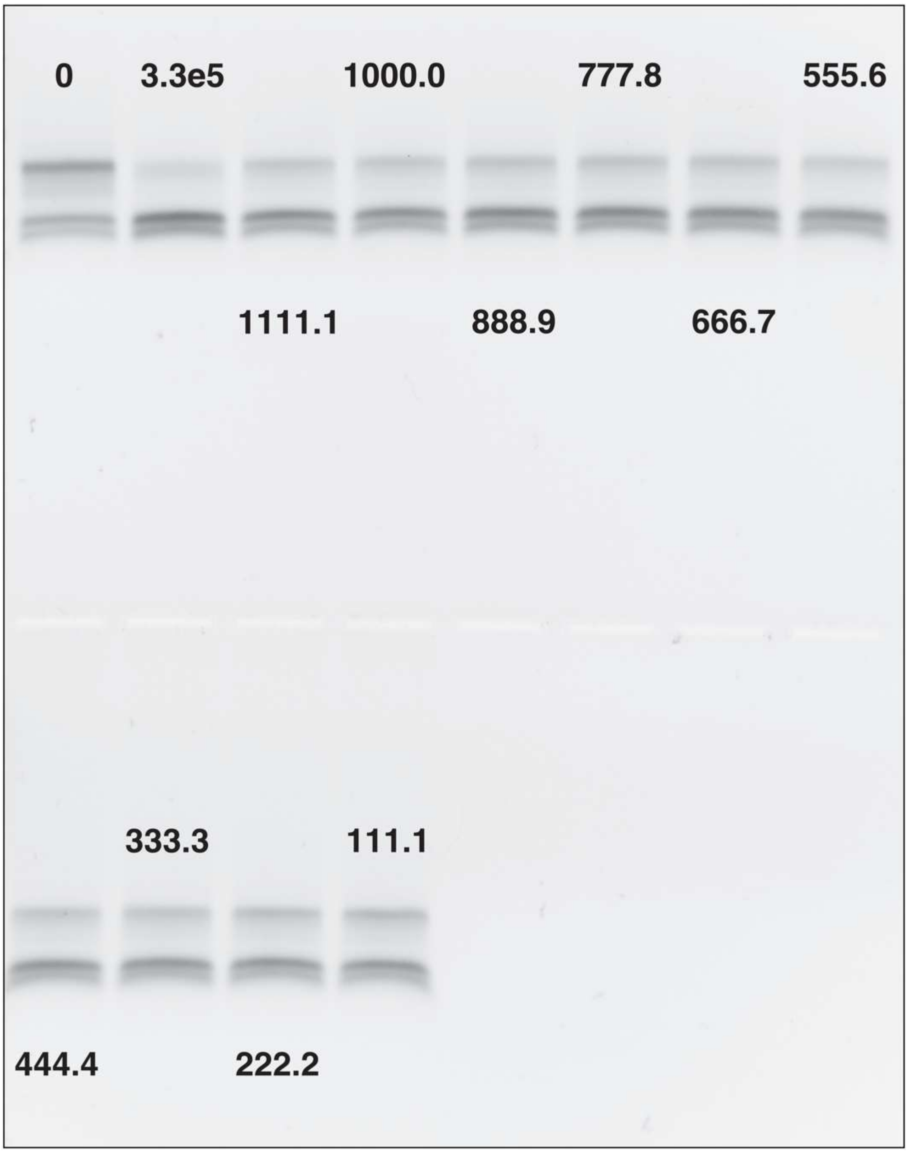


*Figure S5: Raw gel image corresponding to sensitivity profile in Figure 2D. The numbers inserted in the gel indicate the concentration of aldosterone added to each sample in nM.*

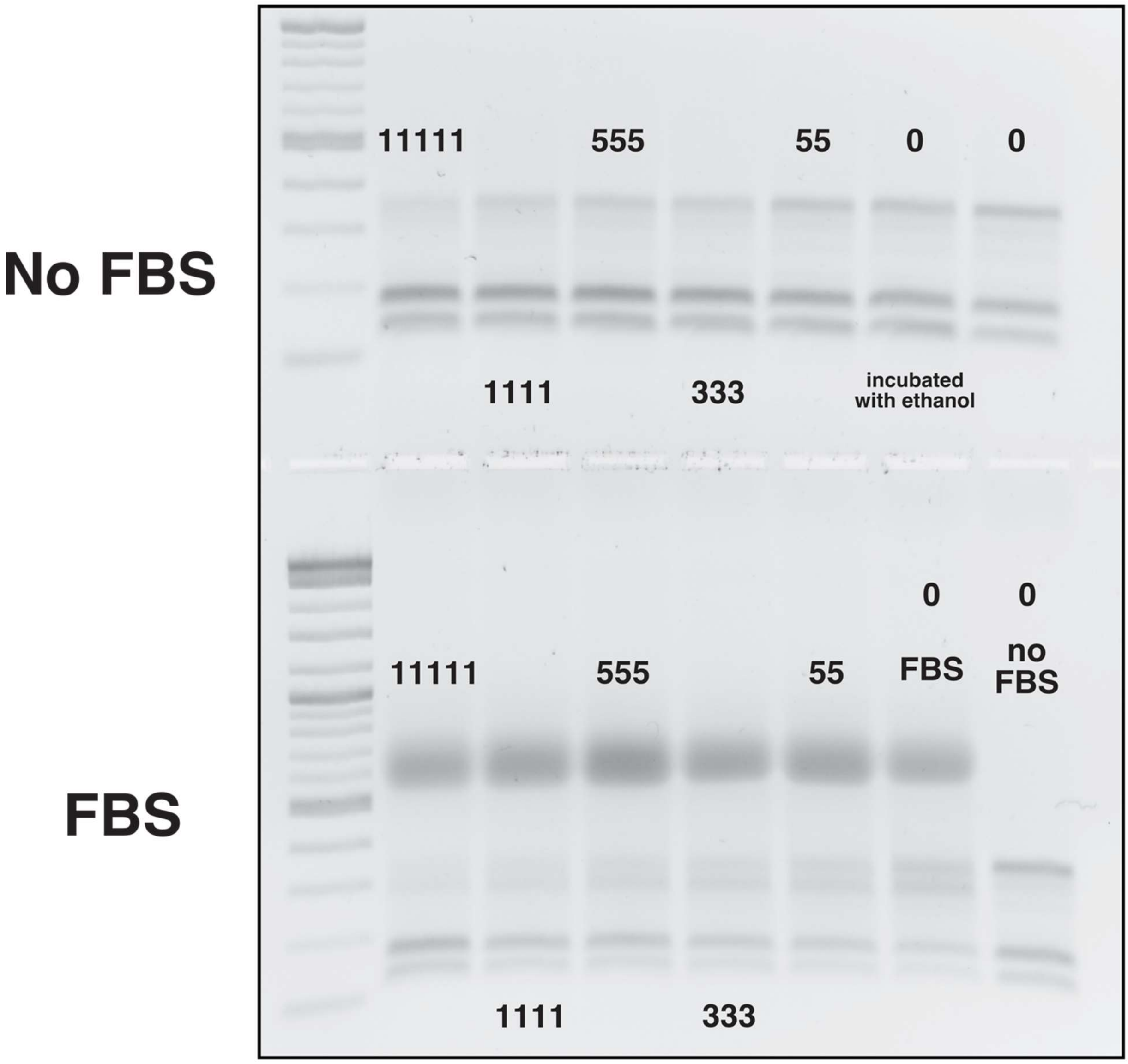


*Figure S6: Raw gel image corresponding to FBS sensitivity profile in the inset of Figure 2D. The numbers inserted in the gel indicate the concentration of aldosterone added to each sample in nM. Band quantitation was done on only one of the two output bands.*

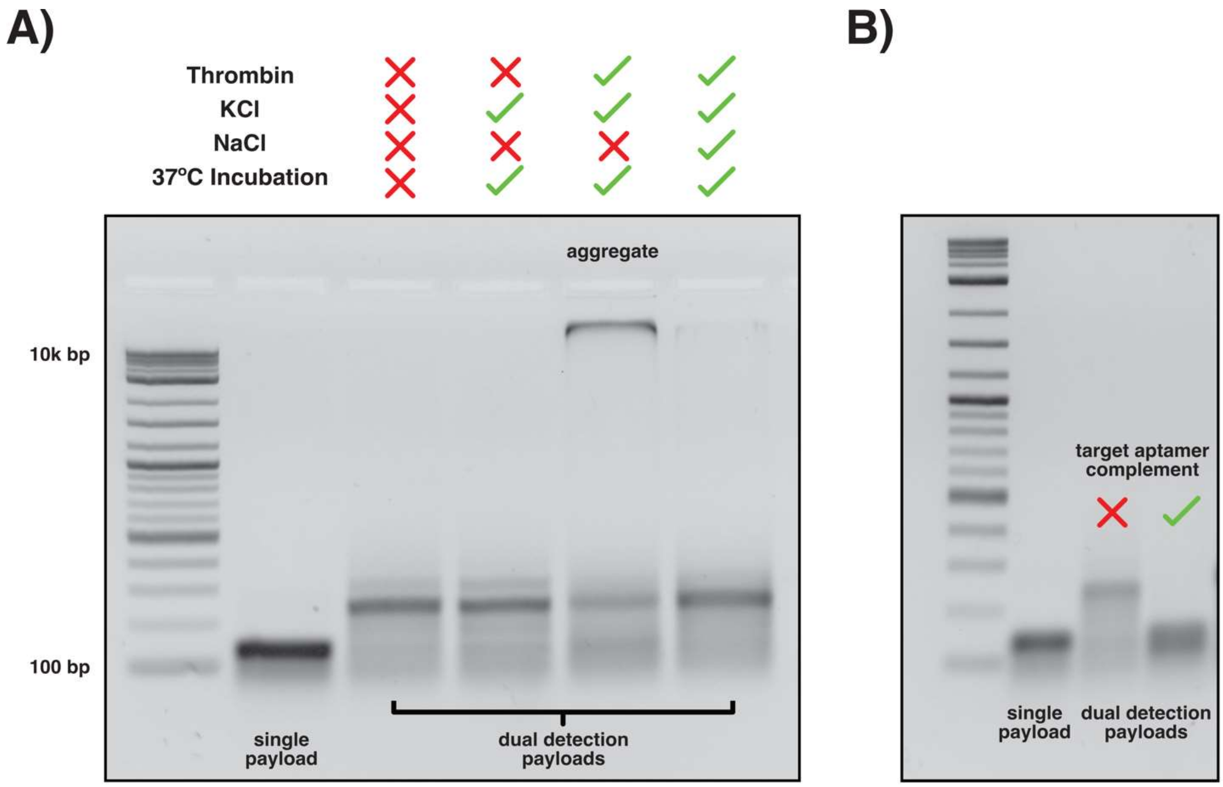


*Figure S7: Gel images obtained after testing action of thrombin detection payloads. Both are TAE-based gels. A) Detection of thrombin under various conditions. All dual detection payloads were used at a final concentration of 111nM (unpurified). When KCl was used, this was added to a final concentration of 133mM. When NaCl was used, this was added to a final concentration of 225mM. Thrombin was added to a final concentration of 2.22µM. All samples were allowed to incubate at room temperature or at 37ºC for 2 hours prior to gel electrophoresis. In the presence of thrombin and KCl, the dual payloads produce a large aggregate structure instead of the expected disassembled payloads. B) Detection of aptamer complement DNA sequence with the standard toehold mediated strand displacement mechanism works as expected without formation of unexpected structures.*

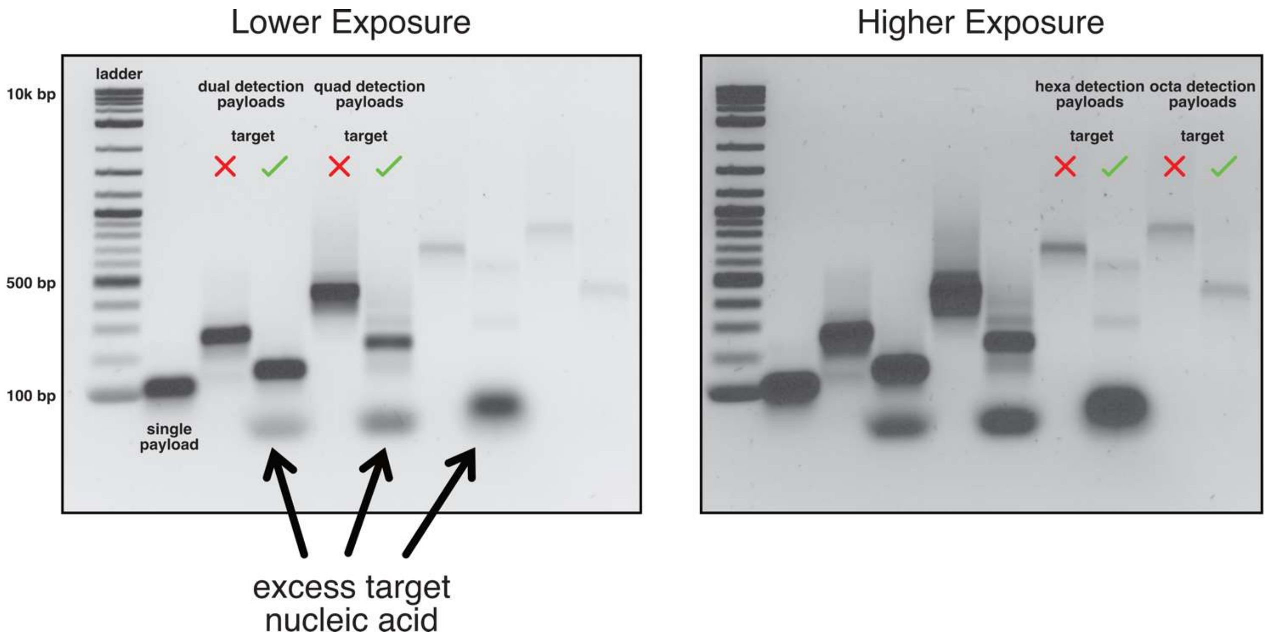


*Figure S8: Detection of various targets using different multi-payloads, all of which produce final output bands at different locations. The two images are identical, save for exposure time when imaging, in order to allow clearer definition of both high concentration and low concentration bands. Quad and dual detection payloads are detecting BRCA1. Hexa detection payloads are detecting miR-141 RNA. Octa detection payloads are detecting the thrombin aptamer complement. All targets are added in excess.*

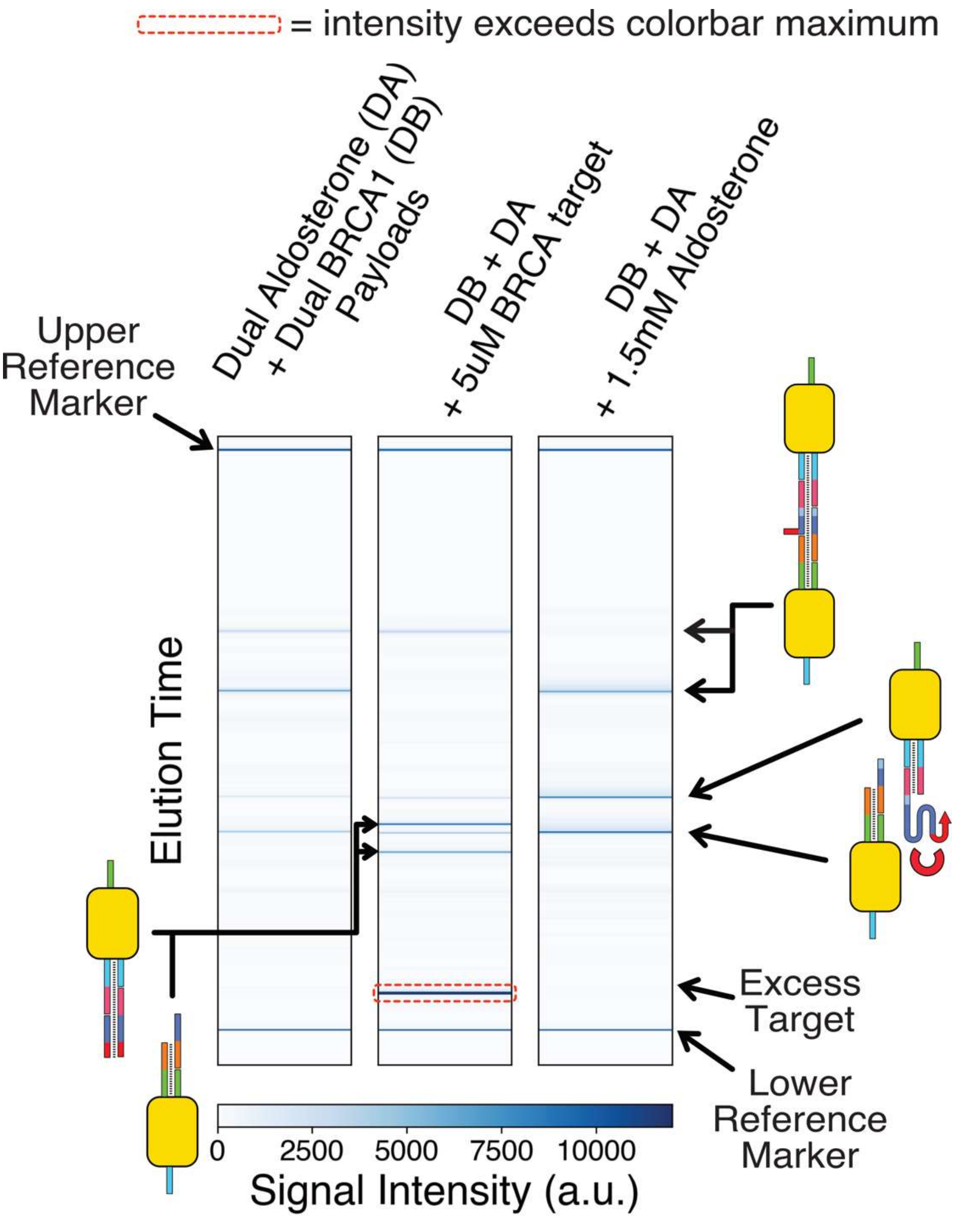


*Figure S9: Result showing that dual aldosterone and BRCA1 payload detection chains can be resolved separately in capillary electrophoresis, allowing for multiplexed detection without the need for multi-payload detection chains.*

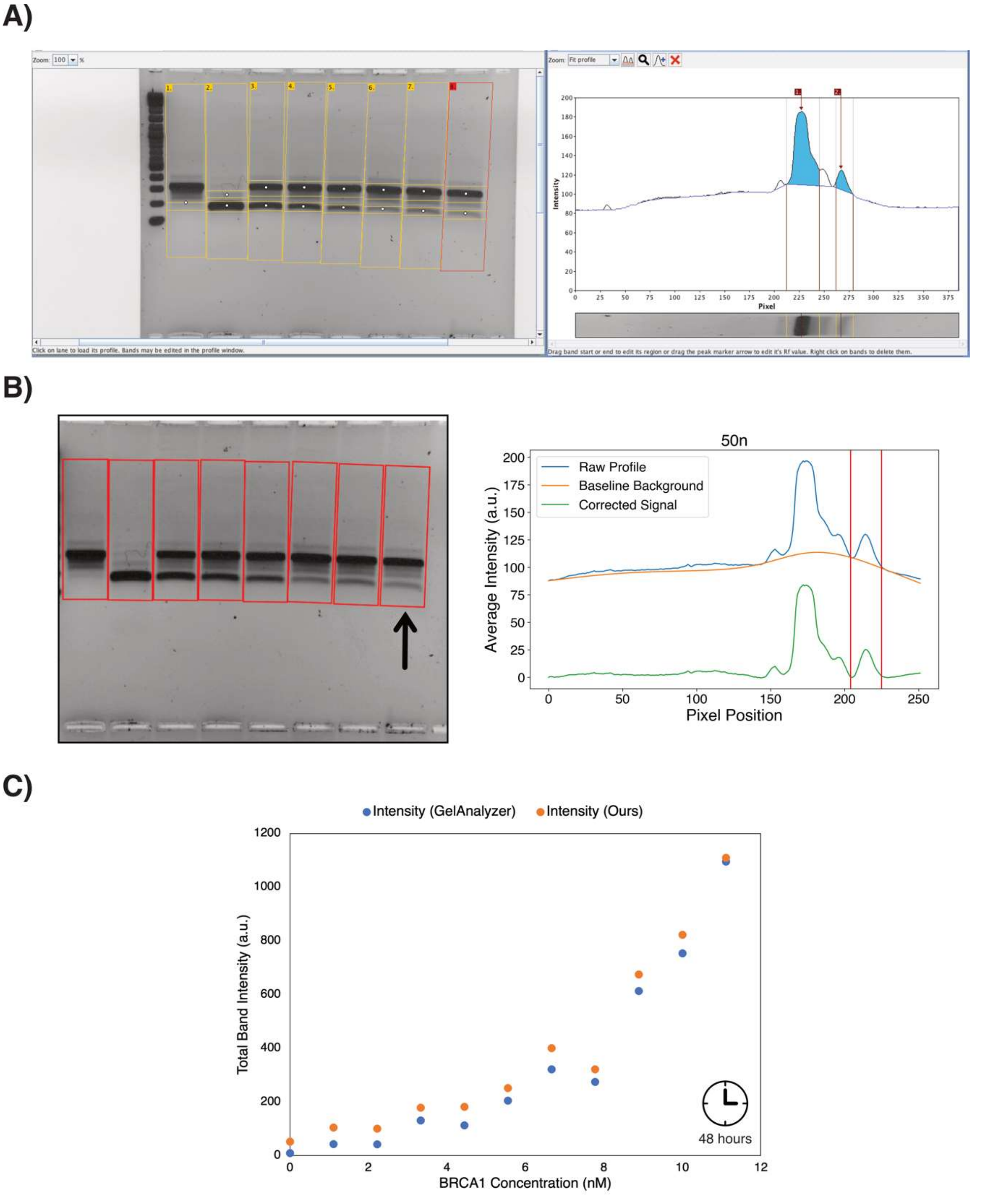


*Figure S10: Comparison between GelAnalyzer image analysis results and our method on the 48-hour BRCA1 incubation gel. A) GelAnalyzer uses a rolling ball background detection method, which can be inaccurate and will require manual tuning for each lane profile. Additionally, it is not possible to rotate lane orientation within GelAnalyzer, which results in bands with incomplete coverage. B) Our method uses baseline correction with*

*asymmetric least squares smoothing, which we found to produce good results for all lane profiles without any manual adjustment required. We use the parameters lambda =1000, p=0.001, and number of iterations =10 for all of our gels. The left image shows the placement of our lanes. We force the lanes to have the same width and height, but allow free rotation. The right image shows the background correction algorithm for the lane selected (identical to the one selected in GelAnalyzer). C) Direct comparison of intensity results from GelAnalyzer and our method. While both produce the same profile shape, GelAnalyzer tends to underestimate the intensity of each band.*
