## Supplementary material for "Multiplexed Label-Free Biomarker Detection by Targeted Disassembly of Variable-Length DNA Payload Chains": Underlying data - gel images etc: report.pdf

### Instrument controller software run summary:

**Filename and data path:** C:\Agilent Technologies\Data\2021 10 11\final_detection_test 14-11-44\2021 10 11 14H 11M.raw
**Created:** Monday, 11 October 2021 14:41:52
**Number of capillaries:** 96
**Array serial number:** 102819-03LFS
**Effect length:** 55 cm
**Array usage count:** 20
**Instrument type:** 5400 Fragment Analyzer
**Instrument controller software version:** 3.1.0.12
**Device serial number:** MY2114AC03

#### Method Information

**Method name:** DNF-905-55 - DNA 1-500bp.mthds
**Gel prime:** No
**Full conditioning:** Yes
**Gel prime to buffer:** No
**Gel selection:** Gel 2
**Perform prerun:** 9.0 kV, 30 sec.
**Rinse:** No
**Marker 1:** Row: A, 7.5 kV, 5 sec.
**Rinse:** No
**Sample injection:** 7.5 kV, 5 sec.
**Separation:** 9.0 kV, 80.0 min.
**Tray name:** final_detection_test

**Analysis mode:** DNA

**Notes**

### Gel Image

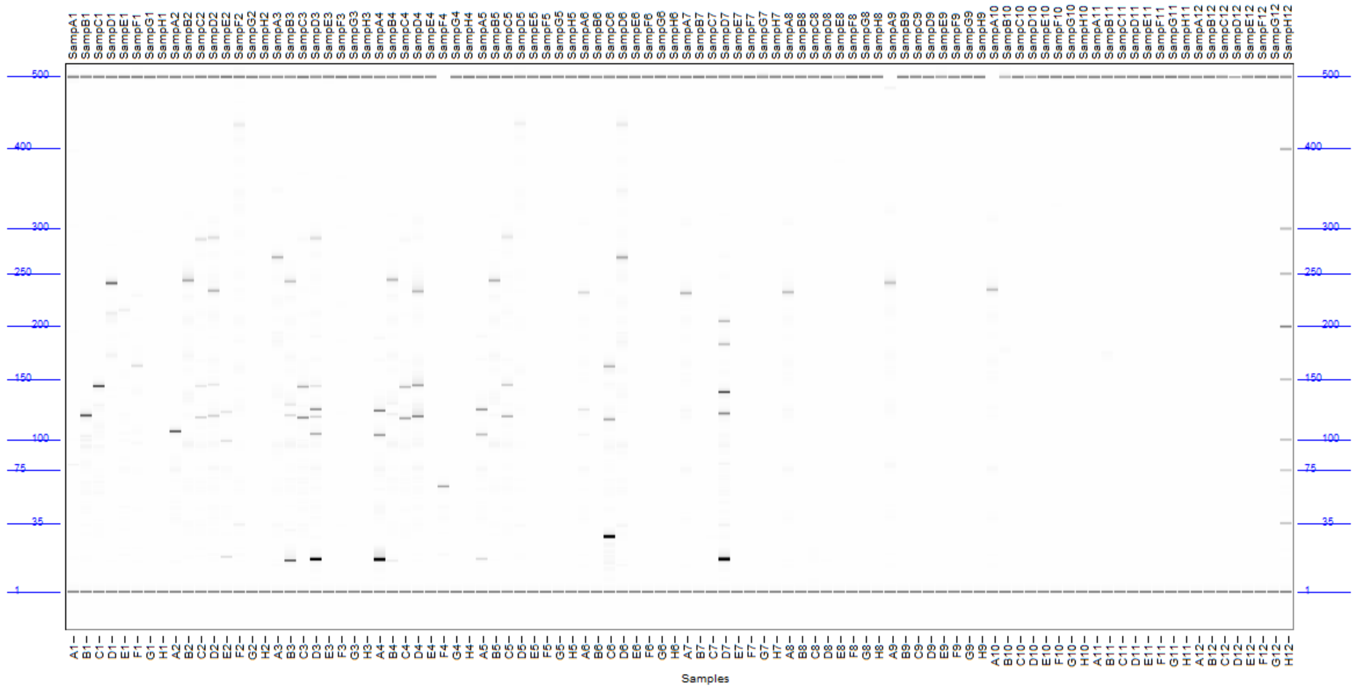
SampA1 SampB1 SampC1 SampD1 SampE1 SampF1 SampG1 SampH1 SampA2 SampB2 SampC2 SampD2 SampE2 SampF2 SampG2 SampH2 SampA3 SampB3 SampC3 SampD3 SampE3 SampF3 SampG3 SampH3 SampA4 SampB4 SampC4 SampD4 SampE4 SampF4 SampG4 SampH4 SampA5 SampB5 SampC5 SampD5 SampE5 SampF5 SampG5 SampH5 SampA6 SampB6 SampC6 SampD6 SampE6 SampF6 SampG6 SampH6 SampA7 SampB7 SampC7 SampD7 SampE7 SampF7 SampG7 SampH7 SampA8 SampB8 SampC8 SampD8 SampE8 SampF8 SampG8 SampH8 SampA9 SampB9 SampC9 SampD9 SampE9 SampF9 SampG9 SampH9 SampA10 SampB10 SampC10 SampD10 SampE10 SampF10 SampG10 SampH10 SampA11 SampB11 SampC11 SampD11 SampE11 SampF11 SampG11 SampH11 SampA12 SampB12 SampC12 SampD12 SampE12 SampF12 SampG12 SampH12
500
400
300
250
200
150
100
75
35
1
A1 B1 C1 D1 E1 F1 G1 H1 A2 B2 C2 D2 E2 F2 G2 H2 A3 B3 C3 D3 E3 F3 G3 H3 A4 B4 C4 D4 E4 F4 G4 H4 A5 B5 C5 D5 E5 F5 G5 H5 A6 B6 C6 D6 E6 F6 G6 H6 A7 B7 C7 D7 E7 F7 G7 H7 A8 B8 C8 D8 E8 F8 G8 H8 A9 B9 C9 D9 E9 F9 G9 H9 A10 B10 C10 D10 E10 F10 G10 H10 A11 B11 C11 D11 E11 F11 G11 H11 A12 B12 C12 D12 E12 F12 G12 H12
Samples

**Filename and data path:** C:\Agilent Technologies\Data\2021 10 11\final_detection_test 14-11-44\2021 10 11 14H 11M.raw

A1: SampA1

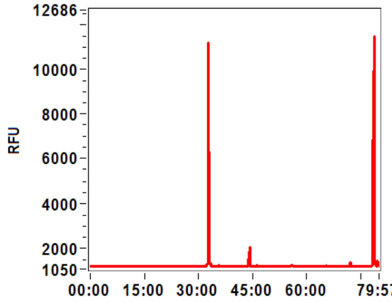


B1: SampB1

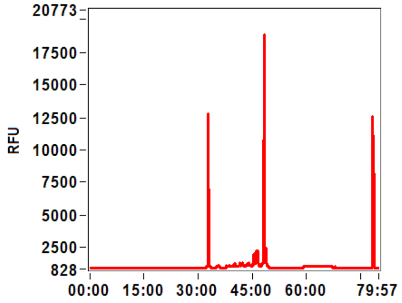


C1: SampC1

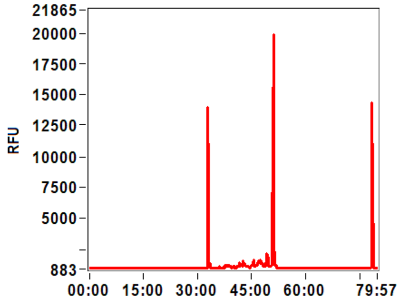


D1: SampD1

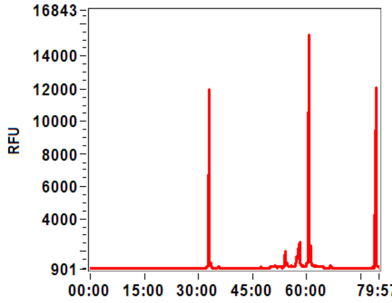


E1: SampE1

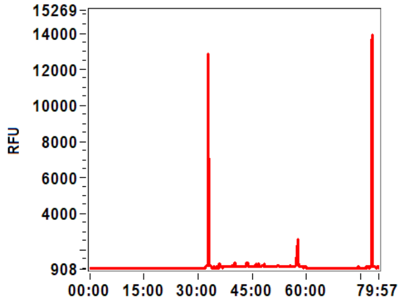


F1: SampF1

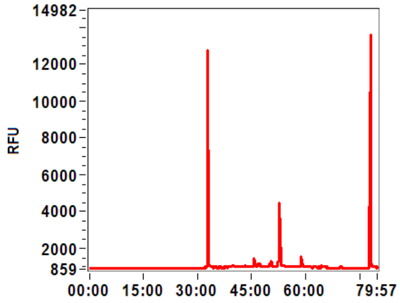


G1: SampG1

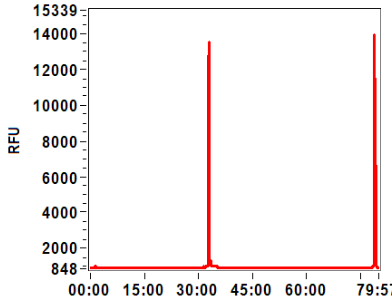


H1: SampH1

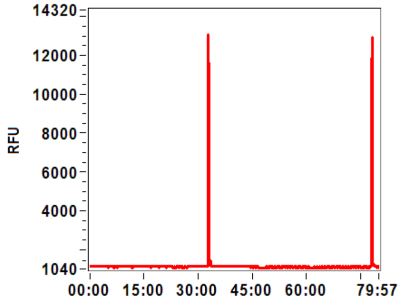


A2: SampA2

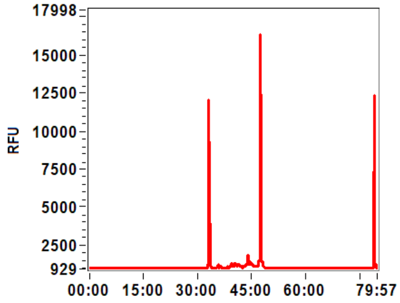


B2: SampB2

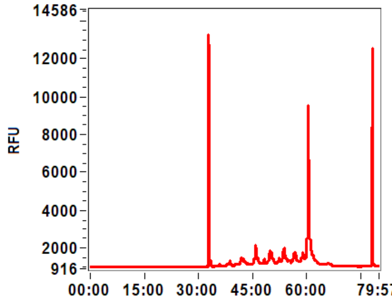


C2: SampC2

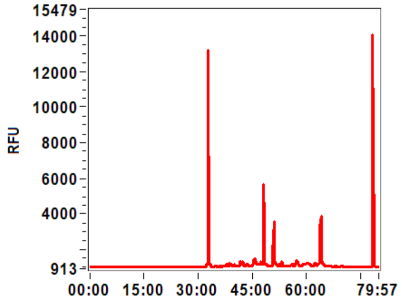


D2: SampD2

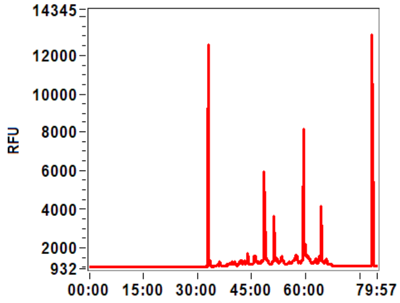

**Filename and data path:** C:\Agilent Technologies\Data\2021 10 11\final_detection_test 14-11-44\2021 10 11 14H 11M.raw

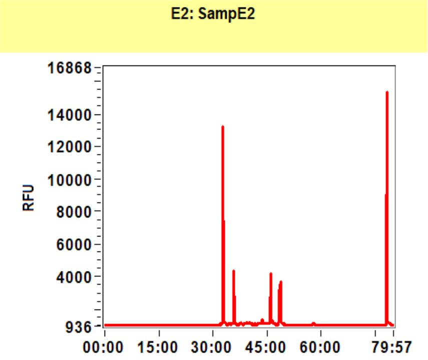


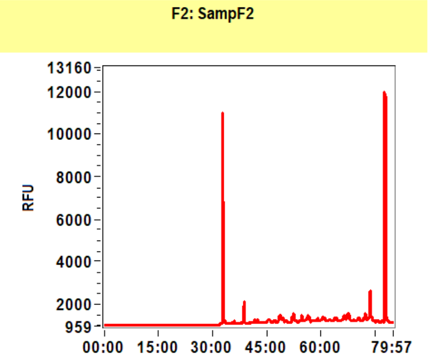


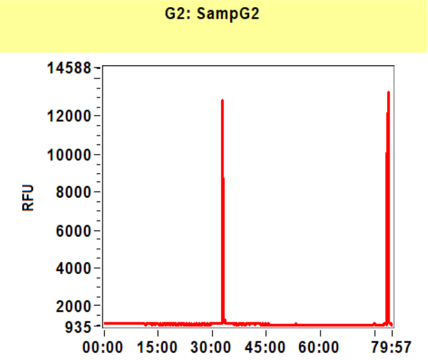


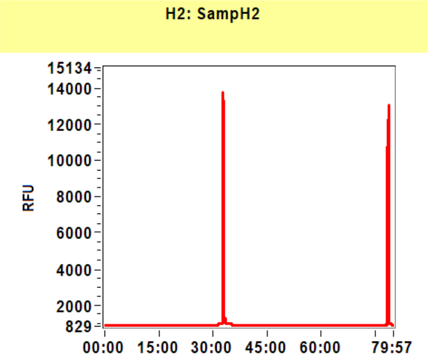


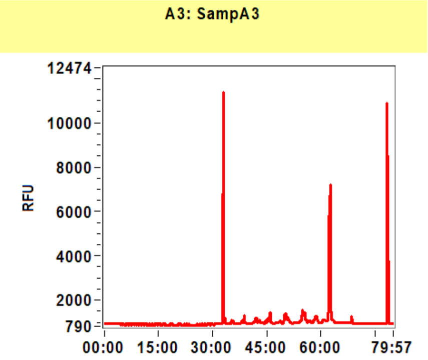


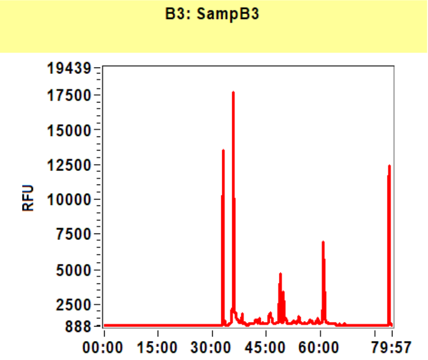


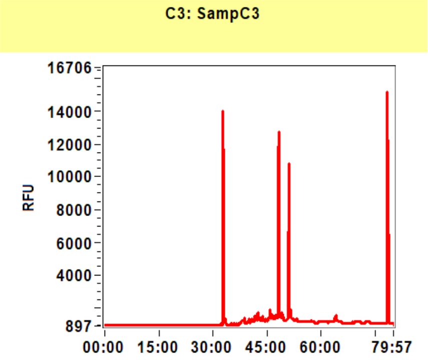


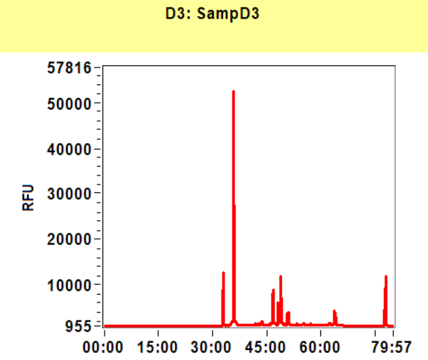


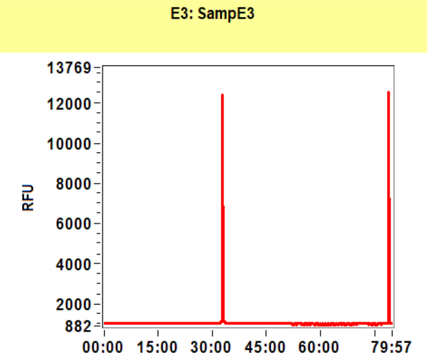


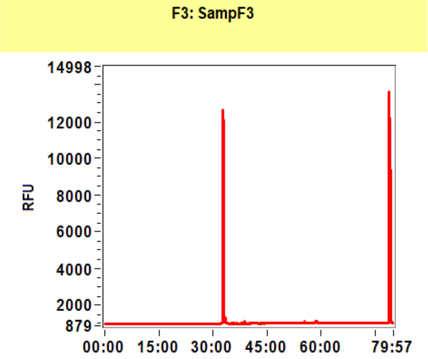


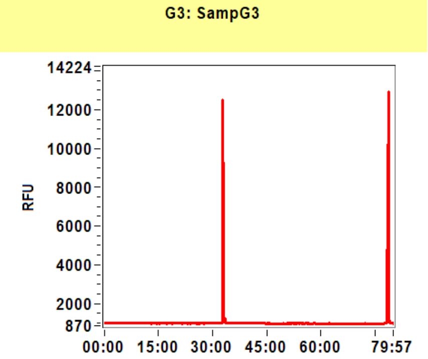


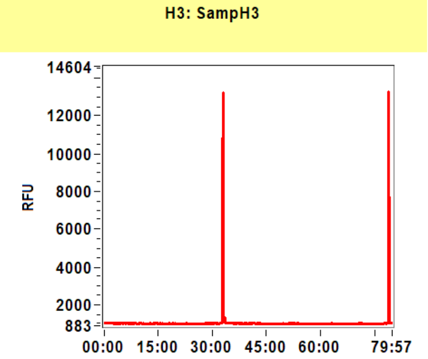

**Filename and data path:** C:\Agilent Technologies\Data\2021 10 11\final_detection_test 14-11-44\2021 10 11 14H 11M.raw

A4: SampA4

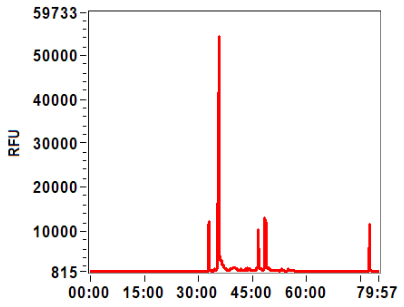


B4: SampB4

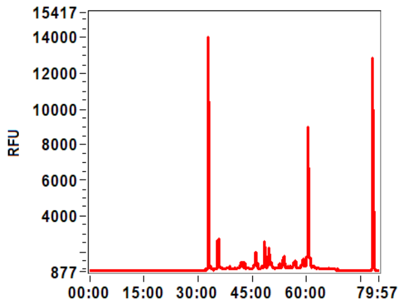


C4: SampC4

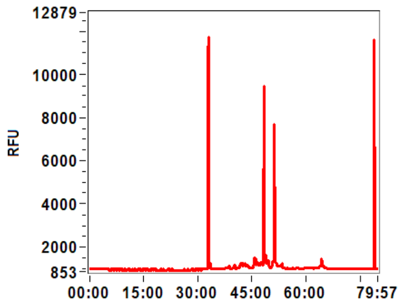


D4: SampD4

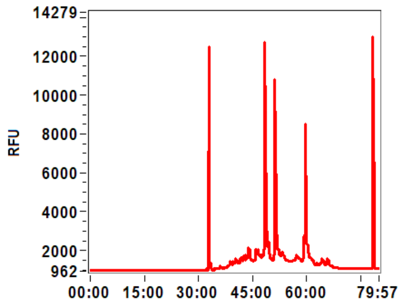


E4: SampE4

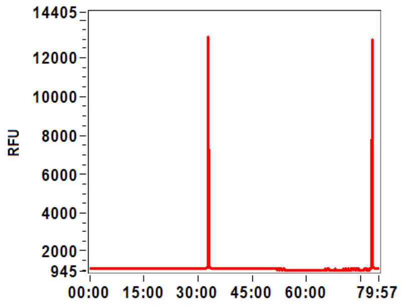


F4: SampF4

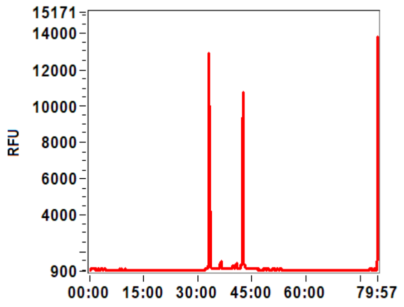


G4: SampG4

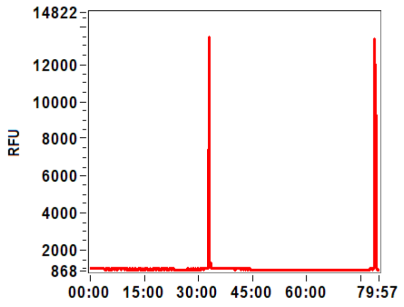


H4: SampH4

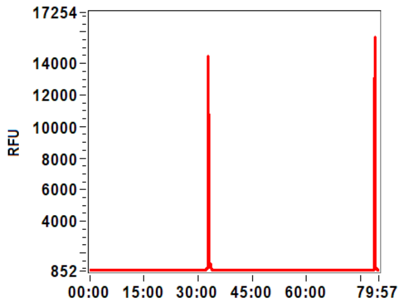


A5: SampA5

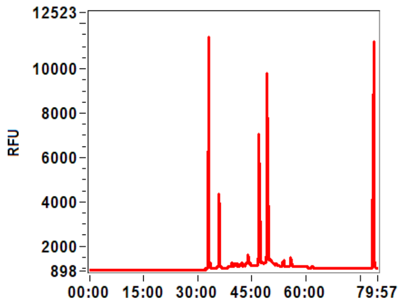


B5: SampB5

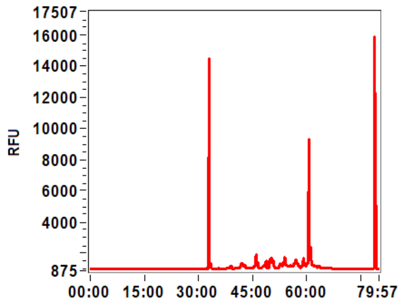


C5: SampC5

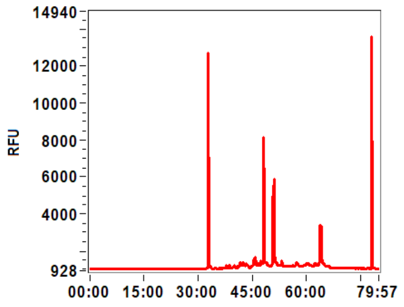


D5: SampD5

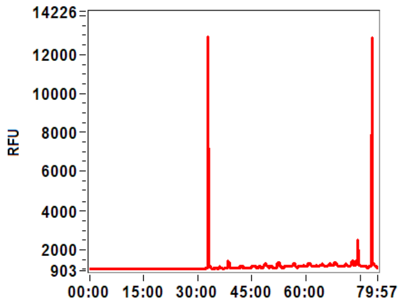

**Filename and data path:** C:\Agilent Technologies\Data\2021 10 11\final_detection_test 14-11-44\2021 10 11 14H 11M.raw

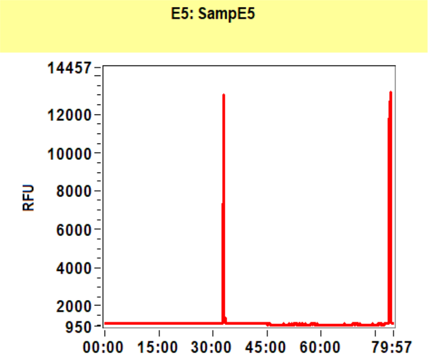


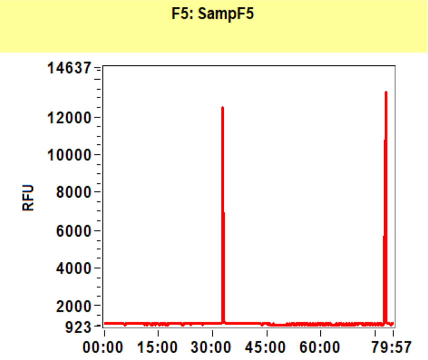


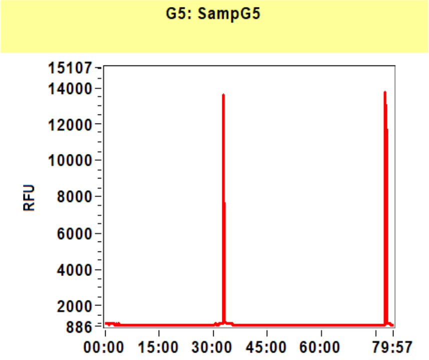


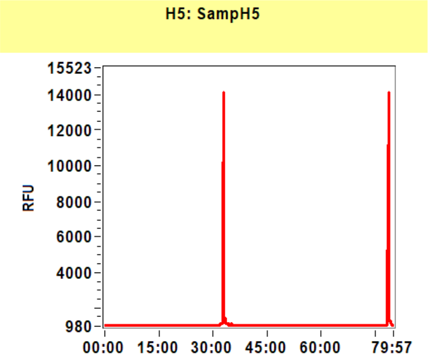


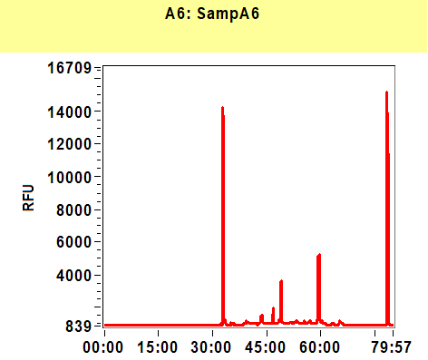


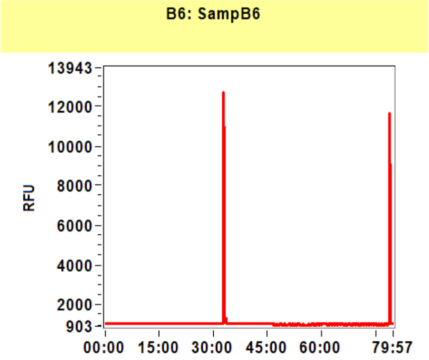


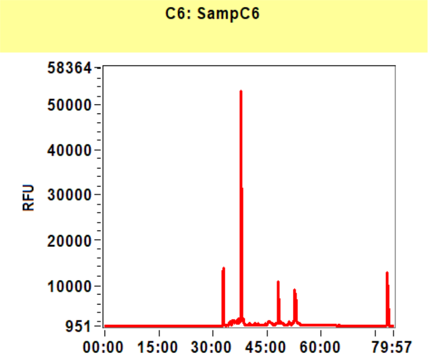


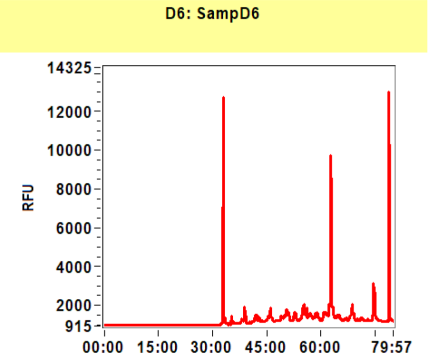


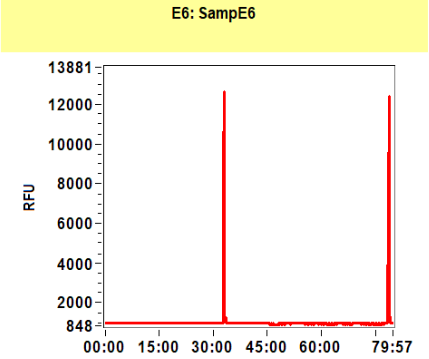


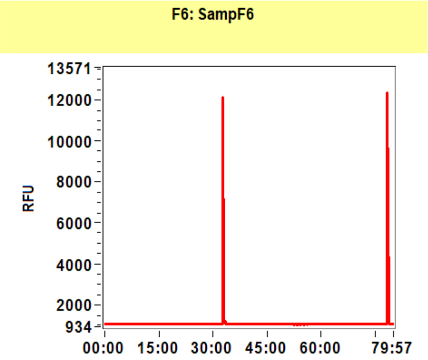


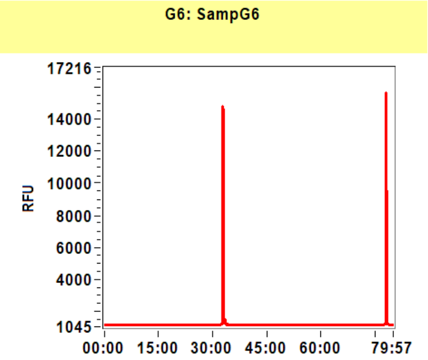


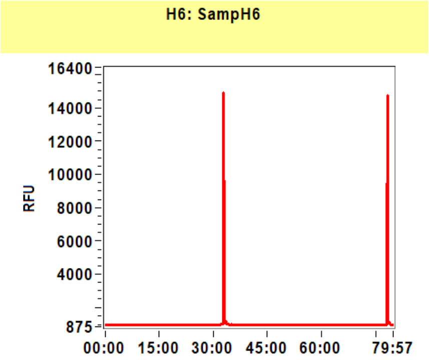

**Filename and data path:** C:\Agilent Technologies\Data\2021 10 11\final_detection_test 14-11-44\2021 10 11 14H 11M.raw

A7: SampA7

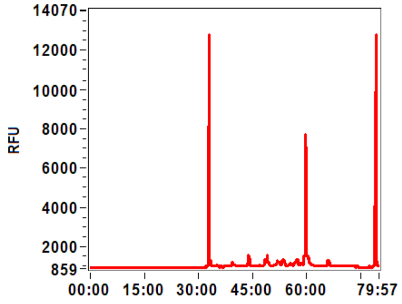


B7: SampB7

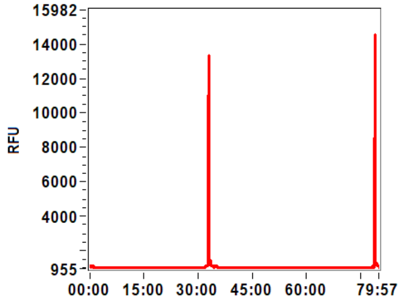


C7: SampC7

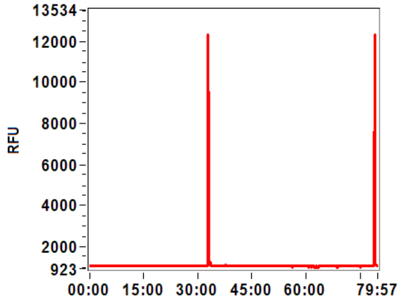


D7: SampD7

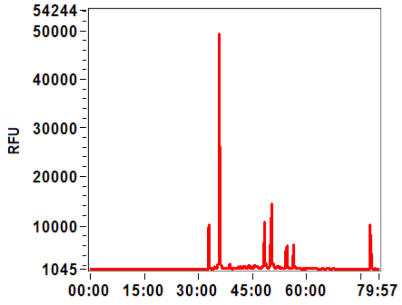


E7: SampE7

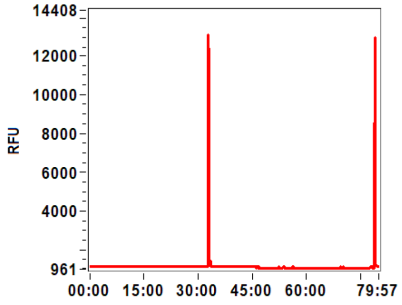


F7: SampF7

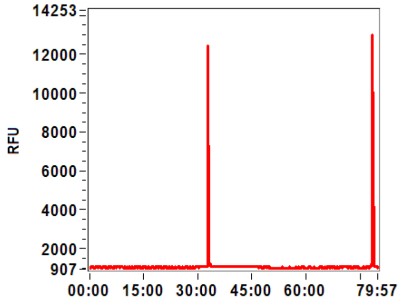


G7: SampG7

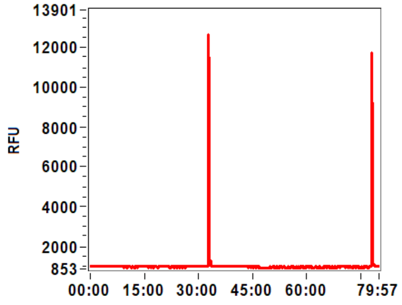


H7: SampH7

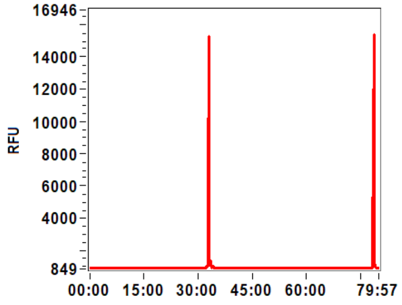


A8: SampA8

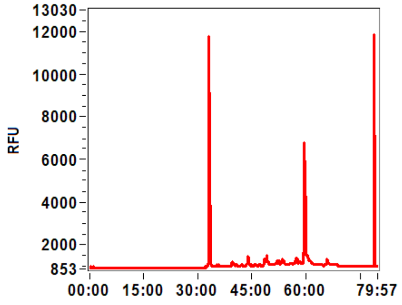


B8: SampB8

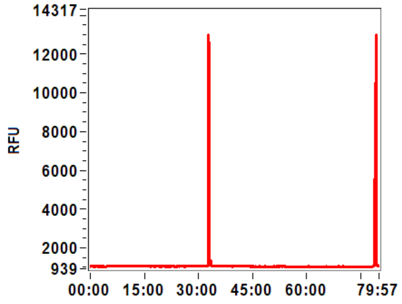


C8: SampC8

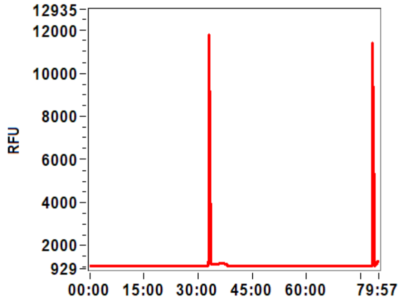


D8: SampD8

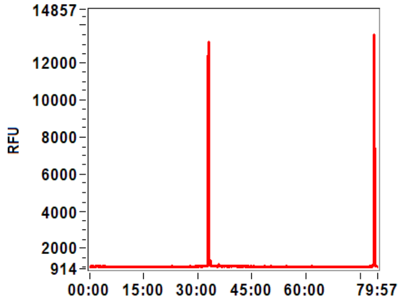

**Filename and data path:** C:\Agilent Technologies\Data\2021 10 11\final_detection_test 14-11-44\2021 10 11 14H 11M.raw

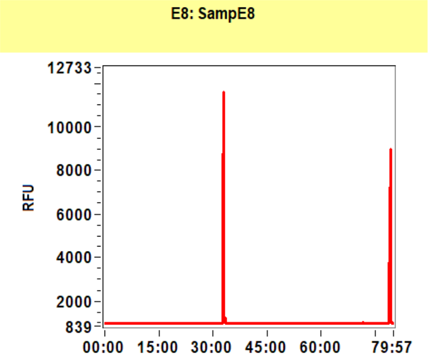


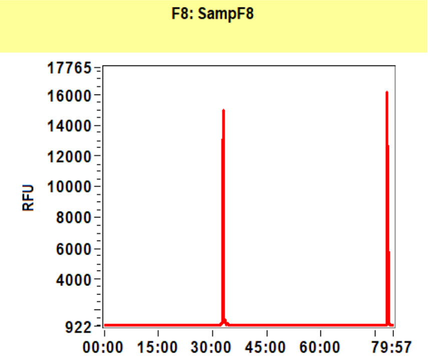


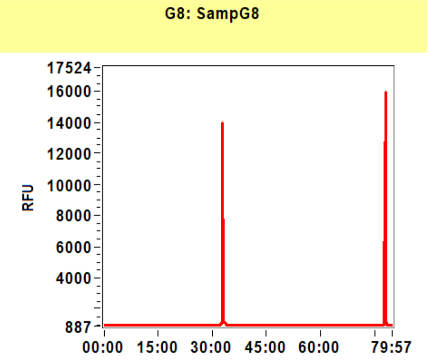


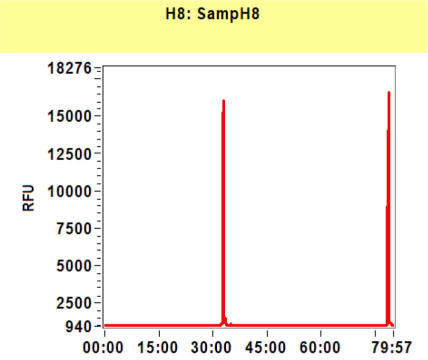


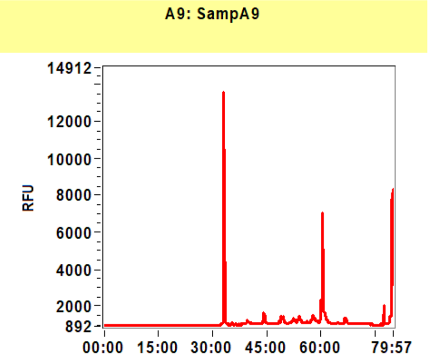


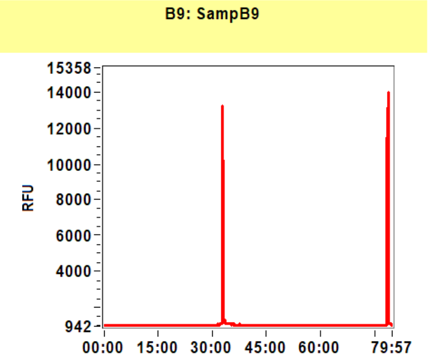


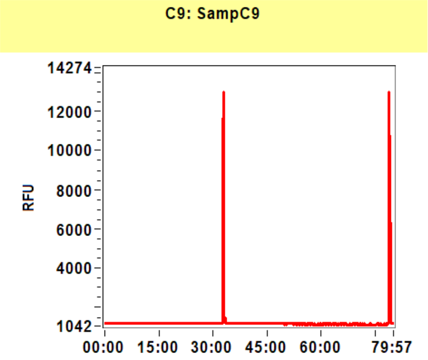


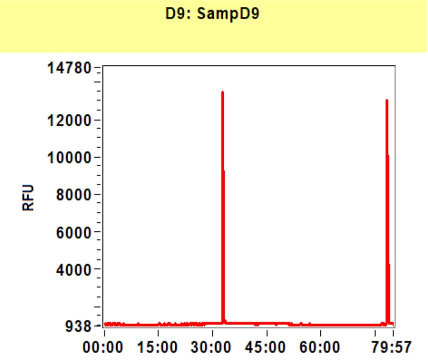


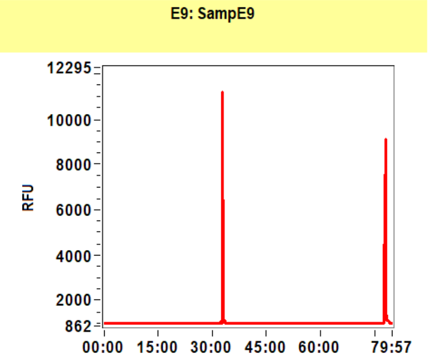


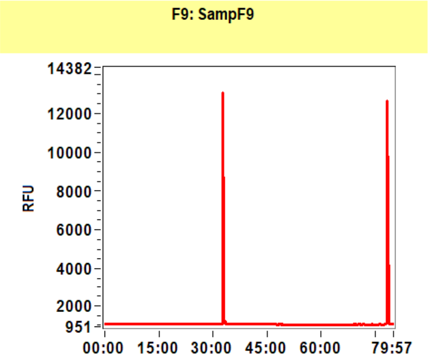


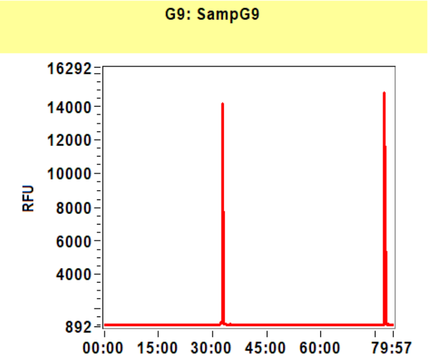


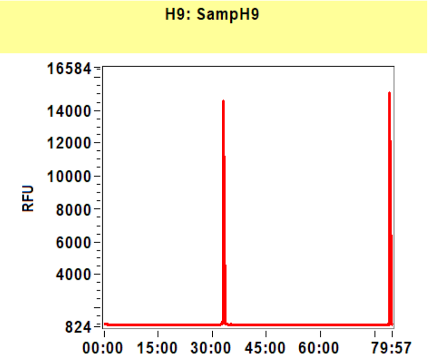

**Filename and data path:** C:\Agilent Technologies\Data\2021 10 11\final_detection_test 14-11-44\2021 10 11 14H 11M.raw

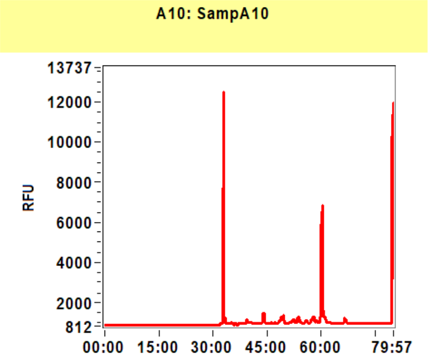


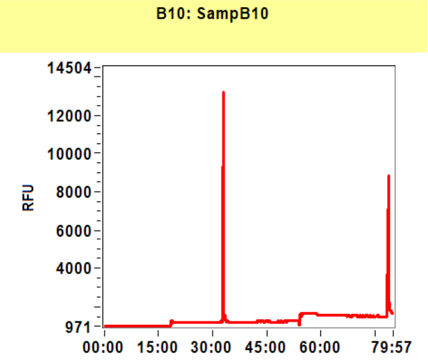


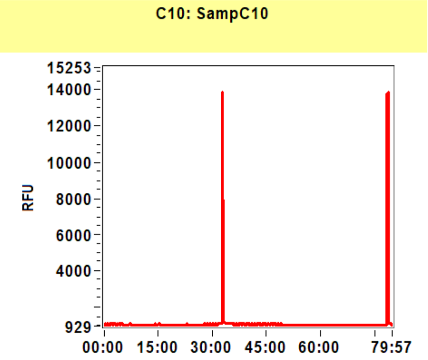


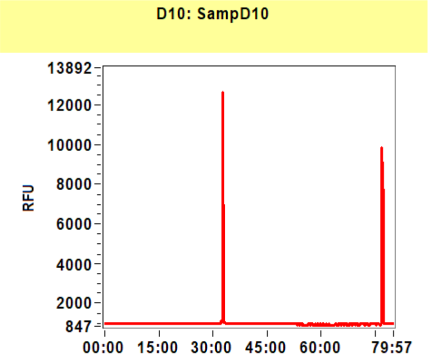


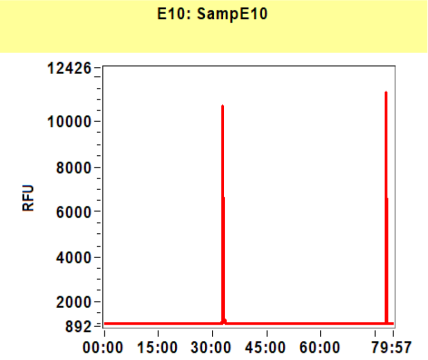


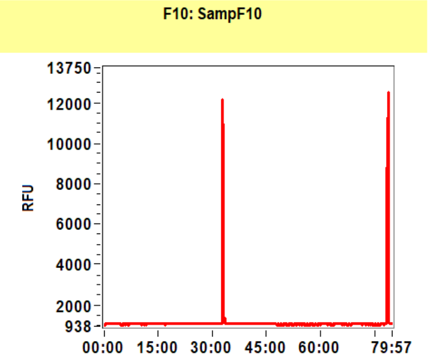


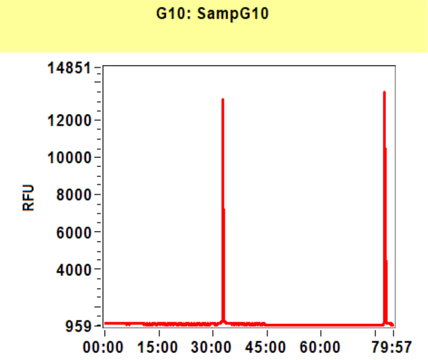


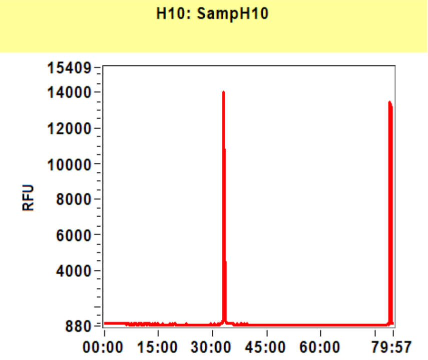


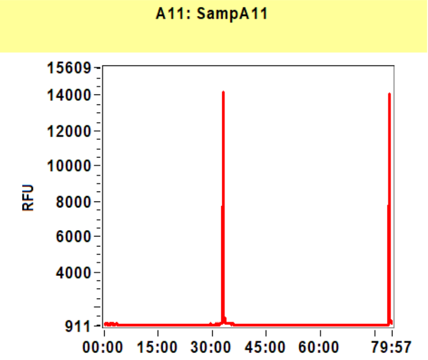


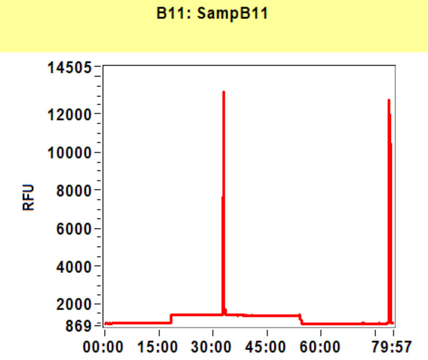


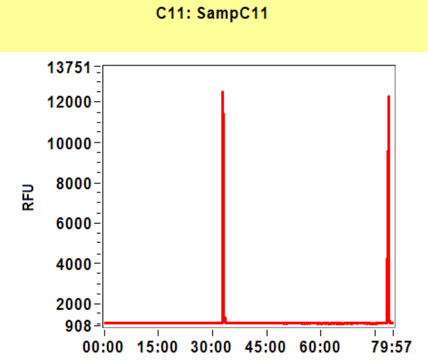


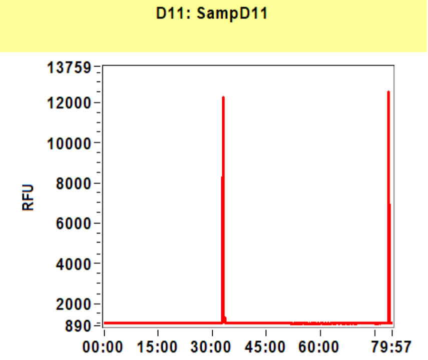

**Filename and data path:** C:\Agilent Technologies\Data\2021 10 11\final_detection_test 14-11-44\2021 10 11 14H 11M.raw

E11: SampE11

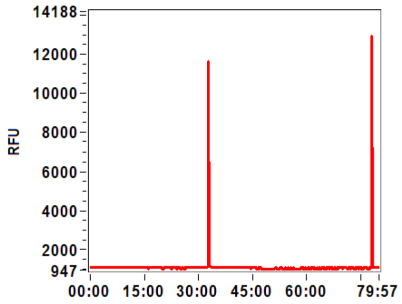


F11: SampF11

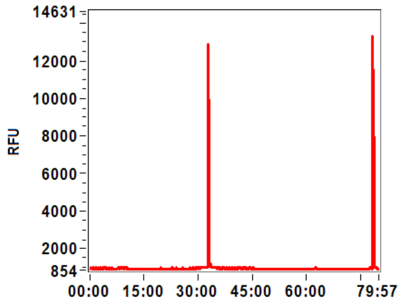


G11: SampG11

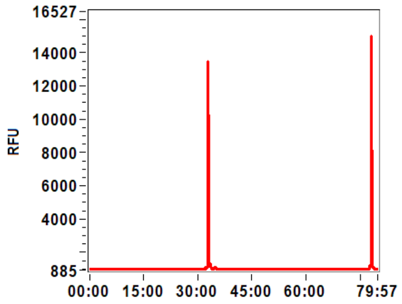


H11: SampH11

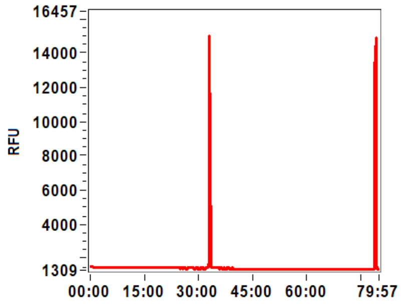


A12: SampA12

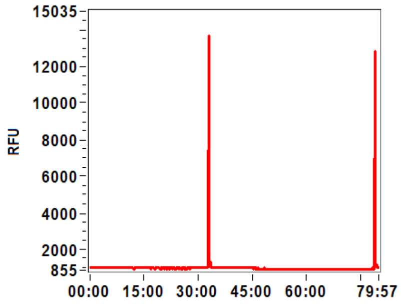


B12: SampB12

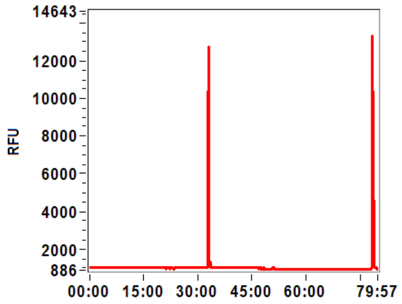


C12: SampC12

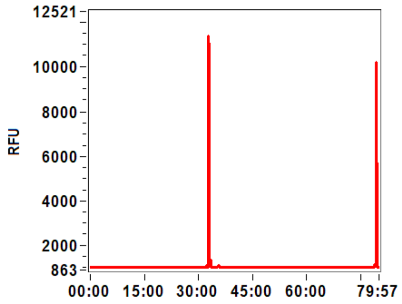


D12: SampD12

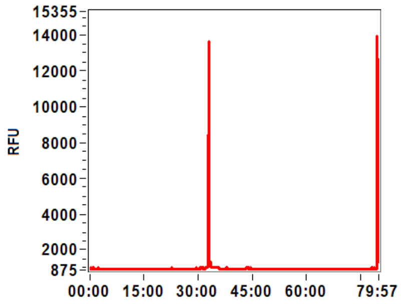


E12: SampE12

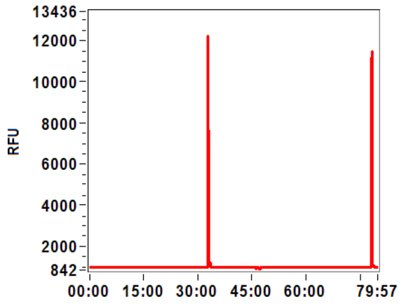


F12: SampF12

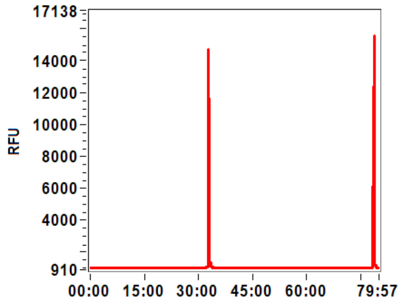


G12: SampG12

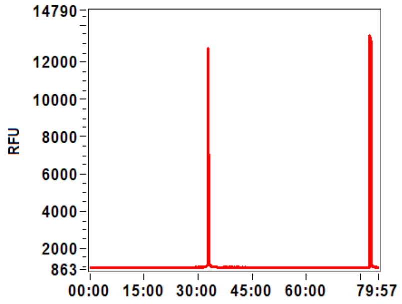


H12: SampH12

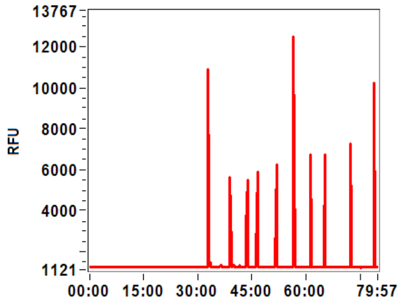

**Sample:** SampA1
**Well location:** A1
**Created:** Monday, 11 October 2021 14:41:52

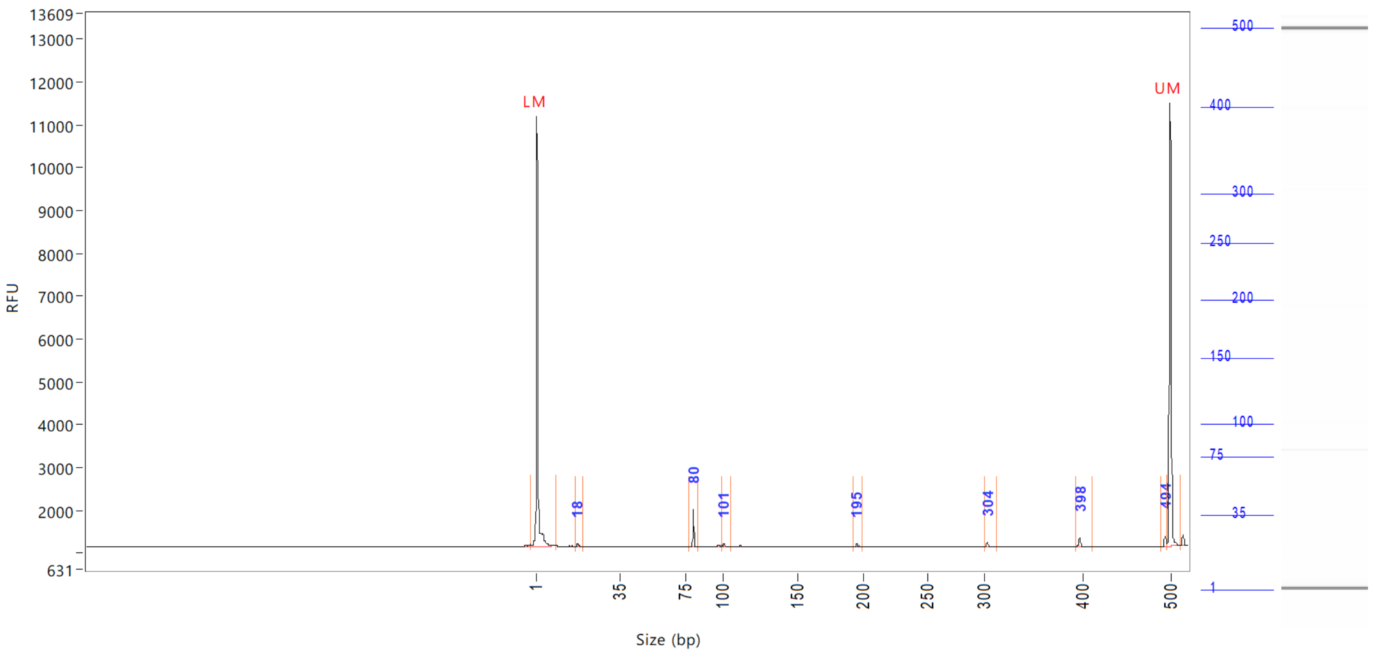


| Peak | Size (bp) | Concentration (ng/uL) | From (bp) | To (bp) | Average size (bp) | CV% | RFU | Corrected peak area |
|---|---|---|---|---|---|---|---|---|
| 1 | 1 (LM) | 0.9464 | 0 | 10 | 2 | 71.13 | 10035 | 40.952 |
| 2 | 18 | 0.0712 | 17 | 20 | 18 | 3.00 | 55 | 0.257 |
| 3 | 80 | 0.4342 | 78 | 83 | 80 | 0.45 | 857 | 1.566 |
| 4 | 101 | 0.0509 | 99 | 105 | 101 | 1.21 | 57 | 0.183 |
| 5 | 195 | 0.0651 | 193 | 199 | 196 | 0.58 | 78 | 0.235 |
| 6 | 304 | 0.0788 | 301 | 313 | 305 | 0.57 | 111 | 0.284 |
| 7 | 398 | 0.1373 | 394 | 412 | 398 | 0.55 | 199 | 0.495 |
| 8 | 494 | 0.1511 | 490 | 497 | 494 | 0.22 | 250 | 0.545 |
| 9 | 500 (UM) | 0.5000 | 497 | 512 | 500 | 0.25 | 10362 | 21.635 |

| | | |
|---|---|---|
| TIC: | 0.9886 | ng/uL |
| TIM: | 18.1980 | nmole/L |
| Total concentration: | 1.5680 | ng/uL |

Sample peak width (sec): 5 Sample min peak height: 50 Sample baseline V to V?: Y Sample baseline V to V points: 3
Sample filter: Binomial Number of points for filter: 3 Sample start region (min): 0 Sample end region (min): 80
Marker peak width (sec): 5 Marker min peak height: 500 Marker baseline V to V?: Y Marker baseline V to V points: 3
Lower marker selection: First peak > 500 RFU Upper marker selection: Last peak > 500 RFU
Ladder size (bp)1, 35, 75, 100, 150, 200, 250, 300, 400, 500
Quantification using: Upper Marker Final concentration (ng/uL): 0.5000 Dilution factor: 12.0

**Sample:** SampB1
**Well location:** B1
**Created:** Monday, 11 October 2021 14:41:52

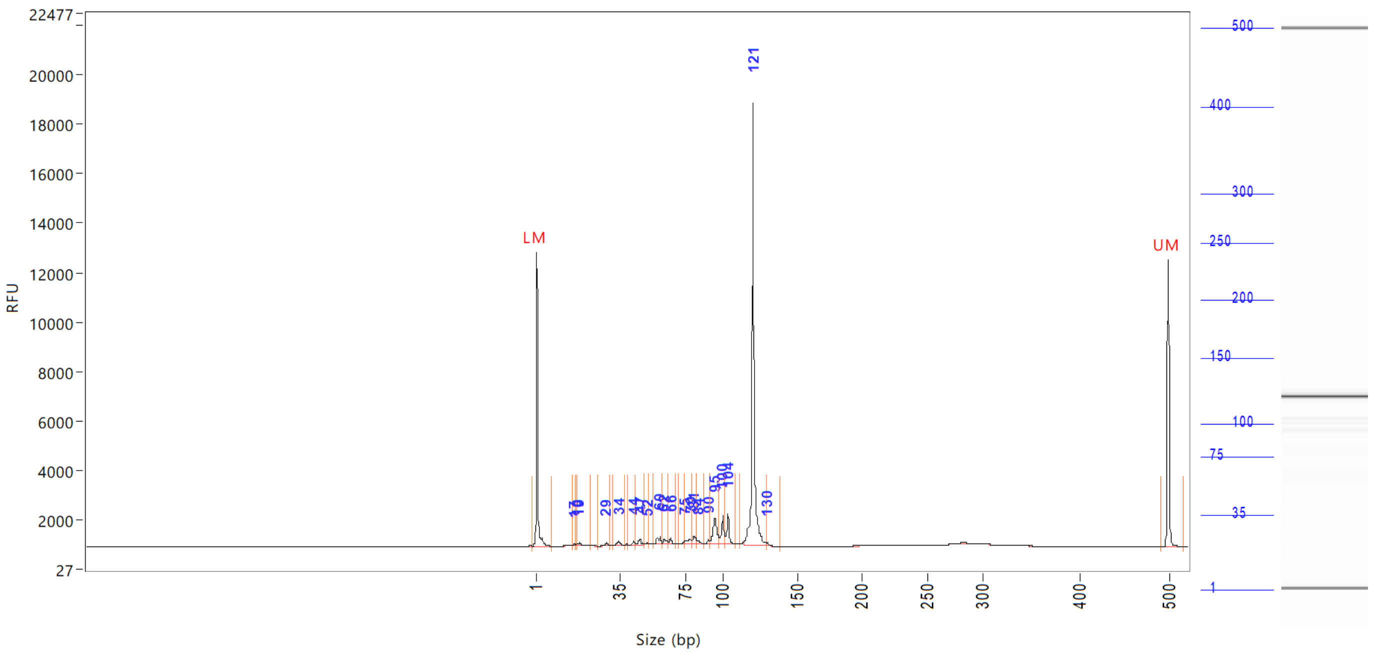


| Peak | Size (bp) | Concentration (ng/uL) | From (bp) | To (bp) | Average size (bp) | CV% | RFU | Corrected peak area |
|---|---|---|---|---|---|---|---|---|
| 1 | 1 (LM) | 0.9128 | 0 | 8 | 2 | 65.57 | 11856 | 45.234 |
| 2 | 17 | 0.0661 | 16 | 17 | 17 | 1.80 | 65 | 0.273 |
| 3 | 18 | 0.0991 | 17 | 18 | 18 | 1.58 | 104 | 0.409 |
| 4 | 19 | 0.2146 | 18 | 23 | 19 | 5.10 | 136 | 0.886 |
| 5 | 29 | 0.2126 | 26 | 31 | 29 | 2.60 | 135 | 0.878 |
| 6 | 34 | 0.2994 | 32 | 38 | 34 | 3.65 | 164 | 1.236 |
| 7 | 44 | 0.1913 | 40 | 45 | 43 | 2.05 | 162 | 0.790 |
| 8 | 47 | 0.3374 | 45 | 50 | 47 | 2.21 | 217 | 1.393 |
| 9 | 52 | 0.0809 | 50 | 53 | 51 | 1.22 | 67 | 0.334 |
| 10 | 60 | 0.5510 | 56 | 61 | 58 | 1.85 | 307 | 2.275 |
| 11 | 62 | 0.3409 | 61 | 64 | 63 | 1.53 | 230 | 1.408 |
| 12 | 66 | 0.3272 | 64 | 69 | 66 | 1.24 | 281 | 1.351 |
| 13 | 75 | 0.1346 | 71 | 75 | 74 | 1.14 | 136 | 0.556 |
| 14 | 79 | 0.4270 | 75 | 80 | 78 | 1.60 | 206 | 1.763 |
| 15 | 81 | 0.5431 | 80 | 83 | 81 | 1.15 | 340 | 2.242 |
| 16 | 84 | 0.1730 | 83 | 87 | 85 | 1.02 | 139 | 0.714 |
| 17 | 90 | 0.1873 | 87 | 91 | 90 | 0.91 | 173 | 0.773 |
| 18 | 95 | 1.9869 | 91 | 98 | 95 | 1.48 | 1049 | 8.205 |
| 19 | 100 | 1.2559 | 98 | 102 | 100 | 1.24 | 1134 | 5.186 |
| 20 | 104 | 1.1177 | 102 | 109 | 104 | 0.84 | 1216 | 4.615 |
| 21 | 121 | 16.0891 | 113 | 130 | 121 | 1.54 | 17882 | 66.438 |
| 22 | 130 | 0.1307 | 130 | 139 | 131 | 0.90 | 117 | 0.540 |
| 23 | 500 (UM) | 0.5000 | 492 | 0 | 500 | 0.28 | 11586 | 24.776 |
| | TIC: | 24.7658 | ng/uL | | | | | |

```
Sample peak width (sec): 5     Sample min peak height: 50    Sample baseline V to V?: Y    Sample baseline V to V points: 3
Sample filter: Binomial        Number of points for filter: 3  Sample start region (min): 0  Sample end region (min): 80
Marker peak width (sec): 5     Marker min peak height: 500   Marker baseline V to V?: Y    Marker baseline V to V points: 3
Lower marker selection: First peak > 500 RFU                 Upper marker selection: Last peak > 500 RFU
Ladder size (bp)1, 35, 75, 100, 150, 200, 250, 300, 400, 500
Quantification using: Upper Marker          Final concentration (ng/uL): 0.5000          Dilution factor: 12.0
```

**Sample:** SampB1
**Well location:** B1
**Created:** Monday, 11 October 2021 14:41:52

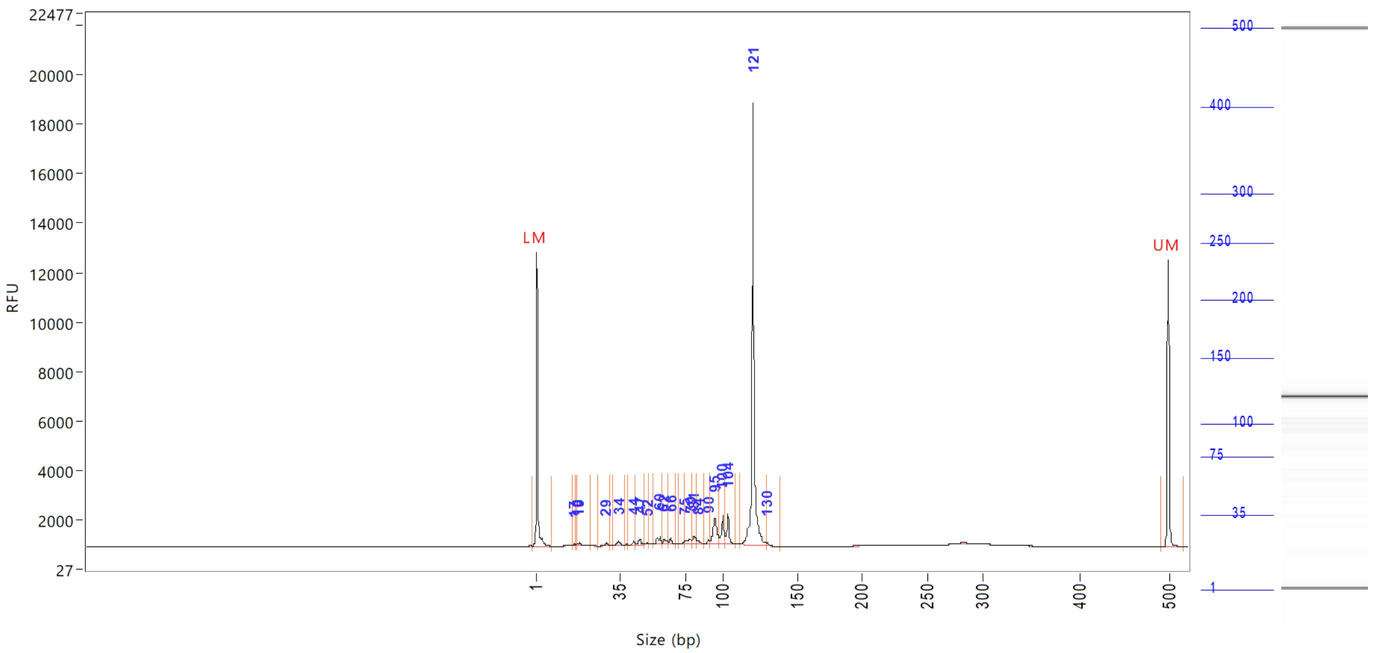


| Peak | Size (bp) | Concentration (ng/uL) | From (bp) | To (bp) | Average size (bp) | CV% | RFU | Corrected peak area |
|---|---|---|---|---|---|---|---|---|

| | | |
|---|---|---|
| TIM: | 436.5648 | nmole/L |
| Total concentration: | 25.2372 | ng/uL |

Sample peak width (sec): 5 Sample min peak height: 50 Sample baseline V to V?: Y Sample baseline V to V points: 3
Sample filter: Binomial Number of points for filter: 3 Sample start region (min): 0 Sample end region (min): 80
Marker peak width (sec): 5 Marker min peak height: 500 Marker baseline V to V?: Y Marker baseline V to V points: 3
Lower marker selection: First peak > 500 RFU Upper marker selection: Last peak > 500 RFU
Ladder size (bp)1, 35, 75, 100, 150, 200, 250, 300, 400, 500
Quantification using: Upper Marker Final concentration (ng/uL): 0.5000 Dilution factor: 12.0

**Sample:** SampC1
**Well location:** C1
**Created:** Monday, 11 October 2021 14:41:52

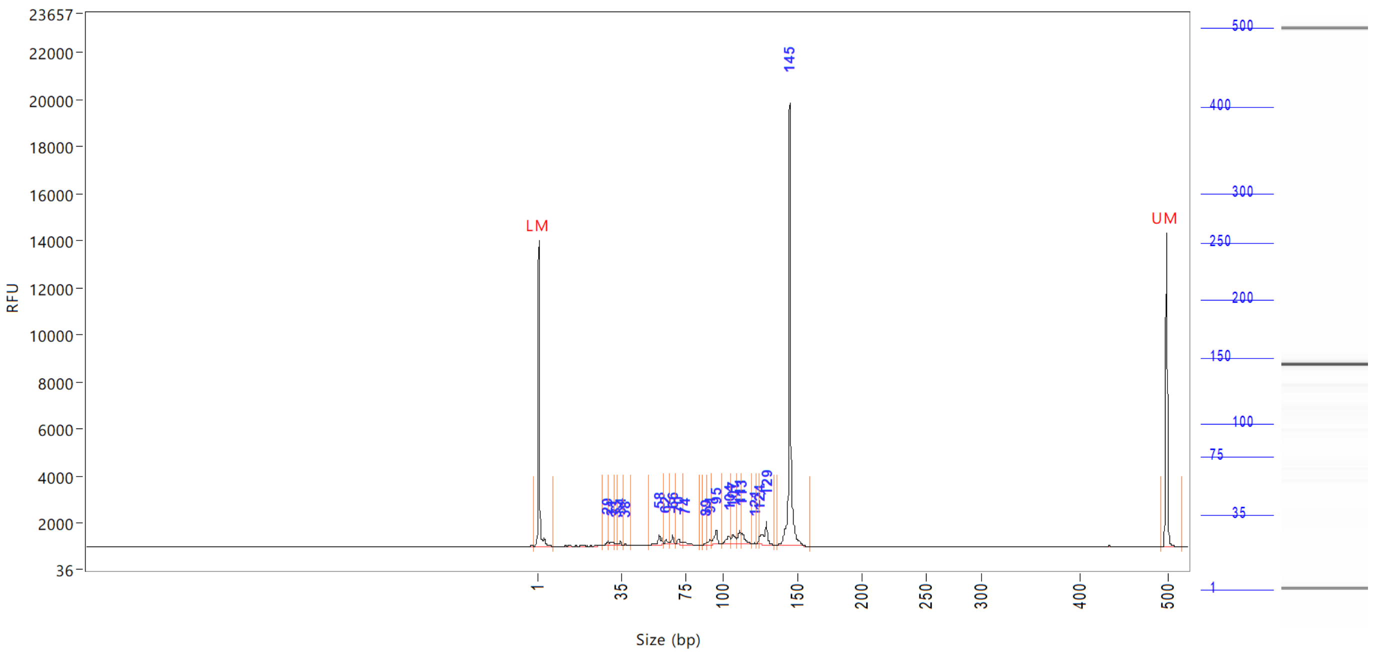

| Peak | Size (bp) | Concentration (ng/uL) | From (bp) | To (bp) | Average size (bp) | CV% | RFU | Corrected peak area |
|---|---|---|---|---|---|---|---|---|
| 1 | 1 (LM) | 0.8842 | 0 | 8 | 1 | 69.12 | 12981 | 48.172 |
| 2 | 29 | 0.1780 | 28 | 30 | 29 | 1.68 | 168 | 0.808 |
| 3 | 31 | 0.2761 | 30 | 32 | 31 | 1.90 | 149 | 1.254 |
| 4 | 33 | 0.0725 | 32 | 33 | 33 | 0.91 | 77 | 0.329 |
| 5 | 34 | 0.1831 | 33 | 36 | 34 | 1.31 | 164 | 0.831 |
| 6 | 38 | 0.0649 | 36 | 41 | 38 | 1.73 | 89 | 0.294 |
| 7 | 58 | 0.6673 | 52 | 61 | 58 | 3.08 | 407 | 3.029 |
| 8 | 62 | 0.2323 | 61 | 65 | 63 | 1.42 | 223 | 1.055 |
| 9 | 66 | 0.3375 | 65 | 68 | 66 | 1.03 | 418 | 1.532 |
| 10 | 70 | 0.3672 | 68 | 73 | 70 | 1.52 | 232 | 1.667 |
| 11 | 74 | 0.2473 | 73 | 84 | 75 | 2.98 | 130 | 1.123 |
| 12 | 89 | 0.1023 | 86 | 89 | 88 | 0.83 | 102 | 0.465 |
| 13 | 91 | 0.2447 | 89 | 92 | 91 | 0.79 | 228 | 1.111 |
| 14 | 95 | 0.8124 | 92 | 98 | 94 | 1.13 | 590 | 3.688 |
| 15 | 104 | 0.5831 | 98 | 105 | 103 | 1.47 | 331 | 2.647 |
| 16 | 107 | 0.6129 | 105 | 109 | 107 | 0.94 | 418 | 2.783 |
| 17 | 111 | 0.6200 | 109 | 112 | 111 | 0.76 | 557 | 2.815 |
| 18 | 113 | 0.7167 | 112 | 120 | 114 | 1.44 | 479 | 3.254 |
| 19 | 121 | 0.0385 | 120 | 122 | 121 | 0.53 | 57 | 0.175 |
| 20 | 124 | 0.1497 | 122 | 125 | 124 | 0.40 | 307 | 0.680 |
| 21 | 129 | 1.4014 | 125 | 135 | 128 | 1.41 | 997 | 6.362 |
| 22 | 145 | 13.5112 | 136 | 160 | 145 | 1.23 | 18834 | 61.341 |
| 23 | 500 (UM) | 0.5000 | 494 | 0 | 500 | 0.26 | 13355 | 27.240 |
| | TIC: | 21.4191 | ng/uL | | | | | |

```
Sample peak width (sec): 5    Sample min peak height: 50    Sample baseline V to V?: Y    Sample baseline V to V points: 3
Sample filter: Binomial       Number of points for filter: 3  Sample start region (min): 0  Sample end region (min): 80
Marker peak width (sec): 5    Marker min peak height: 500   Marker baseline V to V?: Y    Marker baseline V to V points: 3
Lower marker selection: First peak > 500 RFU                Upper marker selection: Last peak > 500 RFU
Ladder size (bp)1, 35, 75, 100, 150, 200, 250, 300, 400, 500
Quantification using: Upper Marker          Final concentration (ng/uL): 0.5000          Dilution factor: 12.0
```

**Sample:** SampC1
**Well location:** C1
**Created:** Monday, 11 October 2021 14:41:52

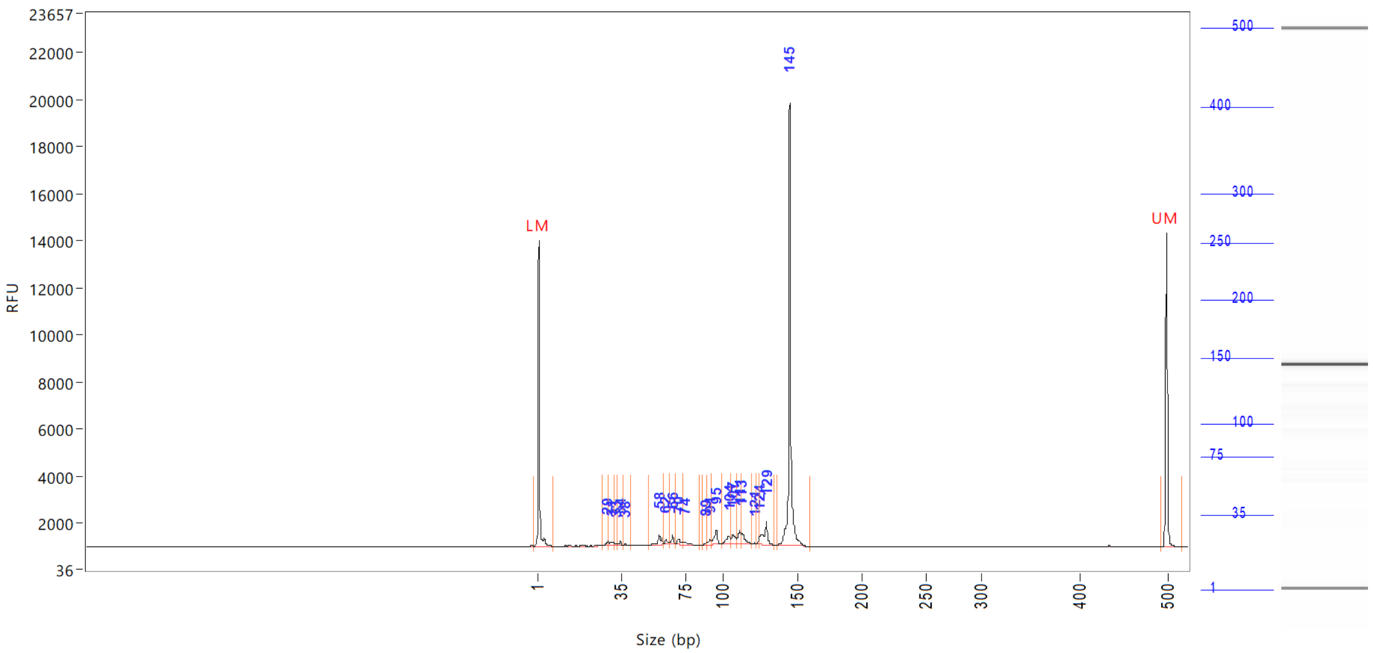


| Peak | Size (bp) | Concentration (ng/uL) | From (bp) | To (bp) | Average size (bp) | CV% | RFU | Corrected peak area |
|---|---|---|---|---|---|---|---|---|

| | | |
|---|---|---|
| TIM: | 319.3842 | nmole/L |
| Total concentration: | 21.9655 | ng/uL |

```
Sample peak width (sec): 5     Sample min peak height: 50    Sample baseline V to V?: Y    Sample baseline V to V points: 3
Sample filter: Binomial        Number of points for filter: 3  Sample start region (min): 0  Sample end region (min): 80
Marker peak width (sec): 5     Marker min peak height: 500   Marker baseline V to V?: Y    Marker baseline V to V points: 3
Lower marker selection: First peak > 500 RFU                 Upper marker selection: Last peak > 500 RFU
Ladder size (bp)1, 35, 75, 100, 150, 200, 250, 300, 400, 500
Quantification using: Upper Marker         Final concentration (ng/uL): 0.5000         Dilution factor: 12.0
```

**Sample:** SampD1
**Well location:** D1
**Created:** Monday, 11 October 2021 14:41:52

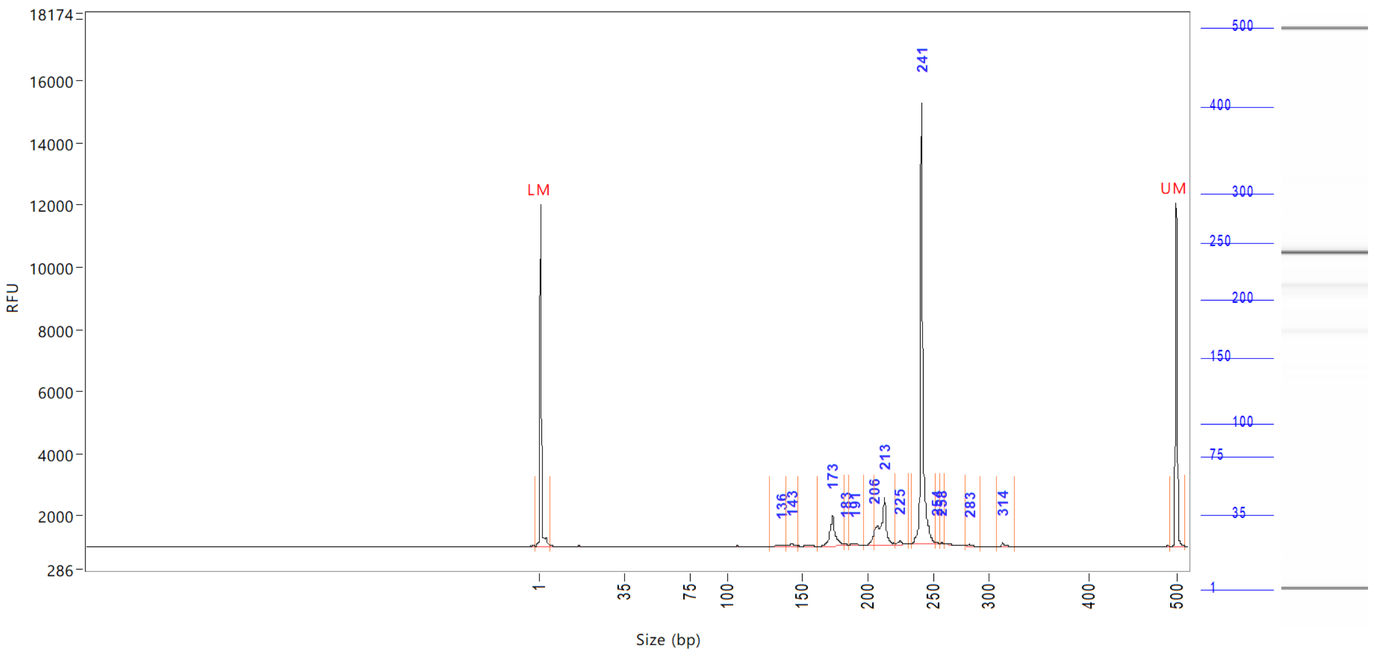


| Peak | Size (bp) | Concentration (ng/uL) | From (bp) | To (bp) | Average size (bp) | CV% | RFU | Corrected peak area |
|---|---|---|---|---|---|---|---|---|
| 1 | 1 (LM) | 0.9180 | 0 | 6 | 1 | 65.32 | 11005 | 41.529 |
| 2 | 136 | 0.1860 | 128 | 139 | 135 | 1.88 | 54 | 0.701 |
| 3 | 143 | 0.1896 | 139 | 147 | 142 | 1.15 | 107 | 0.715 |
| 4 | 173 | 2.1379 | 162 | 182 | 174 | 1.68 | 977 | 8.060 |
| 5 | 183 | 0.0788 | 182 | 185 | 183 | 0.47 | 81 | 0.297 |
| 6 | 191 | 0.2174 | 185 | 197 | 190 | 1.20 | 80 | 0.820 |
| 7 | 206 | 0.4743 | 197 | 206 | 204 | 0.70 | 484 | 1.788 |
| 8 | 213 | 3.3832 | 206 | 222 | 211 | 1.44 | 1554 | 12.754 |
| 9 | 225 | 0.1885 | 222 | 231 | 225 | 0.81 | 103 | 0.711 |
| 10 | 241 | 12.7588 | 233 | 253 | 242 | 0.85 | 14228 | 48.099 |
| 11 | 254 | 0.0608 | 253 | 256 | 254 | 0.32 | 78 | 0.229 |
| 12 | 258 | 0.0447 | 256 | 261 | 258 | 0.32 | 59 | 0.169 |
| 13 | 283 | 0.0768 | 279 | 292 | 282 | 0.74 | 52 | 0.289 |
| 14 | 314 | 0.1378 | 309 | 326 | 316 | 0.80 | 116 | 0.519 |
| 15 | 500 (UM) | 0.5000 | 493 | 510 | 500 | 0.23 | 11027 | 22.619 |

| | | |
|---|---|---|
| TIC: | 19.9346 | ng/uL |
| TIM: | 147.3629 | nmole/L |
| Total concentration: | 20.7151 | ng/uL |

```
Sample peak width (sec): 5     Sample min peak height: 50     Sample baseline V to V?: Y     Sample baseline V to V points: 3
Sample filter: Binomial        Number of points for filter: 3  Sample start region (min): 0  Sample end region (min): 80
Marker peak width (sec): 5     Marker min peak height: 500    Marker baseline V to V?: Y     Marker baseline V to V points: 3
Lower marker selection: First peak > 500 RFU                  Upper marker selection: Last peak > 500 RFU
Ladder size (bp)1, 35, 75, 100, 150, 200, 250, 300, 400, 500
Quantification using: Upper Marker          Final concentration (ng/uL): 0.5000          Dilution factor: 12.0
```

**Sample:** SampE1
**Well location:** E1
**Created:** Monday, 11 October 2021 14:41:52

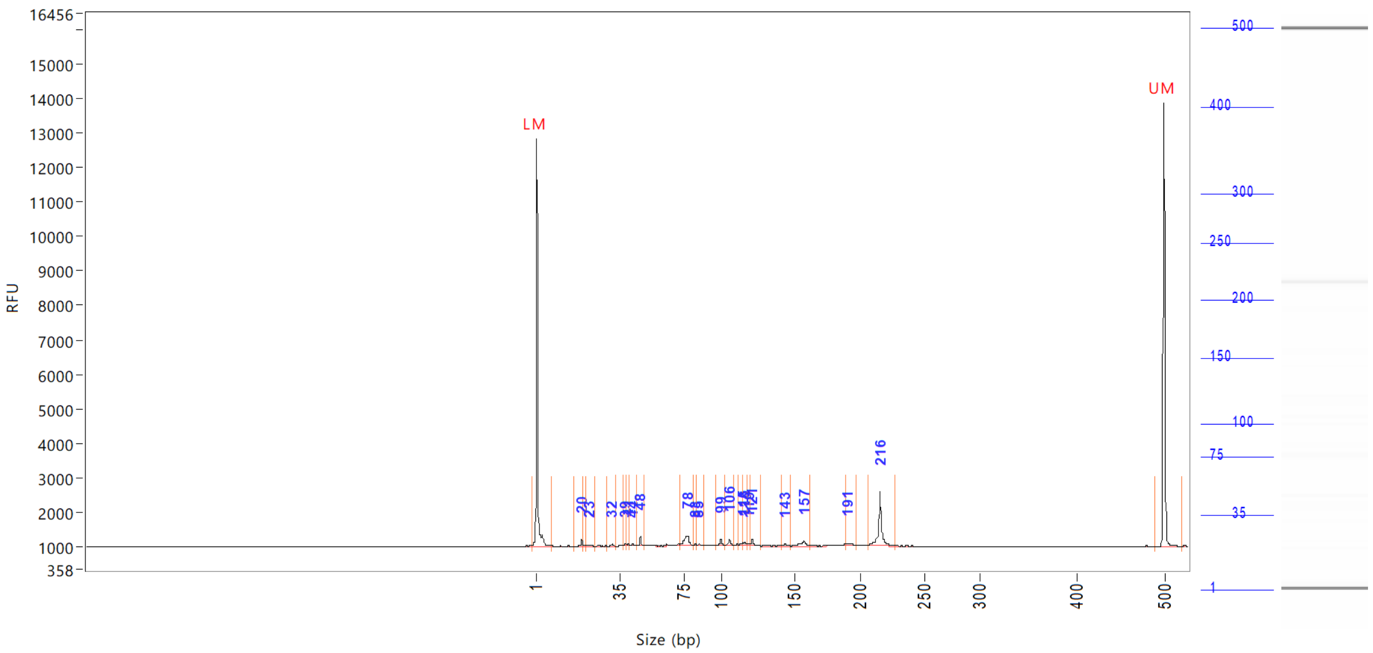


| Peak | Size (bp) | Concentration (ng/uL) | From (bp) | To (bp) | Average size (bp) | CV% | RFU | Corrected peak area |
|---|---|---|---|---|---|---|---|---|
| 1 | 1 (LM) | 0.8595 | 0 | 8 | 1 | 69.28 | 11784 | 44.392 |
| 2 | 20 | 0.1944 | 17 | 21 | 20 | 2.90 | 185 | 0.837 |
| 3 | 23 | 0.0710 | 22 | 25 | 23 | 2.90 | 52 | 0.305 |
| 4 | 32 | 0.0603 | 30 | 33 | 32 | 1.72 | 66 | 0.259 |
| 5 | 39 | 0.0749 | 38 | 40 | 39 | 1.51 | 65 | 0.322 |
| 6 | 41 | 0.0507 | 40 | 42 | 41 | 0.84 | 76 | 0.218 |
| 7 | 44 | 0.0829 | 42 | 46 | 44 | 2.25 | 71 | 0.357 |
| 8 | 48 | 0.2178 | 46 | 51 | 48 | 1.47 | 280 | 0.938 |
| 9 | 78 | 0.6604 | 72 | 81 | 76 | 2.41 | 292 | 2.842 |
| 10 | 83 | 0.0510 | 81 | 84 | 83 | 0.77 | 57 | 0.219 |
| 11 | 85 | 0.0541 | 84 | 89 | 85 | 0.91 | 63 | 0.233 |
| 12 | 99 | 0.1764 | 96 | 103 | 99 | 1.29 | 184 | 0.759 |
| 13 | 106 | 0.1829 | 103 | 109 | 106 | 0.98 | 196 | 0.787 |
| 14 | 115 | 0.0687 | 112 | 115 | 114 | 0.67 | 75 | 0.296 |
| 15 | 116 | 0.0789 | 115 | 118 | 116 | 0.53 | 87 | 0.340 |
| 16 | 119 | 0.0553 | 118 | 120 | 119 | 0.53 | 55 | 0.238 |
| 17 | 121 | 0.1578 | 120 | 127 | 121 | 0.90 | 165 | 0.679 |
| 18 | 143 | 0.0654 | 141 | 147 | 143 | 0.85 | 57 | 0.281 |
| 19 | 157 | 0.3125 | 147 | 162 | 156 | 1.74 | 142 | 1.345 |
| 20 | 191 | 0.1463 | 190 | 197 | 192 | 0.84 | 78 | 0.630 |
| 21 | 216 | 1.6324 | 208 | 228 | 216 | 1.26 | 1569 | 7.026 |
| 22 | 500 (UM) | 0.5000 | 490 | 0 | 500 | 0.28 | 12859 | 25.825 |

| | | |
|---|---|---|
| TIC: | 4.3938 | ng/uL |
| TIM: | 84.3793 | nmole/L |

```
Sample peak width (sec): 5    Sample min peak height: 50    Sample baseline V to V?: Y    Sample baseline V to V points: 3
Sample filter: Binomial       Number of points for filter: 3  Sample start region (min): 0  Sample end region (min): 80
Marker peak width (sec): 5    Marker min peak height: 500   Marker baseline V to V?: Y    Marker baseline V to V points: 3
Lower marker selection: First peak > 500 RFU                Upper marker selection: Last peak > 500 RFU
Ladder size (bp)1, 35, 75, 100, 150, 200, 250, 300, 400, 500
Quantification using: Upper Marker          Final concentration (ng/uL): 0.5000          Dilution factor: 12.0
```

**Sample:** SampE1
**Well location:** E1
**Created:** Monday, 11 October 2021 14:41:52

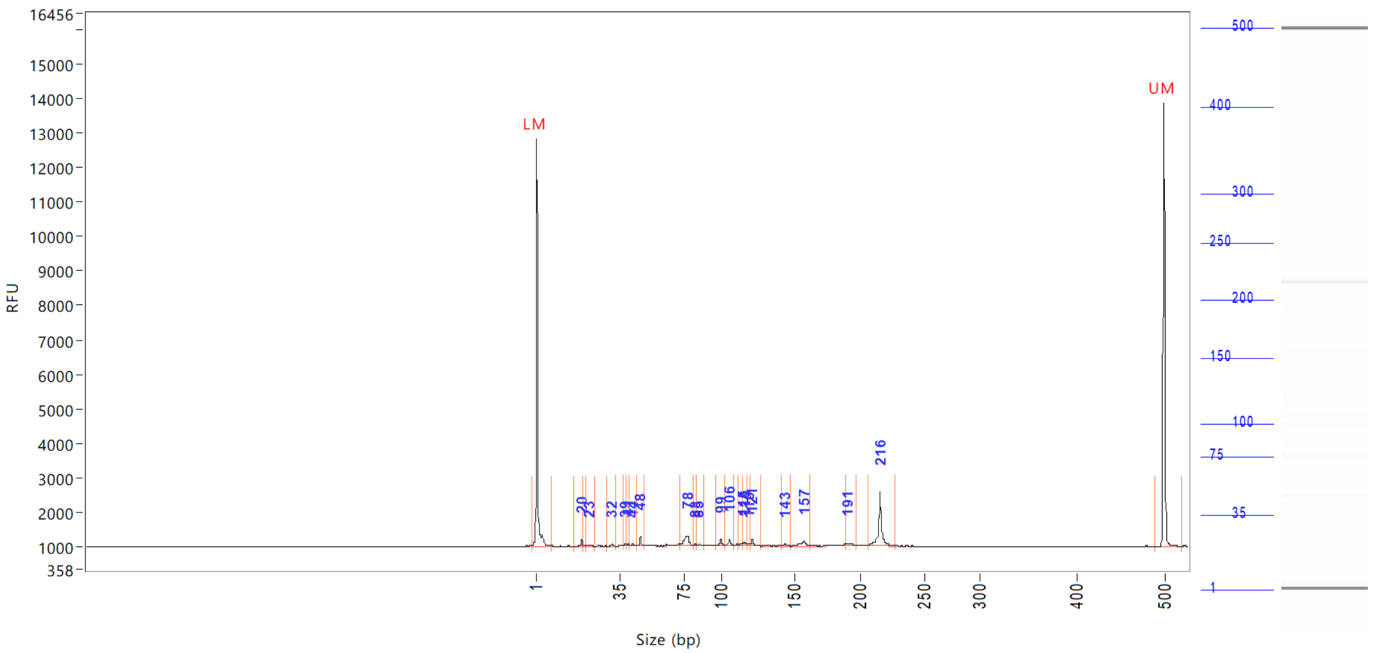


| Peak | Size (bp) | Concentration (ng/uL) | From (bp) | To (bp) | Average size (bp) | CV% | RFU | Corrected peak area |
|---|---|---|---|---|---|---|---|---|
| | Total concentration: | | 5.1583 | ng/uL | | | | |

```
Sample peak width (sec): 5    Sample min peak height: 50    Sample baseline V to V?: Y    Sample baseline V to V points: 3
Sample filter: Binomial       Number of points for filter: 3  Sample start region (min): 0  Sample end region (min): 80
Marker peak width (sec): 5    Marker min peak height: 500   Marker baseline V to V?: Y    Marker baseline V to V points: 3
Lower marker selection: First peak > 500 RFU                Upper marker selection: Last peak > 500 RFU
Ladder size (bp)1, 35, 75, 100, 150, 200, 250, 300, 400, 500
Quantification using: Upper Marker         Final concentration (ng/uL): 0.5000         Dilution factor: 12.0
```

**Sample:** SampF1
**Well location:** F1
**Created:** Monday, 11 October 2021 14:41:52

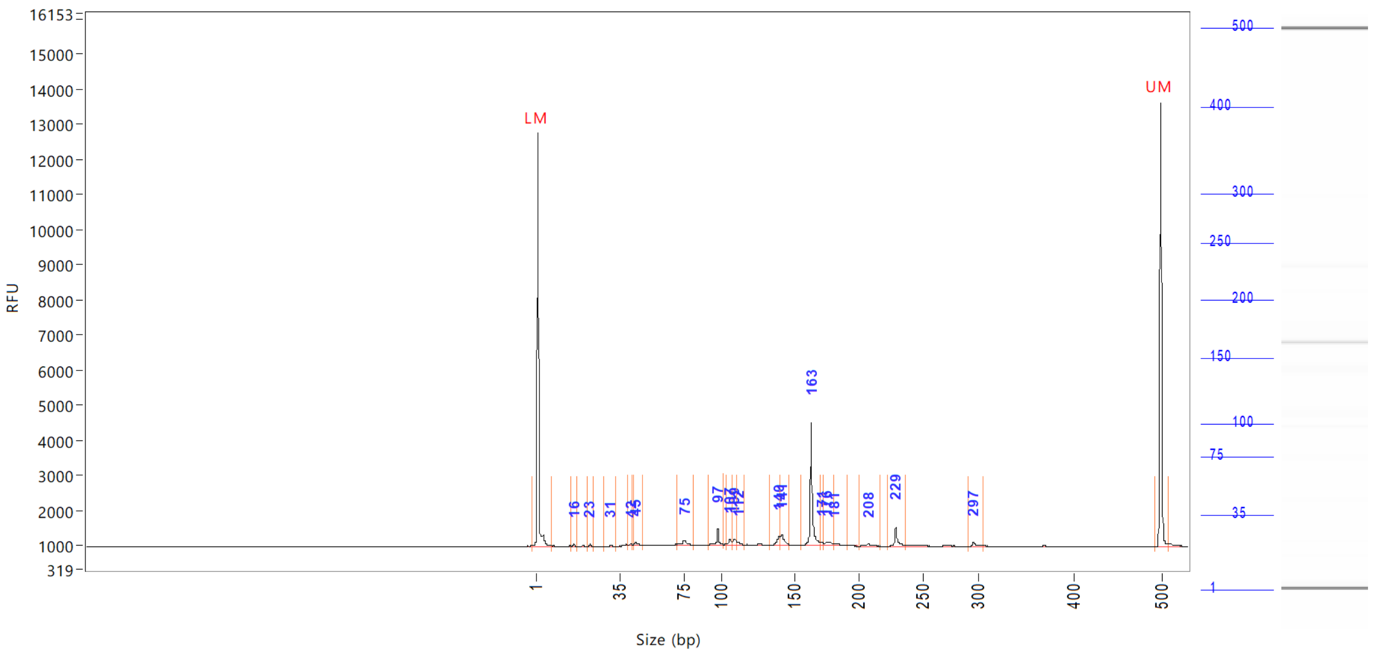


| Peak | Size (bp) | Concentration (ng/uL) | From (bp) | To (bp) | Average size (bp) | CV% | RFU | Corrected peak area |
|---|---|---|---|---|---|---|---|---|
| 1 | 1 (LM) | 0.8441 | 0 | 8 | 2 | 68.23 | 11801 | 44.168 |
| 2 | 16 | 0.0692 | 15 | 18 | 16 | 2.25 | 72 | 0.302 |
| 3 | 23 | 0.0550 | 22 | 25 | 23 | 1.66 | 65 | 0.240 |
| 4 | 31 | 0.0595 | 29 | 33 | 31 | 2.41 | 50 | 0.259 |
| 5 | 42 | 0.0428 | 41 | 43 | 42 | 1.21 | 51 | 0.187 |
| 6 | 45 | 0.1224 | 44 | 49 | 46 | 1.99 | 87 | 0.534 |
| 7 | 75 | 0.3754 | 70 | 81 | 75 | 3.33 | 130 | 1.637 |
| 8 | 97 | 0.3851 | 91 | 102 | 97 | 1.55 | 448 | 1.679 |
| 9 | 107 | 0.1708 | 104 | 108 | 106 | 0.74 | 165 | 0.745 |
| 10 | 109 | 0.2288 | 108 | 111 | 110 | 0.84 | 155 | 0.998 |
| 11 | 112 | 0.1001 | 111 | 116 | 113 | 0.86 | 90 | 0.436 |
| 12 | 140 | 0.3092 | 133 | 140 | 139 | 0.94 | 260 | 1.348 |
| 13 | 141 | 0.4439 | 140 | 147 | 142 | 1.01 | 311 | 1.936 |
| 14 | 163 | 2.6663 | 156 | 170 | 164 | 1.01 | 3497 | 11.626 |
| 15 | 171 | 0.0880 | 170 | 173 | 172 | 0.50 | 84 | 0.384 |
| 16 | 176 | 0.2032 | 173 | 181 | 177 | 1.18 | 76 | 0.886 |
| 17 | 181 | 0.0984 | 181 | 191 | 184 | 1.20 | 54 | 0.429 |
| 18 | 208 | 0.1193 | 202 | 217 | 208 | 1.25 | 57 | 0.520 |
| 19 | 229 | 0.4972 | 224 | 237 | 230 | 0.94 | 551 | 2.168 |
| 20 | 297 | 0.1120 | 292 | 308 | 298 | 1.02 | 117 | 0.489 |
| 21 | 500 (UM) | 0.5000 | 494 | 509 | 500 | 0.26 | 12658 | 26.162 |

| | | |
|---|---|---|
| TIC: | 6.1467 | ng/uL |
| TIM: | 86.5533 | nmole/L |
| Total concentration: | 6.9135 | ng/uL |

```
Sample peak width (sec): 5    Sample min peak height: 50    Sample baseline V to V?: Y    Sample baseline V to V points: 3
Sample filter: Binomial       Number of points for filter: 3  Sample start region (min): 0  Sample end region (min): 80
Marker peak width (sec): 5    Marker min peak height: 500   Marker baseline V to V?: Y    Marker baseline V to V points: 3
Lower marker selection: First peak > 500 RFU                Upper marker selection: Last peak > 500 RFU
Ladder size (bp)1, 35, 75, 100, 150, 200, 250, 300, 400, 500
Quantification using: Upper Marker          Final concentration (ng/uL): 0.5000          Dilution factor: 12.0
```

**Sample:** SampG1
**Well location:** G1
**Created:** Monday, 11 October 2021 14:41:52

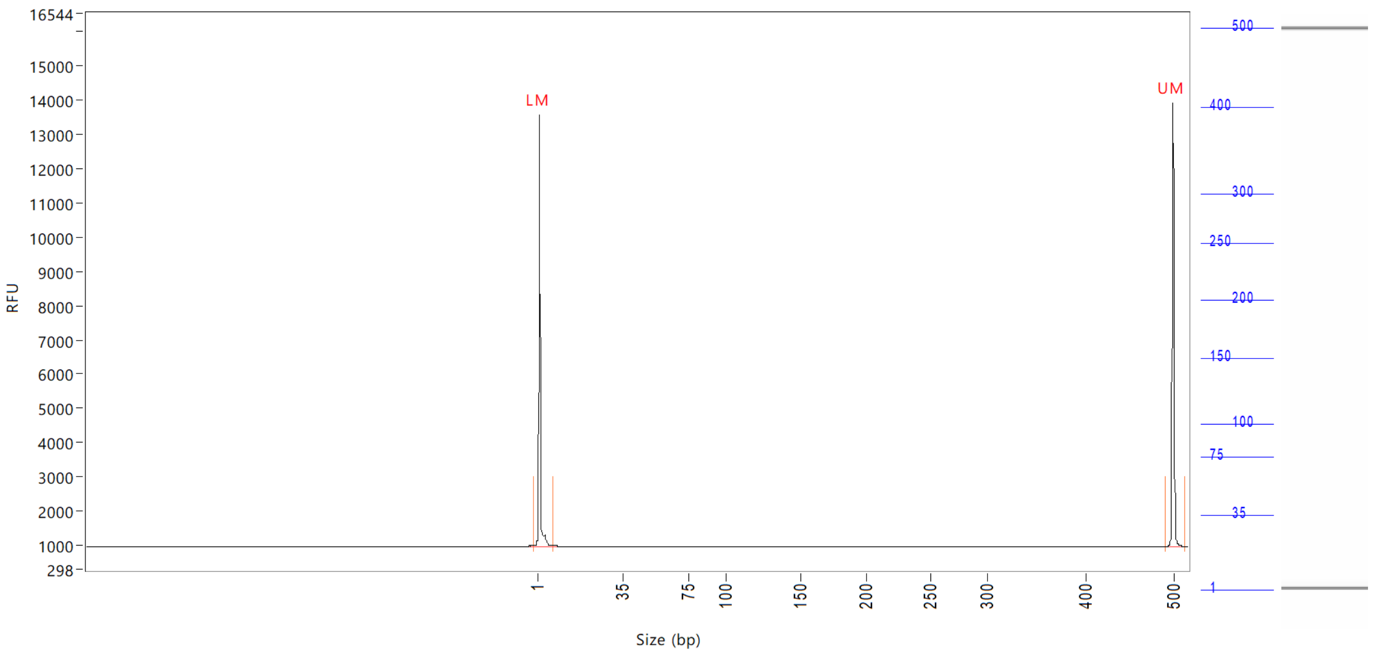


| Peak | Size (bp) | Concentration (ng/uL) | From (bp) | To (bp) | Average size (bp) | CV% | RFU | Corrected peak area |
|---|---|---|---|---|---|---|---|---|
| 1 | 1 (LM) | 0.8859 | 0 | 8 | 2 | 66.67 | 12588 | 47.866 |
| 2 | 500 (UM) | 0.5000 | 491 | 514 | 500 | 0.25 | 12989 | 27.017 |

| | | |
|---|---|---|
| TIC: | 0.0000 | ng/uL |
| TIM: | 0.0000 | nmole/L |
| Total concentration: | 0.4075 | ng/uL |

Sample peak width (sec): 5    Sample min peak height: 50    Sample baseline V to V?: Y    Sample baseline V to V points: 3
Sample filter: Binomial    Number of points for filter: 3  Sample start region (min): 0  Sample end region (min): 80
Marker peak width (sec): 5    Marker min peak height: 500   Marker baseline V to V?: Y    Marker baseline V to V points: 3
Lower marker selection: First peak > 500 RFU    Upper marker selection: Last peak > 500 RFU
Ladder size (bp)1, 35, 75, 100, 150, 200, 250, 300, 400, 500
Quantification using: Upper Marker    Final concentration (ng/uL): 0.5000    Dilution factor: 12.0

**Sample:** SampH1
**Well location:** H1
**Created:** Monday, 11 October 2021 14:41:52

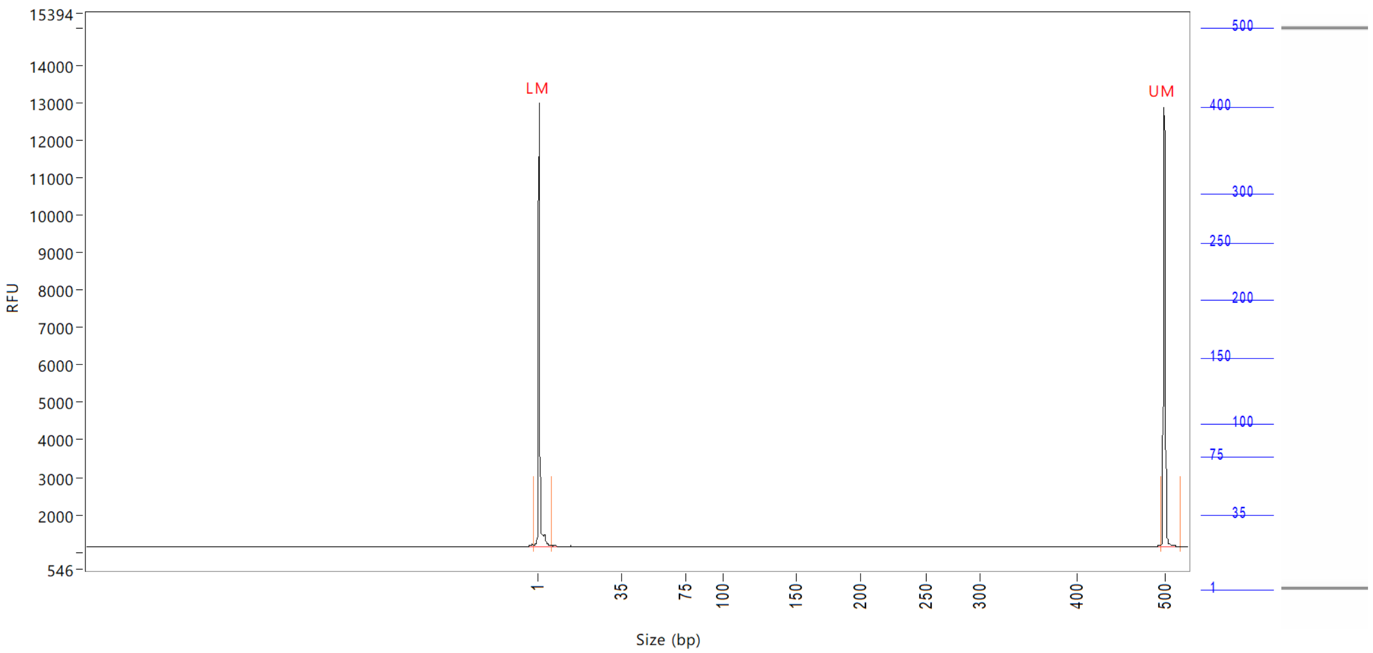


| Peak | Size (bp) | Concentration (ng/uL) | From (bp) | To (bp) | Average size (bp) | CV% | RFU | Corrected peak area |
|---|---|---|---|---|---|---|---|---|
| 1 | 1 (LM) | 0.9082 | 0 | 7 | 1 | 69.92 | 11859 | 44.762 |
| 2 | 500 (UM) | 0.5000 | 495 | 0 | 500 | 0.29 | 11746 | 24.643 |

| | | |
|---|---|---|
| TIC: | 0.0000 | ng/uL |
| TIM: | 0.0000 | nmole/L |
| Total concentration: | 0.5653 | ng/uL |

Sample peak width (sec): 5    Sample min peak height: 50    Sample baseline V to V?: Y    Sample baseline V to V points: 3
Sample filter: Binomial    Number of points for filter: 3  Sample start region (min): 0  Sample end region (min): 80
Marker peak width (sec): 5    Marker min peak height: 500    Marker baseline V to V?: Y    Marker baseline V to V points: 3
Lower marker selection: First peak > 500 RFU    Upper marker selection: Last peak > 500 RFU
Ladder size (bp)1, 35, 75, 100, 150, 200, 250, 300, 400, 500
Quantification using: Upper Marker    Final concentration (ng/uL): 0.5000    Dilution factor: 12.0

**Sample:** SampA2
**Well location:** A2
**Created:** Monday, 11 October 2021 14:41:52

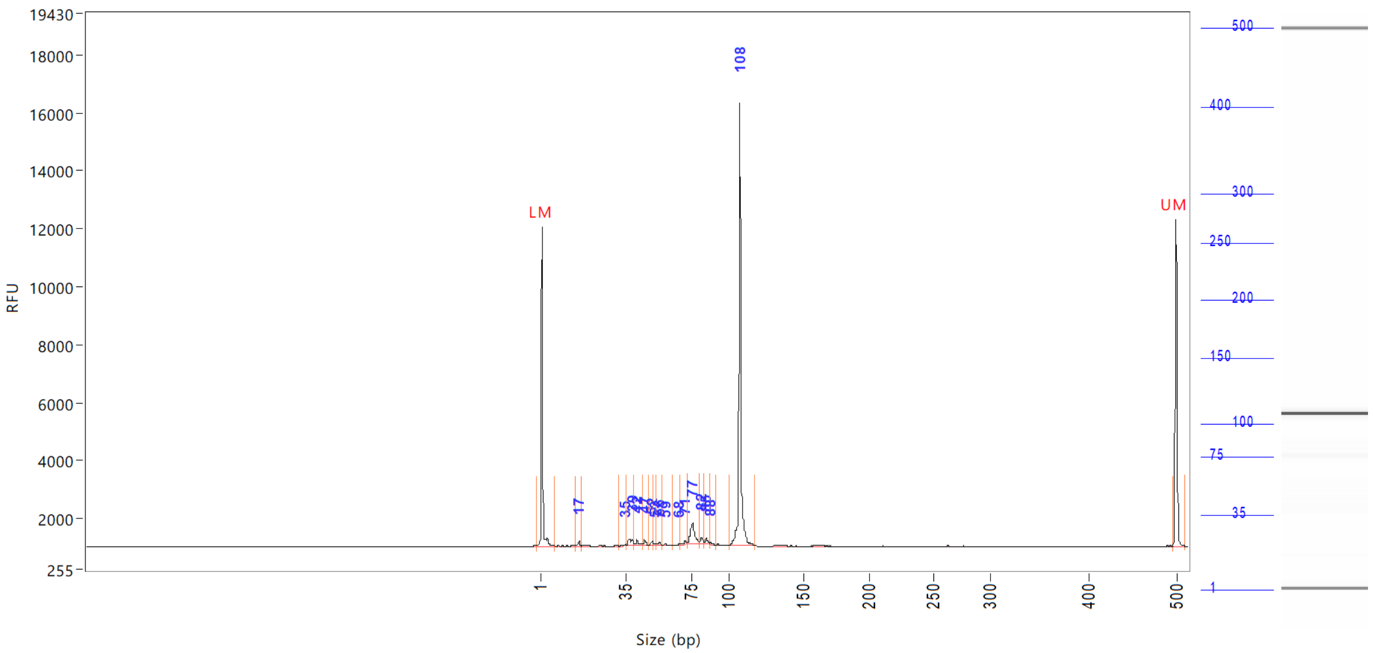


| Peak | Size (bp) | Concentration (ng/uL) | From (bp) | To (bp) | Average size (bp) | CV% | RFU | Corrected peak area |
|---|---|---|---|---|---|---|---|---|
| 1 | 1 (LM) | 0.8760 | 0 | 7 | 1 | 70.18 | 11006 | 40.554 |
| 2 | 17 | 0.2253 | 15 | 17 | 16 | 2.44 | 192 | 0.869 |
| 3 | 35 | 0.0774 | 32 | 35 | 34 | 1.94 | 51 | 0.299 |
| 4 | 39 | 0.5796 | 35 | 41 | 38 | 3.13 | 248 | 2.236 |
| 5 | 42 | 0.3485 | 41 | 46 | 43 | 3.07 | 199 | 1.344 |
| 6 | 47 | 0.2478 | 46 | 50 | 47 | 1.53 | 196 | 0.956 |
| 7 | 52 | 0.1779 | 50 | 53 | 51 | 1.25 | 146 | 0.686 |
| 8 | 53 | 0.0594 | 53 | 54 | 53 | 0.86 | 54 | 0.229 |
| 9 | 56 | 0.1413 | 54 | 57 | 56 | 1.33 | 97 | 0.545 |
| 10 | 59 | 0.1122 | 57 | 64 | 59 | 2.01 | 69 | 0.433 |
| 11 | 68 | 0.0591 | 64 | 69 | 67 | 1.12 | 50 | 0.228 |
| 12 | 71 | 0.2074 | 69 | 73 | 71 | 1.41 | 136 | 0.800 |
| 13 | 77 | 1.5901 | 73 | 81 | 76 | 2.42 | 713 | 6.134 |
| 14 | 82 | 0.2657 | 81 | 84 | 82 | 0.82 | 226 | 1.025 |
| 15 | 85 | 0.2694 | 84 | 87 | 85 | 1.01 | 202 | 1.039 |
| 16 | 88 | 0.0922 | 87 | 91 | 89 | 1.00 | 71 | 0.356 |
| 17 | 108 | 13.0648 | 100 | 118 | 108 | 1.32 | 15281 | 50.402 |
| 18 | 500 (UM) | 0.5000 | 496 | 510 | 500 | 0.23 | 11265 | 23.147 |

| | | |
|---|---|---|
| TIC: | 17.5181 | ng/uL |
| TIM: | 339.6368 | nmole/L |
| Total concentration: | 18.2557 | ng/uL |

```
Sample peak width (sec): 5    Sample min peak height: 50    Sample baseline V to V?: Y    Sample baseline V to V points: 3
Sample filter: Binomial       Number of points for filter: 3  Sample start region (min): 0  Sample end region (min): 80
Marker peak width (sec): 5    Marker min peak height: 500   Marker baseline V to V?: Y    Marker baseline V to V points: 3
Lower marker selection: First peak > 500 RFU                Upper marker selection: Last peak > 500 RFU
Ladder size (bp)1, 35, 75, 100, 150, 200, 250, 300, 400, 500
Quantification using: Upper Marker          Final concentration (ng/uL): 0.5000          Dilution factor: 12.0
```

**Sample:** SampB2
**Well location:** B2
**Created:** Monday, 11 October 2021 14:41:52

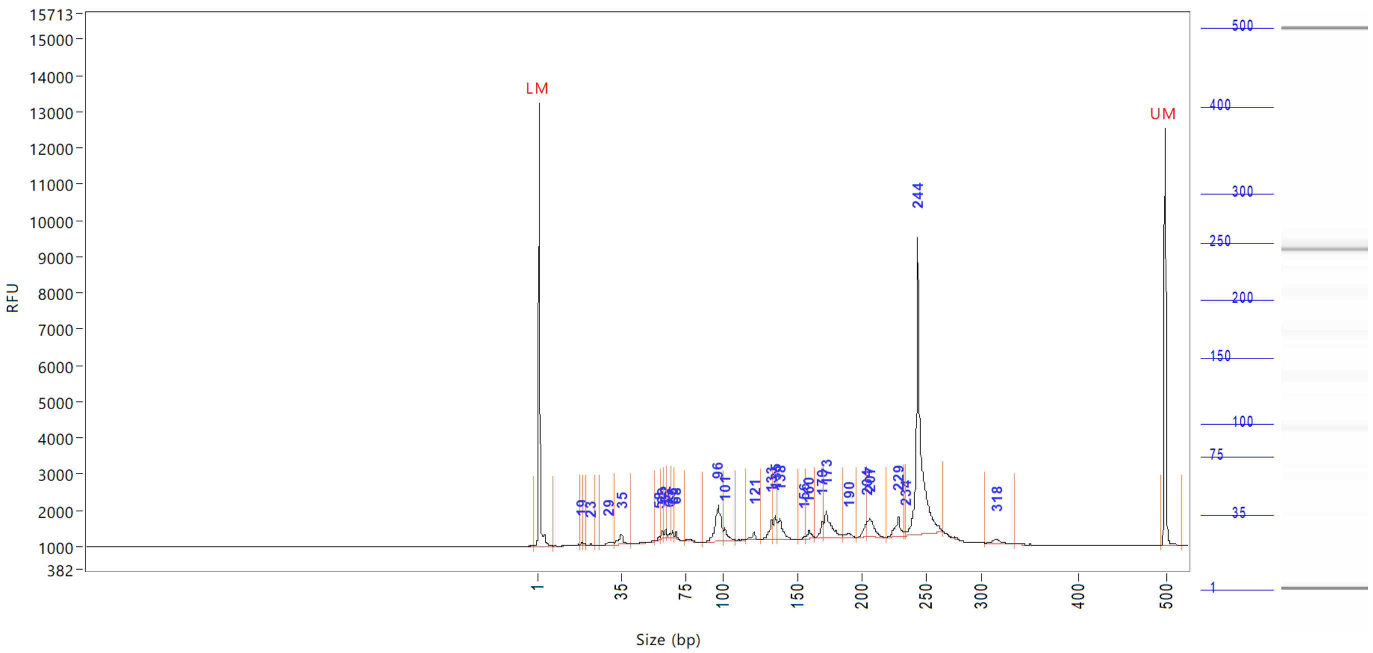

| Peak | Size (bp) | Concentration (ng/uL) | From (bp) | To (bp) | Average size (bp) | CV% | RFU | Corrected peak area |
|---|---|---|---|---|---|---|---|---|
| 1 | 1 (LM) | 0.9343 | 0 | 8 | 1 | 69.90 | 12250 | 44.708 |
| 2 | 19 | 0.0782 | 18 | 19 | 19 | 1.63 | 81 | 0.312 |
| 3 | 23 | 0.0650 | 21 | 24 | 22 | 2.59 | 52 | 0.259 |
| 4 | 29 | 0.2745 | 26 | 32 | 30 | 3.81 | 83 | 1.094 |
| 5 | 35 | 0.7105 | 32 | 41 | 35 | 4.77 | 266 | 2.833 |
| 6 | 58 | 0.1249 | 56 | 59 | 58 | 1.01 | 113 | 0.498 |
| 7 | 60 | 0.2374 | 59 | 61 | 60 | 0.93 | 239 | 0.947 |
| 8 | 62 | 0.1857 | 61 | 63 | 62 | 0.69 | 234 | 0.741 |
| 9 | 65 | 0.1034 | 63 | 65 | 64 | 0.84 | 106 | 0.412 |
| 10 | 66 | 0.1485 | 65 | 67 | 66 | 0.68 | 194 | 0.592 |
| 11 | 68 | 0.1605 | 67 | 74 | 68 | 1.10 | 207 | 0.640 |
| 12 | 96 | 2.7381 | 86 | 100 | 96 | 2.26 | 984 | 10.919 |
| 13 | 101 | 0.5861 | 100 | 109 | 103 | 1.90 | 359 | 2.337 |
| 14 | 121 | 0.3759 | 115 | 126 | 120 | 1.59 | 243 | 1.499 |
| 15 | 133 | 0.8650 | 126 | 134 | 132 | 1.24 | 552 | 3.449 |
| 16 | 135 | 0.7324 | 134 | 136 | 135 | 0.52 | 645 | 2.921 |
| 17 | 138 | 1.2751 | 136 | 152 | 139 | 1.60 | 598 | 5.085 |
| 18 | 156 | 0.0912 | 152 | 157 | 156 | 0.59 | 89 | 0.364 |
| 19 | 160 | 0.3889 | 157 | 165 | 160 | 1.05 | 243 | 1.551 |
| 20 | 170 | 0.5308 | 165 | 171 | 169 | 0.79 | 429 | 2.116 |
| 21 | 173 | 1.8403 | 171 | 185 | 175 | 1.93 | 734 | 7.338 |
| 22 | 190 | 0.2930 | 185 | 195 | 190 | 1.15 | 124 | 1.168 |
| 23 | 204 | 0.4581 | 195 | 205 | 203 | 0.89 | 391 | 1.827 |
| 24 | 207 | 1.2821 | 205 | 220 | 208 | 1.26 | 509 | 5.113 |
| 25 | 229 | 0.9481 | 220 | 233 | 228 | 1.07 | 536 | 3.781 |

Sample peak width (sec): 5    Sample min peak height: 50    Sample baseline V to V?: Y    Sample baseline V to V points: 3
Sample filter: Binomial    Number of points for filter: 3  Sample start region (min): 0  Sample end region (min): 80
Marker peak width (sec): 5    Marker min peak height: 500    Marker baseline V to V?: Y    Marker baseline V to V points: 3
Lower marker selection: First peak > 500 RFU    Upper marker selection: Last peak > 500 RFU
Ladder size (bp)1, 35, 75, 100, 150, 200, 250, 300, 400, 500
Quantification using: Upper Marker    Final concentration (ng/uL): 0.5000    Dilution factor: 12.0

**Sample:** SampB2
**Well location:** B2
**Created:** Monday, 11 October 2021 14:41:52

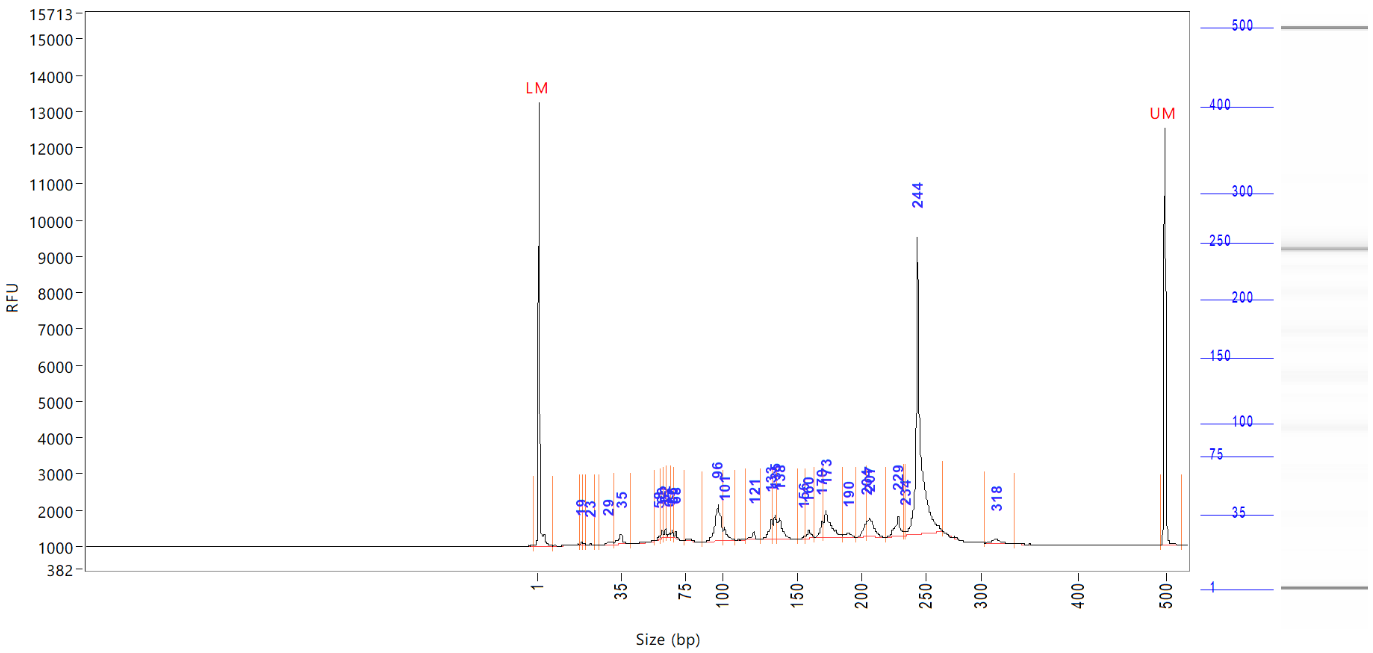


| Peak | Size (bp) | Concentration (ng/uL) | From (bp) | To (bp) | Average size (bp) | CV% | RFU | Corrected peak area |
|---|---|---|---|---|---|---|---|---|
| 26 | 234 | 0.0835 | 233 | 235 | 234 | 0.25 | 103 | |
| 27 | 244 | 12.3772 | 235 | 266 | 246 | 1.94 | 8185 | |
| 28 | 318 | 0.3296 | 306 | 336 | 319 | 1.70 | 135 | |
| 29 | 500 (UM) | 0.5000 | 495 | 0 | 500 | 0.32 | 11531 | |

| | | |
|---|---|---|
| TIC: | 27.2838 | ng/uL |
| TIM: | 316.4662 | nmole/L |
| Total concentration: | 27.7780 | ng/uL |

```
Sample peak width (sec): 5    Sample min peak height: 50    Sample baseline V to V?: Y    Sample baseline V to V points: 3
Sample filter: Binomial       Number of points for filter: 3  Sample start region (min): 0  Sample end region (min): 80
Marker peak width (sec): 5    Marker min peak height: 500   Marker baseline V to V?: Y    Marker baseline V to V points: 3
Lower marker selection: First peak > 500 RFU               Upper marker selection: Last peak > 500 RFU
Ladder size (bp)1, 35, 75, 100, 150, 200, 250, 300, 400, 500
Quantification using: Upper Marker         Final concentration (ng/uL): 0.5000         Dilution factor: 12.0
```

**Sample:** SampC2
**Well location:** C2
**Created:** Monday, 11 October 2021 14:41:52

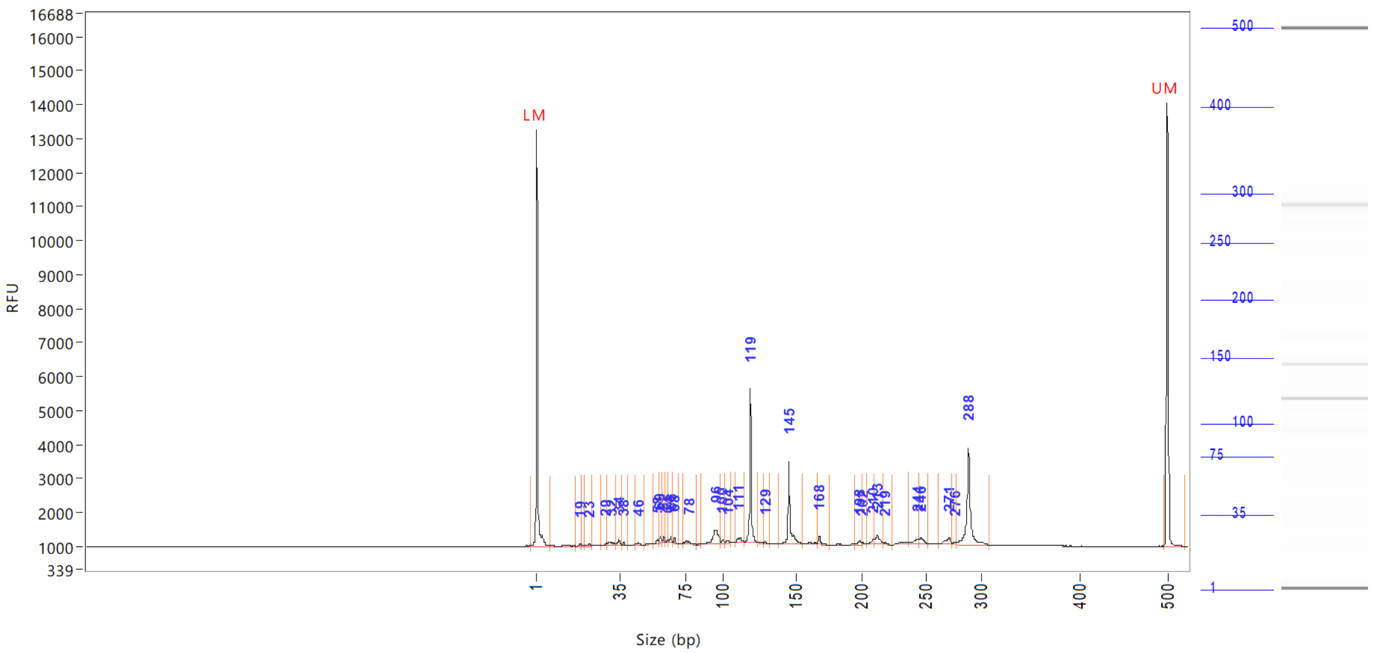

| Peak | Size (bp) | Concentration (ng/uL) | From (bp) | To (bp) | Average size (bp) | CV% | RFU | Corrected peak area |
|---|---|---|---|---|---|---|---|---|
| 1 | 1 (LM) | 0.8581 | 0 | 8 | 1 | 68.50 | 12233 | 46.489 |
| 2 | 19 | 0.0612 | 18 | 20 | 19 | 2.04 | 69 | 0.276 |
| 3 | 23 | 0.0567 | 21 | 24 | 22 | 2.25 | 62 | 0.256 |
| 4 | 29 | 0.0951 | 27 | 30 | 29 | 1.98 | 82 | 0.429 |
| 5 | 32 | 0.2453 | 30 | 33 | 31 | 2.92 | 92 | 1.107 |
| 6 | 34 | 0.2409 | 33 | 37 | 34 | 2.11 | 178 | 1.088 |
| 7 | 38 | 0.0651 | 37 | 40 | 38 | 1.59 | 83 | 0.294 |
| 8 | 46 | 0.0957 | 45 | 50 | 47 | 1.90 | 77 | 0.432 |
| 9 | 58 | 0.0991 | 55 | 59 | 58 | 0.88 | 115 | 0.447 |
| 10 | 60 | 0.1576 | 59 | 61 | 60 | 0.92 | 183 | 0.712 |
| 11 | 62 | 0.1075 | 61 | 63 | 62 | 0.60 | 162 | 0.485 |
| 12 | 64 | 0.0597 | 63 | 65 | 64 | 0.77 | 77 | 0.270 |
| 13 | 66 | 0.1121 | 65 | 67 | 66 | 0.65 | 174 | 0.506 |
| 14 | 68 | 0.0981 | 67 | 71 | 68 | 0.78 | 158 | 0.443 |
| 15 | 78 | 0.1776 | 74 | 83 | 78 | 2.41 | 82 | 0.802 |
| 16 | 96 | 0.9636 | 86 | 99 | 95 | 2.17 | 406 | 4.351 |
| 17 | 100 | 0.0913 | 99 | 102 | 100 | 0.98 | 107 | 0.412 |
| 18 | 104 | 0.1190 | 102 | 106 | 104 | 1.00 | 75 | 0.537 |
| 19 | 111 | 0.2838 | 109 | 116 | 112 | 1.34 | 165 | 1.281 |
| 20 | 119 | 2.8966 | 116 | 124 | 119 | 0.73 | 4523 | 13.078 |
| 21 | 129 | 0.0490 | 128 | 132 | 129 | 0.56 | 75 | 0.221 |
| 22 | 145 | 1.9135 | 138 | 155 | 145 | 1.34 | 2447 | 8.639 |
| 23 | 168 | 0.1905 | 166 | 176 | 168 | 0.77 | 251 | 0.860 |
| 24 | 198 | 0.1371 | 195 | 201 | 198 | 0.81 | 108 | 0.619 |
| 25 | 202 | 0.0354 | 201 | 204 | 202 | 0.30 | 54 | 0.160 |

```
Sample peak width (sec): 5     Sample min peak height: 50     Sample baseline V to V?: Y     Sample baseline V to V points: 3
Sample filter: Binomial        Number of points for filter: 3  Sample start region (min): 0  Sample end region (min): 80
Marker peak width (sec): 5     Marker min peak height: 500    Marker baseline V to V?: Y     Marker baseline V to V points: 3
Lower marker selection: First peak > 500 RFU                  Upper marker selection: Last peak > 500 RFU
Ladder size (bp)1, 35, 75, 100, 150, 200, 250, 300, 400, 500
Quantification using: Upper Marker          Final concentration (ng/uL): 0.5000          Dilution factor: 12.0
```

**Sample:** SampC2
**Well location:** C2
**Created:** Monday, 11 October 2021 14:41:52

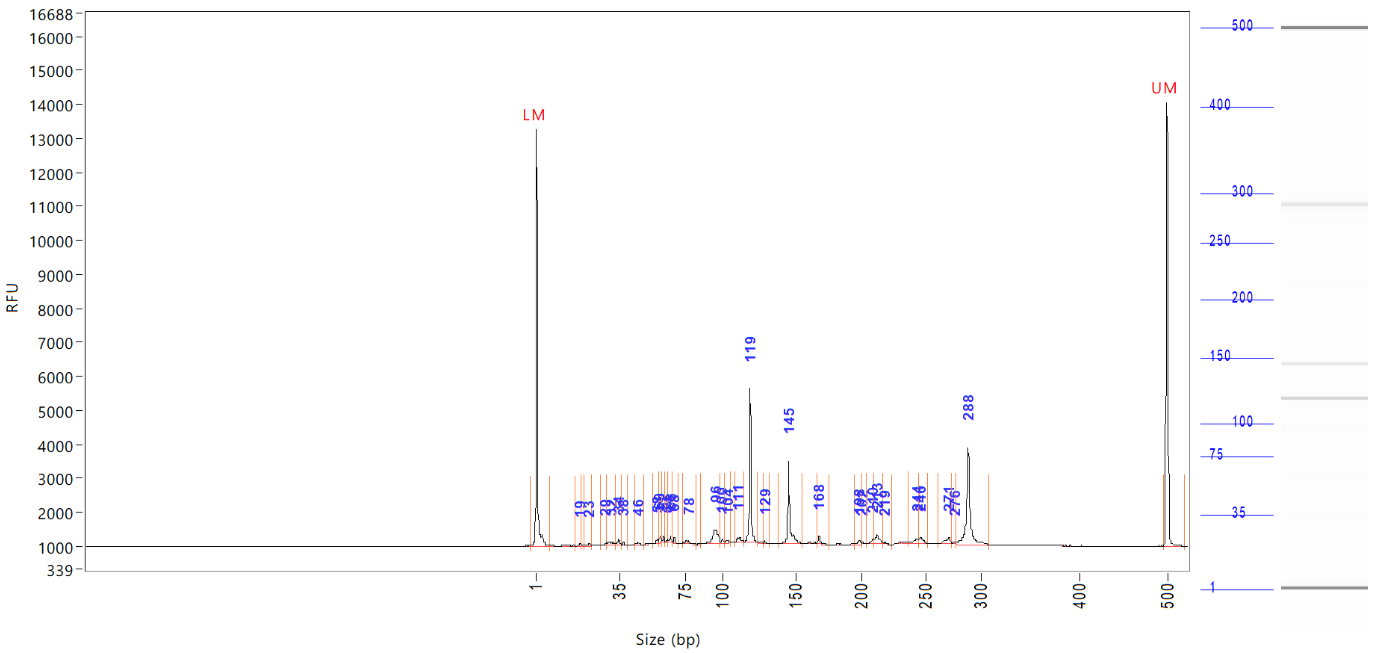


| Peak | Size (bp) | Concentration (ng/uL) | From (bp) | To (bp) | Average size (bp) | CV% | RFU | Corrected peak area |
|---|---|---|---|---|---|---|---|---|
| 26 | 210 | 0.1296 | 204 | 211 | 209 | 0.58 | 137 | |
| 27 | 213 | 0.3792 | 211 | 218 | 213 | 0.82 | 267 | |
| 28 | 219 | 0.0377 | 218 | 224 | 219 | 0.46 | 53 | |
| 29 | 244 | 0.1935 | 236 | 244 | 242 | 0.80 | 137 | |
| 30 | 246 | 0.2167 | 244 | 253 | 247 | 0.68 | 156 | |
| 31 | 271 | 0.2996 | 262 | 274 | 269 | 0.94 | 180 | |
| 32 | 276 | 0.0518 | 274 | 277 | 276 | 0.32 | 70 | |
| 33 | 288 | 3.0214 | 277 | 309 | 289 | 1.39 | 2836 | |
| 34 | 500 (UM) | 0.5000 | 496 | 0 | 500 | 0.27 | 13056 | |

| | | |
|---|---|---|
| TIC: | 12.6809 | ng/uL |
| TIM: | 181.8766 | nmole/L |
| Total concentration: | 13.4119 | ng/uL |

Sample peak width (sec): 5 Sample min peak height: 50 Sample baseline V to V?: Y Sample baseline V to V points: 3
Sample filter: Binomial Number of points for filter: 3 Sample start region (min): 0 Sample end region (min): 80
Marker peak width (sec): 5 Marker min peak height: 500 Marker baseline V to V?: Y Marker baseline V to V points: 3
Lower marker selection: First peak > 500 RFU Upper marker selection: Last peak > 500 RFU
Ladder size (bp)1, 35, 75, 100, 150, 200, 250, 300, 400, 500
Quantification using: Upper Marker Final concentration (ng/uL): 0.5000 Dilution factor: 12.0

**Sample:** SampD2
**Well location:** D2
**Created:** Monday, 11 October 2021 14:41:52

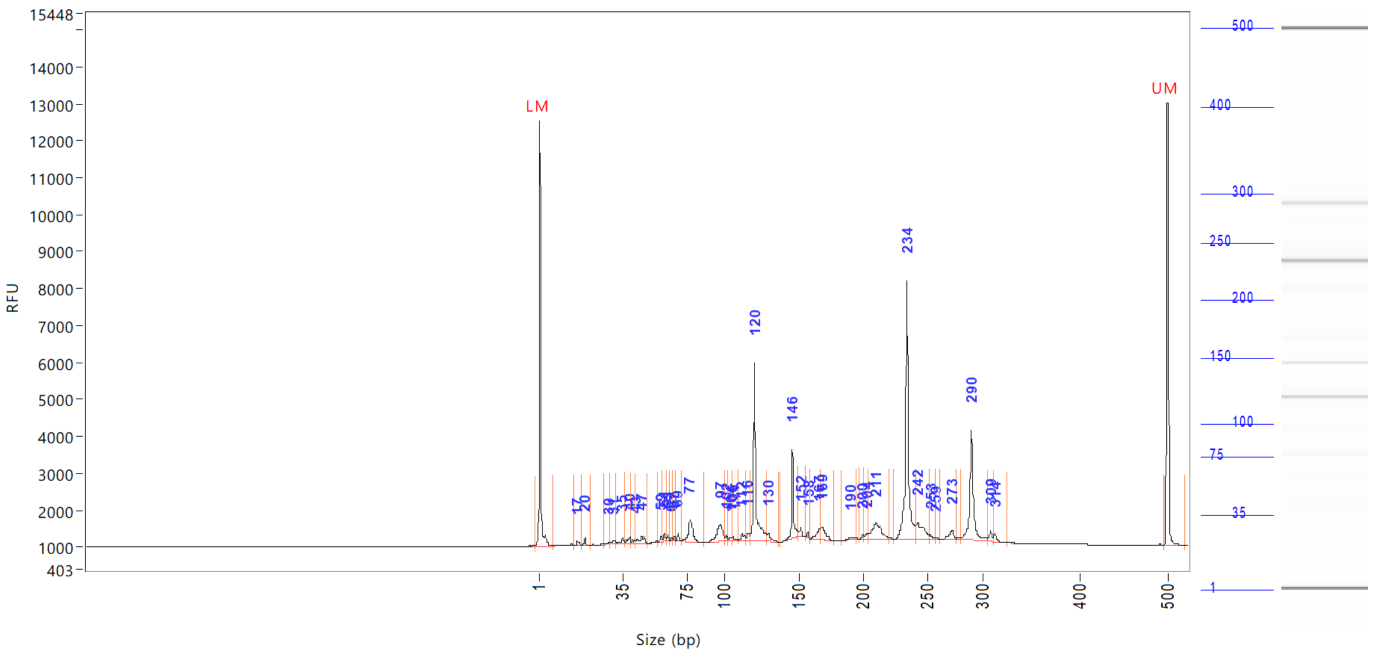


| Peak | Size (bp) | Concentration (ng/uL) | From (bp) | To (bp) | Average size (bp) | CV% | RFU | Corrected peak area |
|---|---|---|---|---|---|---|---|---|
| 1 | 1 (LM) | 0.8587 | 0 | 8 | 1 | 67.04 | 11521 | 43.220 |
| 2 | 17 | 0.1957 | 15 | 19 | 17 | 3.30 | 117 | 0.821 |
| 3 | 20 | 0.1905 | 19 | 22 | 20 | 2.09 | 219 | 0.799 |
| 4 | 30 | 0.0869 | 28 | 30 | 29 | 1.94 | 70 | 0.365 |
| 5 | 31 | 0.1607 | 30 | 32 | 31 | 1.99 | 78 | 0.674 |
| 6 | 35 | 0.1722 | 32 | 37 | 34 | 2.52 | 130 | 0.722 |
| 7 | 40 | 0.2589 | 37 | 41 | 39 | 2.55 | 169 | 1.086 |
| 8 | 43 | 0.1455 | 41 | 43 | 42 | 1.98 | 95 | 0.610 |
| 9 | 47 | 0.4659 | 43 | 51 | 47 | 3.54 | 199 | 1.954 |
| 10 | 59 | 0.1417 | 57 | 60 | 59 | 1.01 | 135 | 0.594 |
| 11 | 61 | 0.2042 | 60 | 62 | 61 | 0.99 | 200 | 0.856 |
| 12 | 63 | 0.1226 | 62 | 64 | 63 | 0.63 | 176 | 0.514 |
| 13 | 65 | 0.0626 | 64 | 66 | 65 | 0.77 | 78 | 0.263 |
| 14 | 67 | 0.0941 | 66 | 68 | 67 | 0.53 | 143 | 0.395 |
| 15 | 69 | 0.1381 | 68 | 71 | 69 | 0.68 | 213 | 0.579 |
| 16 | 77 | 1.2530 | 71 | 87 | 77 | 2.70 | 593 | 5.256 |
| 17 | 97 | 1.2397 | 87 | 100 | 96 | 2.24 | 456 | 5.200 |
| 18 | 102 | 0.1376 | 100 | 103 | 102 | 0.75 | 150 | 0.577 |
| 19 | 104 | 0.0914 | 103 | 105 | 104 | 0.61 | 80 | 0.383 |
| 20 | 106 | 0.1178 | 105 | 110 | 107 | 1.01 | 102 | 0.494 |
| 21 | 112 | 0.2633 | 110 | 115 | 113 | 0.96 | 174 | 1.104 |
| 22 | 116 | 0.2370 | 115 | 117 | 116 | 0.61 | 211 | 0.994 |
| 23 | 120 | 4.6780 | 117 | 129 | 121 | 1.86 | 4810 | 19.622 |
| 24 | 130 | 0.3384 | 129 | 137 | 130 | 1.16 | 238 | 1.419 |
| 25 | 146 | 1.7739 | 138 | 151 | 146 | 0.93 | 2394 | 7.441 |

```
Sample peak width (sec): 5     Sample min peak height: 50    Sample baseline V to V?: Y    Sample baseline V to V points: 3
Sample filter: Binomial        Number of points for filter: 3  Sample start region (min): 0  Sample end region (min): 80
Marker peak width (sec): 5     Marker min peak height: 500   Marker baseline V to V?: Y    Marker baseline V to V points: 3
Lower marker selection: First peak > 500 RFU                 Upper marker selection: Last peak > 500 RFU
Ladder size (bp)1, 35, 75, 100, 150, 200, 250, 300, 400, 500
Quantification using: Upper Marker         Final concentration (ng/uL): 0.5000         Dilution factor: 12.0
```

**Sample:** SampD2
**Well location:** D2
**Created:** Monday, 11 October 2021 14:41:52

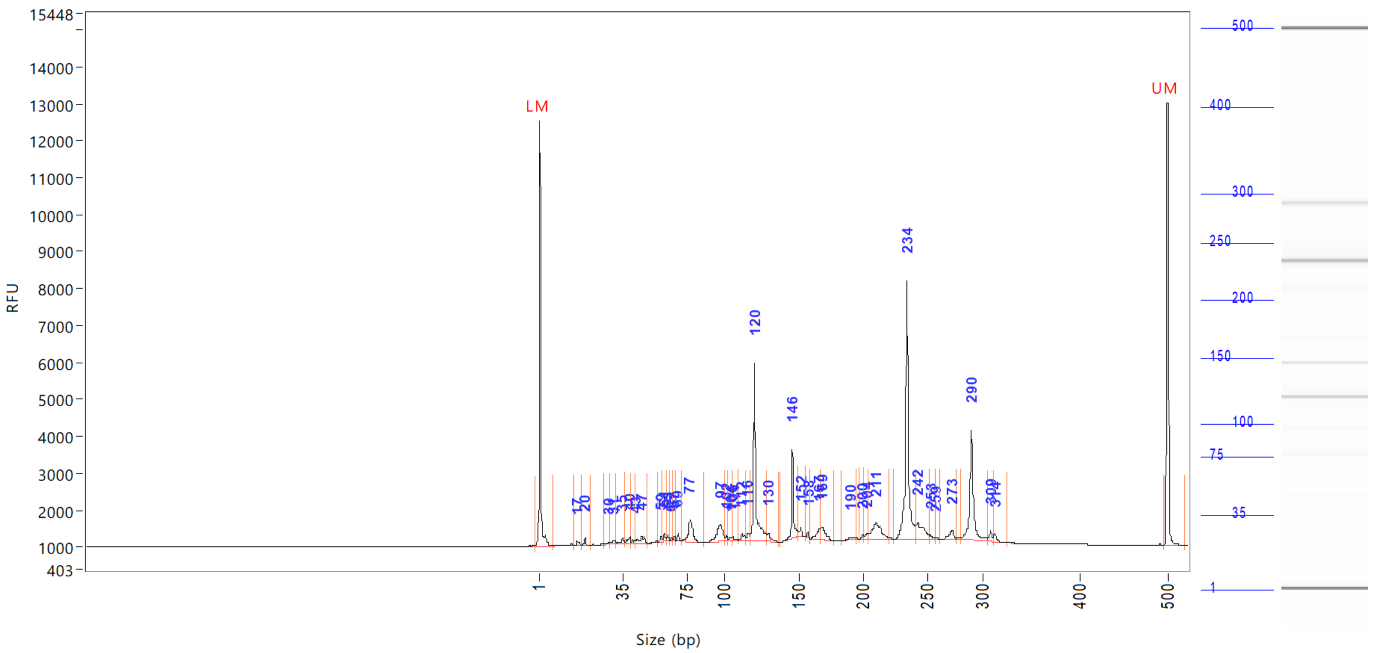


| Peak | Size (bp) | Concentration (ng/uL) | From (bp) | To (bp) | Average size (bp) | CV% | RFU | Corrected peak area |
|---|---|---|---|---|---|---|---|---|
| 26 | 152 | 0.1828 | 151 | 156 | 152 | 0.61 | 223 | |
| 27 | 158 | 0.0915 | 156 | 160 | 158 | 0.33 | 169 | |
| 28 | 167 | 0.4416 | 160 | 167 | 165 | 1.06 | 307 | |
| 29 | 169 | 0.6450 | 167 | 178 | 170 | 1.18 | 354 | |
| 30 | 190 | 0.1136 | 184 | 195 | 190 | 1.14 | 64 | |
| 31 | 200 | 0.0704 | 197 | 202 | 200 | 0.54 | 93 | |
| 32 | 204 | 0.1356 | 202 | 205 | 203 | 0.40 | 145 | |
| 33 | 211 | 1.3993 | 205 | 221 | 211 | 1.65 | 435 | |
| 34 | 234 | 6.7479 | 224 | 241 | 234 | 0.96 | 6991 | |
| 35 | 242 | 1.1736 | 241 | 252 | 245 | 1.16 | 466 | |
| 36 | 253 | 0.1614 | 252 | 258 | 254 | 0.58 | 133 | |
| 37 | 259 | 0.0590 | 258 | 263 | 259 | 0.41 | 60 | |
| 38 | 273 | 0.4017 | 263 | 277 | 271 | 0.89 | 238 | |
| 39 | 290 | 3.3988 | 280 | 306 | 290 | 1.18 | 2983 | |
| 40 | 309 | 0.2425 | 306 | 311 | 309 | 0.44 | 296 | |
| 41 | 314 | 0.1980 | 311 | 325 | 314 | 0.63 | 226 | |
| 42 | 500 (UM) | 0.5000 | 496 | 0 | 500 | 0.29 | 11986 | |

| | | |
|---|---|---|
| TIC: | 28.0325 | ng/uL |
| TIM: | 364.5267 | nmole/L |
| Total concentration: | 28.3679 | ng/uL |

```
Sample peak width (sec): 5    Sample min peak height: 50    Sample baseline V to V?: Y    Sample baseline V to V points: 3
Sample filter: Binomial       Number of points for filter: 3  Sample start region (min): 0  Sample end region (min): 80
Marker peak width (sec): 5    Marker min peak height: 500   Marker baseline V to V?: Y    Marker baseline V to V points: 3
Lower marker selection: First peak > 500 RFU                Upper marker selection: Last peak > 500 RFU
Ladder size (bp)1, 35, 75, 100, 150, 200, 250, 300, 400, 500
Quantification using: Upper Marker         Final concentration (ng/uL): 0.5000         Dilution factor: 12.0
```

**Sample:** SampE2
**Well location:** E2
**Created:** Monday, 11 October 2021 14:41:52

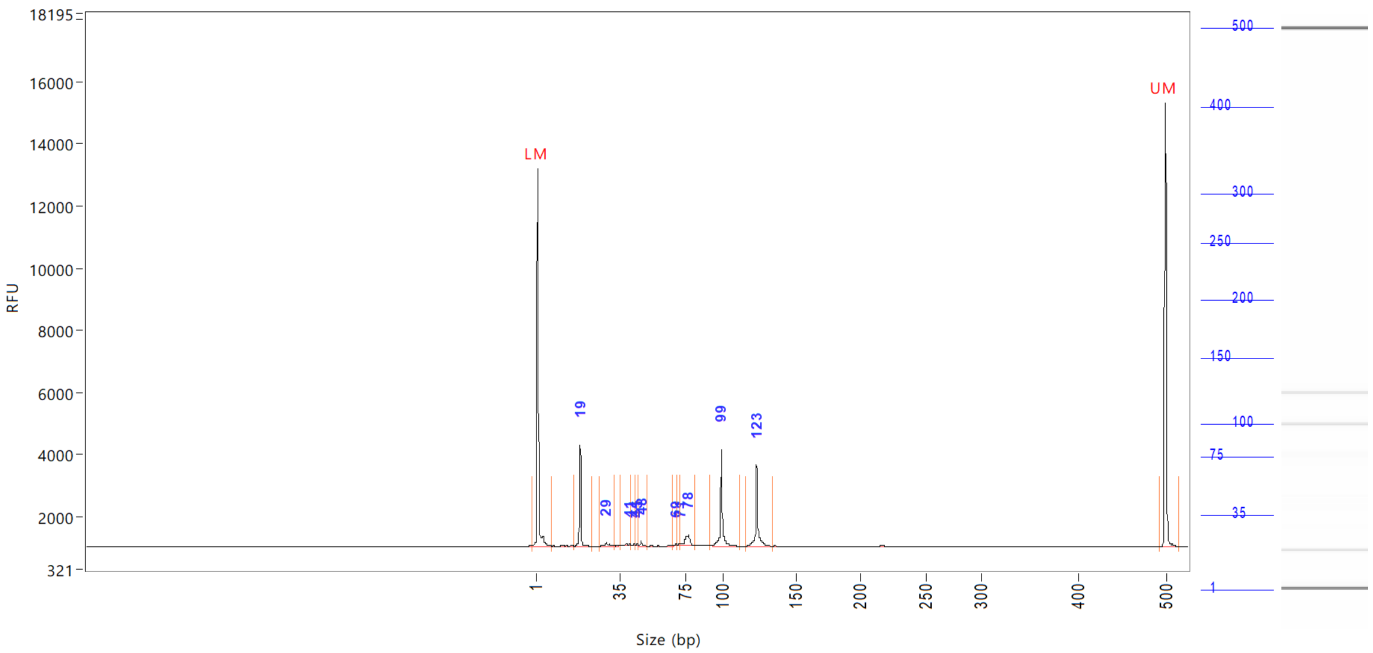


| Peak | Size (bp) | Concentration (ng/uL) | From (bp) | To (bp) | Average size (bp) | CV% | RFU | Corrected peak area |
|---|---|---|---|---|---|---|---|---|
| 1 | 1 (LM) | 0.7934 | 0 | 8 | 1 | 68.57 | 12154 | 45.879 |
| 2 | 19 | 2.4287 | 17 | 23 | 19 | 2.80 | 3257 | 11.703 |
| 3 | 29 | 0.1749 | 27 | 33 | 30 | 3.78 | 113 | 0.843 |
| 4 | 41 | 0.1540 | 35 | 42 | 40 | 3.00 | 84 | 0.742 |
| 5 | 44 | 0.0704 | 42 | 45 | 44 | 1.36 | 80 | 0.339 |
| 6 | 45 | 0.0623 | 45 | 46 | 45 | 1.10 | 70 | 0.300 |
| 7 | 48 | 0.1245 | 46 | 52 | 48 | 1.62 | 159 | 0.600 |
| 8 | 69 | 0.0608 | 67 | 70 | 69 | 1.13 | 59 | 0.293 |
| 9 | 71 | 0.0513 | 70 | 72 | 71 | 0.71 | 54 | 0.247 |
| 10 | 78 | 0.7383 | 72 | 82 | 76 | 2.60 | 336 | 3.558 |
| 11 | 99 | 2.3876 | 91 | 113 | 99 | 2.18 | 3095 | 11.505 |
| 12 | 123 | 2.0927 | 116 | 134 | 124 | 1.68 | 2605 | 10.084 |
| 13 | 500 (UM) | 0.5000 | 493 | 516 | 500 | 0.27 | 14291 | 28.913 |

| | | |
|---|---|---|
| TIC: | 8.3454 | ng/uL |
| TIM: | 318.4492 | nmole/L |
| Total concentration: | 8.9187 | ng/uL |

```
Sample peak width (sec): 5    Sample min peak height: 50    Sample baseline V to V?: Y    Sample baseline V to V points: 3
Sample filter: Binomial       Number of points for filter: 3  Sample start region (min): 0  Sample end region (min): 80
Marker peak width (sec): 5    Marker min peak height: 500   Marker baseline V to V?: Y    Marker baseline V to V points: 3
Lower marker selection: First peak > 500 RFU                Upper marker selection: Last peak > 500 RFU
Ladder size (bp)1, 35, 75, 100, 150, 200, 250, 300, 400, 500
Quantification using: Upper Marker         Final concentration (ng/uL): 0.5000         Dilution factor: 12.0
```

**Sample:** SampF2
**Well location:** F2
**Created:** Monday, 11 October 2021 14:41:52

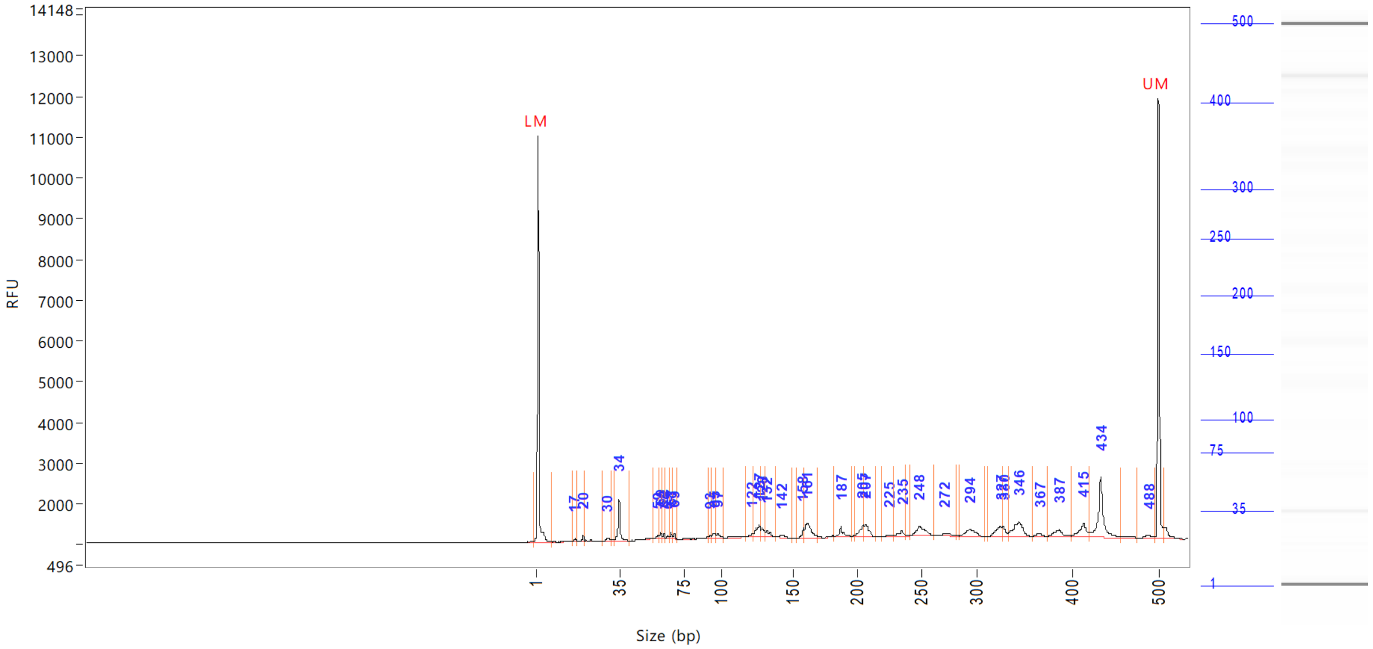


| Peak | Size (bp) | Concentration (ng/uL) | From (bp) | To (bp) | Average size (bp) | CV% | RFU | Corrected peak area |
|---|---|---|---|---|---|---|---|---|
| 1 | 1 (LM) | 0.8364 | 0 | 8 | 1 | 74.57 | 9974 | 37.688 |
| 2 | 17 | 0.0810 | 16 | 18 | 17 | 2.06 | 81 | 0.304 |
| 3 | 20 | 0.1802 | 18 | 21 | 20 | 2.26 | 158 | 0.677 |
| 4 | 30 | 0.1953 | 28 | 31 | 30 | 2.56 | 73 | 0.733 |
| 5 | 34 | 1.2982 | 32 | 42 | 34 | 3.52 | 1038 | 4.875 |
| 6 | 59 | 0.0874 | 56 | 59 | 58 | 1.35 | 66 | 0.328 |
| 7 | 60 | 0.1339 | 59 | 61 | 60 | 0.84 | 139 | 0.503 |
| 8 | 62 | 0.0885 | 61 | 63 | 62 | 0.60 | 111 | 0.332 |
| 9 | 65 | 0.0551 | 63 | 66 | 65 | 0.76 | 60 | 0.207 |
| 10 | 67 | 0.1218 | 66 | 68 | 67 | 0.80 | 137 | 0.457 |
| 11 | 69 | 0.0897 | 68 | 70 | 69 | 0.55 | 119 | 0.337 |
| 12 | 93 | 0.0585 | 92 | 93 | 93 | 0.52 | 61 | 0.220 |
| 13 | 95 | 0.1737 | 93 | 96 | 95 | 0.89 | 117 | 0.652 |
| 14 | 97 | 0.1780 | 96 | 103 | 98 | 1.25 | 118 | 0.668 |
| 15 | 122 | 0.1290 | 117 | 123 | 121 | 1.07 | 83 | 0.484 |
| 16 | 127 | 0.5815 | 123 | 127 | 125 | 1.02 | 294 | 2.184 |
| 17 | 128 | 0.3592 | 127 | 131 | 129 | 0.82 | 228 | 1.349 |
| 18 | 132 | 0.2589 | 131 | 137 | 133 | 0.85 | 171 | 0.972 |
| 19 | 142 | 0.1507 | 137 | 149 | 142 | 1.33 | 59 | 0.566 |
| 20 | 158 | 0.2342 | 153 | 159 | 157 | 0.72 | 232 | 0.879 |
| 21 | 161 | 0.9898 | 159 | 170 | 162 | 1.55 | 368 | 3.717 |
| 22 | 187 | 0.3726 | 182 | 195 | 188 | 1.30 | 257 | 1.399 |
| 23 | 205 | 0.4263 | 198 | 206 | 203 | 0.96 | 271 | 1.601 |
| 24 | 207 | 0.5676 | 206 | 215 | 208 | 0.85 | 290 | 2.131 |
| 25 | 225 | 0.1005 | 220 | 228 | 225 | 0.81 | 52 | 0.378 |

```
Sample peak width (sec): 5     Sample min peak height: 50    Sample baseline V to V?: Y    Sample baseline V to V points: 3
Sample filter: Binomial        Number of points for filter: 3  Sample start region (min): 0  Sample end region (min): 80
Marker peak width (sec): 5     Marker min peak height: 500   Marker baseline V to V?: Y    Marker baseline V to V points: 3
Lower marker selection: First peak > 500 RFU                 Upper marker selection: Last peak > 500 RFU
Ladder size (bp)1, 35, 75, 100, 150, 200, 250, 300, 400, 500
Quantification using: Upper Marker          Final concentration (ng/uL): 0.5000          Dilution factor: 12.0
```

**Sample:** SampF2
**Well location:** F2
**Created:** Monday, 11 October 2021 14:41:52

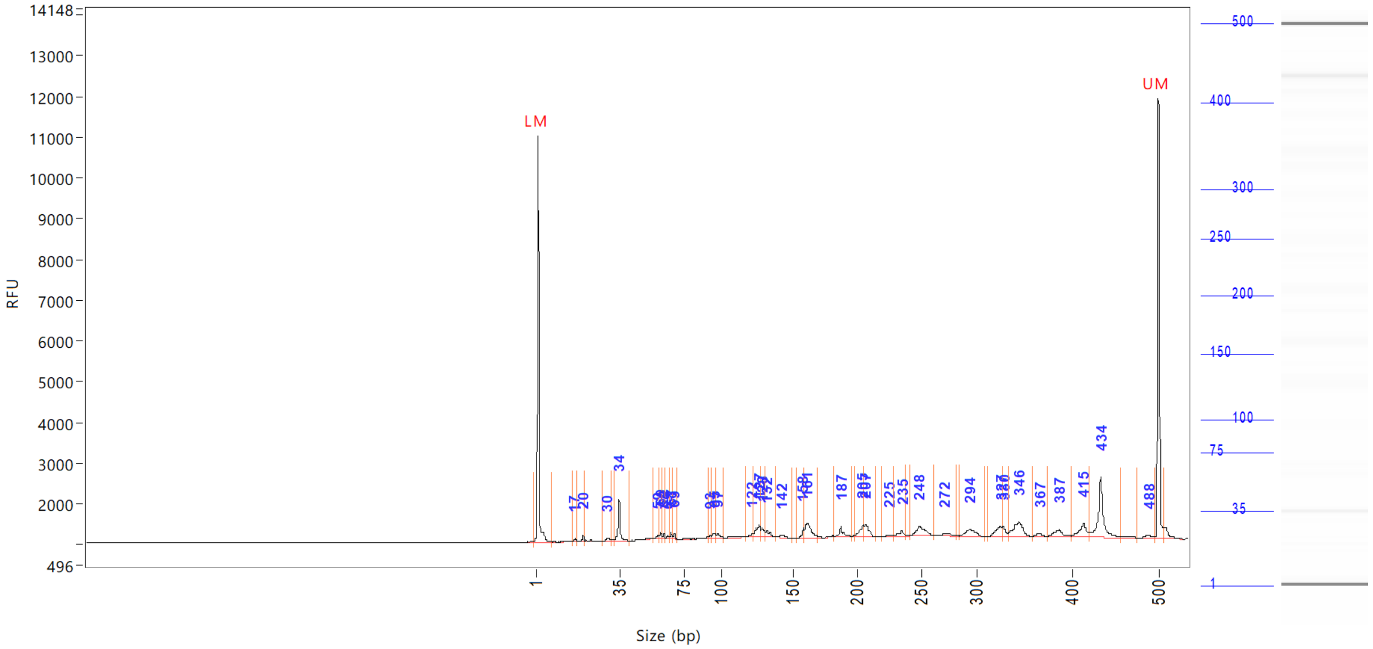


| Peak | Size (bp) | Concentration (ng/uL) | From (bp) | To (bp) | Average size (bp) | CV% | RFU | Corrected peak area |
|---|---|---|---|---|---|---|---|---|
| 26 | 235 | 0.2564 | 228 | 238 | 233 | 0.98 | 143 | |
| 27 | 248 | 0.7438 | 241 | 262 | 250 | 1.58 | 214 | |
| 28 | 272 | 0.1652 | 263 | 283 | 274 | 1.30 | 51 | |
| 29 | 294 | 0.5744 | 285 | 310 | 296 | 1.48 | 165 | |
| 30 | 327 | 0.5447 | 314 | 328 | 324 | 0.96 | 248 | |
| 31 | 330 | 0.2929 | 328 | 334 | 331 | 0.51 | 261 | |
| 32 | 346 | 1.1041 | 334 | 359 | 344 | 1.33 | 347 | |
| 33 | 367 | 0.1294 | 359 | 374 | 366 | 0.78 | 69 | |
| 34 | 387 | 0.4608 | 374 | 400 | 386 | 1.29 | 165 | |
| 35 | 415 | 0.7247 | 400 | 421 | 413 | 1.08 | 342 | |
| 36 | 434 | 2.2525 | 421 | 457 | 434 | 1.26 | 1474 | |
| 37 | 488 | 0.1632 | 476 | 496 | 488 | 0.83 | 77 | |
| 38 | 500 (UM) | 0.5000 | 496 | 506 | 500 | 0.26 | 10815 | |

| | | |
|---|---|---|
| TIC: | 14.3236 | ng/uL |
| TIM: | 195.6127 | nmole/L |
| Total concentration: | 15.1195 | ng/uL |

```
Sample peak width (sec): 5    Sample min peak height: 50    Sample baseline V to V?: Y    Sample baseline V to V points: 3
Sample filter: Binomial       Number of points for filter: 3  Sample start region (min): 0  Sample end region (min): 80
Marker peak width (sec): 5    Marker min peak height: 500   Marker baseline V to V?: Y    Marker baseline V to V points: 3
Lower marker selection: First peak > 500 RFU                Upper marker selection: Last peak > 500 RFU
Ladder size (bp)1, 35, 75, 100, 150, 200, 250, 300, 400, 500
Quantification using: Upper Marker         Final concentration (ng/uL): 0.5000          Dilution factor: 12.0
```

**Sample:** SampG2
**Well location:** G2
**Created:** Monday, 11 October 2021 14:41:52

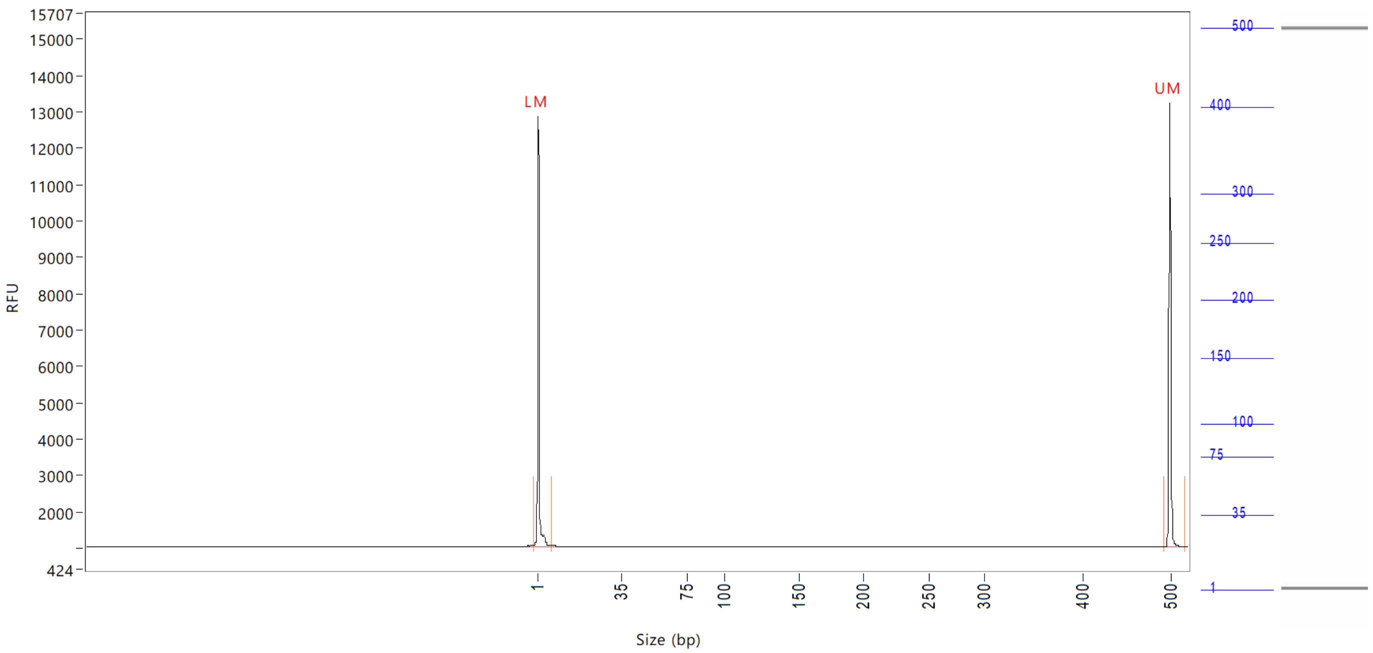


| Peak | Size (bp) | Concentration (ng/uL) | From (bp) | To (bp) | Average size (bp) | CV% | RFU | Corrected peak area |
|---|---|---|---|---|---|---|---|---|
| 1 | 1 (LM) | 0.8750 | 0 | 8 | 1 | 72.97 | 11810 | 45.228 |
| 2 | 500 (UM) | 0.5000 | 494 | 0 | 500 | 0.26 | 12220 | 25.845 |

| | | |
|---|---|---|
| TIC: | 0.0000 | ng/uL |
| TIM: | 0.0000 | nmole/L |
| Total concentration: | 0.4960 | ng/uL |

Sample peak width (sec): 5    Sample min peak height: 50    Sample baseline V to V?: Y    Sample baseline V to V points: 3
Sample filter: Binomial    Number of points for filter: 3  Sample start region (min): 0  Sample end region (min): 80
Marker peak width (sec): 5    Marker min peak height: 500    Marker baseline V to V?: Y    Marker baseline V to V points: 3
Lower marker selection: First peak > 500 RFU    Upper marker selection: Last peak > 500 RFU
Ladder size (bp)1, 35, 75, 100, 150, 200, 250, 300, 400, 500
Quantification using: Upper Marker    Final concentration (ng/uL): 0.5000    Dilution factor: 12.0

**Sample:** SampH2
**Well location:** H2
**Created:** Monday, 11 October 2021 14:41:52

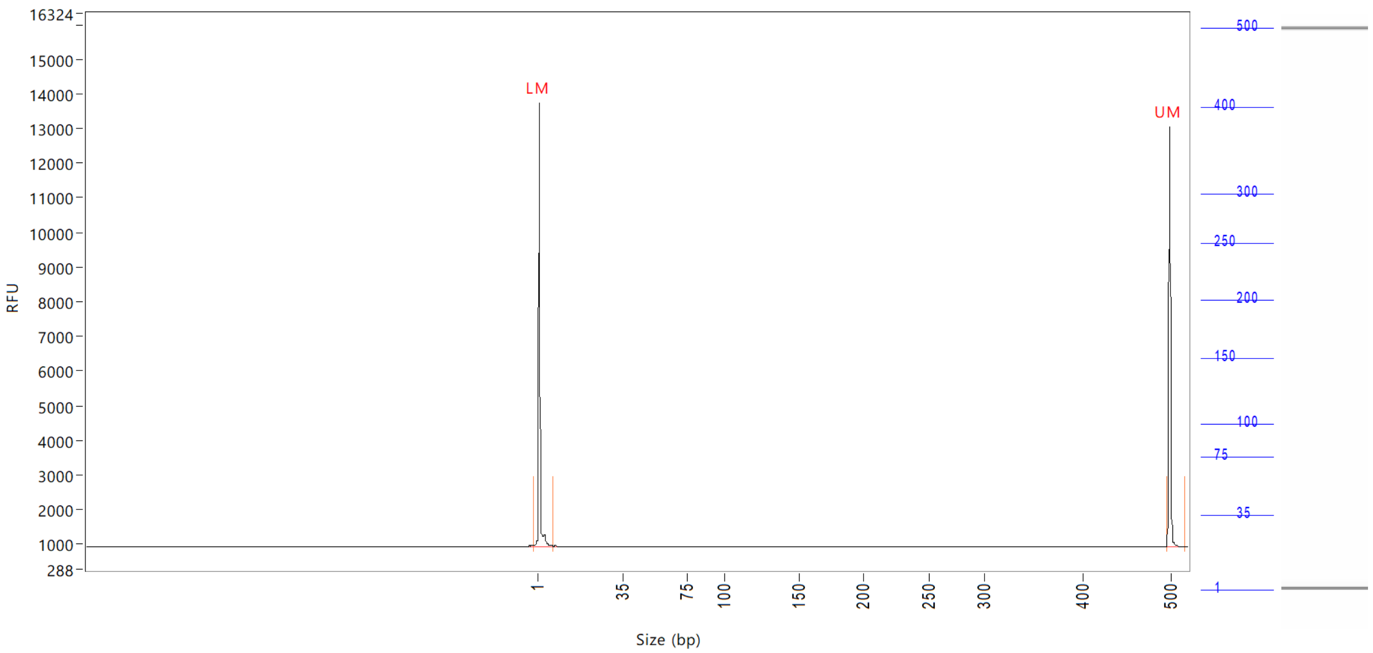


| Peak | Size (bp) | Concentration (ng/uL) | From (bp) | To (bp) | Average size (bp) | CV% | RFU | Corrected peak area |
|---|---|---|---|---|---|---|---|---|
| 1 | 1 (LM) | 0.9699 | 0 | 8 | 1 | 69.20 | 12812 | 48.177 |
| 2 | 500 (UM) | 0.5000 | 496 | 0 | 500 | 0.26 | 12115 | 24.835 |

| | | |
|---|---|---|
| TIC: | 0.0000 | ng/uL |
| TIM: | 0.0000 | nmole/L |
| Total concentration: | 0.4906 | ng/uL |

Sample peak width (sec): 5 Sample min peak height: 50 Sample baseline V to V?: Y Sample baseline V to V points: 3
Sample filter: Binomial Number of points for filter: 3 Sample start region (min): 0 Sample end region (min): 80
Marker peak width (sec): 5 Marker min peak height: 500 Marker baseline V to V?: Y Marker baseline V to V points: 3
Lower marker selection: First peak > 500 RFU Upper marker selection: Last peak > 500 RFU
Ladder size (bp)1, 35, 75, 100, 150, 200, 250, 300, 400, 500
Quantification using: Upper Marker Final concentration (ng/uL): 0.5000 Dilution factor: 12.0

**Sample:** SampA3
**Well location:** A3
**Created:** Monday, 11 October 2021 14:41:52

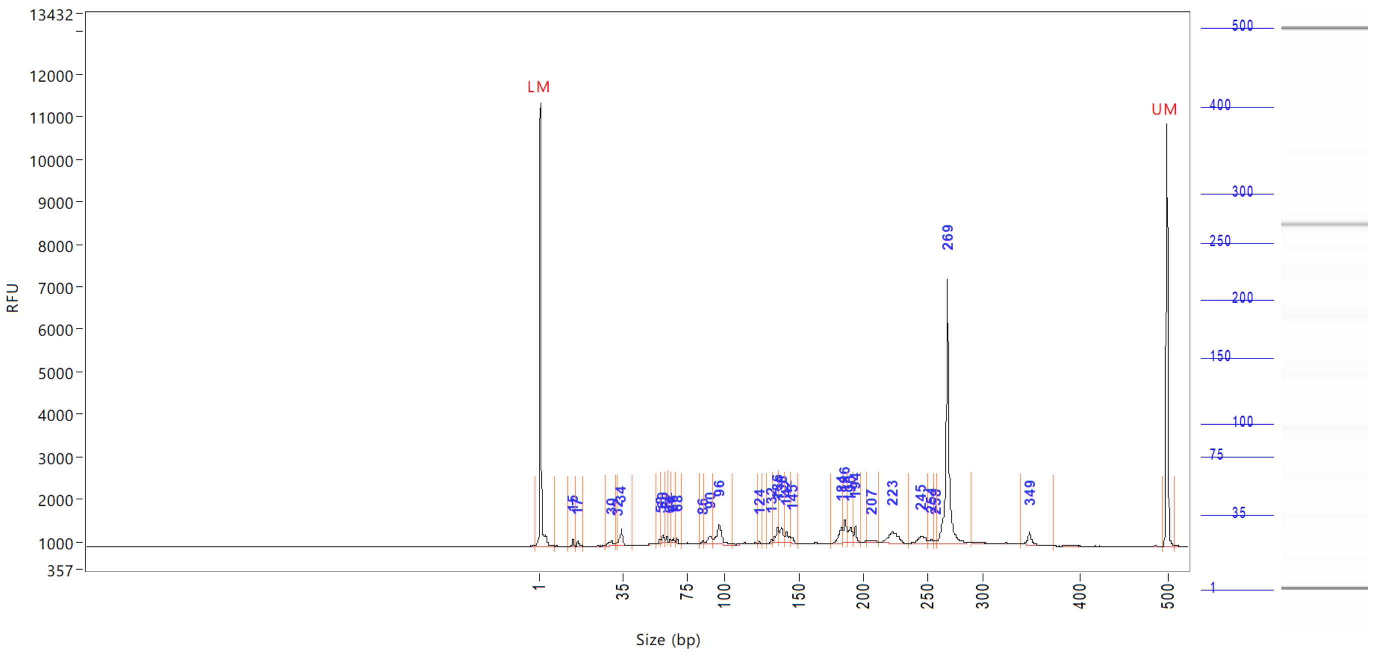

| Peak | Size (bp) | Concentration (ng/uL) | From (bp) | To (bp) | Average size (bp) | CV% | RFU | Corrected peak area |
|---|---|---|---|---|---|---|---|---|
| 1 | 1 (LM) | 0.9286 | 0 | 8 | 1 | 71.61 | 10449 | 38.361 |
| 2 | 15 | 0.1962 | 13 | 16 | 15 | 2.43 | 180 | 0.675 |
| 3 | 17 | 0.1693 | 16 | 19 | 17 | 3.21 | 128 | 0.583 |
| 4 | 30 | 0.4142 | 28 | 32 | 30 | 3.31 | 129 | 1.426 |
| 5 | 32 | 0.1148 | 32 | 33 | 32 | 0.91 | 90 | 0.395 |
| 6 | 34 | 0.6871 | 33 | 41 | 34 | 2.87 | 396 | 2.365 |
| 7 | 58 | 0.1381 | 56 | 59 | 58 | 1.14 | 102 | 0.476 |
| 8 | 60 | 0.2068 | 59 | 61 | 60 | 0.92 | 187 | 0.712 |
| 9 | 62 | 0.1370 | 61 | 63 | 62 | 0.62 | 153 | 0.471 |
| 10 | 64 | 0.0712 | 63 | 65 | 64 | 0.84 | 70 | 0.245 |
| 11 | 66 | 0.0918 | 65 | 67 | 66 | 0.68 | 106 | 0.316 |
| 12 | 68 | 0.0960 | 67 | 71 | 68 | 0.67 | 114 | 0.330 |
| 13 | 86 | 0.1156 | 83 | 86 | 85 | 0.89 | 93 | 0.398 |
| 14 | 90 | 0.5249 | 86 | 92 | 90 | 1.59 | 203 | 1.807 |
| 15 | 96 | 1.3117 | 92 | 106 | 96 | 2.17 | 490 | 4.516 |
| 16 | 124 | 0.0516 | 122 | 126 | 124 | 0.45 | 67 | 0.178 |
| 17 | 132 | 0.0924 | 128 | 133 | 132 | 0.52 | 95 | 0.318 |
| 18 | 136 | 0.5423 | 133 | 137 | 135 | 0.83 | 354 | 1.867 |
| 19 | 138 | 0.5155 | 137 | 141 | 139 | 0.65 | 333 | 1.774 |
| 20 | 142 | 0.4163 | 141 | 145 | 142 | 0.79 | 241 | 1.433 |
| 21 | 145 | 0.1898 | 145 | 150 | 146 | 0.60 | 138 | 0.653 |
| 22 | 184 | 0.7119 | 176 | 185 | 183 | 0.93 | 365 | 2.451 |
| 23 | 186 | 0.7352 | 185 | 189 | 187 | 0.54 | 561 | 2.531 |
| 24 | 190 | 0.6164 | 189 | 193 | 191 | 0.62 | 374 | 2.122 |
| 25 | 194 | 0.3817 | 193 | 198 | 194 | 0.46 | 414 | 1.314 |

```
Sample peak width (sec): 5    Sample min peak height: 50    Sample baseline V to V?: Y    Sample baseline V to V points: 3
Sample filter: Binomial       Number of points for filter: 3  Sample start region (min): 0  Sample end region (min): 80
Marker peak width (sec): 5    Marker min peak height: 500   Marker baseline V to V?: Y    Marker baseline V to V points: 3
Lower marker selection: First peak > 500 RFU                Upper marker selection: Last peak > 500 RFU
Ladder size (bp)1, 35, 75, 100, 150, 200, 250, 300, 400, 500
Quantification using: Upper Marker          Final concentration (ng/uL): 0.5000          Dilution factor: 12.0
```

**Sample:** SampA3
**Well location:** A3
**Created:** Monday, 11 October 2021 14:41:52

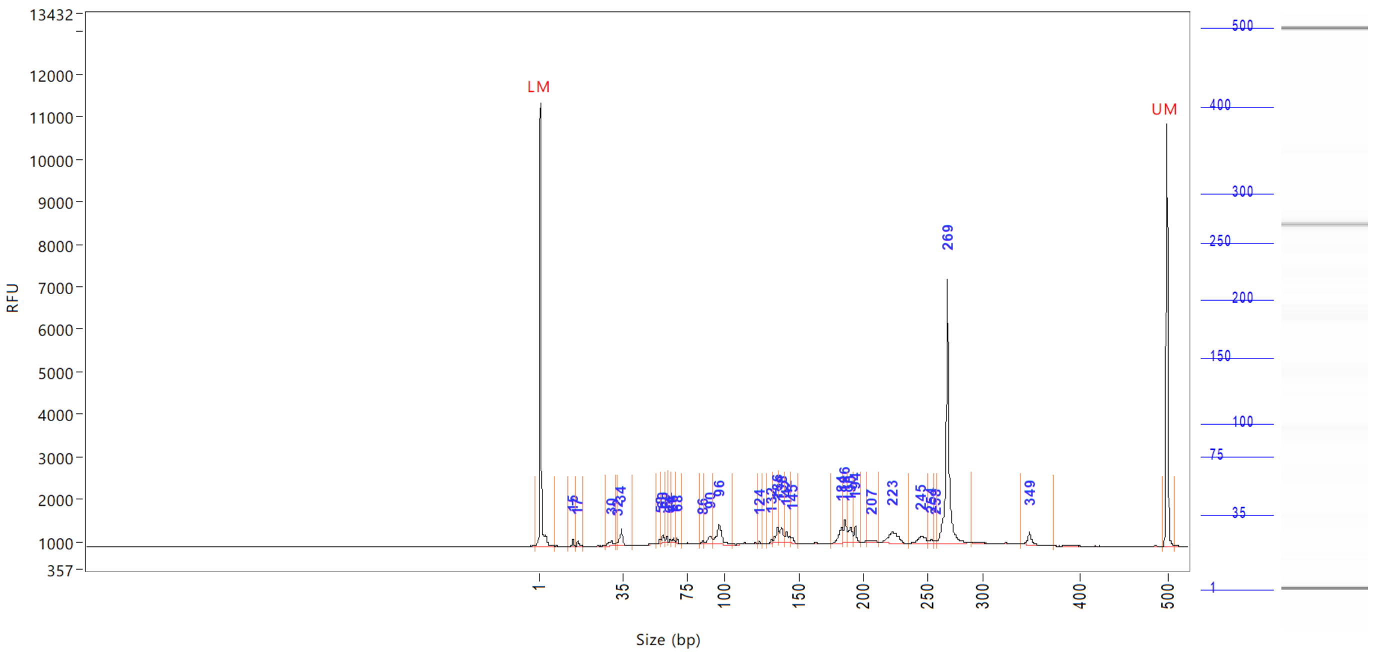


| Peak | Size (bp) | Concentration (ng/uL) | From (bp) | To (bp) | Average size (bp) | CV% | RFU | Corrected peak area |
|---|---|---|---|---|---|---|---|---|
| 26 | 207 | 0.1739 | 203 | 213 | 208 | 1.08 | 55 | |
| 27 | 223 | 1.0920 | 213 | 235 | 224 | 1.65 | 257 | |
| 28 | 245 | 0.5689 | 235 | 252 | 245 | 1.23 | 173 | |
| 29 | 254 | 0.1208 | 252 | 256 | 254 | 0.46 | 101 | |
| 30 | 258 | 0.0661 | 256 | 259 | 258 | 0.36 | 62 | |
| 31 | 269 | 7.3607 | 259 | 290 | 269 | 1.14 | 6218 | |
| 32 | 349 | 0.4092 | 340 | 374 | 349 | 0.94 | 294 | |
| 33 | 500 (UM) | 0.5000 | 495 | 509 | 500 | 0.24 | 9953 | |

| | | |
|---|---|---|
| TIC: | 18.3192 | ng/uL |
| TIM: | 256.8022 | nmole/L |
| Total concentration: | 18.9180 | ng/uL |

```
Sample peak width (sec): 5    Sample min peak height: 50    Sample baseline V to V?: Y    Sample baseline V to V points: 3
Sample filter: Binomial       Number of points for filter: 3  Sample start region (min): 0  Sample end region (min): 80
Marker peak width (sec): 5    Marker min peak height: 500   Marker baseline V to V?: Y    Marker baseline V to V points: 3
Lower marker selection: First peak > 500 RFU                Upper marker selection: Last peak > 500 RFU
Ladder size (bp)1, 35, 75, 100, 150, 200, 250, 300, 400, 500
Quantification using: Upper Marker         Final concentration (ng/uL): 0.5000         Dilution factor: 12.0
```

**Sample:** SampB3
**Well location:** B3
**Created:** Monday, 11 October 2021 14:41:52

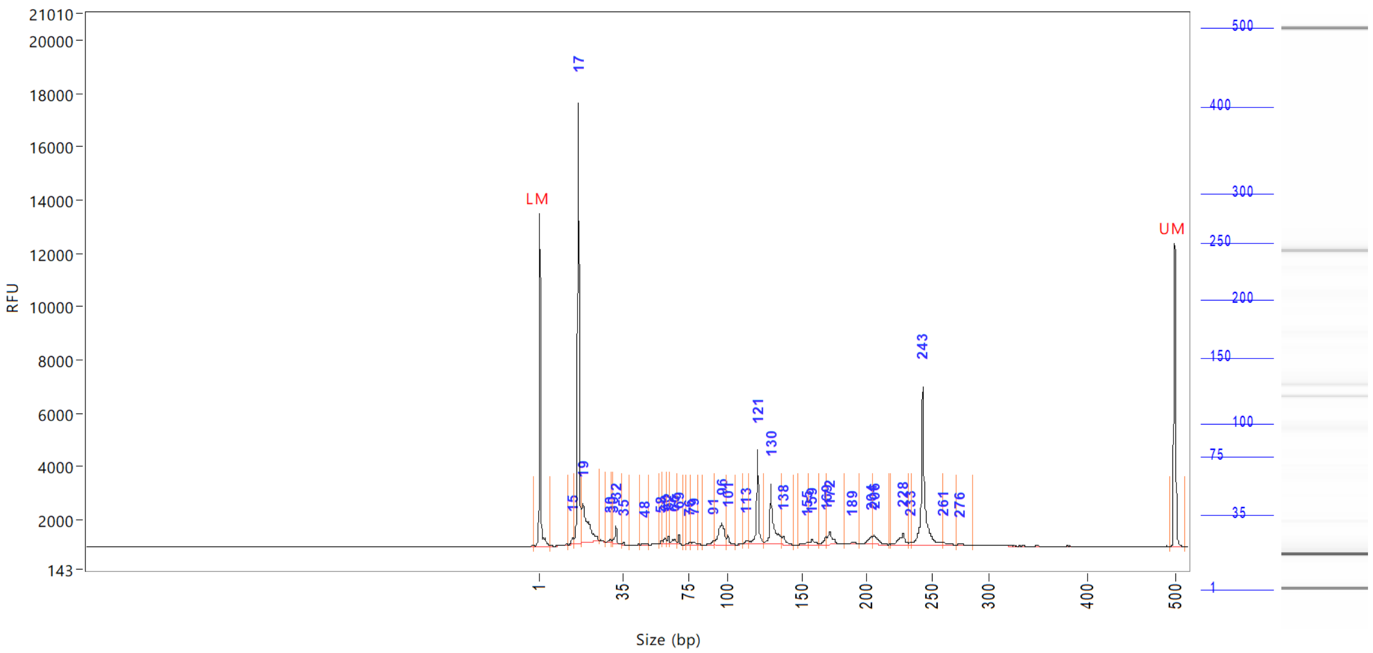

| Peak | Size (bp) | Concentration (ng/uL) | From (bp) | To (bp) | Average size (bp) | CV% | RFU | Corrected peak area |
|---|---|---|---|---|---|---|---|---|
| 1 | 1 (LM) | 0.9553 | 0 | 6 | 1 | 66.22 | 12506 | 46.525 |
| 2 | 15 | 0.3317 | 13 | 15 | 15 | 3.16 | 221 | 1.346 |
| 3 | 17 | 16.0961 | 15 | 18 | 17 | 2.46 | 16561 | 65.324 |
| 4 | 19 | 5.3433 | 18 | 25 | 20 | 7.98 | 1450 | 21.685 |
| 5 | 30 | 0.1347 | 28 | 30 | 29 | 1.69 | 100 | 0.547 |
| 6 | 30 | 0.1378 | 30 | 31 | 30 | 0.91 | 128 | 0.559 |
| 7 | 32 | 0.6767 | 31 | 34 | 32 | 1.76 | 656 | 2.746 |
| 8 | 35 | 0.1044 | 34 | 39 | 35 | 1.73 | 98 | 0.424 |
| 9 | 48 | 0.1016 | 45 | 50 | 48 | 2.25 | 62 | 0.412 |
| 10 | 58 | 0.1249 | 57 | 59 | 58 | 0.94 | 115 | 0.507 |
| 11 | 60 | 0.2494 | 59 | 61 | 60 | 1.02 | 217 | 1.012 |
| 12 | 62 | 0.1941 | 61 | 63 | 62 | 0.72 | 239 | 0.788 |
| 13 | 66 | 0.2731 | 63 | 67 | 65 | 1.46 | 190 | 1.108 |
| 14 | 69 | 0.2522 | 67 | 71 | 68 | 0.63 | 357 | 1.024 |
| 15 | 76 | 0.1162 | 73 | 77 | 75 | 1.48 | 89 | 0.471 |
| 16 | 79 | 0.1906 | 77 | 81 | 79 | 1.28 | 117 | 0.774 |
| 17 | 91 | 0.1793 | 85 | 92 | 90 | 1.66 | 138 | 0.728 |
| 18 | 96 | 2.1994 | 92 | 99 | 96 | 1.82 | 845 | 8.926 |
| 19 | 101 | 0.3958 | 99 | 107 | 101 | 1.31 | 398 | 1.606 |
| 20 | 113 | 0.1922 | 111 | 115 | 114 | 0.88 | 134 | 0.780 |
| 21 | 121 | 2.9816 | 115 | 125 | 121 | 1.14 | 3520 | 12.100 |
| 22 | 130 | 3.4230 | 125 | 137 | 131 | 1.83 | 2278 | 13.892 |
| 23 | 138 | 0.4448 | 137 | 145 | 139 | 1.06 | 297 | 1.805 |
| 24 | 155 | 0.1324 | 148 | 156 | 154 | 1.15 | 88 | 0.537 |
| 25 | 159 | 0.3340 | 156 | 164 | 159 | 1.15 | 191 | 1.355 |

Sample peak width (sec): 5 Sample min peak height: 50 Sample baseline V to V?: Y Sample baseline V to V points: 3
Sample filter: Binomial Number of points for filter: 3 Sample start region (min): 0 Sample end region (min): 80
Marker peak width (sec): 5 Marker min peak height: 500 Marker baseline V to V?: Y Marker baseline V to V points: 3
Lower marker selection: First peak > 500 RFU Upper marker selection: Last peak > 500 RFU
Ladder size (bp)1, 35, 75, 100, 150, 200, 250, 300, 400, 500
Quantification using: Upper Marker Final concentration (ng/uL): 0.5000 Dilution factor: 12.0

**Sample:** SampB3
**Well location:** B3
**Created:** Monday, 11 October 2021 14:41:52

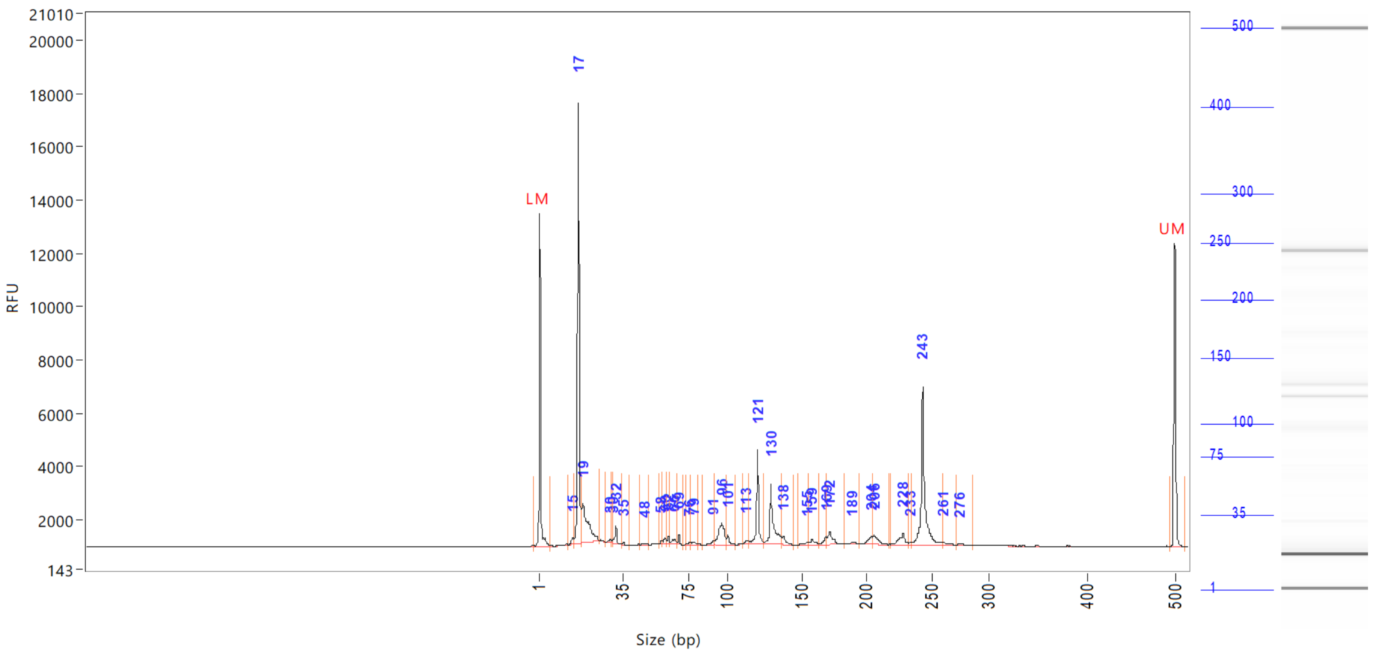


| Peak | Size (bp) | Concentration (ng/uL) | From (bp) | To (bp) | Average size (bp) | CV% | RFU | Corrected peak area |
|---|---|---|---|---|---|---|---|---|
| 26 | 169 | 0.3937 | 164 | 170 | 168 | 0.85 | 306 | |
| 27 | 172 | 0.8780 | 170 | 184 | 173 | 1.29 | 493 | |
| 28 | 189 | 0.2084 | 184 | 194 | 190 | 1.19 | 88 | |
| 29 | 204 | 0.4896 | 194 | 205 | 202 | 1.19 | 288 | |
| 30 | 206 | 0.6893 | 205 | 217 | 208 | 1.06 | 346 | |
| 31 | 228 | 0.7979 | 219 | 232 | 227 | 1.11 | 439 | |
| 32 | 233 | 0.0459 | 232 | 234 | 233 | 0.23 | 63 | |
| 33 | 243 | 6.7030 | 234 | 260 | 244 | 1.38 | 5907 | |
| 34 | 261 | 0.1518 | 260 | 272 | 264 | 1.13 | 84 | |
| 35 | 276 | 0.0871 | 272 | 286 | 277 | 0.89 | 53 | |
| 36 | 500 (UM) | 0.5000 | 495 | 512 | 500 | 0.24 | 11404 | |

| | | |
|---|---|---|
| TIC: | 45.0539 | ng/uL |
| TIM: | 2318.0474 | nmole/L |
| Total concentration: | 45.2298 | ng/uL |

Sample peak width (sec): 5 Sample min peak height: 50 Sample baseline V to V?: Y Sample baseline V to V points: 3
Sample filter: Binomial Number of points for filter: 3 Sample start region (min): 0 Sample end region (min): 80
Marker peak width (sec): 5 Marker min peak height: 500 Marker baseline V to V?: Y Marker baseline V to V points: 3
Lower marker selection: First peak > 500 RFU Upper marker selection: Last peak > 500 RFU
Ladder size (bp)1, 35, 75, 100, 150, 200, 250, 300, 400, 500
Quantification using: Upper Marker Final concentration (ng/uL): 0.5000 Dilution factor: 12.0

**Sample:** SampC3
**Well location:** C3
**Created:** Monday, 11 October 2021 14:41:52

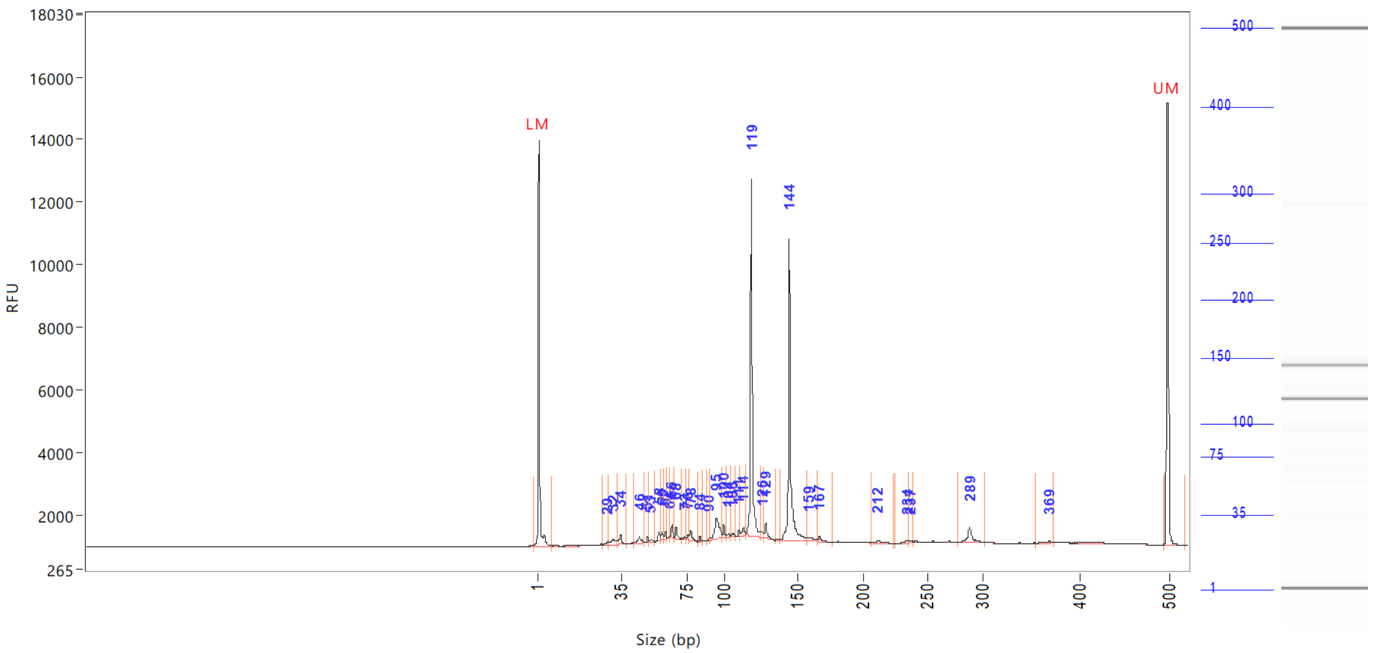


| Peak | Size (bp) | Concentration (ng/uL) | From (bp) | To (bp) | Average size (bp) | CV% | RFU | Corrected peak area |
|---|---|---|---|---|---|---|---|---|
| 1 | 1 (LM) | 0.8094 | 0 | 7 | 2 | 65.38 | 12974 | 49.134 |
| 2 | 29 | 0.1530 | 27 | 30 | 29 | 2.25 | 115 | 0.774 |
| 3 | 32 | 0.4037 | 30 | 33 | 31 | 2.89 | 193 | 2.042 |
| 4 | 34 | 0.3933 | 33 | 39 | 34 | 2.97 | 326 | 1.990 |
| 5 | 46 | 0.2953 | 43 | 50 | 46 | 2.67 | 205 | 1.494 |
| 6 | 51 | 0.1221 | 50 | 52 | 51 | 0.96 | 182 | 0.618 |
| 7 | 53 | 0.0665 | 52 | 56 | 53 | 1.06 | 87 | 0.336 |
| 8 | 58 | 0.1988 | 56 | 59 | 58 | 0.86 | 291 | 1.006 |
| 9 | 60 | 0.1992 | 59 | 61 | 60 | 0.98 | 240 | 1.008 |
| 10 | 62 | 0.1325 | 61 | 63 | 62 | 0.59 | 227 | 0.670 |
| 11 | 64 | 0.0549 | 63 | 65 | 64 | 0.67 | 89 | 0.278 |
| 12 | 66 | 0.2774 | 65 | 67 | 66 | 0.75 | 459 | 1.403 |
| 13 | 68 | 0.2859 | 67 | 72 | 69 | 1.24 | 374 | 1.446 |
| 14 | 73 | 0.0550 | 72 | 74 | 73 | 0.70 | 78 | 0.278 |
| 15 | 76 | 0.1399 | 74 | 77 | 76 | 1.20 | 173 | 0.707 |
| 16 | 78 | 0.2913 | 77 | 83 | 79 | 1.37 | 307 | 1.474 |
| 17 | 84 | 0.0932 | 83 | 86 | 84 | 0.68 | 150 | 0.471 |
| 18 | 90 | 0.0460 | 89 | 91 | 90 | 0.48 | 80 | 0.233 |
| 19 | 95 | 1.0959 | 91 | 98 | 95 | 1.59 | 666 | 5.544 |
| 20 | 100 | 0.2562 | 98 | 102 | 100 | 0.83 | 420 | 1.296 |
| 21 | 104 | 0.1035 | 102 | 105 | 103 | 0.73 | 103 | 0.524 |
| 22 | 106 | 0.1170 | 105 | 109 | 106 | 0.81 | 113 | 0.592 |
| 23 | 111 | 0.1199 | 109 | 112 | 111 | 0.52 | 191 | 0.606 |
| 24 | 114 | 0.2625 | 112 | 115 | 113 | 0.86 | 261 | 1.328 |
| 25 | 119 | 7.1333 | 115 | 125 | 119 | 1.03 | 11389 | 36.085 |

```
Sample peak width (sec): 5     Sample min peak height: 50    Sample baseline V to V?: Y    Sample baseline V to V points: 3
Sample filter: Binomial        Number of points for filter: 3  Sample start region (min): 0  Sample end region (min): 80
Marker peak width (sec): 5     Marker min peak height: 500   Marker baseline V to V?: Y    Marker baseline V to V points: 3
Lower marker selection: First peak > 500 RFU                 Upper marker selection: Last peak > 500 RFU
Ladder size (bp)1, 35, 75, 100, 150, 200, 250, 300, 400, 500
Quantification using: Upper Marker          Final concentration (ng/uL): 0.5000          Dilution factor: 12.0
```

**Sample:** SampC3
**Well location:** C3
**Created:** Monday, 11 October 2021 14:41:52

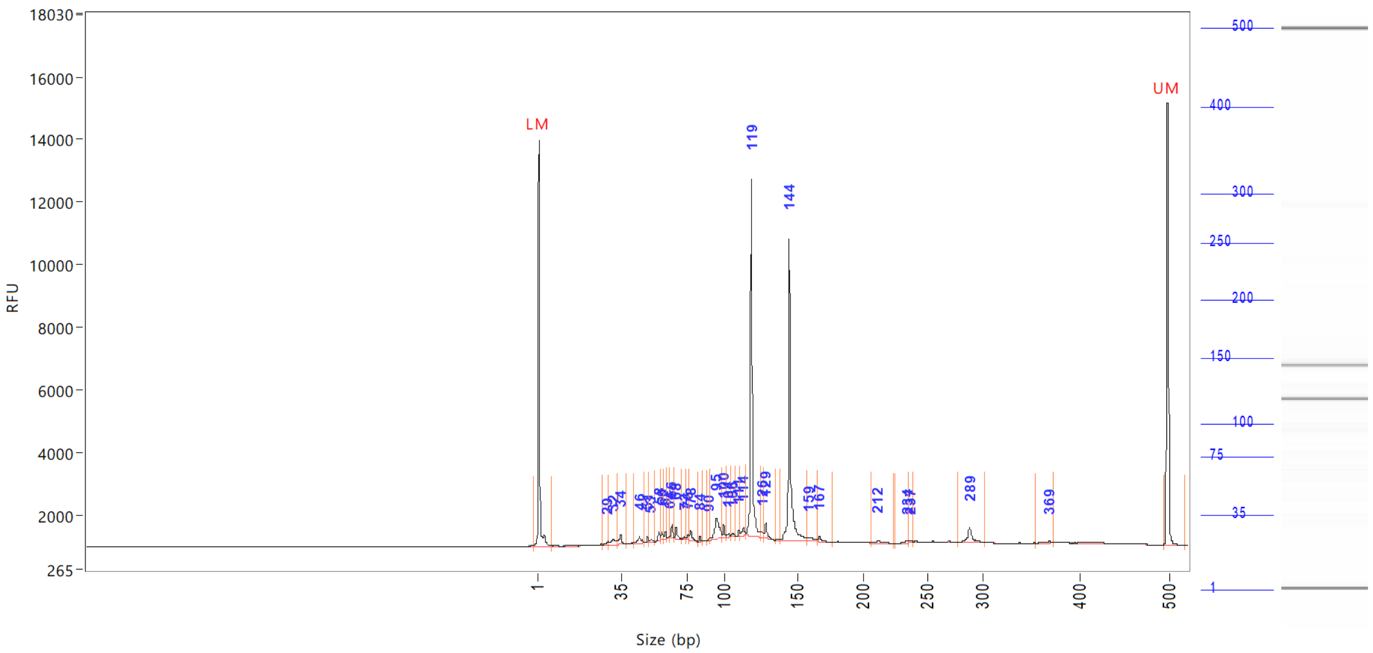


| Peak | Size (bp) | Concentration (ng/uL) | From (bp) | To (bp) | Average size (bp) | CV% | RFU | Corrected peak area |
|---|---|---|---|---|---|---|---|---|
| 26 | 126 | 0.1457 | 125 | 127 | 126 | 0.48 | 185 | |
| 27 | 129 | 0.3953 | 127 | 135 | 129 | 0.84 | 470 | |
| 28 | 144 | 6.8420 | 138 | 158 | 145 | 1.82 | 9630 | |
| 29 | 159 | 0.2529 | 158 | 166 | 162 | 1.41 | 98 | |
| 30 | 167 | 0.1532 | 166 | 178 | 168 | 1.05 | 149 | |
| 31 | 212 | 0.1595 | 207 | 224 | 214 | 1.60 | 101 | |
| 32 | 234 | 0.0901 | 226 | 235 | 233 | 0.78 | 78 | |
| 33 | 237 | 0.0797 | 235 | 240 | 237 | 0.53 | 71 | |
| 34 | 289 | 0.5564 | 278 | 303 | 289 | 1.16 | 469 | |
| 35 | 369 | 0.0769 | 356 | 373 | 367 | 1.07 | 75 | |
| 36 | 500 (UM) | 0.5000 | 496 | 0 | 500 | 0.27 | 14151 | |

| | | |
|---|---|---|
| TIC: | 21.0481 | ng/uL |
| TIM: | 333.0319 | nmole/L |
| Total concentration: | 21.7913 | ng/uL |

Sample peak width (sec): 5    Sample min peak height: 50    Sample baseline V to V?: Y    Sample baseline V to V points: 3
Sample filter: Binomial    Number of points for filter: 3  Sample start region (min): 0  Sample end region (min): 80
Marker peak width (sec): 5    Marker min peak height: 500    Marker baseline V to V?: Y    Marker baseline V to V points: 3
Lower marker selection: First peak > 500 RFU    Upper marker selection: Last peak > 500 RFU
Ladder size (bp)1, 35, 75, 100, 150, 200, 250, 300, 400, 500
Quantification using: Upper Marker    Final concentration (ng/uL): 0.5000    Dilution factor: 12.0

**Sample:** SampD3
**Well location:** D3
**Created:** Monday, 11 October 2021 14:41:52

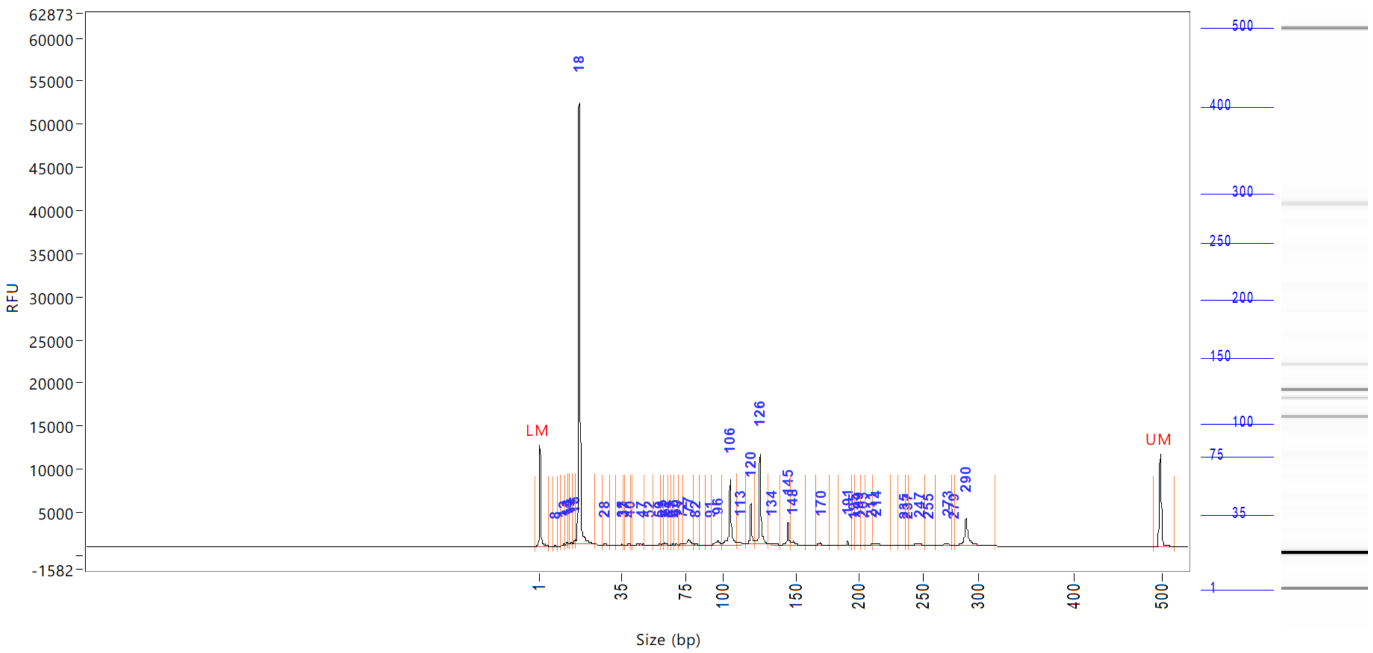

| Peak | Size (bp) | Concentration (ng/uL) | From (bp) | To (bp) | Average size (bp) | CV% | RFU | Corrected peak area |
|---|---|---|---|---|---|---|---|---|
| 1 | 1 (LM) | 0.8777 | 0 | 6 | 1 | 69.36 | 11798 | 43.595 |
| 2 | 8 | 0.0968 | 7 | 9 | 8 | 5.11 | 82 | 0.401 |
| 3 | 12 | 0.1494 | 10 | 12 | 11 | 2.91 | 198 | 0.618 |
| 4 | 13 | 0.1868 | 12 | 13 | 13 | 2.17 | 237 | 0.773 |
| 5 | 14 | 0.0795 | 13 | 14 | 14 | 1.41 | 124 | 0.329 |
| 6 | 14 | 0.2244 | 14 | 15 | 14 | 1.87 | 272 | 0.929 |
| 7 | 16 | 0.6383 | 15 | 16 | 16 | 2.28 | 509 | 2.642 |
| 8 | 18 | 51.5506 | 16 | 24 | 18 | 3.92 | 51233 | 213.372 |
| 9 | 28 | 0.2317 | 27 | 30 | 28 | 1.69 | 228 | 0.959 |
| 10 | 35 | 0.2015 | 32 | 36 | 34 | 2.55 | 133 | 0.834 |
| 11 | 37 | 0.0758 | 36 | 37 | 37 | 0.83 | 105 | 0.314 |
| 12 | 40 | 0.3094 | 37 | 41 | 39 | 2.33 | 205 | 1.281 |
| 13 | 47 | 0.5252 | 42 | 50 | 46 | 3.82 | 177 | 2.174 |
| 14 | 52 | 0.1104 | 50 | 55 | 52 | 1.58 | 109 | 0.457 |
| 15 | 59 | 0.1559 | 55 | 60 | 58 | 1.37 | 139 | 0.645 |
| 16 | 61 | 0.1854 | 60 | 61 | 60 | 0.71 | 219 | 0.768 |
| 17 | 62 | 0.3198 | 61 | 64 | 62 | 0.95 | 265 | 1.324 |
| 18 | 65 | 0.0427 | 64 | 66 | 65 | 0.73 | 59 | 0.177 |
| 19 | 67 | 0.0737 | 66 | 68 | 67 | 0.51 | 119 | 0.305 |
| 20 | 69 | 0.1729 | 68 | 70 | 69 | 0.63 | 231 | 0.716 |
| 21 | 72 | 0.2068 | 70 | 73 | 72 | 0.83 | 218 | 0.856 |
| 22 | 77 | 1.3411 | 73 | 81 | 77 | 2.39 | 626 | 5.551 |
| 23 | 82 | 0.1926 | 81 | 85 | 82 | 1.01 | 140 | 0.797 |
| 24 | 91 | 0.0806 | 89 | 92 | 91 | 0.83 | 72 | 0.334 |
| 25 | 96 | 0.9893 | 92 | 99 | 96 | 1.66 | 409 | 4.095 |

```
Sample peak width (sec): 5     Sample min peak height: 50    Sample baseline V to V?: Y     Sample baseline V to V points: 3
Sample filter: Binomial        Number of points for filter: 3  Sample start region (min): 0  Sample end region (min): 80
Marker peak width (sec): 5     Marker min peak height: 500   Marker baseline V to V?: Y     Marker baseline V to V points: 3
Lower marker selection: First peak > 500 RFU                 Upper marker selection: Last peak > 500 RFU
Ladder size (bp)1, 35, 75, 100, 150, 200, 250, 300, 400, 500
Quantification using: Upper Marker          Final concentration (ng/uL): 0.5000          Dilution factor: 12.0
```

**Sample:** SampD3
**Well location:** D3
**Created:** Monday, 11 October 2021 14:41:52

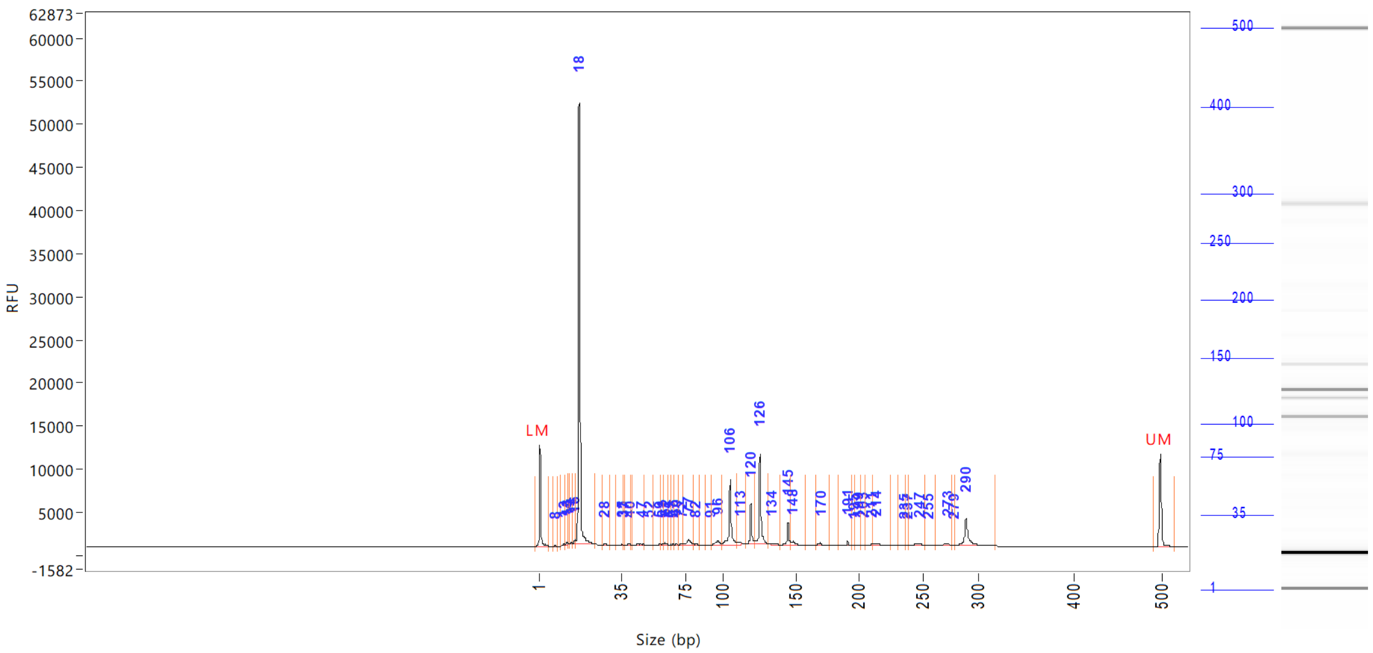


| Peak | Size (bp) | Concentration (ng/uL) | From (bp) | To (bp) | Average size (bp) | CV% | RFU | Corrected peak area |
|---|---|---|---|---|---|---|---|---|
| 26 | 106 | 6.7538 | 99 | 110 | 105 | 1.55 | 7560 | |
| 27 | 113 | 0.5265 | 110 | 116 | 112 | 1.14 | 281 | |
| 28 | 120 | 3.3841 | 116 | 122 | 120 | 0.76 | 4710 | |
| 29 | 126 | 8.0484 | 122 | 132 | 126 | 1.00 | 10458 | |
| 30 | 134 | 0.3497 | 132 | 140 | 134 | 1.32 | 166 | |
| 31 | 145 | 1.9137 | 140 | 147 | 145 | 0.72 | 2636 | |
| 32 | 148 | 0.6069 | 147 | 158 | 149 | 1.40 | 423 | |
| 33 | 170 | 0.3728 | 166 | 177 | 169 | 0.84 | 329 | |
| 34 | 191 | 0.4002 | 184 | 194 | 191 | 0.69 | 510 | |
| 35 | 195 | 0.0673 | 194 | 196 | 195 | 0.35 | 74 | |
| 36 | 199 | 0.1642 | 196 | 202 | 199 | 0.73 | 134 | |
| 37 | 203 | 0.0845 | 202 | 205 | 203 | 0.34 | 109 | |
| 38 | 211 | 0.1766 | 205 | 211 | 210 | 0.55 | 165 | |
| 39 | 214 | 0.5010 | 211 | 225 | 215 | 1.02 | 293 | |
| 40 | 235 | 0.1372 | 230 | 236 | 233 | 0.72 | 81 | |
| 41 | 237 | 0.0597 | 236 | 239 | 238 | 0.36 | 60 | |
| 42 | 247 | 0.5389 | 239 | 253 | 246 | 1.24 | 197 | |
| 43 | 255 | 0.1059 | 253 | 262 | 256 | 0.81 | 59 | |
| 44 | 273 | 0.5505 | 262 | 277 | 271 | 0.98 | 291 | |
| 45 | 279 | 0.0756 | 277 | 280 | 278 | 0.35 | 85 | |
| 46 | 290 | 4.1532 | 280 | 318 | 291 | 1.63 | 3148 | |
| 47 | 500 (UM) | 0.5000 | 491 | 517 | 500 | 0.35 | 10746 | |

| | | | |
|---|---|---|---|
| | TIC: | 87.1113 | ng/uL |
| | TIM: | 5299.2075 | nmole/L |

```
Sample peak width (sec): 5    Sample min peak height: 50    Sample baseline V to V?: Y    Sample baseline V to V points: 3
Sample filter: Binomial       Number of points for filter: 3  Sample start region (min): 0  Sample end region (min): 80
Marker peak width (sec): 5    Marker min peak height: 500   Marker baseline V to V?: Y    Marker baseline V to V points: 3
Lower marker selection: First peak > 500 RFU                Upper marker selection: Last peak > 500 RFU
Ladder size (bp)1, 35, 75, 100, 150, 200, 250, 300, 400, 500
Quantification using: Upper Marker          Final concentration (ng/uL): 0.5000          Dilution factor: 12.0
```

**Sample:** SampD3
**Well location:** D3
**Created:** Monday, 11 October 2021 14:41:52

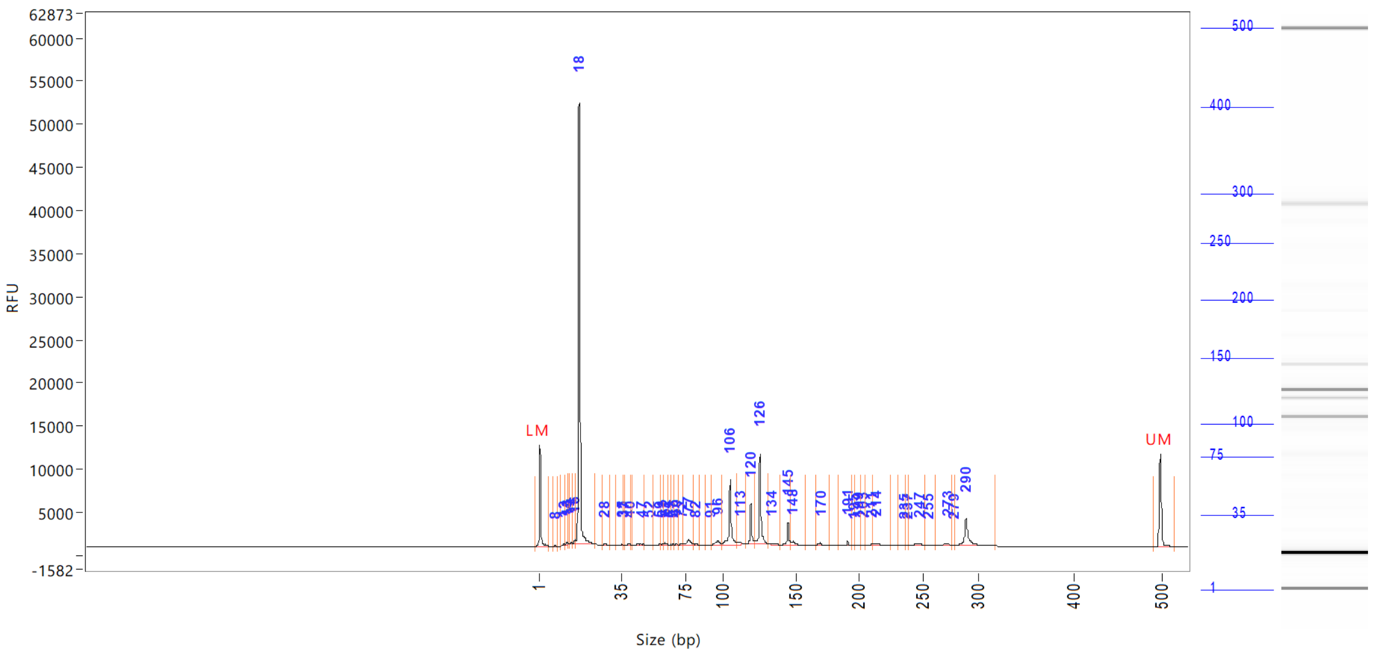


| Peak | Size (bp) | Concentration (ng/uL) | From (bp) | To (bp) | Average size (bp) | CV% | RFU | Corrected peak area |
|---|---|---|---|---|---|---|---|---|

Total concentration: 87.2511 ng/uL

```
Sample peak width (sec): 5    Sample min peak height: 50    Sample baseline V to V?: Y    Sample baseline V to V points: 3
Sample filter: Binomial       Number of points for filter: 3  Sample start region (min): 0  Sample end region (min): 80
Marker peak width (sec): 5    Marker min peak height: 500   Marker baseline V to V?: Y    Marker baseline V to V points: 3
Lower marker selection: First peak > 500 RFU                Upper marker selection: Last peak > 500 RFU
Ladder size (bp)1, 35, 75, 100, 150, 200, 250, 300, 400, 500
Quantification using: Upper Marker          Final concentration (ng/uL): 0.5000          Dilution factor: 12.0
```

**Sample:** SampE3
**Well location:** E3
**Created:** Monday, 11 October 2021 14:41:52

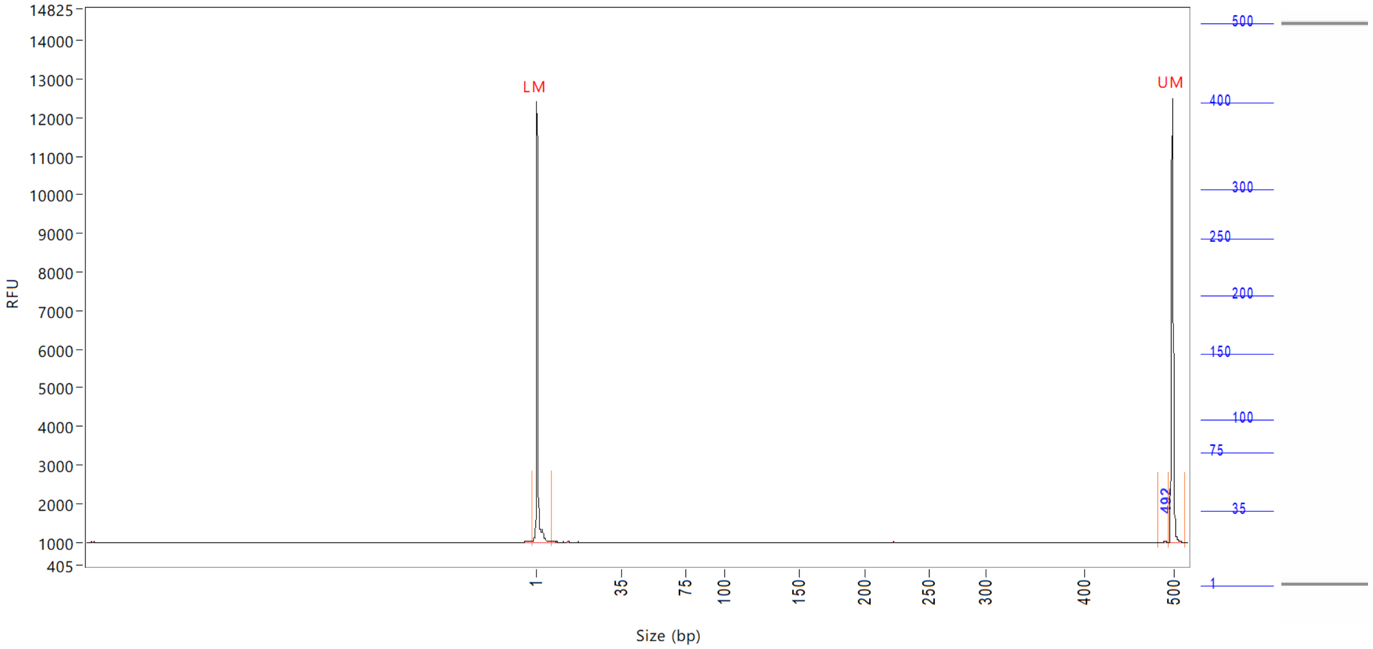


| Peak | Size (bp) | Concentration (ng/uL) | From (bp) | To (bp) | Average size (bp) | CV% | RFU | Corrected peak area |
|---|---|---|---|---|---|---|---|---|
| 1 | 1 (LM) | 0.8589 | 0 | 8 | 1 | 72.29 | 11402 | 43.290 |
| 2 | 492 | 0.0330 | 483 | 496 | 492 | 0.39 | 50 | 0.139 |
| 3 | 500 (UM) | 0.5000 | 496 | 515 | 500 | 0.26 | 11525 | 25.201 |

| | | |
|---|---|---|
| TIC: | 0.0330 | ng/uL |
| TIM: | 0.1104 | nmole/L |
| Total concentration: | 0.4723 | ng/uL |

Sample peak width (sec): 5 Sample min peak height: 50 Sample baseline V to V?: Y Sample baseline V to V points: 3
Sample filter: Binomial Number of points for filter: 3 Sample start region (min): 0 Sample end region (min): 80
Marker peak width (sec): 5 Marker min peak height: 500 Marker baseline V to V?: Y Marker baseline V to V points: 3
Lower marker selection: First peak > 500 RFU Upper marker selection: Last peak > 500 RFU
Ladder size (bp)1, 35, 75, 100, 150, 200, 250, 300, 400, 500
Quantification using: Upper Marker Final concentration (ng/uL): 0.5000 Dilution factor: 12.0

**Sample:** SampF3
**Well location:** F3
**Created:** Monday, 11 October 2021 14:41:52

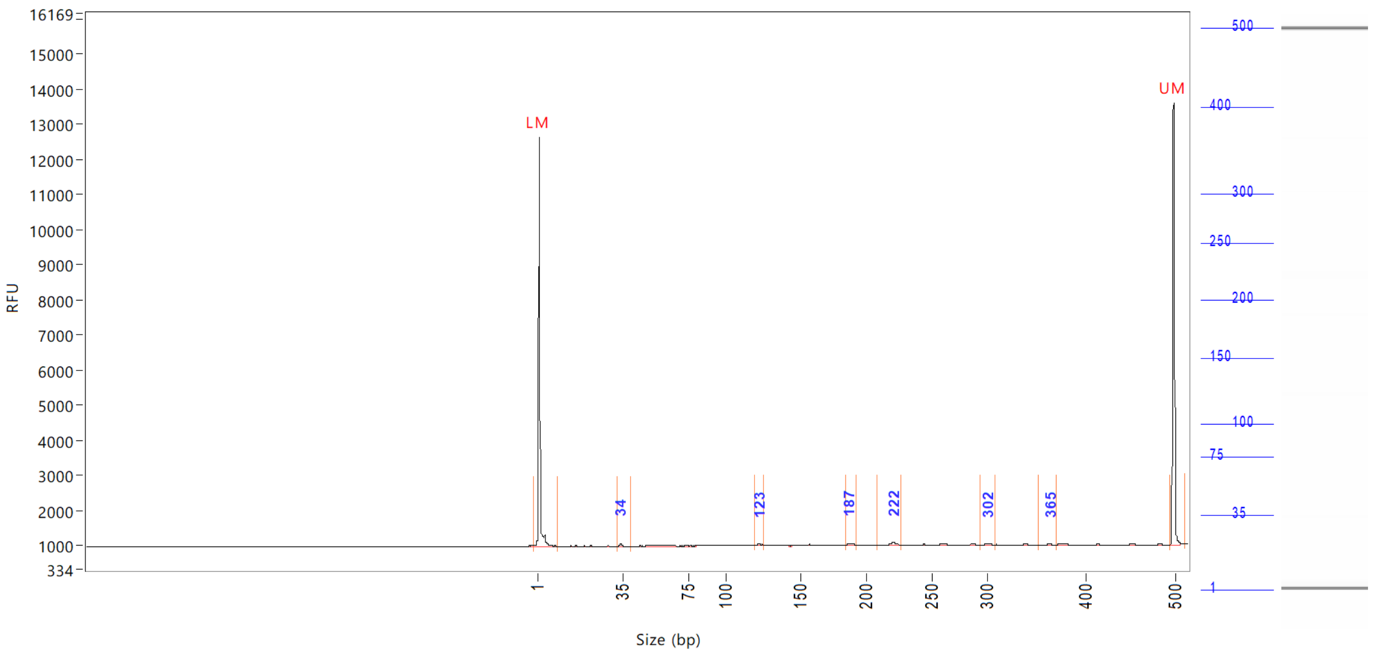


| Peak | Size (bp) | Concentration (ng/uL) | From (bp) | To (bp) | Average size (bp) | CV% | RFU | Corrected peak area |
|---|---|---|---|---|---|---|---|---|
| 1 | 1 (LM) | 0.8290 | 0 | 9 | 1 | 72.10 | 11657 | 43.588 |
| 2 | 34 | 0.1031 | 33 | 40 | 34 | 4.13 | 93 | 0.452 |
| 3 | 123 | 0.0985 | 120 | 125 | 123 | 1.09 | 60 | 0.431 |
| 4 | 187 | 0.1143 | 185 | 193 | 188 | 1.00 | 70 | 0.501 |
| 5 | 222 | 0.2032 | 210 | 228 | 221 | 1.30 | 94 | 0.890 |
| 6 | 302 | 0.1045 | 295 | 309 | 302 | 1.15 | 54 | 0.458 |
| 7 | 365 | 0.0899 | 354 | 371 | 364 | 1.00 | 58 | 0.394 |
| 8 | 500 (UM) | 0.5000 | 495 | 513 | 500 | 0.25 | 12610 | 26.290 |

| | | |
|---|---|---|
| TIC: | 0.7134 | ng/uL |
| TIM: | 9.7563 | nmole/L |
| Total concentration: | 2.2997 | ng/uL |

Sample peak width (sec): 5    Sample min peak height: 50    Sample baseline V to V?: Y    Sample baseline V to V points: 3
Sample filter: Binomial    Number of points for filter: 3  Sample start region (min): 0  Sample end region (min): 80
Marker peak width (sec): 5    Marker min peak height: 500    Marker baseline V to V?: Y    Marker baseline V to V points: 3
Lower marker selection: First peak > 500 RFU    Upper marker selection: Last peak > 500 RFU
Ladder size (bp)1, 35, 75, 100, 150, 200, 250, 300, 400, 500
Quantification using: Upper Marker    Final concentration (ng/uL): 0.5000    Dilution factor: 12.0

**Sample:** SampG3
**Well location:** G3
**Created:** Monday, 11 October 2021 14:41:52

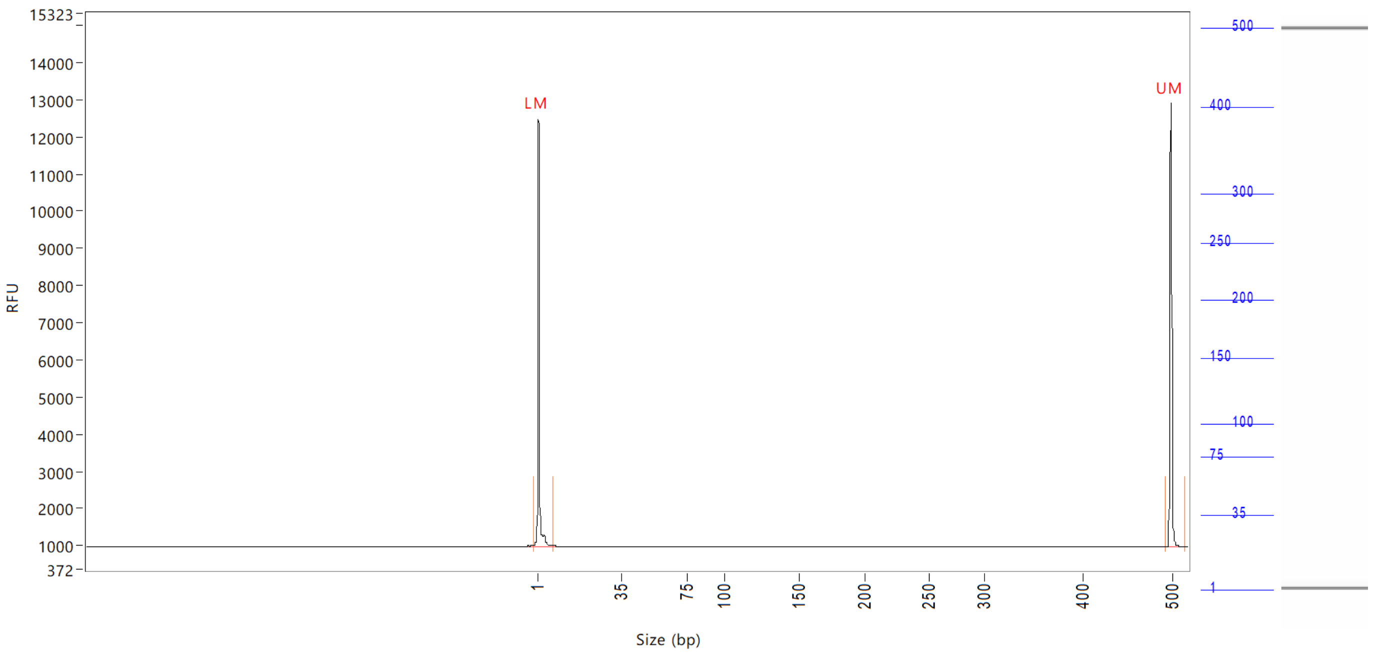


| Peak | Size (bp) | Concentration (ng/uL) | From (bp) | To (bp) | Average size (bp) | CV% | RFU | Corrected peak area |
|---|---|---|---|---|---|---|---|---|
| 1 | 1 (LM) | 0.8741 | 0 | 8 | 2 | 66.00 | 11496 | 44.151 |
| 2 | 500 (UM) | 0.5000 | 493 | 517 | 500 | 0.26 | 11955 | 25.254 |

| | | |
|---|---|---|
| TIC: | 0.0000 | ng/uL |
| TIM: | 0.0000 | nmole/L |
| Total concentration: | 0.4465 | ng/uL |

Sample peak width (sec): 5    Sample min peak height: 50    Sample baseline V to V?: Y    Sample baseline V to V points: 3
Sample filter: Binomial    Number of points for filter: 3  Sample start region (min): 0  Sample end region (min): 80
Marker peak width (sec): 5    Marker min peak height: 500   Marker baseline V to V?: Y    Marker baseline V to V points: 3
Lower marker selection: First peak > 500 RFU    Upper marker selection: Last peak > 500 RFU
Ladder size (bp)1, 35, 75, 100, 150, 200, 250, 300, 400, 500
Quantification using: Upper Marker    Final concentration (ng/uL): 0.5000    Dilution factor: 12.0

**Sample:** SampH3
**Well location:** H3
**Created:** Monday, 11 October 2021 14:41:52

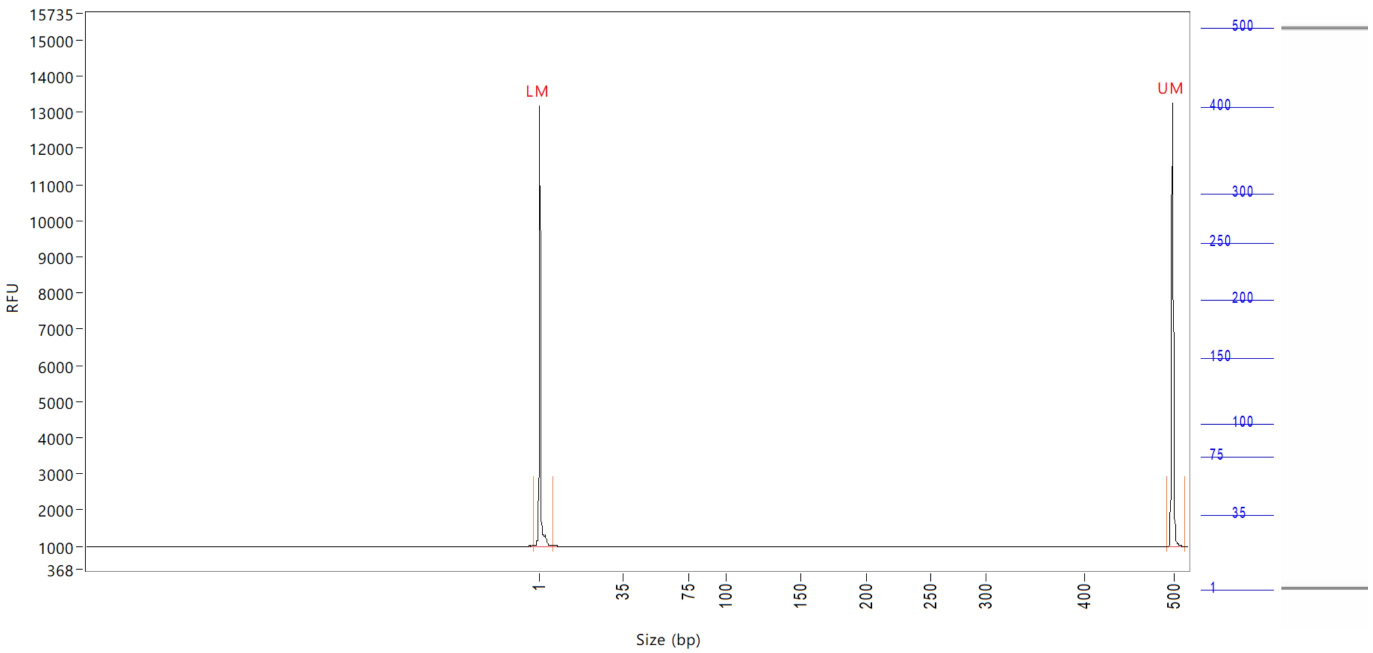


| Peak | Size (bp) | Concentration (ng/uL) | From (bp) | To (bp) | Average size (bp) | CV% | RFU | Corrected peak area |
|---|---|---|---|---|---|---|---|---|
| 1 | 1 (LM) | 0.8784 | 0 | 7 | 1 | 74.56 | 12192 | 46.367 |
| 2 | 500 (UM) | 0.5000 | 494 | 515 | 500 | 0.26 | 12283 | 26.394 |

| | | |
|---|---|---|
| TIC: | 0.0000 | ng/uL |
| TIM: | 0.0000 | nmole/L |
| Total concentration: | 0.4181 | ng/uL |

```
Sample peak width (sec): 5    Sample min peak height: 50    Sample baseline V to V?: Y    Sample baseline V to V points: 3
Sample filter: Binomial       Number of points for filter: 3  Sample start region (min): 0  Sample end region (min): 80
Marker peak width (sec): 5    Marker min peak height: 500   Marker baseline V to V?: Y    Marker baseline V to V points: 3
Lower marker selection: First peak > 500 RFU                Upper marker selection: Last peak > 500 RFU
Ladder size (bp)1, 35, 75, 100, 150, 200, 250, 300, 400, 500
Quantification using: Upper Marker          Final concentration (ng/uL): 0.5000          Dilution factor: 12.0
```

**Sample:** SampA4
**Well location:** A4
**Created:** Monday, 11 October 2021 14:41:52

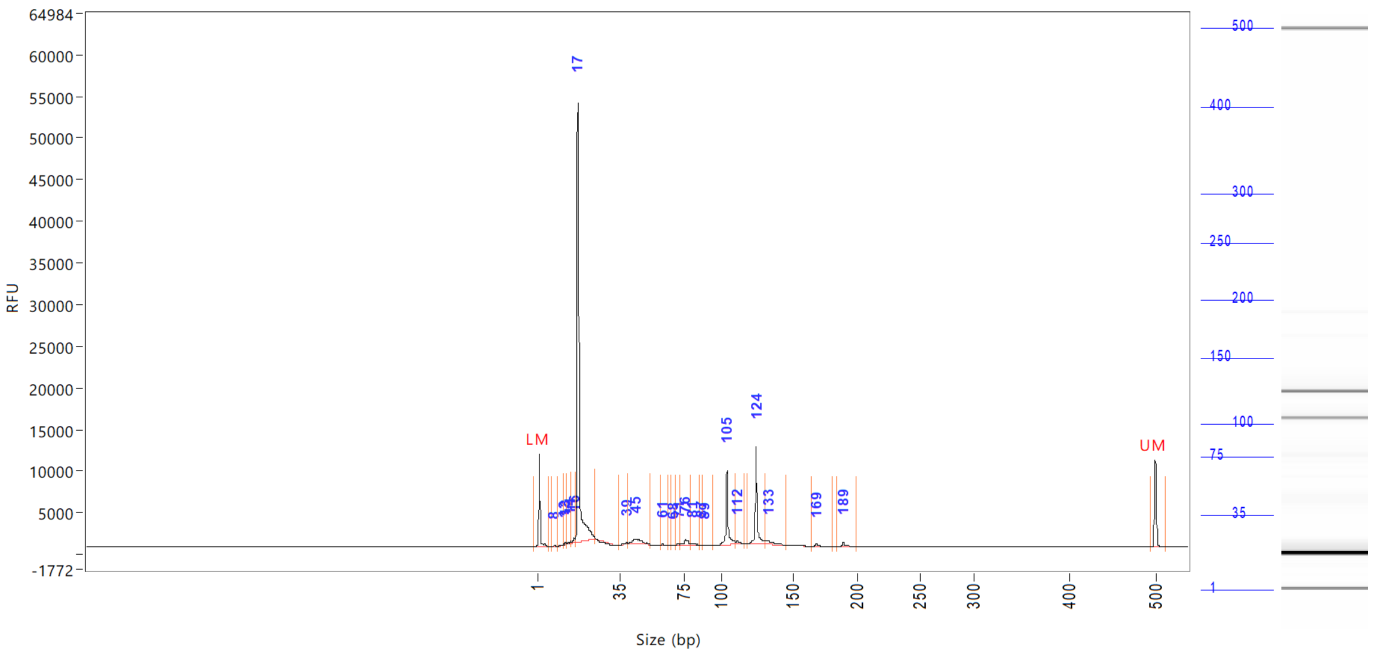


| Peak | Size (bp) | Concentration (ng/uL) | From (bp) | To (bp) | Average size (bp) | CV% | RFU | Corrected peak area |
|---|---|---|---|---|---|---|---|---|
| 1 | 1 (LM) | 0.8877 | 0 | 6 | 1 | 64.89 | 11101 | 39.895 |
| 2 | 8 | 0.1169 | 7 | 9 | 8 | 4.60 | 90 | 0.438 |
| 3 | 12 | 0.1539 | 9 | 12 | 11 | 5.89 | 164 | 0.576 |
| 4 | 13 | 0.1873 | 12 | 13 | 13 | 1.50 | 259 | 0.701 |
| 5 | 14 | 0.2628 | 13 | 15 | 14 | 2.75 | 239 | 0.984 |
| 6 | 16 | 0.6766 | 15 | 16 | 16 | 2.32 | 510 | 2.534 |
| 7 | 17 | 70.1290 | 16 | 25 | 18 | 7.17 | 52827 | 262.631 |
| 8 | 39 | 0.3603 | 34 | 40 | 38 | 3.30 | 168 | 1.349 |
| 9 | 45 | 2.9022 | 40 | 54 | 46 | 5.71 | 531 | 10.869 |
| 10 | 61 | 0.2003 | 60 | 65 | 61 | 1.15 | 245 | 0.750 |
| 11 | 68 | 0.0904 | 66 | 69 | 68 | 0.61 | 115 | 0.339 |
| 12 | 71 | 0.2867 | 69 | 72 | 71 | 0.87 | 256 | 1.074 |
| 13 | 76 | 1.8280 | 72 | 80 | 76 | 2.59 | 690 | 6.846 |
| 14 | 81 | 0.4616 | 80 | 86 | 82 | 1.74 | 209 | 1.729 |
| 15 | 87 | 0.0670 | 86 | 88 | 87 | 0.72 | 60 | 0.251 |
| 16 | 89 | 0.0868 | 88 | 95 | 90 | 1.86 | 59 | 0.325 |
| 17 | 105 | 8.7969 | 95 | 110 | 105 | 1.58 | 8961 | 32.944 |
| 18 | 112 | 0.6953 | 110 | 117 | 112 | 1.35 | 331 | 2.604 |
| 19 | 124 | 10.7138 | 119 | 131 | 125 | 1.36 | 11798 | 40.123 |
| 20 | 133 | 1.3752 | 131 | 145 | 134 | 2.06 | 412 | 5.150 |
| 21 | 169 | 0.3373 | 165 | 182 | 170 | 1.48 | 326 | 1.263 |
| 22 | 189 | 0.5320 | 185 | 201 | 190 | 1.23 | 554 | 1.992 |
| 23 | 500 (UM) | 0.5000 | 494 | 511 | 500 | 0.25 | 10480 | 22.470 |
| | TIC: | 100.2604 | ng/uL | | | | | |

Sample peak width (sec): 5 Sample min peak height: 50 Sample baseline V to V?: Y Sample baseline V to V points: 3
Sample filter: Binomial Number of points for filter: 3 Sample start region (min): 0 Sample end region (min): 80
Marker peak width (sec): 5 Marker min peak height: 500 Marker baseline V to V?: Y Marker baseline V to V points: 3
Lower marker selection: First peak > 500 RFU Upper marker selection: Last peak > 500 RFU
Ladder size (bp)1, 35, 75, 100, 150, 200, 250, 300, 400, 500
Quantification using: Upper Marker Final concentration (ng/uL): 0.5000 Dilution factor: 12.0

**Sample:** SampA4
**Well location:** A4
**Created:** Monday, 11 October 2021 14:41:52

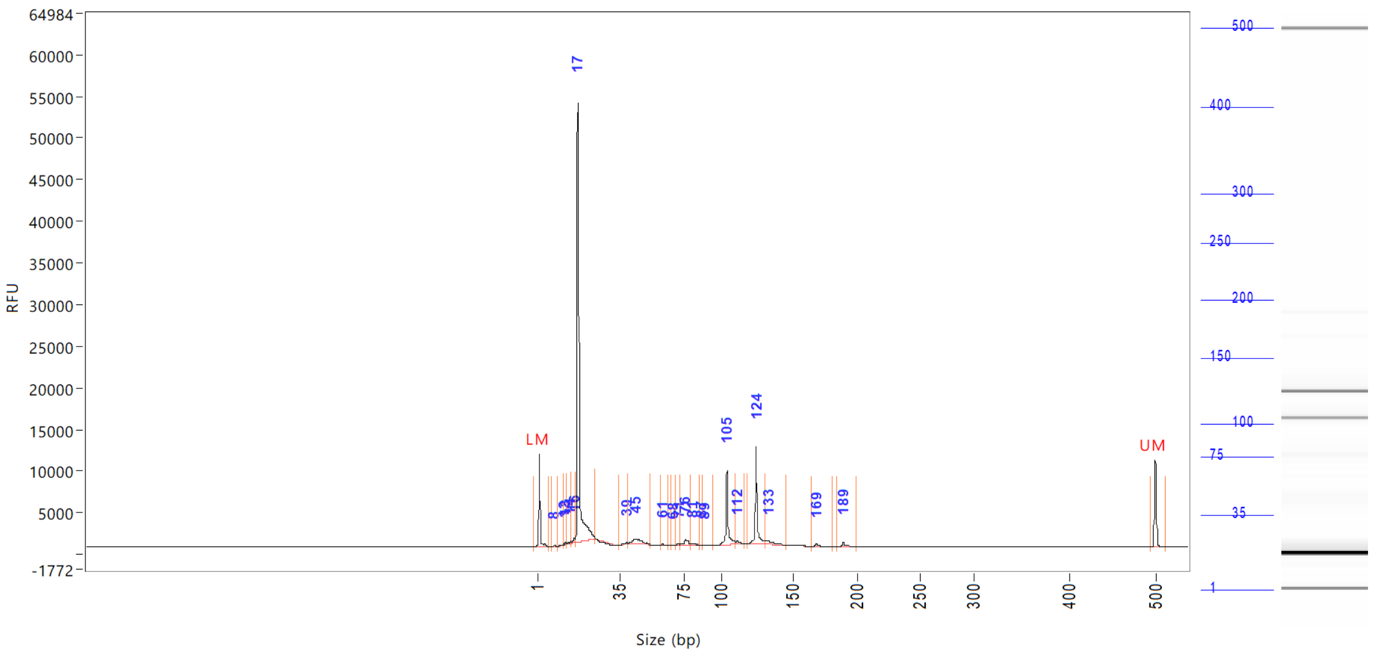

| Peak | Size | Concentration | From | To | Average size | CV% | RFU | Corrected peak area |
|---|---|---|---|---|---|---|---|---|
| | (bp) | (ng/uL) | (bp) | (bp) | (bp) | | | |

| | | |
|---|---|---|
| TIM: | 6988.7700 | nmole/L |
| Total concentration: | 100.6372 | ng/uL |

Sample peak width (sec): 5 Sample min peak height: 50 Sample baseline V to V?: Y Sample baseline V to V points: 3
Sample filter: Binomial Number of points for filter: 3 Sample start region (min): 0 Sample end region (min): 80
Marker peak width (sec): 5 Marker min peak height: 500 Marker baseline V to V?: Y Marker baseline V to V points: 3
Lower marker selection: First peak > 500 RFU Upper marker selection: Last peak > 500 RFU
Ladder size (bp)1, 35, 75, 100, 150, 200, 250, 300, 400, 500
Quantification using: Upper Marker Final concentration (ng/uL): 0.5000 Dilution factor: 12.0

**Sample:** SampB4
**Well location:** B4
**Created:** Monday, 11 October 2021 14:41:52

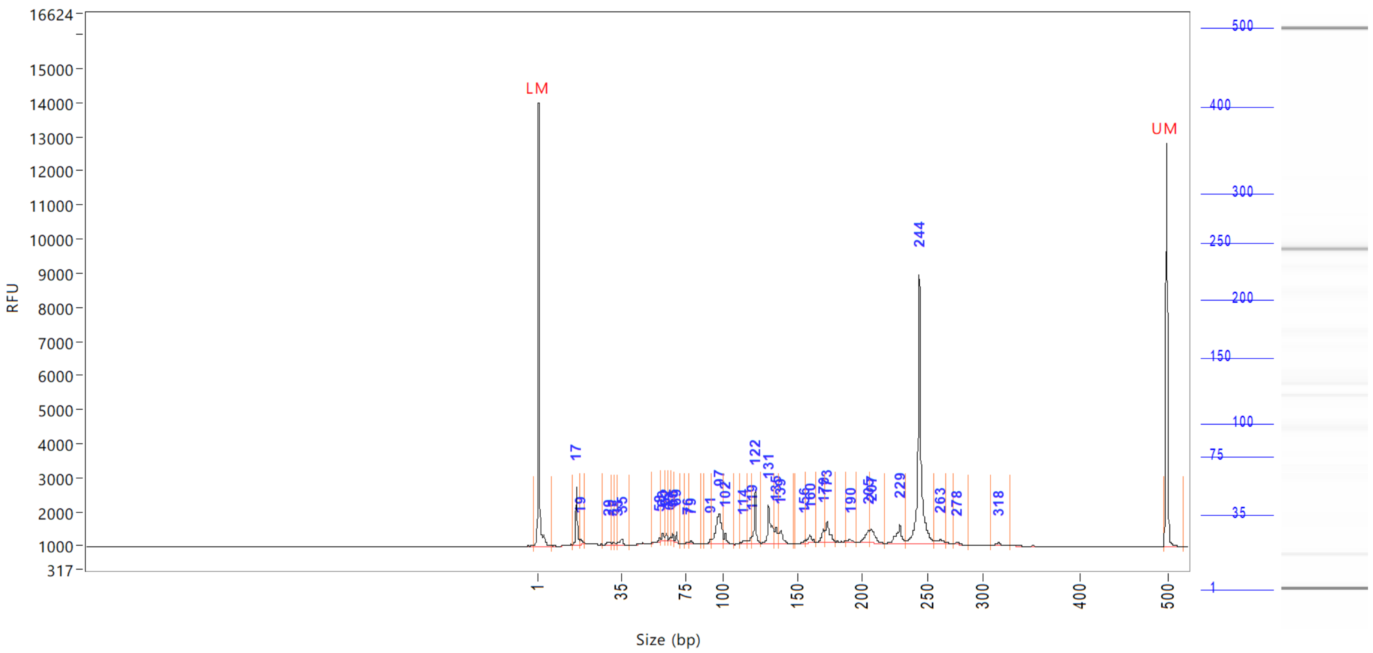


| Peak | Size (bp) | Concentration (ng/uL) | From (bp) | To (bp) | Average size (bp) | CV% | RFU | Corrected peak area |
|---|---|---|---|---|---|---|---|---|
| 1 | 1 (LM) | 0.9859 | 0 | 7 | 1 | 72.59 | 13028 | 48.721 |
| 2 | 17 | 1.6302 | 15 | 18 | 17 | 2.61 | 1692 | 6.713 |
| 3 | 19 | 0.2420 | 18 | 20 | 19 | 3.04 | 151 | 0.997 |
| 4 | 29 | 0.1678 | 28 | 31 | 30 | 2.39 | 81 | 0.691 |
| 5 | 32 | 0.1084 | 31 | 32 | 32 | 1.13 | 91 | 0.446 |
| 6 | 33 | 0.0692 | 32 | 33 | 33 | 0.87 | 70 | 0.285 |
| 7 | 35 | 0.3135 | 33 | 40 | 35 | 3.48 | 170 | 1.291 |
| 8 | 58 | 0.1300 | 54 | 59 | 58 | 1.49 | 123 | 0.535 |
| 9 | 60 | 0.2368 | 59 | 62 | 60 | 0.98 | 228 | 0.975 |
| 10 | 62 | 0.1731 | 62 | 64 | 62 | 0.64 | 231 | 0.713 |
| 11 | 65 | 0.1020 | 64 | 66 | 65 | 0.81 | 122 | 0.420 |
| 12 | 66 | 0.1553 | 66 | 67 | 66 | 0.60 | 215 | 0.640 |
| 13 | 69 | 0.2100 | 67 | 71 | 69 | 0.63 | 313 | 0.865 |
| 14 | 76 | 0.0932 | 73 | 77 | 76 | 1.51 | 69 | 0.384 |
| 15 | 79 | 0.1312 | 77 | 85 | 79 | 1.85 | 79 | 0.540 |
| 16 | 91 | 0.1958 | 87 | 92 | 91 | 1.41 | 148 | 0.806 |
| 17 | 97 | 2.2796 | 92 | 100 | 96 | 1.79 | 907 | 9.387 |
| 18 | 102 | 0.3303 | 100 | 108 | 102 | 1.26 | 334 | 1.360 |
| 19 | 114 | 0.1958 | 112 | 116 | 114 | 0.99 | 106 | 0.806 |
| 20 | 119 | 0.2706 | 116 | 120 | 118 | 0.88 | 213 | 1.114 |
| 21 | 122 | 1.1759 | 120 | 126 | 122 | 0.64 | 1514 | 4.842 |
| 22 | 131 | 1.6555 | 126 | 134 | 131 | 1.20 | 1125 | 6.817 |
| 23 | 135 | 0.6455 | 134 | 138 | 136 | 0.72 | 480 | 2.658 |
| 24 | 139 | 0.4929 | 138 | 147 | 139 | 0.91 | 378 | 2.030 |
| 25 | 156 | 0.1187 | 149 | 157 | 156 | 1.03 | 95 | 0.489 |

```
Sample peak width (sec): 5     Sample min peak height: 50     Sample baseline V to V?: Y     Sample baseline V to V points: 3
Sample filter: Binomial        Number of points for filter: 3  Sample start region (min): 0  Sample end region (min): 80
Marker peak width (sec): 5     Marker min peak height: 500    Marker baseline V to V?: Y     Marker baseline V to V points: 3
Lower marker selection: First peak > 500 RFU                  Upper marker selection: Last peak > 500 RFU
Ladder size (bp)1, 35, 75, 100, 150, 200, 250, 300, 400, 500
Quantification using: Upper Marker         Final concentration (ng/uL): 0.5000         Dilution factor: 12.0
```

**Sample:** SampB4
**Well location:** B4
**Created:** Monday, 11 October 2021 14:41:52

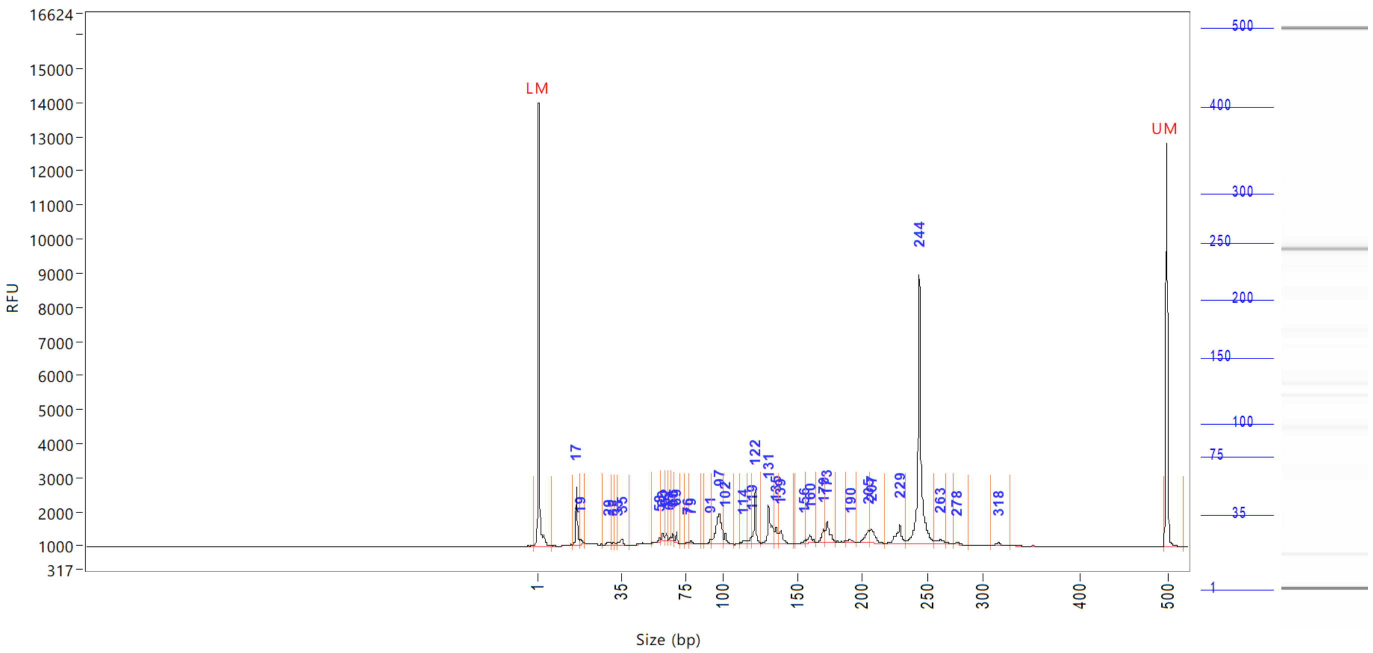


| Peak | Size (bp) | Concentration (ng/uL) | From (bp) | To (bp) | Average size (bp) | CV% | RFU | Corrected peak area |
|---|---|---|---|---|---|---|---|---|
| 26 | 160 | 0.3339 | 157 | 165 | 160 | 1.01 | 223 | |
| 27 | 170 | 0.4525 | 165 | 171 | 170 | 0.74 | 382 | |
| 28 | 173 | 0.9586 | 171 | 180 | 174 | 1.04 | 605 | |
| 29 | 190 | 0.1811 | 187 | 196 | 191 | 0.96 | 96 | |
| 30 | 205 | 0.6067 | 196 | 206 | 204 | 1.01 | 366 | |
| 31 | 207 | 0.7463 | 206 | 218 | 209 | 0.94 | 434 | |
| 32 | 229 | 0.9659 | 218 | 234 | 228 | 1.18 | 537 | |
| 33 | 244 | 8.2850 | 234 | 258 | 245 | 1.24 | 7872 | |
| 34 | 263 | 0.2466 | 258 | 268 | 262 | 0.99 | 112 | |
| 35 | 278 | 0.0983 | 274 | 288 | 278 | 0.81 | 64 | |
| 36 | 318 | 0.1430 | 310 | 330 | 318 | 1.08 | 89 | |
| 37 | 500 (UM) | 0.5000 | 496 | 0 | 500 | 0.29 | 11841 | |

| | | |
|---|---|---|
| TIC: | 24.1411 | ng/uL |
| TIM: | 440.3306 | nmole/L |
| Total concentration: | 24.5015 | ng/uL |

Sample peak width (sec): 5 Sample min peak height: 50 Sample baseline V to V?: Y Sample baseline V to V points: 3
Sample filter: Binomial Number of points for filter: 3 Sample start region (min): 0 Sample end region (min): 80
Marker peak width (sec): 5 Marker min peak height: 500 Marker baseline V to V?: Y Marker baseline V to V points: 3
Lower marker selection: First peak > 500 RFU Upper marker selection: Last peak > 500 RFU
Ladder size (bp)1, 35, 75, 100, 150, 200, 250, 300, 400, 500
Quantification using: Upper Marker Final concentration (ng/uL): 0.5000 Dilution factor: 12.0

**Sample:** SampC4
**Well location:** C4
**Created:** Monday, 11 October 2021 14:41:52

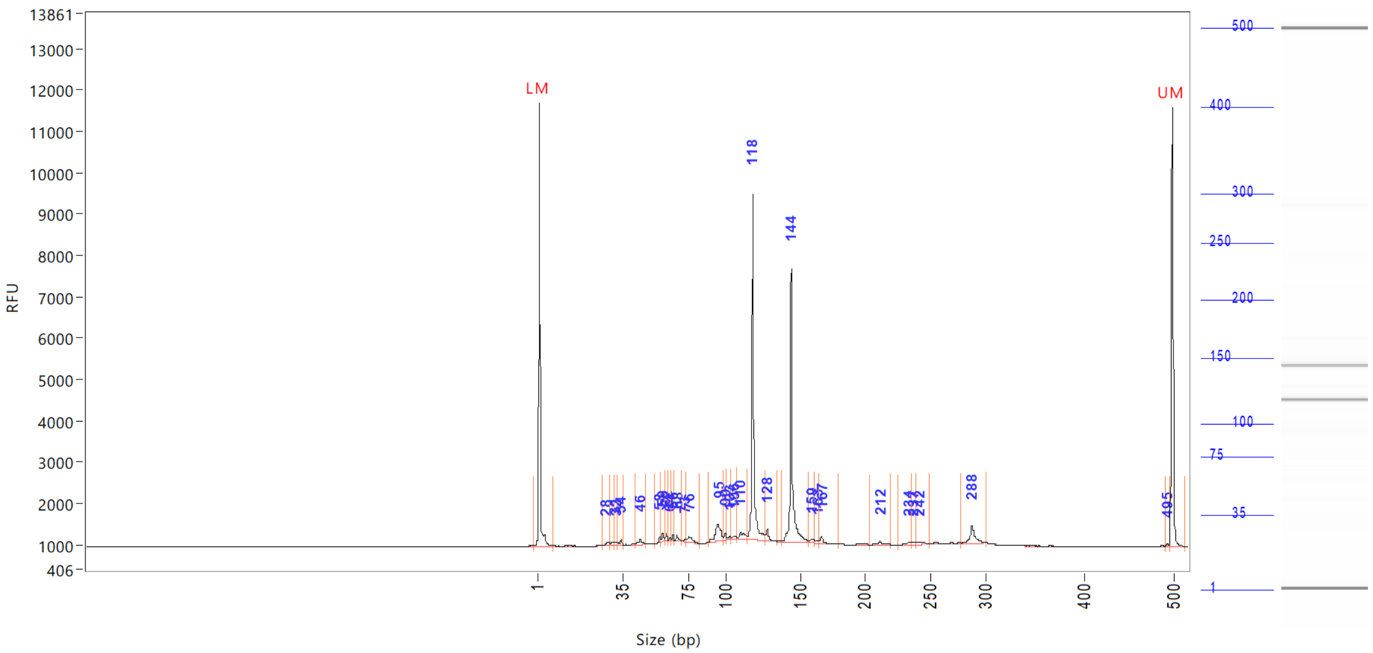

| Peak | Size (bp) | Concentration (ng/uL) | From (bp) | To (bp) | Average size (bp) | CV% | RFU | Corrected peak area |
|---|---|---|---|---|---|---|---|---|
| 1 | 1 (LM) | 0.8808 | 0 | 8 | 1 | 69.75 | 10750 | 40.016 |
| 2 | 28 | 0.1405 | 27 | 30 | 29 | 1.91 | 74 | 0.532 |
| 3 | 31 | 0.1840 | 30 | 32 | 31 | 2.10 | 83 | 0.697 |
| 4 | 32 | 0.0765 | 32 | 33 | 32 | 0.91 | 65 | 0.290 |
| 5 | 34 | 0.1683 | 33 | 36 | 34 | 1.46 | 121 | 0.637 |
| 6 | 46 | 0.2644 | 43 | 50 | 46 | 2.65 | 142 | 1.001 |
| 7 | 58 | 0.1338 | 55 | 58 | 58 | 0.89 | 132 | 0.507 |
| 8 | 59 | 0.1989 | 58 | 61 | 59 | 0.95 | 186 | 0.753 |
| 9 | 62 | 0.1362 | 61 | 63 | 61 | 0.64 | 166 | 0.516 |
| 10 | 64 | 0.0540 | 63 | 65 | 64 | 0.71 | 66 | 0.204 |
| 11 | 66 | 0.1016 | 65 | 67 | 65 | 0.56 | 141 | 0.385 |
| 12 | 68 | 0.1576 | 67 | 71 | 68 | 1.25 | 147 | 0.597 |
| 13 | 73 | 0.0897 | 71 | 74 | 73 | 1.10 | 69 | 0.340 |
| 14 | 76 | 0.3517 | 74 | 83 | 77 | 2.42 | 138 | 1.331 |
| 15 | 95 | 1.1180 | 88 | 98 | 94 | 2.04 | 414 | 4.233 |
| 16 | 99 | 0.1651 | 98 | 101 | 99 | 0.82 | 179 | 0.625 |
| 17 | 103 | 0.1156 | 101 | 104 | 103 | 0.77 | 81 | 0.437 |
| 18 | 106 | 0.1172 | 104 | 108 | 106 | 0.81 | 80 | 0.444 |
| 19 | 110 | 0.3737 | 108 | 115 | 112 | 1.49 | 154 | 1.415 |
| 20 | 118 | 7.3534 | 115 | 127 | 119 | 1.52 | 8365 | 27.840 |
| 21 | 128 | 0.3119 | 127 | 134 | 128 | 0.90 | 285 | 1.181 |
| 22 | 144 | 6.2125 | 138 | 157 | 145 | 1.73 | 6602 | 23.520 |
| 23 | 159 | 0.1599 | 157 | 162 | 160 | 0.85 | 78 | 0.605 |
| 24 | 163 | 0.0859 | 162 | 165 | 164 | 0.53 | 65 | 0.325 |
| 25 | 167 | 0.2087 | 165 | 179 | 168 | 1.07 | 188 | 0.790 |

Sample peak width (sec): 5    Sample min peak height: 50    Sample baseline V to V?: Y    Sample baseline V to V points: 3
Sample filter: Binomial    Number of points for filter: 3    Sample start region (min): 0    Sample end region (min): 80
Marker peak width (sec): 5    Marker min peak height: 500    Marker baseline V to V?: Y    Marker baseline V to V points: 3
Lower marker selection: First peak > 500 RFU    Upper marker selection: Last peak > 500 RFU
Ladder size (bp)1, 35, 75, 100, 150, 200, 250, 300, 400, 500
Quantification using: Upper Marker    Final concentration (ng/uL): 0.5000    Dilution factor: 12.0

**Sample:** SampC4
**Well location:** C4
**Created:** Monday, 11 October 2021 14:41:52

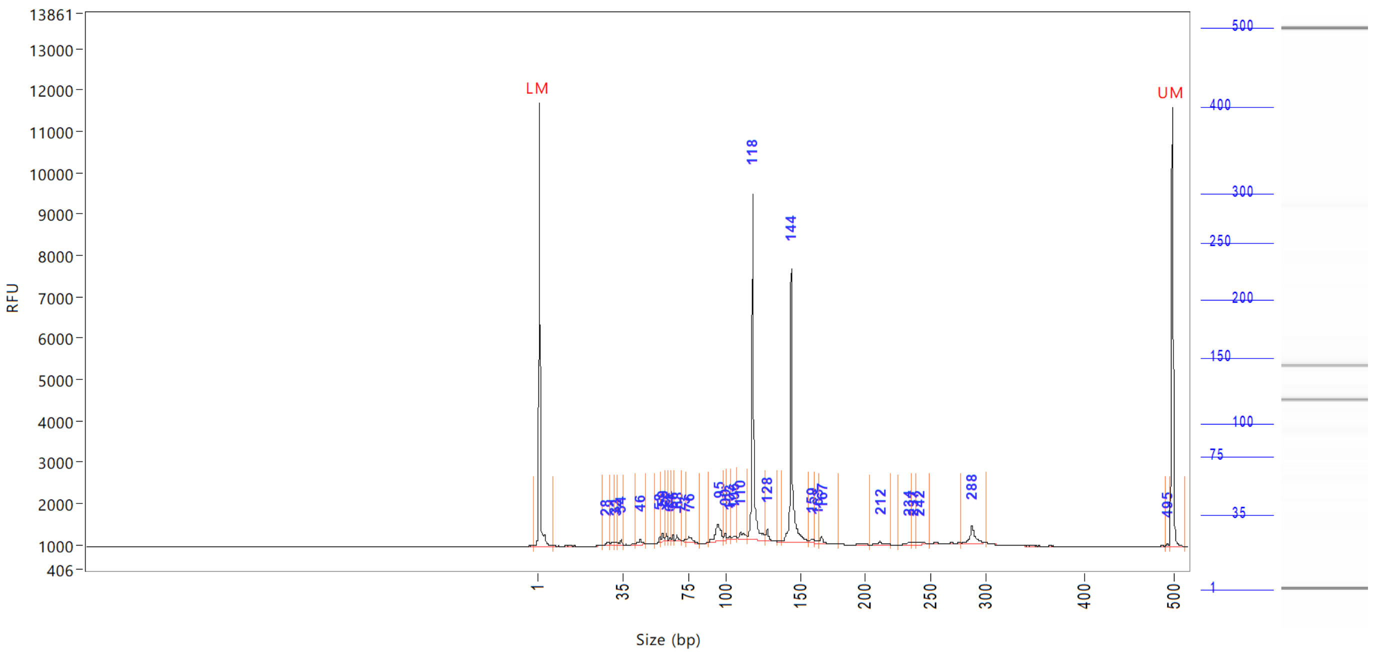


| Peak | Size (bp) | Concentration (ng/uL) | From (bp) | To (bp) | Average size (bp) | CV% | RFU | Corrected peak area |
|---|---|---|---|---|---|---|---|---|
| 26 | 212 | 0.1377 | 204 | 220 | 212 | 1.28 | 77 | |
| 27 | 234 | 0.1080 | 226 | 236 | 233 | 0.81 | 66 | |
| 28 | 237 | 0.0890 | 236 | 239 | 237 | 0.45 | 64 | |
| 29 | 242 | 0.1762 | 239 | 250 | 244 | 1.17 | 57 | |
| 30 | 288 | 0.7578 | 278 | 301 | 288 | 1.19 | 426 | |
| 31 | 495 | 0.0329 | 492 | 496 | 494 | 0.22 | 52 | |
| 32 | 500 (UM) | 0.5000 | 496 | 515 | 500 | 0.26 | 10612 | |

| | | |
|---|---|---|
| TIC: | 19.5808 | ng/uL |
| TIM: | 289.1783 | nmole/L |
| Total concentration: | 20.1704 | ng/uL |

```
Sample peak width (sec): 5    Sample min peak height: 50    Sample baseline V to V?: Y    Sample baseline V to V points: 3
Sample filter: Binomial       Number of points for filter: 3  Sample start region (min): 0  Sample end region (min): 80
Marker peak width (sec): 5    Marker min peak height: 500    Marker baseline V to V?: Y    Marker baseline V to V points: 3
Lower marker selection: First peak > 500 RFU                 Upper marker selection: Last peak > 500 RFU
Ladder size (bp)1, 35, 75, 100, 150, 200, 250, 300, 400, 500
Quantification using: Upper Marker          Final concentration (ng/uL): 0.5000          Dilution factor: 12.0
```

**Sample:** SampD4
**Well location:** D4
**Created:** Monday, 11 October 2021 14:41:52

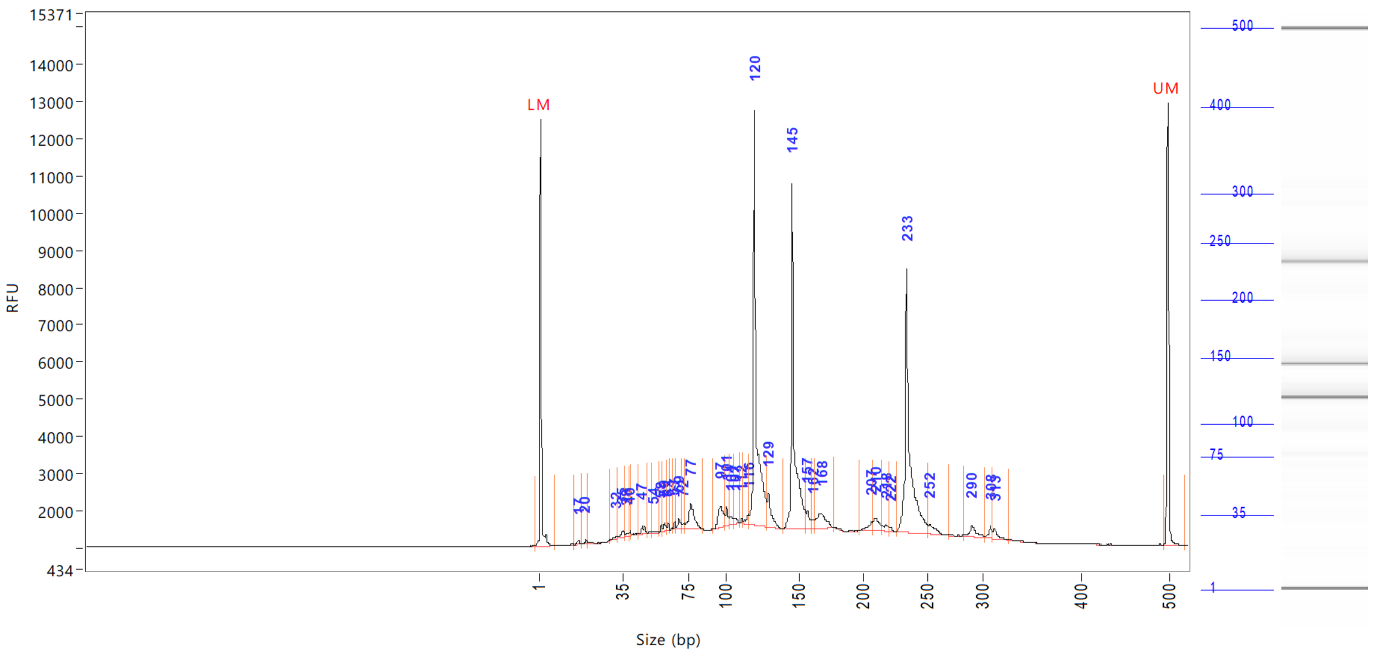


| Peak | Size (bp) | Concentration (ng/uL) | From (bp) | To (bp) | Average size (bp) | CV% | RFU | Corrected peak area |
|---|---|---|---|---|---|---|---|---|
| 1 | 1 (LM) | 0.8689 | 0 | 8 | 1 | 71.19 | 11464 | 42.839 |
| 2 | 17 | 0.1537 | 15 | 19 | 17 | 2.90 | 108 | 0.631 |
| 3 | 20 | 0.1216 | 19 | 21 | 20 | 1.98 | 120 | 0.499 |
| 4 | 32 | 0.1117 | 30 | 32 | 31 | 1.88 | 84 | 0.459 |
| 5 | 35 | 0.3170 | 32 | 37 | 34 | 2.93 | 176 | 1.302 |
| 6 | 38 | 0.1503 | 37 | 39 | 38 | 1.75 | 102 | 0.617 |
| 7 | 40 | 0.0999 | 39 | 41 | 39 | 0.86 | 130 | 0.410 |
| 8 | 47 | 0.3806 | 45 | 51 | 47 | 2.58 | 214 | 1.564 |
| 9 | 54 | 0.0888 | 53 | 57 | 55 | 1.61 | 53 | 0.365 |
| 10 | 58 | 0.1561 | 57 | 59 | 58 | 0.80 | 187 | 0.641 |
| 11 | 60 | 0.2287 | 59 | 62 | 60 | 1.06 | 211 | 0.940 |
| 12 | 63 | 0.1502 | 62 | 64 | 62 | 0.61 | 199 | 0.617 |
| 13 | 67 | 0.1197 | 66 | 68 | 67 | 0.59 | 186 | 0.492 |
| 14 | 69 | 0.3782 | 68 | 71 | 69 | 1.30 | 267 | 1.554 |
| 15 | 72 | 0.1206 | 71 | 73 | 72 | 0.58 | 136 | 0.496 |
| 16 | 77 | 1.7717 | 73 | 85 | 78 | 2.90 | 690 | 7.279 |
| 17 | 97 | 1.3651 | 92 | 99 | 97 | 1.60 | 573 | 5.609 |
| 18 | 101 | 0.5473 | 99 | 103 | 101 | 1.03 | 516 | 2.249 |
| 19 | 104 | 0.3653 | 103 | 106 | 105 | 0.98 | 211 | 1.501 |
| 20 | 107 | 0.1929 | 106 | 110 | 108 | 0.70 | 160 | 0.792 |
| 21 | 112 | 0.1033 | 110 | 113 | 112 | 0.47 | 145 | 0.424 |
| 22 | 116 | 0.4063 | 113 | 117 | 115 | 1.01 | 271 | 1.670 |
| 23 | 120 | 13.4142 | 117 | 128 | 121 | 2.07 | 11123 | 55.114 |
| 24 | 129 | 1.5020 | 128 | 139 | 131 | 1.50 | 908 | 6.171 |
| 25 | 145 | 10.9704 | 139 | 156 | 147 | 2.12 | 9299 | 45.073 |

Sample peak width (sec): 5 Sample min peak height: 50 Sample baseline V to V?: Y Sample baseline V to V points: 3
Sample filter: Binomial Number of points for filter: 3 Sample start region (min): 0 Sample end region (min): 80
Marker peak width (sec): 5 Marker min peak height: 500 Marker baseline V to V?: Y Marker baseline V to V points: 3
Lower marker selection: First peak > 500 RFU Upper marker selection: Last peak > 500 RFU
Ladder size (bp)1, 35, 75, 100, 150, 200, 250, 300, 400, 500
Quantification using: Upper Marker Final concentration (ng/uL): 0.5000 Dilution factor: 12.0

**Sample:** SampD4
**Well location:** D4
**Created:** Monday, 11 October 2021 14:41:52

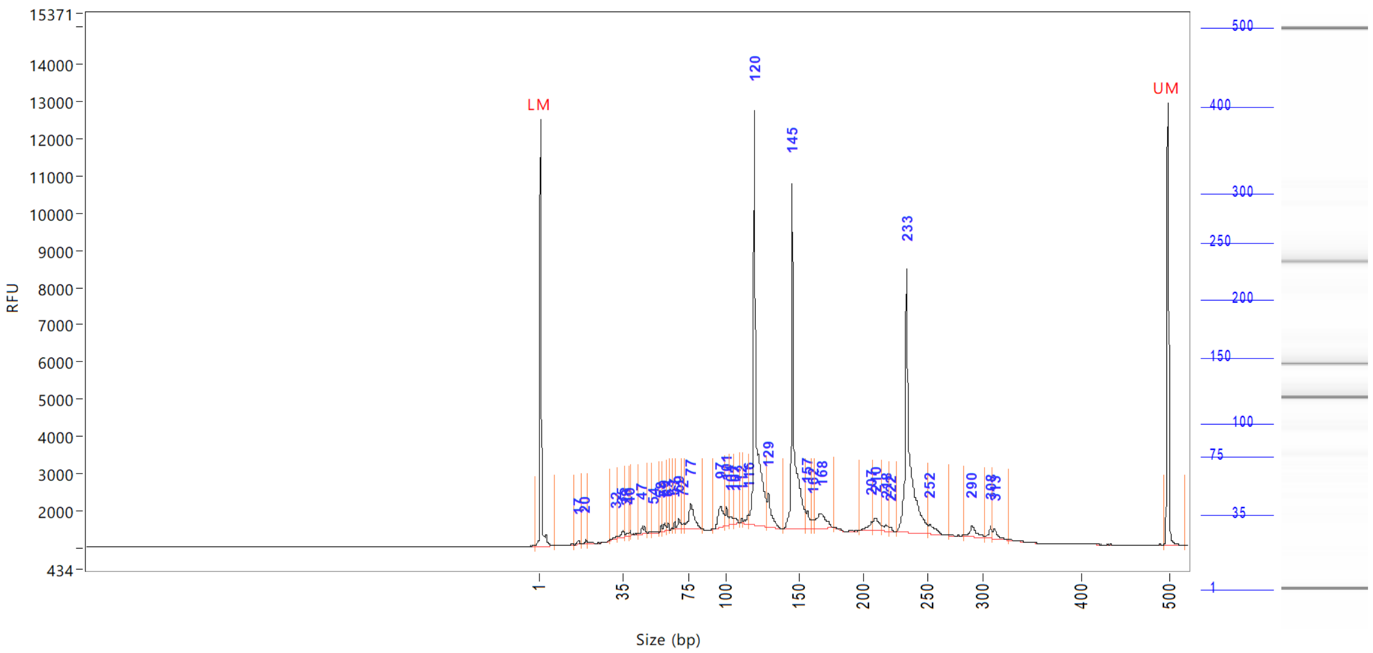


| Peak | Size (bp) | Concentration (ng/uL) | From (bp) | To (bp) | Average size (bp) | CV% | RFU | Corrected peak area |
|---|---|---|---|---|---|---|---|---|
| 26 | 157 | 0.5702 | 156 | 160 | 158 | 0.67 | 493 | |
| 27 | 162 | 0.2971 | 160 | 163 | 161 | 0.52 | 240 | |
| 28 | 168 | 1.3959 | 163 | 178 | 168 | 1.83 | 404 | |
| 29 | 207 | 0.3579 | 197 | 207 | 204 | 1.30 | 227 | |
| 30 | 210 | 0.7382 | 207 | 215 | 211 | 0.93 | 328 | |
| 31 | 218 | 0.3298 | 215 | 221 | 218 | 0.81 | 165 | |
| 32 | 222 | 0.0851 | 221 | 226 | 222 | 0.43 | 113 | |
| 33 | 233 | 10.9318 | 226 | 251 | 236 | 1.88 | 7076 | |
| 34 | 252 | 0.5630 | 251 | 270 | 256 | 1.28 | 251 | |
| 35 | 290 | 0.6598 | 283 | 303 | 292 | 1.40 | 308 | |
| 36 | 308 | 0.3335 | 303 | 311 | 308 | 0.64 | 327 | |
| 37 | 313 | 0.3749 | 311 | 327 | 315 | 1.07 | 284 | |
| 38 | 500 (UM) | 0.5000 | 495 | 0 | 500 | 0.28 | 11899 | |

| | | |
|---|---|---|
| TIC: | 49.8526 | ng/uL |
| TIM: | 631.9974 | nmole/L |
| Total concentration: | 50.2970 | ng/uL |

Sample peak width (sec): 5 Sample min peak height: 50 Sample baseline V to V?: Y Sample baseline V to V points: 3
Sample filter: Binomial Number of points for filter: 3 Sample start region (min): 0 Sample end region (min): 80
Marker peak width (sec): 5 Marker min peak height: 500 Marker baseline V to V?: Y Marker baseline V to V points: 3
Lower marker selection: First peak > 500 RFU Upper marker selection: Last peak > 500 RFU
Ladder size (bp)1, 35, 75, 100, 150, 200, 250, 300, 400, 500
Quantification using: Upper Marker Final concentration (ng/uL): 0.5000 Dilution factor: 12.0

**Sample:** SampE4
**Well location:** E4
**Created:** Monday, 11 October 2021 14:41:52

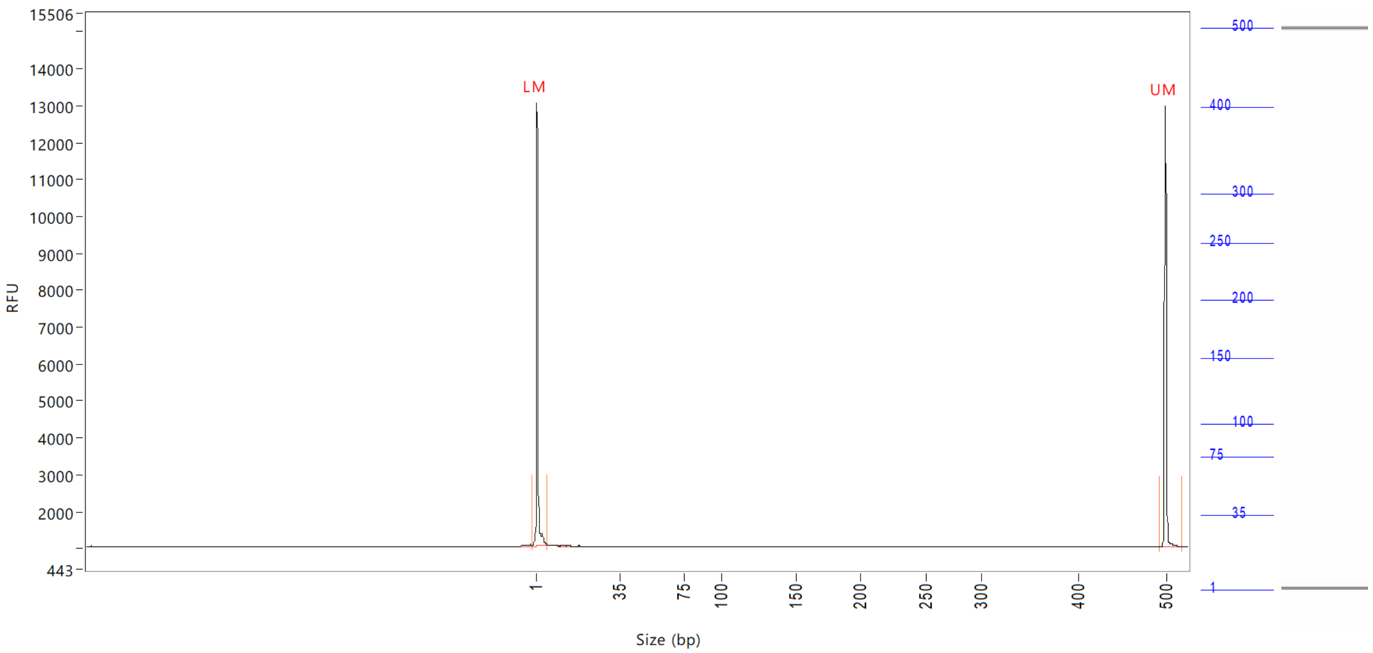


| Peak | Size (bp) | Concentration (ng/uL) | From (bp) | To (bp) | Average size (bp) | CV% | RFU | Corrected peak area |
|---|---|---|---|---|---|---|---|---|
| 1 | 1 (LM) | 0.9336 | 0 | 6 | 1 | 67.34 | 12025 | 45.326 |
| 2 | 500 (UM) | 0.5000 | 493 | 0 | 500 | 0.29 | 11923 | 24.275 |

| | | |
|---|---|---|
| TIC: | 0.0000 | ng/uL |
| TIM: | 0.0000 | nmole/L |
| Total concentration: | 0.5688 | ng/uL |

Sample peak width (sec): 5    Sample min peak height: 50    Sample baseline V to V?: Y    Sample baseline V to V points: 3
Sample filter: Binomial    Number of points for filter: 3    Sample start region (min): 0    Sample end region (min): 80
Marker peak width (sec): 5    Marker min peak height: 500    Marker baseline V to V?: Y    Marker baseline V to V points: 3
Lower marker selection: First peak > 500 RFU    Upper marker selection: Last peak > 500 RFU
Ladder size (bp)1, 35, 75, 100, 150, 200, 250, 300, 400, 500
Quantification using: Upper Marker    Final concentration (ng/uL): 0.5000    Dilution factor: 12.0

**Sample:** SampF4
**Well location:** F4
**Created:** Monday, 11 October 2021 14:41:52

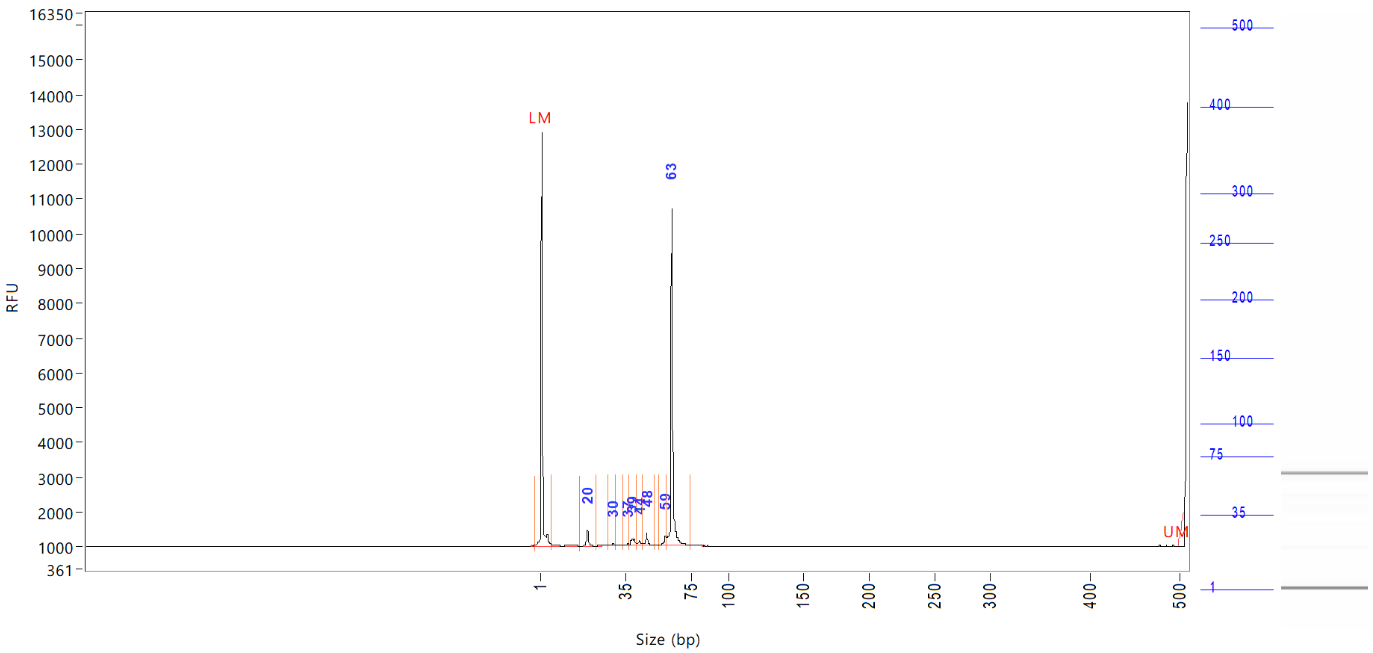


| Peak | Size (bp) | Concentration (ng/uL) | From (bp) | To (bp) | Average size (bp) | CV% | RFU | Corrected peak area |
|---|---|---|---|---|---|---|---|---|
| 1 | 1 (LM) | Inf | 0 | 6 | 1 | 67.30 | 11897 | 44.663 |
| 2 | 20 | Inf | 17 | 23 | 20 | 2.92 | 437 | 2.074 |
| 3 | 30 | Inf | 28 | 31 | 30 | 1.29 | 54 | 0.176 |
| 4 | 37 | Inf | 34 | 38 | 36 | 2.51 | 68 | 0.286 |
| 5 | 39 | Inf | 38 | 42 | 40 | 2.53 | 194 | 1.653 |
| 6 | 44 | Inf | 42 | 46 | 44 | 2.30 | 138 | 0.904 |
| 7 | 48 | Inf | 46 | 53 | 48 | 1.99 | 355 | 1.468 |
| 8 | 59 | Inf | 56 | 60 | 59 | 1.47 | 273 | 1.328 |
| 9 | 500 (UM) | Inf | 60 | 75 | 63 | 2.16 | 9677 | 32.587 |

| | | |
|---|---|---|
| TIC: | Inf | ng/uL |
| TIM: | Inf | nmole/L |
| Total concentration: | 0.0000 | ng/uL |

Sample peak width (sec): 5  Sample min peak height: 50  Sample baseline V to V?: Y  Sample baseline V to V points: 3
Sample filter: Binomial  Number of points for filter: 3  Sample start region (min): 0  Sample end region (min): 80
Marker peak width (sec): 5  Marker min peak height: 500  Marker baseline V to V?: Y  Marker baseline V to V points: 3
Lower marker selection: First peak > 500 RFU  Upper marker selection: Last peak > 500 RFU
Ladder size (bp)1, 35, 75, 100, 150, 200, 250, 300, 400, 500
Quantification using: Upper Marker  Final concentration (ng/uL): 0.5000  Dilution factor: 12.0

**Sample:** SampG4
**Well location:** G4
**Created:** Monday, 11 October 2021 14:41:52

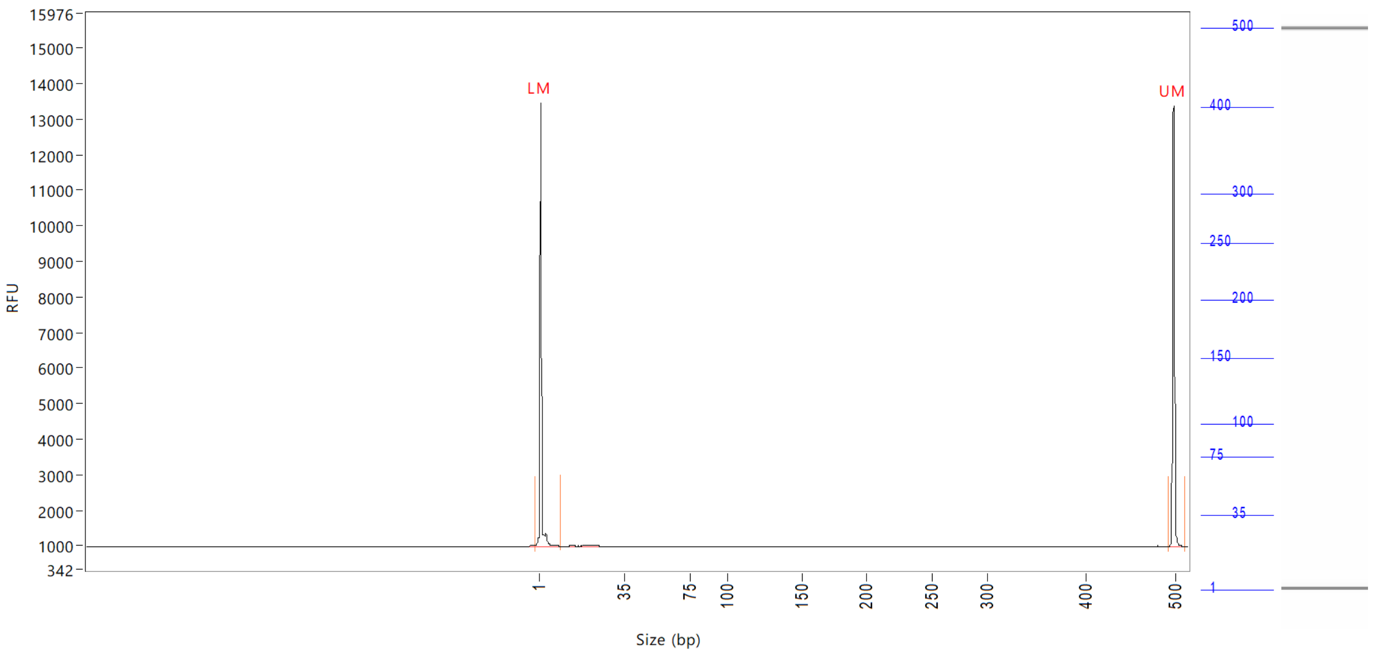


| Peak | Size (bp) | Concentration (ng/uL) | From (bp) | To (bp) | Average size (bp) | CV% | RFU | Corrected peak area |
|---|---|---|---|---|---|---|---|---|
| 1 | 1 (LM) | 0.8975 | 0 | 10 | 1 | 76.33 | 12487 | 47.287 |
| 2 | 500 (UM) | 0.5000 | 493 | 513 | 500 | 0.25 | 12398 | 26.343 |

| | | |
|---|---|---|
| TIC: | 0.0000 | ng/uL |
| TIM: | 0.0000 | nmole/L |
| Total concentration: | 0.4542 | ng/uL |

```
Sample peak width (sec): 5    Sample min peak height: 50    Sample baseline V to V?: Y    Sample baseline V to V points: 3
Sample filter: Binomial       Number of points for filter: 3  Sample start region (min): 0  Sample end region (min): 80
Marker peak width (sec): 5    Marker min peak height: 500   Marker baseline V to V?: Y    Marker baseline V to V points: 3
Lower marker selection: First peak > 500 RFU                Upper marker selection: Last peak > 500 RFU
Ladder size (bp)1, 35, 75, 100, 150, 200, 250, 300, 400, 500
Quantification using: Upper Marker          Final concentration (ng/uL): 0.5000          Dilution factor: 12.0
```

**Sample:** SampH4
**Well location:** H4
**Created:** Monday, 11 October 2021 14:41:52

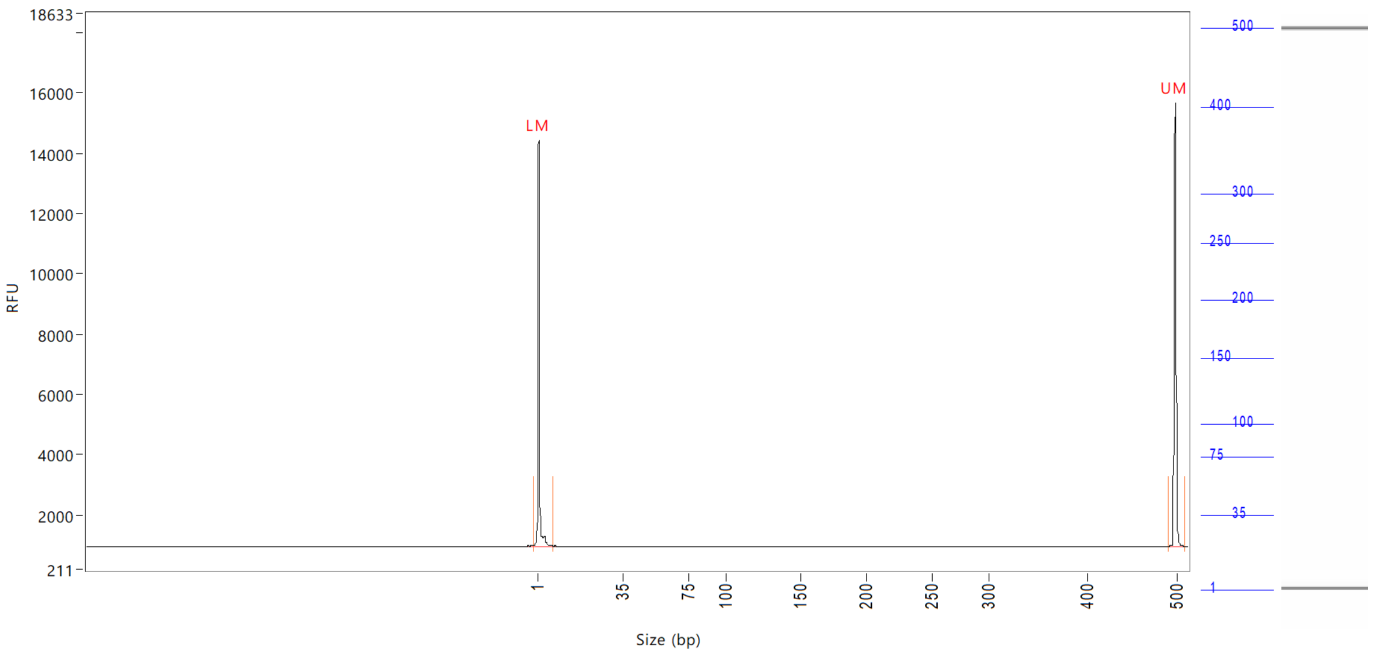


| Peak | Size (bp) | Concentration (ng/uL) | From (bp) | To (bp) | Average size (bp) | CV% | RFU | Corrected peak area |
|---|---|---|---|---|---|---|---|---|
| 1 | 1 (LM) | 0.8079 | 0 | 8 | 1 | 73.45 | 13455 | 50.984 |
| 2 | 500 (UM) | 0.5000 | 492 | 512 | 500 | 0.24 | 14724 | 31.554 |

| | | |
|---|---|---|
| TIC: | 0.0000 | ng/uL |
| TIM: | 0.0000 | nmole/L |
| Total concentration: | 0.3467 | ng/uL |

```
Sample peak width (sec): 5    Sample min peak height: 50    Sample baseline V to V?: Y    Sample baseline V to V points: 3
Sample filter: Binomial       Number of points for filter: 3  Sample start region (min): 0  Sample end region (min): 80
Marker peak width (sec): 5    Marker min peak height: 500   Marker baseline V to V?: Y    Marker baseline V to V points: 3
Lower marker selection: First peak > 500 RFU                Upper marker selection: Last peak > 500 RFU
Ladder size (bp)1, 35, 75, 100, 150, 200, 250, 300, 400, 500
Quantification using: Upper Marker          Final concentration (ng/uL): 0.5000          Dilution factor: 12.0
```

**Sample:** SampA5
**Well location:** A5
**Created:** Monday, 11 October 2021 14:41:52

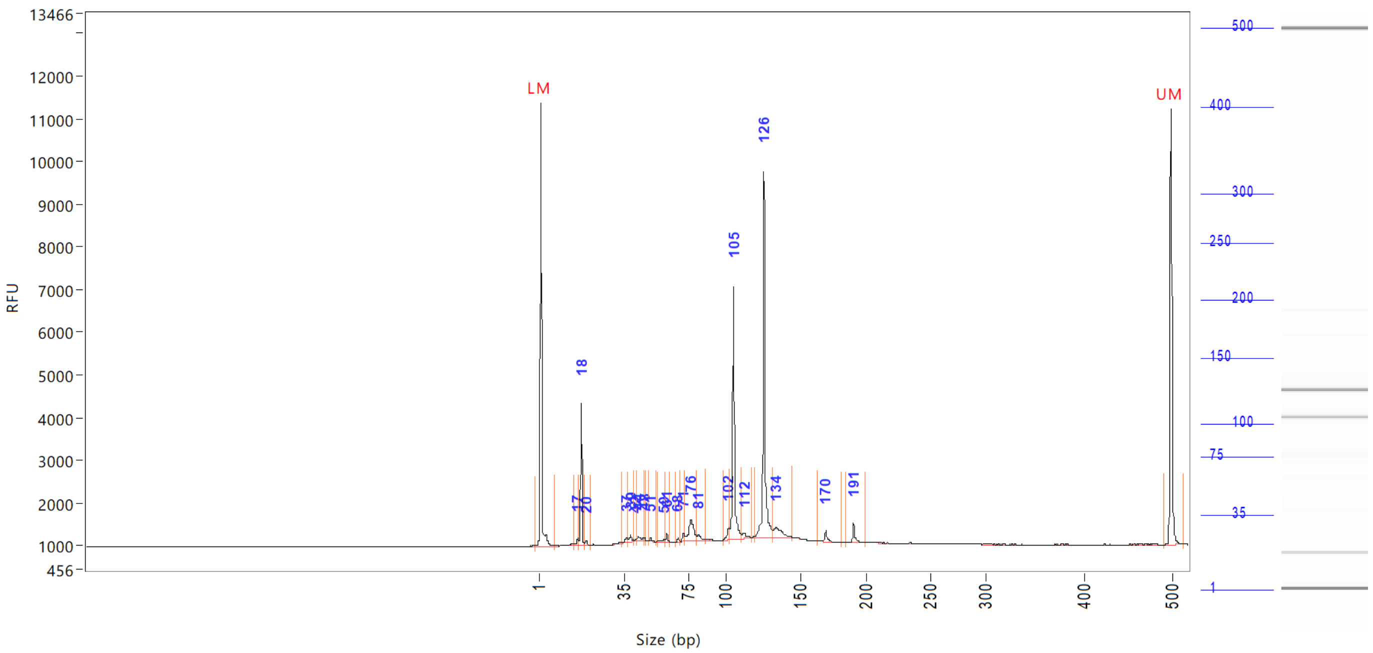


| Peak | Size<br>(bp) | Concentration<br>(ng/uL) | From<br>(bp) | To<br>(bp) | Average size<br>(bp) | CV% | RFU | Corrected peak area |
|---|---|---|---|---|---|---|---|---|
| 1 | 1 (LM) | 0.8737 | 0 | 7 | 1 | 72.84 | 10393 | 39.037 |
| 2 | 17 | 0.2023 | 15 | 17 | 16 | 2.57 | 158 | 0.753 |
| 3 | 18 | 2.9435 | 17 | 19 | 18 | 1.64 | 3331 | 10.959 |
| 4 | 20 | 0.1353 | 19 | 22 | 20 | 2.41 | 119 | 0.504 |
| 5 | 37 | 0.1698 | 34 | 38 | 36 | 2.60 | 99 | 0.632 |
| 6 | 39 | 0.2255 | 38 | 42 | 39 | 2.13 | 153 | 0.839 |
| 7 | 42 | 0.0409 | 42 | 43 | 42 | 0.81 | 55 | 0.152 |
| 8 | 44 | 0.2507 | 43 | 47 | 45 | 2.74 | 106 | 0.934 |
| 9 | 48 | 0.0635 | 47 | 49 | 48 | 0.64 | 92 | 0.237 |
| 10 | 51 | 0.0801 | 50 | 55 | 52 | 1.32 | 79 | 0.298 |
| 11 | 59 | 0.1249 | 56 | 60 | 58 | 2.12 | 54 | 0.465 |
| 12 | 61 | 0.1650 | 60 | 63 | 61 | 0.77 | 193 | 0.614 |
| 13 | 68 | 0.0723 | 66 | 70 | 68 | 0.75 | 85 | 0.269 |
| 14 | 71 | 0.2239 | 70 | 72 | 71 | 0.86 | 193 | 0.834 |
| 15 | 76 | 1.3209 | 72 | 80 | 77 | 2.48 | 513 | 4.918 |
| 16 | 81 | 0.2541 | 80 | 86 | 82 | 1.58 | 131 | 0.946 |
| 17 | 102 | 0.3367 | 97 | 103 | 101 | 1.24 | 287 | 1.254 |
| 18 | 105 | 5.3924 | 103 | 111 | 105 | 1.22 | 5924 | 20.077 |
| 19 | 112 | 0.2402 | 111 | 117 | 113 | 1.23 | 131 | 0.894 |
| 20 | 126 | 7.5883 | 120 | 131 | 126 | 1.19 | 8563 | 28.253 |
| 21 | 134 | 0.8696 | 131 | 144 | 135 | 2.07 | 238 | 3.238 |
| 22 | 170 | 0.2861 | 164 | 181 | 170 | 1.25 | 265 | 1.065 |
| 23 | 191 | 0.4213 | 186 | 201 | 191 | 0.95 | 447 | 1.569 |
| 24 | 500 (UM) | 0.5000 | 492 | 513 | 500 | 0.28 | 10187 | 22.340 |

Sample peak width (sec): 5  Sample min peak height: 50  Sample baseline V to V?: Y  Sample baseline V to V points: 3
Sample filter: Binomial  Number of points for filter: 3  Sample start region (min): 0  Sample end region (min): 80
Marker peak width (sec): 5  Marker min peak height: 500  Marker baseline V to V?: Y  Marker baseline V to V points: 3
Lower marker selection: First peak > 500 RFU  Upper marker selection: Last peak > 500 RFU
Ladder size (bp)1, 35, 75, 100, 150, 200, 250, 300, 400, 500
Quantification using: Upper Marker  Final concentration (ng/uL): 0.5000  Dilution factor: 12.0

**Sample:** SampA5
**Well location:** A5
**Created:** Monday, 11 October 2021 14:41:52

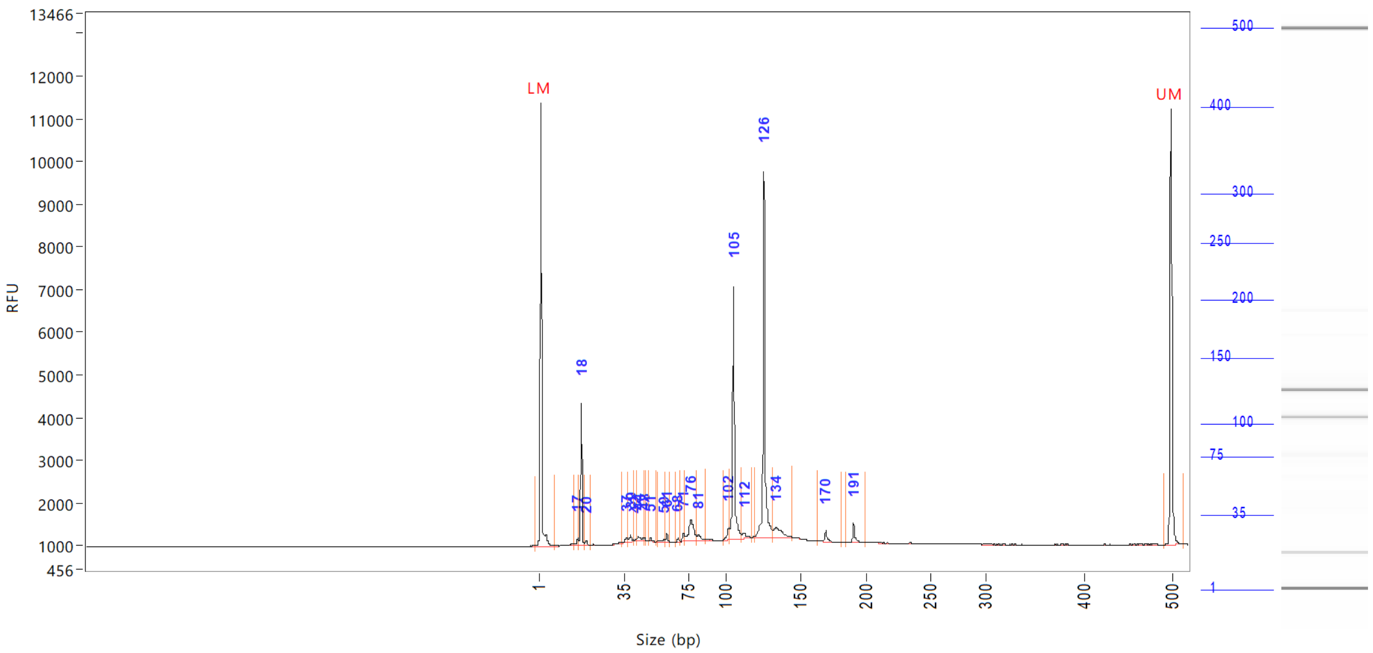

| Peak | Size (bp) | Concentration (ng/uL) | From (bp) | To (bp) | Average size (bp) | CV% | RFU | Corrected peak area |
|---|---|---|---|---|---|---|---|---|

| | | |
|---|---|---|
| TIC: | 21.4072 | ng/uL |
| TIM: | 586.7840 | nmole/L |
| Total concentration: | 21.9745 | ng/uL |

```
Sample peak width (sec): 5    Sample min peak height: 50    Sample baseline V to V?: Y    Sample baseline V to V points: 3
Sample filter: Binomial       Number of points for filter: 3  Sample start region (min): 0  Sample end region (min): 80
Marker peak width (sec): 5    Marker min peak height: 500   Marker baseline V to V?: Y    Marker baseline V to V points: 3
Lower marker selection: First peak > 500 RFU                Upper marker selection: Last peak > 500 RFU
Ladder size (bp)1, 35, 75, 100, 150, 200, 250, 300, 400, 500
Quantification using: Upper Marker         Final concentration (ng/uL): 0.5000         Dilution factor: 12.0
```

**Sample:** SampB5
**Well location:** B5
**Created:** Monday, 11 October 2021 14:41:52

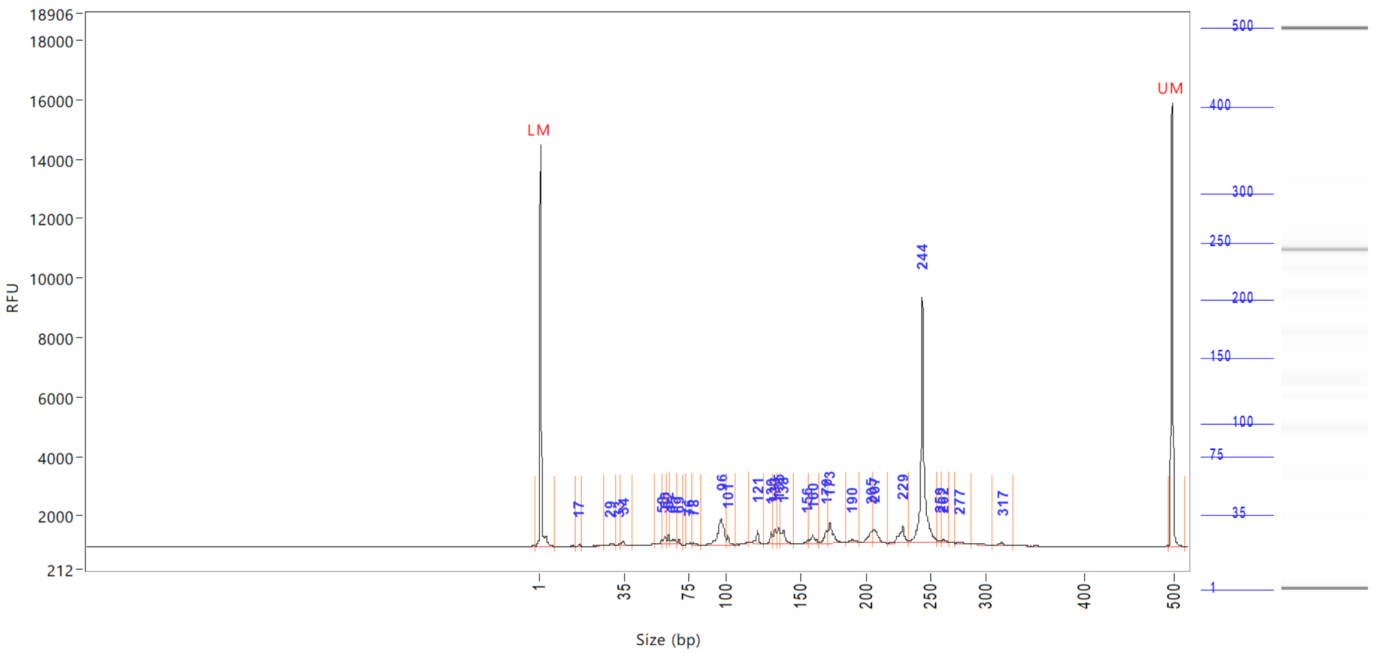

| Peak | Size (bp) | Concentration (ng/uL) | From (bp) | To (bp) | Average size (bp) | CV% | RFU | Corrected peak area |
|---|---|---|---|---|---|---|---|---|
| 1 | 1 (LM) | 0.8294 | 0 | 8 | 1 | 72.58 | 13500 | 51.001 |
| 2 | 17 | 0.0637 | 16 | 18 | 17 | 2.29 | 81 | 0.327 |
| 3 | 29 | 0.1538 | 27 | 32 | 30 | 3.39 | 73 | 0.788 |
| 4 | 33 | 0.0628 | 32 | 33 | 33 | 1.13 | 67 | 0.322 |
| 5 | 34 | 0.2145 | 33 | 40 | 35 | 3.82 | 160 | 1.099 |
| 6 | 58 | 0.1241 | 54 | 59 | 58 | 1.54 | 135 | 0.636 |
| 7 | 60 | 0.2330 | 59 | 61 | 60 | 1.03 | 263 | 1.194 |
| 8 | 62 | 0.1812 | 61 | 63 | 62 | 0.77 | 266 | 0.929 |
| 9 | 65 | 0.2068 | 63 | 67 | 65 | 1.55 | 147 | 1.059 |
| 10 | 69 | 0.0931 | 67 | 71 | 68 | 0.64 | 157 | 0.477 |
| 11 | 76 | 0.0515 | 73 | 77 | 76 | 1.42 | 53 | 0.264 |
| 12 | 78 | 0.0671 | 77 | 84 | 79 | 1.78 | 54 | 0.344 |
| 13 | 96 | 1.9356 | 84 | 100 | 96 | 2.36 | 907 | 9.918 |
| 14 | 101 | 0.2247 | 100 | 106 | 102 | 1.14 | 315 | 1.151 |
| 15 | 121 | 0.3623 | 116 | 125 | 120 | 1.13 | 458 | 1.857 |
| 16 | 130 | 0.3401 | 125 | 131 | 130 | 0.79 | 431 | 1.743 |
| 17 | 133 | 0.4579 | 131 | 134 | 133 | 0.55 | 505 | 2.346 |
| 18 | 135 | 0.4908 | 134 | 136 | 135 | 0.51 | 580 | 2.515 |
| 19 | 138 | 0.6536 | 136 | 146 | 139 | 1.10 | 480 | 3.349 |
| 20 | 156 | 0.1479 | 146 | 157 | 154 | 1.62 | 127 | 0.758 |
| 21 | 160 | 0.3810 | 157 | 165 | 160 | 1.13 | 274 | 1.953 |
| 22 | 170 | 0.4449 | 165 | 171 | 169 | 0.89 | 430 | 2.280 |
| 23 | 173 | 0.9612 | 171 | 185 | 174 | 1.30 | 691 | 4.925 |
| 24 | 190 | 0.1845 | 185 | 196 | 190 | 1.09 | 112 | 0.945 |
| 25 | 205 | 0.4624 | 196 | 206 | 203 | 0.95 | 374 | 2.369 |

Sample peak width (sec): 5 Sample min peak height: 50 Sample baseline V to V?: Y Sample baseline V to V points: 3
Sample filter: Binomial Number of points for filter: 3 Sample start region (min): 0 Sample end region (min): 80
Marker peak width (sec): 5 Marker min peak height: 500 Marker baseline V to V?: Y Marker baseline V to V points: 3
Lower marker selection: First peak > 500 RFU Upper marker selection: Last peak > 500 RFU
Ladder size (bp)1, 35, 75, 100, 150, 200, 250, 300, 400, 500
Quantification using: Upper Marker Final concentration (ng/uL): 0.5000 Dilution factor: 12.0

**Sample:** SampB5
**Well location:** B5
**Created:** Monday, 11 October 2021 14:41:52

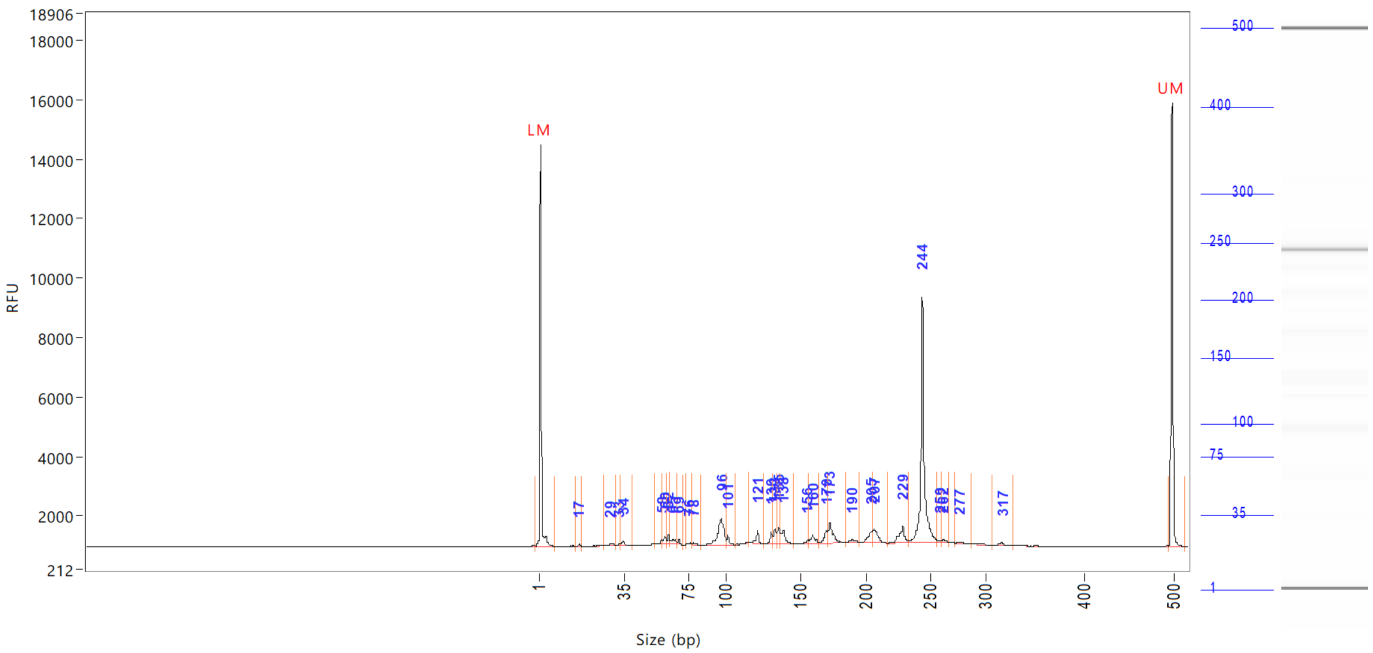


| Peak | Size (bp) | Concentration (ng/uL) | From (bp) | To (bp) | Average size (bp) | CV% | RFU | Corrected peak area |
|---|---|---|---|---|---|---|---|---|
| 26 | 207 | 0.6569 | 206 | 217 | 208 | 0.97 | 455 | |
| 27 | 229 | 0.8157 | 217 | 233 | 228 | 1.09 | 580 | |
| 28 | 244 | 7.0531 | 233 | 257 | 244 | 1.22 | 8230 | |
| 29 | 259 | 0.0681 | 257 | 261 | 259 | 0.42 | 72 | |
| 30 | 262 | 0.1002 | 261 | 267 | 263 | 0.62 | 99 | |
| 31 | 277 | 0.0664 | 273 | 287 | 278 | 0.74 | 61 | |
| 32 | 317 | 0.1069 | 308 | 328 | 317 | 0.96 | 86 | |
| 33 | 500 (UM) | 0.5000 | 496 | 515 | 500 | 0.25 | 14936 | |

| | | |
|---|---|---|
| TIC: | 17.3658 | ng/uL |
| TIM: | 202.4147 | nmole/L |
| Total concentration: | 17.5167 | ng/uL |

Sample peak width (sec): 5 Sample min peak height: 50 Sample baseline V to V?: Y Sample baseline V to V points: 3
Sample filter: Binomial Number of points for filter: 3 Sample start region (min): 0 Sample end region (min): 80
Marker peak width (sec): 5 Marker min peak height: 500 Marker baseline V to V?: Y Marker baseline V to V points: 3
Lower marker selection: First peak > 500 RFU Upper marker selection: Last peak > 500 RFU
Ladder size (bp)1, 35, 75, 100, 150, 200, 250, 300, 400, 500
Quantification using: Upper Marker Final concentration (ng/uL): 0.5000 Dilution factor: 12.0

**Sample:** SampC5
**Well location:** C5
**Created:** Monday, 11 October 2021 14:41:52

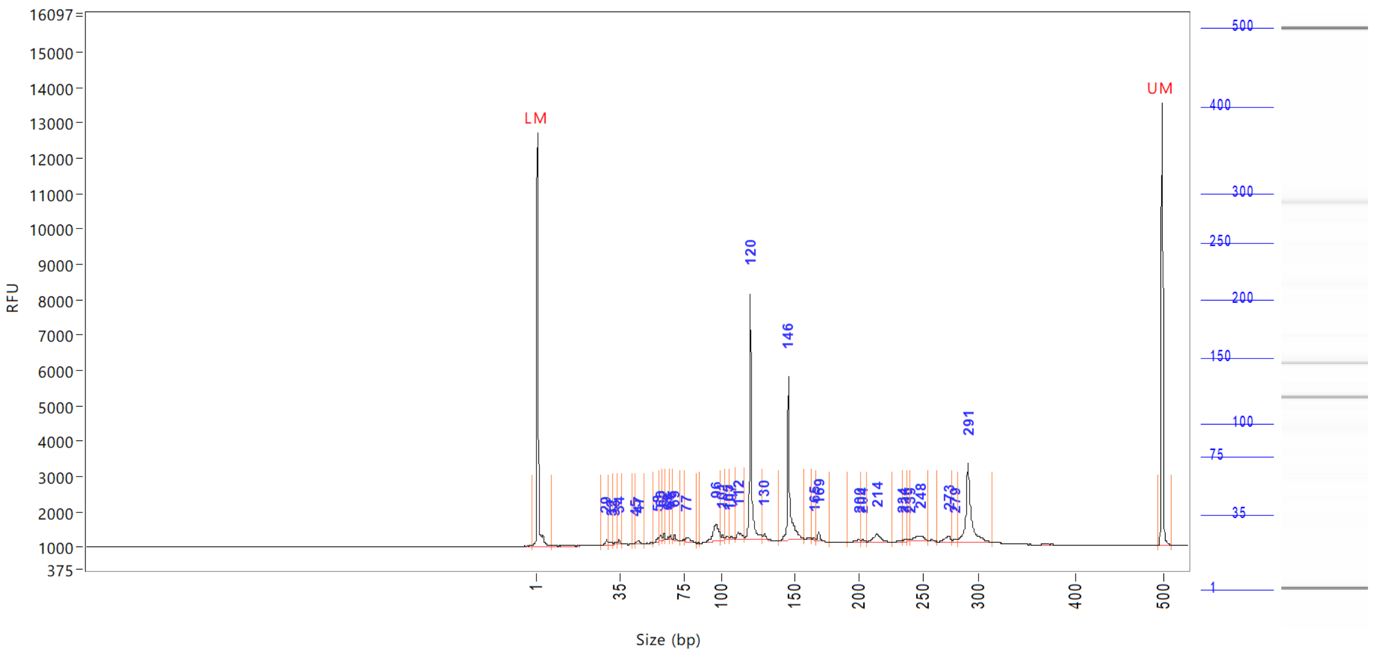


| Peak | Size (bp) | Concentration (ng/uL) | From (bp) | To (bp) | Average size (bp) | CV% | RFU | Corrected peak area |
|---|---|---|---|---|---|---|---|---|
| 1 | 1 (LM) | 0.8858 | 0 | 8 | 2 | 65.42 | 11693 | 44.474 |
| 2 | 29 | 0.1988 | 27 | 30 | 29 | 1.99 | 151 | 0.832 |
| 3 | 32 | 0.1321 | 30 | 32 | 31 | 1.57 | 77 | 0.553 |
| 4 | 33 | 0.0969 | 32 | 34 | 33 | 1.46 | 55 | 0.405 |
| 5 | 34 | 0.1311 | 34 | 37 | 34 | 1.18 | 125 | 0.548 |
| 6 | 45 | 0.0568 | 44 | 46 | 45 | 1.13 | 57 | 0.238 |
| 7 | 47 | 0.1599 | 46 | 50 | 47 | 2.01 | 116 | 0.669 |
| 8 | 58 | 0.1153 | 56 | 59 | 58 | 0.93 | 121 | 0.482 |
| 9 | 60 | 0.1963 | 59 | 62 | 60 | 1.00 | 190 | 0.821 |
| 10 | 62 | 0.1330 | 62 | 64 | 62 | 0.61 | 186 | 0.556 |
| 11 | 65 | 0.0628 | 64 | 66 | 65 | 0.78 | 77 | 0.263 |
| 12 | 66 | 0.0929 | 66 | 68 | 66 | 0.52 | 143 | 0.389 |
| 13 | 69 | 0.1319 | 68 | 72 | 69 | 1.02 | 170 | 0.552 |
| 14 | 77 | 0.2765 | 75 | 84 | 78 | 2.20 | 117 | 1.157 |
| 15 | 96 | 1.2584 | 86 | 100 | 96 | 2.15 | 480 | 5.265 |
| 16 | 101 | 0.1462 | 100 | 103 | 101 | 0.93 | 164 | 0.612 |
| 17 | 105 | 0.1346 | 103 | 106 | 104 | 0.87 | 91 | 0.563 |
| 18 | 107 | 0.0797 | 106 | 110 | 107 | 0.73 | 61 | 0.333 |
| 19 | 112 | 0.3519 | 110 | 117 | 113 | 1.35 | 182 | 1.472 |
| 20 | 120 | 5.5646 | 117 | 129 | 120 | 1.45 | 6924 | 23.282 |
| 21 | 130 | 0.2055 | 129 | 140 | 130 | 1.33 | 200 | 0.860 |
| 22 | 146 | 4.1605 | 140 | 158 | 146 | 1.66 | 4636 | 17.407 |
| 23 | 165 | 0.0588 | 164 | 167 | 165 | 0.48 | 60 | 0.246 |
| 24 | 169 | 0.2295 | 167 | 177 | 169 | 0.80 | 264 | 0.960 |
| 25 | 200 | 0.1786 | 192 | 202 | 198 | 1.24 | 93 | 0.747 |

```
Sample peak width (sec): 5    Sample min peak height: 50    Sample baseline V to V?: Y    Sample baseline V to V points: 3
Sample filter: Binomial       Number of points for filter: 3  Sample start region (min): 0  Sample end region (min): 80
Marker peak width (sec): 5    Marker min peak height: 500   Marker baseline V to V?: Y    Marker baseline V to V points: 3
Lower marker selection: First peak > 500 RFU                Upper marker selection: Last peak > 500 RFU
Ladder size (bp)1, 35, 75, 100, 150, 200, 250, 300, 400, 500
Quantification using: Upper Marker          Final concentration (ng/uL): 0.5000          Dilution factor: 12.0
```

**Sample:** SampC5
**Well location:** C5
**Created:** Monday, 11 October 2021 14:41:52

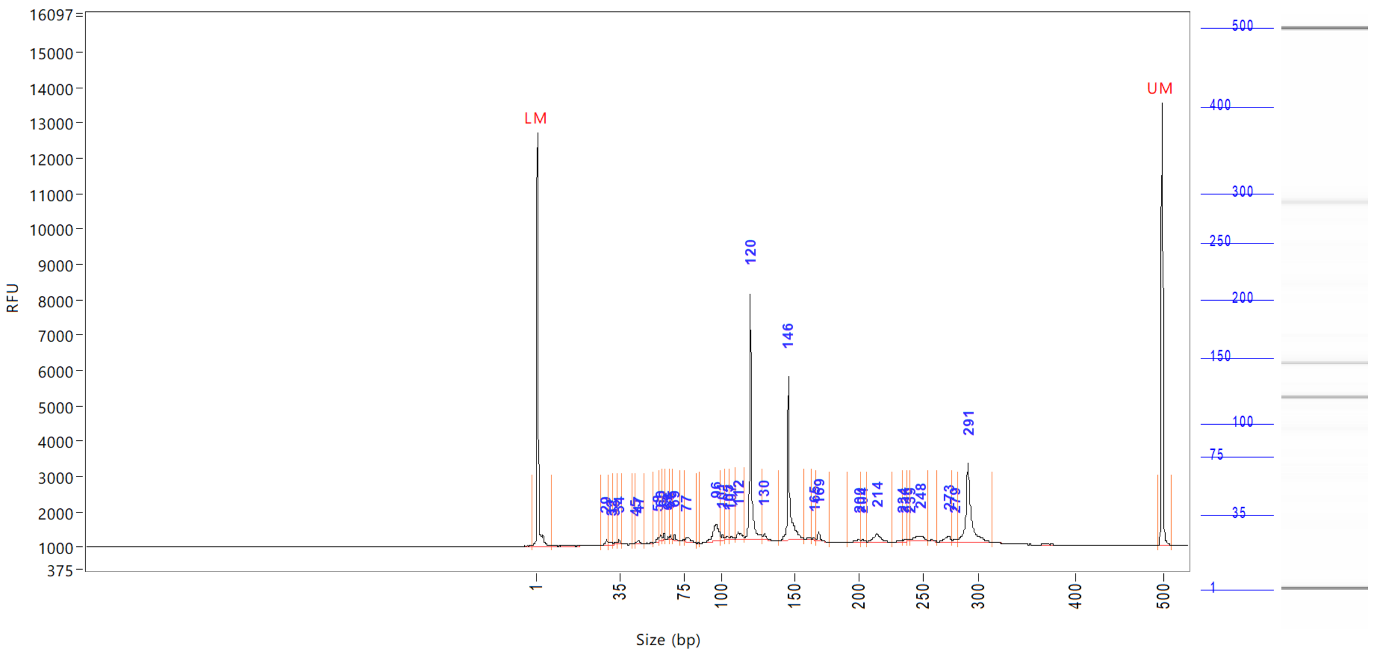


| Peak | Size (bp) | Concentration (ng/uL) | From (bp) | To (bp) | Average size (bp) | CV% | RFU | Corrected peak area |
|---|---|---|---|---|---|---|---|---|
| 26 | 204 | 0.0890 | 202 | 207 | 204 | 0.57 | 77 | |
| 27 | 214 | 0.7053 | 207 | 226 | 215 | 1.66 | 262 | |
| 28 | 234 | 0.0752 | 226 | 235 | 232 | 0.75 | 51 | |
| 29 | 236 | 0.0747 | 235 | 238 | 236 | 0.36 | 72 | |
| 30 | 239 | 0.0620 | 238 | 240 | 239 | 0.33 | 64 | |
| 31 | 248 | 0.4977 | 240 | 255 | 247 | 1.36 | 159 | |
| 32 | 273 | 0.3005 | 264 | 277 | 272 | 0.94 | 166 | |
| 33 | 279 | 0.1023 | 277 | 281 | 279 | 0.48 | 82 | |
| 34 | 291 | 3.5050 | 281 | 316 | 292 | 1.58 | 2228 | |
| 35 | 500 (UM) | 0.5000 | 496 | 510 | 500 | 0.25 | 12545 | |

| | | |
|---|---|---|
| TIC: | 19.5642 | ng/uL |
| TIM: | 257.2003 | nmole/L |
| Total concentration: | 20.0823 | ng/uL |

Sample peak width (sec): 5    Sample min peak height: 50    Sample baseline V to V?: Y    Sample baseline V to V points: 3
Sample filter: Binomial    Number of points for filter: 3  Sample start region (min): 0  Sample end region (min): 80
Marker peak width (sec): 5    Marker min peak height: 500    Marker baseline V to V?: Y    Marker baseline V to V points: 3
Lower marker selection: First peak > 500 RFU    Upper marker selection: Last peak > 500 RFU
Ladder size (bp)1, 35, 75, 100, 150, 200, 250, 300, 400, 500
Quantification using: Upper Marker    Final concentration (ng/uL): 0.5000    Dilution factor: 12.0

**Sample:** SampD5
**Well location:** D5
**Created:** Monday, 11 October 2021 14:41:52

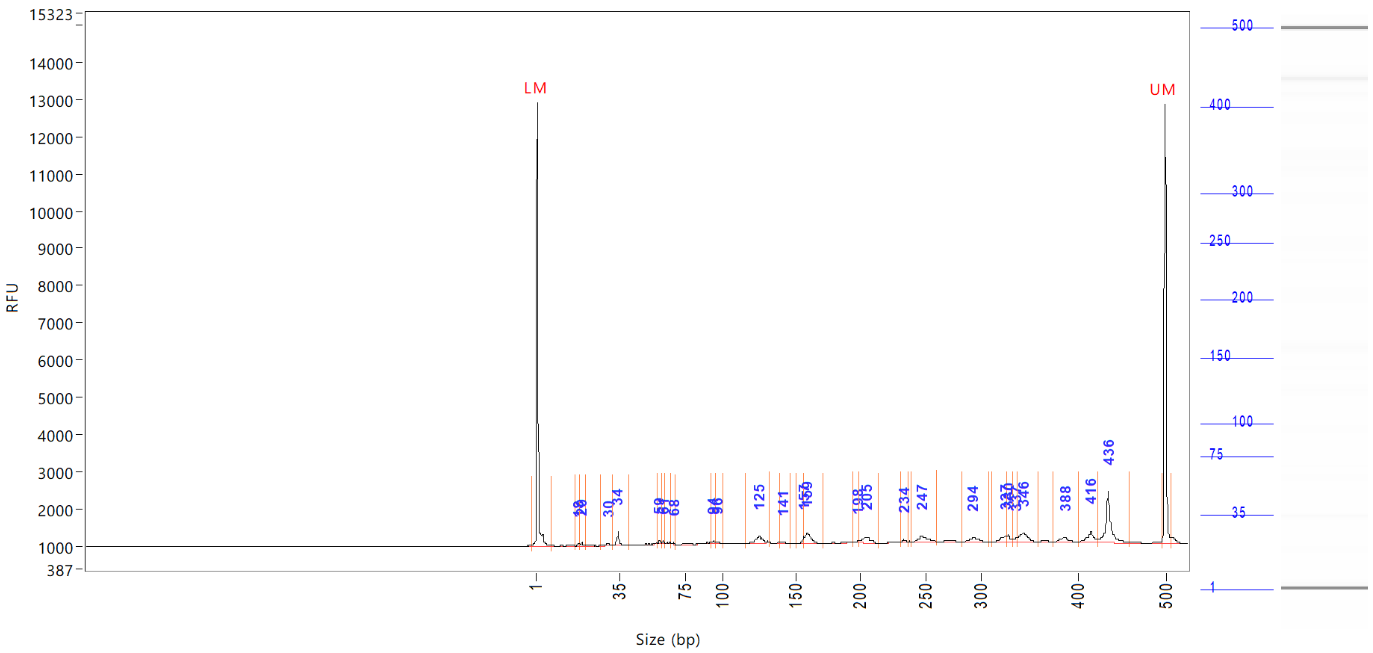


| Peak | Size (bp) | Concentration (ng/uL) | From (bp) | To (bp) | Average size (bp) | CV% | RFU | Corrected peak area |
|---|---|---|---|---|---|---|---|---|
| 1 | 1 (LM) | 0.8723 | 0 | 8 | 1 | 72.50 | 11936 | 44.210 |
| 2 | 18 | 0.0926 | 17 | 19 | 18 | 2.21 | 69 | 0.391 |
| 3 | 20 | 0.1023 | 19 | 22 | 20 | 2.56 | 104 | 0.432 |
| 4 | 30 | 0.1489 | 27 | 32 | 30 | 3.45 | 51 | 0.629 |
| 5 | 34 | 0.4909 | 32 | 41 | 34 | 3.95 | 372 | 2.073 |
| 6 | 59 | 0.0860 | 58 | 61 | 59 | 0.95 | 90 | 0.363 |
| 7 | 61 | 0.0557 | 61 | 63 | 61 | 0.61 | 77 | 0.235 |
| 8 | 68 | 0.0331 | 67 | 69 | 68 | 0.57 | 52 | 0.140 |
| 9 | 94 | 0.1040 | 92 | 95 | 94 | 0.85 | 78 | 0.439 |
| 10 | 96 | 0.1108 | 95 | 101 | 97 | 1.18 | 72 | 0.468 |
| 11 | 125 | 0.5447 | 116 | 132 | 125 | 2.23 | 177 | 2.301 |
| 12 | 141 | 0.1079 | 139 | 146 | 141 | 1.04 | 55 | 0.456 |
| 13 | 157 | 0.1848 | 151 | 157 | 156 | 0.77 | 198 | 0.781 |
| 14 | 159 | 0.6533 | 157 | 171 | 161 | 1.59 | 275 | 2.759 |
| 15 | 198 | 0.0608 | 194 | 199 | 198 | 0.48 | 69 | 0.257 |
| 16 | 205 | 0.4291 | 199 | 215 | 205 | 1.39 | 156 | 1.812 |
| 17 | 234 | 0.0801 | 231 | 237 | 234 | 0.64 | 65 | 0.338 |
| 18 | 247 | 0.4258 | 240 | 261 | 249 | 1.55 | 152 | 1.798 |
| 19 | 294 | 0.3496 | 284 | 310 | 295 | 1.40 | 115 | 1.476 |
| 20 | 327 | 0.3005 | 314 | 328 | 323 | 0.94 | 166 | 1.269 |
| 21 | 330 | 0.2105 | 328 | 335 | 330 | 0.58 | 186 | 0.889 |
| 22 | 337 | 0.0970 | 335 | 339 | 337 | 0.34 | 112 | 0.410 |
| 23 | 346 | 0.6040 | 339 | 360 | 345 | 1.13 | 235 | 2.551 |
| 24 | 388 | 0.3501 | 375 | 402 | 388 | 1.33 | 134 | 1.479 |
| 25 | 416 | 0.5490 | 402 | 424 | 415 | 1.12 | 308 | 2.319 |

```
Sample peak width (sec): 5     Sample min peak height: 50    Sample baseline V to V?: Y    Sample baseline V to V points: 3
Sample filter: Binomial        Number of points for filter: 3  Sample start region (min): 0  Sample end region (min): 80
Marker peak width (sec): 5     Marker min peak height: 500   Marker baseline V to V?: Y    Marker baseline V to V points: 3
Lower marker selection: First peak > 500 RFU                 Upper marker selection: Last peak > 500 RFU
Ladder size (bp)1, 35, 75, 100, 150, 200, 250, 300, 400, 500
Quantification using: Upper Marker          Final concentration (ng/uL): 0.5000          Dilution factor: 12.0
```

**Sample:** SampD5
**Well location:** D5
**Created:** Monday, 11 October 2021 14:41:52

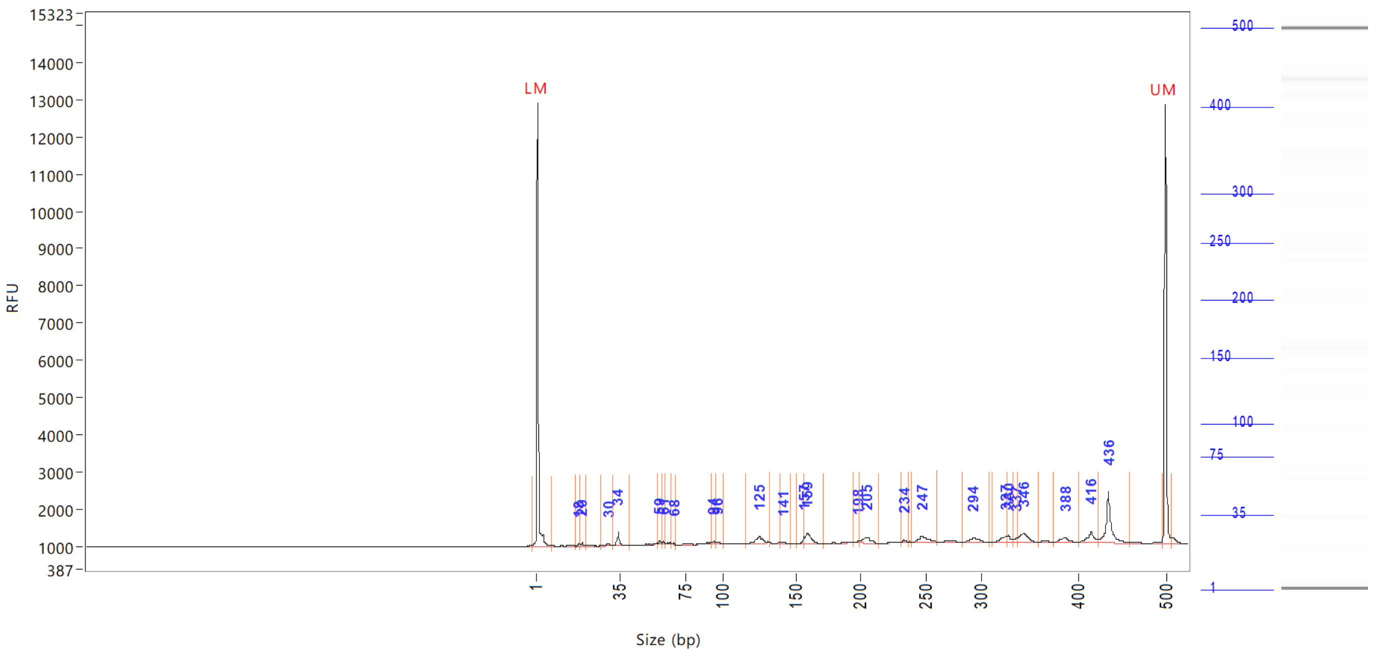


| Peak | Size (bp) | Concentration (ng/uL) | From (bp) | To (bp) | Average size (bp) | CV% | RFU | Corrected peak area |
|---|---|---|---|---|---|---|---|---|
| 26 | 436 | 1.8406 | 424 | 459 | 437 | 1.28 | 1368 | |
| 27 | 500 (UM) | 0.5000 | 496 | 506 | 500 | 0.25 | 11783 | |

| | | |
|---|---|---|
| TIC: | 8.0121 | ng/uL |
| TIM: | 99.5597 | nmole/L |
| Total concentration: | 8.9587 | ng/uL |

```
Sample peak width (sec): 5    Sample min peak height: 50    Sample baseline V to V?: Y    Sample baseline V to V points: 3
Sample filter: Binomial       Number of points for filter: 3  Sample start region (min): 0  Sample end region (min): 80
Marker peak width (sec): 5    Marker min peak height: 500   Marker baseline V to V?: Y    Marker baseline V to V points: 3
Lower marker selection: First peak > 500 RFU                Upper marker selection: Last peak > 500 RFU
Ladder size (bp)1, 35, 75, 100, 150, 200, 250, 300, 400, 500
Quantification using: Upper Marker          Final concentration (ng/uL): 0.5000          Dilution factor: 12.0
```

**Sample:** SampE5
**Well location:** E5
**Created:** Monday, 11 October 2021 14:41:52

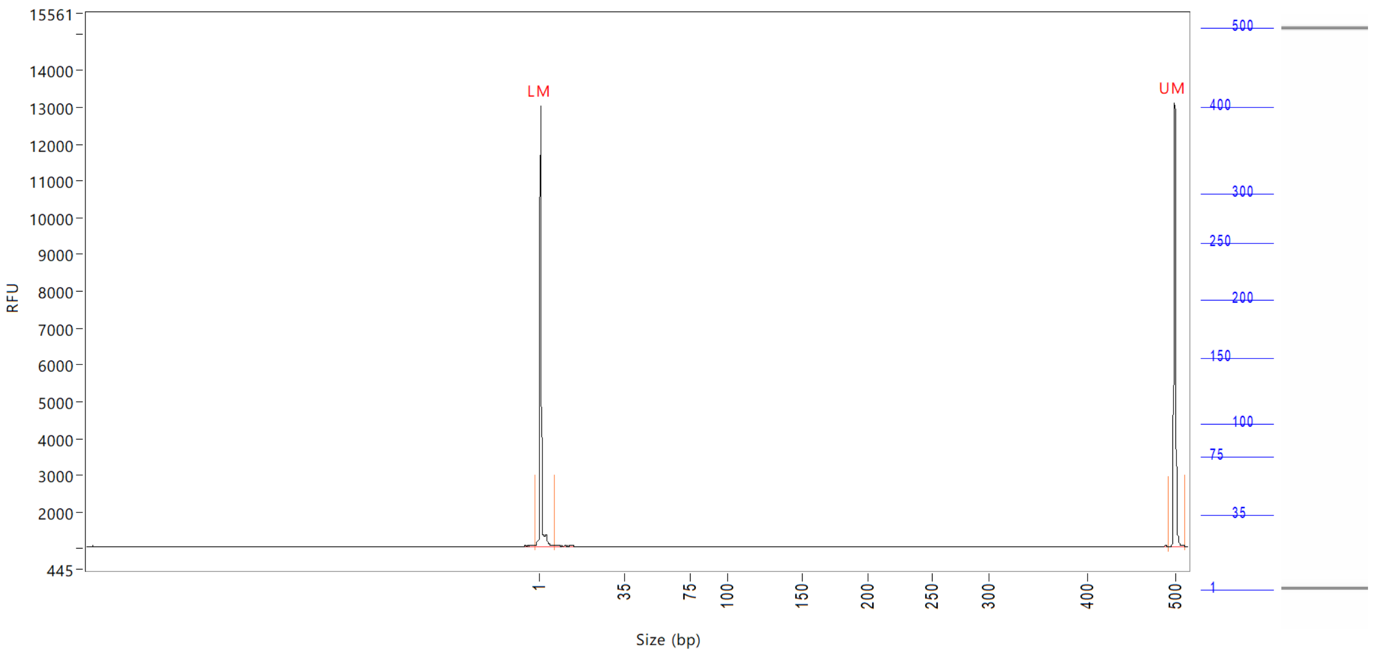


| Peak | Size (bp) | Concentration (ng/uL) | From (bp) | To (bp) | Average size (bp) | CV% | RFU | Corrected peak area |
|---|---|---|---|---|---|---|---|---|
| 1 | 1 (LM) | 0.8786 | 0 | 8 | 1 | 71.44 | 11963 | 44.707 |
| 2 | 500 (UM) | 0.5000 | 493 | 512 | 500 | 0.25 | 12078 | 25.442 |

| | | |
|---|---|---|
| TIC: | 0.0000 | ng/uL |
| TIM: | 0.0000 | nmole/L |
| Total concentration: | 0.4880 | ng/uL |

Sample peak width (sec): 5 Sample min peak height: 50 Sample baseline V to V?: Y Sample baseline V to V points: 3
Sample filter: Binomial Number of points for filter: 3 Sample start region (min): 0 Sample end region (min): 80
Marker peak width (sec): 5 Marker min peak height: 500 Marker baseline V to V?: Y Marker baseline V to V points: 3
Lower marker selection: First peak > 500 RFU Upper marker selection: Last peak > 500 RFU
Ladder size (bp)1, 35, 75, 100, 150, 200, 250, 300, 400, 500
Quantification using: Upper Marker Final concentration (ng/uL): 0.5000 Dilution factor: 12.0

**Sample:** SampF5
**Well location:** F5
**Created:** Monday, 11 October 2021 14:41:52

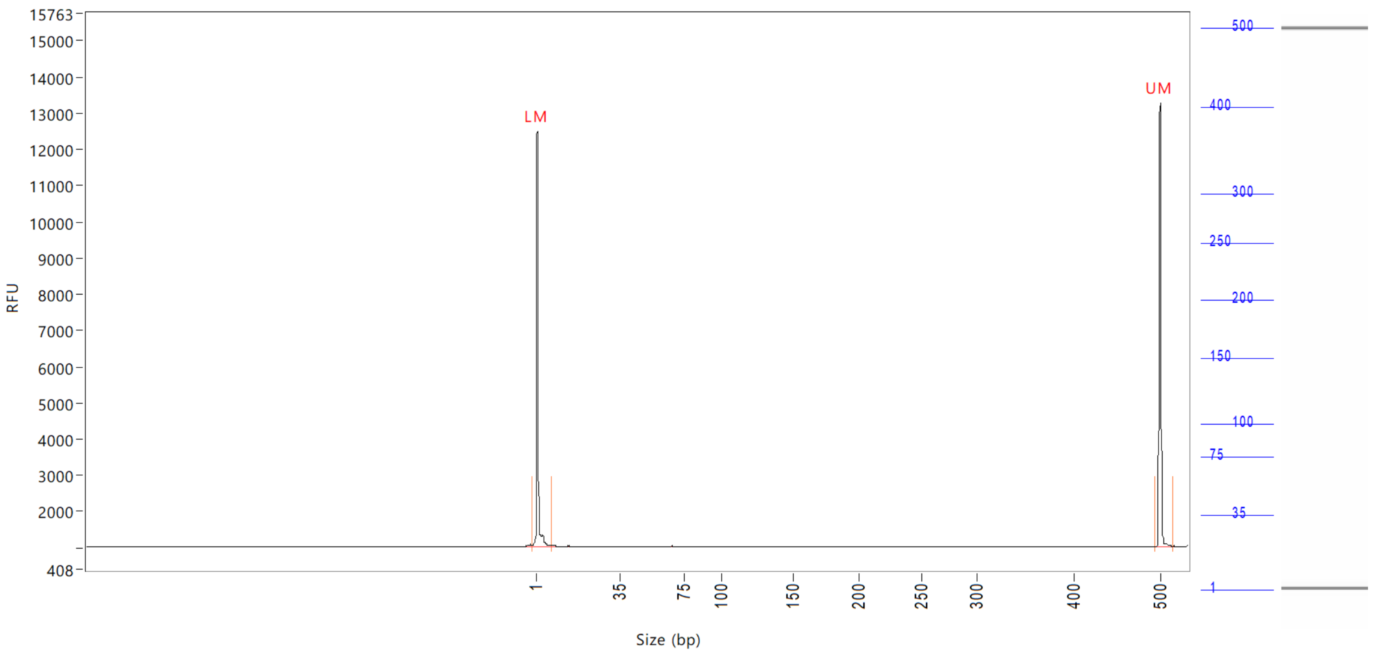


| Peak | Size (bp) | Concentration (ng/uL) | From (bp) | To (bp) | Average size (bp) | CV% | RFU | Corrected peak area |
|---|---|---|---|---|---|---|---|---|
| 1 | 1 (LM) | 0.8566 | 0 | 8 | 1 | 75.40 | 11466 | 44.057 |
| 2 | 500 (UM) | 0.5000 | 494 | 514 | 500 | 0.28 | 12275 | 25.717 |

| | | |
|---|---|---|
| TIC: | 0.0000 | ng/uL |
| TIM: | 0.0000 | nmole/L |
| Total concentration: | 0.4921 | ng/uL |

Sample peak width (sec): 5    Sample min peak height: 50    Sample baseline V to V?: Y    Sample baseline V to V points: 3
Sample filter: Binomial    Number of points for filter: 3  Sample start region (min): 0  Sample end region (min): 80
Marker peak width (sec): 5    Marker min peak height: 500    Marker baseline V to V?: Y    Marker baseline V to V points: 3
Lower marker selection: First peak > 500 RFU    Upper marker selection: Last peak > 500 RFU
Ladder size (bp)1, 35, 75, 100, 150, 200, 250, 300, 400, 500
Quantification using: Upper Marker    Final concentration (ng/uL): 0.5000    Dilution factor: 12.0

**Sample:** SampG5
**Well location:** G5
**Created:** Monday, 11 October 2021 14:41:52

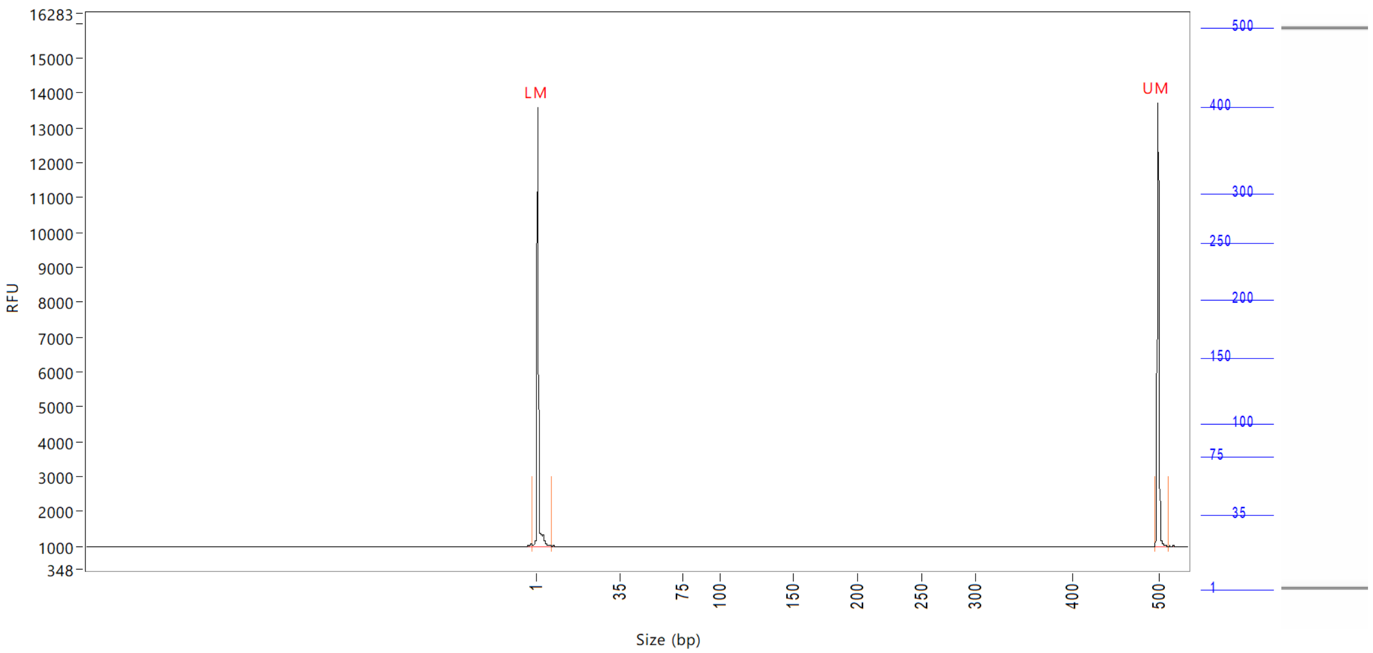


| Peak | Size (bp) | Concentration (ng/uL) | From (bp) | To (bp) | Average size (bp) | CV% | RFU | Corrected peak area |
|---|---|---|---|---|---|---|---|---|
| 1 | 1 (LM) | 0.9004 | 0 | 8 | 2 | 66.28 | 12560 | 47.605 |
| 2 | 500 (UM) | 0.5000 | 496 | 513 | 500 | 0.26 | 12743 | 26.436 |

| | | |
|---|---|---|
| TIC: | 0.0000 | ng/uL |
| TIM: | 0.0000 | nmole/L |
| Total concentration: | 0.4209 | ng/uL |

```
Sample peak width (sec): 5    Sample min peak height: 50    Sample baseline V to V?: Y    Sample baseline V to V points: 3
Sample filter: Binomial       Number of points for filter: 3  Sample start region (min): 0  Sample end region (min): 80
Marker peak width (sec): 5    Marker min peak height: 500   Marker baseline V to V?: Y    Marker baseline V to V points: 3
Lower marker selection: First peak > 500 RFU                Upper marker selection: Last peak > 500 RFU
Ladder size (bp)1, 35, 75, 100, 150, 200, 250, 300, 400, 500
Quantification using: Upper Marker          Final concentration (ng/uL): 0.5000          Dilution factor: 12.0
```

**Sample:** SampH5
**Well location:** H5
**Created:** Monday, 11 October 2021 14:41:52

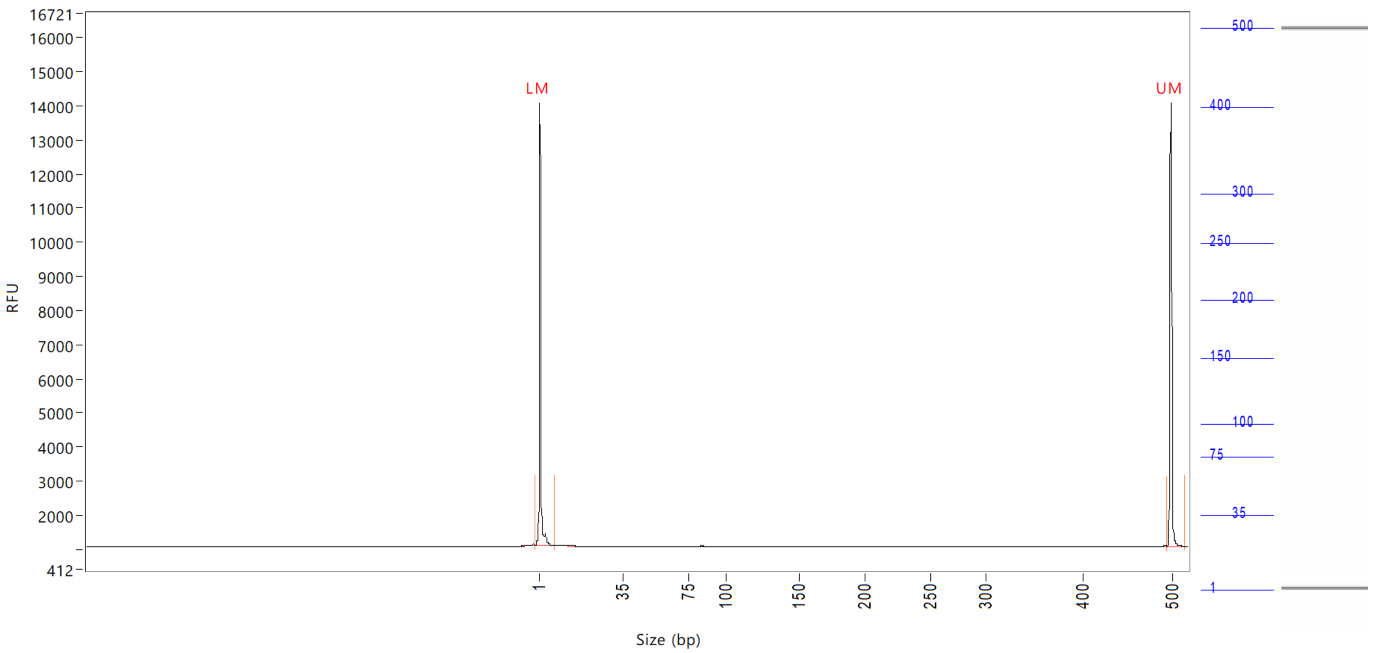


| Peak | Size (bp) | Concentration (ng/uL) | From (bp) | To (bp) | Average size (bp) | CV% | RFU | Corrected peak area |
|---|---|---|---|---|---|---|---|---|
| 1 | 1 (LM) | 0.9030 | 0 | 8 | 2 | 65.95 | 13004 | 49.554 |
| 2 | 500 (UM) | 0.5000 | 496 | 517 | 500 | 0.26 | 13027 | 27.437 |

| | | |
|---|---|---|
| TIC: | 0.0000 | ng/uL |
| TIM: | 0.0000 | nmole/L |
| Total concentration: | 0.4807 | ng/uL |

Sample peak width (sec): 5    Sample min peak height: 50    Sample baseline V to V?: Y    Sample baseline V to V points: 3
Sample filter: Binomial    Number of points for filter: 3    Sample start region (min): 0    Sample end region (min): 80
Marker peak width (sec): 5    Marker min peak height: 500    Marker baseline V to V?: Y    Marker baseline V to V points: 3
Lower marker selection: First peak > 500 RFU    Upper marker selection: Last peak > 500 RFU
Ladder size (bp)1, 35, 75, 100, 150, 200, 250, 300, 400, 500
Quantification using: Upper Marker    Final concentration (ng/uL): 0.5000    Dilution factor: 12.0

**Sample:** SampA6
**Well location:** A6
**Created:** Monday, 11 October 2021 14:41:52

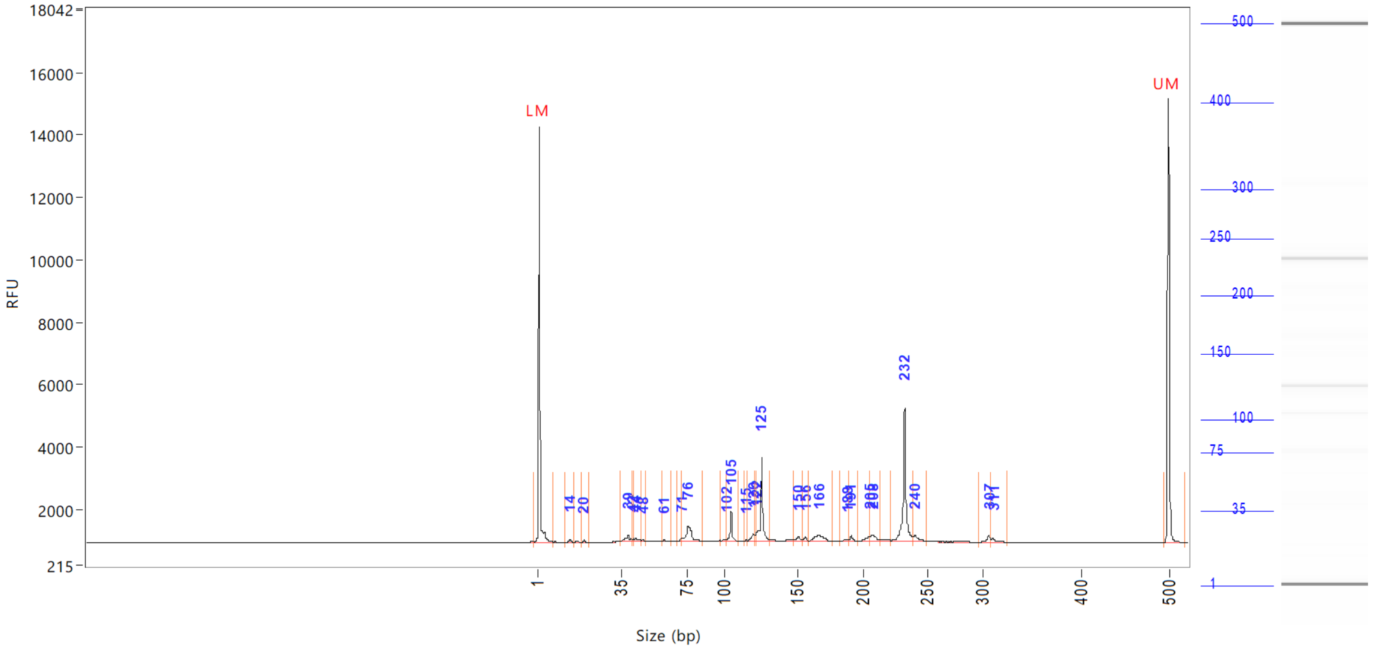


| Peak | Size<br>(bp) | Concentration<br>(ng/uL) | From<br>(bp) | To<br>(bp) | Average size<br>(bp) | CV% | RFU | Corrected peak area |
|---|---|---|---|---|---|---|---|---|
| 1 | 1 (LM) | 0.8062 | 0 | 7 | 1 | 68.13 | 13296 | 48.295 |
| 2 | 14 | 0.1037 | 13 | 16 | 14 | 3.95 | 107 | 0.518 |
| 3 | 20 | 0.0455 | 19 | 22 | 20 | 2.24 | 66 | 0.227 |
| 4 | 39 | 0.3647 | 34 | 41 | 38 | 4.37 | 183 | 1.821 |
| 5 | 42 | 0.0670 | 41 | 43 | 42 | 0.96 | 93 | 0.335 |
| 6 | 44 | 0.1900 | 43 | 47 | 45 | 2.82 | 113 | 0.948 |
| 7 | 48 | 0.0379 | 47 | 50 | 48 | 0.80 | 66 | 0.189 |
| 8 | 61 | 0.0606 | 60 | 65 | 61 | 1.34 | 82 | 0.303 |
| 9 | 71 | 0.1082 | 69 | 72 | 71 | 1.08 | 115 | 0.540 |
| 10 | 76 | 1.0227 | 72 | 85 | 76 | 3.01 | 513 | 5.106 |
| 11 | 102 | 0.0684 | 97 | 102 | 101 | 1.46 | 68 | 0.341 |
| 12 | 105 | 0.6594 | 102 | 110 | 105 | 1.03 | 953 | 3.292 |
| 13 | 115 | 0.0358 | 114 | 116 | 115 | 0.50 | 54 | 0.179 |
| 14 | 120 | 0.2518 | 116 | 121 | 119 | 0.92 | 234 | 1.257 |
| 15 | 122 | 0.1702 | 121 | 122 | 121 | 0.39 | 292 | 0.849 |
| 16 | 125 | 1.9627 | 122 | 131 | 125 | 1.07 | 2667 | 9.798 |
| 17 | 150 | 0.1855 | 147 | 154 | 150 | 1.33 | 144 | 0.926 |
| 18 | 156 | 0.0868 | 154 | 159 | 156 | 0.54 | 126 | 0.433 |
| 19 | 166 | 0.5775 | 159 | 176 | 167 | 1.85 | 201 | 2.883 |
| 20 | 188 | 0.0959 | 182 | 189 | 187 | 0.89 | 77 | 0.479 |
| 21 | 191 | 0.1300 | 189 | 196 | 191 | 0.60 | 174 | 0.649 |
| 22 | 205 | 0.2143 | 196 | 206 | 204 | 1.04 | 160 | 1.070 |
| 23 | 208 | 0.3013 | 206 | 213 | 209 | 0.78 | 198 | 1.504 |
| 24 | 232 | 3.2907 | 222 | 239 | 232 | 0.93 | 4280 | 16.428 |
| 25 | 240 | 0.2334 | 239 | 249 | 242 | 0.95 | 174 | 1.165 |

```
Sample peak width (sec): 5    Sample min peak height: 50    Sample baseline V to V?: Y    Sample baseline V to V points: 3
Sample filter: Binomial       Number of points for filter: 3  Sample start region (min): 0  Sample end region (min): 80
Marker peak width (sec): 5    Marker min peak height: 500   Marker baseline V to V?: Y    Marker baseline V to V points: 3
Lower marker selection: First peak > 500 RFU                Upper marker selection: Last peak > 500 RFU
Ladder size (bp)1, 35, 75, 100, 150, 200, 250, 300, 400, 500
Quantification using: Upper Marker          Final concentration (ng/uL): 0.5000          Dilution factor: 12.0
```

**Sample:** SampA6
**Well location:** A6
**Created:** Monday, 11 October 2021 14:41:52

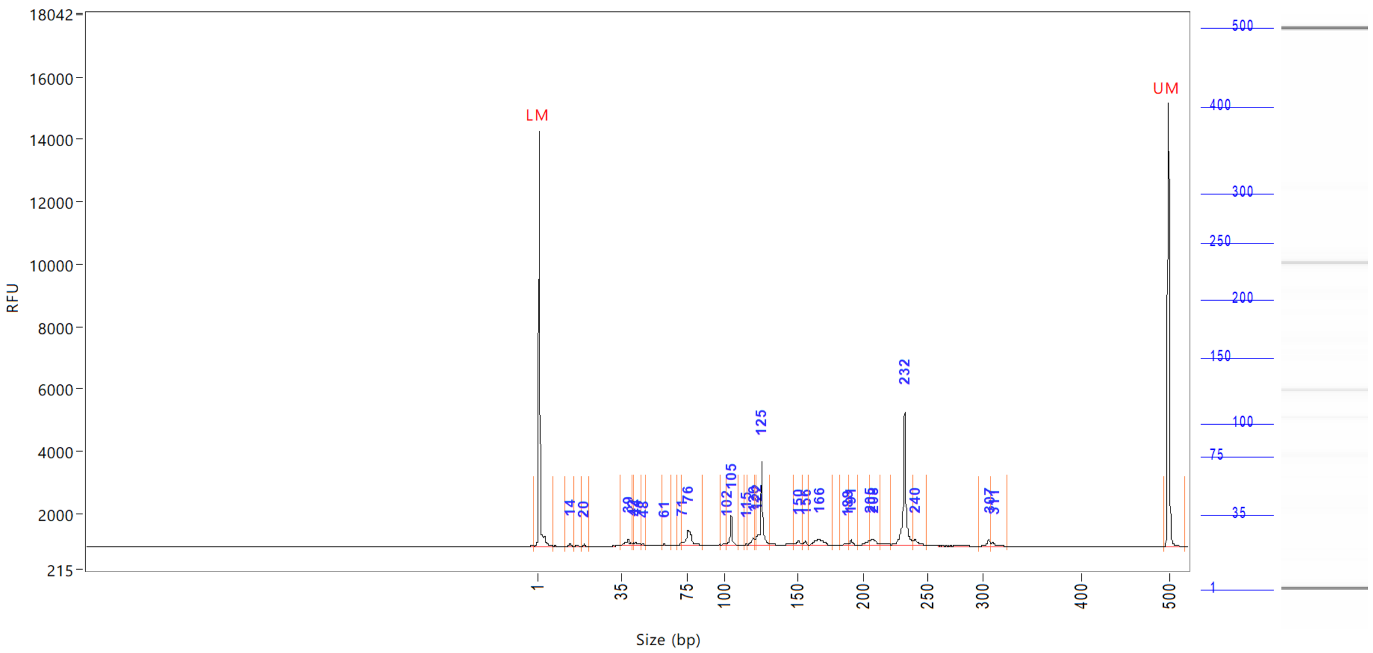


| Peak | Size (bp) | Concentration (ng/uL) | From (bp) | To (bp) | Average size (bp) | CV% | RFU | Corrected peak area |
|---|---|---|---|---|---|---|---|---|
| 26 | 307 | 0.1638 | 297 | 309 | 306 | 0.70 | 209 | |
| 27 | 311 | 0.1304 | 309 | 326 | 312 | 0.97 | 147 | |
| 28 | 500 (UM) | 0.5000 | 496 | 0 | 500 | 0.27 | 14244 | |

| | | |
|---|---|---|
| TIC: | 10.5581 | ng/uL |
| TIM: | 153.0741 | nmole/L |
| Total concentration: | 11.1972 | ng/uL |

Sample peak width (sec): 5    Sample min peak height: 50    Sample baseline V to V?: Y    Sample baseline V to V points: 3
Sample filter: Binomial    Number of points for filter: 3  Sample start region (min): 0  Sample end region (min): 80
Marker peak width (sec): 5    Marker min peak height: 500    Marker baseline V to V?: Y    Marker baseline V to V points: 3
Lower marker selection: First peak > 500 RFU    Upper marker selection: Last peak > 500 RFU
Ladder size (bp)1, 35, 75, 100, 150, 200, 250, 300, 400, 500
Quantification using: Upper Marker    Final concentration (ng/uL): 0.5000    Dilution factor: 12.0

**Sample:** SampB6
**Well location:** B6
**Created:** Monday, 11 October 2021 14:41:52

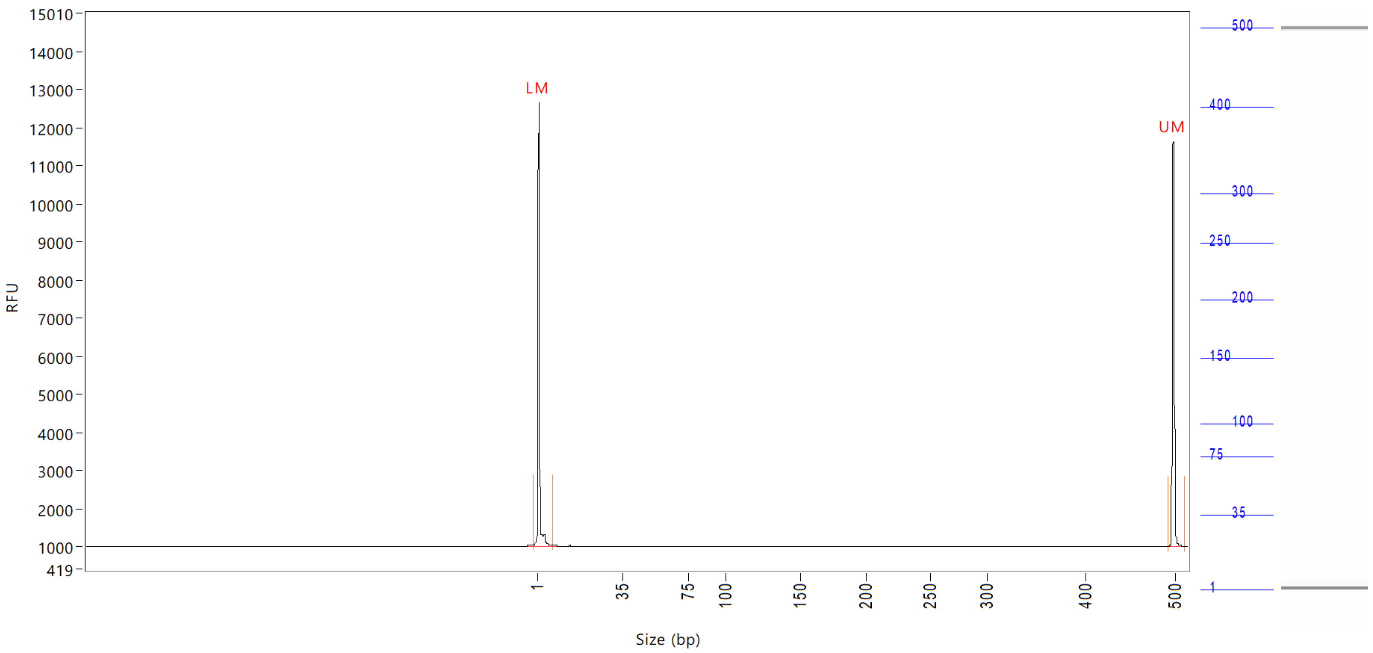


| Peak | Size (bp) | Concentration (ng/uL) | From (bp) | To (bp) | Average size (bp) | CV% | RFU | Corrected peak area |
|---|---|---|---|---|---|---|---|---|
| 1 | 1 (LM) | 0.9917 | 0 | 8 | 1 | 71.95 | 11653 | 44.275 |
| 2 | 500 (UM) | 0.5000 | 495 | 513 | 500 | 0.24 | 10636 | 22.322 |

| | | |
|---|---|---|
| TIC: | 0.0000 | ng/uL |
| TIM: | 0.0000 | nmole/L |
| Total concentration: | 0.5366 | ng/uL |

Sample peak width (sec): 5 Sample min peak height: 50 Sample baseline V to V?: Y Sample baseline V to V points: 3
Sample filter: Binomial Number of points for filter: 3 Sample start region (min): 0 Sample end region (min): 80
Marker peak width (sec): 5 Marker min peak height: 500 Marker baseline V to V?: Y Marker baseline V to V points: 3
Lower marker selection: First peak > 500 RFU Upper marker selection: Last peak > 500 RFU
Ladder size (bp)1, 35, 75, 100, 150, 200, 250, 300, 400, 500
Quantification using: Upper Marker Final concentration (ng/uL): 0.5000 Dilution factor: 12.0

**Sample:** SampC6
**Well location:** C6
**Created:** Monday, 11 October 2021 14:41:52

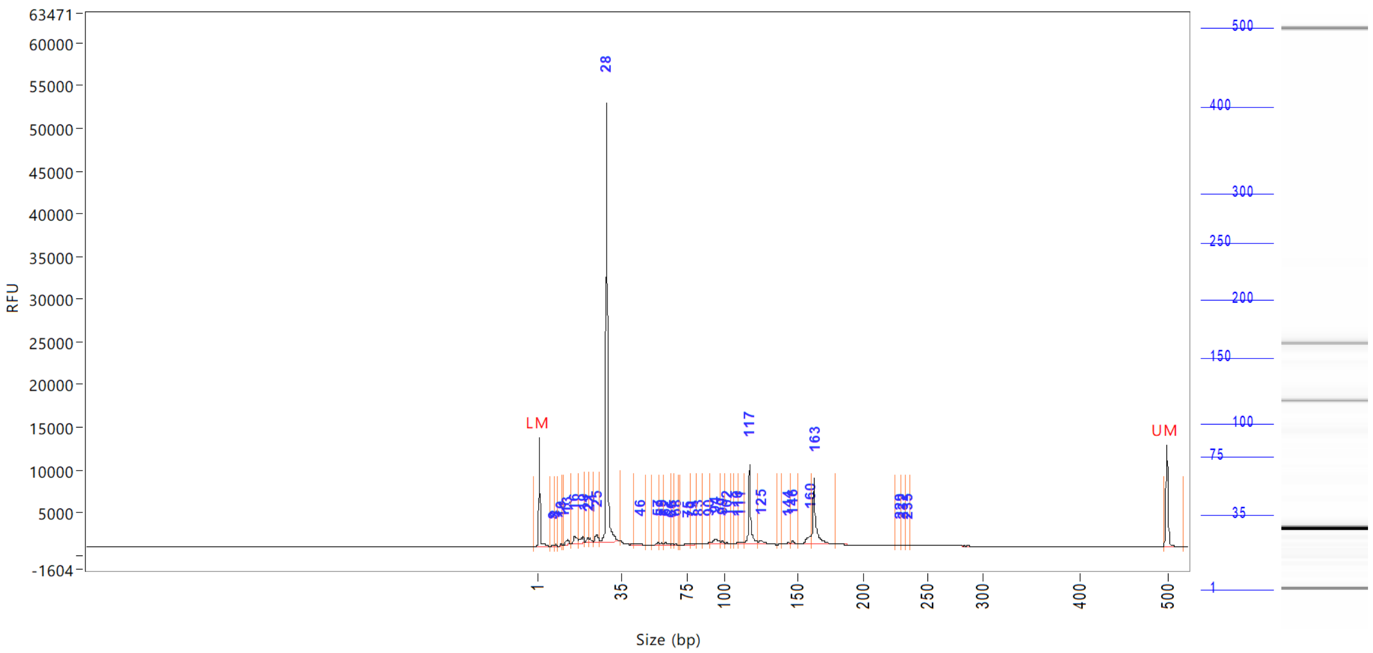


| Peak | Size (bp) | Concentration (ng/uL) | From (bp) | To (bp) | Average size (bp) | CV% | RFU | Corrected peak area |
|---|---|---|---|---|---|---|---|---|
| 1 | 1 (LM) | 0.8293 | 0 | 6 | 1 | 67.23 | 12762 | 47.076 |
| 2 | 8 | 0.1948 | 6 | 8 | 7 | 6.13 | 191 | 0.921 |
| 3 | 8 | 0.1755 | 8 | 9 | 8 | 2.81 | 229 | 0.830 |
| 4 | 10 | 0.0886 | 9 | 11 | 10 | 1.78 | 156 | 0.419 |
| 5 | 11 | 0.0507 | 11 | 12 | 11 | 1.57 | 107 | 0.240 |
| 6 | 13 | 0.9616 | 12 | 14 | 13 | 4.17 | 538 | 4.549 |
| 7 | 16 | 1.6285 | 14 | 17 | 16 | 4.00 | 838 | 7.703 |
| 8 | 19 | 1.4162 | 17 | 20 | 19 | 3.71 | 693 | 6.699 |
| 9 | 21 | 0.5781 | 20 | 22 | 21 | 1.69 | 626 | 2.734 |
| 10 | 22 | 0.4694 | 22 | 23 | 22 | 1.93 | 476 | 2.220 |
| 11 | 25 | 1.4238 | 23 | 26 | 25 | 2.46 | 900 | 6.735 |
| 12 | 28 | 43.4211 | 26 | 34 | 29 | 2.38 | 51445 | 205.392 |
| 13 | 46 | 0.3476 | 43 | 50 | 46 | 2.88 | 195 | 1.644 |
| 14 | 57 | 0.2781 | 54 | 58 | 57 | 1.71 | 227 | 1.315 |
| 15 | 59 | 0.2295 | 58 | 61 | 59 | 1.09 | 224 | 1.085 |
| 16 | 62 | 0.2719 | 61 | 65 | 62 | 1.67 | 266 | 1.286 |
| 17 | 66 | 0.0646 | 65 | 67 | 66 | 0.49 | 126 | 0.305 |
| 18 | 68 | 0.1091 | 67 | 70 | 68 | 0.53 | 203 | 0.516 |
| 19 | 75 | 0.2776 | 71 | 78 | 76 | 2.25 | 154 | 1.313 |
| 20 | 79 | 0.1164 | 78 | 82 | 80 | 1.32 | 84 | 0.550 |
| 21 | 83 | 0.1032 | 82 | 86 | 83 | 0.76 | 139 | 0.488 |
| 22 | 90 | 0.1948 | 86 | 91 | 90 | 1.12 | 192 | 0.922 |
| 23 | 94 | 1.4291 | 91 | 98 | 94 | 1.72 | 621 | 6.760 |
| 24 | 99 | 0.2973 | 98 | 101 | 99 | 0.74 | 357 | 1.406 |
| 25 | 102 | 0.1664 | 101 | 105 | 102 | 0.79 | 159 | 0.787 |

Sample peak width (sec): 5 Sample min peak height: 50 Sample baseline V to V?: Y Sample baseline V to V points: 3
Sample filter: Binomial Number of points for filter: 3 Sample start region (min): 0 Sample end region (min): 80
Marker peak width (sec): 5 Marker min peak height: 500 Marker baseline V to V?: Y Marker baseline V to V points: 3
Lower marker selection: First peak > 500 RFU Upper marker selection: Last peak > 500 RFU
Ladder size (bp)1, 35, 75, 100, 150, 200, 250, 300, 400, 500
Quantification using: Upper Marker Final concentration (ng/uL): 0.5000 Dilution factor: 12.0

**Sample:** SampC6
**Well location:** C6
**Created:** Monday, 11 October 2021 14:41:52

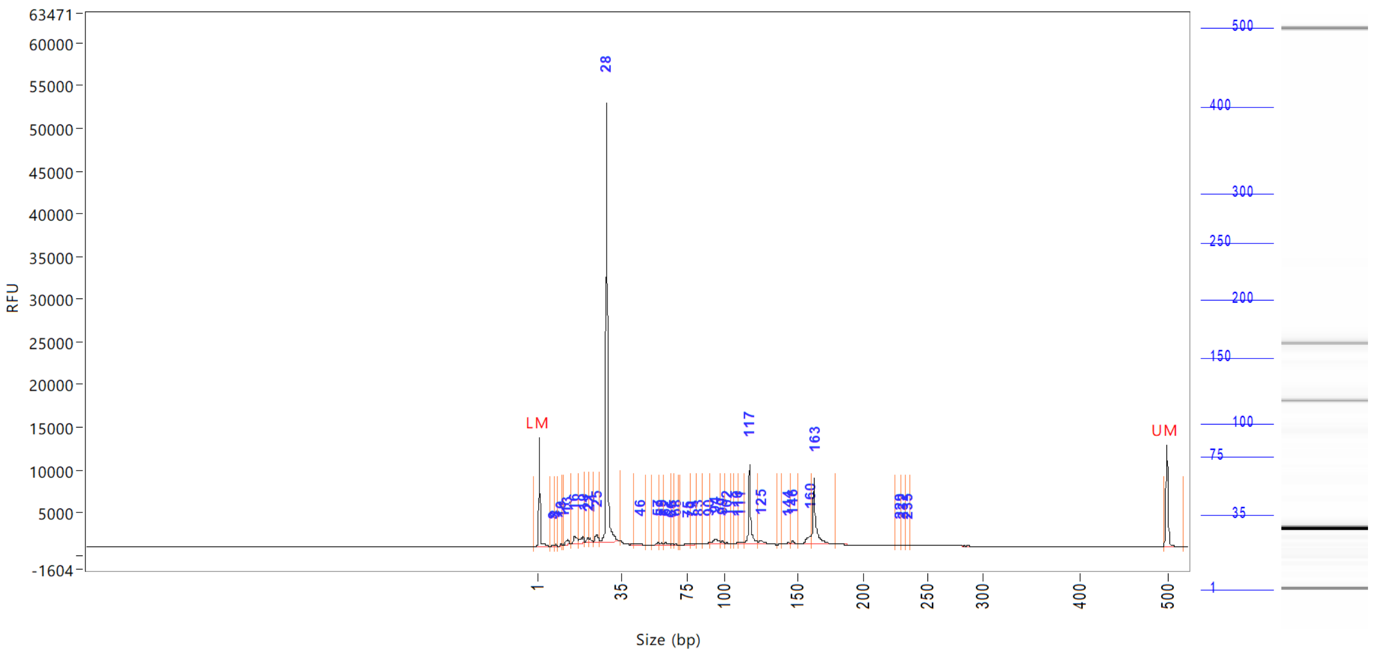


| Peak | Size (bp) | Concentration (ng/uL) | From (bp) | To (bp) | Average size (bp) | CV% | RFU | Corrected peak area |
|---|---|---|---|---|---|---|---|---|
| 26 | 110 | 0.1571 | 108 | 110 | 109 | 0.53 | 230 | |
| 27 | 111 | 0.4384 | 110 | 114 | 112 | 1.03 | 265 | |
| 28 | 117 | 7.4956 | 114 | 124 | 118 | 1.30 | 9401 | |
| 29 | 125 | 0.8978 | 124 | 136 | 126 | 1.64 | 437 | |
| 30 | 144 | 0.2703 | 139 | 145 | 143 | 0.88 | 184 | |
| 31 | 146 | 0.4027 | 145 | 151 | 147 | 0.86 | 423 | |
| 32 | 160 | 1.4341 | 151 | 161 | 158 | 1.13 | 989 | |
| 33 | 163 | 7.2926 | 161 | 180 | 164 | 1.63 | 7720 | |
| 34 | 229 | 0.0515 | 225 | 230 | 229 | 0.45 | 70 | |
| 35 | 232 | 0.0806 | 230 | 234 | 232 | 0.42 | 77 | |
| 36 | 235 | 0.0699 | 234 | 238 | 235 | 0.48 | 68 | |
| 37 | 500 (UM) | 0.5000 | 496 | 0 | 500 | 0.31 | 11861 | |

| | | |
|---|---|---|
| TIC: | 72.8844 | ng/uL |
| TIM: | 3428.1873 | nmole/L |
| Total concentration: | 73.0537 | ng/uL |

Sample peak width (sec): 5 Sample min peak height: 50 Sample baseline V to V?: Y Sample baseline V to V points: 3
Sample filter: Binomial Number of points for filter: 3 Sample start region (min): 0 Sample end region (min): 80
Marker peak width (sec): 5 Marker min peak height: 500 Marker baseline V to V?: Y Marker baseline V to V points: 3
Lower marker selection: First peak > 500 RFU Upper marker selection: Last peak > 500 RFU
Ladder size (bp)1, 35, 75, 100, 150, 200, 250, 300, 400, 500
Quantification using: Upper Marker Final concentration (ng/uL): 0.5000 Dilution factor: 12.0

**Sample:** SampD6
**Well location:** D6
**Created:** Monday, 11 October 2021 14:41:52

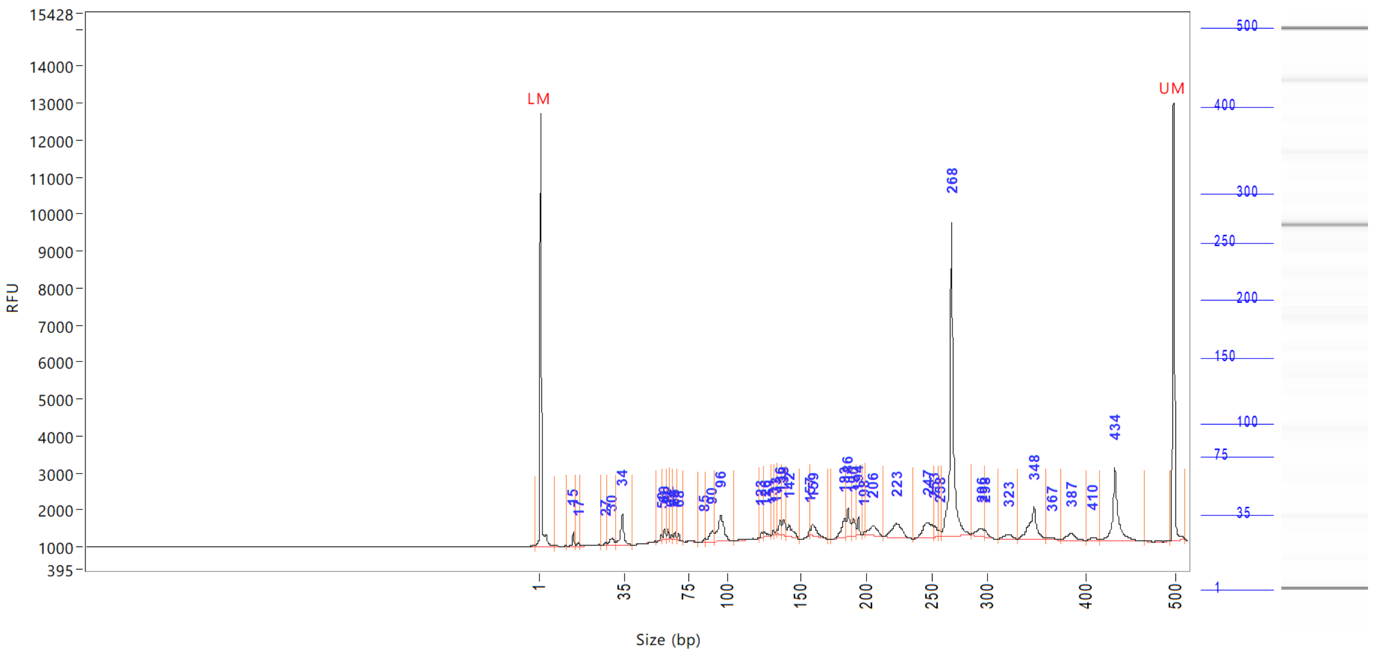


| Peak | Size<br>(bp) | Concentration<br>(ng/uL) | From<br>(bp) | To<br>(bp) | Average size<br>(bp) | CV% | RFU | Corrected peak area |
|---|---|---|---|---|---|---|---|---|
| 1 | 1 (LM) | 0.8476 | 0 | 7 | 2 | 66.78 | 11708 | 44.167 |
| 2 | 15 | 0.3398 | 13 | 16 | 15 | 2.58 | 399 | 1.476 |
| 3 | 17 | 0.1021 | 16 | 18 | 17 | 2.18 | 97 | 0.443 |
| 4 | 27 | 0.1022 | 26 | 28 | 27 | 1.78 | 85 | 0.444 |
| 5 | 30 | 0.5512 | 28 | 32 | 30 | 3.21 | 206 | 2.393 |
| 6 | 34 | 1.3008 | 32 | 40 | 34 | 3.41 | 857 | 5.648 |
| 7 | 58 | 0.1674 | 55 | 59 | 58 | 1.14 | 158 | 0.727 |
| 8 | 60 | 0.2573 | 59 | 61 | 60 | 0.87 | 303 | 1.117 |
| 9 | 62 | 0.1857 | 61 | 63 | 62 | 0.64 | 264 | 0.806 |
| 10 | 64 | 0.1043 | 63 | 65 | 64 | 0.81 | 125 | 0.453 |
| 11 | 66 | 0.1368 | 65 | 67 | 66 | 0.72 | 187 | 0.594 |
| 12 | 68 | 0.1365 | 67 | 72 | 68 | 0.67 | 200 | 0.593 |
| 13 | 85 | 0.1087 | 81 | 86 | 85 | 1.11 | 95 | 0.472 |
| 14 | 90 | 0.6343 | 86 | 92 | 90 | 1.57 | 309 | 2.754 |
| 15 | 96 | 1.7877 | 92 | 106 | 96 | 2.25 | 717 | 7.763 |
| 16 | 123 | 0.1339 | 122 | 125 | 124 | 0.49 | 169 | 0.582 |
| 17 | 126 | 0.1929 | 125 | 130 | 126 | 1.02 | 144 | 0.838 |
| 18 | 131 | 0.0863 | 130 | 132 | 131 | 0.37 | 136 | 0.375 |
| 19 | 133 | 0.0720 | 132 | 134 | 133 | 0.34 | 124 | 0.313 |
| 20 | 136 | 0.3802 | 134 | 137 | 136 | 0.53 | 414 | 1.651 |
| 21 | 138 | 0.5464 | 137 | 140 | 138 | 0.68 | 434 | 2.373 |
| 22 | 142 | 0.6300 | 140 | 149 | 143 | 1.27 | 327 | 2.735 |
| 23 | 157 | 0.1127 | 149 | 158 | 156 | 0.75 | 182 | 0.489 |
| 24 | 159 | 0.6058 | 158 | 171 | 161 | 1.34 | 307 | 2.631 |
| 25 | 183 | 0.8536 | 173 | 185 | 182 | 1.10 | 512 | 3.706 |

Sample peak width (sec): 5    Sample min peak height: 50    Sample baseline V to V?: Y    Sample baseline V to V points: 3
Sample filter: Binomial    Number of points for filter: 3    Sample start region (min): 0    Sample end region (min): 80
Marker peak width (sec): 5    Marker min peak height: 500    Marker baseline V to V?: Y    Marker baseline V to V points: 3
Lower marker selection: First peak > 500 RFU    Upper marker selection: Last peak > 500 RFU
Ladder size (bp)1, 35, 75, 100, 150, 200, 250, 300, 400, 500
Quantification using: Upper Marker    Final concentration (ng/uL): 0.5000    Dilution factor: 12.0

**Sample:** SampD6
**Well location:** D6
**Created:** Monday, 11 October 2021 14:41:52

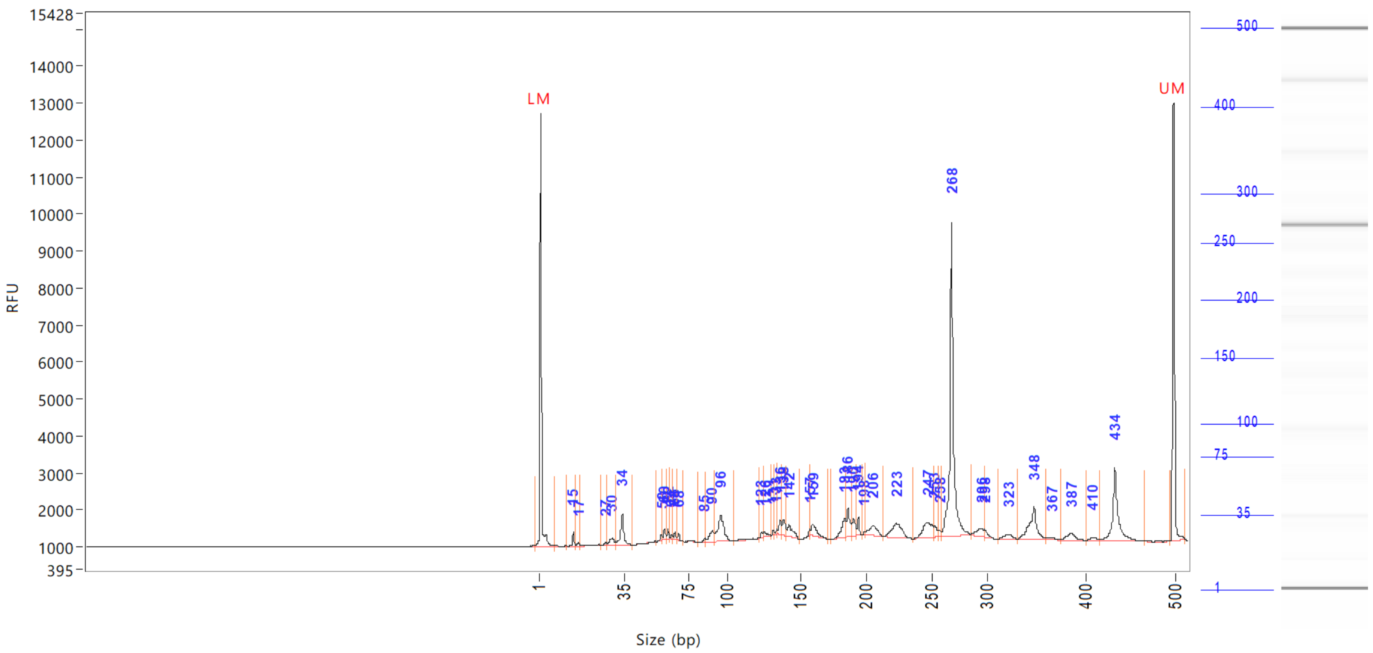


| Peak | Size (bp) | Concentration (ng/uL) | From (bp) | To (bp) | Average size (bp) | CV% | RFU | Corrected peak area |
|---|---|---|---|---|---|---|---|---|
| 26 | 186 | 0.7971 | 185 | 188 | 186 | 0.57 | 768 | |
| 27 | 190 | 0.5676 | 188 | 192 | 190 | 0.57 | 458 | |
| 28 | 194 | 0.3192 | 192 | 197 | 194 | 0.35 | 502 | |
| 29 | 198 | 0.0520 | 197 | 199 | 198 | 0.32 | 78 | |
| 30 | 206 | 0.7142 | 199 | 213 | 206 | 1.32 | 268 | |
| 31 | 223 | 1.3033 | 213 | 237 | 223 | 1.61 | 387 | |
| 32 | 247 | 1.2189 | 237 | 252 | 247 | 1.29 | 412 | |
| 33 | 253 | 0.3411 | 252 | 257 | 254 | 0.49 | 316 | |
| 34 | 258 | 0.1476 | 257 | 260 | 258 | 0.33 | 189 | |
| 35 | 268 | 8.8115 | 260 | 286 | 269 | 1.23 | 8469 | |
| 36 | 296 | 0.4975 | 286 | 298 | 293 | 1.04 | 212 | |
| 37 | 298 | 0.1899 | 298 | 312 | 300 | 0.76 | 196 | |
| 38 | 323 | 0.3019 | 312 | 331 | 322 | 1.18 | 131 | |
| 39 | 348 | 1.7974 | 331 | 361 | 346 | 1.46 | 888 | |
| 40 | 367 | 0.1409 | 361 | 375 | 366 | 0.87 | 68 | |
| 41 | 387 | 0.4256 | 375 | 402 | 386 | 1.17 | 179 | |
| 42 | 410 | 0.1586 | 402 | 417 | 411 | 0.82 | 92 | |
| 43 | 434 | 2.8003 | 417 | 467 | 436 | 1.61 | 1995 | |
| 44 | 500 (UM) | 0.5000 | 496 | 513 | 500 | 0.33 | 11864 | |

| | | |
|---|---|---|
| TIC: | 30.1134 | ng/uL |
| TIM: | 378.1637 | nmole/L |
| Total concentration: | 30.2948 | ng/uL |

```
Sample peak width (sec): 5    Sample min peak height: 50    Sample baseline V to V?: Y    Sample baseline V to V points: 3
Sample filter: Binomial       Number of points for filter: 3  Sample start region (min): 0  Sample end region (min): 80
Marker peak width (sec): 5    Marker min peak height: 500   Marker baseline V to V?: Y    Marker baseline V to V points: 3
Lower marker selection: First peak > 500 RFU                Upper marker selection: Last peak > 500 RFU
Ladder size (bp)1, 35, 75, 100, 150, 200, 250, 300, 400, 500
Quantification using: Upper Marker         Final concentration (ng/uL): 0.5000         Dilution factor: 12.0
```

**Sample:** SampE6
**Well location:** E6
**Created:** Monday, 11 October 2021 14:41:52

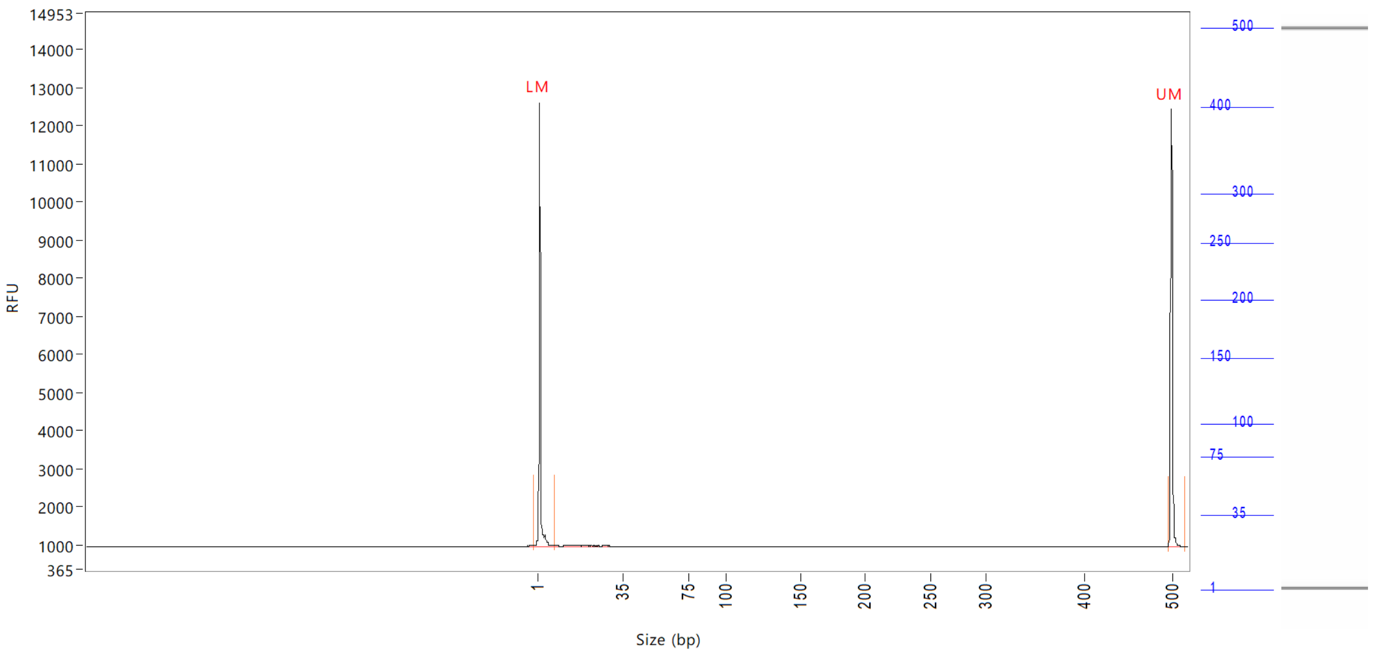


| Peak | Size (bp) | Concentration (ng/uL) | From (bp) | To (bp) | Average size (bp) | CV% | RFU | Corrected peak area |
|---|---|---|---|---|---|---|---|---|
| 1 | 1 (LM) | 0.8834 | 0 | 8 | 1 | 67.23 | 11649 | 43.538 |
| 2 | 500 (UM) | 0.5000 | 496 | 516 | 500 | 0.26 | 11491 | 24.642 |

| | | |
|---|---|---|
| TIC: | 0.0000 | ng/uL |
| TIM: | 0.0000 | nmole/L |
| Total concentration: | 0.4725 | ng/uL |

```
Sample peak width (sec): 5    Sample min peak height: 50    Sample baseline V to V?: Y    Sample baseline V to V points: 3
Sample filter: Binomial       Number of points for filter: 3  Sample start region (min): 0  Sample end region (min): 80
Marker peak width (sec): 5    Marker min peak height: 500   Marker baseline V to V?: Y    Marker baseline V to V points: 3
Lower marker selection: First peak > 500 RFU                Upper marker selection: Last peak > 500 RFU
Ladder size (bp)1, 35, 75, 100, 150, 200, 250, 300, 400, 500
Quantification using: Upper Marker          Final concentration (ng/uL): 0.5000          Dilution factor: 12.0
```

**Sample:** SampF6
**Well location:** F6
**Created:** Monday, 11 October 2021 14:41:52

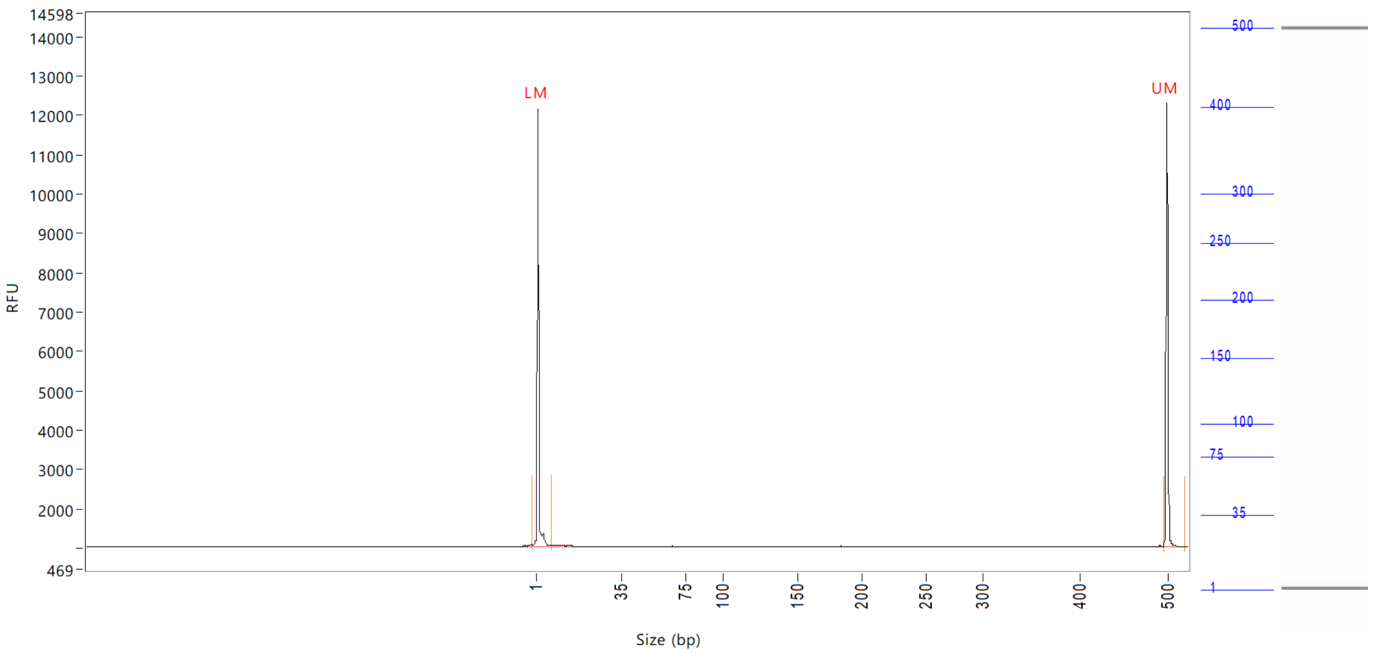


| Peak | Size<br>(bp) | Concentration<br>(ng/uL) | From<br>(bp) | To<br>(bp) | Average size<br>(bp) | CV% | RFU | Corrected peak area |
|---|---|---|---|---|---|---|---|---|
| 1 | 1 (LM) | 0.9110 | 0 | 8 | 1 | 72.72 | 11128 | 42.200 |
| 2 | 500 (UM) | 0.5000 | 496 | 0 | 500 | 0.27 | 11295 | 23.160 |

| | | |
|---|---|---|
| TIC: | 0.0000 | ng/uL |
| TIM: | 0.0000 | nmole/L |
| Total concentration: | 0.5935 | ng/uL |

```
Sample peak width (sec): 5    Sample min peak height: 50    Sample baseline V to V?: Y    Sample baseline V to V points: 3
Sample filter: Binomial       Number of points for filter: 3  Sample start region (min): 0  Sample end region (min): 80
Marker peak width (sec): 5    Marker min peak height: 500   Marker baseline V to V?: Y    Marker baseline V to V points: 3
Lower marker selection: First peak > 500 RFU                Upper marker selection: Last peak > 500 RFU
Ladder size (bp)1, 35, 75, 100, 150, 200, 250, 300, 400, 500
Quantification using: Upper Marker          Final concentration (ng/uL): 0.5000          Dilution factor: 12.0
```

**Sample:** SampG6
**Well location:** G6
**Created:** Monday, 11 October 2021 14:41:52

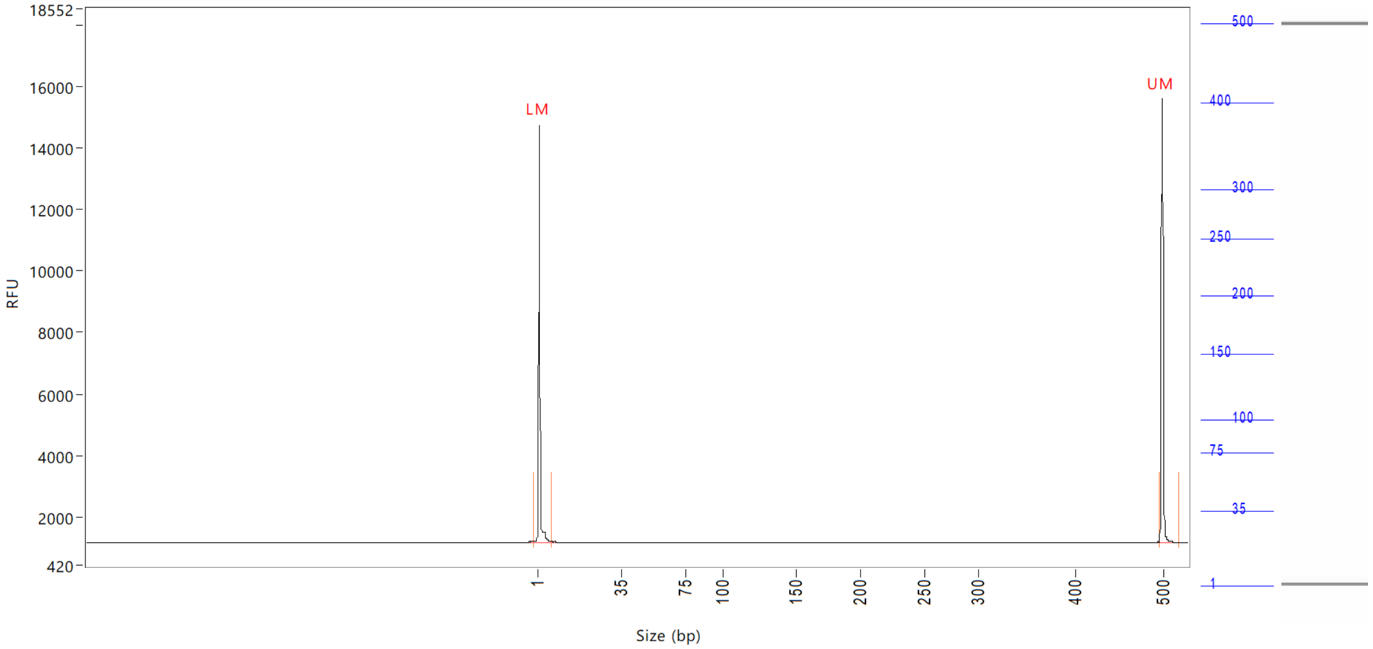


| Peak | Size (bp) | Concentration (ng/uL) | From (bp) | To (bp) | Average size (bp) | CV% | RFU | Corrected peak area |
|---|---|---|---|---|---|---|---|---|
| 1 | 1 (LM) | 0.7969 | 0 | 7 | 2 | 66.53 | 13604 | 48.924 |
| 2 | 500 (UM) | 0.5000 | 496 | 0 | 500 | 0.29 | 14502 | 30.696 |

| | | |
|---|---|---|
| TIC: | 0.0000 | ng/uL |
| TIM: | 0.0000 | nmole/L |
| Total concentration: | 0.4427 | ng/uL |

Sample peak width (sec): 5 Sample min peak height: 50 Sample baseline V to V?: Y Sample baseline V to V points: 3
Sample filter: Binomial Number of points for filter: 3 Sample start region (min): 0 Sample end region (min): 80
Marker peak width (sec): 5 Marker min peak height: 500 Marker baseline V to V?: Y Marker baseline V to V points: 3
Lower marker selection: First peak > 500 RFU Upper marker selection: Last peak > 500 RFU
Ladder size (bp)1, 35, 75, 100, 150, 200, 250, 300, 400, 500
Quantification using: Upper Marker Final concentration (ng/uL): 0.5000 Dilution factor: 12.0

**Sample:** SampH6
**Well location:** H6
**Created:** Monday, 11 October 2021 14:41:52

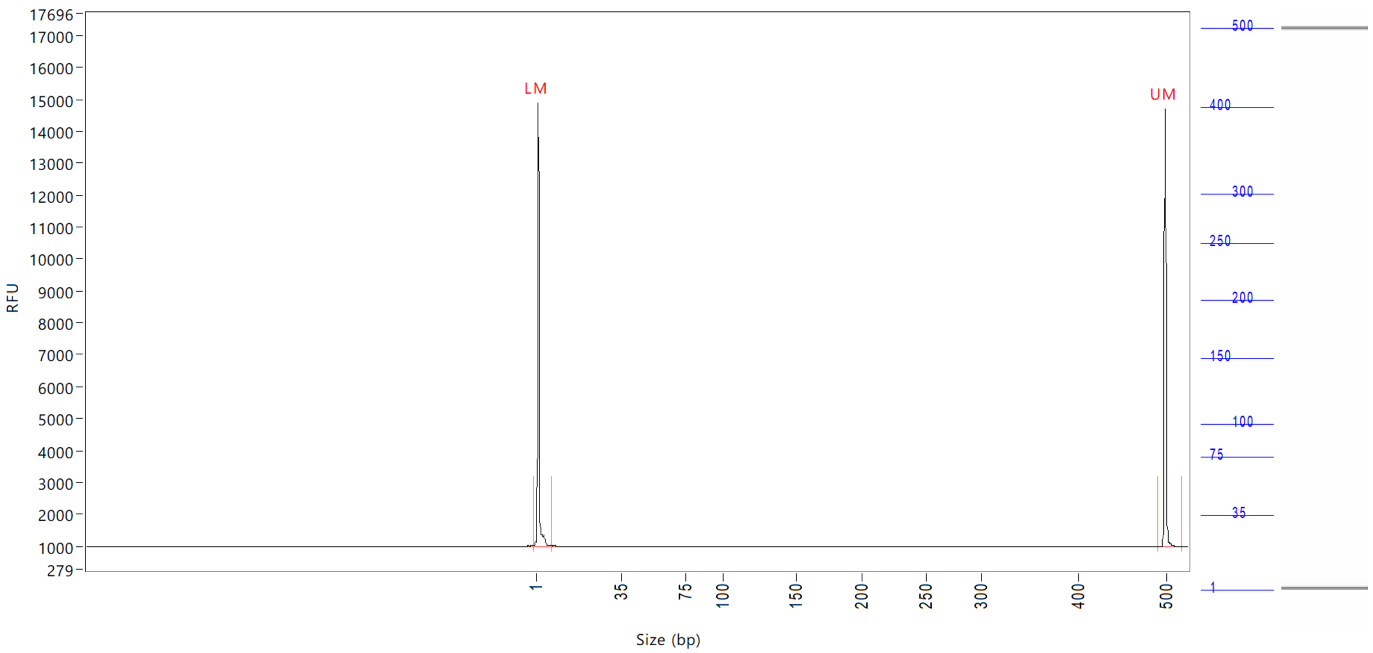


| Peak | Size (bp) | Concentration (ng/uL) | From (bp) | To (bp) | Average size (bp) | CV% | RFU | Corrected peak area |
|---|---|---|---|---|---|---|---|---|
| 1 | 1 (LM) | 0.9370 | 0 | 8 | 1 | 67.90 | 13916 | 52.534 |
| 2 | 500 (UM) | 0.5000 | 493 | 0 | 500 | 0.29 | 13722 | 28.034 |

| | | |
|---|---|---|
| TIC: | 0.0000 | ng/uL |
| TIM: | 0.0000 | nmole/L |
| Total concentration: | 0.4414 | ng/uL |

Sample peak width (sec): 5    Sample min peak height: 50    Sample baseline V to V?: Y    Sample baseline V to V points: 3
Sample filter: Binomial    Number of points for filter: 3    Sample start region (min): 0    Sample end region (min): 80
Marker peak width (sec): 5    Marker min peak height: 500    Marker baseline V to V?: Y    Marker baseline V to V points: 3
Lower marker selection: First peak > 500 RFU    Upper marker selection: Last peak > 500 RFU
Ladder size (bp)1, 35, 75, 100, 150, 200, 250, 300, 400, 500
Quantification using: Upper Marker    Final concentration (ng/uL): 0.5000    Dilution factor: 12.0

**Sample:** SampA7
**Well location:** A7
**Created:** Monday, 11 October 2021 14:41:52

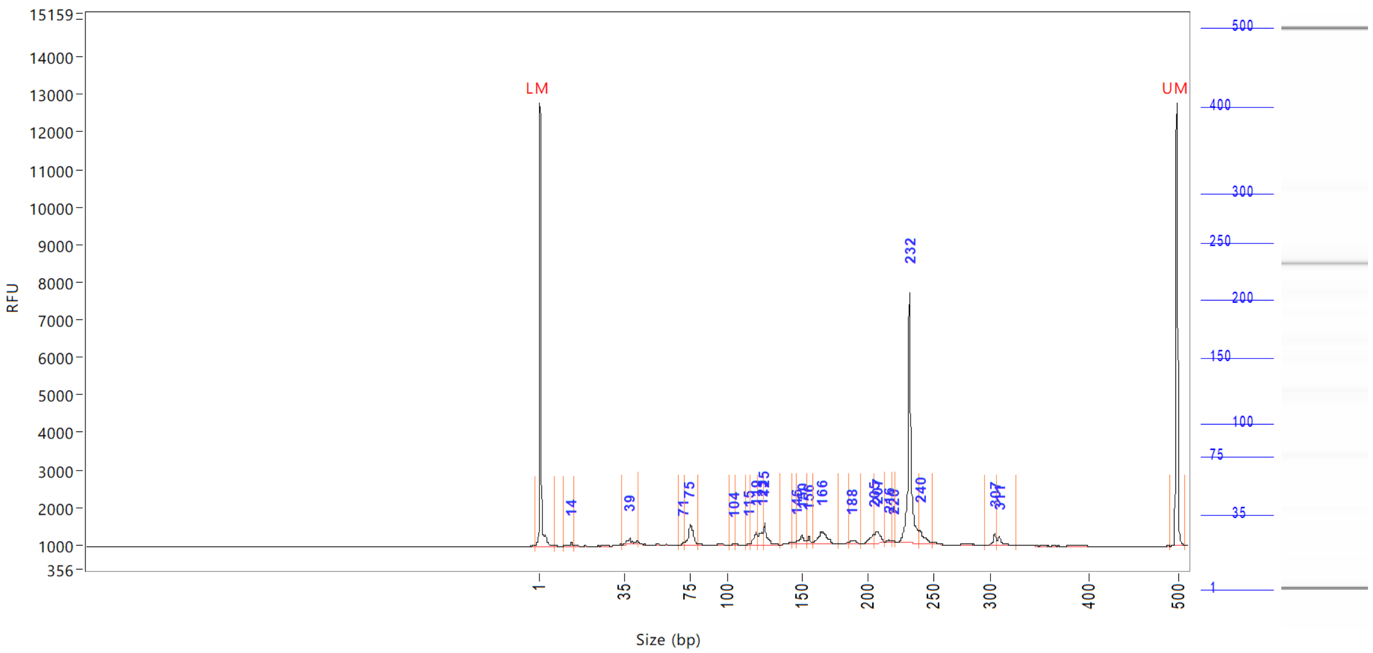


| Peak | Size (bp) | Concentration (ng/uL) | From (bp) | To (bp) | Average size (bp) | CV% | RFU | Corrected peak area |
|---|---|---|---|---|---|---|---|---|
| 1 | 1 (LM) | 0.8955 | 0 | 8 | 1 | 66.00 | 11810 | 44.218 |
| 2 | 14 | 0.1304 | 12 | 15 | 14 | 5.62 | 97 | 0.537 |
| 3 | 39 | 0.4005 | 34 | 44 | 39 | 5.43 | 168 | 1.648 |
| 4 | 71 | 0.0938 | 68 | 71 | 70 | 1.10 | 91 | 0.386 |
| 5 | 75 | 1.2931 | 71 | 81 | 76 | 2.67 | 551 | 5.321 |
| 6 | 104 | 0.0500 | 102 | 106 | 104 | 0.80 | 50 | 0.206 |
| 7 | 115 | 0.0489 | 113 | 116 | 115 | 0.53 | 57 | 0.201 |
| 8 | 119 | 0.5371 | 116 | 120 | 119 | 1.02 | 363 | 2.210 |
| 9 | 123 | 0.6222 | 120 | 124 | 122 | 0.83 | 334 | 2.560 |
| 10 | 125 | 0.7720 | 124 | 136 | 126 | 1.48 | 583 | 3.177 |
| 11 | 146 | 0.0864 | 143 | 147 | 145 | 0.71 | 69 | 0.356 |
| 12 | 150 | 0.3664 | 147 | 154 | 150 | 1.37 | 234 | 1.508 |
| 13 | 156 | 0.1583 | 154 | 158 | 156 | 0.55 | 191 | 0.651 |
| 14 | 166 | 1.0510 | 158 | 178 | 166 | 1.89 | 317 | 4.325 |
| 15 | 188 | 0.2037 | 186 | 194 | 189 | 0.98 | 92 | 0.838 |
| 16 | 205 | 0.4392 | 194 | 206 | 203 | 1.14 | 256 | 1.807 |
| 17 | 207 | 0.5719 | 206 | 213 | 208 | 0.81 | 316 | 2.353 |
| 18 | 216 | 0.0903 | 213 | 219 | 216 | 0.64 | 66 | 0.372 |
| 19 | 220 | 0.0357 | 219 | 222 | 220 | 0.28 | 52 | 0.147 |
| 20 | 232 | 6.7948 | 222 | 239 | 232 | 0.96 | 6661 | 27.960 |
| 21 | 240 | 0.6441 | 239 | 249 | 242 | 1.04 | 361 | 2.650 |
| 22 | 307 | 0.3118 | 296 | 309 | 306 | 0.79 | 311 | 1.283 |
| 23 | 311 | 0.3118 | 309 | 328 | 313 | 1.15 | 248 | 1.283 |
| 24 | 500 (UM) | 0.5000 | 493 | 510 | 500 | 0.23 | 11810 | 24.690 |

Sample peak width (sec): 5 Sample min peak height: 50 Sample baseline V to V?: Y Sample baseline V to V points: 3
Sample filter: Binomial Number of points for filter: 3 Sample start region (min): 0 Sample end region (min): 80
Marker peak width (sec): 5 Marker min peak height: 500 Marker baseline V to V?: Y Marker baseline V to V points: 3
Lower marker selection: First peak > 500 RFU Upper marker selection: Last peak > 500 RFU
Ladder size (bp)1, 35, 75, 100, 150, 200, 250, 300, 400, 500
Quantification using: Upper Marker Final concentration (ng/uL): 0.5000 Dilution factor: 12.0

**Sample:** SampA7
**Well location:** A7
**Created:** Monday, 11 October 2021 14:41:52

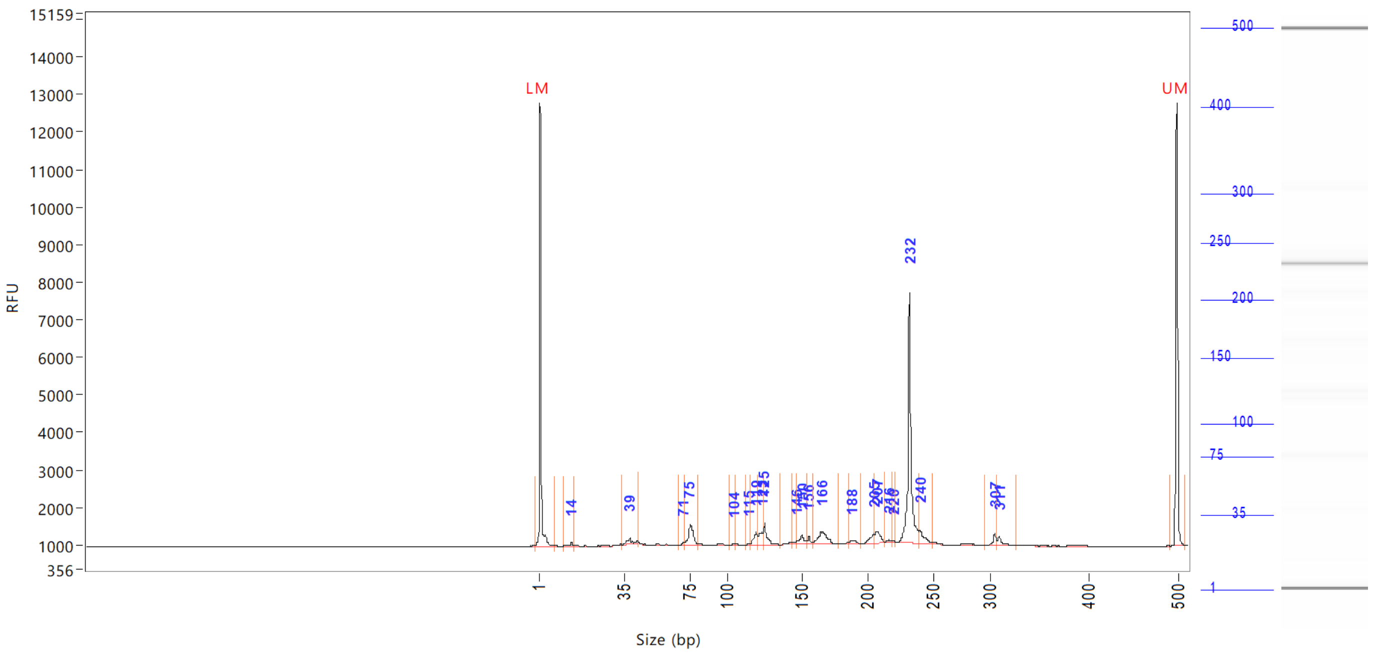


| Peak | Size (bp) | Concentration (ng/uL) | From (bp) | To (bp) | Average size (bp) | CV% | RFU | Corrected peak area |
|---|---|---|---|---|---|---|---|---|

| | | |
|---|---|---|
| TIC: | 15.0134 | ng/uL |
| TIM: | 173.0449 | nmole/L |
| Total concentration: | 15.9652 | ng/uL |

```
Sample peak width (sec): 5    Sample min peak height: 50    Sample baseline V to V?: Y    Sample baseline V to V points: 3
Sample filter: Binomial       Number of points for filter: 3  Sample start region (min): 0  Sample end region (min): 80
Marker peak width (sec): 5    Marker min peak height: 500   Marker baseline V to V?: Y    Marker baseline V to V points: 3
Lower marker selection: First peak > 500 RFU                Upper marker selection: Last peak > 500 RFU
Ladder size (bp)1, 35, 75, 100, 150, 200, 250, 300, 400, 500
Quantification using: Upper Marker          Final concentration (ng/uL): 0.5000          Dilution factor: 12.0
```

**Sample:** SampB7
**Well location:** B7
**Created:** Monday, 11 October 2021 14:41:52

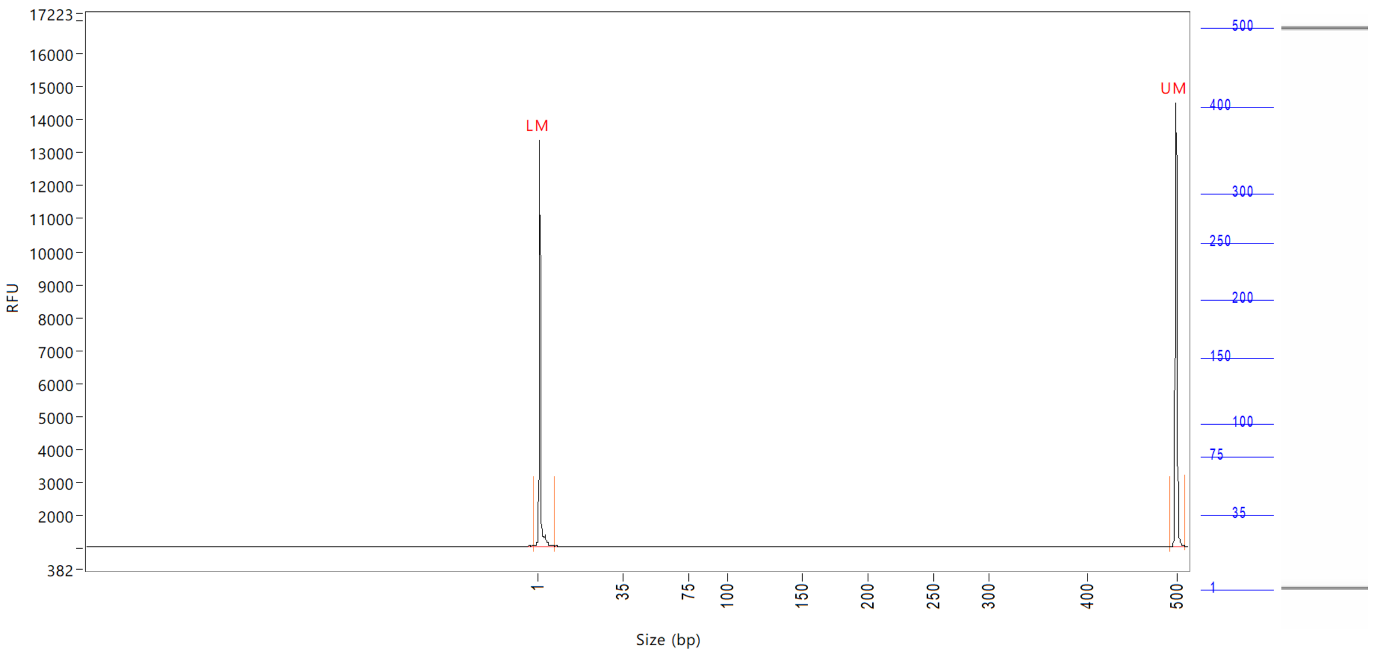


| Peak | Size | Concentration | From | To | Average size | CV% | RFU | Corrected peak area |
|---|---|---|---|---|---|---|---|---|
| | (bp) | (ng/uL) | (bp) | (bp) | (bp) | | | |
| 1 | 1 (LM) | 0.8354 | 0 | 8 | 2 | 65.50 | 12288 | 47.134 |
| 2 | 500 (UM) | 0.5000 | 494 | 510 | 500 | 0.24 | 13458 | 28.211 |

| | | |
|---|---|---|
| TIC: | 0.0000 | ng/uL |
| TIM: | 0.0000 | nmole/L |
| Total concentration: | 0.4940 | ng/uL |

```
Sample peak width (sec): 5    Sample min peak height: 50    Sample baseline V to V?: Y    Sample baseline V to V points: 3
Sample filter: Binomial       Number of points for filter: 3  Sample start region (min): 0  Sample end region (min): 80
Marker peak width (sec): 5    Marker min peak height: 500   Marker baseline V to V?: Y    Marker baseline V to V points: 3
Lower marker selection: First peak > 500 RFU                Upper marker selection: Last peak > 500 RFU
Ladder size (bp)1, 35, 75, 100, 150, 200, 250, 300, 400, 500
Quantification using: Upper Marker          Final concentration (ng/uL): 0.5000          Dilution factor: 12.0
```

**Sample:** SampC7
**Well location:** C7
**Created:** Monday, 11 October 2021 14:41:52

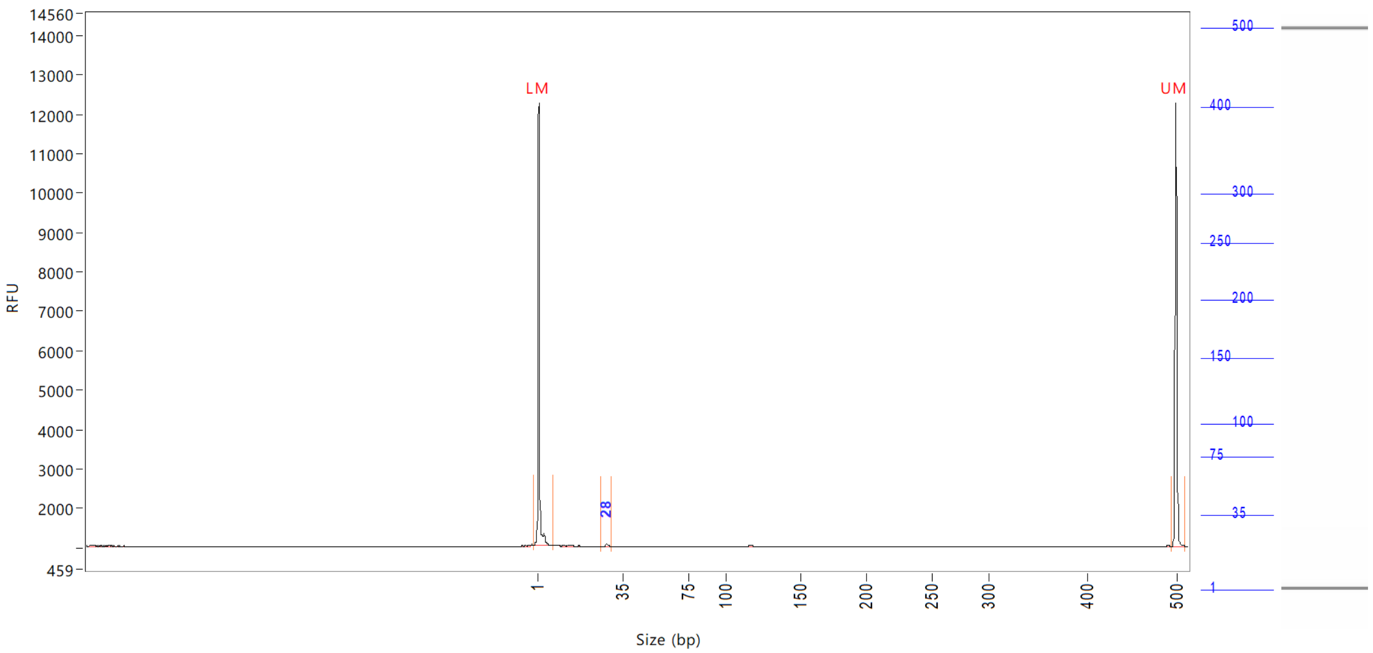


| Peak | Size (bp) | Concentration (ng/uL) | From (bp) | To (bp) | Average size (bp) | CV% | RFU | Corrected peak area |
|---|---|---|---|---|---|---|---|---|
| 1 | 1 (LM) | 0.9231 | 0 | 8 | 1 | 70.36 | 11254 | 42.617 |
| 2 | 28 | 0.0895 | 26 | 30 | 28 | 1.99 | 78 | 0.344 |
| 3 | 500 (UM) | 0.5000 | 494 | 511 | 500 | 0.23 | 11260 | 23.083 |

| | | |
|---|---|---|
| TIC: | 0.0895 | ng/uL |
| TIM: | 5.2136 | nmole/L |
| Total concentration: | 0.6455 | ng/uL |

Sample peak width (sec): 5 Sample min peak height: 50 Sample baseline V to V?: Y Sample baseline V to V points: 3
Sample filter: Binomial Number of points for filter: 3 Sample start region (min): 0 Sample end region (min): 80
Marker peak width (sec): 5 Marker min peak height: 500 Marker baseline V to V?: Y Marker baseline V to V points: 3
Lower marker selection: First peak > 500 RFU Upper marker selection: Last peak > 500 RFU
Ladder size (bp)1, 35, 75, 100, 150, 200, 250, 300, 400, 500
Quantification using: Upper Marker Final concentration (ng/uL): 0.5000 Dilution factor: 12.0

**Sample:** SampD7
**Well location:** D7
**Created:** Monday, 11 October 2021 14:41:52

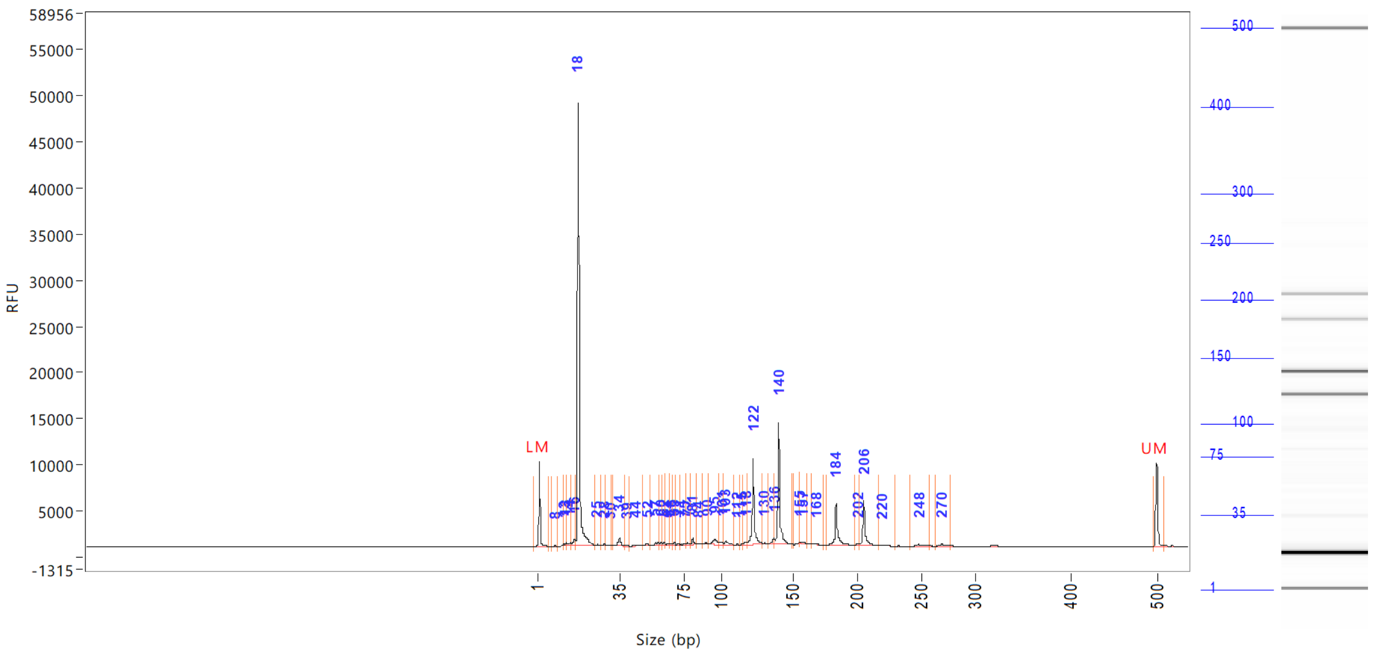

| Peak | Size (bp) | Concentration (ng/uL) | From (bp) | To (bp) | Average size (bp) | CV% | RFU | Corrected peak area |
|---|---|---|---|---|---|---|---|---|
| 1 | 1 (LM) | 0.8427 | 0 | 6 | 1 | 63.05 | 9156 | 33.965 |
| 2 | 8 | 0.1071 | 7 | 9 | 8 | 4.59 | 77 | 0.360 |
| 3 | 12 | 0.1583 | 9 | 12 | 11 | 5.26 | 145 | 0.532 |
| 4 | 13 | 0.2226 | 12 | 13 | 13 | 1.99 | 239 | 0.748 |
| 5 | 14 | 0.4847 | 13 | 15 | 14 | 3.38 | 274 | 1.628 |
| 6 | 16 | 1.0447 | 15 | 17 | 16 | 2.41 | 620 | 3.509 |
| 7 | 18 | 63.1048 | 17 | 25 | 18 | 4.81 | 47996 | 211.955 |
| 8 | 25 | 0.1482 | 25 | 27 | 26 | 2.00 | 81 | 0.498 |
| 9 | 28 | 0.2269 | 27 | 29 | 28 | 1.23 | 194 | 0.762 |
| 10 | 30 | 0.1456 | 29 | 31 | 30 | 1.55 | 81 | 0.489 |
| 11 | 34 | 1.5313 | 32 | 38 | 34 | 3.02 | 843 | 5.143 |
| 12 | 39 | 0.1944 | 38 | 41 | 39 | 1.65 | 120 | 0.653 |
| 13 | 44 | 0.3496 | 41 | 49 | 45 | 3.14 | 149 | 1.174 |
| 14 | 52 | 0.3514 | 49 | 54 | 52 | 1.86 | 204 | 1.180 |
| 15 | 57 | 0.6131 | 54 | 59 | 57 | 2.03 | 304 | 2.059 |
| 16 | 60 | 0.3277 | 59 | 61 | 60 | 0.80 | 315 | 1.101 |
| 17 | 62 | 0.2044 | 61 | 63 | 62 | 0.69 | 222 | 0.686 |
| 18 | 64 | 0.1305 | 63 | 65 | 64 | 0.79 | 130 | 0.438 |
| 19 | 66 | 0.1915 | 65 | 68 | 67 | 0.88 | 175 | 0.643 |
| 20 | 69 | 0.1951 | 68 | 69 | 68 | 0.52 | 234 | 0.655 |
| 21 | 71 | 0.2174 | 69 | 72 | 71 | 0.68 | 201 | 0.730 |
| 22 | 75 | 0.2293 | 72 | 76 | 74 | 1.63 | 143 | 0.770 |
| 23 | 77 | 0.2968 | 76 | 80 | 78 | 1.20 | 185 | 0.997 |
| 24 | 81 | 0.6865 | 80 | 84 | 81 | 0.87 | 682 | 2.306 |
| 25 | 84 | 0.0857 | 84 | 87 | 85 | 1.14 | 68 | 0.288 |

```
Sample peak width (sec): 5     Sample min peak height: 50    Sample baseline V to V?: Y    Sample baseline V to V points: 3
Sample filter: Binomial        Number of points for filter: 3  Sample start region (min): 0  Sample end region (min): 80
Marker peak width (sec): 5     Marker min peak height: 500   Marker baseline V to V?: Y    Marker baseline V to V points: 3
Lower marker selection: First peak > 500 RFU                 Upper marker selection: Last peak > 500 RFU
Ladder size (bp)1, 35, 75, 100, 150, 200, 250, 300, 400, 500
Quantification using: Upper Marker          Final concentration (ng/uL): 0.5000          Dilution factor: 12.0
```

**Sample:** SampD7
**Well location:** D7
**Created:** Monday, 11 October 2021 14:41:52

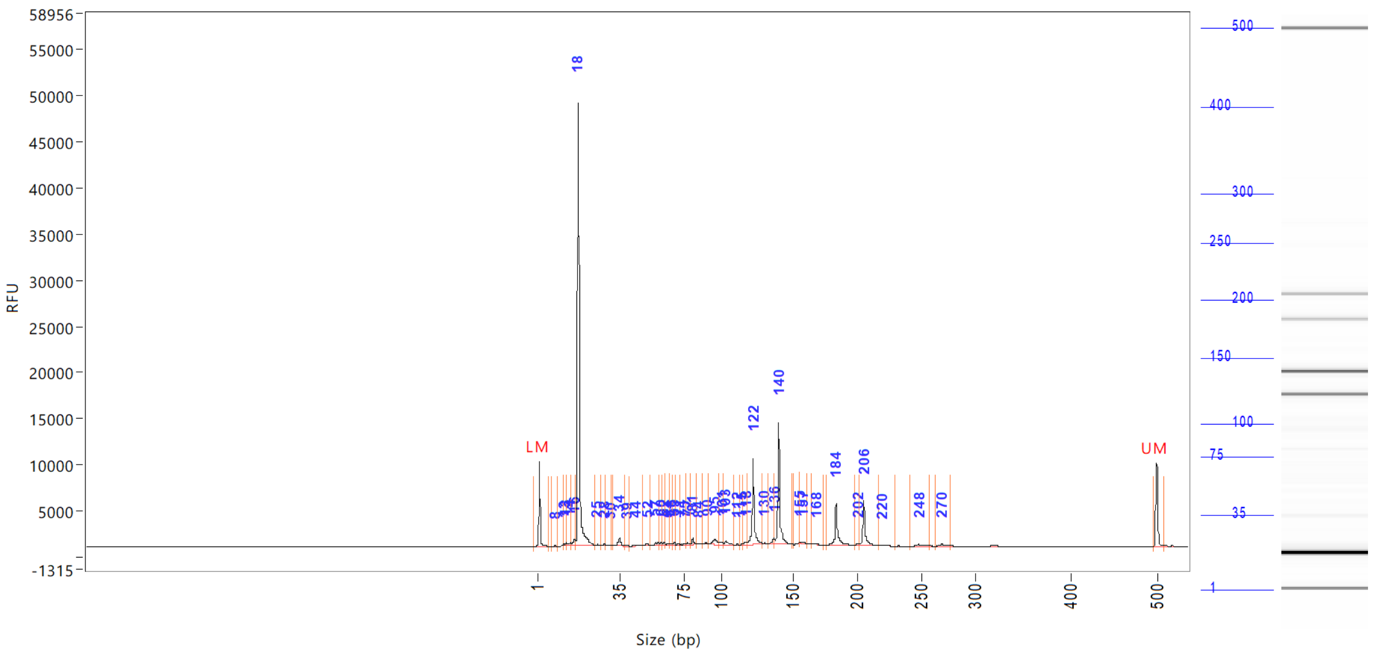


| Peak | Size (bp) | Concentration (ng/uL) | From (bp) | To (bp) | Average size (bp) | CV% | RFU | Corrected peak area |
|---|---|---|---|---|---|---|---|---|
| 26 | 90 | 0.3512 | 87 | 92 | 90 | 1.12 | 193 | |
| 27 | 95 | 1.5309 | 92 | 99 | 95 | 1.64 | 578 | |
| 28 | 101 | 0.4241 | 99 | 102 | 100 | 1.15 | 241 | |
| 29 | 103 | 0.8009 | 102 | 110 | 104 | 1.51 | 349 | |
| 30 | 112 | 0.0669 | 110 | 113 | 111 | 0.44 | 85 | |
| 31 | 115 | 0.0754 | 113 | 115 | 114 | 0.41 | 89 | |
| 32 | 118 | 0.5198 | 115 | 119 | 117 | 0.76 | 310 | |
| 33 | 122 | 9.9250 | 119 | 129 | 122 | 1.18 | 9387 | |
| 34 | 130 | 0.1745 | 129 | 132 | 130 | 0.62 | 159 | |
| 35 | 136 | 0.8427 | 132 | 137 | 136 | 0.71 | 601 | |
| 36 | 140 | 12.3715 | 137 | 149 | 140 | 1.01 | 13134 | |
| 37 | 155 | 0.0587 | 150 | 156 | 154 | 0.50 | 93 | |
| 38 | 157 | 0.2239 | 156 | 161 | 158 | 0.68 | 152 | |
| 39 | 168 | 0.1324 | 164 | 174 | 167 | 1.00 | 71 | |
| 40 | 184 | 5.3930 | 176 | 198 | 184 | 1.47 | 4458 | |
| 41 | 202 | 0.2588 | 198 | 203 | 201 | 0.65 | 209 | |
| 42 | 206 | 4.9234 | 203 | 218 | 207 | 1.03 | 4811 | |
| 43 | 220 | 0.1078 | 218 | 231 | 221 | 0.99 | 60 | |
| 44 | 248 | 0.3449 | 242 | 259 | 249 | 1.29 | 181 | |
| 45 | 270 | 0.3015 | 264 | 277 | 271 | 0.96 | 201 | |
| 46 | 500 (UM) | 0.5000 | 496 | 509 | 500 | 0.26 | 9087 | |

| | | |
|---|---|---|
| TIC: | 110.0749 | ng/uL |
| TIM: | 6587.8340 | nmole/L |
| Total concentration: | 110.2051 | ng/uL |

```
Sample peak width (sec): 5     Sample min peak height: 50     Sample baseline V to V?: Y     Sample baseline V to V points: 3
Sample filter: Binomial        Number of points for filter: 3  Sample start region (min): 0  Sample end region (min): 80
Marker peak width (sec): 5     Marker min peak height: 500    Marker baseline V to V?: Y     Marker baseline V to V points: 3
Lower marker selection: First peak > 500 RFU                  Upper marker selection: Last peak > 500 RFU
Ladder size (bp)1, 35, 75, 100, 150, 200, 250, 300, 400, 500
Quantification using: Upper Marker          Final concentration (ng/uL): 0.5000          Dilution factor: 12.0
```

**Sample:** SampE7
**Well location:** E7
**Created:** Monday, 11 October 2021 14:41:52

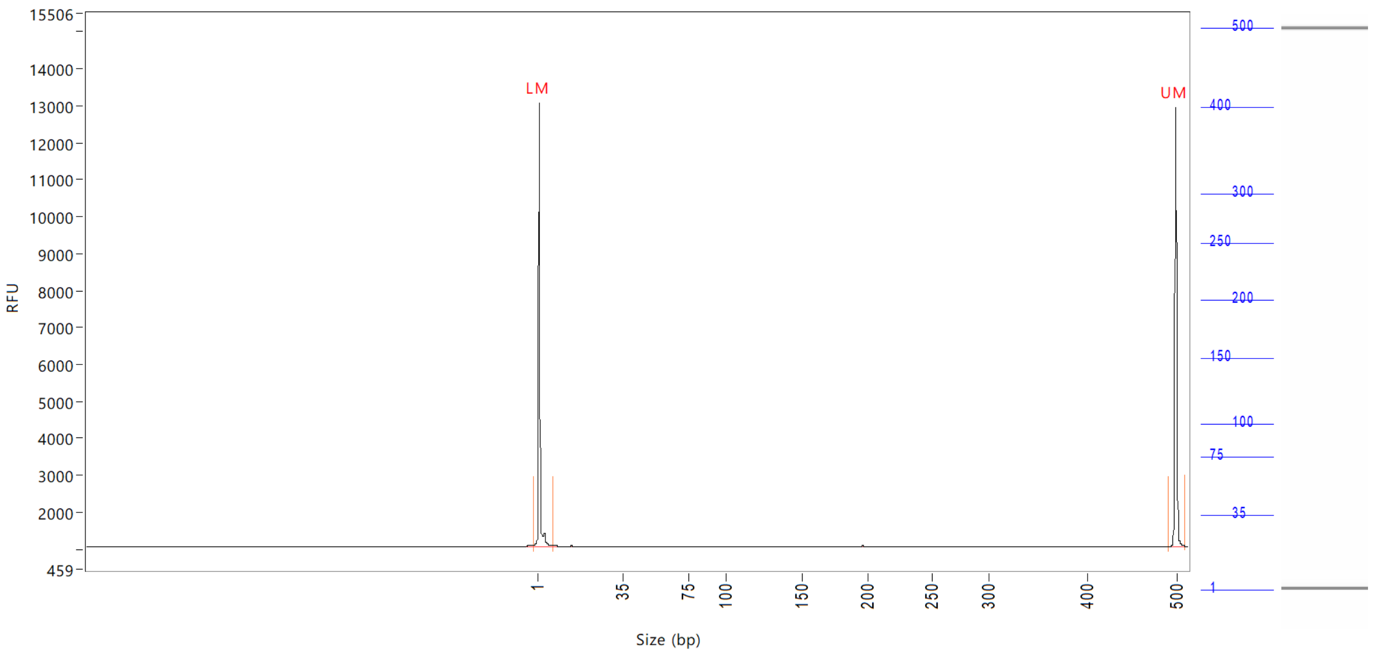


| Peak | Size (bp) | Concentration (ng/uL) | From (bp) | To (bp) | Average size (bp) | CV% | RFU | Corrected peak area |
|---|---|---|---|---|---|---|---|---|
| 1 | 1 (LM) | 0.8995 | 0 | 8 | 1 | 73.02 | 12017 | 45.519 |
| 2 | 500 (UM) | 0.5000 | 492 | 511 | 500 | 0.24 | 11898 | 25.302 |

| | | |
|---|---|---|
| TIC: | 0.0000 | ng/uL |
| TIM: | 0.0000 | nmole/L |
| Total concentration: | 0.5009 | ng/uL |

Sample peak width (sec): 5 Sample min peak height: 50 Sample baseline V to V?: Y Sample baseline V to V points: 3
Sample filter: Binomial Number of points for filter: 3 Sample start region (min): 0 Sample end region (min): 80
Marker peak width (sec): 5 Marker min peak height: 500 Marker baseline V to V?: Y Marker baseline V to V points: 3
Lower marker selection: First peak > 500 RFU Upper marker selection: Last peak > 500 RFU
Ladder size (bp)1, 35, 75, 100, 150, 200, 250, 300, 400, 500
Quantification using: Upper Marker Final concentration (ng/uL): 0.5000 Dilution factor: 12.0

**Sample:** SampF7
**Well location:** F7
**Created:** Monday, 11 October 2021 14:41:52

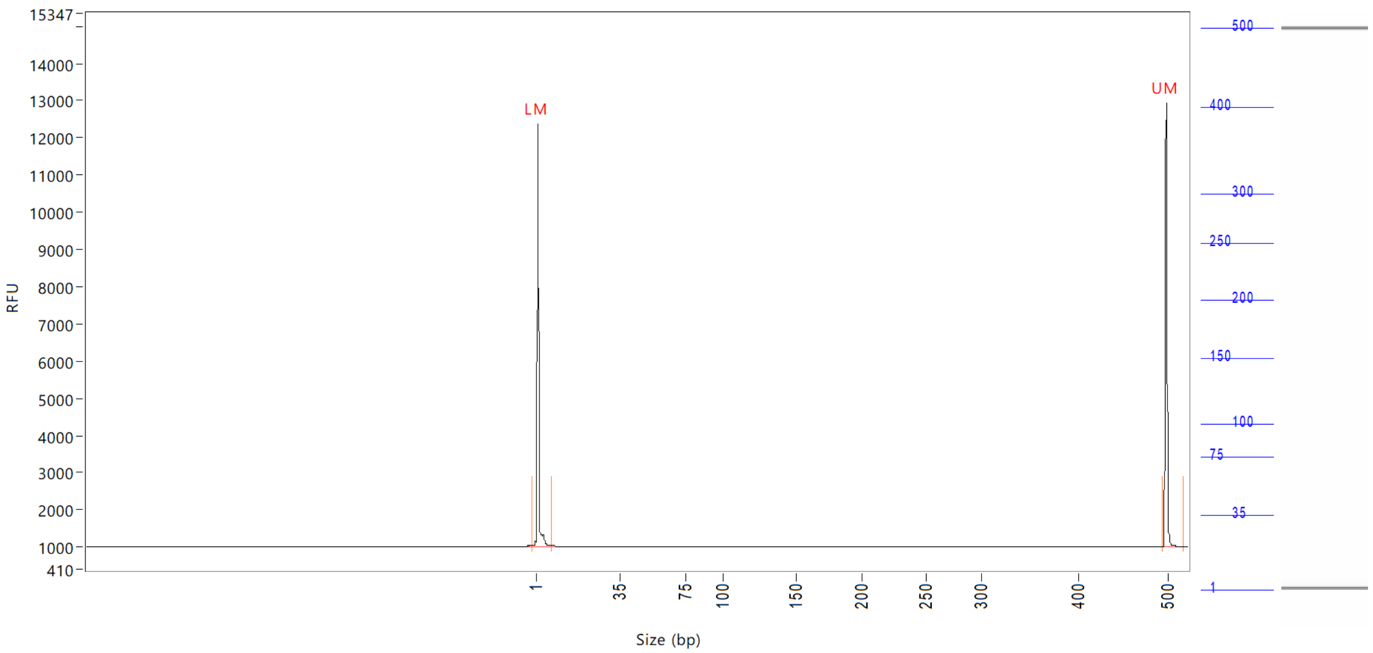


| Peak | Size (bp) | Concentration (ng/uL) | From (bp) | To (bp) | Average size (bp) | CV% | RFU | Corrected peak area |
|---|---|---|---|---|---|---|---|---|
| 1 | 1 (LM) | 0.8739 | 0 | 8 | 2 | 65.33 | 11364 | 43.413 |
| 2 | 500 (UM) | 0.5000 | 496 | 0 | 500 | 0.27 | 11943 | 24.840 |

| | | |
|---|---|---|
| TIC: | 0.0000 | ng/uL |
| TIM: | 0.0000 | nmole/L |
| Total concentration: | 0.5254 | ng/uL |

Sample peak width (sec): 5 Sample min peak height: 50 Sample baseline V to V?: Y Sample baseline V to V points: 3
Sample filter: Binomial Number of points for filter: 3 Sample start region (min): 0 Sample end region (min): 80
Marker peak width (sec): 5 Marker min peak height: 500 Marker baseline V to V?: Y Marker baseline V to V points: 3
Lower marker selection: First peak > 500 RFU Upper marker selection: Last peak > 500 RFU
Ladder size (bp)1, 35, 75, 100, 150, 200, 250, 300, 400, 500
Quantification using: Upper Marker Final concentration (ng/uL): 0.5000 Dilution factor: 12.0

**Sample:** SampG7
**Well location:** G7
**Created:** Monday, 11 October 2021 14:41:52

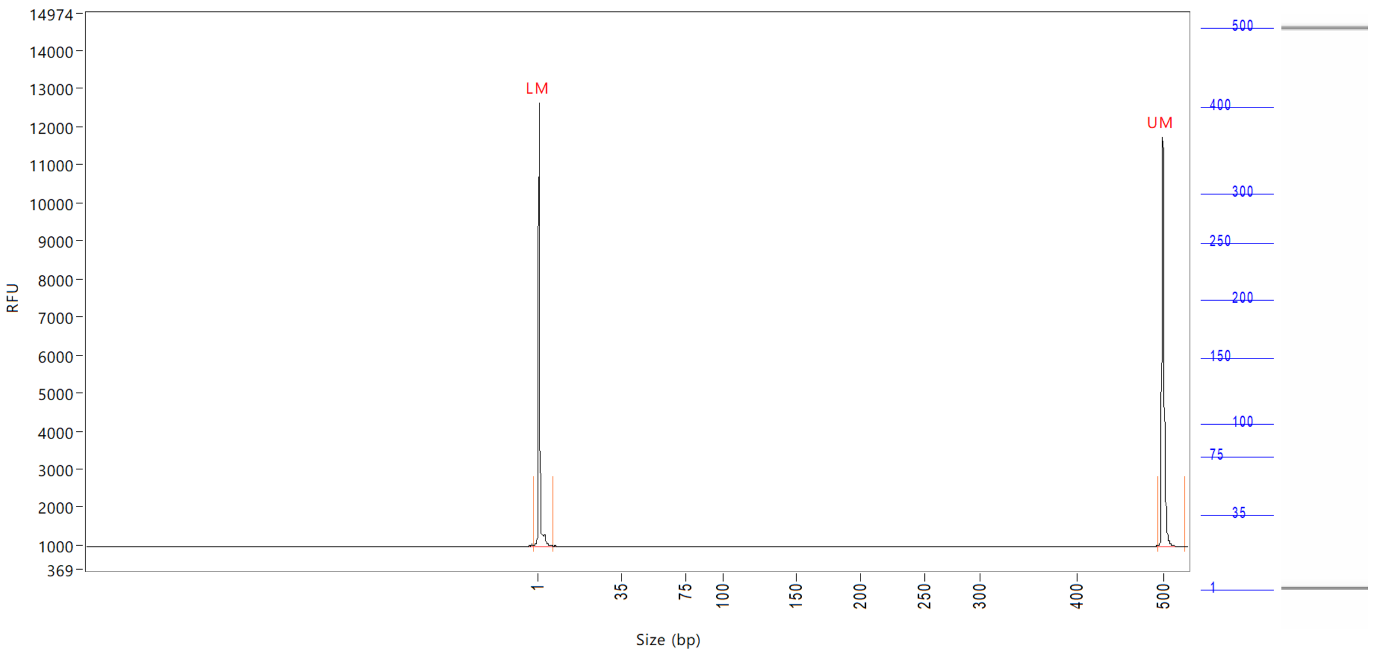


| Peak | Size (bp) | Concentration (ng/uL) | From (bp) | To (bp) | Average size (bp) | CV% | RFU | Corrected peak area |
|---|---|---|---|---|---|---|---|---|
| 1 | 1 (LM) | 0.7863 | 0 | 8 | 1 | 68.25 | 11670 | 43.710 |
| 2 | 500 (UM) | 0.5000 | 495 | 0 | 500 | 0.38 | 10755 | 27.793 |

| | | |
|---|---|---|
| TIC: | 0.0000 | ng/uL |
| TIM: | 0.0000 | nmole/L |
| Total concentration: | 0.3879 | ng/uL |

Sample peak width (sec): 5 Sample min peak height: 50 Sample baseline V to V?: Y Sample baseline V to V points: 3
Sample filter: Binomial Number of points for filter: 3 Sample start region (min): 0 Sample end region (min): 80
Marker peak width (sec): 5 Marker min peak height: 500 Marker baseline V to V?: Y Marker baseline V to V points: 3
Lower marker selection: First peak > 500 RFU Upper marker selection: Last peak > 500 RFU
Ladder size (bp)1, 35, 75, 100, 150, 200, 250, 300, 400, 500
Quantification using: Upper Marker Final concentration (ng/uL): 0.5000 Dilution factor: 12.0

**Sample:** SampH7
**Well location:** H7
**Created:** Monday, 11 October 2021 14:41:52

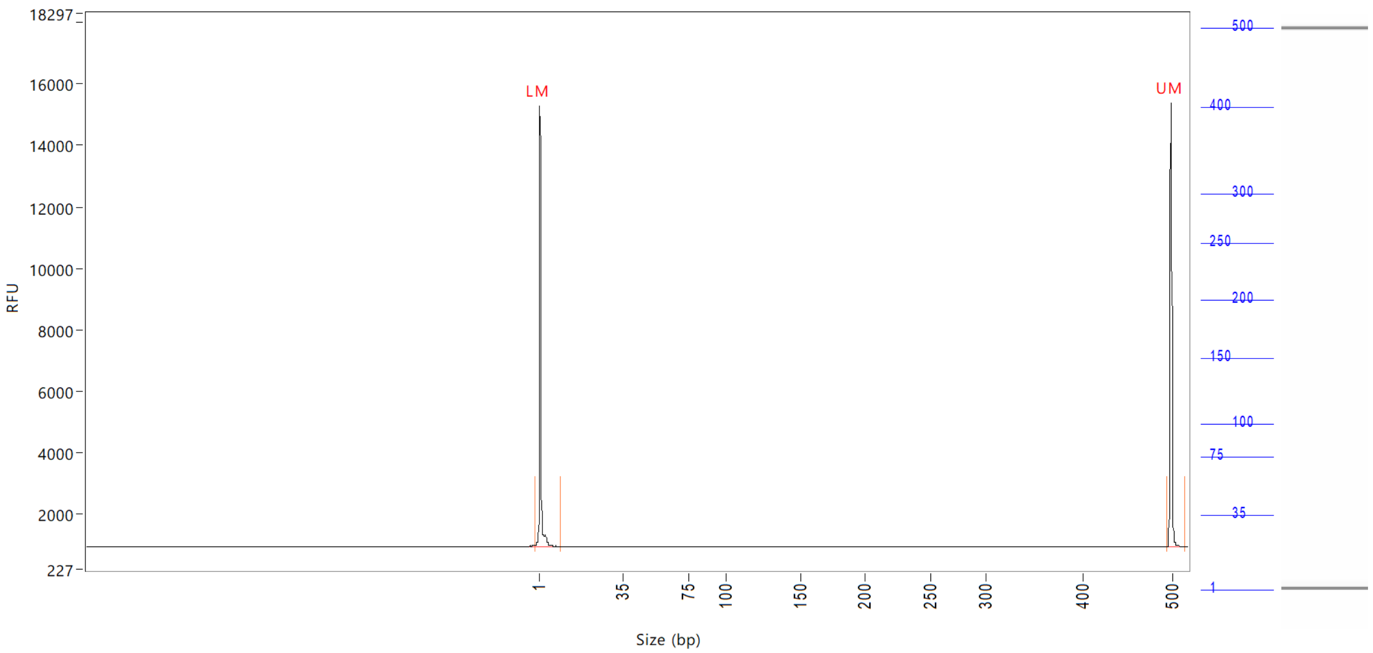


| Peak | Size (bp) | Concentration (ng/uL) | From (bp) | To (bp) | Average size (bp) | CV% | RFU | Corrected peak area |
|---|---|---|---|---|---|---|---|---|
| 1 | 1 (LM) | 0.8848 | 0 | 10 | 1 | 72.88 | 14332 | 53.752 |
| 2 | 500 (UM) | 0.5000 | 496 | 517 | 500 | 0.26 | 14452 | 30.375 |

| | | |
|---|---|---|
| TIC: | 0.0000 | ng/uL |
| TIM: | 0.0000 | nmole/L |
| Total concentration: | 0.3401 | ng/uL |

Sample peak width (sec): 5    Sample min peak height: 50    Sample baseline V to V?: Y    Sample baseline V to V points: 3
Sample filter: Binomial    Number of points for filter: 3  Sample start region (min): 0  Sample end region (min): 80
Marker peak width (sec): 5    Marker min peak height: 500    Marker baseline V to V?: Y    Marker baseline V to V points: 3
Lower marker selection: First peak > 500 RFU    Upper marker selection: Last peak > 500 RFU
Ladder size (bp)1, 35, 75, 100, 150, 200, 250, 300, 400, 500
Quantification using: Upper Marker    Final concentration (ng/uL): 0.5000    Dilution factor: 12.0

**Sample:** SampA8
**Well location:** A8
**Created:** Monday, 11 October 2021 14:41:52

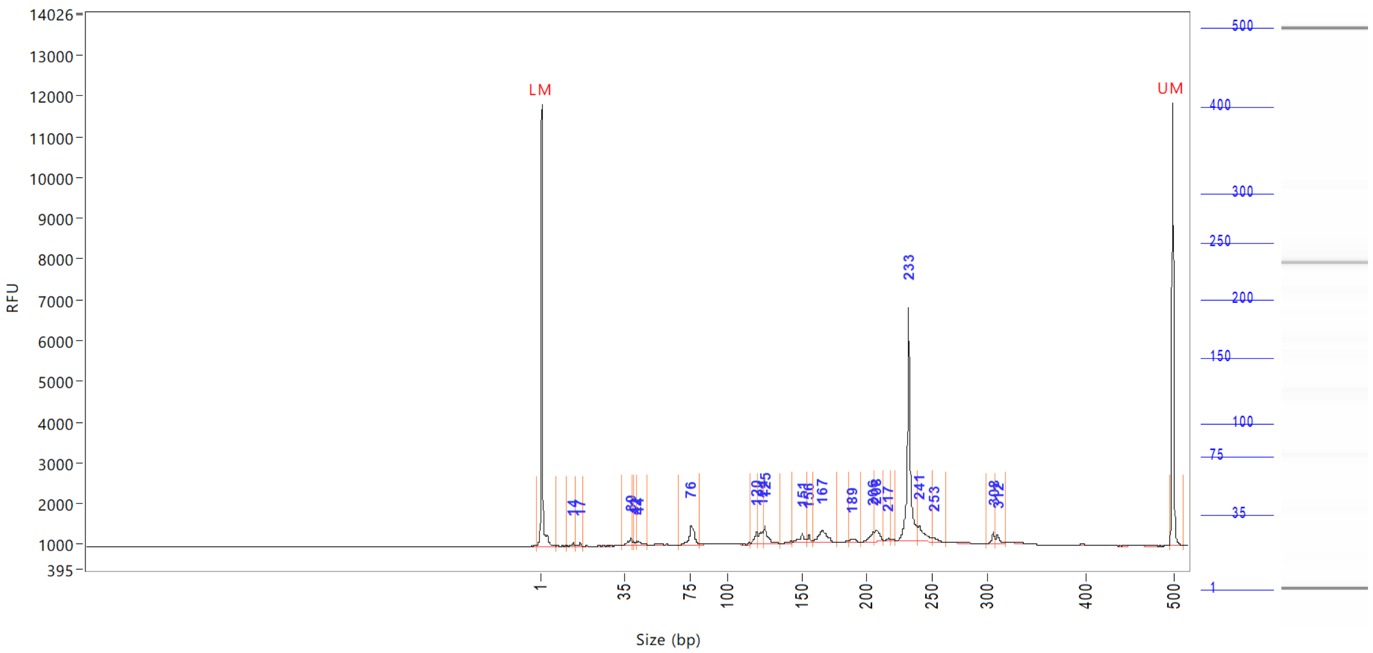

| Peak | Size (bp) | Concentration (ng/uL) | From (bp) | To (bp) | Average size (bp) | CV% | RFU | Corrected peak area |
|---|---|---|---|---|---|---|---|---|
| 1 | 1 (LM) | 0.9082 | 0 | 8 | 1 | 71.39 | 10841 | 40.683 |
| 2 | 14 | 0.1233 | 12 | 15 | 14 | 5.60 | 82 | 0.460 |
| 3 | 17 | 0.1157 | 15 | 18 | 17 | 3.14 | 85 | 0.432 |
| 4 | 39 | 0.3558 | 34 | 40 | 38 | 3.67 | 165 | 1.328 |
| 5 | 41 | 0.0890 | 40 | 41 | 41 | 0.98 | 96 | 0.332 |
| 6 | 42 | 0.0921 | 41 | 43 | 42 | 1.05 | 89 | 0.344 |
| 7 | 44 | 0.2074 | 43 | 49 | 45 | 3.48 | 98 | 0.774 |
| 8 | 76 | 1.3057 | 68 | 82 | 76 | 3.14 | 477 | 4.874 |
| 9 | 120 | 0.5307 | 116 | 121 | 119 | 1.04 | 312 | 1.981 |
| 10 | 124 | 0.5355 | 121 | 124 | 123 | 0.76 | 291 | 1.999 |
| 11 | 125 | 0.7538 | 124 | 136 | 127 | 1.58 | 458 | 2.814 |
| 12 | 151 | 0.5205 | 143 | 155 | 150 | 1.93 | 223 | 1.943 |
| 13 | 156 | 0.1994 | 155 | 159 | 156 | 0.66 | 188 | 0.744 |
| 14 | 167 | 1.0548 | 159 | 177 | 166 | 1.91 | 285 | 3.938 |
| 15 | 189 | 0.1838 | 186 | 195 | 190 | 0.95 | 79 | 0.686 |
| 16 | 206 | 0.4297 | 195 | 207 | 204 | 1.13 | 229 | 1.604 |
| 17 | 208 | 0.5483 | 207 | 214 | 209 | 0.80 | 278 | 2.047 |
| 18 | 217 | 0.0892 | 214 | 220 | 217 | 0.64 | 59 | 0.333 |
| 19 | 233 | 6.5325 | 222 | 239 | 233 | 0.96 | 5711 | 24.386 |
| 20 | 241 | 0.9523 | 239 | 251 | 243 | 1.25 | 386 | 3.555 |
| 21 | 253 | 0.1603 | 251 | 264 | 255 | 0.98 | 86 | 0.599 |
| 22 | 308 | 0.2569 | 299 | 310 | 307 | 0.55 | 268 | 0.959 |
| 23 | 312 | 0.2334 | 310 | 320 | 313 | 0.72 | 203 | 0.871 |
| 24 | 500 (UM) | 0.5000 | 496 | 512 | 500 | 0.24 | 10872 | 22.398 |

```
Sample peak width (sec): 5    Sample min peak height: 50    Sample baseline V to V?: Y    Sample baseline V to V points: 3
Sample filter: Binomial       Number of points for filter: 3  Sample start region (min): 0  Sample end region (min): 80
Marker peak width (sec): 5    Marker min peak height: 500   Marker baseline V to V?: Y    Marker baseline V to V points: 3
Lower marker selection: First peak > 500 RFU                Upper marker selection: Last peak > 500 RFU
Ladder size (bp)1, 35, 75, 100, 150, 200, 250, 300, 400, 500
Quantification using: Upper Marker          Final concentration (ng/uL): 0.5000          Dilution factor: 12.0
```

**Sample:** SampA8
**Well location:** A8
**Created:** Monday, 11 October 2021 14:41:52

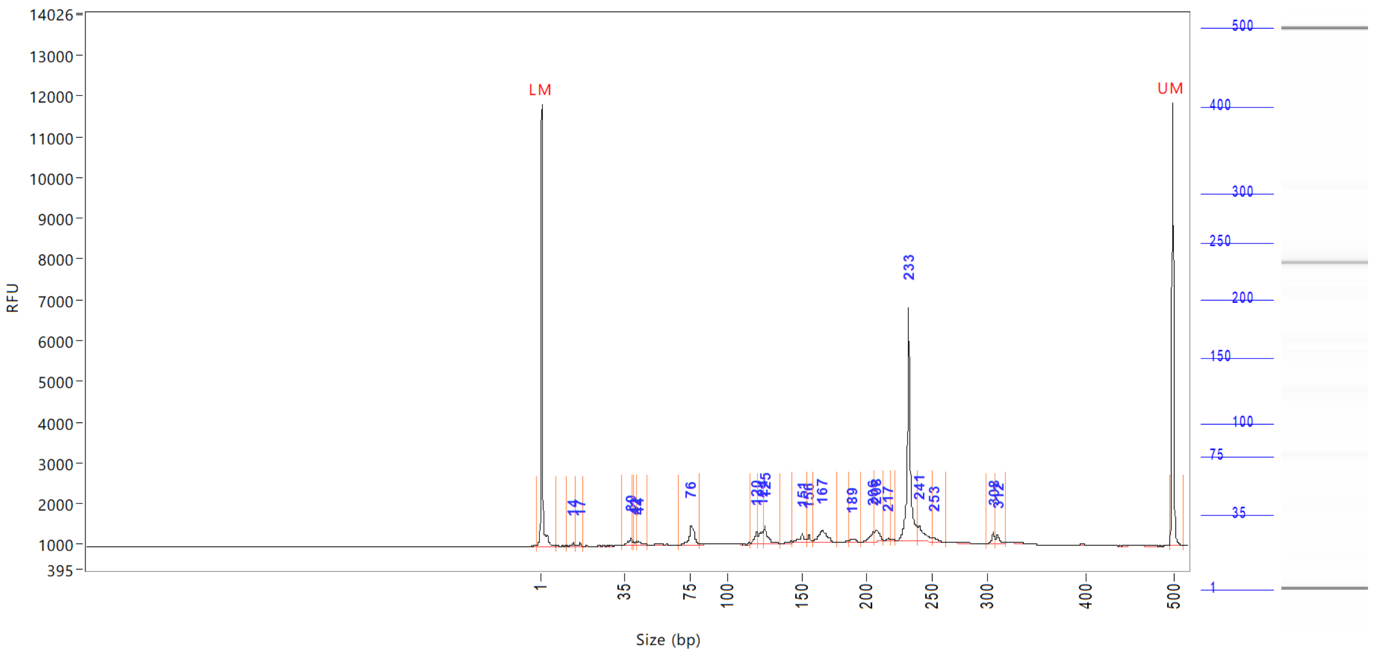


| Peak | Size (bp) | Concentration (ng/uL) | From (bp) | To (bp) | Average size (bp) | CV% | RFU | Corrected peak area |
|---|---|---|---|---|---|---|---|---|

| | | |
|---|---|---|
| TIC: | 15.2701 | ng/uL |
| TIM: | 192.1725 | nmole/L |
| Total concentration: | 16.1611 | ng/uL |

Sample peak width (sec): 5 Sample min peak height: 50 Sample baseline V to V?: Y Sample baseline V to V points: 3
Sample filter: Binomial Number of points for filter: 3 Sample start region (min): 0 Sample end region (min): 80
Marker peak width (sec): 5 Marker min peak height: 500 Marker baseline V to V?: Y Marker baseline V to V points: 3
Lower marker selection: First peak > 500 RFU Upper marker selection: Last peak > 500 RFU
Ladder size (bp)1, 35, 75, 100, 150, 200, 250, 300, 400, 500
Quantification using: Upper Marker Final concentration (ng/uL): 0.5000 Dilution factor: 12.0

**Sample:** SampB8
**Well location:** B8
**Created:** Monday, 11 October 2021 14:41:52

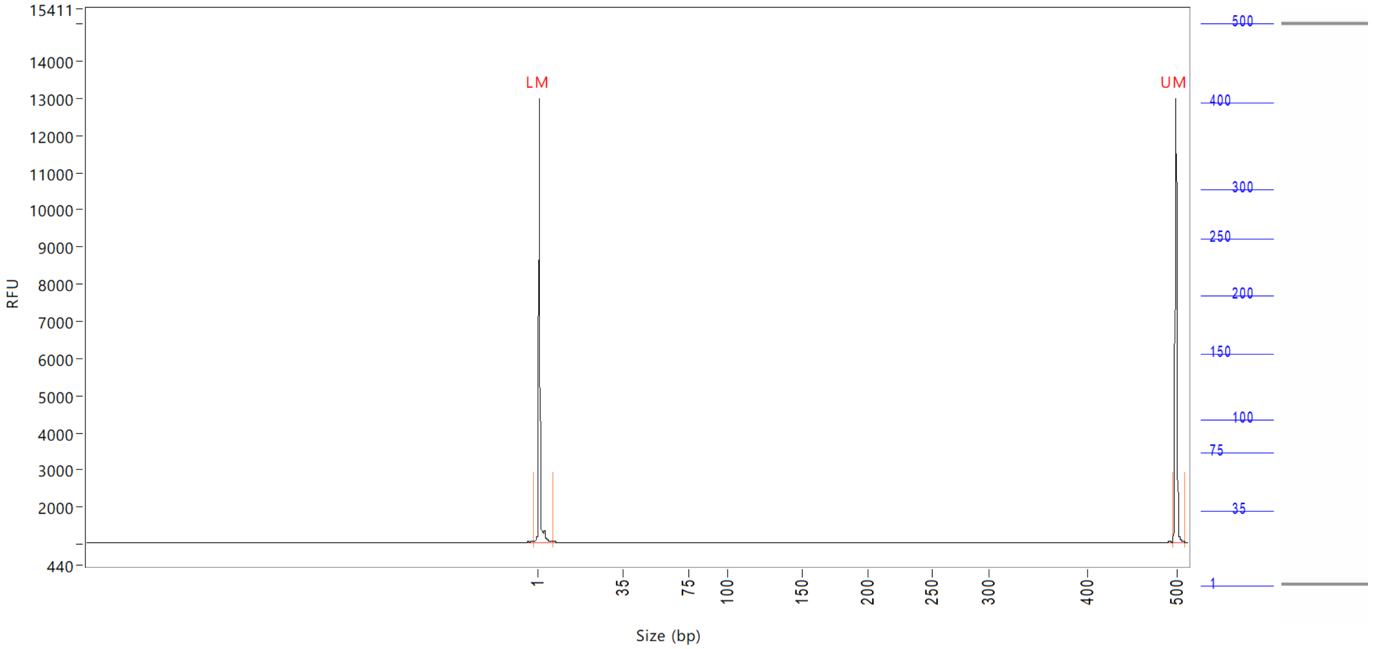


| Peak | Size (bp) | Concentration (ng/uL) | From (bp) | To (bp) | Average size (bp) | CV% | RFU | Corrected peak area |
|---|---|---|---|---|---|---|---|---|
| 1 | 1 (LM) | 0.8822 | 0 | 8 | 1 | 68.15 | 11959 | 44.513 |
| 2 | 500 (UM) | 0.5000 | 496 | 511 | 500 | 0.23 | 11962 | 25.228 |

| | | |
|---|---|---|
| TIC: | 0.0000 | ng/uL |
| TIM: | 0.0000 | nmole/L |
| Total concentration: | 0.5502 | ng/uL |

```
Sample peak width (sec): 5    Sample min peak height: 50    Sample baseline V to V?: Y    Sample baseline V to V points: 3
Sample filter: Binomial       Number of points for filter: 3  Sample start region (min): 0  Sample end region (min): 80
Marker peak width (sec): 5    Marker min peak height: 500   Marker baseline V to V?: Y    Marker baseline V to V points: 3
Lower marker selection: First peak > 500 RFU                Upper marker selection: Last peak > 500 RFU
Ladder size (bp)1, 35, 75, 100, 150, 200, 250, 300, 400, 500
Quantification using: Upper Marker          Final concentration (ng/uL): 0.5000          Dilution factor: 12.0
```

**Sample:** SampC8
**Well location:** C8
**Created:** Monday, 11 October 2021 14:41:52

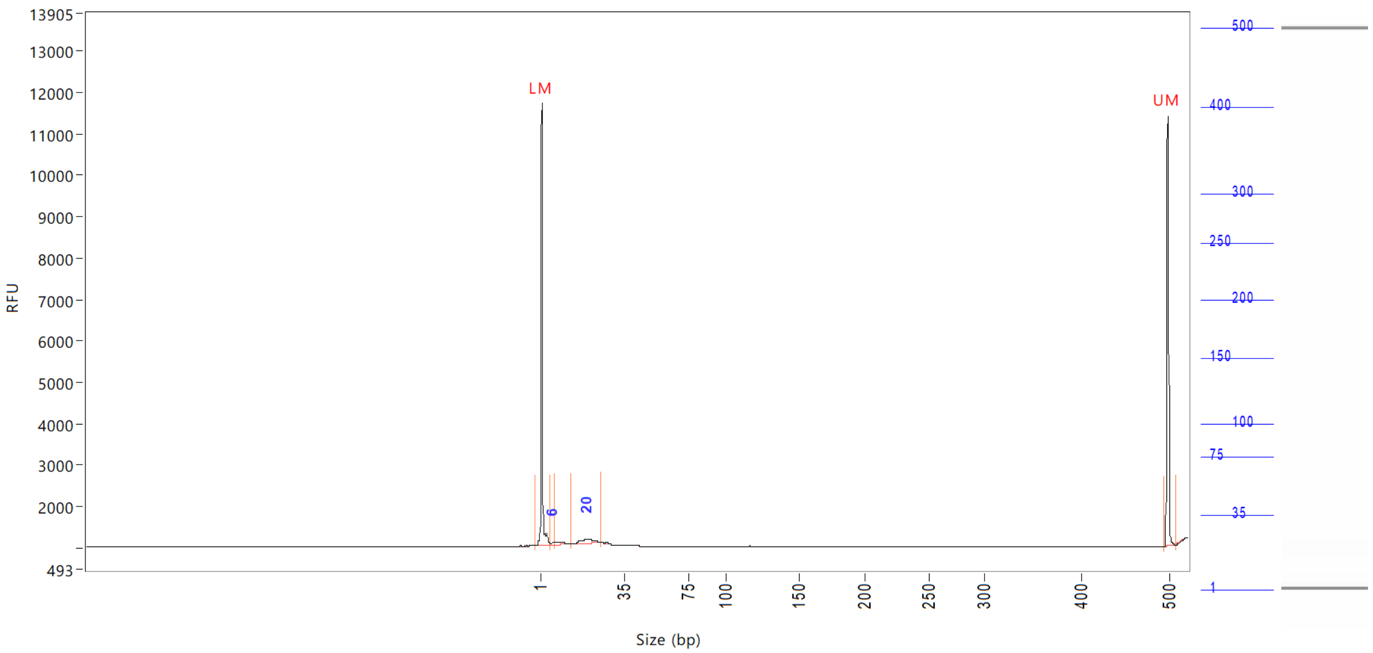


| Peak | Size (bp) | Concentration (ng/uL) | From (bp) | To (bp) | Average size (bp) | CV% | RFU | Corrected peak area |
|---|---|---|---|---|---|---|---|---|
| 1 | 1 (LM) | 0.9134 | 0 | 6 | 1 | 66.89 | 10687 | 38.141 |
| 2 | 6 | 0.1391 | 6 | 8 | 7 | 8.23 | 51 | 0.484 |
| 3 | 20 | 0.8774 | 14 | 25 | 20 | 11.69 | 94 | 3.053 |
| 4 | 500 (UM) | 0.5000 | 496 | 508 | 500 | 0.21 | 10350 | 20.879 |

| | | |
|---|---|---|
| TIC: | 1.0165 | ng/uL |
| TIM: | 102.8446 | nmole/L |
| Total concentration: | 1.8310 | ng/uL |

```
Sample peak width (sec): 5    Sample min peak height: 50    Sample baseline V to V?: Y    Sample baseline V to V points: 3
Sample filter: Binomial       Number of points for filter: 3  Sample start region (min): 0  Sample end region (min): 80
Marker peak width (sec): 5    Marker min peak height: 500   Marker baseline V to V?: Y    Marker baseline V to V points: 3
Lower marker selection: First peak > 500 RFU                Upper marker selection: Last peak > 500 RFU
Ladder size (bp)1, 35, 75, 100, 150, 200, 250, 300, 400, 500
Quantification using: Upper Marker          Final concentration (ng/uL): 0.5000          Dilution factor: 12.0
```

**Sample:** SampD8
**Well location:** D8
**Created:** Monday, 11 October 2021 14:41:52

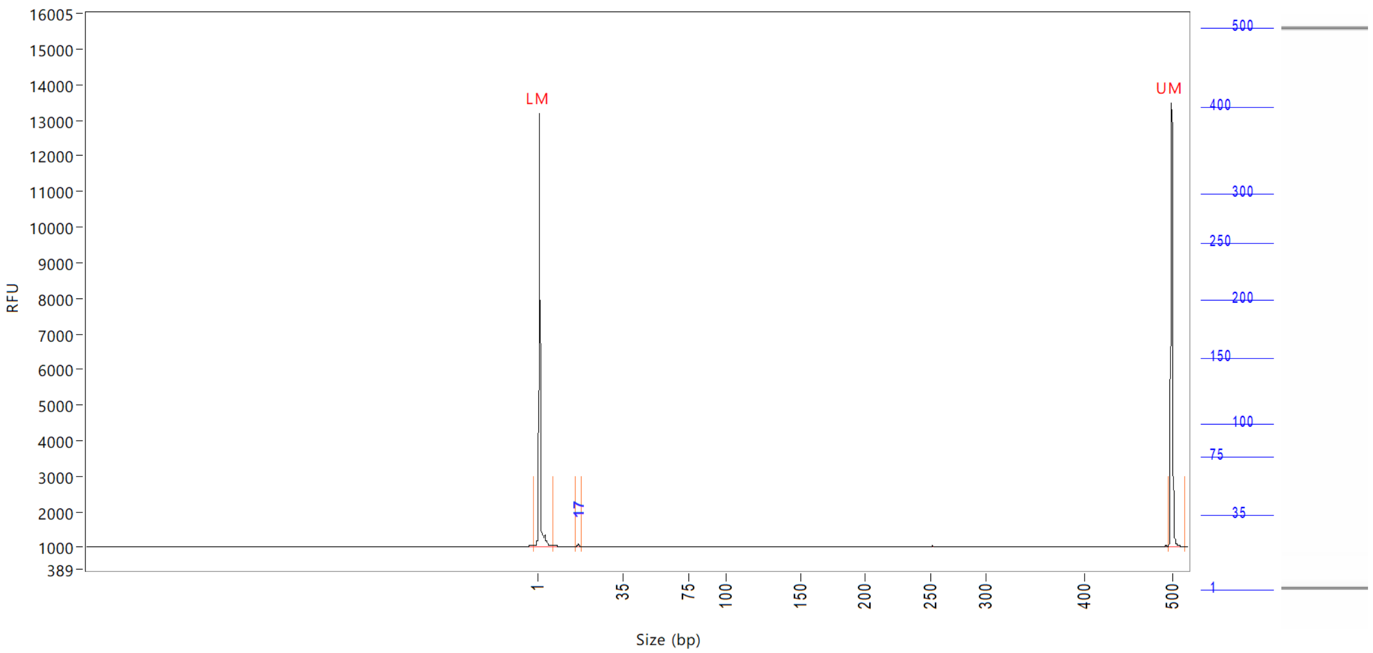


| Peak | Size (bp) | Concentration (ng/uL) | From (bp) | To (bp) | Average size (bp) | CV% | RFU | Corrected peak area |
|---|---|---|---|---|---|---|---|---|
| 1 | 1 (LM) | 0.8777 | 0 | 8 | 1 | 67.32 | 12146 | 45.810 |
| 2 | 17 | 0.0807 | 16 | 19 | 17 | 2.35 | 77 | 0.351 |
| 3 | 500 (UM) | 0.5000 | 496 | 515 | 500 | 0.25 | 12485 | 26.097 |

| | | |
|---|---|---|
| TIC: | 0.0807 | ng/uL |
| TIM: | 7.6977 | nmole/L |
| Total concentration: | 0.6035 | ng/uL |

```
Sample peak width (sec): 5    Sample min peak height: 50    Sample baseline V to V?: Y    Sample baseline V to V points: 3
Sample filter: Binomial       Number of points for filter: 3  Sample start region (min): 0  Sample end region (min): 80
Marker peak width (sec): 5    Marker min peak height: 500   Marker baseline V to V?: Y    Marker baseline V to V points: 3
Lower marker selection: First peak > 500 RFU                Upper marker selection: Last peak > 500 RFU
Ladder size (bp)1, 35, 75, 100, 150, 200, 250, 300, 400, 500
Quantification using: Upper Marker          Final concentration (ng/uL): 0.5000          Dilution factor: 12.0
```

**Sample:** SampE8
**Well location:** E8
**Created:** Monday, 11 October 2021 14:41:52

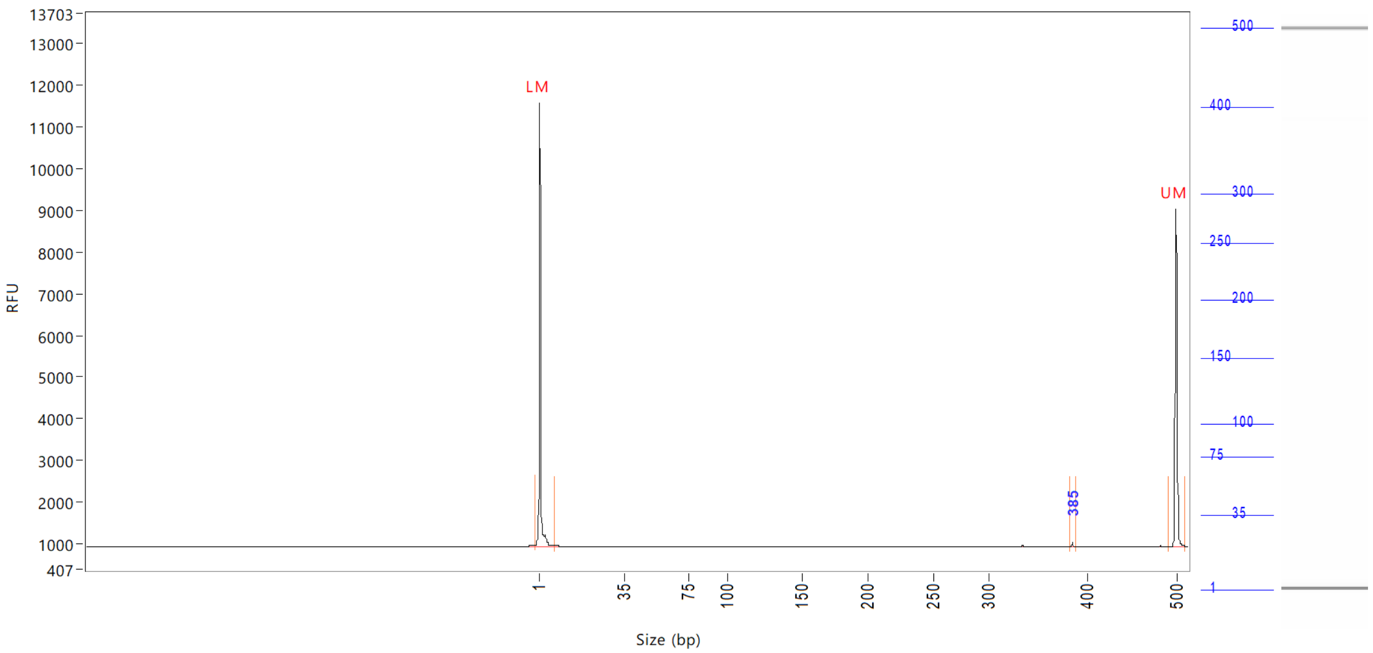


| Peak | Size (bp) | Concentration (ng/uL) | From (bp) | To (bp) | Average size (bp) | CV% | RFU | Corrected peak area |
|---|---|---|---|---|---|---|---|---|
| 1 | 1 (LM) | 1.1496 | 0 | 8 | 1 | 69.92 | 10618 | 39.850 |
| 2 | 385 | 0.0521 | 383 | 388 | 385 | 0.16 | 104 | 0.150 |
| 3 | 500 (UM) | 0.5000 | 492 | 510 | 500 | 0.25 | 8059 | 17.332 |

| | | |
|---|---|---|
| TIC: | 0.0521 | ng/uL |
| TIM: | 0.2224 | nmole/L |
| Total concentration: | 0.7205 | ng/uL |

Sample peak width (sec): 5    Sample min peak height: 50    Sample baseline V to V?: Y    Sample baseline V to V points: 3
Sample filter: Binomial    Number of points for filter: 3    Sample start region (min): 0    Sample end region (min): 80
Marker peak width (sec): 5    Marker min peak height: 500    Marker baseline V to V?: Y    Marker baseline V to V points: 3
Lower marker selection: First peak > 500 RFU    Upper marker selection: Last peak > 500 RFU
Ladder size (bp)1, 35, 75, 100, 150, 200, 250, 300, 400, 500
Quantification using: Upper Marker    Final concentration (ng/uL): 0.5000    Dilution factor: 12.0

**Sample:** SampF8
**Well location:** F8
**Created:** Monday, 11 October 2021 14:41:52

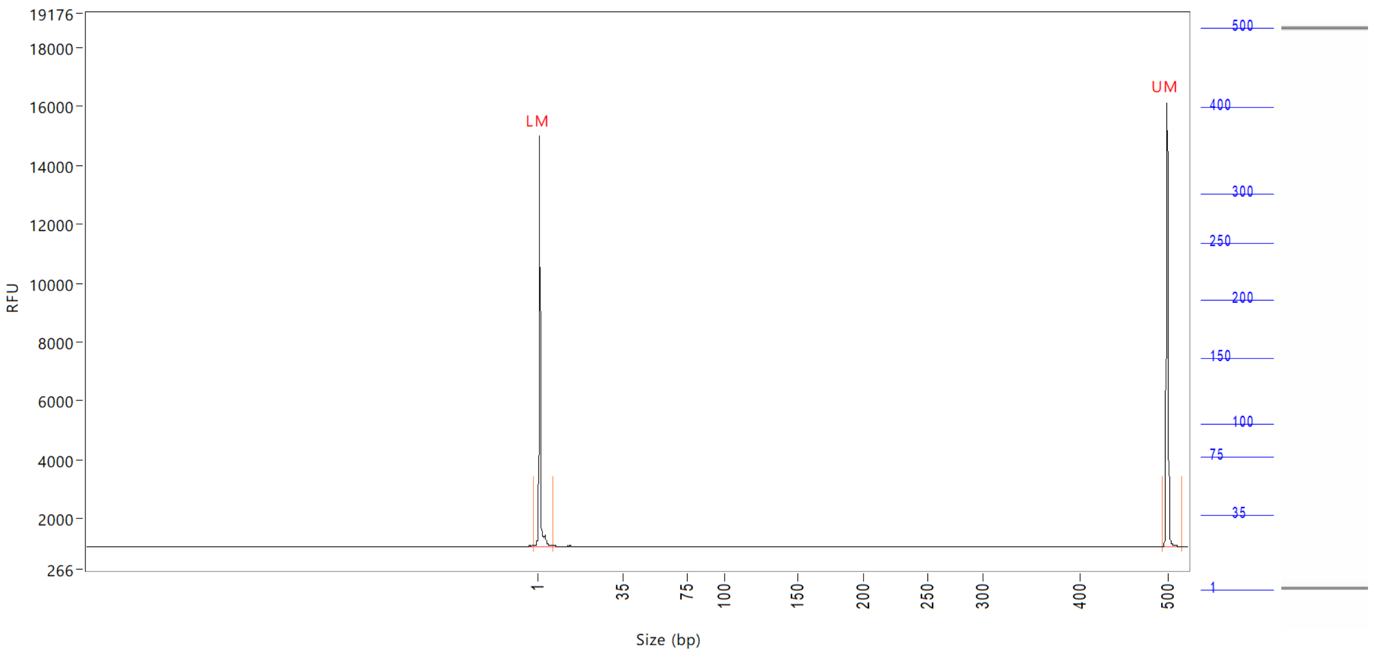


| Peak | Size (bp) | Concentration (ng/uL) | From (bp) | To (bp) | Average size (bp) | CV% | RFU | Corrected peak area |
|---|---|---|---|---|---|---|---|---|
| 1 | 1 (LM) | 0.7895 | 0 | 8 | 1 | 70.97 | 13946 | 51.086 |
| 2 | 500 (UM) | 0.5000 | 494 | 517 | 500 | 0.27 | 15117 | 32.353 |

| | | |
|---|---|---|
| TIC: | 0.0000 | ng/uL |
| TIM: | 0.0000 | nmole/L |
| Total concentration: | 0.3955 | ng/uL |

Sample peak width (sec): 5 Sample min peak height: 50 Sample baseline V to V?: Y Sample baseline V to V points: 3
Sample filter: Binomial Number of points for filter: 3 Sample start region (min): 0 Sample end region (min): 80
Marker peak width (sec): 5 Marker min peak height: 500 Marker baseline V to V?: Y Marker baseline V to V points: 3
Lower marker selection: First peak > 500 RFU Upper marker selection: Last peak > 500 RFU
Ladder size (bp)1, 35, 75, 100, 150, 200, 250, 300, 400, 500
Quantification using: Upper Marker Final concentration (ng/uL): 0.5000 Dilution factor: 12.0

**Sample:** SampG8
**Well location:** G8
**Created:** Monday, 11 October 2021 14:41:52

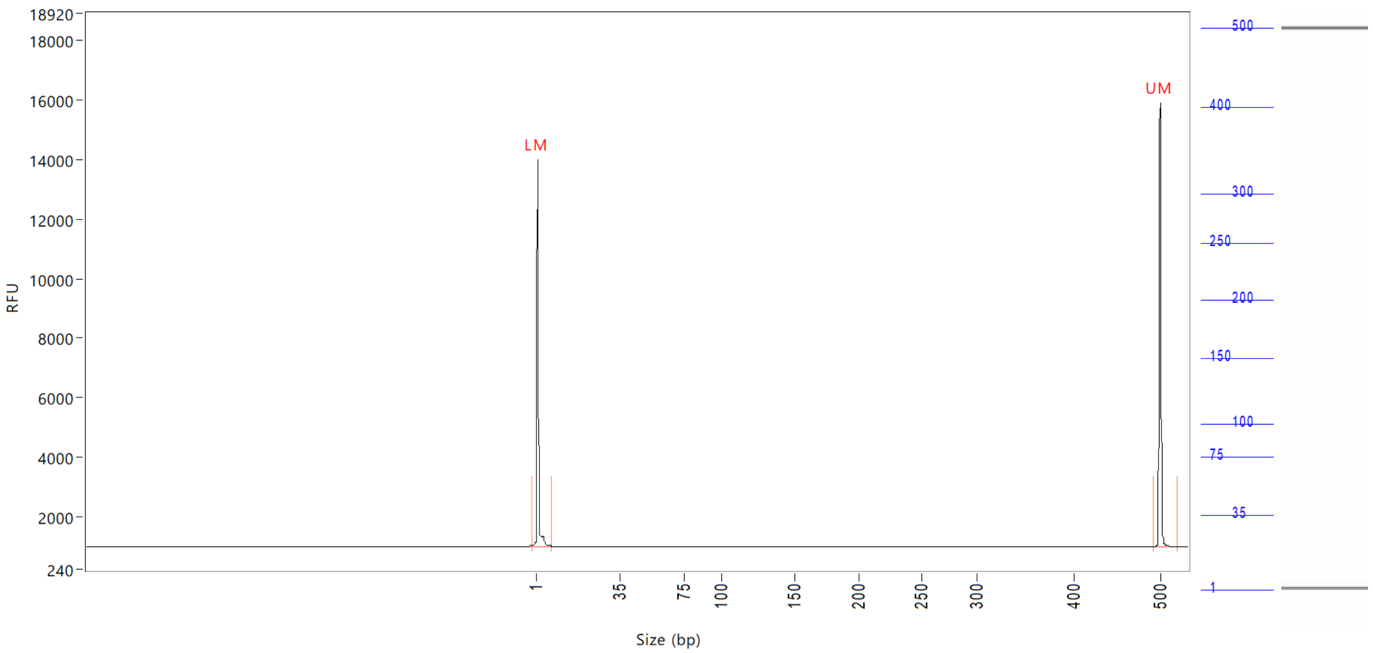


| Peak | Size<br>(bp) | Concentration<br>(ng/uL) | From<br>(bp) | To<br>(bp) | Average size<br>(bp) | CV% | RFU | Corrected peak area |
|---|---|---|---|---|---|---|---|---|
| 1 | 1 (LM) | 0.7925 | 0 | 7 | 1 | 69.63 | 12989 | 48.956 |
| 2 | 500 (UM) | 0.5000 | 493 | 0 | 500 | 0.28 | 14937 | 30.886 |

| | | |
|---|---|---|
| TIC: | 0.0000 | ng/uL |
| TIM: | 0.0000 | nmole/L |
| Total concentration: | 0.3949 | ng/uL |

```
Sample peak width (sec): 5    Sample min peak height: 50    Sample baseline V to V?: Y    Sample baseline V to V points: 3
Sample filter: Binomial       Number of points for filter: 3  Sample start region (min): 0  Sample end region (min): 80
Marker peak width (sec): 5    Marker min peak height: 500   Marker baseline V to V?: Y    Marker baseline V to V points: 3
Lower marker selection: First peak > 500 RFU                Upper marker selection: Last peak > 500 RFU
Ladder size (bp)1, 35, 75, 100, 150, 200, 250, 300, 400, 500
Quantification using: Upper Marker          Final concentration (ng/uL): 0.5000          Dilution factor: 12.0
```

**Sample:** SampH8
**Well location:** H8
**Created:** Monday, 11 October 2021 14:41:52

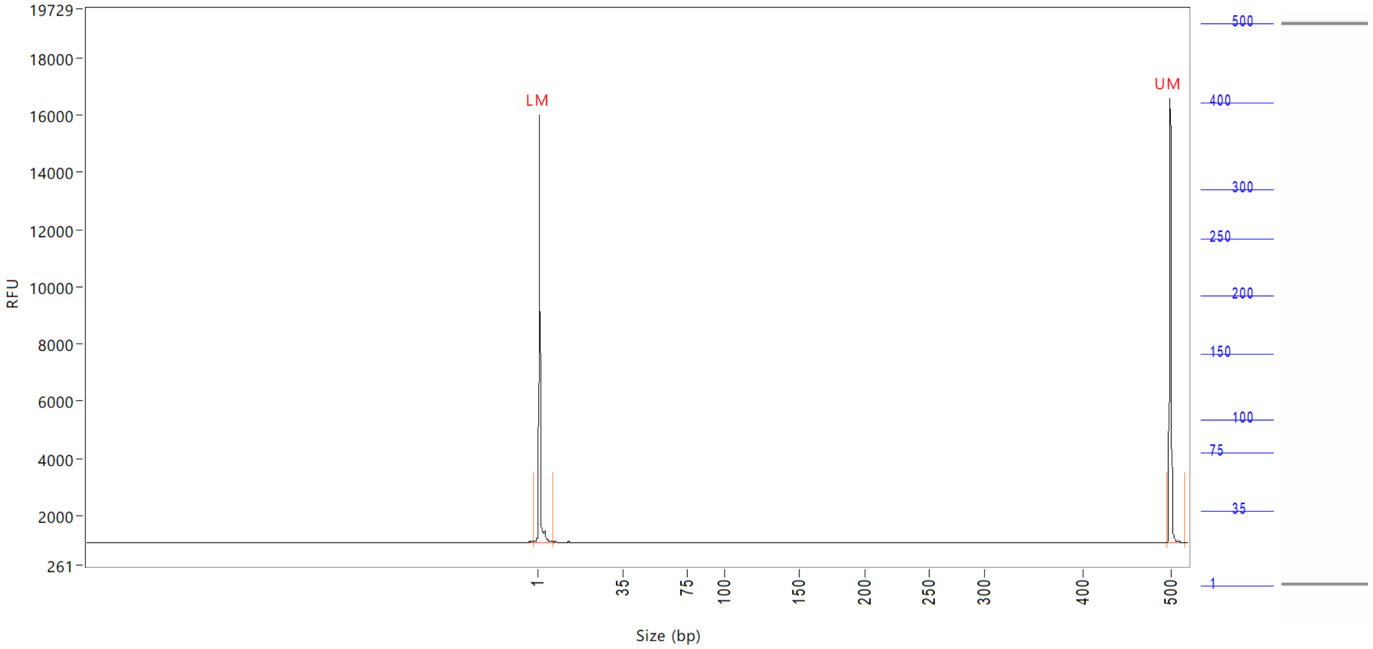


| Peak | Size (bp) | Concentration (ng/uL) | From (bp) | To (bp) | Average size (bp) | CV% | RFU | Corrected peak area |
|---|---|---|---|---|---|---|---|---|
| 1 | 1 (LM) | 0.8680 | 0 | 8 | 1 | 67.80 | 14963 | 55.682 |
| 2 | 500 (UM) | 0.5000 | 496 | 517 | 500 | 0.28 | 15569 | 32.076 |

| | | |
|---|---|---|
| TIC: | 0.0000 | ng/uL |
| TIM: | 0.0000 | nmole/L |
| Total concentration: | 0.4226 | ng/uL |

Sample peak width (sec): 5    Sample min peak height: 50    Sample baseline V to V?: Y    Sample baseline V to V points: 3
Sample filter: Binomial    Number of points for filter: 3  Sample start region (min): 0  Sample end region (min): 80
Marker peak width (sec): 5    Marker min peak height: 500    Marker baseline V to V?: Y    Marker baseline V to V points: 3
Lower marker selection: First peak > 500 RFU    Upper marker selection: Last peak > 500 RFU
Ladder size (bp)1, 35, 75, 100, 150, 200, 250, 300, 400, 500
Quantification using: Upper Marker    Final concentration (ng/uL): 0.5000    Dilution factor: 12.0

**Sample:** SampA9
**Well location:** A9
**Created:** Monday, 11 October 2021 14:41:52

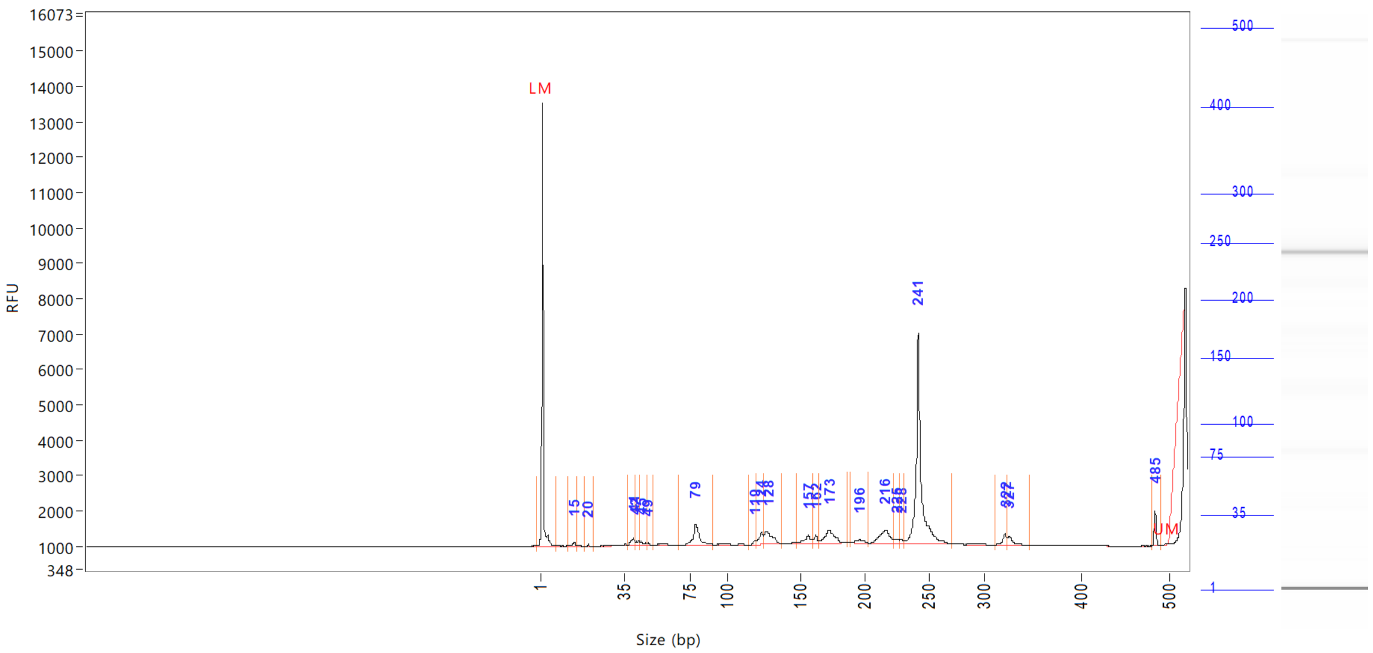


| Peak | Size (bp) | Concentration (ng/uL) | From (bp) | To (bp) | Average size (bp) | CV% | RFU | Corrected peak area |
|---|---|---|---|---|---|---|---|---|
| 1 | 1 (LM) | Inf | 0 | 8 | 1 | 73.75 | 12564 | 47.509 |
| 2 | 15 | Inf | 12 | 16 | 14 | 4.76 | 117 | 0.733 |
| 3 | 20 | Inf | 19 | 22 | 20 | 2.11 | 58 | 0.204 |
| 4 | 41 | Inf | 37 | 41 | 40 | 2.50 | 198 | 1.343 |
| 5 | 42 | Inf | 41 | 44 | 43 | 1.99 | 110 | 0.779 |
| 6 | 45 | Inf | 44 | 49 | 46 | 2.90 | 101 | 0.780 |
| 7 | 49 | Inf | 49 | 53 | 49 | 1.32 | 59 | 0.231 |
| 8 | 79 | Inf | 68 | 91 | 80 | 4.07 | 597 | 7.275 |
| 9 | 119 | Inf | 116 | 120 | 119 | 0.69 | 122 | 0.550 |
| 10 | 124 | Inf | 120 | 125 | 123 | 1.00 | 352 | 2.302 |
| 11 | 128 | Inf | 125 | 138 | 129 | 2.11 | 353 | 5.167 |
| 12 | 157 | Inf | 148 | 160 | 155 | 2.01 | 247 | 2.537 |
| 13 | 162 | Inf | 160 | 165 | 162 | 0.76 | 222 | 1.207 |
| 14 | 173 | Inf | 165 | 186 | 173 | 2.08 | 358 | 5.423 |
| 15 | 196 | Inf | 189 | 203 | 196 | 1.42 | 92 | 1.060 |
| 16 | 216 | Inf | 203 | 223 | 215 | 1.72 | 377 | 5.550 |
| 17 | 225 | Inf | 223 | 227 | 225 | 0.50 | 120 | 0.703 |
| 18 | 228 | Inf | 227 | 231 | 229 | 0.51 | 104 | 0.568 |
| 19 | 241 | Inf | 231 | 271 | 243 | 2.05 | 5958 | 39.332 |
| 20 | 322 | Inf | 313 | 324 | 321 | 0.73 | 317 | 1.567 |
| 21 | 327 | Inf | 324 | 347 | 330 | 1.44 | 264 | 1.724 |
| 22 | 500 (UM) | Inf | 481 | 491 | 485 | 0.23 | 1003 | 2.269 |

| | | |
|---|---|---|
| TIC: | Inf | ng/uL |
| TIM: | Inf | nmole/L |

```
Sample peak width (sec): 5     Sample min peak height: 50    Sample baseline V to V?: Y    Sample baseline V to V points: 3
Sample filter: Binomial        Number of points for filter: 3  Sample start region (min): 0  Sample end region (min): 80
Marker peak width (sec): 5     Marker min peak height: 500   Marker baseline V to V?: Y    Marker baseline V to V points: 3
Lower marker selection: First peak > 500 RFU                 Upper marker selection: Last peak > 500 RFU
Ladder size (bp)1, 35, 75, 100, 150, 200, 250, 300, 400, 500
Quantification using: Upper Marker          Final concentration (ng/uL): 0.5000          Dilution factor: 12.0
```

**Sample:** SampA9
**Well location:** A9
**Created:** Monday, 11 October 2021 14:41:52

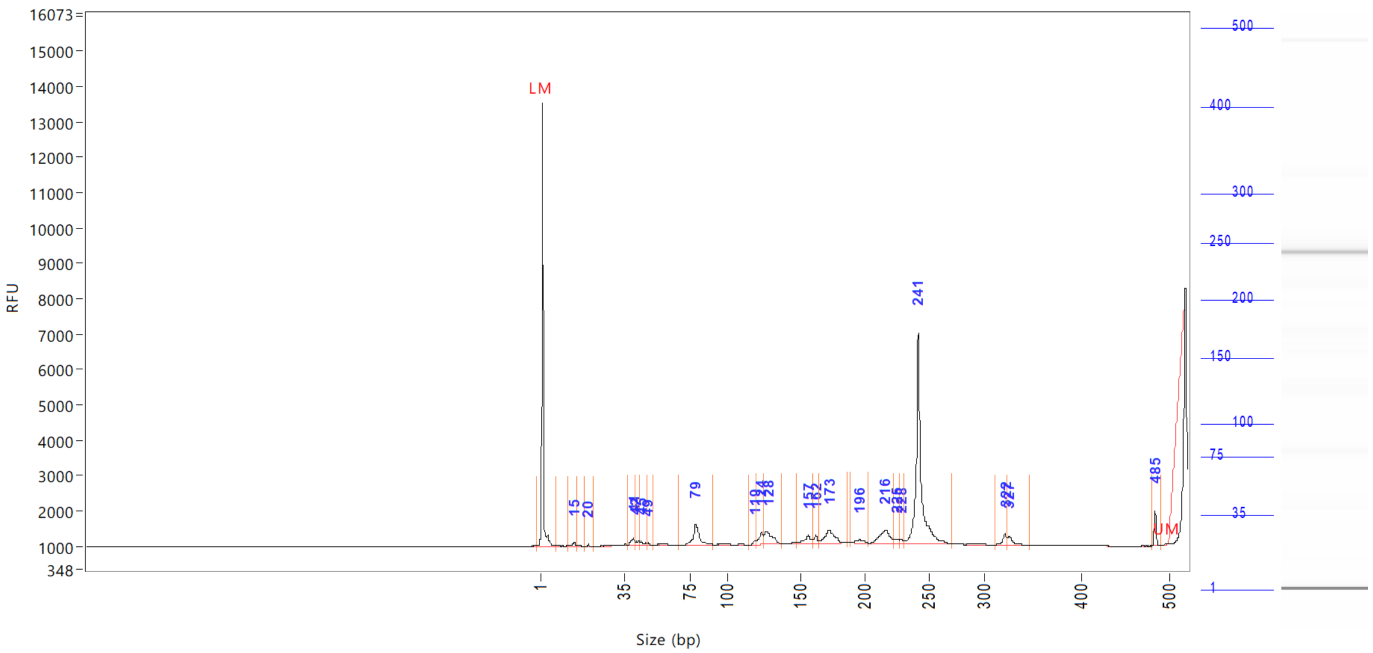

| Peak | Size | Concentration | From | To | Average size | CV% | RFU | Corrected peak area |
|---|---|---|---|---|---|---|---|---|
| | (bp) | (ng/uL) | (bp) | (bp) | (bp) | | | |

Total concentration: 0.0000 ng/uL

```
Sample peak width (sec): 5    Sample min peak height: 50    Sample baseline V to V?: Y    Sample baseline V to V points: 3
Sample filter: Binomial       Number of points for filter: 3  Sample start region (min): 0  Sample end region (min): 80
Marker peak width (sec): 5    Marker min peak height: 500   Marker baseline V to V?: Y    Marker baseline V to V points: 3
Lower marker selection: First peak > 500 RFU                Upper marker selection: Last peak > 500 RFU
Ladder size (bp)1, 35, 75, 100, 150, 200, 250, 300, 400, 500
Quantification using: Upper Marker         Final concentration (ng/uL): 0.5000         Dilution factor: 12.0
```

**Sample:** SampB9
**Well location:** B9
**Created:** Monday, 11 October 2021 14:41:52

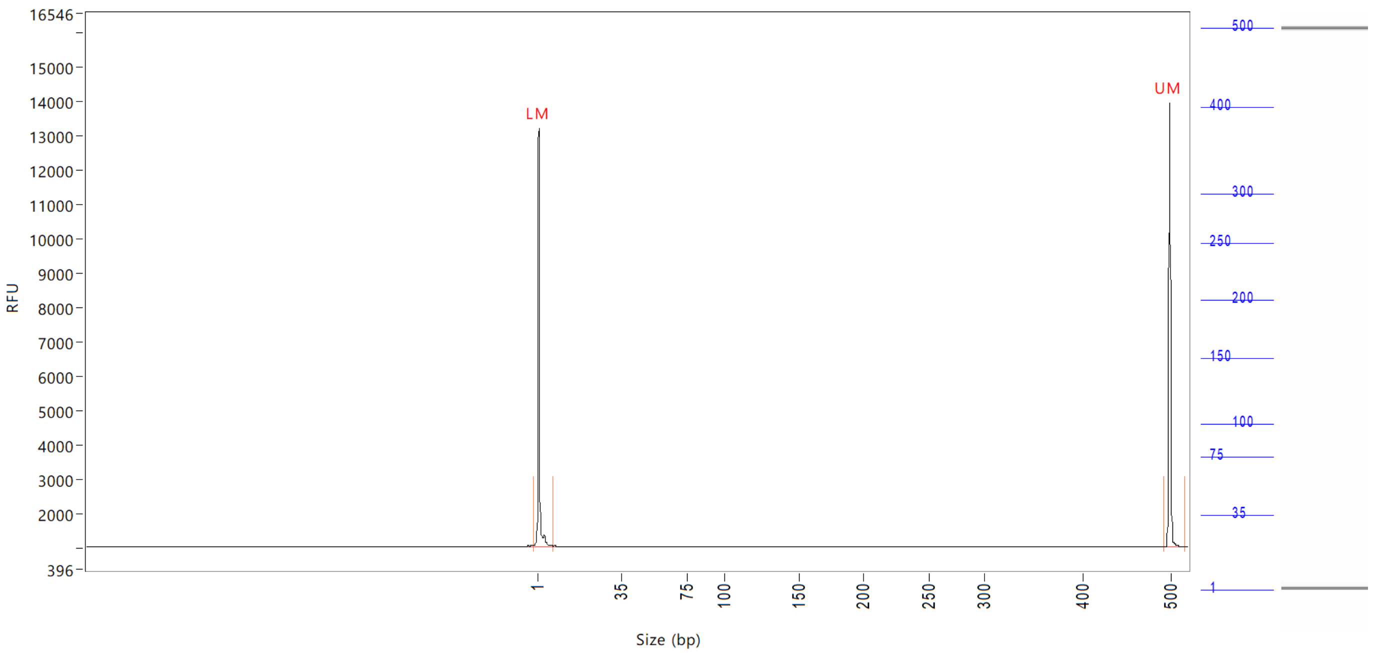


| Peak | Size | Concentration | From | To | Average size | CV% | RFU | Corrected peak area |
|---|---|---|---|---|---|---|---|---|
| | (bp) | (ng/uL) | (bp) | (bp) | (bp) | | | |
| 1 | 1 (LM) | 0.8689 | 0 | 8 | 1 | 72.51 | 12151 | 46.405 |
| 2 | 500 (UM) | 0.5000 | 493 | 0 | 500 | 0.26 | 12915 | 26.703 |

| | | |
|---|---|---|
| TIC: | 0.0000 | ng/uL |
| TIM: | 0.0000 | nmole/L |
| Total concentration: | 0.4752 | ng/uL |

```
Sample peak width (sec): 5    Sample min peak height: 50    Sample baseline V to V?: Y    Sample baseline V to V points: 3
Sample filter: Binomial       Number of points for filter: 3  Sample start region (min): 0  Sample end region (min): 80
Marker peak width (sec): 5    Marker min peak height: 500   Marker baseline V to V?: Y    Marker baseline V to V points: 3
Lower marker selection: First peak > 500 RFU                Upper marker selection: Last peak > 500 RFU
Ladder size (bp)1, 35, 75, 100, 150, 200, 250, 300, 400, 500
Quantification using: Upper Marker          Final concentration (ng/uL): 0.5000          Dilution factor: 12.0
```

**Sample:** SampC9
**Well location:** C9
**Created:** Monday, 11 October 2021 14:41:52

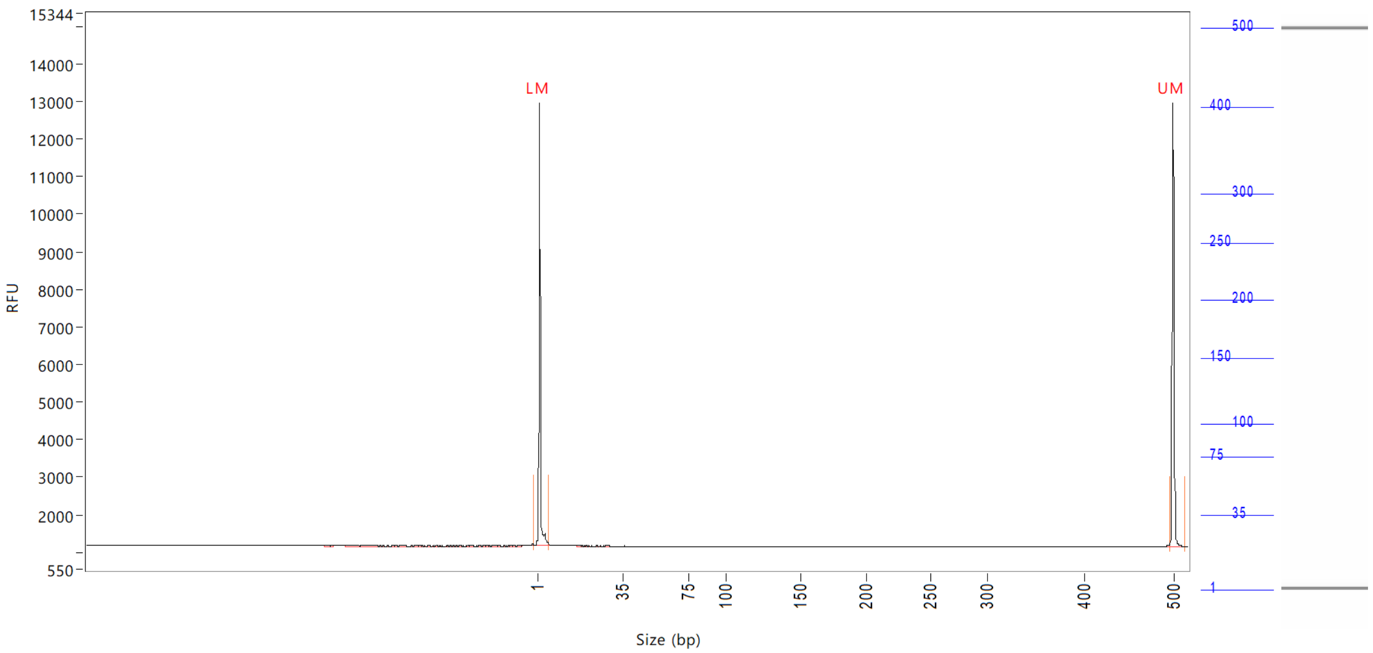


| Peak | Size (bp) | Concentration (ng/uL) | From (bp) | To (bp) | Average size (bp) | CV% | RFU | Corrected peak area |
|---|---|---|---|---|---|---|---|---|
| 1 | 1 (LM) | 0.8785 | 0 | 6 | 1 | 68.64 | 11806 | 43.760 |
| 2 | 500 (UM) | 0.5000 | 496 | 514 | 500 | 0.25 | 11819 | 24.905 |

| | | |
|---|---|---|
| TIC: | 0.0000 | ng/uL |
| TIM: | 0.0000 | nmole/L |
| Total concentration: | 0.6909 | ng/uL |

Sample peak width (sec): 5    Sample min peak height: 50    Sample baseline V to V?: Y    Sample baseline V to V points: 3
Sample filter: Binomial    Number of points for filter: 3  Sample start region (min): 0  Sample end region (min): 80
Marker peak width (sec): 5    Marker min peak height: 500    Marker baseline V to V?: Y    Marker baseline V to V points: 3
Lower marker selection: First peak > 500 RFU    Upper marker selection: Last peak > 500 RFU
Ladder size (bp)1, 35, 75, 100, 150, 200, 250, 300, 400, 500
Quantification using: Upper Marker    Final concentration (ng/uL): 0.5000    Dilution factor: 12.0

**Sample:** SampD9
**Well location:** D9
**Created:** Monday, 11 October 2021 14:41:52

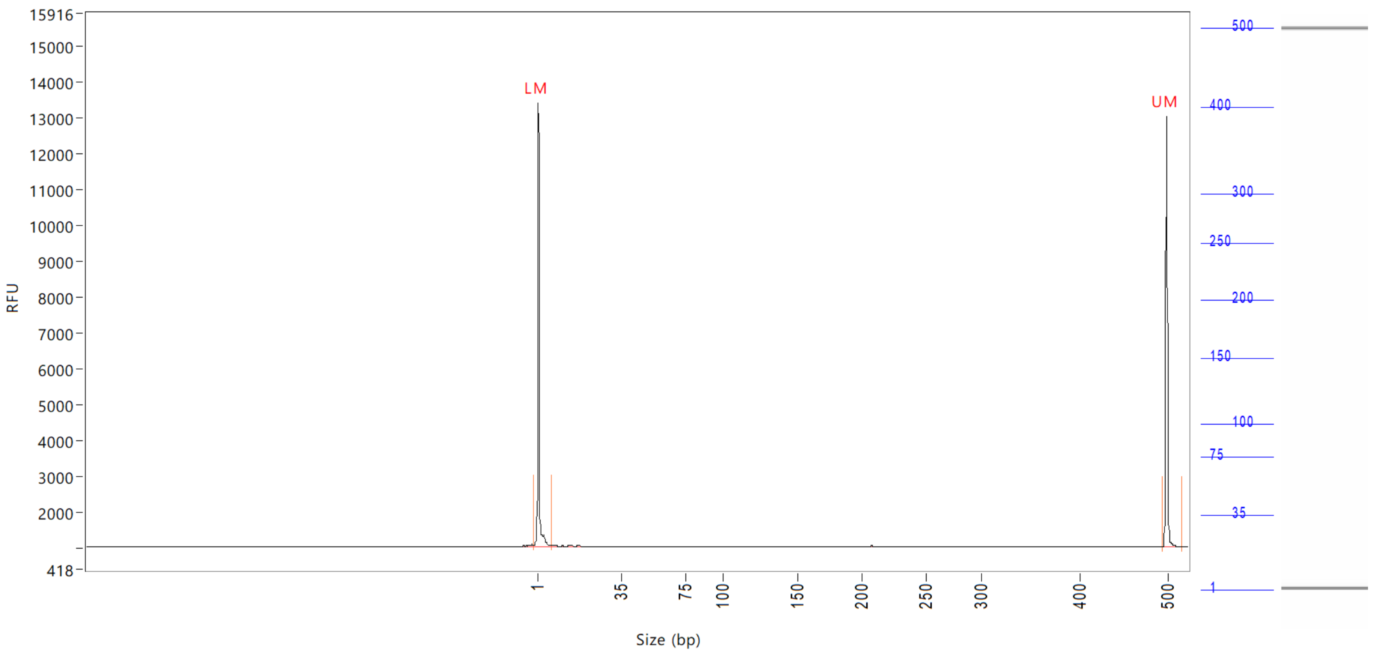


| Peak | Size (bp) | Concentration (ng/uL) | From (bp) | To (bp) | Average size (bp) | CV% | RFU | Corrected peak area |
|---|---|---|---|---|---|---|---|---|
| 1 | 1 (LM) | 0.9332 | 0 | 8 | 1 | 71.18 | 12375 | 45.367 |
| 2 | 500 (UM) | 0.5000 | 496 | 0 | 500 | 0.26 | 11976 | 24.308 |

| | | |
|---|---|---|
| TIC: | 0.0000 | ng/uL |
| TIM: | 0.0000 | nmole/L |
| Total concentration: | 0.5806 | ng/uL |

Sample peak width (sec): 5 Sample min peak height: 50 Sample baseline V to V?: Y Sample baseline V to V points: 3
Sample filter: Binomial Number of points for filter: 3 Sample start region (min): 0 Sample end region (min): 80
Marker peak width (sec): 5 Marker min peak height: 500 Marker baseline V to V?: Y Marker baseline V to V points: 3
Lower marker selection: First peak > 500 RFU Upper marker selection: Last peak > 500 RFU
Ladder size (bp)1, 35, 75, 100, 150, 200, 250, 300, 400, 500
Quantification using: Upper Marker Final concentration (ng/uL): 0.5000 Dilution factor: 12.0

**Sample:** SampE9
**Well location:** E9
**Created:** Monday, 11 October 2021 14:41:52

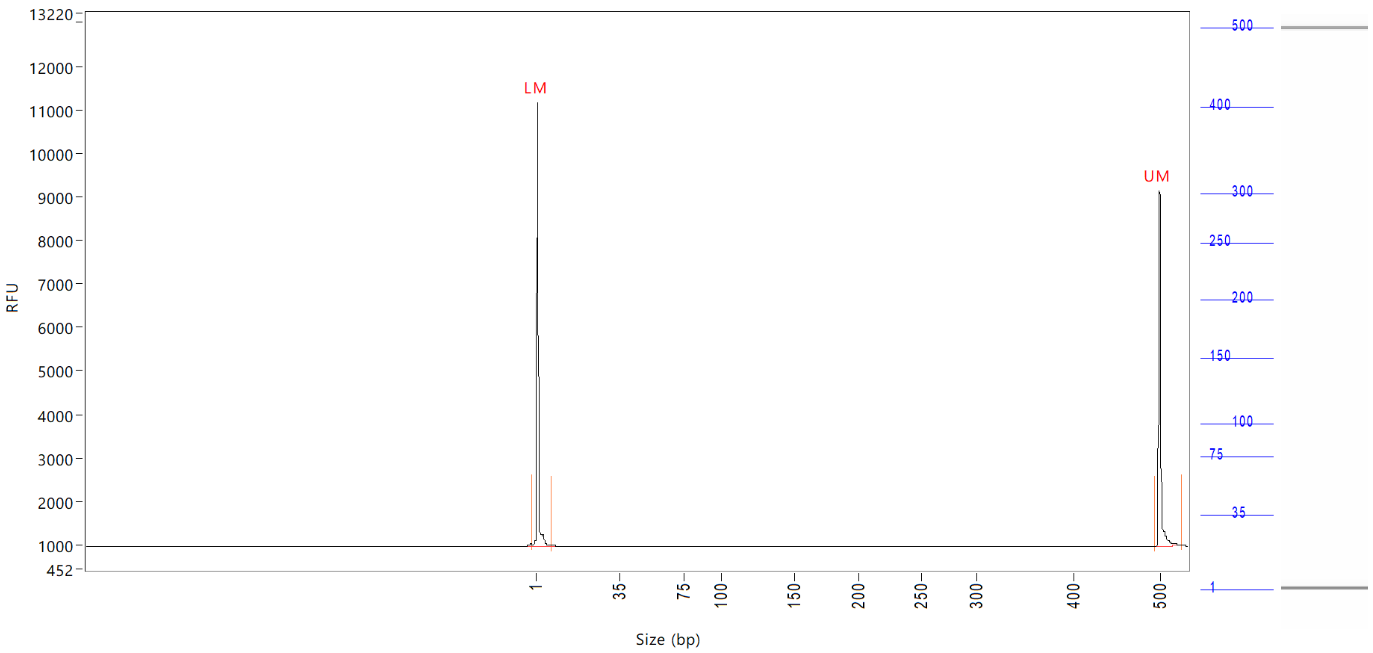


| Peak | Size (bp) | Concentration (ng/uL) | From (bp) | To (bp) | Average size (bp) | CV% | RFU | Corrected peak area |
|---|---|---|---|---|---|---|---|---|
| 1 | 1 (LM) | 1.0064 | 0 | 8 | 1 | 71.82 | 10198 | 38.155 |
| 2 | 500 (UM) | 0.5000 | 494 | 0 | 501 | 0.53 | 8165 | 18.957 |

| | | |
|---|---|---|
| TIC: | 0.0000 | ng/uL |
| TIM: | 0.0000 | nmole/L |
| Total concentration: | 0.5891 | ng/uL |

Sample peak width (sec): 5 Sample min peak height: 50 Sample baseline V to V?: Y Sample baseline V to V points: 3
Sample filter: Binomial Number of points for filter: 3 Sample start region (min): 0 Sample end region (min): 80
Marker peak width (sec): 5 Marker min peak height: 500 Marker baseline V to V?: Y Marker baseline V to V points: 3
Lower marker selection: First peak > 500 RFU Upper marker selection: Last peak > 500 RFU
Ladder size (bp)1, 35, 75, 100, 150, 200, 250, 300, 400, 500
Quantification using: Upper Marker Final concentration (ng/uL): 0.5000 Dilution factor: 12.0

**Sample:** SampF9
**Well location:** F9
**Created:** Monday, 11 October 2021 14:41:52

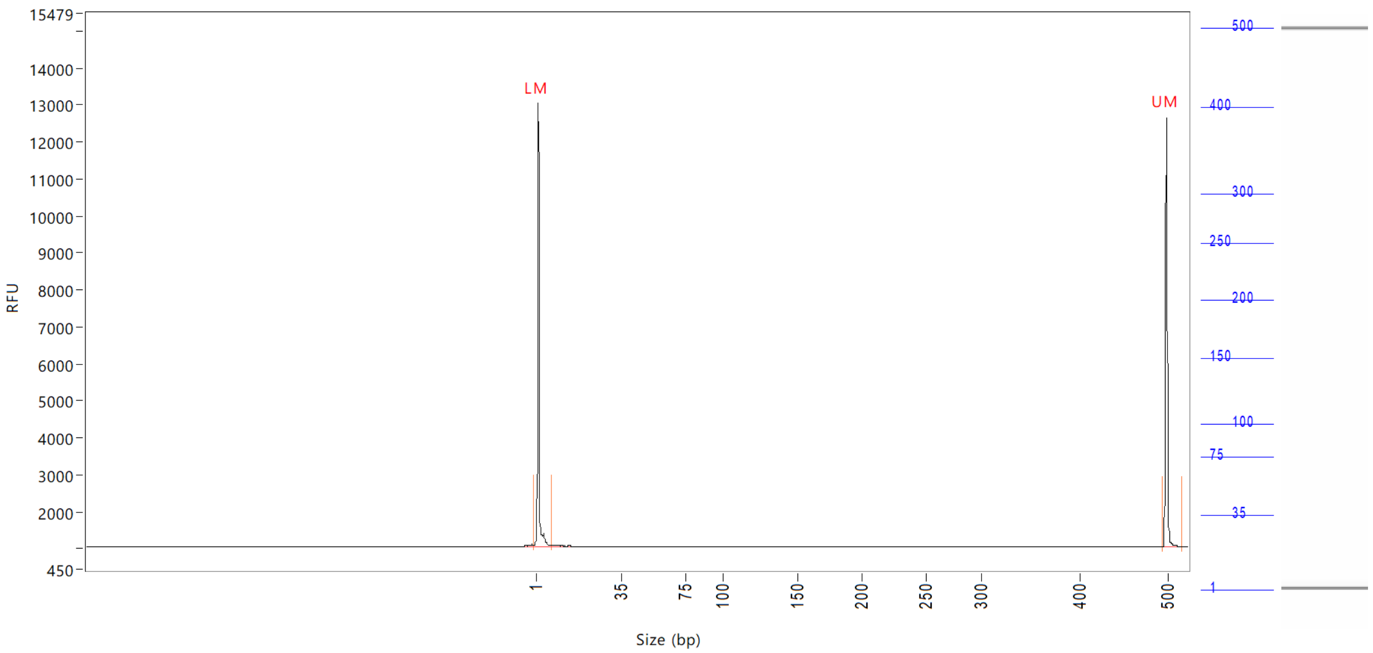


| Peak | Size (bp) | Concentration (ng/uL) | From (bp) | To (bp) | Average size (bp) | CV% | RFU | Corrected peak area |
|---|---|---|---|---|---|---|---|---|
| 1 | 1 (LM) | 0.9397 | 0 | 8 | 2 | 65.69 | 12000 | 45.275 |
| 2 | 500 (UM) | 0.5000 | 496 | 517 | 500 | 0.28 | 11592 | 24.090 |

| | | |
|---|---|---|
| TIC: | 0.0000 | ng/uL |
| TIM: | 0.0000 | nmole/L |
| Total concentration: | 0.4783 | ng/uL |

Sample peak width (sec): 5 Sample min peak height: 50 Sample baseline V to V?: Y Sample baseline V to V points: 3
Sample filter: Binomial Number of points for filter: 3 Sample start region (min): 0 Sample end region (min): 80
Marker peak width (sec): 5 Marker min peak height: 500 Marker baseline V to V?: Y Marker baseline V to V points: 3
Lower marker selection: First peak > 500 RFU Upper marker selection: Last peak > 500 RFU
Ladder size (bp)1, 35, 75, 100, 150, 200, 250, 300, 400, 500
Quantification using: Upper Marker Final concentration (ng/uL): 0.5000 Dilution factor: 12.0

**Sample:** SampG9
**Well location:** G9
**Created:** Monday, 11 October 2021 14:41:52

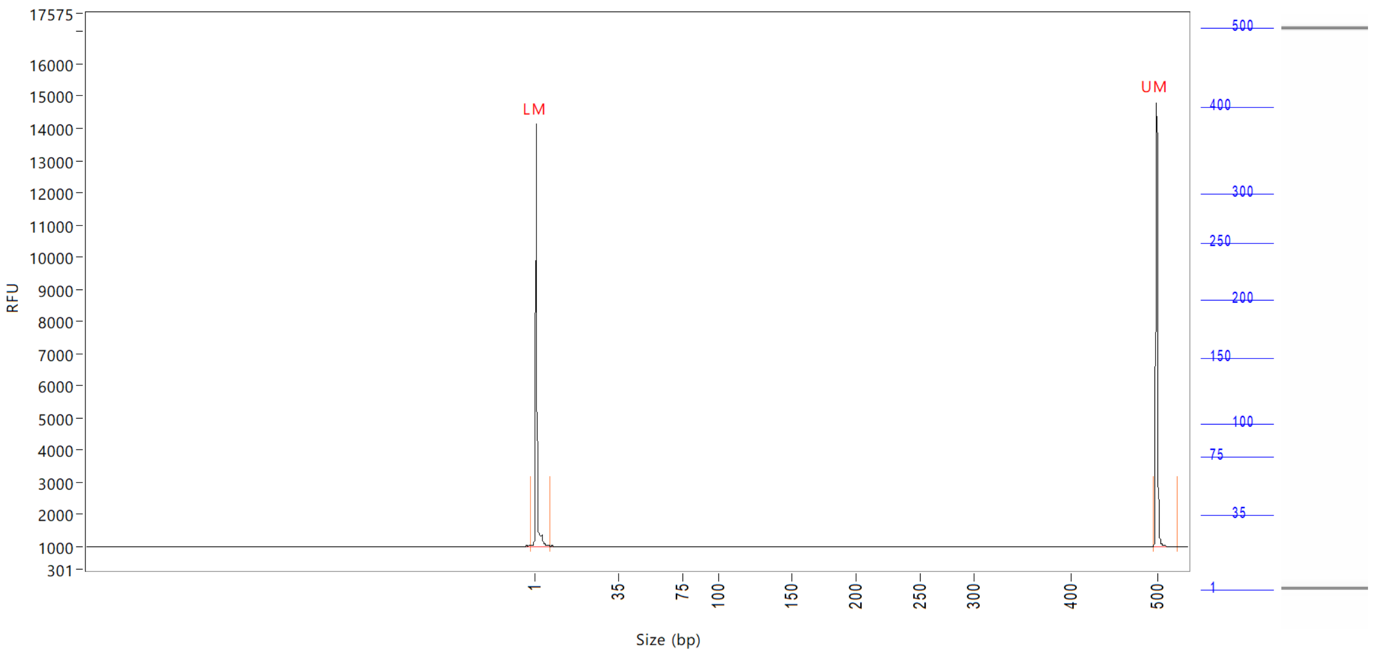


| Peak | Size (bp) | Concentration (ng/uL) | From (bp) | To (bp) | Average size (bp) | CV% | RFU | Corrected peak area |
|---|---|---|---|---|---|---|---|---|
| 1 | 1 (LM) | 0.8563 | 0 | 7 | 1 | 70.92 | 13139 | 49.762 |
| 2 | 500 (UM) | 0.5000 | 496 | 0 | 500 | 0.28 | 13809 | 29.058 |

| | | |
|---|---|---|
| TIC: | 0.0000 | ng/uL |
| TIM: | 0.0000 | nmole/L |
| Total concentration: | 0.4026 | ng/uL |

Sample peak width (sec): 5 Sample min peak height: 50 Sample baseline V to V?: Y Sample baseline V to V points: 3
Sample filter: Binomial Number of points for filter: 3 Sample start region (min): 0 Sample end region (min): 80
Marker peak width (sec): 5 Marker min peak height: 500 Marker baseline V to V?: Y Marker baseline V to V points: 3
Lower marker selection: First peak > 500 RFU Upper marker selection: Last peak > 500 RFU
Ladder size (bp)1, 35, 75, 100, 150, 200, 250, 300, 400, 500
Quantification using: Upper Marker Final concentration (ng/uL): 0.5000 Dilution factor: 12.0

**Sample:** SampH9
**Well location:** H9
**Created:** Monday, 11 October 2021 14:41:52

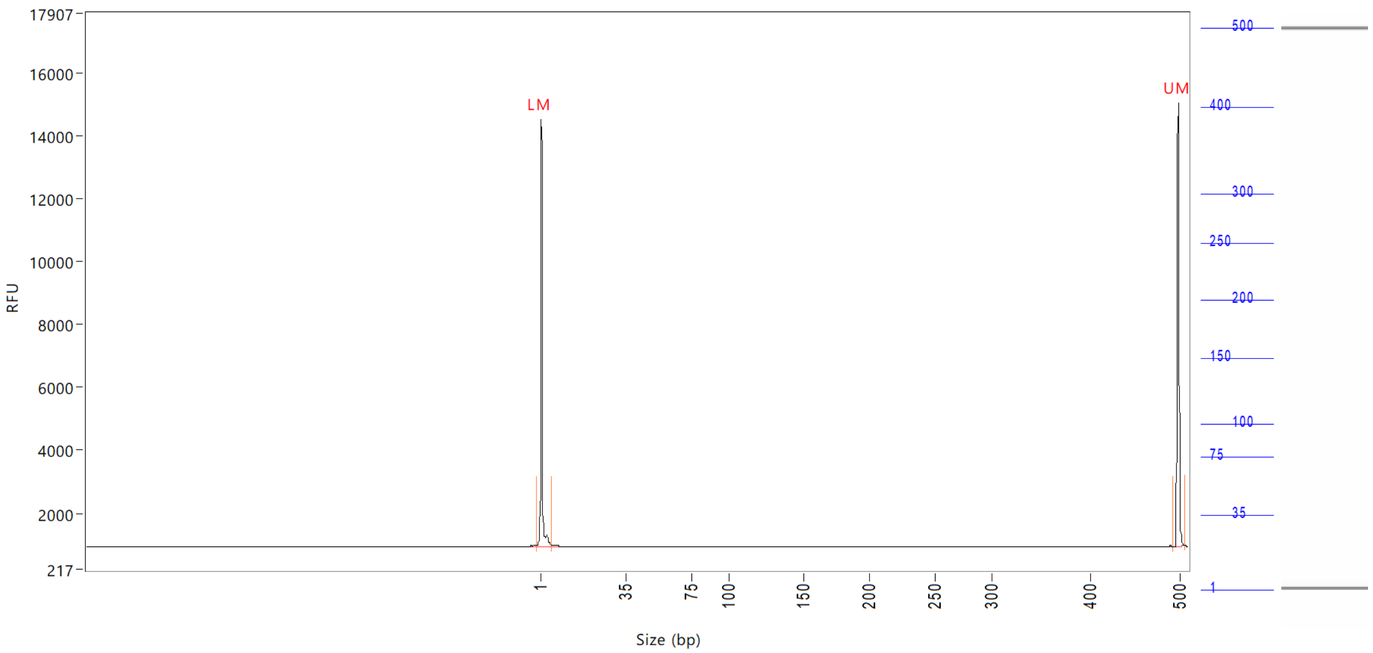


| Peak | Size (bp) | Concentration (ng/uL) | From (bp) | To (bp) | Average size (bp) | CV% | RFU | Corrected peak area |
|---|---|---|---|---|---|---|---|---|
| 1 | 1 (LM) | 0.8719 | 0 | 6 | 1 | 66.92 | 13597 | 50.984 |
| 2 | 500 (UM) | 0.5000 | 494 | 508 | 500 | 0.22 | 14129 | 29.239 |

| | | |
|---|---|---|
| TIC: | 0.0000 | ng/uL |
| TIM: | 0.0000 | nmole/L |
| Total concentration: | 0.4328 | ng/uL |

Sample peak width (sec): 5 Sample min peak height: 50 Sample baseline V to V?: Y Sample baseline V to V points: 3
Sample filter: Binomial Number of points for filter: 3 Sample start region (min): 0 Sample end region (min): 80
Marker peak width (sec): 5 Marker min peak height: 500 Marker baseline V to V?: Y Marker baseline V to V points: 3
Lower marker selection: First peak > 500 RFU Upper marker selection: Last peak > 500 RFU
Ladder size (bp)1, 35, 75, 100, 150, 200, 250, 300, 400, 500
Quantification using: Upper Marker Final concentration (ng/uL): 0.5000 Dilution factor: 12.0

**Sample:** SampA10
**Well location:** A10
**Created:** Monday, 11 October 2021 14:41:52

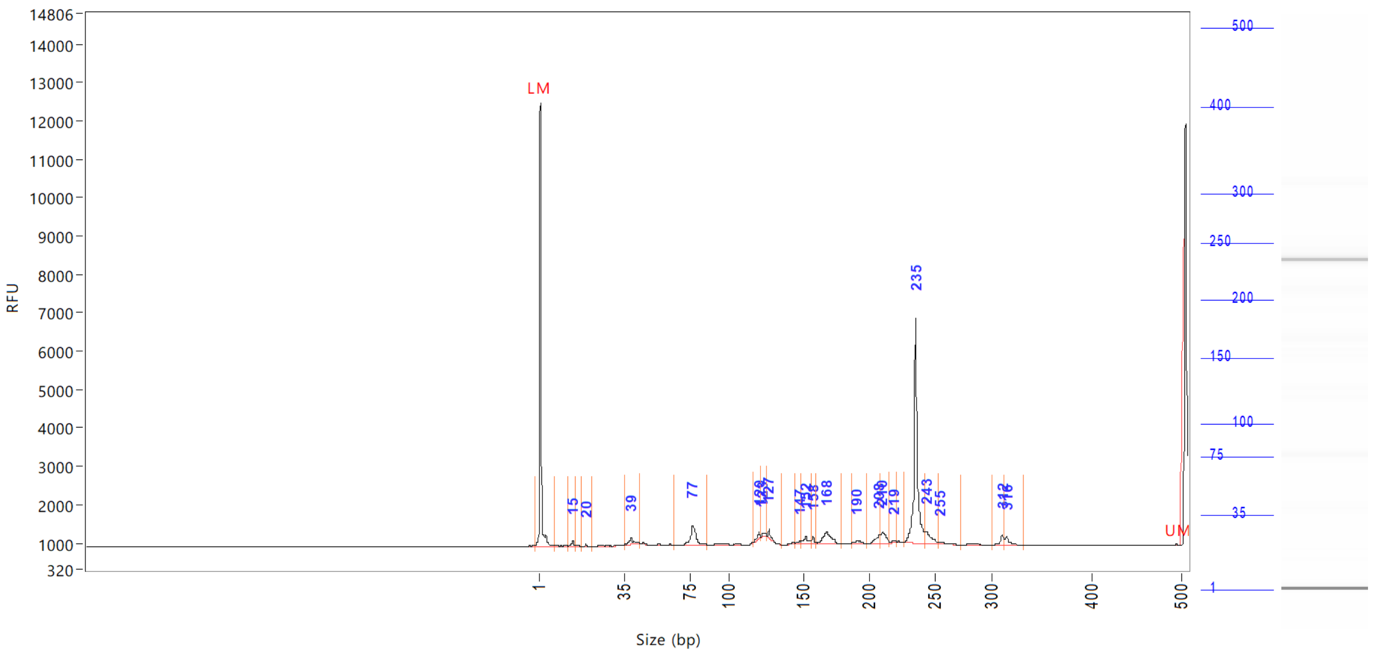


| Peak | Size (bp) | Concentration (ng/uL) | From (bp) | To (bp) | Average size (bp) | CV% | RFU | Corrected peak area |
|---|---|---|---|---|---|---|---|---|
| 1 | 1 (LM) | Inf | 0 | 7 | 1 | 71.69 | 11573 | 44.237 |
| 2 | 15 | Inf | 13 | 16 | 14 | 3.68 | 128 | 0.804 |
| 3 | 20 | Inf | 18 | 22 | 20 | 2.26 | 52 | 0.186 |
| 4 | 39 | Inf | 36 | 45 | 40 | 4.88 | 169 | 1.712 |
| 5 | 77 | Inf | 65 | 86 | 77 | 3.53 | 521 | 5.549 |
| 6 | 120 | Inf | 117 | 122 | 120 | 0.81 | 175 | 0.766 |
| 7 | 123 | Inf | 122 | 126 | 124 | 0.75 | 96 | 0.579 |
| 8 | 127 | Inf | 126 | 136 | 128 | 0.88 | 232 | 0.837 |
| 9 | 147 | Inf | 144 | 148 | 146 | 0.76 | 65 | 0.378 |
| 10 | 152 | Inf | 148 | 156 | 152 | 1.41 | 207 | 1.424 |
| 11 | 158 | Inf | 156 | 160 | 158 | 0.56 | 167 | 0.589 |
| 12 | 168 | Inf | 160 | 179 | 168 | 1.89 | 298 | 3.989 |
| 13 | 190 | Inf | 187 | 198 | 191 | 1.17 | 85 | 0.894 |
| 14 | 208 | Inf | 198 | 209 | 206 | 0.99 | 232 | 1.521 |
| 15 | 210 | Inf | 209 | 216 | 211 | 0.83 | 289 | 2.342 |
| 16 | 219 | Inf | 216 | 222 | 219 | 0.69 | 58 | 0.388 |
| 17 | 235 | Inf | 227 | 242 | 235 | 0.96 | 5846 | 25.987 |
| 18 | 243 | Inf | 242 | 253 | 246 | 1.10 | 322 | 2.619 |
| 19 | 255 | Inf | 253 | 274 | 257 | 1.22 | 59 | 0.446 |
| 20 | 312 | Inf | 302 | 314 | 311 | 0.71 | 280 | 1.182 |
| 21 | 500 (UM) | Inf | 314 | 332 | 318 | 1.03 | 222 | 1.154 |

| | | |
|---|---|---|
| TIC: | Inf | ng/uL |
| TIM: | Inf | nmole/L |
| Total concentration: | 0.0000 | ng/uL |

Sample peak width (sec): 5    Sample min peak height: 50    Sample baseline V to V?: Y    Sample baseline V to V points: 3
Sample filter: Binomial    Number of points for filter: 3    Sample start region (min): 0    Sample end region (min): 80
Marker peak width (sec): 5    Marker min peak height: 500    Marker baseline V to V?: Y    Marker baseline V to V points: 3
Lower marker selection: First peak > 500 RFU    Upper marker selection: Last peak > 500 RFU
Ladder size (bp)1, 35, 75, 100, 150, 200, 250, 300, 400, 500
Quantification using: Upper Marker    Final concentration (ng/uL): 0.5000    Dilution factor: 12.0

**Sample:** SampB10
**Well location:** B10
**Created:** Monday, 11 October 2021 14:41:52

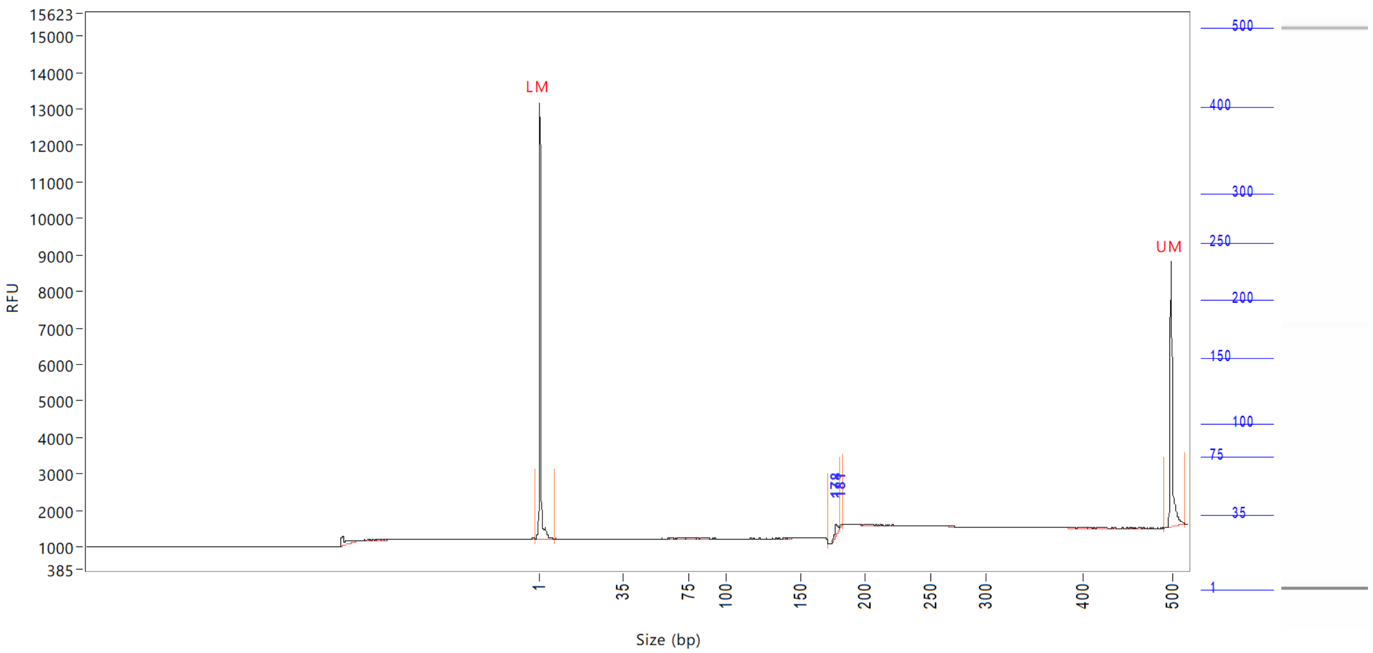


| Peak | Size (bp) | Concentration (ng/uL) | From (bp) | To (bp) | Average size (bp) | CV% | RFU | Corrected peak area |
|---|---|---|---|---|---|---|---|---|
| 1 | 1 (LM) | 1.2348 | 0 | 8 | 1 | 74.59 | 11979 | 45.044 |
| 2 | 178 | 0.5023 | 172 | 181 | 178 | 0.68 | 317 | 1.527 |
| 3 | 181 | 0.0449 | 181 | 183 | 181 | 0.26 | 63 | 0.136 |
| 4 | 500 (UM) | 0.5000 | 492 | 516 | 501 | 0.51 | 7295 | 18.240 |

| | | |
|---|---|---|
| TIC: | 0.5472 | ng/uL |
| TIM: | 5.0471 | nmole/L |
| Total concentration: | 1.4969 | ng/uL |

```
Sample peak width (sec): 5    Sample min peak height: 50    Sample baseline V to V?: Y    Sample baseline V to V points: 3
Sample filter: Binomial       Number of points for filter: 3  Sample start region (min): 0  Sample end region (min): 80
Marker peak width (sec): 5    Marker min peak height: 500   Marker baseline V to V?: Y    Marker baseline V to V points: 3
Lower marker selection: First peak > 500 RFU                Upper marker selection: Last peak > 500 RFU
Ladder size (bp)1, 35, 75, 100, 150, 200, 250, 300, 400, 500
Quantification using: Upper Marker         Final concentration (ng/uL): 0.5000         Dilution factor: 12.0
```

**Sample:** SampC10
**Well location:** C10
**Created:** Monday, 11 October 2021 14:41:52

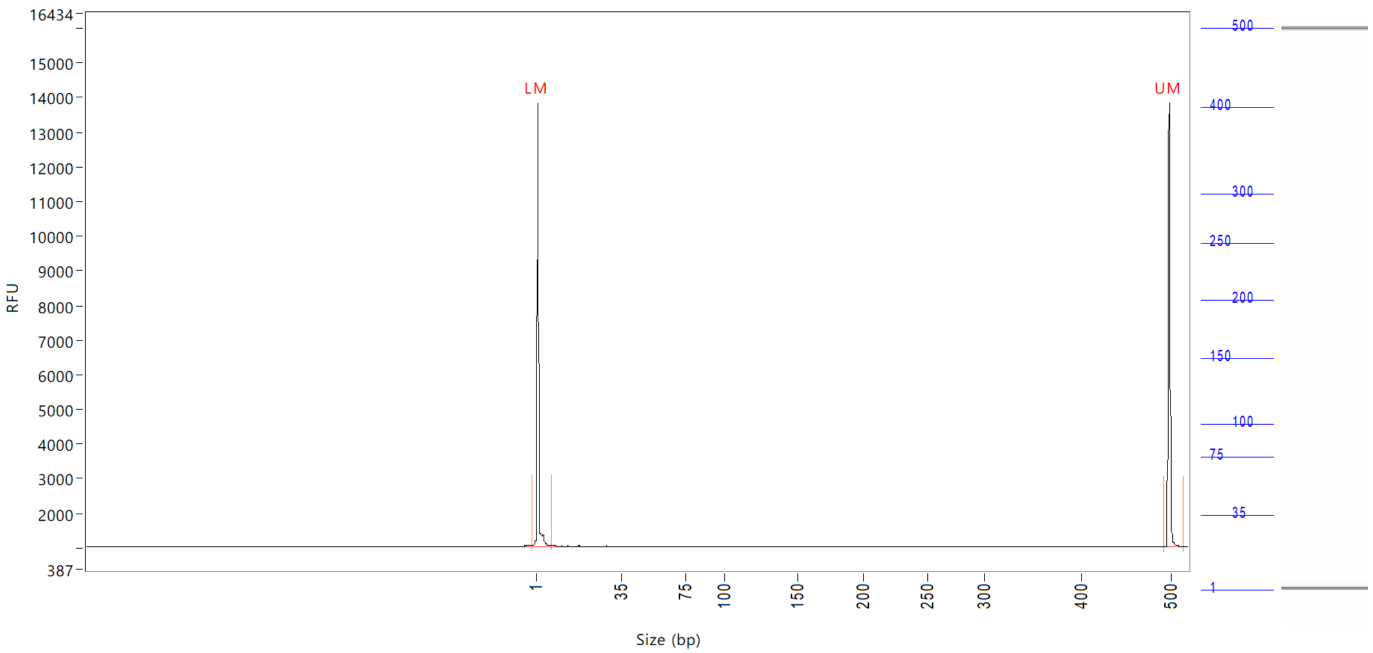


| Peak | Size<br>(bp) | Concentration<br>(ng/uL) | From<br>(bp) | To<br>(bp) | Average size<br>(bp) | CV% | RFU | Corrected peak area |
|---|---|---|---|---|---|---|---|---|
| 1 | 1 (LM) | 0.8762 | 0 | 8 | 1 | 68.84 | 12814 | 47.461 |
| 2 | 500 (UM) | 0.5000 | 493 | 516 | 500 | 0.27 | 12817 | 27.085 |

| | | |
|---|---|---|
| TIC: | 0.0000 | ng/uL |
| TIM: | 0.0000 | nmole/L |
| Total concentration: | 0.4756 | ng/uL |

Sample peak width (sec): 5 Sample min peak height: 50 Sample baseline V to V?: Y Sample baseline V to V points: 3
Sample filter: Binomial Number of points for filter: 3 Sample start region (min): 0 Sample end region (min): 80
Marker peak width (sec): 5 Marker min peak height: 500 Marker baseline V to V?: Y Marker baseline V to V points: 3
Lower marker selection: First peak > 500 RFU Upper marker selection: Last peak > 500 RFU
Ladder size (bp)1, 35, 75, 100, 150, 200, 250, 300, 400, 500
Quantification using: Upper Marker Final concentration (ng/uL): 0.5000 Dilution factor: 12.0

**Sample:** SampD10
**Well location:** D10
**Created:** Monday, 11 October 2021 14:41:52

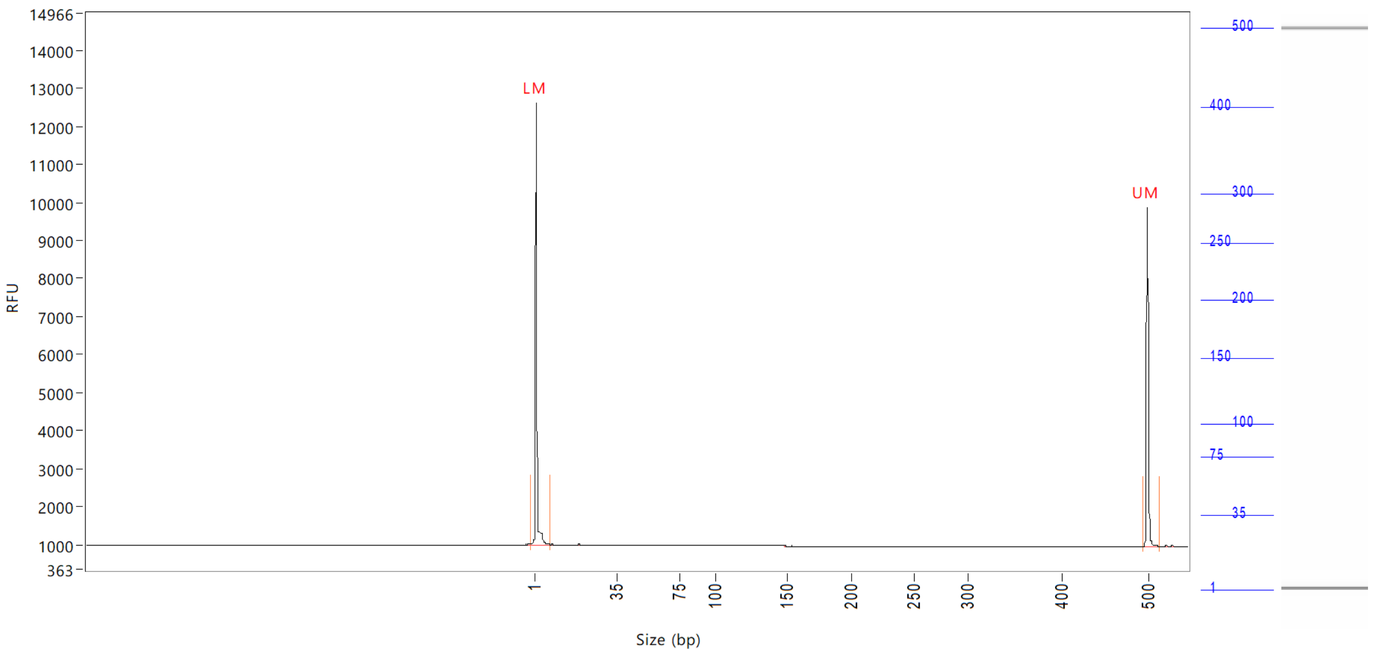


| Peak | Size (bp) | Concentration (ng/uL) | From (bp) | To (bp) | Average size (bp) | CV% | RFU | Corrected peak area |
|---|---|---|---|---|---|---|---|---|
| 1 | 1 (LM) | 1.1672 | 0 | 8 | 2 | 67.48 | 11638 | 44.926 |
| 2 | 500 (UM) | 0.5000 | 495 | 514 | 500 | 0.28 | 8890 | 19.245 |

| | | |
|---|---|---|
| TIC: | 0.0000 | ng/uL |
| TIM: | 0.0000 | nmole/L |
| Total concentration: | 0.5991 | ng/uL |

Sample peak width (sec): 5    Sample min peak height: 50    Sample baseline V to V?: Y    Sample baseline V to V points: 3
Sample filter: Binomial    Number of points for filter: 3  Sample start region (min): 0  Sample end region (min): 80
Marker peak width (sec): 5    Marker min peak height: 500    Marker baseline V to V?: Y    Marker baseline V to V points: 3
Lower marker selection: First peak > 500 RFU    Upper marker selection: Last peak > 500 RFU
Ladder size (bp)1, 35, 75, 100, 150, 200, 250, 300, 400, 500
Quantification using: Upper Marker    Final concentration (ng/uL): 0.5000    Dilution factor: 12.0

**Sample:** SampE10
**Well location:** E10
**Created:** Monday, 11 October 2021 14:41:52

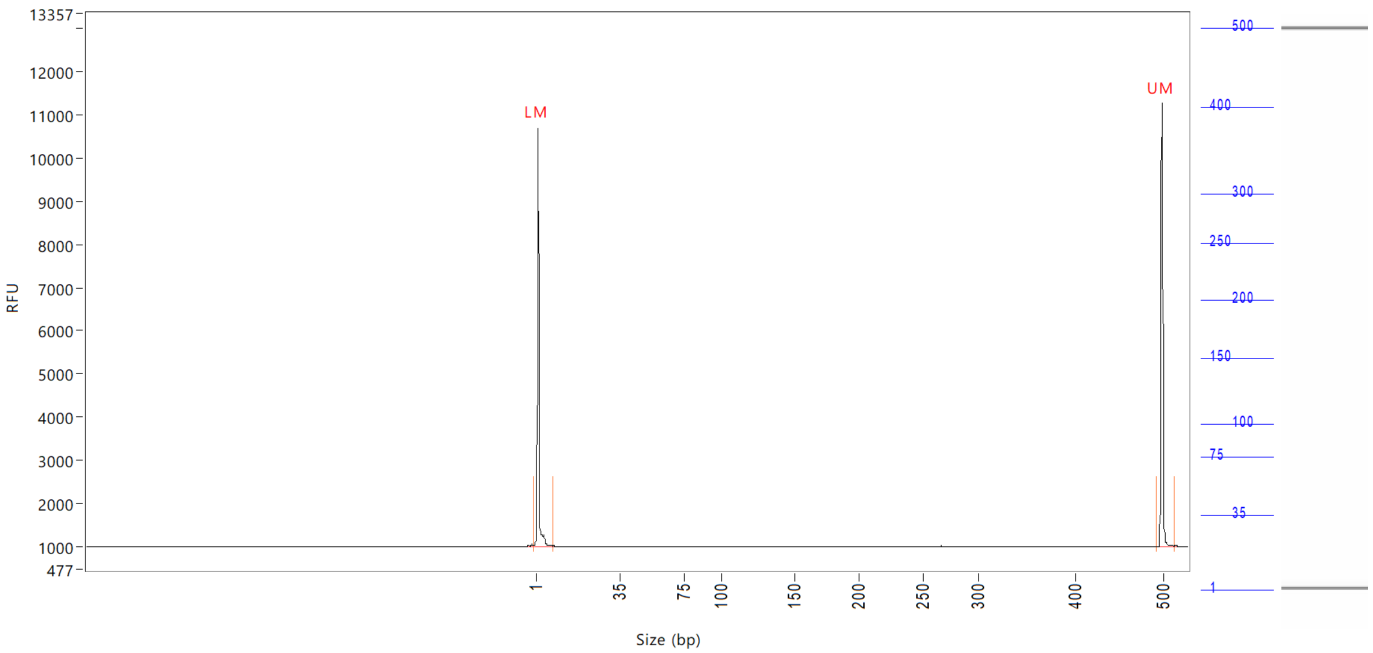


| Peak | Size | Concentration | From | To | Average size | CV% | RFU | Corrected peak area |
|---|---|---|---|---|---|---|---|---|
| | (bp) | (ng/uL) | (bp) | (bp) | (bp) | | | |
| 1 | 1 (LM) | 0.8579 | 0 | 8 | 2 | 67.04 | 9697 | 37.105 |
| 2 | 500 (UM) | 0.5000 | 494 | 514 | 500 | 0.28 | 10298 | 21.627 |

| | | |
|---|---|---|
| TIC: | 0.0000 | ng/uL |
| TIM: | 0.0000 | nmole/L |
| Total concentration: | 0.5171 | ng/uL |

```
Sample peak width (sec): 5    Sample min peak height: 50    Sample baseline V to V?: Y    Sample baseline V to V points: 3
Sample filter: Binomial       Number of points for filter: 3  Sample start region (min): 0  Sample end region (min): 80
Marker peak width (sec): 5    Marker min peak height: 500   Marker baseline V to V?: Y    Marker baseline V to V points: 3
Lower marker selection: First peak > 500 RFU                Upper marker selection: Last peak > 500 RFU
Ladder size (bp)1, 35, 75, 100, 150, 200, 250, 300, 400, 500
Quantification using: Upper Marker          Final concentration (ng/uL): 0.5000          Dilution factor: 12.0
```

**Sample:** SampF10
**Well location:** F10
**Created:** Monday, 11 October 2021 14:41:52

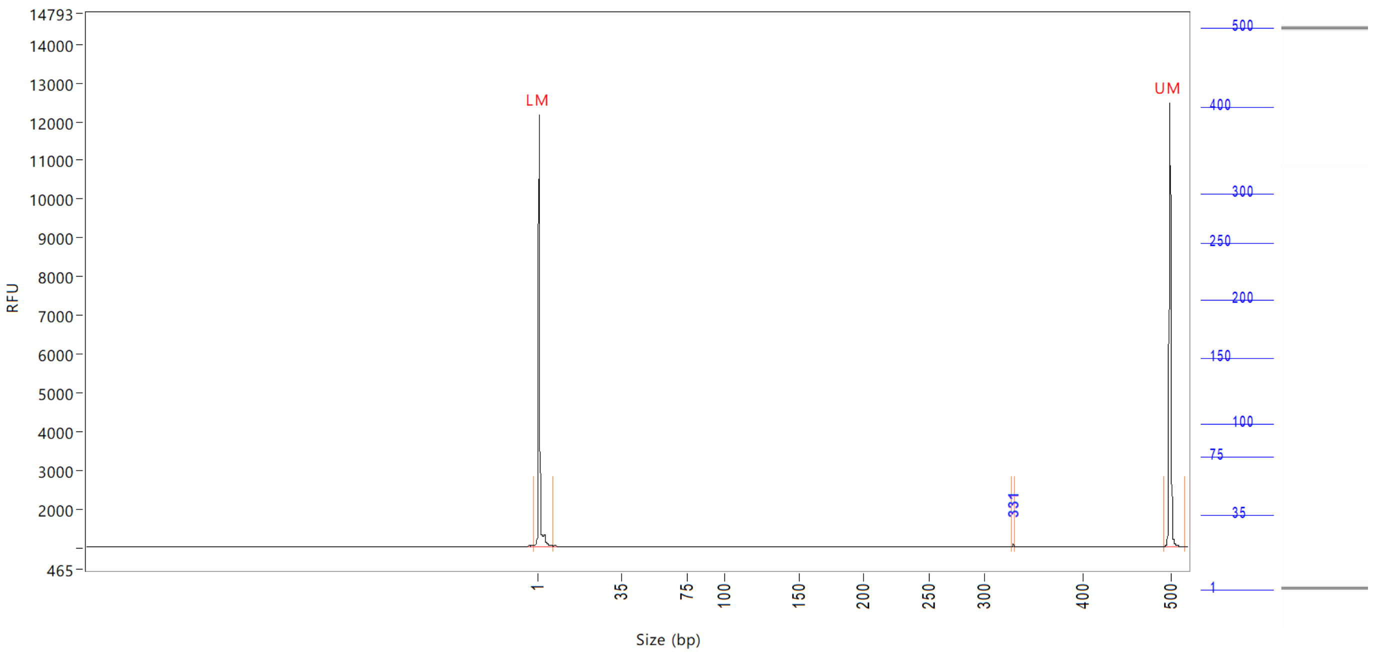


| Peak | Size (bp) | Concentration (ng/uL) | From (bp) | To (bp) | Average size (bp) | CV% | RFU | Corrected peak area |
|---|---|---|---|---|---|---|---|---|
| 1 | 1 (LM) | 0.8556 | 0 | 8 | 1 | 68.34 | 11101 | 41.971 |
| 2 | 331 | 0.0246 | 329 | 333 | 330 | 0.16 | 68 | 0.100 |
| 3 | 500 (UM) | 0.5000 | 492 | 0 | 500 | 0.27 | 11457 | 24.526 |

| | | |
|---|---|---|
| TIC: | 0.0246 | ng/uL |
| TIM: | 0.1224 | nmole/L |
| Total concentration: | 0.4950 | ng/uL |

Sample peak width (sec): 5    Sample min peak height: 50    Sample baseline V to V?: Y    Sample baseline V to V points: 3
Sample filter: Binomial    Number of points for filter: 3    Sample start region (min): 0    Sample end region (min): 80
Marker peak width (sec): 5    Marker min peak height: 500    Marker baseline V to V?: Y    Marker baseline V to V points: 3
Lower marker selection: First peak > 500 RFU    Upper marker selection: Last peak > 500 RFU
Ladder size (bp)1, 35, 75, 100, 150, 200, 250, 300, 400, 500
Quantification using: Upper Marker    Final concentration (ng/uL): 0.5000    Dilution factor: 12.0

**Sample:** SampG10
**Well location:** G10
**Created:** Monday, 11 October 2021 14:41:52

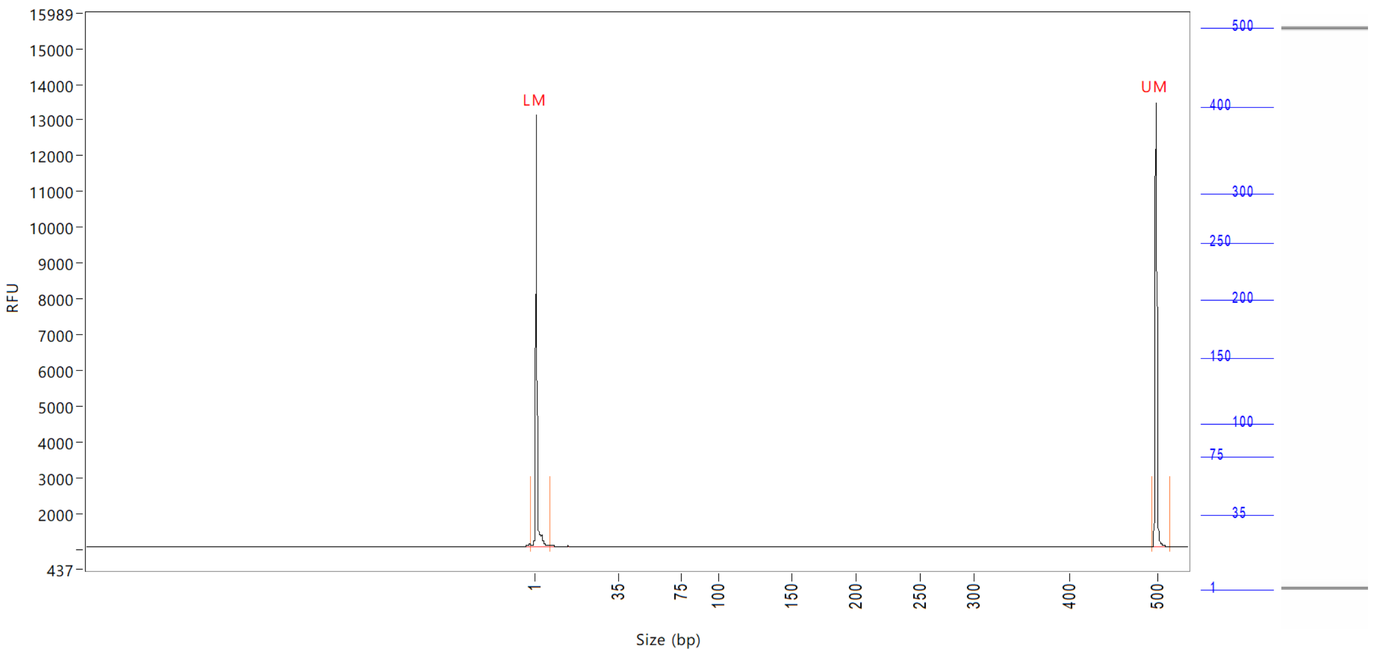


| Peak | Size (bp) | Concentration (ng/uL) | From (bp) | To (bp) | Average size (bp) | CV% | RFU | Corrected peak area |
|---|---|---|---|---|---|---|---|---|
| 1 | 1 (LM) | 0.8825 | 0 | 8 | 2 | 66.75 | 12088 | 45.938 |
| 2 | 500 (UM) | 0.5000 | 495 | 515 | 500 | 0.28 | 12434 | 26.028 |

| | | |
|---|---|---|
| TIC: | 0.0000 | ng/uL |
| TIM: | 0.0000 | nmole/L |
| Total concentration: | 0.4784 | ng/uL |

```
Sample peak width (sec): 5    Sample min peak height: 50    Sample baseline V to V?: Y    Sample baseline V to V points: 3
Sample filter: Binomial       Number of points for filter: 3  Sample start region (min): 0  Sample end region (min): 80
Marker peak width (sec): 5    Marker min peak height: 500   Marker baseline V to V?: Y    Marker baseline V to V points: 3
Lower marker selection: First peak > 500 RFU                Upper marker selection: Last peak > 500 RFU
Ladder size (bp)1, 35, 75, 100, 150, 200, 250, 300, 400, 500
Quantification using: Upper Marker         Final concentration (ng/uL): 0.5000         Dilution factor: 12.0
```

**Sample:** SampH10
**Well location:** H10
**Created:** Monday, 11 October 2021 14:41:52

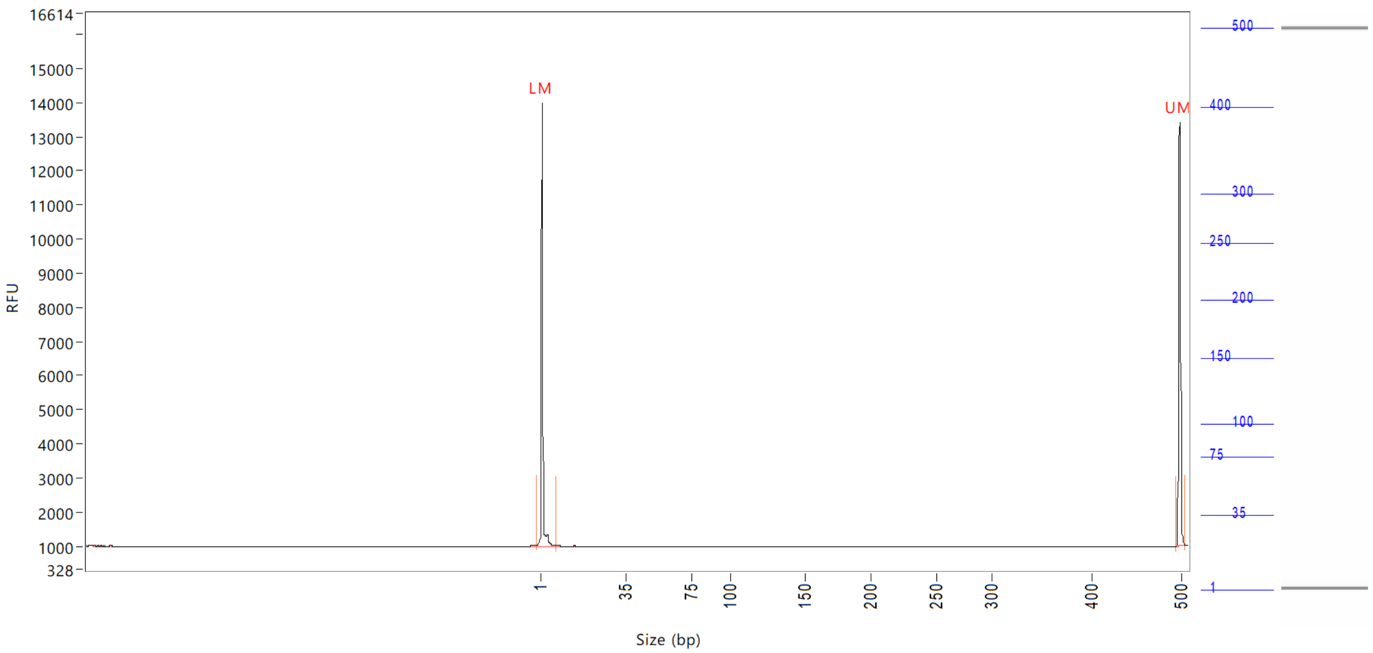


| Peak | Size (bp) | Concentration (ng/uL) | From (bp) | To (bp) | Average size (bp) | CV% | RFU | Corrected peak area |
|---|---|---|---|---|---|---|---|---|
| 1 | 1 (LM) | 0.9600 | 0 | 8 | 1 | 67.25 | 13007 | 49.456 |
| 2 | 500 (UM) | 0.5000 | 496 | 507 | 500 | 0.22 | 12418 | 25.758 |

| | | |
|---|---|---|
| TIC: | 0.0000 | ng/uL |
| TIM: | 0.0000 | nmole/L |
| Total concentration: | 0.4665 | ng/uL |

Sample peak width (sec): 5    Sample min peak height: 50    Sample baseline V to V?: Y    Sample baseline V to V points: 3
Sample filter: Binomial    Number of points for filter: 3  Sample start region (min): 0  Sample end region (min): 80
Marker peak width (sec): 5    Marker min peak height: 500   Marker baseline V to V?: Y    Marker baseline V to V points: 3
Lower marker selection: First peak > 500 RFU    Upper marker selection: Last peak > 500 RFU
Ladder size (bp)1, 35, 75, 100, 150, 200, 250, 300, 400, 500
Quantification using: Upper Marker    Final concentration (ng/uL): 0.5000    Dilution factor: 12.0

**Sample:** SampA11
**Well location:** A11
**Created:** Monday, 11 October 2021 14:41:52

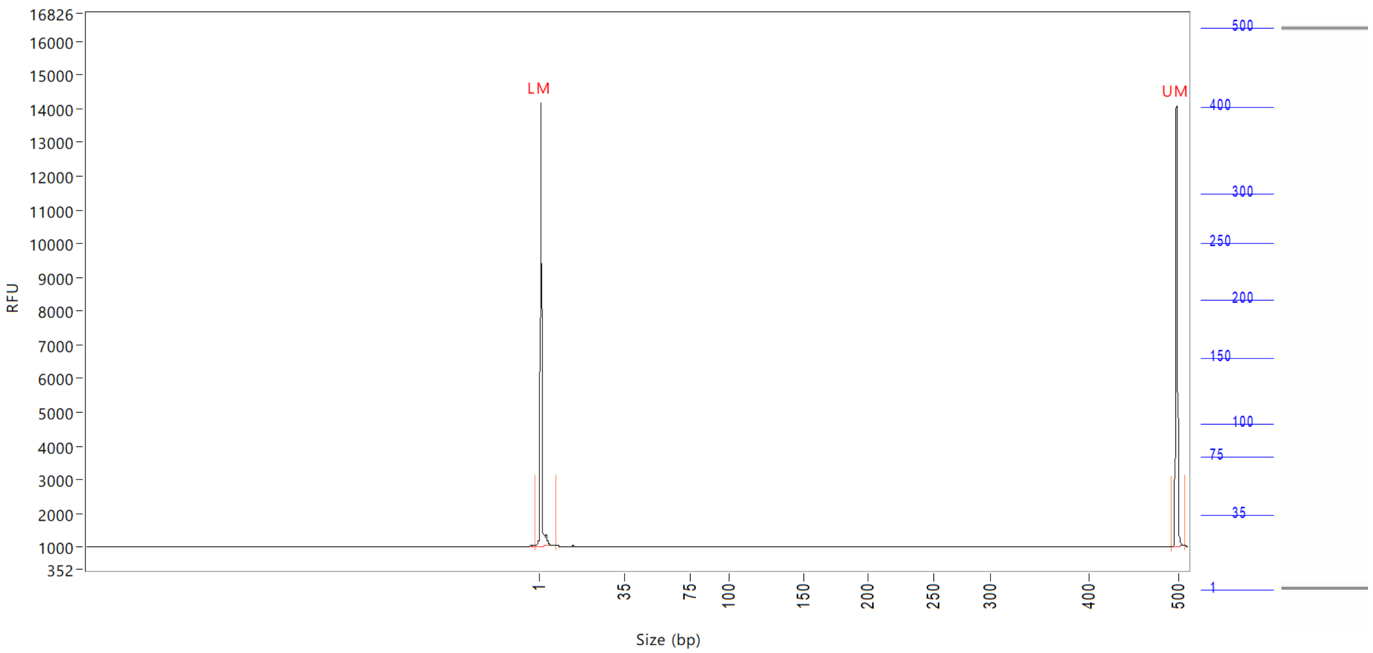


| Peak | Size (bp) | Concentration (ng/uL) | From (bp) | To (bp) | Average size (bp) | CV% | RFU | Corrected peak area |
|---|---|---|---|---|---|---|---|---|
| 1 | 1 (LM) | 0.9012 | 0 | 8 | 1 | 70.57 | 13152 | 48.897 |
| 2 | 500 (UM) | 0.5000 | 493 | 510 | 500 | 0.22 | 13047 | 27.128 |

| | | |
|---|---|---|
| TIC: | 0.0000 | ng/uL |
| TIM: | 0.0000 | nmole/L |
| Total concentration: | 0.4099 | ng/uL |

Sample peak width (sec): 5 Sample min peak height: 50 Sample baseline V to V?: Y Sample baseline V to V points: 3
Sample filter: Binomial Number of points for filter: 3 Sample start region (min): 0 Sample end region (min): 80
Marker peak width (sec): 5 Marker min peak height: 500 Marker baseline V to V?: Y Marker baseline V to V points: 3
Lower marker selection: First peak > 500 RFU Upper marker selection: Last peak > 500 RFU
Ladder size (bp)1, 35, 75, 100, 150, 200, 250, 300, 400, 500
Quantification using: Upper Marker Final concentration (ng/uL): 0.5000 Dilution factor: 12.0

**Sample:** SampB11
**Well location:** B11
**Created:** Monday, 11 October 2021 14:41:52

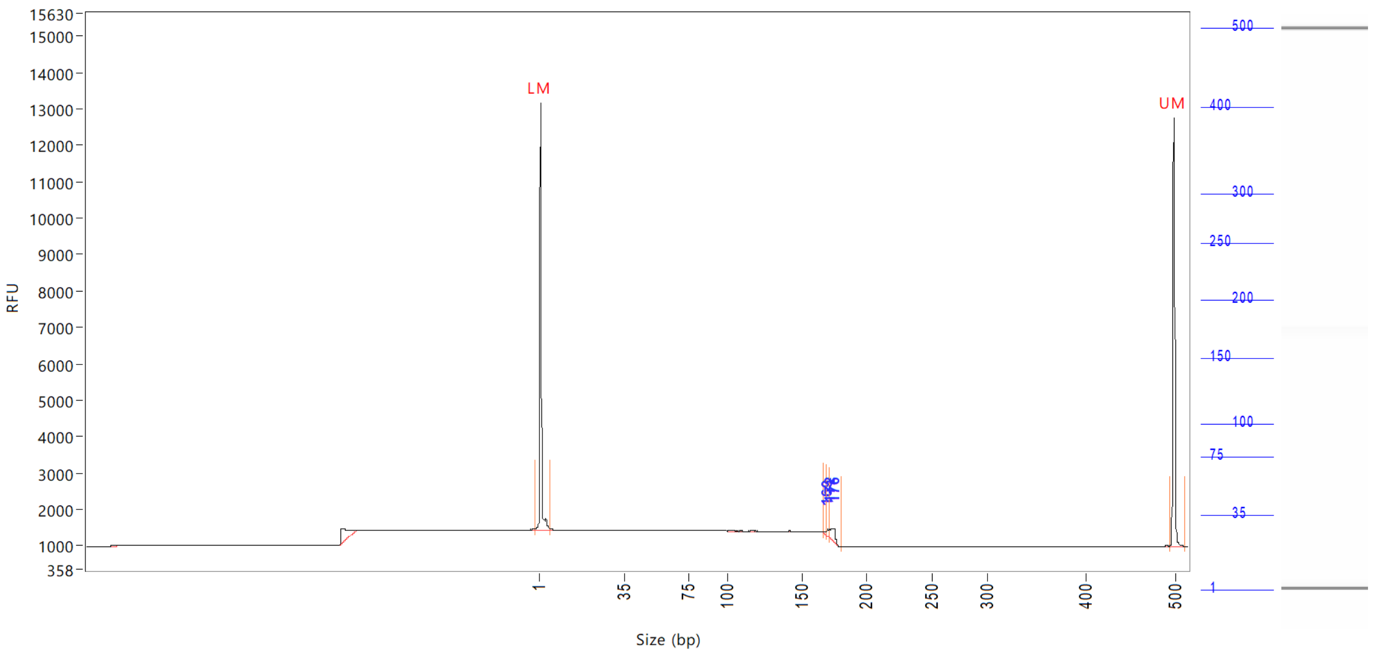


| Peak | Size (bp) | Concentration (ng/uL) | From (bp) | To (bp) | Average size (bp) | CV% | RFU | Corrected peak area |
|---|---|---|---|---|---|---|---|---|
| 1 | 1 (LM) | 0.8761 | 0 | 6 | 1 | 68.15 | 11756 | 43.553 |
| 2 | 169 | 0.0750 | 167 | 170 | 169 | 0.37 | 100 | 0.311 |
| 3 | 172 | 0.2528 | 170 | 172 | 171 | 0.50 | 237 | 1.047 |
| 4 | 176 | 0.5910 | 172 | 181 | 175 | 0.74 | 366 | 2.448 |
| 5 | 500 (UM) | 0.5000 | 495 | 513 | 500 | 0.25 | 11764 | 24.857 |

| | | |
|---|---|---|
| TIC: | 0.9188 | ng/uL |
| TIM: | 8.7118 | nmole/L |
| Total concentration: | 1.4991 | ng/uL |

Sample peak width (sec): 5 Sample min peak height: 50 Sample baseline V to V?: Y Sample baseline V to V points: 3
Sample filter: Binomial Number of points for filter: 3 Sample start region (min): 0 Sample end region (min): 80
Marker peak width (sec): 5 Marker min peak height: 500 Marker baseline V to V?: Y Marker baseline V to V points: 3
Lower marker selection: First peak > 500 RFU Upper marker selection: Last peak > 500 RFU
Ladder size (bp)1, 35, 75, 100, 150, 200, 250, 300, 400, 500
Quantification using: Upper Marker Final concentration (ng/uL): 0.5000 Dilution factor: 12.0

**Sample:** SampC11
**Well location:** C11
**Created:** Monday, 11 October 2021 14:41:52

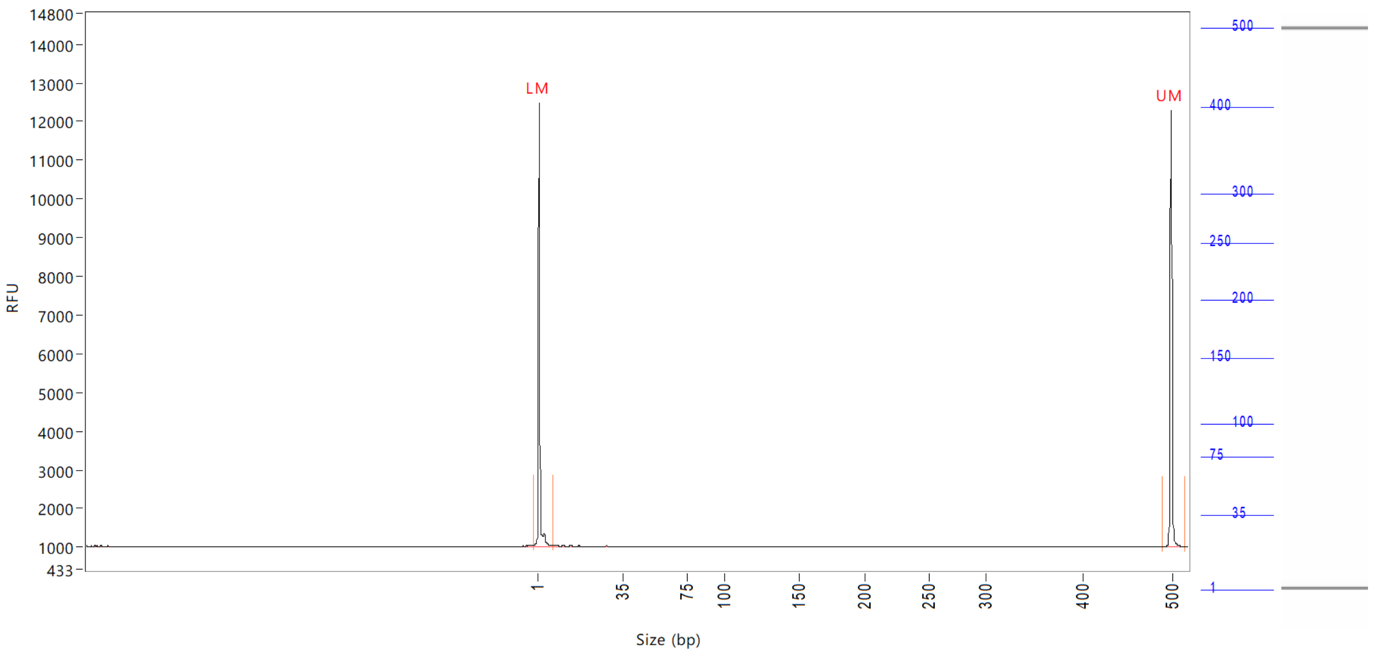


| Peak | Size (bp) | Concentration (ng/uL) | From (bp) | To (bp) | Average size (bp) | CV% | RFU | Corrected peak area |
|---|---|---|---|---|---|---|---|---|
| 1 | 1 (LM) | 0.9082 | 0 | 8 | 1 | 67.09 | 11471 | 42.878 |
| 2 | 500 (UM) | 0.5000 | 491 | 516 | 500 | 0.27 | 11280 | 23.607 |

| | | |
|---|---|---|
| TIC: | 0.0000 | ng/uL |
| TIM: | 0.0000 | nmole/L |
| Total concentration: | 0.5787 | ng/uL |

Sample peak width (sec): 5    Sample min peak height: 50    Sample baseline V to V?: Y    Sample baseline V to V points: 3
Sample filter: Binomial    Number of points for filter: 3  Sample start region (min): 0  Sample end region (min): 80
Marker peak width (sec): 5    Marker min peak height: 500    Marker baseline V to V?: Y    Marker baseline V to V points: 3
Lower marker selection: First peak > 500 RFU    Upper marker selection: Last peak > 500 RFU
Ladder size (bp)1, 35, 75, 100, 150, 200, 250, 300, 400, 500
Quantification using: Upper Marker    Final concentration (ng/uL): 0.5000    Dilution factor: 12.0

**Sample:** SampD11
**Well location:** D11
**Created:** Monday, 11 October 2021 14:41:52

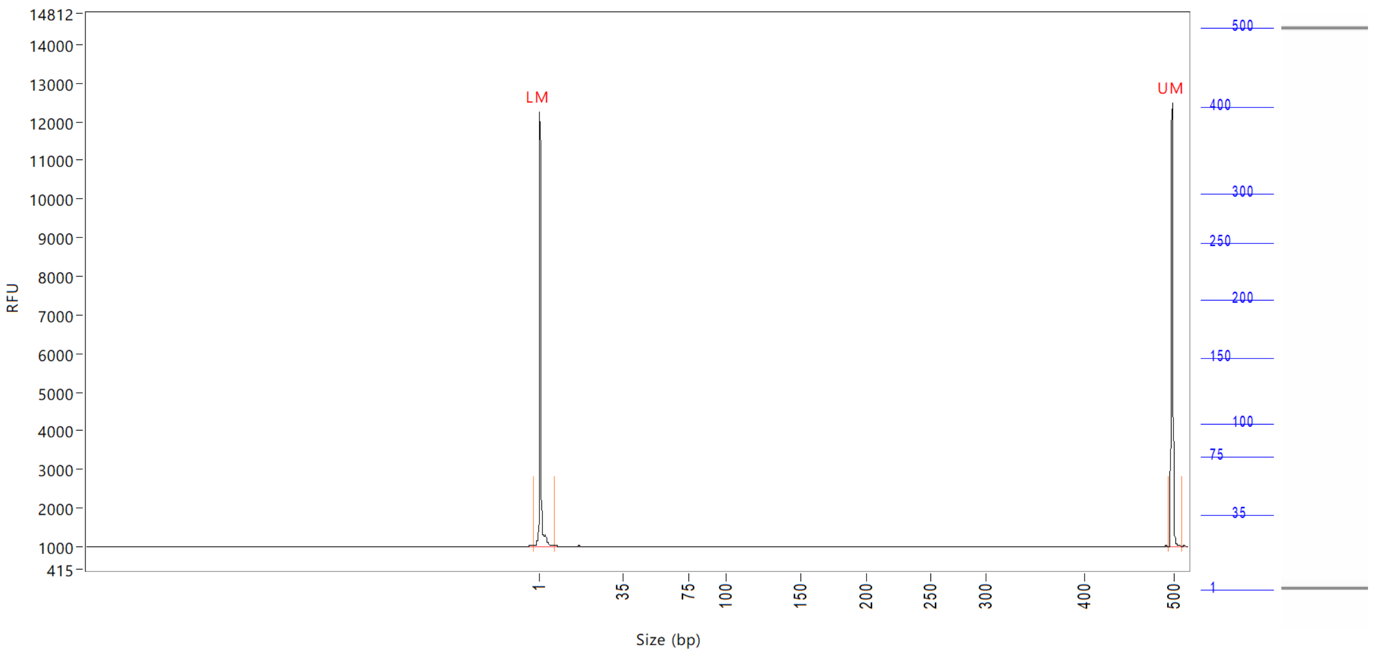


| Peak | Size (bp) | Concentration (ng/uL) | From (bp) | To (bp) | Average size (bp) | CV% | RFU | Corrected peak area |
|---|---|---|---|---|---|---|---|---|
| 1 | 1 (LM) | 0.9046 | 0 | 8 | 1 | 72.37 | 11256 | 42.475 |
| 2 | 500 (UM) | 0.5000 | 496 | 511 | 500 | 0.23 | 11503 | 23.479 |

| | | |
|---|---|---|
| TIC: | 0.0000 | ng/uL |
| TIM: | 0.0000 | nmole/L |
| Total concentration: | 0.5462 | ng/uL |

Sample peak width (sec): 5  Sample min peak height: 50  Sample baseline V to V?: Y  Sample baseline V to V points: 3
Sample filter: Binomial  Number of points for filter: 3  Sample start region (min): 0  Sample end region (min): 80
Marker peak width (sec): 5  Marker min peak height: 500  Marker baseline V to V?: Y  Marker baseline V to V points: 3
Lower marker selection: First peak > 500 RFU  Upper marker selection: Last peak > 500 RFU
Ladder size (bp)1, 35, 75, 100, 150, 200, 250, 300, 400, 500
Quantification using: Upper Marker  Final concentration (ng/uL): 0.5000  Dilution factor: 12.0

**Sample:** SampE11
**Well location:** E11
**Created:** Monday, 11 October 2021 14:41:52

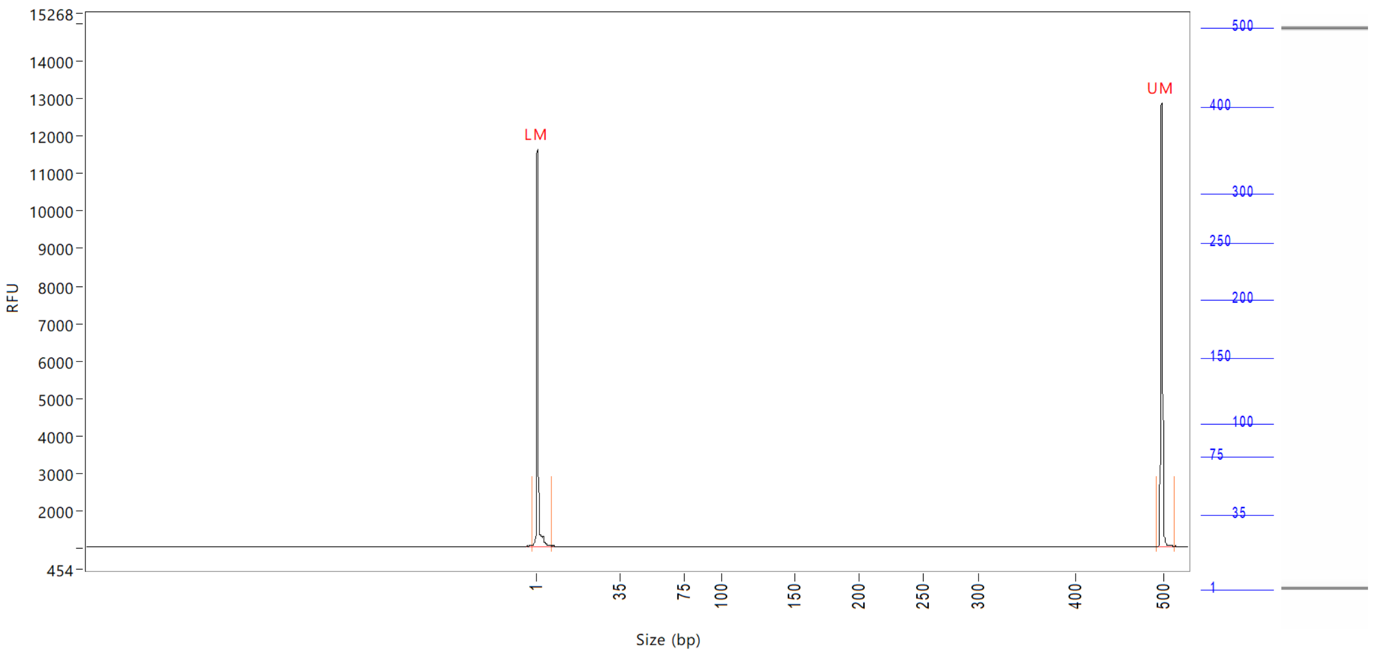


| Peak | Size (bp) | Concentration (ng/uL) | From (bp) | To (bp) | Average size (bp) | CV% | RFU | Corrected peak area |
|---|---|---|---|---|---|---|---|---|
| 1 | 1 (LM) | 0.8282 | 0 | 8 | 1 | 74.47 | 10572 | 41.065 |
| 2 | 500 (UM) | 0.5000 | 494 | 514 | 500 | 0.27 | 11845 | 24.792 |

| | | |
|---|---|---|
| TIC: | 0.0000 | ng/uL |
| TIM: | 0.0000 | nmole/L |
| Total concentration: | 0.4534 | ng/uL |

Sample peak width (sec): 5    Sample min peak height: 50    Sample baseline V to V?: Y    Sample baseline V to V points: 3
Sample filter: Binomial    Number of points for filter: 3  Sample start region (min): 0  Sample end region (min): 80
Marker peak width (sec): 5    Marker min peak height: 500   Marker baseline V to V?: Y    Marker baseline V to V points: 3
Lower marker selection: First peak > 500 RFU    Upper marker selection: Last peak > 500 RFU
Ladder size (bp)1, 35, 75, 100, 150, 200, 250, 300, 400, 500
Quantification using: Upper Marker    Final concentration (ng/uL): 0.5000    Dilution factor: 12.0

**Sample:** SampF11
**Well location:** F11
**Created:** Monday, 11 October 2021 14:41:52

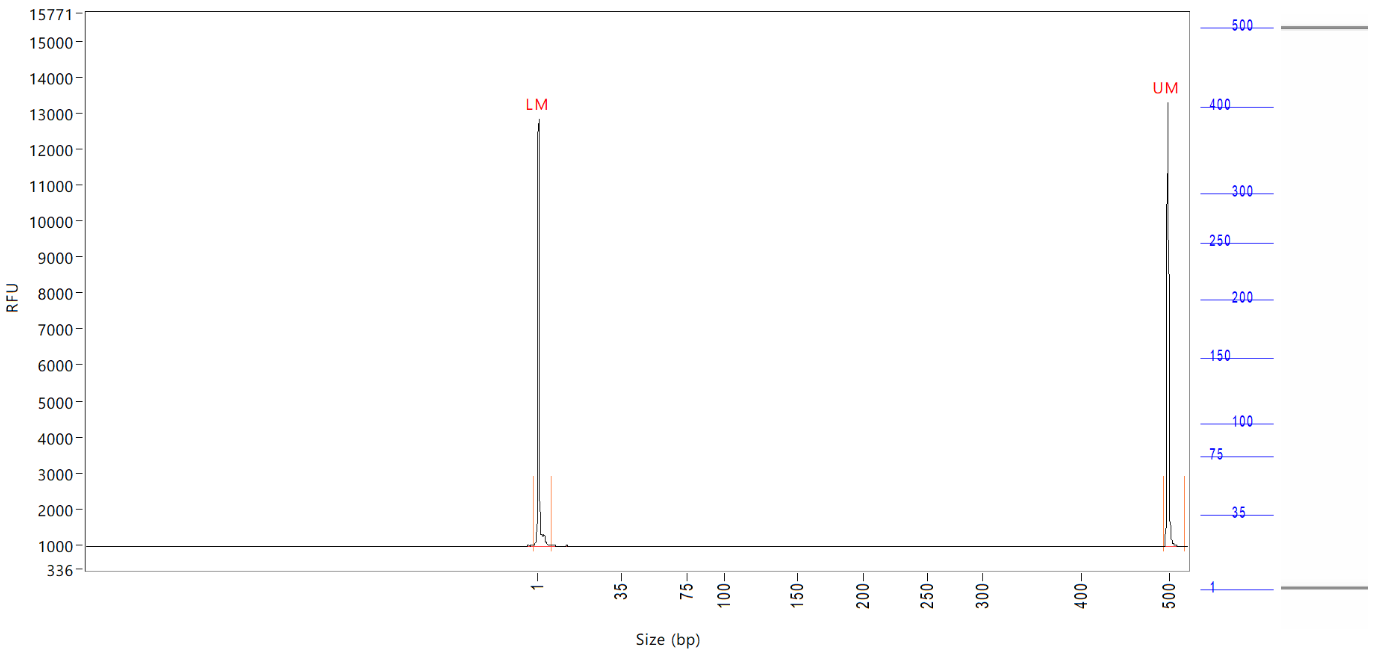


| Peak | Size (bp) | Concentration (ng/uL) | From (bp) | To (bp) | Average size (bp) | CV% | RFU | Corrected peak area |
|---|---|---|---|---|---|---|---|---|
| 1 | 1 (LM) | 0.8699 | 0 | 7 | 1 | 71.56 | 11861 | 44.882 |
| 2 | 500 (UM) | 0.5000 | 495 | 0 | 500 | 0.27 | 12343 | 25.796 |

| | | |
|---|---|---|
| TIC: | 0.0000 | ng/uL |
| TIM: | 0.0000 | nmole/L |
| Total concentration: | 0.4553 | ng/uL |

Sample peak width (sec): 5    Sample min peak height: 50    Sample baseline V to V?: Y    Sample baseline V to V points: 3
Sample filter: Binomial    Number of points for filter: 3  Sample start region (min): 0  Sample end region (min): 80
Marker peak width (sec): 5    Marker min peak height: 500    Marker baseline V to V?: Y    Marker baseline V to V points: 3
Lower marker selection: First peak > 500 RFU    Upper marker selection: Last peak > 500 RFU
Ladder size (bp)1, 35, 75, 100, 150, 200, 250, 300, 400, 500
Quantification using: Upper Marker    Final concentration (ng/uL): 0.5000    Dilution factor: 12.0

**Sample:** SampG11
**Well location:** G11
**Created:** Monday, 11 October 2021 14:41:52

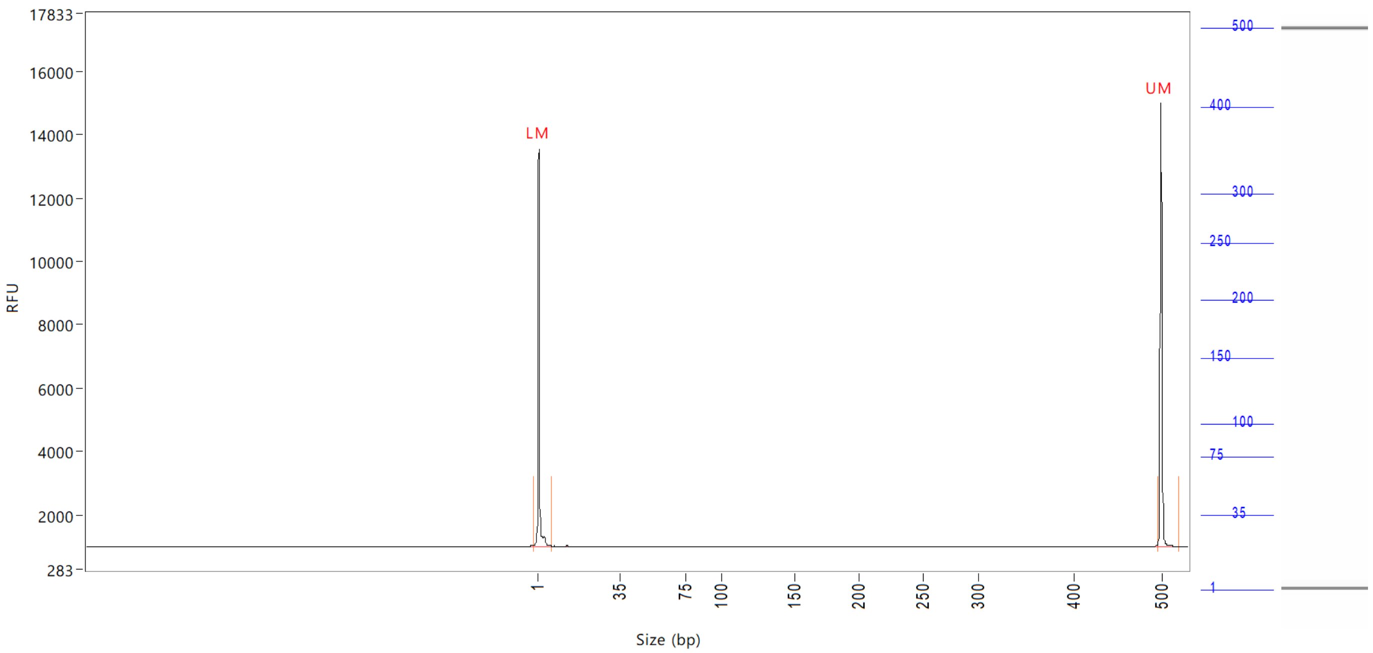


| Peak | Size (bp) | Concentration (ng/uL) | From (bp) | To (bp) | Average size (bp) | CV% | RFU | Corrected peak area |
|---|---|---|---|---|---|---|---|---|
| 1 | 1 (LM) | 0.7955 | 0 | 8 | 1 | 73.11 | 12549 | 46.881 |
| 2 | 500 (UM) | 0.5000 | 496 | 0 | 500 | 0.30 | 14033 | 29.466 |

| | | |
|---|---|---|
| TIC: | 0.0000 | ng/uL |
| TIM: | 0.0000 | nmole/L |
| Total concentration: | 0.4141 | ng/uL |

Sample peak width (sec): 5    Sample min peak height: 50    Sample baseline V to V?: Y    Sample baseline V to V points: 3
Sample filter: Binomial    Number of points for filter: 3  Sample start region (min): 0  Sample end region (min): 80
Marker peak width (sec): 5    Marker min peak height: 500    Marker baseline V to V?: Y    Marker baseline V to V points: 3
Lower marker selection: First peak > 500 RFU    Upper marker selection: Last peak > 500 RFU
Ladder size (bp)1, 35, 75, 100, 150, 200, 250, 300, 400, 500
Quantification using: Upper Marker    Final concentration (ng/uL): 0.5000    Dilution factor: 12.0

**Sample:** SampH11
**Well location:** H11
**Created:** Monday, 11 October 2021 14:41:52

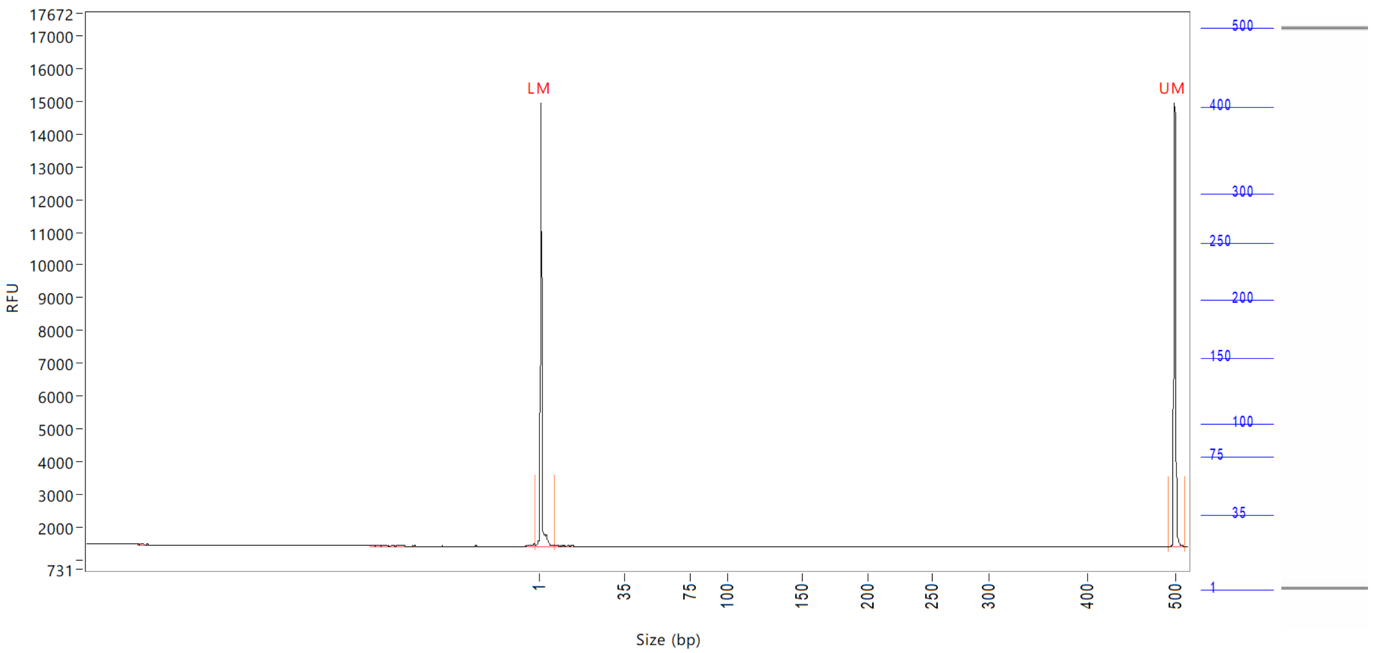


| Peak | Size (bp) | Concentration (ng/uL) | From (bp) | To (bp) | Average size (bp) | CV% | RFU | Corrected peak area |
|---|---|---|---|---|---|---|---|---|
| 1 | 1 (LM) | 0.8905 | 0 | 8 | 1 | 69.76 | 13526 | 49.938 |
| 2 | 500 (UM) | 0.5000 | 493 | 512 | 500 | 0.25 | 13518 | 28.038 |

| | | |
|---|---|---|
| TIC: | 0.0000 | ng/uL |
| TIM: | 0.0000 | nmole/L |
| Total concentration: | 0.5799 | ng/uL |

Sample peak width (sec): 5    Sample min peak height: 50    Sample baseline V to V?: Y    Sample baseline V to V points: 3
Sample filter: Binomial    Number of points for filter: 3    Sample start region (min): 0    Sample end region (min): 80
Marker peak width (sec): 5    Marker min peak height: 500    Marker baseline V to V?: Y    Marker baseline V to V points: 3
Lower marker selection: First peak > 500 RFU    Upper marker selection: Last peak > 500 RFU
Ladder size (bp)1, 35, 75, 100, 150, 200, 250, 300, 400, 500
Quantification using: Upper Marker    Final concentration (ng/uL): 0.5000    Dilution factor: 12.0

**Sample:** SampA12
**Well location:** A12
**Created:** Monday, 11 October 2021 14:41:52

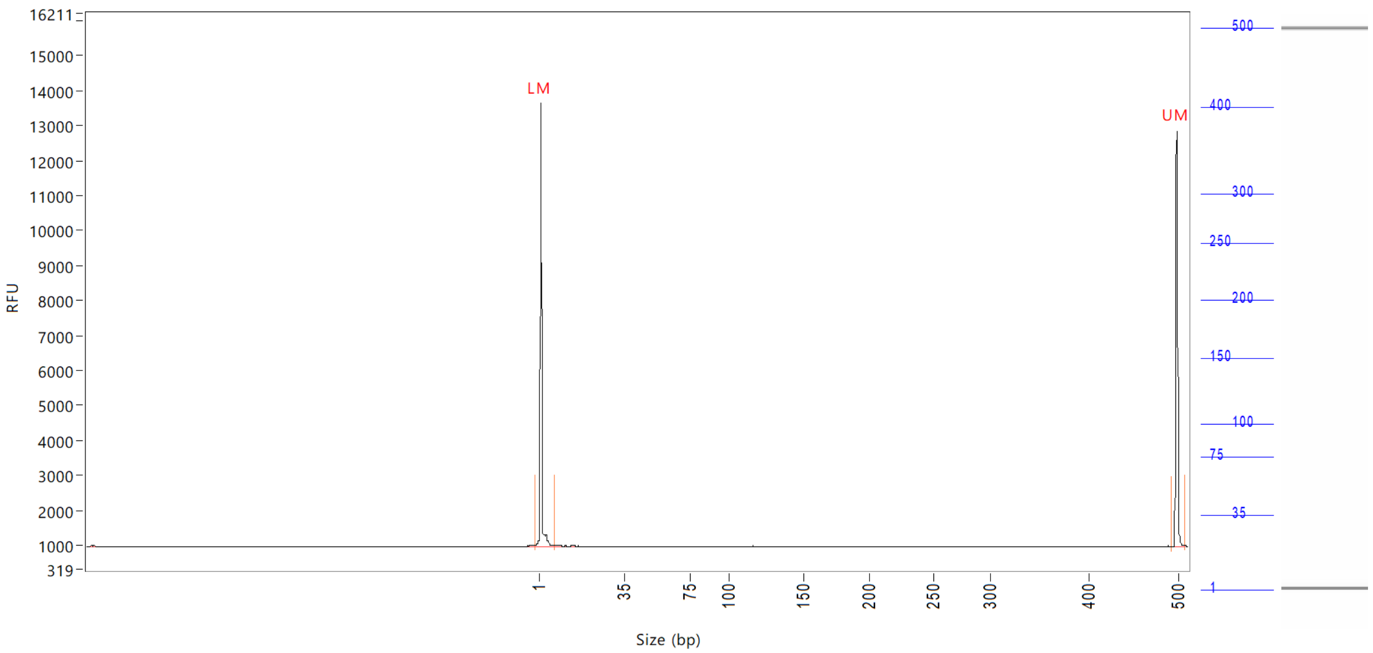


| Peak | Size (bp) | Concentration (ng/uL) | From (bp) | To (bp) | Average size (bp) | CV% | RFU | Corrected peak area |
|---|---|---|---|---|---|---|---|---|
| 1 | 1 (LM) | 0.9672 | 0 | 7 | 1 | 73.28 | 12691 | 47.727 |
| 2 | 500 (UM) | 0.5000 | 494 | 510 | 500 | 0.23 | 11848 | 24.672 |

| | | |
|---|---|---|
| TIC: | 0.0000 | ng/uL |
| TIM: | 0.0000 | nmole/L |
| Total concentration: | 0.5156 | ng/uL |

Sample peak width (sec): 5    Sample min peak height: 50    Sample baseline V to V?: Y    Sample baseline V to V points: 3
Sample filter: Binomial    Number of points for filter: 3    Sample start region (min): 0    Sample end region (min): 80
Marker peak width (sec): 5    Marker min peak height: 500    Marker baseline V to V?: Y    Marker baseline V to V points: 3
Lower marker selection: First peak > 500 RFU    Upper marker selection: Last peak > 500 RFU
Ladder size (bp)1, 35, 75, 100, 150, 200, 250, 300, 400, 500
Quantification using: Upper Marker    Final concentration (ng/uL): 0.5000    Dilution factor: 12.0

**Sample:** SampB12
**Well location:** B12
**Created:** Monday, 11 October 2021 14:41:52

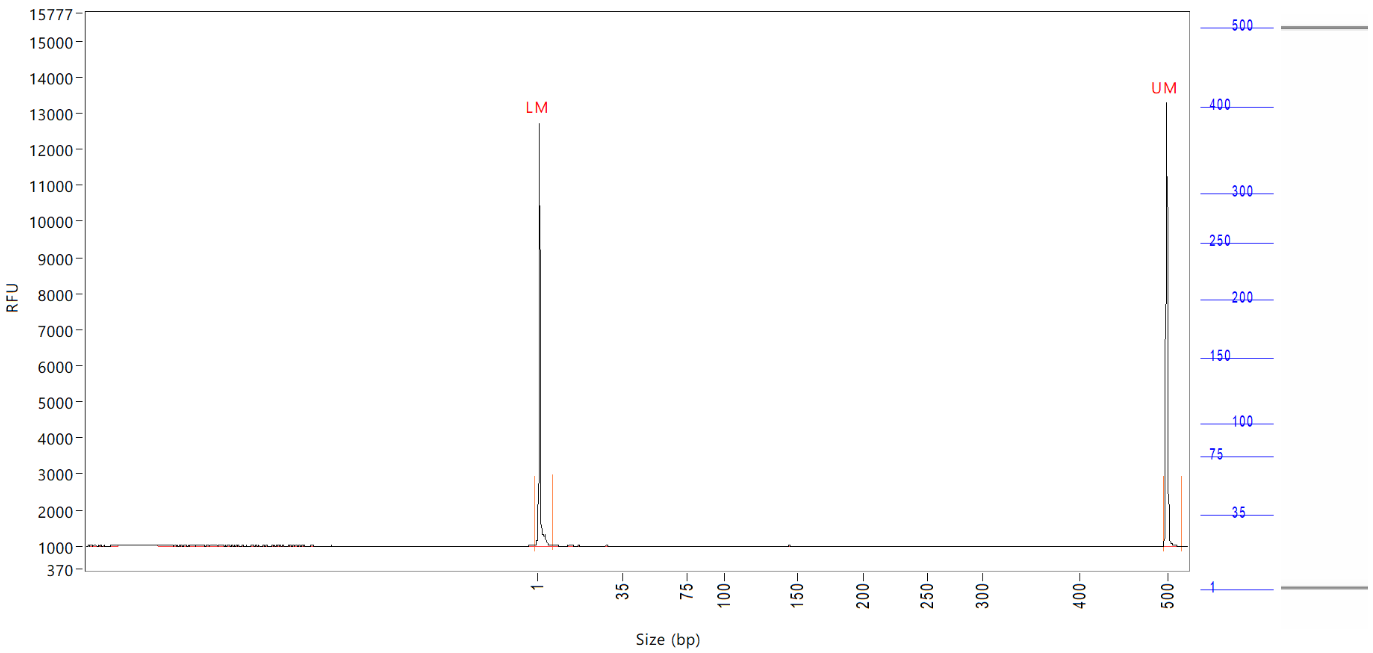


| Peak | Size (bp) | Concentration (ng/uL) | From (bp) | To (bp) | Average size (bp) | CV% | RFU | Corrected peak area |
|---|---|---|---|---|---|---|---|---|
| 1 | 1 (LM) | 0.8501 | 0 | 7 | 2 | 65.12 | 11720 | 43.656 |
| 2 | 500 (UM) | 0.5000 | 496 | 0 | 500 | 0.27 | 12320 | 25.676 |

| | | |
|---|---|---|
| TIC: | 0.0000 | ng/uL |
| TIM: | 0.0000 | nmole/L |
| Total concentration: | 0.4928 | ng/uL |

Sample peak width (sec): 5 Sample min peak height: 50 Sample baseline V to V?: Y Sample baseline V to V points: 3
Sample filter: Binomial Number of points for filter: 3 Sample start region (min): 0 Sample end region (min): 80
Marker peak width (sec): 5 Marker min peak height: 500 Marker baseline V to V?: Y Marker baseline V to V points: 3
Lower marker selection: First peak > 500 RFU Upper marker selection: Last peak > 500 RFU
Ladder size (bp)1, 35, 75, 100, 150, 200, 250, 300, 400, 500
Quantification using: Upper Marker Final concentration (ng/uL): 0.5000 Dilution factor: 12.0

**Sample:** SampC12
**Well location:** C12
**Created:** Monday, 11 October 2021 14:41:52

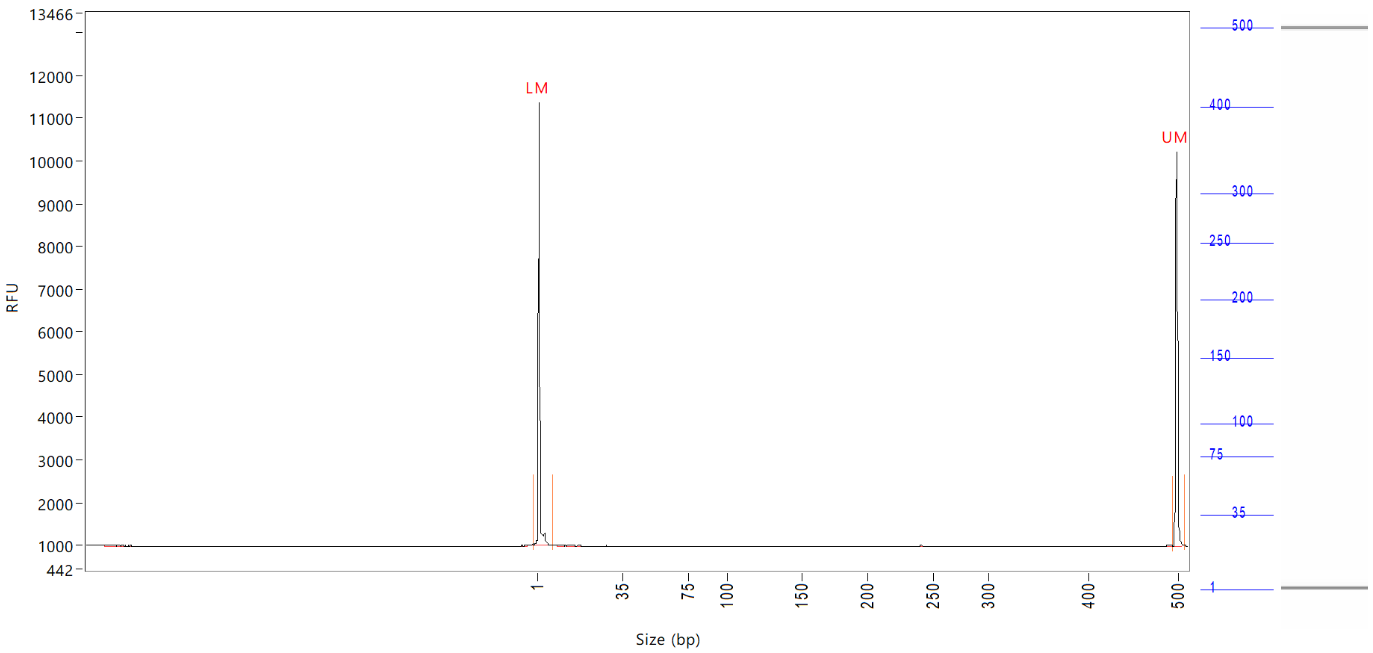


| Peak | Size<br>(bp) | Concentration<br>(ng/uL) | From<br>(bp) | To<br>(bp) | Average size<br>(bp) | CV% | RFU | Corrected peak area |
|---|---|---|---|---|---|---|---|---|
| 1 | 1 (LM) | 1.0002 | 0 | 7 | 1 | 68.37 | 10398 | 39.417 |
| 2 | 500 (UM) | 0.5000 | 496 | 510 | 500 | 0.24 | 9230 | 19.705 |

| | | |
|---|---|---|
| TIC: | 0.0000 | ng/uL |
| TIM: | 0.0000 | nmole/L |
| Total concentration: | 0.7427 | ng/uL |

Sample peak width (sec): 5 Sample min peak height: 50 Sample baseline V to V?: Y Sample baseline V to V points: 3
Sample filter: Binomial Number of points for filter: 3 Sample start region (min): 0 Sample end region (min): 80
Marker peak width (sec): 5 Marker min peak height: 500 Marker baseline V to V?: Y Marker baseline V to V points: 3
Lower marker selection: First peak > 500 RFU Upper marker selection: Last peak > 500 RFU
Ladder size (bp)1, 35, 75, 100, 150, 200, 250, 300, 400, 500
Quantification using: Upper Marker Final concentration (ng/uL): 0.5000 Dilution factor: 12.0

**Sample:** SampD12
**Well location:** D12
**Created:** Monday, 11 October 2021 14:41:52

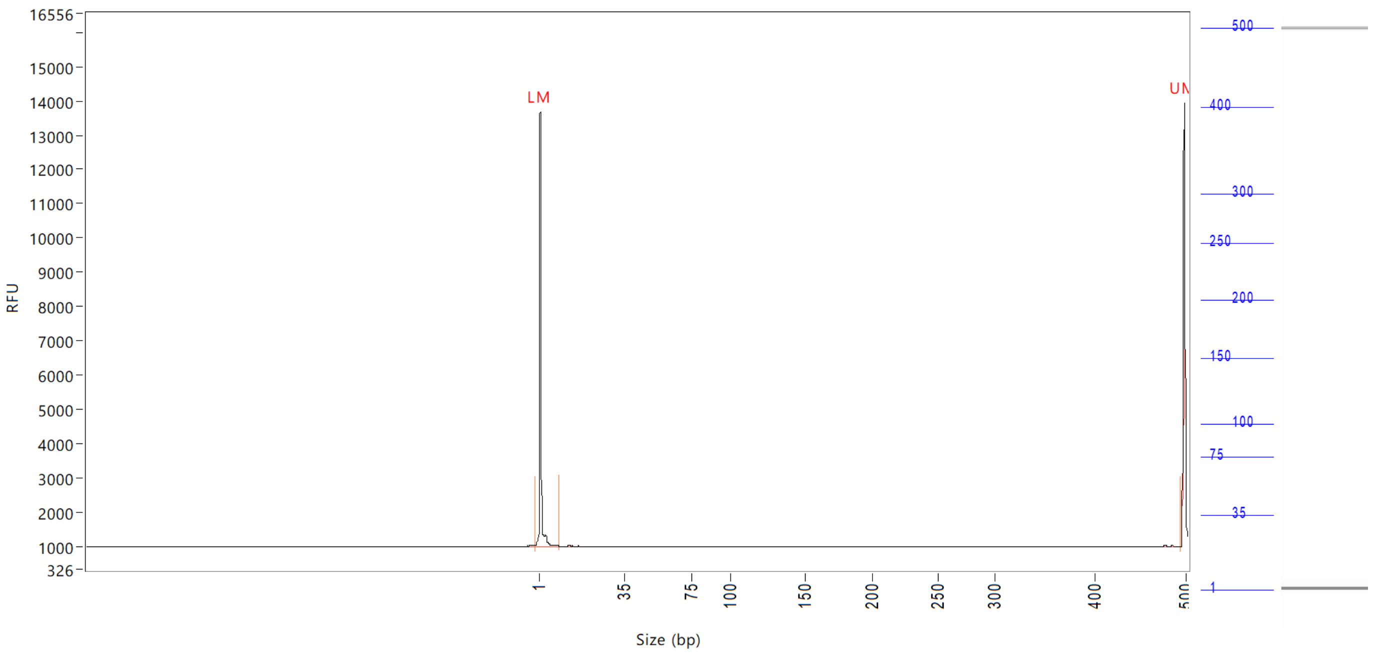


| Peak | Size (bp) | Concentration (ng/uL) | From (bp) | To (bp) | Average size (bp) | CV% | RFU | Corrected peak area |
|---|---|---|---|---|---|---|---|---|
| 1 | 1 (LM) | 1.7794 | 0 | 9 | 1 | 76.11 | 12688 | 48.009 |
| 2 | 500 (UM) | 0.5000 | 495 | 501 | 500 | 0.14 | 8752 | 13.491 |

| | | |
|---|---|---|
| TIC: | 0.0000 | ng/uL |
| TIM: | 0.0000 | nmole/L |
| Total concentration: | 0.9795 | ng/uL |

Sample peak width (sec): 5 Sample min peak height: 50 Sample baseline V to V?: Y Sample baseline V to V points: 3
Sample filter: Binomial Number of points for filter: 3 Sample start region (min): 0 Sample end region (min): 80
Marker peak width (sec): 5 Marker min peak height: 500 Marker baseline V to V?: Y Marker baseline V to V points: 3
Lower marker selection: First peak > 500 RFU Upper marker selection: Last peak > 500 RFU
Ladder size (bp)1, 35, 75, 100, 150, 200, 250, 300, 400, 500
Quantification using: Upper Marker Final concentration (ng/uL): 0.5000 Dilution factor: 12.0

**Sample:** SampE12
**Well location:** E12
**Created:** Monday, 11 October 2021 14:41:52

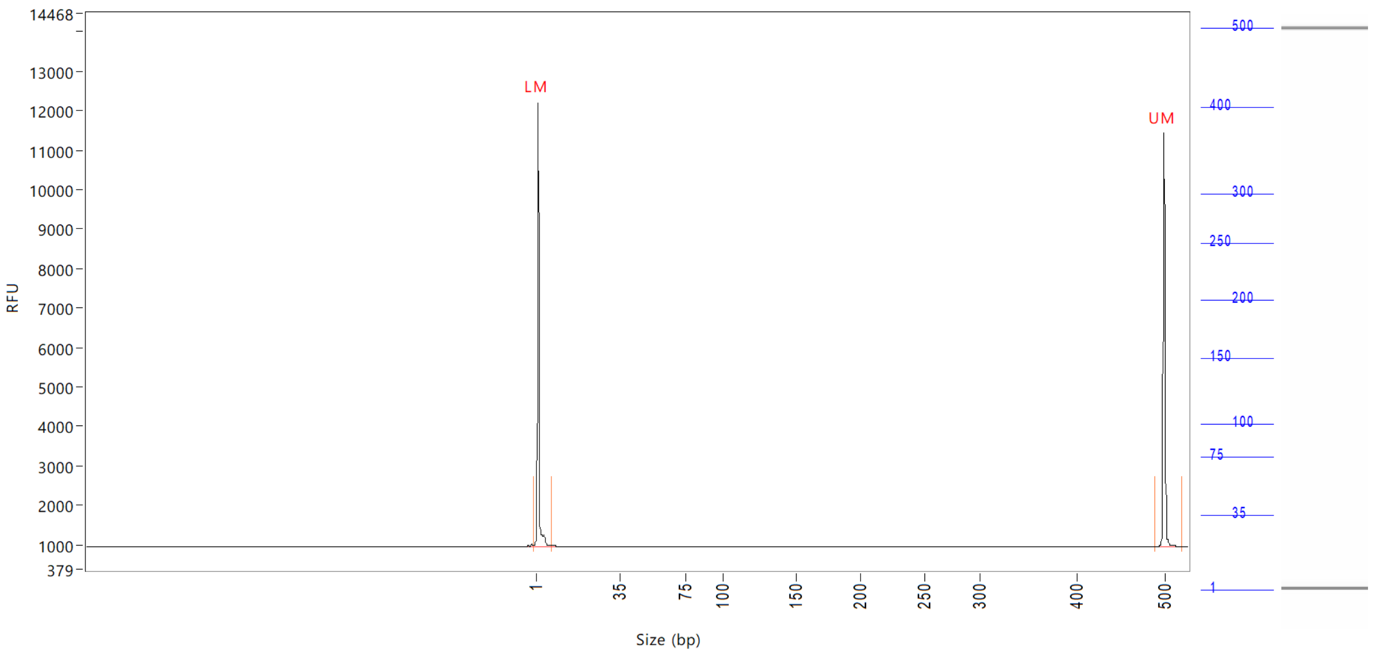


| Peak | Size (bp) | Concentration (ng/uL) | From (bp) | To (bp) | Average size (bp) | CV% | RFU | Corrected peak area |
|---|---|---|---|---|---|---|---|---|
| 1 | 1 (LM) | 0.9387 | 0 | 8 | 1 | 73.01 | 11257 | 42.635 |
| 2 | 500 (UM) | 0.5000 | 491 | 0 | 500 | 0.31 | 10507 | 22.710 |

| | | |
|---|---|---|
| TIC: | 0.0000 | ng/uL |
| TIM: | 0.0000 | nmole/L |
| Total concentration: | 0.4595 | ng/uL |

Sample peak width (sec): 5    Sample min peak height: 50    Sample baseline V to V?: Y    Sample baseline V to V points: 3
Sample filter: Binomial    Number of points for filter: 3  Sample start region (min): 0  Sample end region (min): 80
Marker peak width (sec): 5    Marker min peak height: 500   Marker baseline V to V?: Y    Marker baseline V to V points: 3
Lower marker selection: First peak > 500 RFU    Upper marker selection: Last peak > 500 RFU
Ladder size (bp)1, 35, 75, 100, 150, 200, 250, 300, 400, 500
Quantification using: Upper Marker    Final concentration (ng/uL): 0.5000    Dilution factor: 12.0

**Sample:** SampF12
**Well location:** F12
**Created:** Monday, 11 October 2021 14:41:52

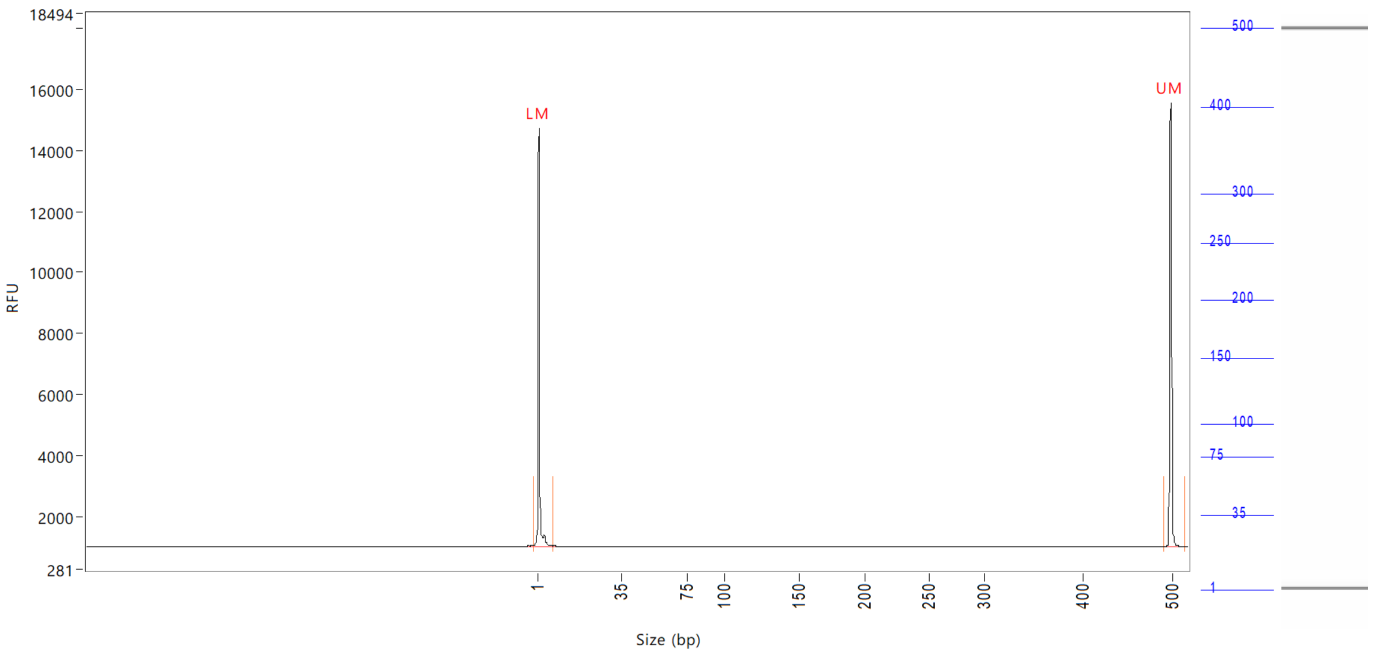


| Peak | Size<br>(bp) | Concentration<br>(ng/uL) | From<br>(bp) | To<br>(bp) | Average size<br>(bp) | CV% | RFU | Corrected peak area |
|---|---|---|---|---|---|---|---|---|
| 1 | 1 (LM) | 0.8590 | 0 | 8 | 1 | 69.35 | 13713 | 51.365 |
| 2 | 500 (UM) | 0.5000 | 493 | 517 | 500 | 0.25 | 14560 | 29.896 |

| | | |
|---|---|---|
| TIC: | 0.0000 | ng/uL |
| TIM: | 0.0000 | nmole/L |
| Total concentration: | 0.4011 | ng/uL |

Sample peak width (sec): 5    Sample min peak height: 50    Sample baseline V to V?: Y    Sample baseline V to V points: 3
Sample filter: Binomial    Number of points for filter: 3  Sample start region (min): 0  Sample end region (min): 80
Marker peak width (sec): 5    Marker min peak height: 500    Marker baseline V to V?: Y    Marker baseline V to V points: 3
Lower marker selection: First peak > 500 RFU    Upper marker selection: Last peak > 500 RFU
Ladder size (bp)1, 35, 75, 100, 150, 200, 250, 300, 400, 500
Quantification using: Upper Marker    Final concentration (ng/uL): 0.5000    Dilution factor: 12.0

**Sample:** SampG12
**Well location:** G12
**Created:** Monday, 11 October 2021 14:41:52

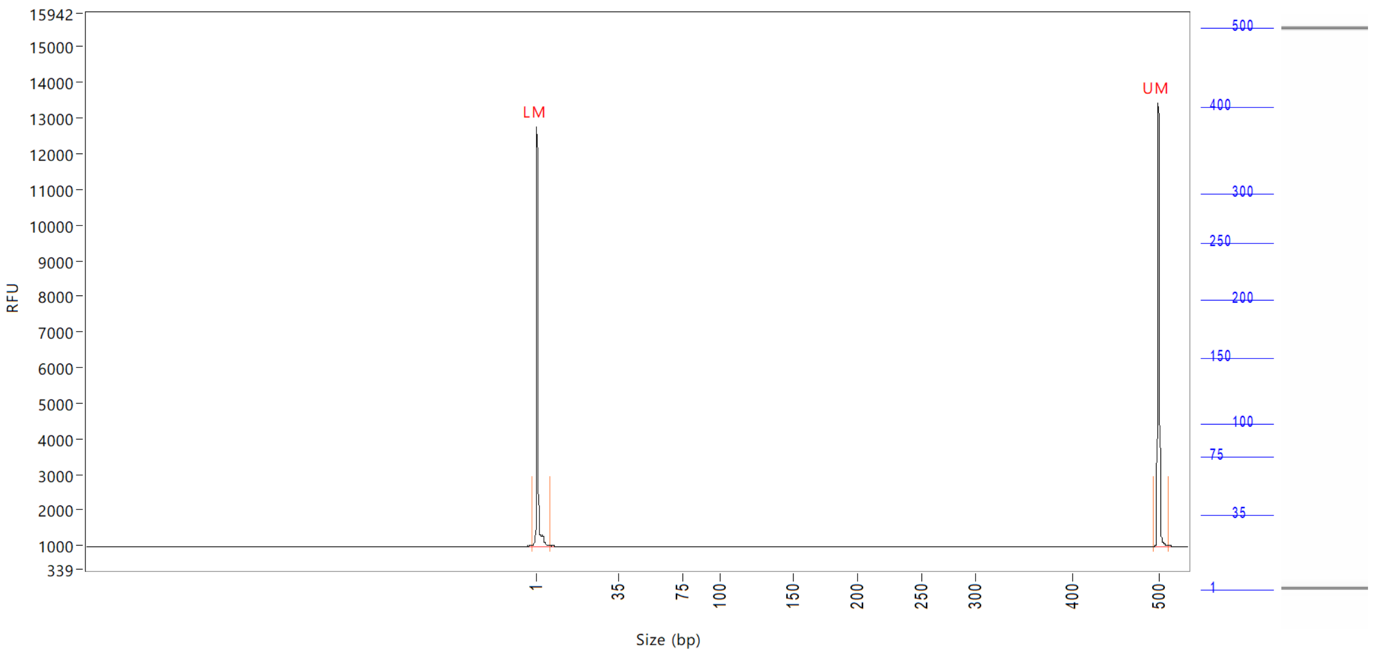


| Peak | Size (bp) | Concentration (ng/uL) | From (bp) | To (bp) | Average size (bp) | CV% | RFU | Corrected peak area |
|---|---|---|---|---|---|---|---|---|
| 1 | 1 (LM) | 0.8594 | 0 | 7 | 1 | 69.84 | 11788 | 44.835 |
| 2 | 500 (UM) | 0.5000 | 494 | 511 | 500 | 0.26 | 12477 | 26.086 |

| | | |
|---|---|---|
| TIC: | 0.0000 | ng/uL |
| TIM: | 0.0000 | nmole/L |
| Total concentration: | 0.4257 | ng/uL |

```
Sample peak width (sec): 5    Sample min peak height: 50    Sample baseline V to V?: Y    Sample baseline V to V points: 3
Sample filter: Binomial       Number of points for filter: 3  Sample start region (min): 0  Sample end region (min): 80
Marker peak width (sec): 5    Marker min peak height: 500   Marker baseline V to V?: Y    Marker baseline V to V points: 3
Lower marker selection: First peak > 500 RFU                Upper marker selection: Last peak > 500 RFU
Ladder size (bp)1, 35, 75, 100, 150, 200, 250, 300, 400, 500
Quantification using: Upper Marker          Final concentration (ng/uL): 0.5000          Dilution factor: 12.0
```

**Sample:** SampH12
**Well location:** H12
**Created:** Monday, 11 October 2021 14:41:52

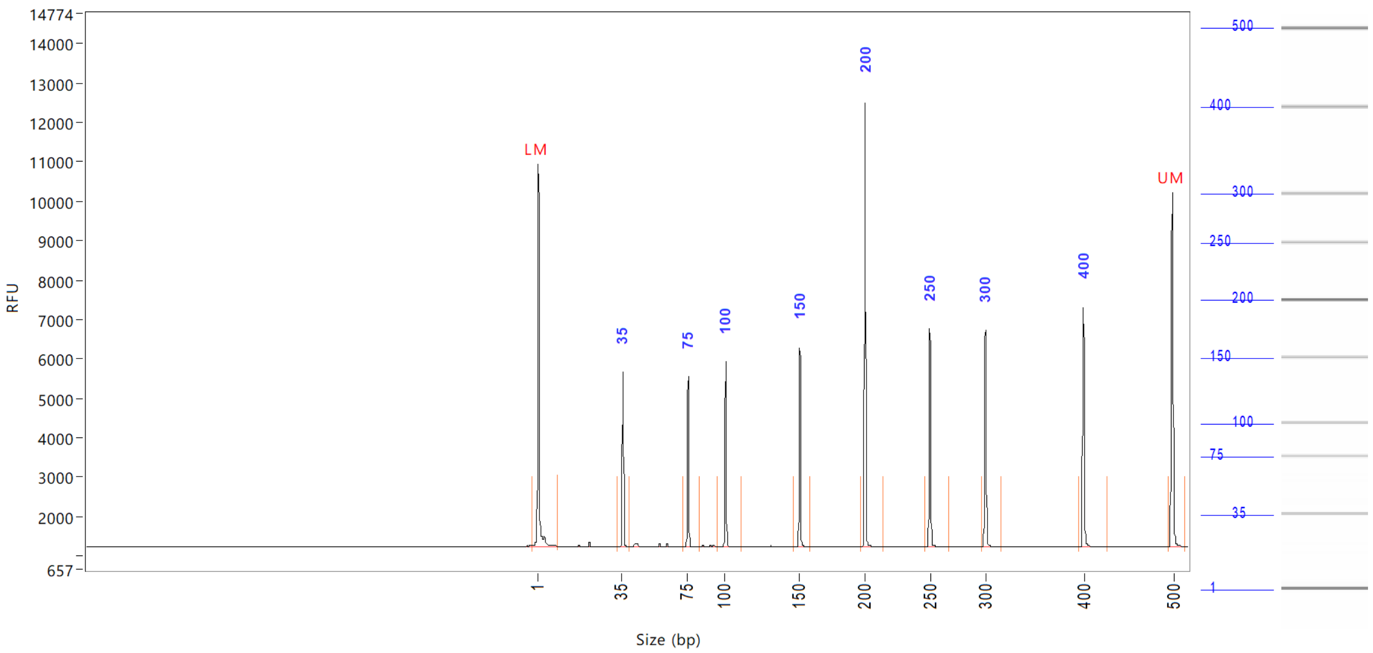


| Peak | Size (bp) | Concentration (ng/uL) | From (bp) | To (bp) | Average size (bp) | CV% | RFU | Corrected peak area |
|---|---|---|---|---|---|---|---|---|
| 1 | 1 (LM) | 1.0169 | 0 | 10 | 1 | 76.29 | 9721 | 37.878 |
| 2 | 35 | 4.2349 | 33 | 40 | 35 | 2.04 | 4444 | 13.146 |
| 3 | 75 | 3.5433 | 73 | 84 | 75 | 1.13 | 4316 | 10.999 |
| 4 | 100 | 3.5680 | 95 | 111 | 100 | 0.86 | 4686 | 11.076 |
| 5 | 150 | 3.5348 | 146 | 159 | 150 | 0.59 | 5062 | 10.973 |
| 6 | 200 | 7.3911 | 197 | 215 | 200 | 0.50 | 11288 | 22.943 |
| 7 | 250 | 3.7261 | 247 | 268 | 250 | 0.49 | 5538 | 11.566 |
| 8 | 300 | 3.6982 | 297 | 317 | 300 | 0.43 | 5503 | 11.480 |
| 9 | 400 | 3.9486 | 396 | 427 | 400 | 0.43 | 6081 | 12.257 |
| 10 | 500 (UM) | 0.5000 | 496 | 515 | 500 | 0.25 | 8997 | 18.625 |

| | | |
|---|---|---|
| TIC: | 33.6450 | ng/uL |
| TIM: | 494.3617 | nmole/L |
| Total concentration: | 34.5977 | ng/uL |

```
Sample peak width (sec): 10   Sample min peak height: 500   Sample baseline V to V?: Y    Sample baseline V to V points: 3
Sample filter: Binomial       Number of points for filter: 3  Sample start region (min): 0  Sample end region (min): 80
Marker peak width (sec): 5    Marker min peak height: 500   Marker baseline V to V?: Y    Marker baseline V to V points: 3
Lower marker selection: First peak > 500 RFU                Upper marker selection: Last peak > 500 RFU
Ladder size (bp)1, 35, 75, 100, 150, 200, 250, 300, 400, 500
Quantification using: Upper Marker          Final concentration (ng/uL): 0.5000          Dilution factor: 12.0
```

**Sample:** SampH12
**Well location:** H12
**Created:** Monday, 11 October 2021 14:41:52
**Fit type:** Point to point

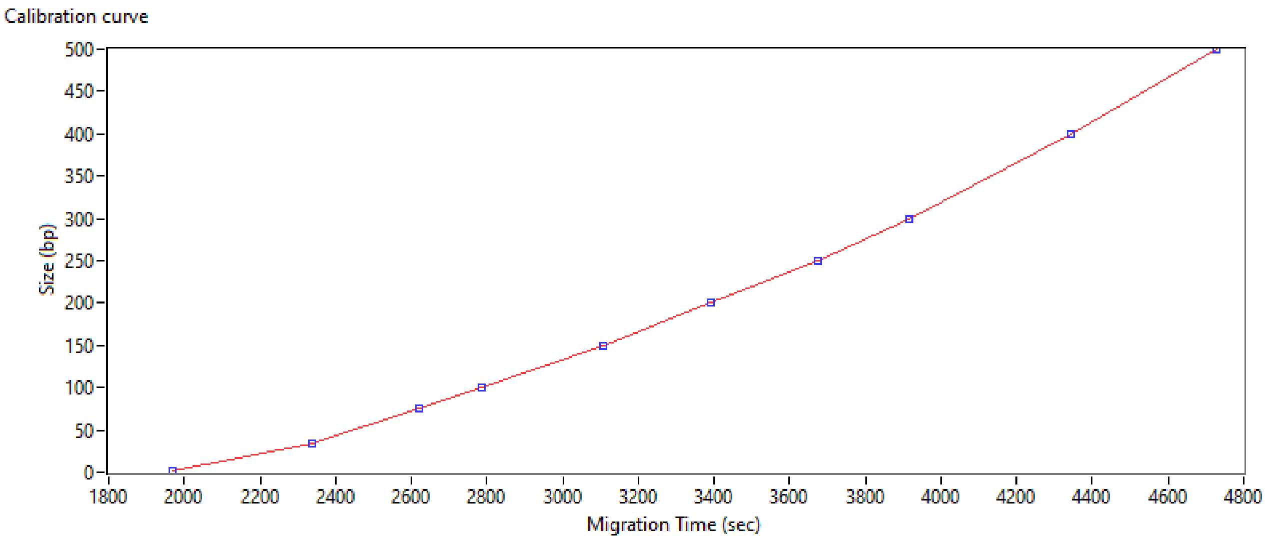


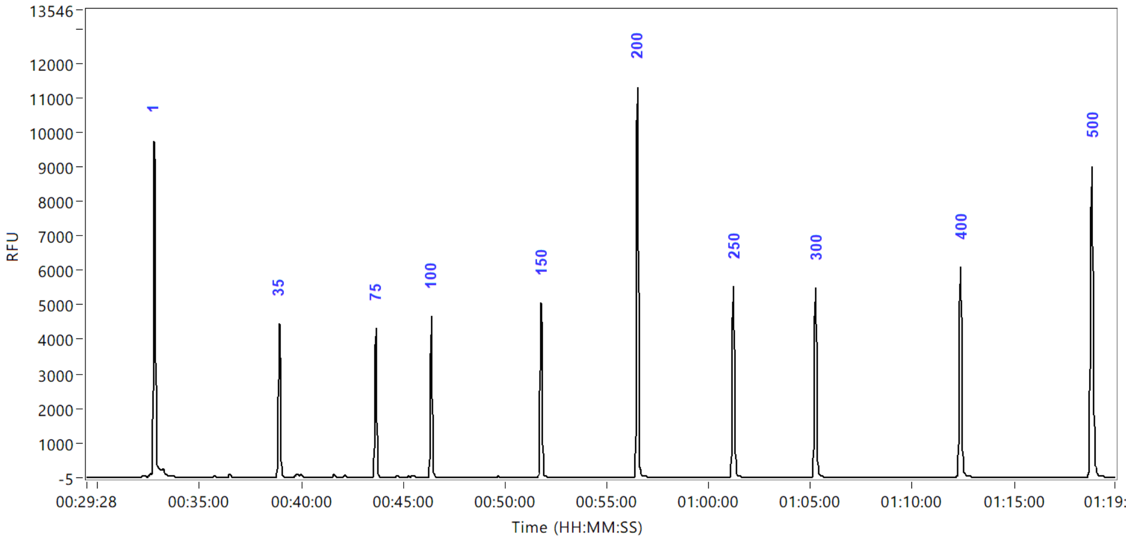


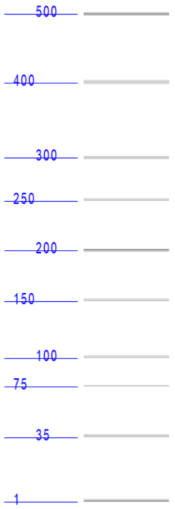
