## Supplementary figures and images for "Multiplexed Label-Free Biomarker Detection by Targeted Disassembly of Variable-Length DNA Payload Chains"

### 2021 10 11 14H 11M A1 SampA1.PNG

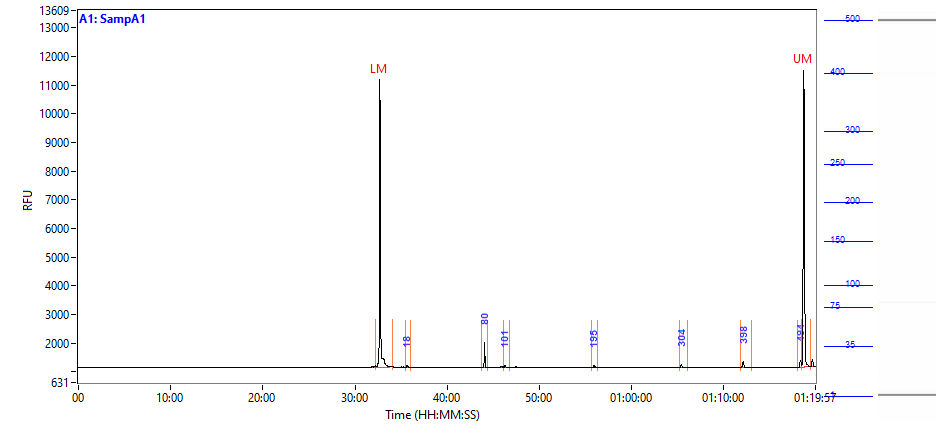

### 2021 10 11 14H 11M A2 SampA2.PNG

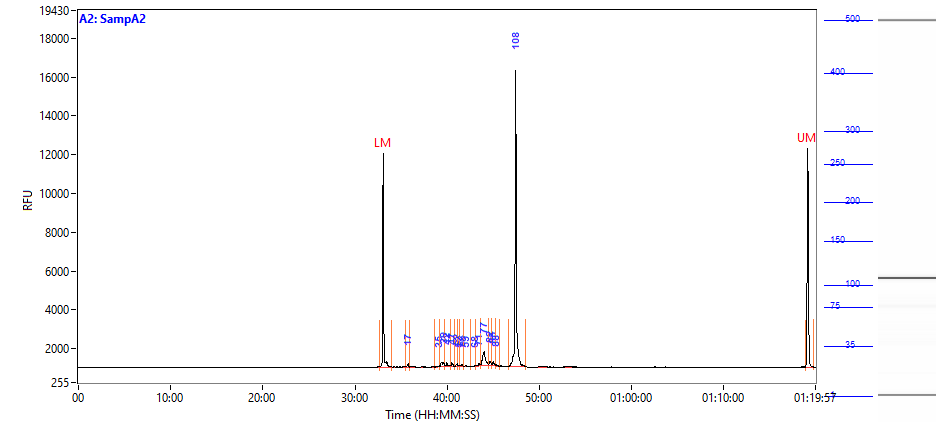

### 2021 10 11 14H 11M A3 SampA3.PNG

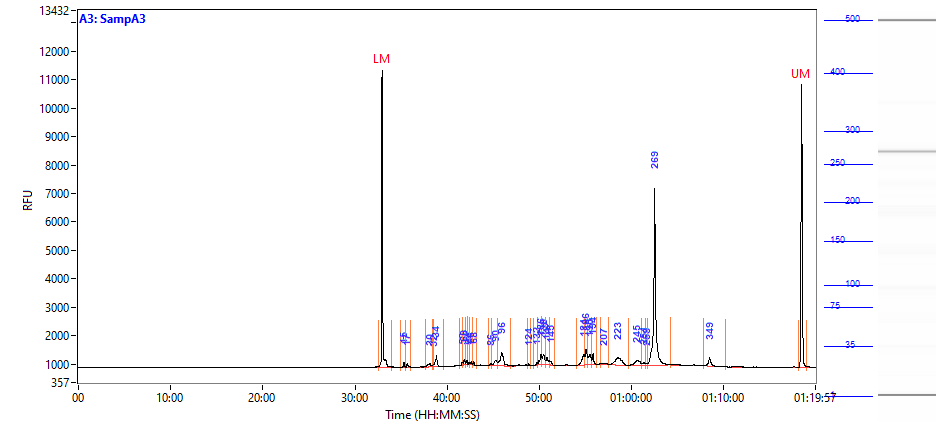

### 2021 10 11 14H 11M A4 SampA4.PNG

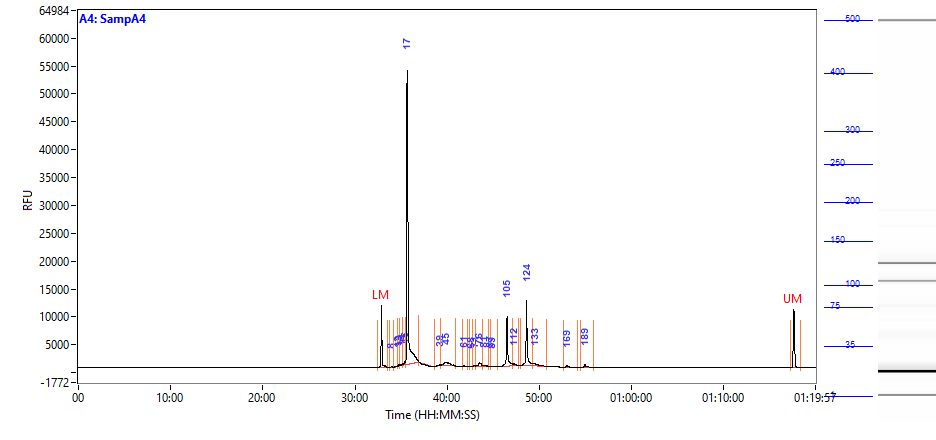

### 2021 10 11 14H 11M A5 SampA5.PNG

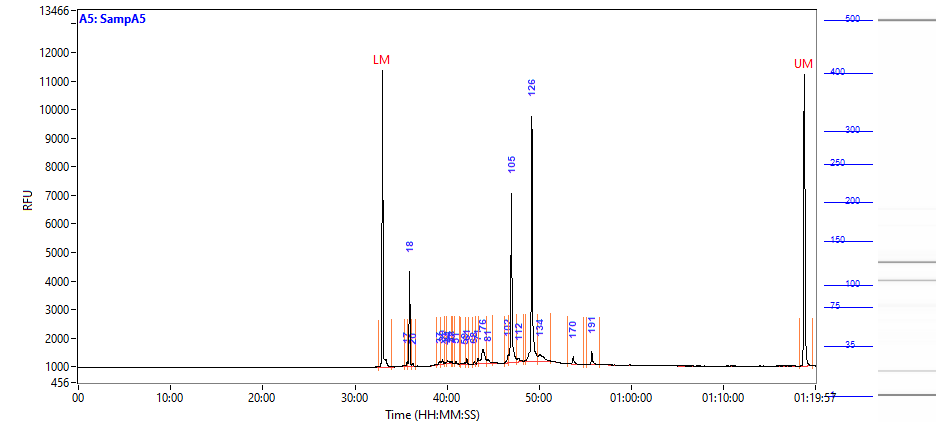

### 2021 10 11 14H 11M A6 SampA6.PNG

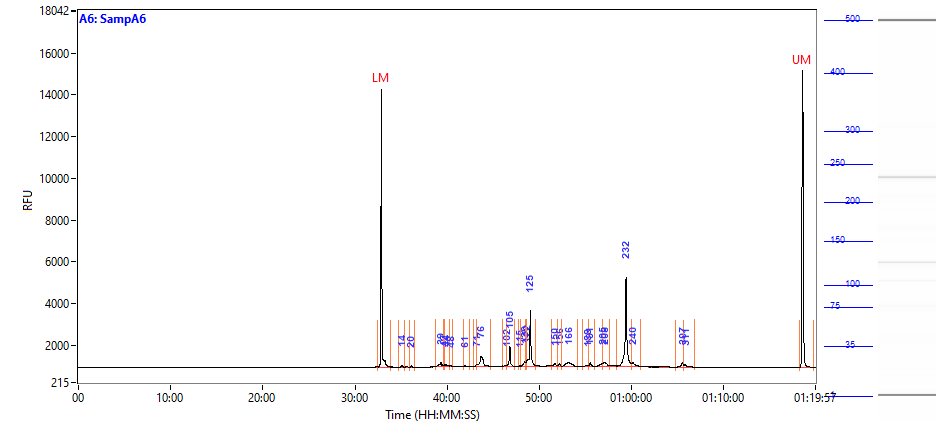

### 2021 10 11 14H 11M A7 SampA7.PNG

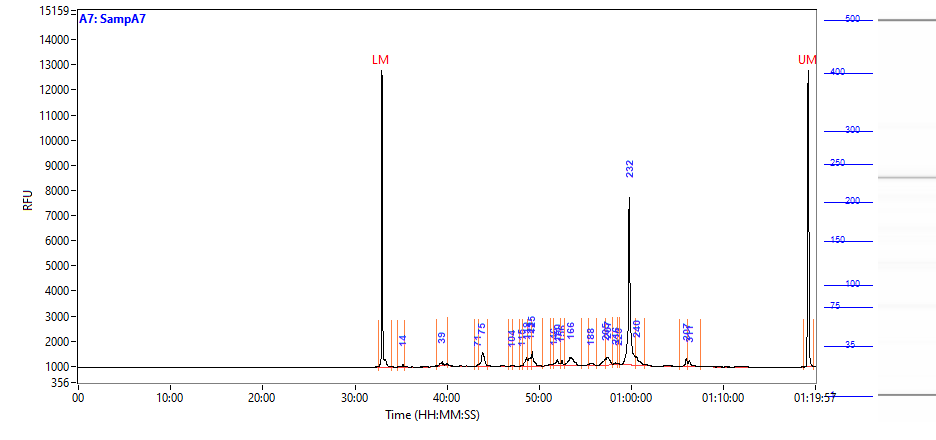

### 2021 10 11 14H 11M A8 SampA8.PNG

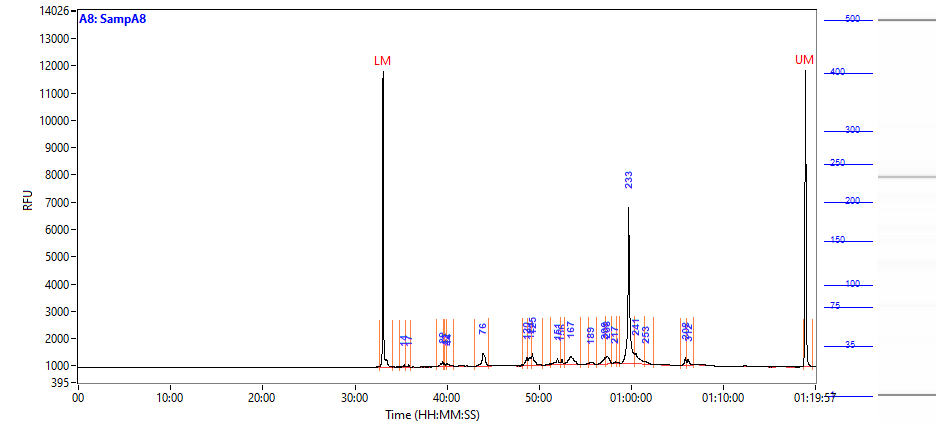

### 2021 10 11 14H 11M A9 SampA9.PNG

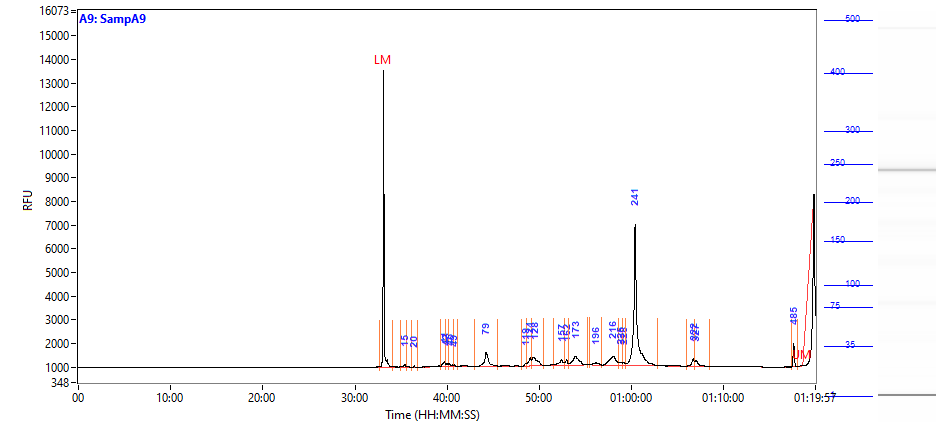

### 2021 10 11 14H 11M A10 SampA10.PNG

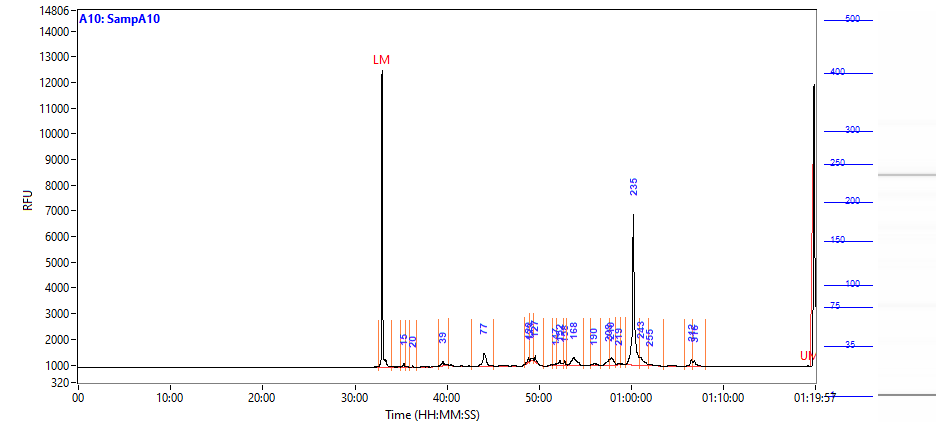

### 2021 10 11 14H 11M A11 SampA11.PNG

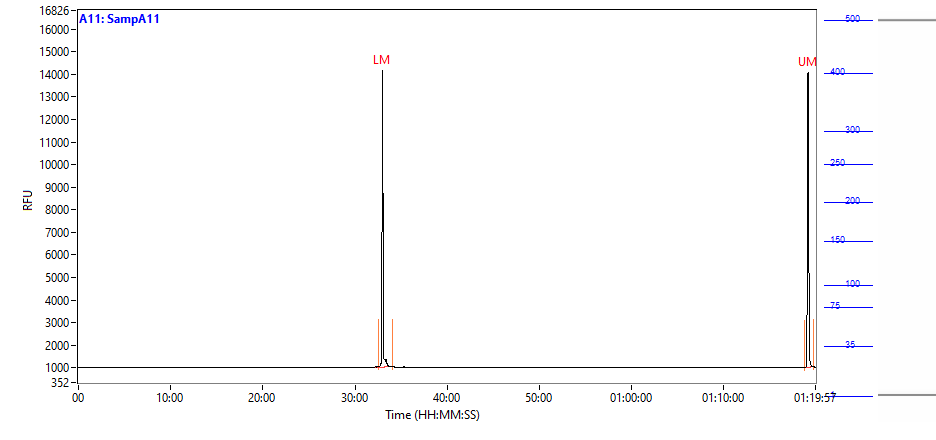

### 2021 10 11 14H 11M A12 SampA12.PNG

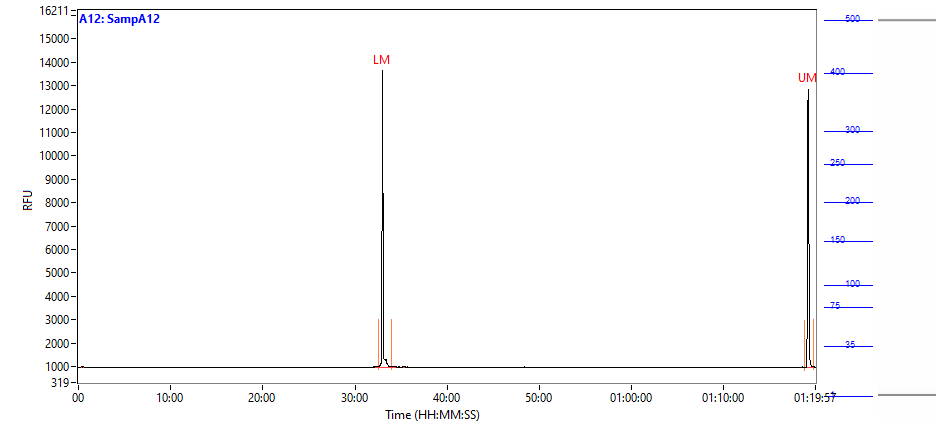

### 2021 10 11 14H 11M B1 SampB1.PNG

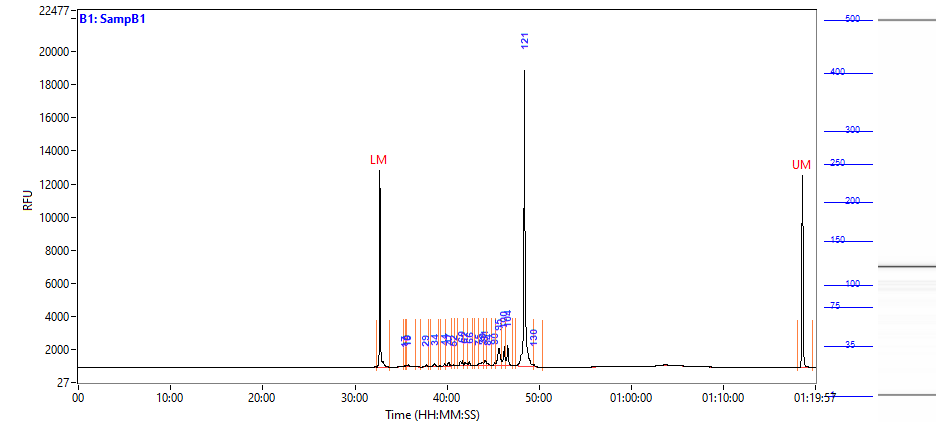

### 2021 10 11 14H 11M B2 SampB2.PNG

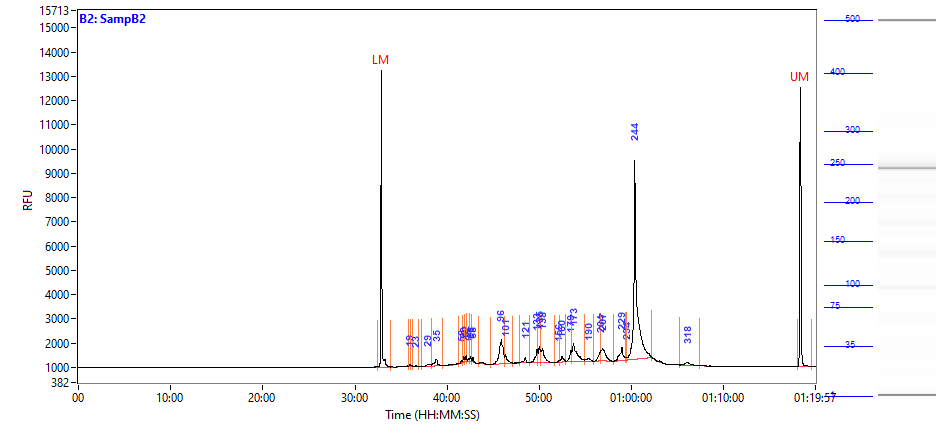

### 2021 10 11 14H 11M B3 SampB3.PNG

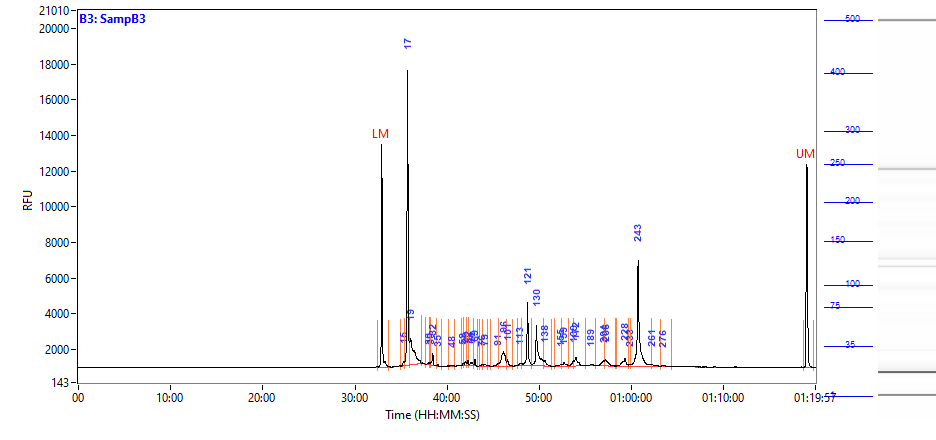

### 2021 10 11 14H 11M B4 SampB4.PNG

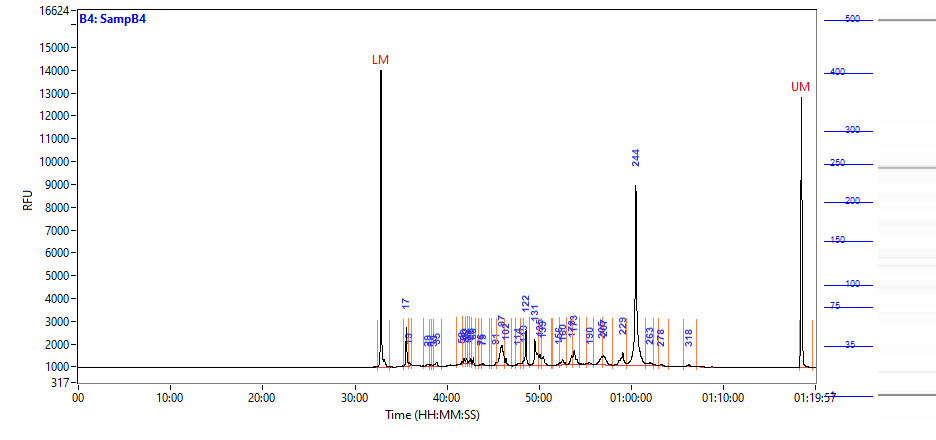

### 2021 10 11 14H 11M B5 SampB5.PNG

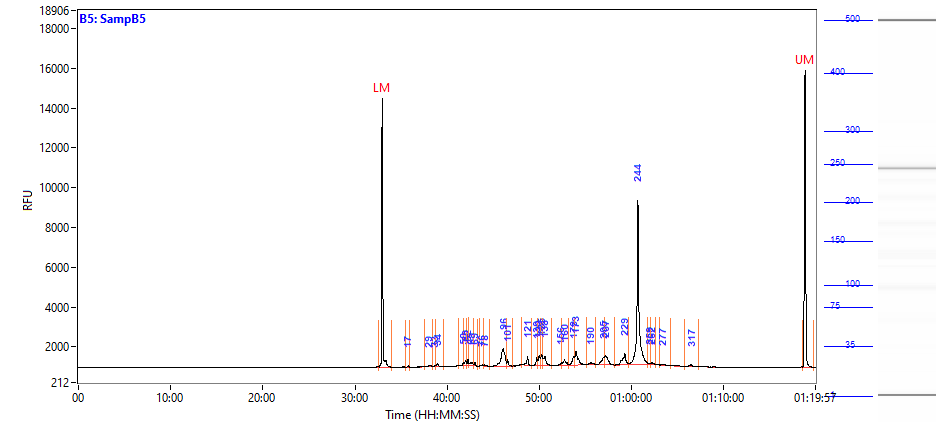

### 2021 10 11 14H 11M B6 SampB6.PNG

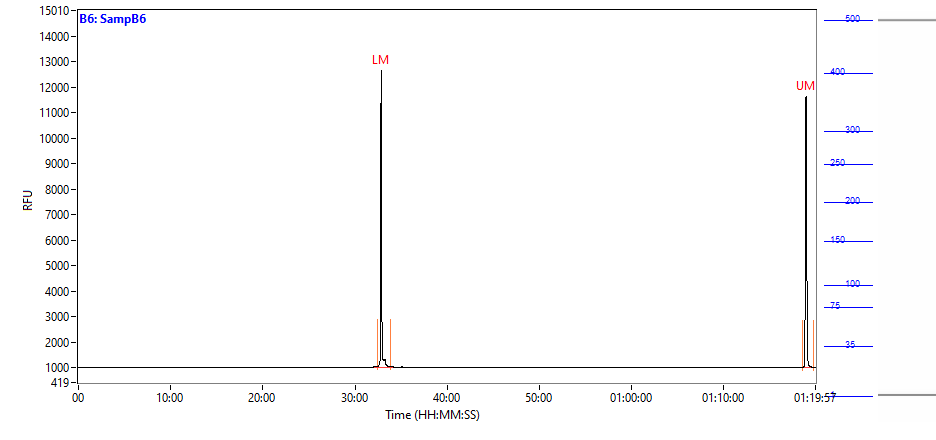

### 2021 10 11 14H 11M B7 SampB7.PNG

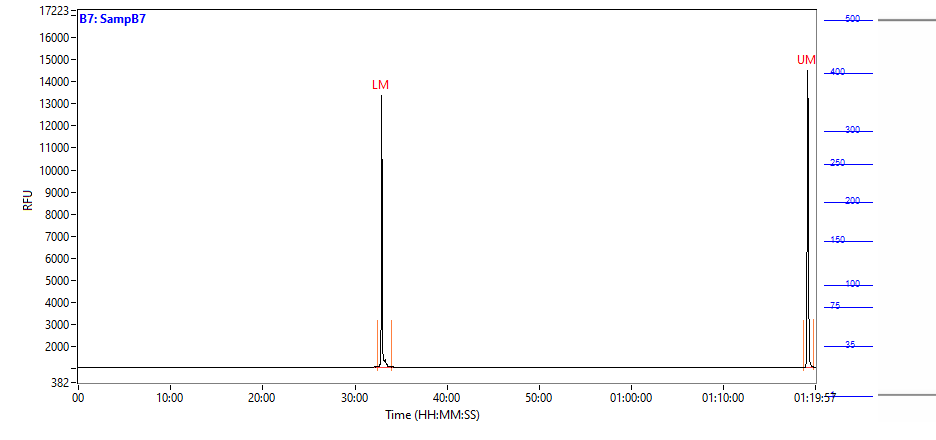

### 2021 10 11 14H 11M B8 SampB8.PNG

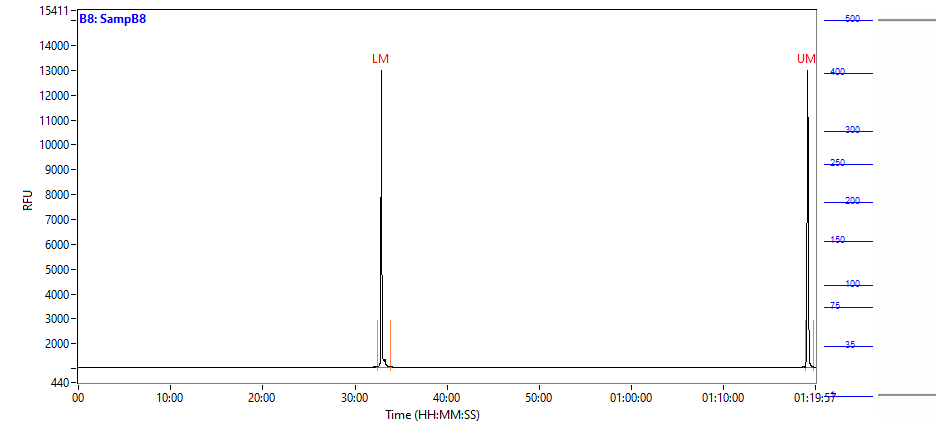

### 2021 10 11 14H 11M B9 SampB9.PNG

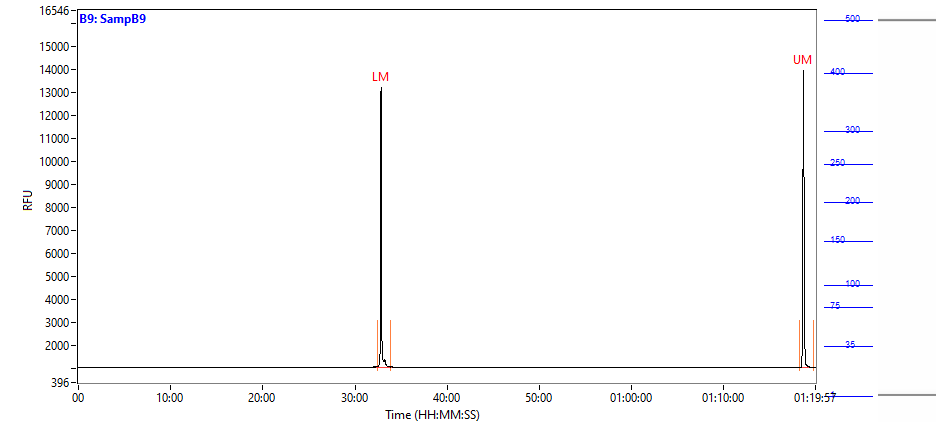

### 2021 10 11 14H 11M B10 SampB10.PNG

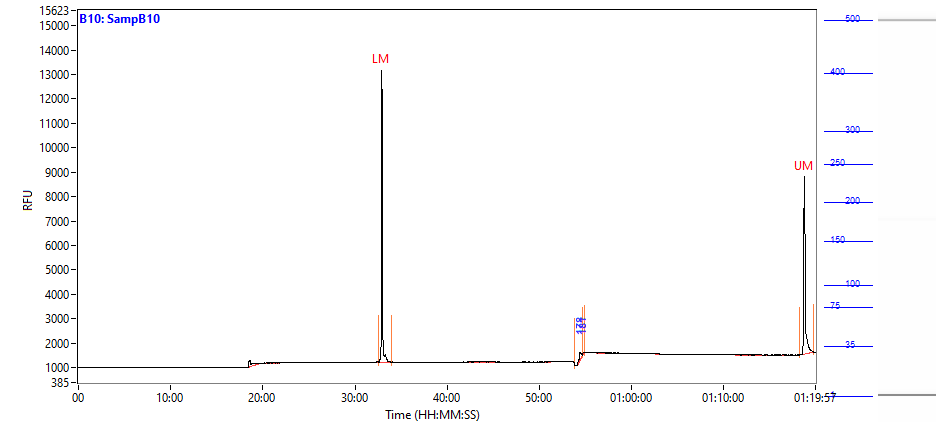

### 2021 10 11 14H 11M B11 SampB11.PNG

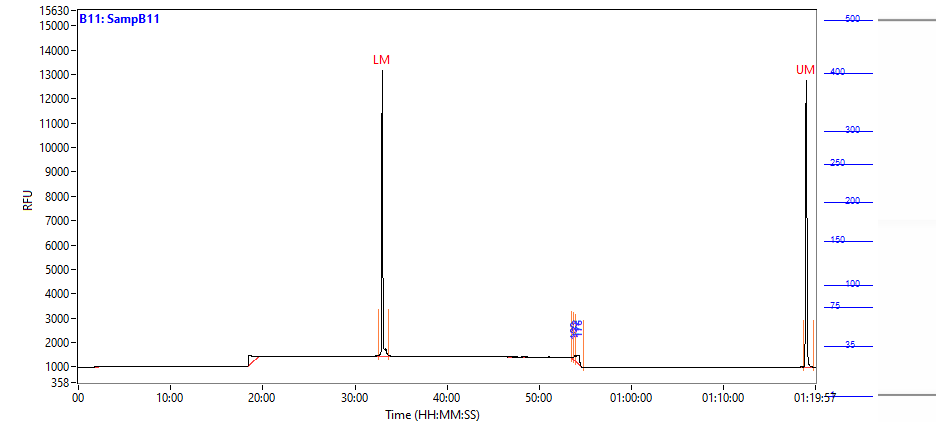

### 2021 10 11 14H 11M B12 SampB12.PNG

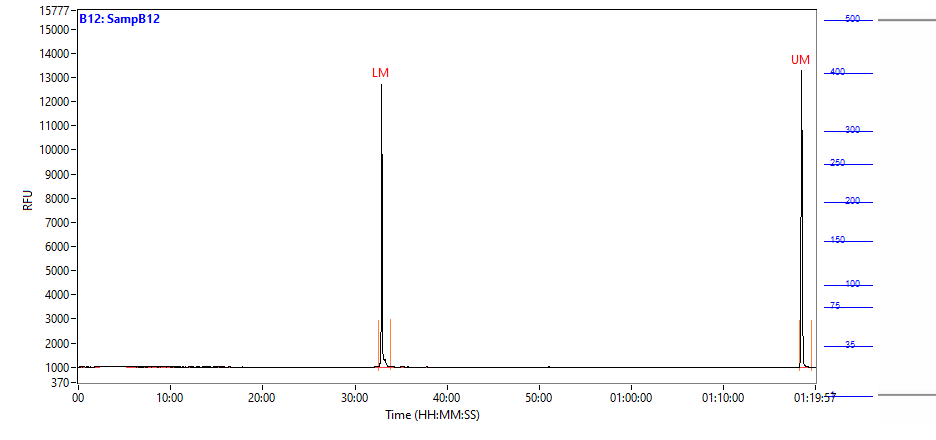

### 2021 10 11 14H 11M C1 SampC1.PNG

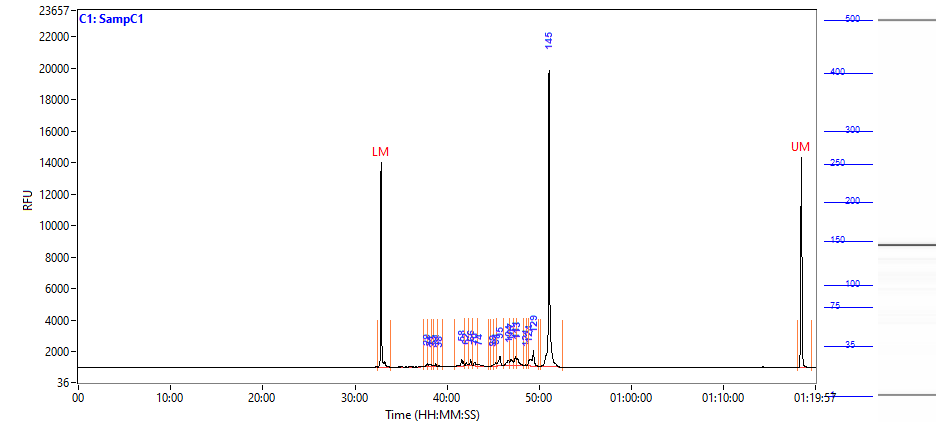

### 2021 10 11 14H 11M C2 SampC2.PNG

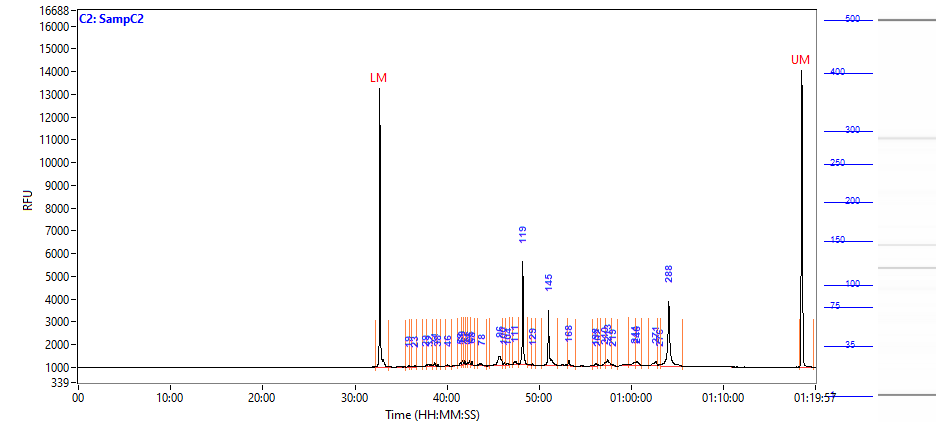

### 2021 10 11 14H 11M C3 SampC3.PNG

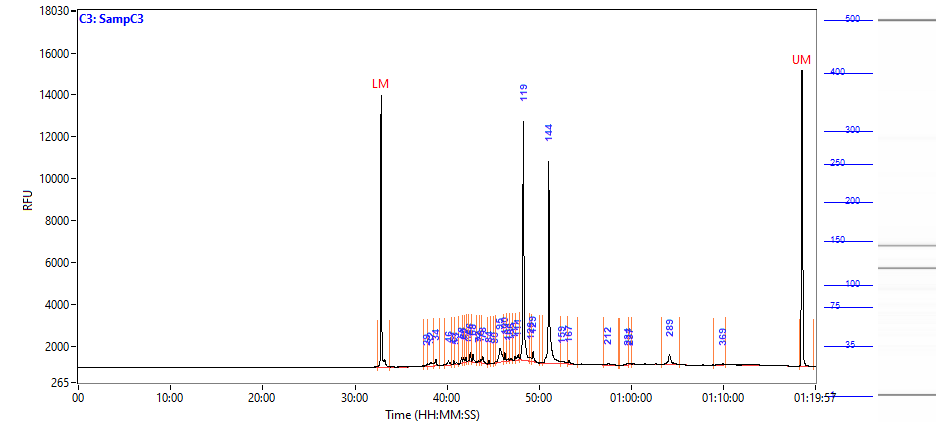

### 2021 10 11 14H 11M C4 SampC4.PNG

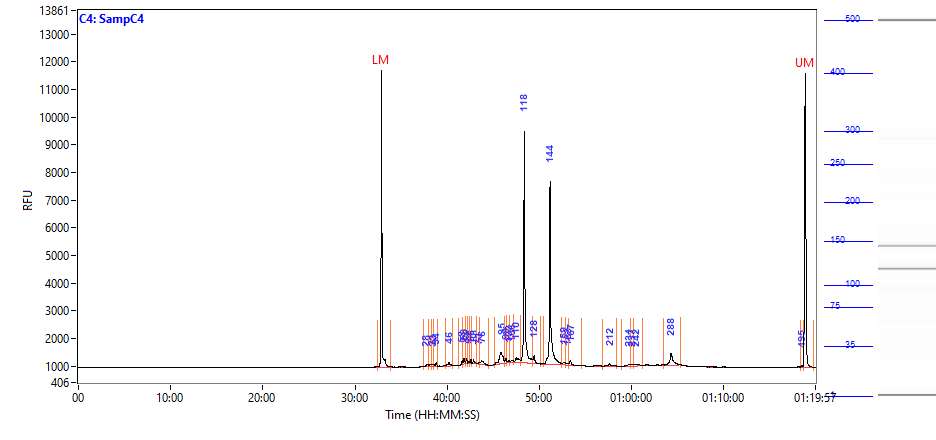

### 2021 10 11 14H 11M C5 SampC5.PNG

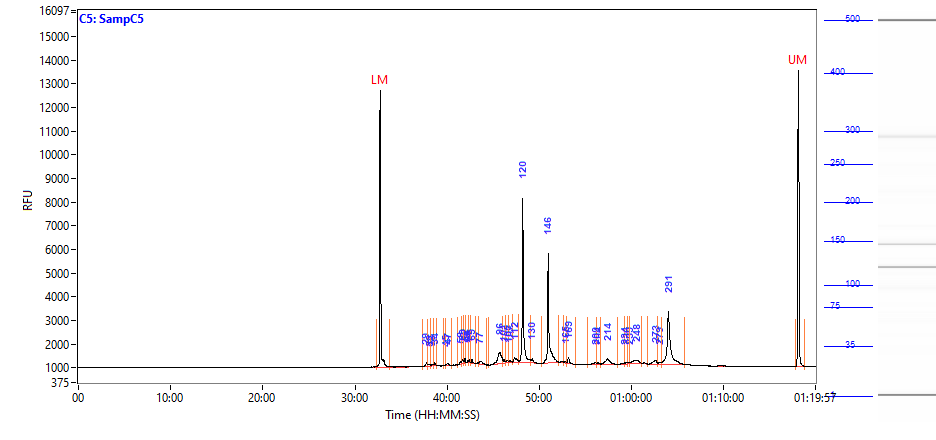

### 2021 10 11 14H 11M C6 SampC6.PNG

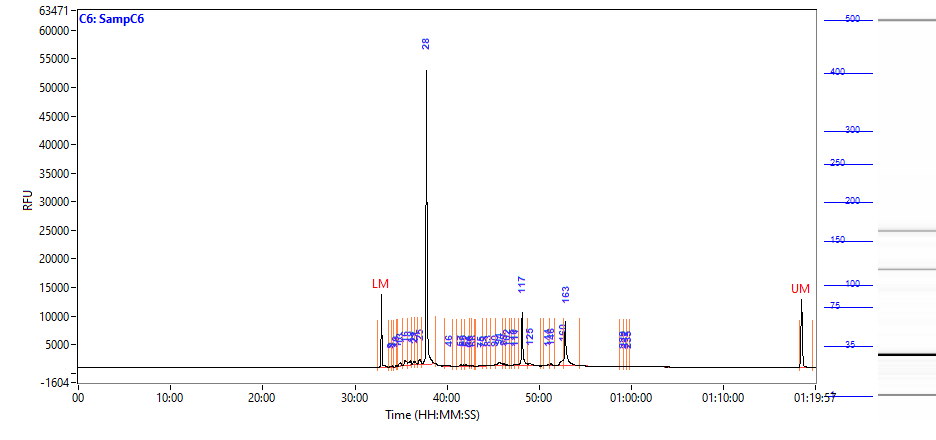

### 2021 10 11 14H 11M C7 SampC7.PNG

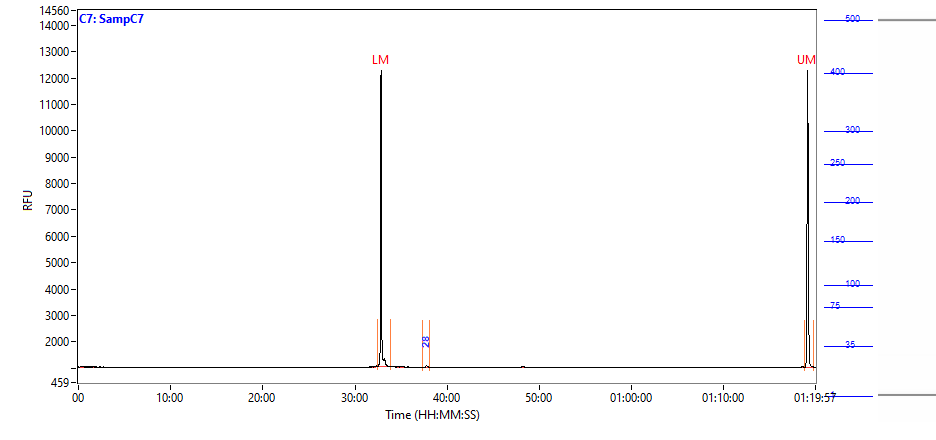

### 2021 10 11 14H 11M C8 SampC8.PNG

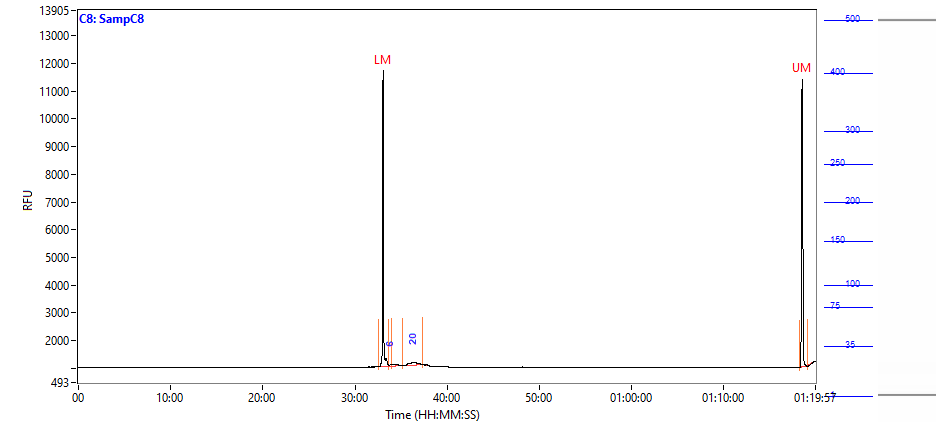

### 2021 10 11 14H 11M C9 SampC9.PNG

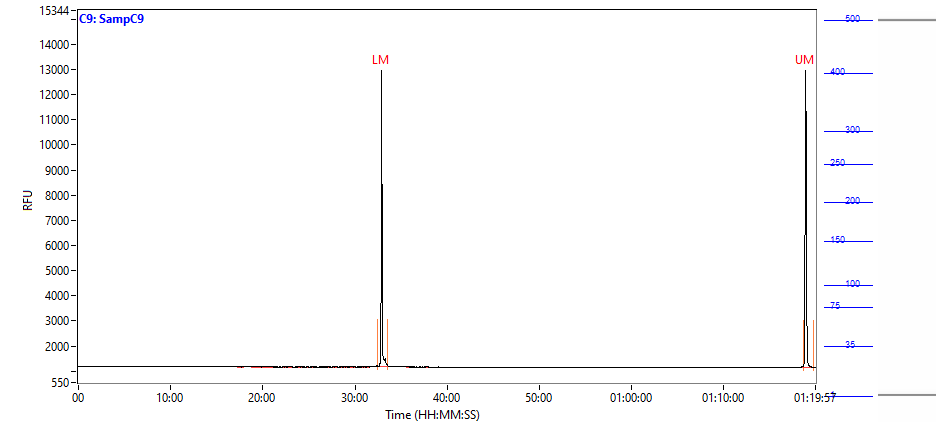

### 2021 10 11 14H 11M C10 SampC10.PNG

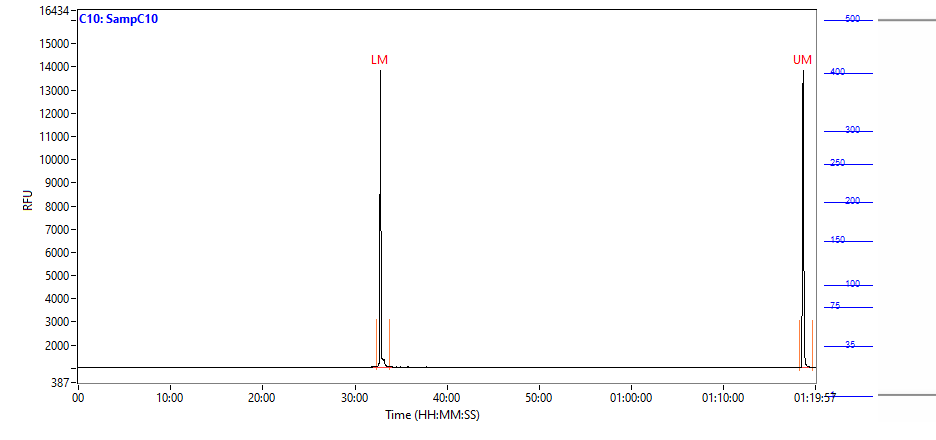

### 2021 10 11 14H 11M C12 SampC12.PNG

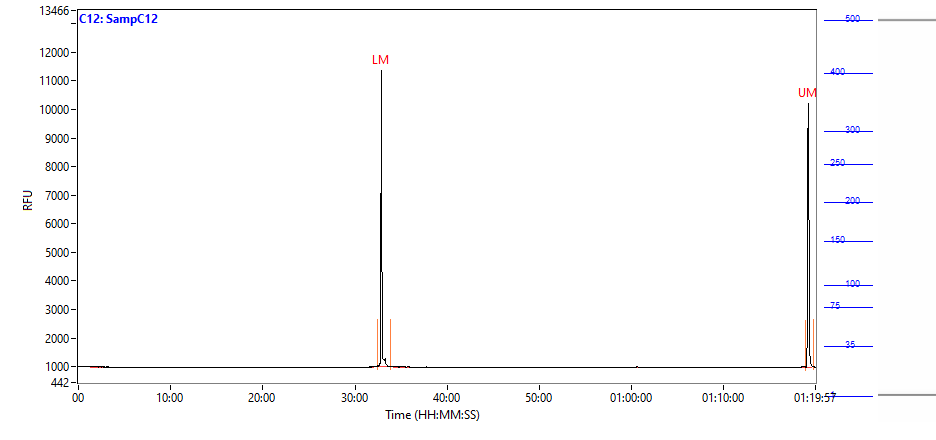

### 2021 10 11 14H 11M D1 SampD1.PNG

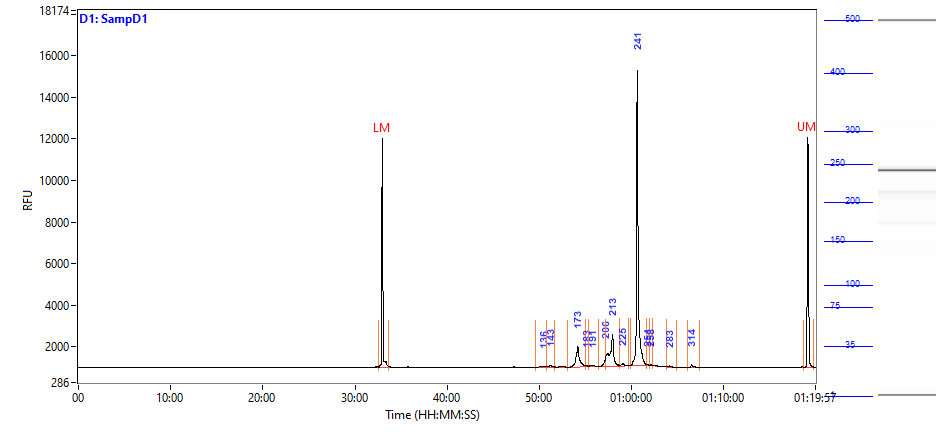

### 2021 10 11 14H 11M D2 SampD2.PNG

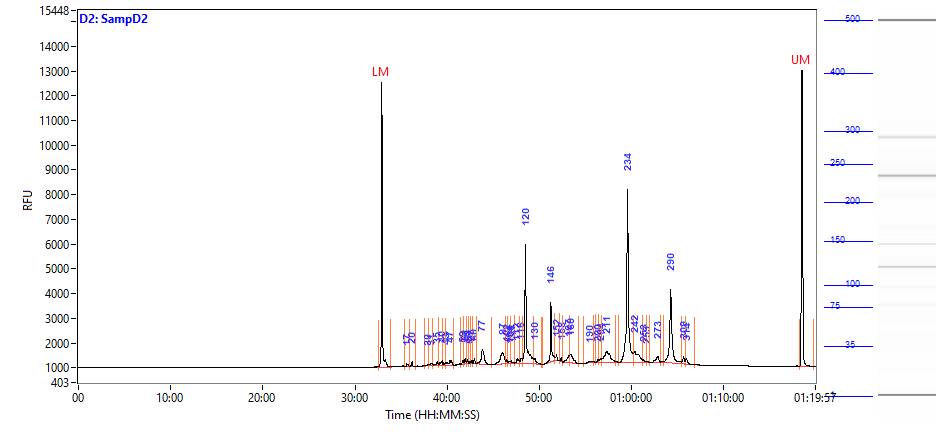

### 2021 10 11 14H 11M D3 SampD3.PNG

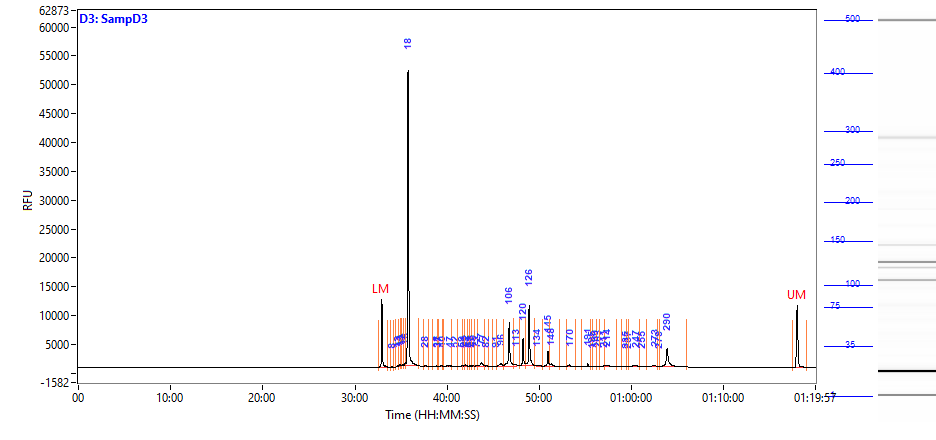

### 2021 10 11 14H 11M D4 SampD4.PNG

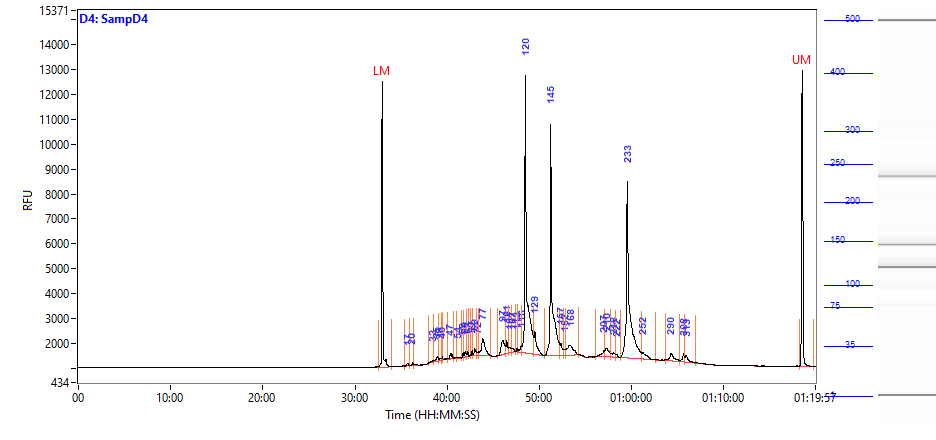

### 2021 10 11 14H 11M D5 SampD5.PNG

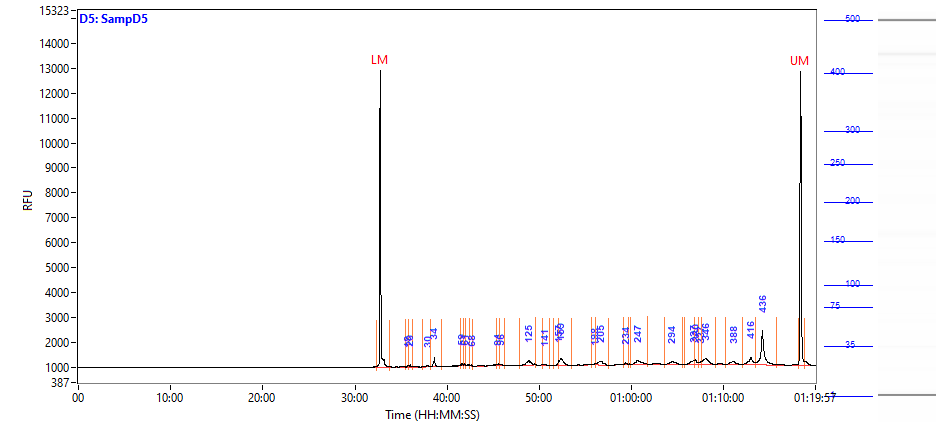

### 2021 10 11 14H 11M D6 SampD6.PNG

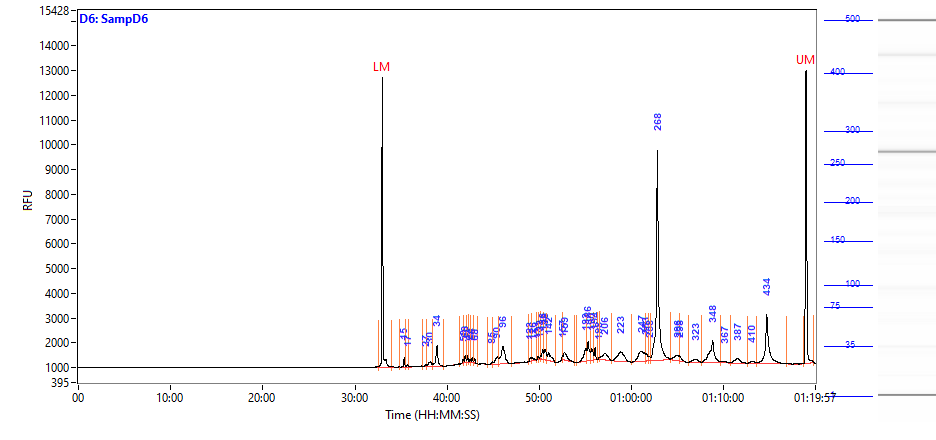

### 2021 10 11 14H 11M D7 SampD7.PNG

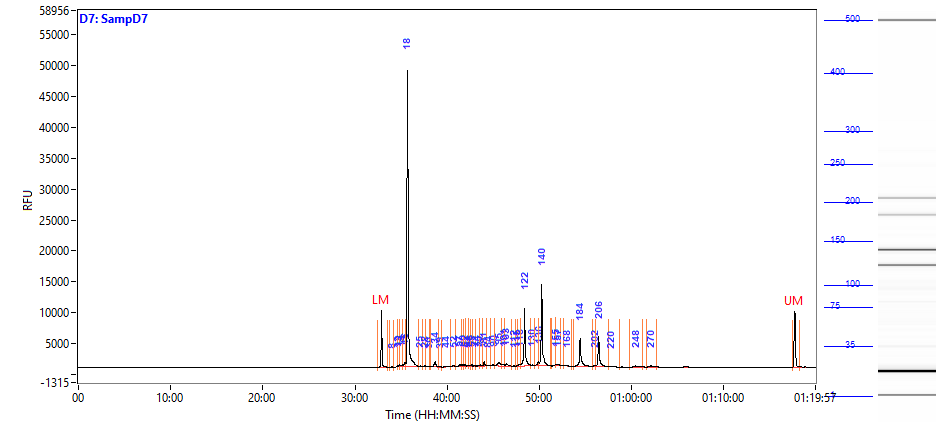

### 2021 10 11 14H 11M D8 SampD8.PNG

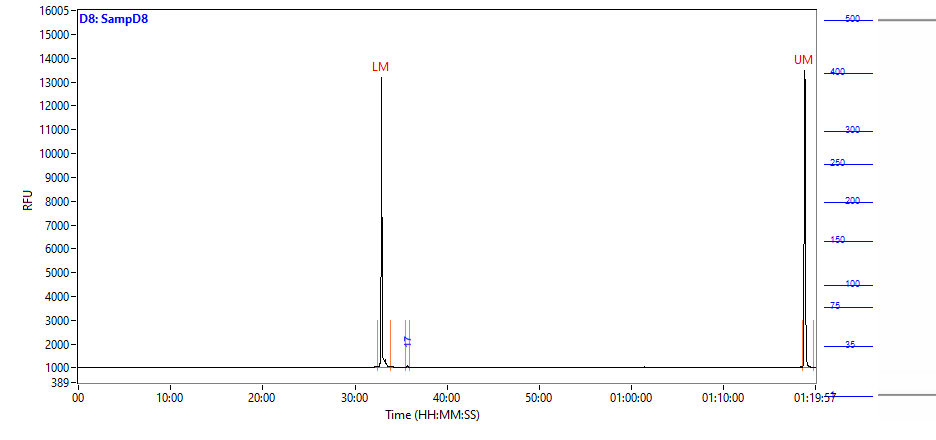

### 2021 10 11 14H 11M D9 SampD9.PNG

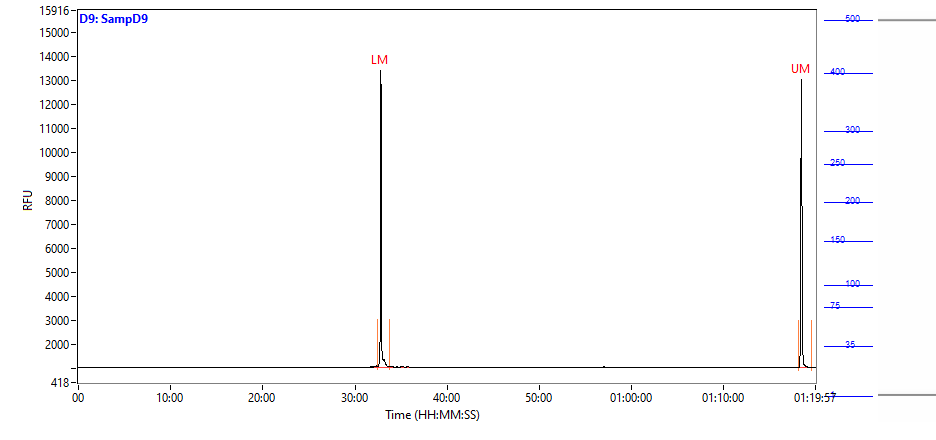

### 2021 10 11 14H 11M D10 SampD10.PNG

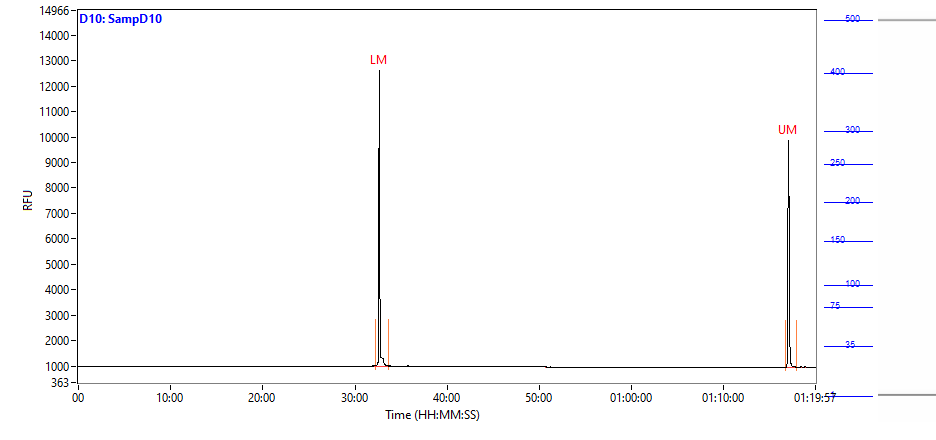

### 2021 10 11 14H 11M D11 SampD11.PNG

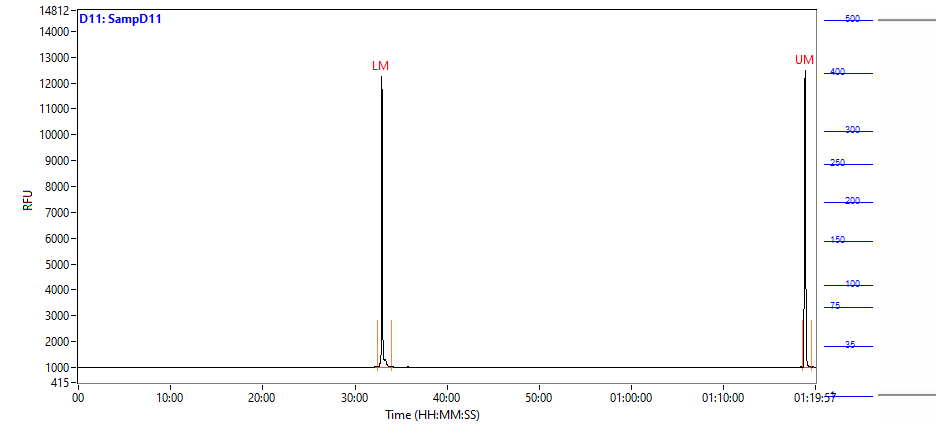

### 2021 10 11 14H 11M D12 SampD12.PNG

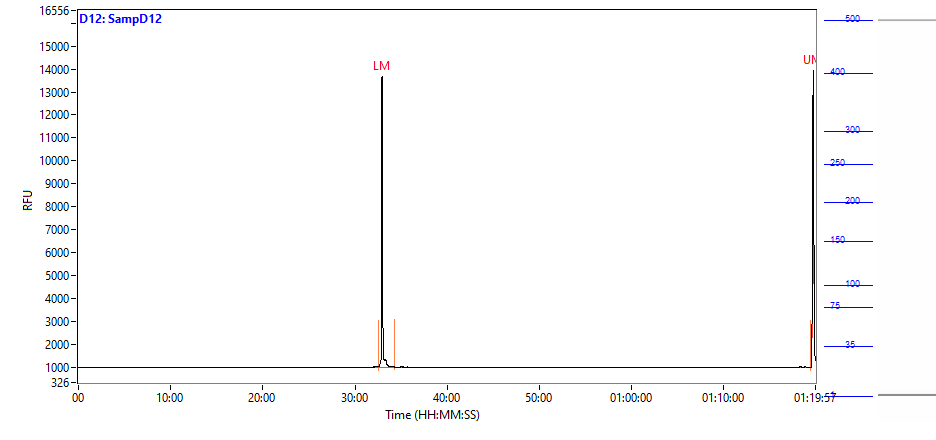

### 2021 10 11 14H 11M E1 SampE1.PNG

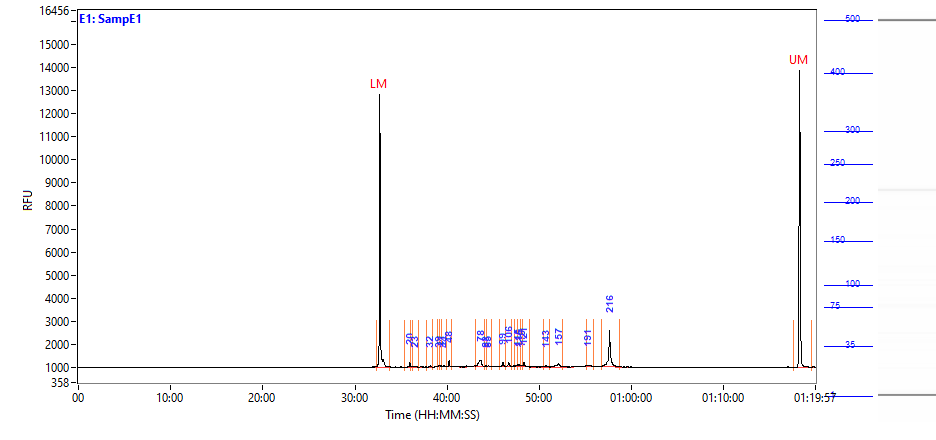

### 2021 10 11 14H 11M E2 SampE2.PNG

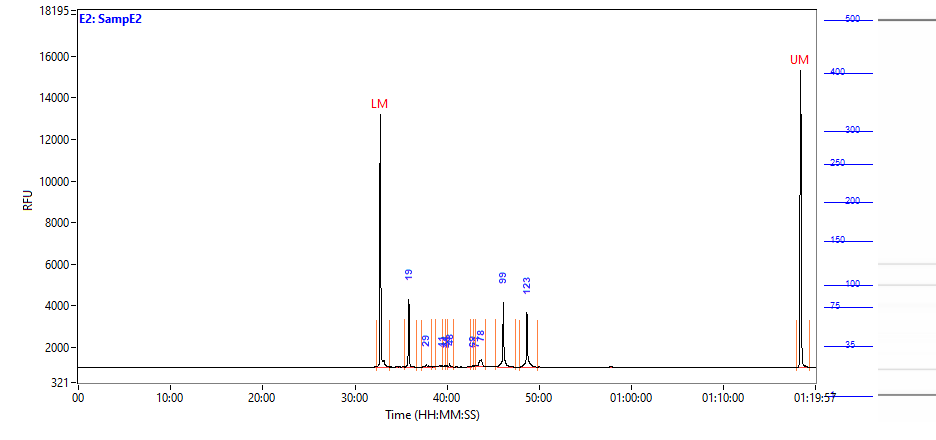

### 2021 10 11 14H 11M E3 SampE3.PNG

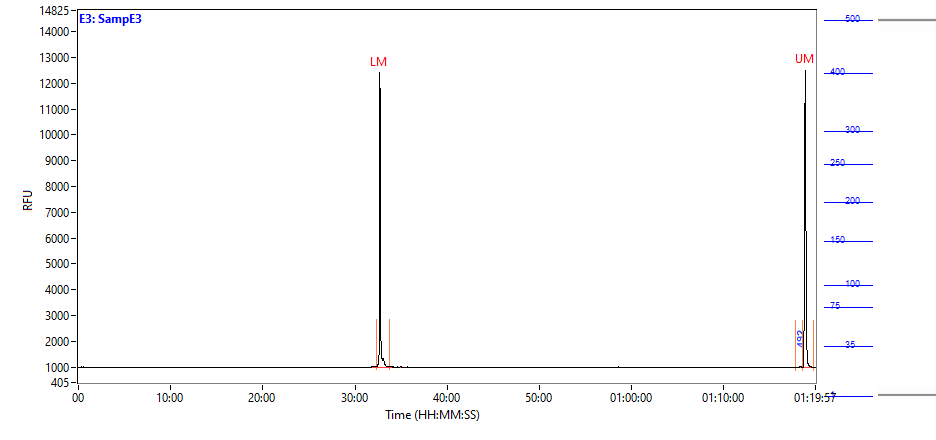

### 2021 10 11 14H 11M E4 SampE4.PNG

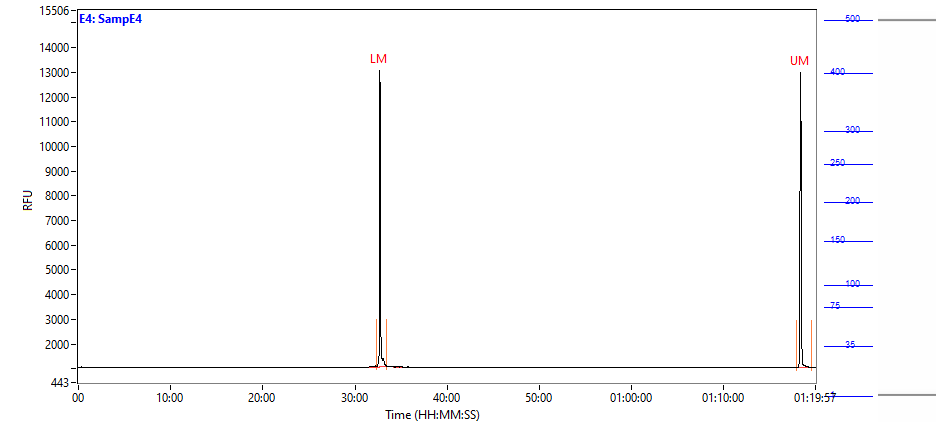

### 2021 10 11 14H 11M E5 SampE5.PNG

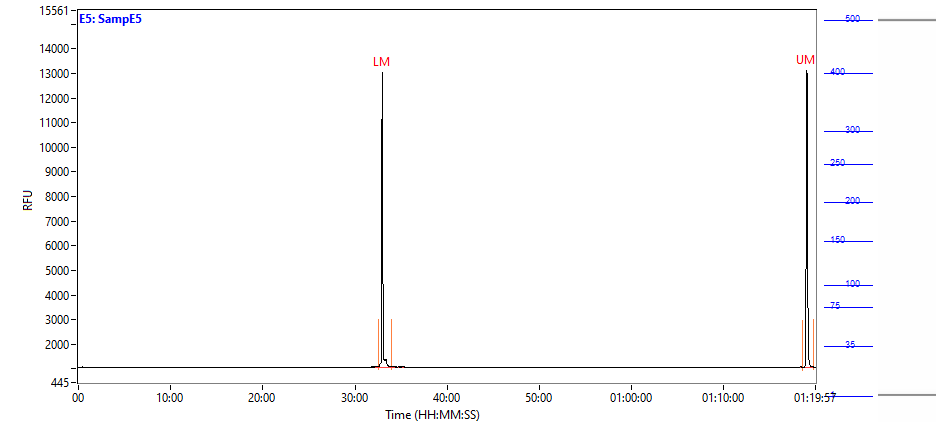

### 2021 10 11 14H 11M E6 SampE6.PNG

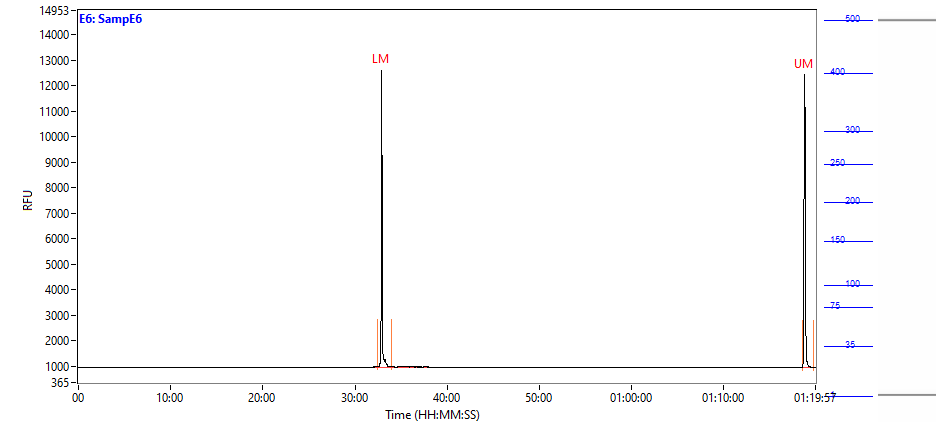

### 2021 10 11 14H 11M E7 SampE7.PNG

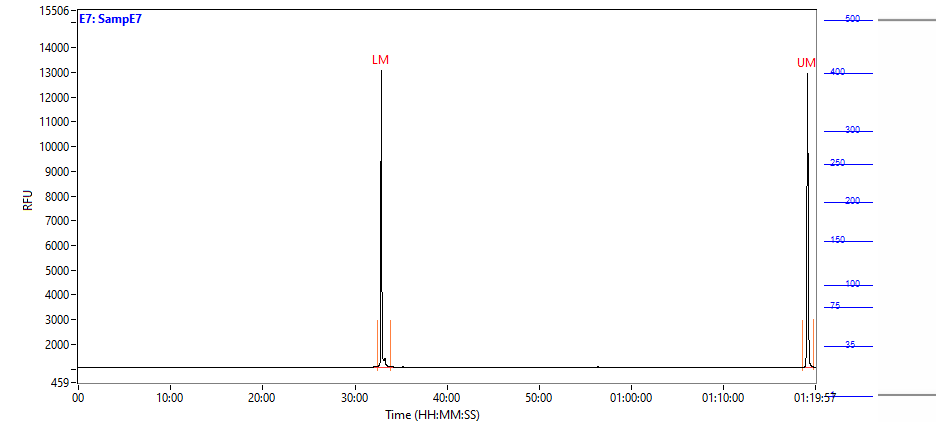

### 2021 10 11 14H 11M E8 SampE8.PNG

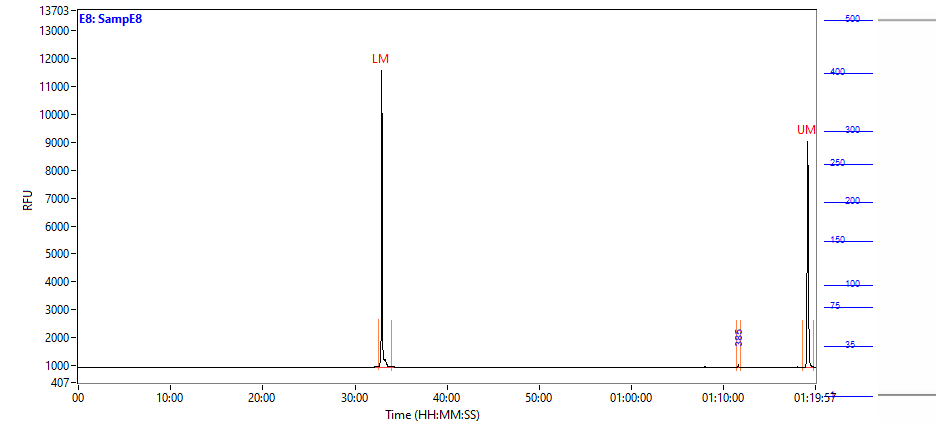

### 2021 10 11 14H 11M E9 SampE9.PNG

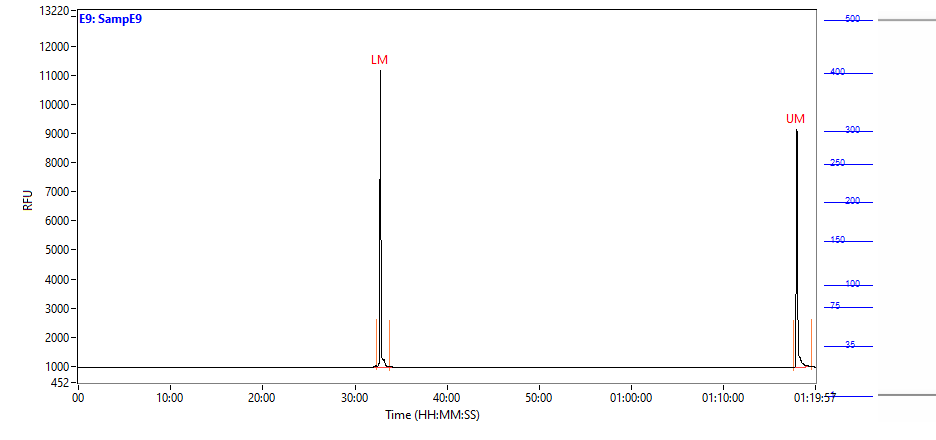

### 2021 10 11 14H 11M E10 SampE10.PNG

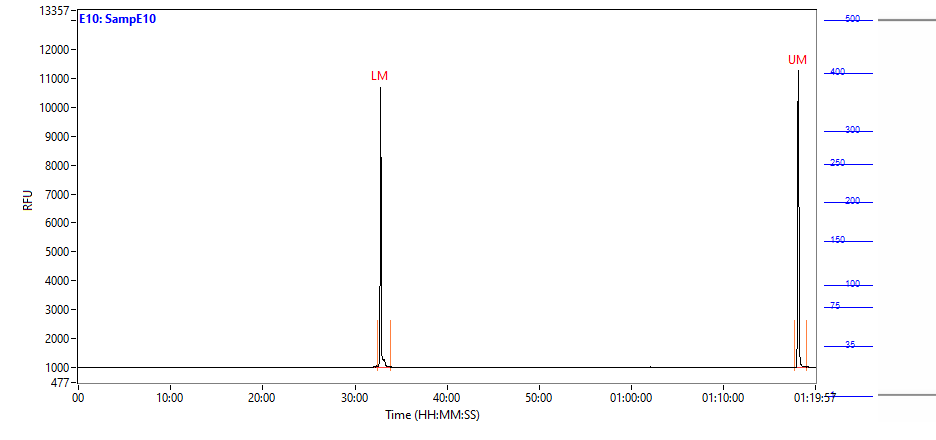

### 2021 10 11 14H 11M E12 SampE12.PNG

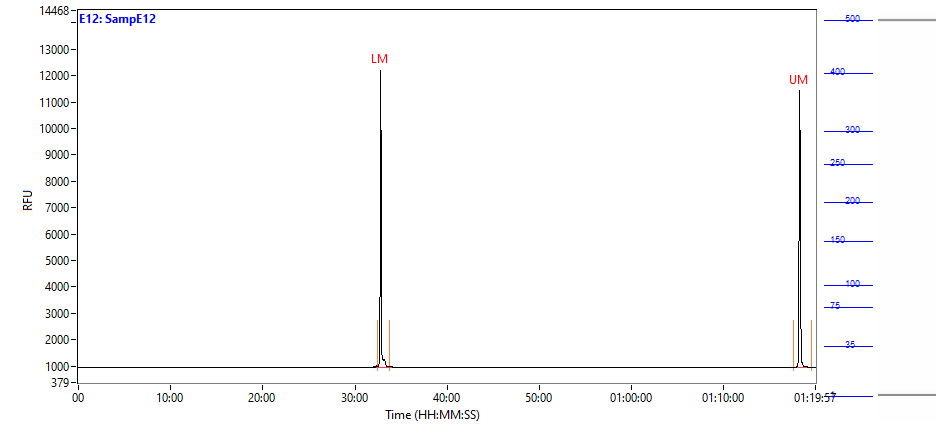

### 2021 10 11 14H 11M F1 SampF1.PNG

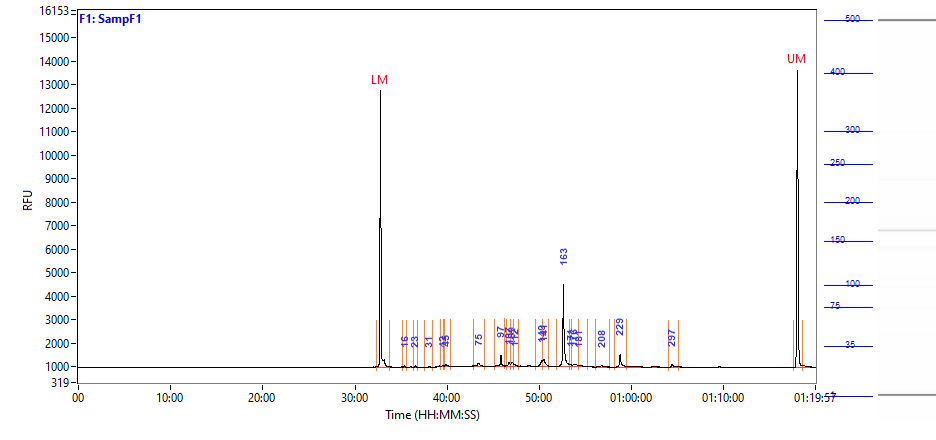

### 2021 10 11 14H 11M F2 SampF2.PNG

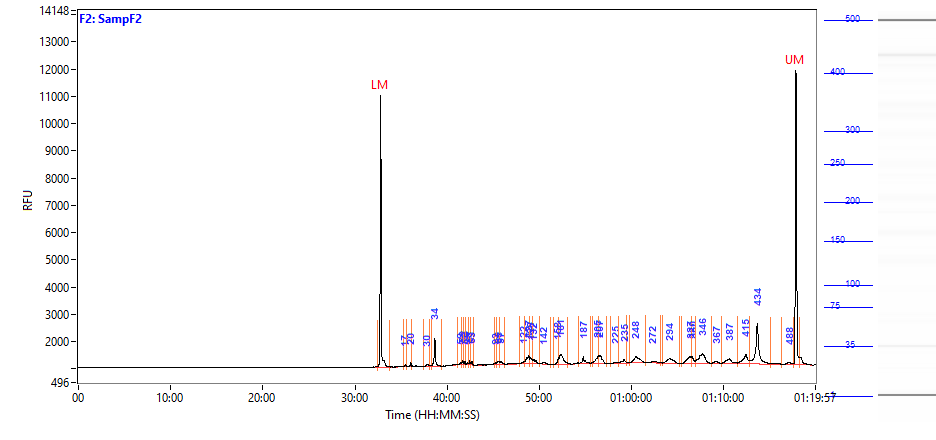

### 2021 10 11 14H 11M F3 SampF3.PNG

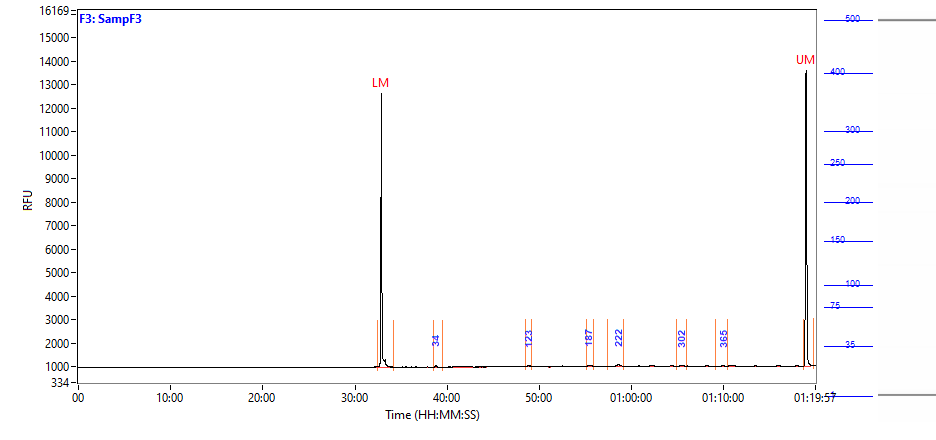

### 2021 10 11 14H 11M F4 SampF4.PNG

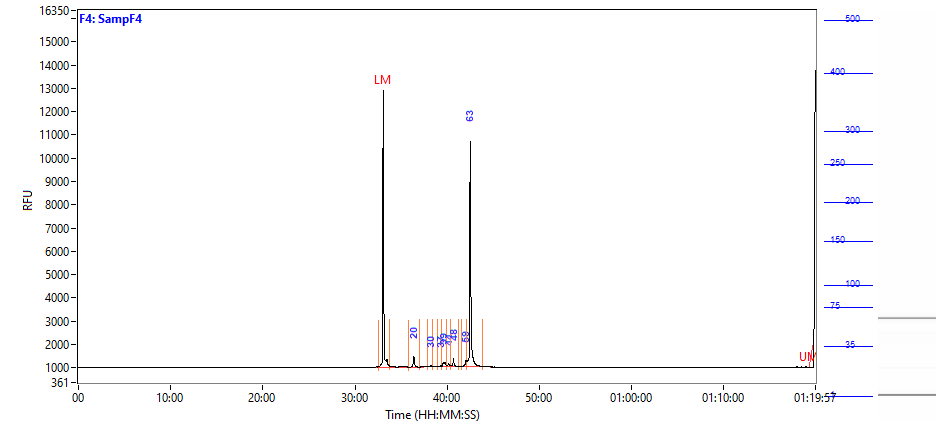

### 2021 10 11 14H 11M F5 SampF5.PNG

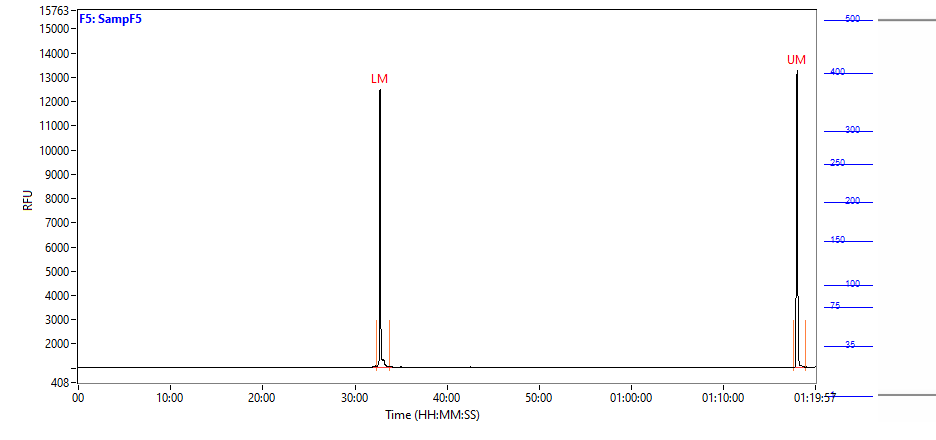

### 2021 10 11 14H 11M F6 SampF6.PNG

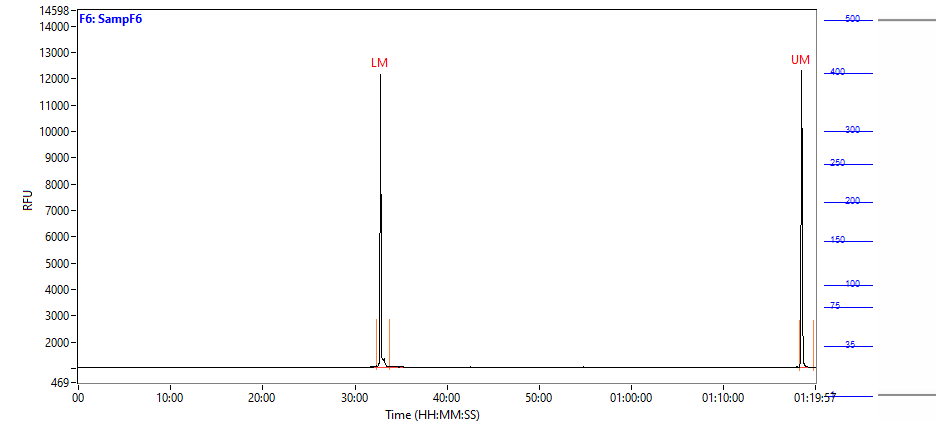

### 2021 10 11 14H 11M F7 SampF7.PNG

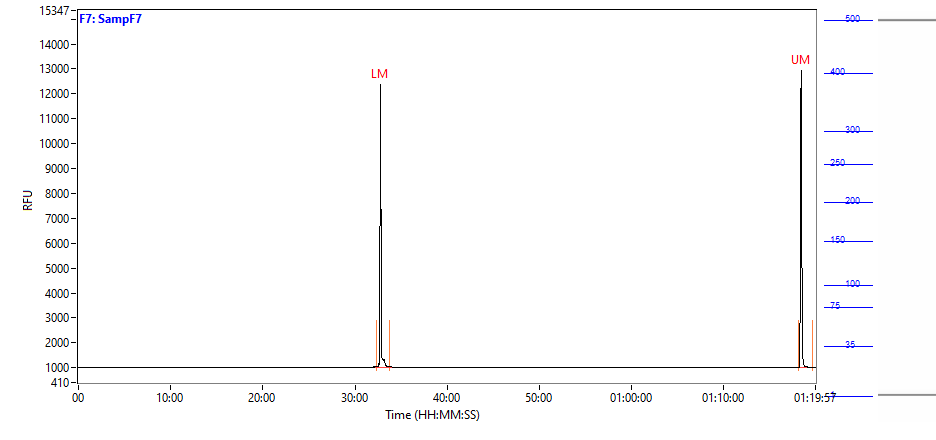

### 2021 10 11 14H 11M F8 SampF8.PNG

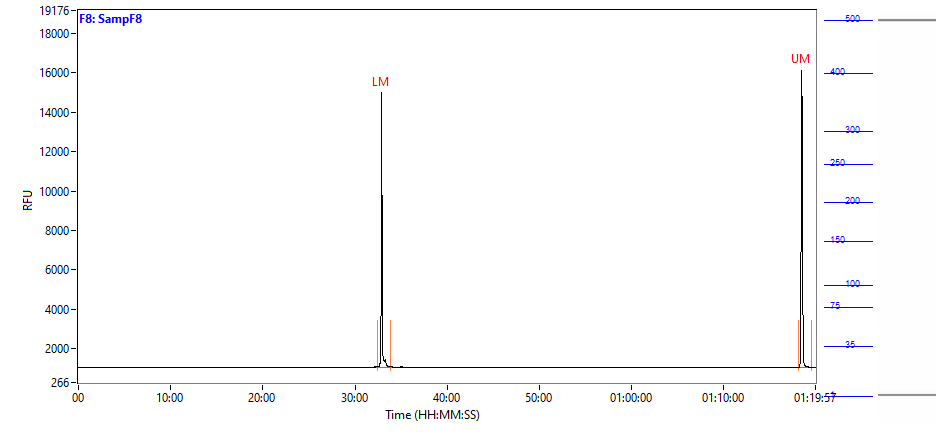

### 2021 10 11 14H 11M F9 SampF9.PNG

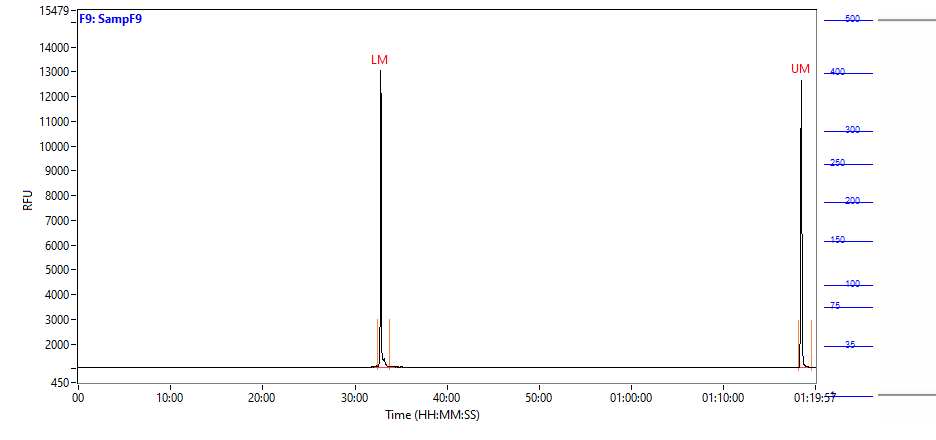

### 2021 10 11 14H 11M F10 SampF10.PNG

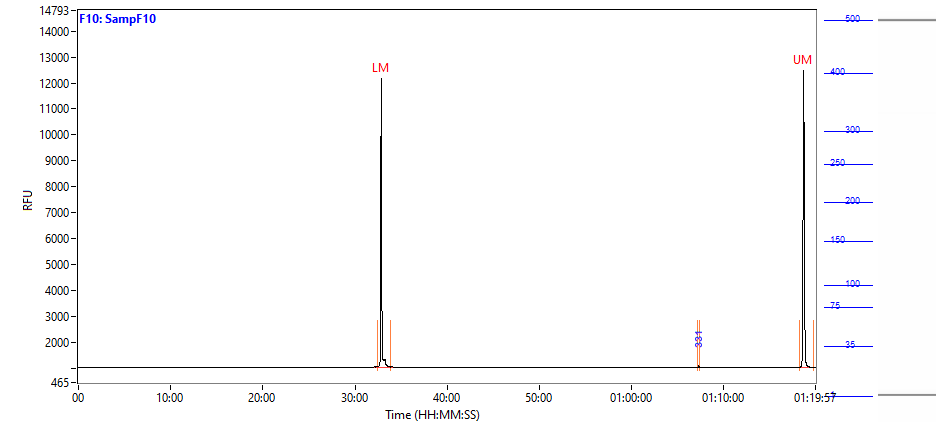

### 2021 10 11 14H 11M F12 SampF12.PNG

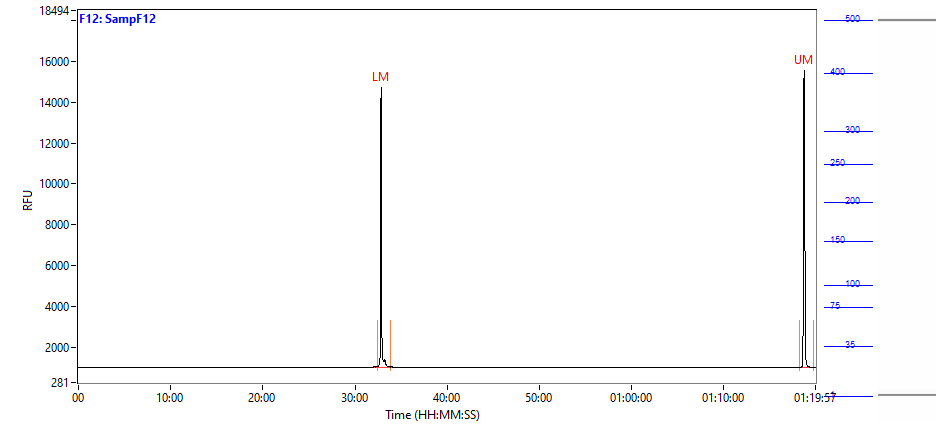

### 2021 10 11 14H 11M G1 SampG1.PNG

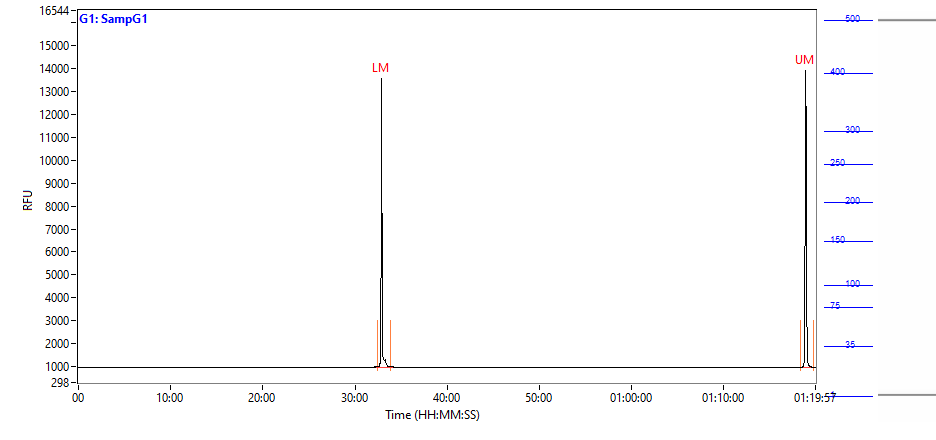

### 2021 10 11 14H 11M G2 SampG2.PNG

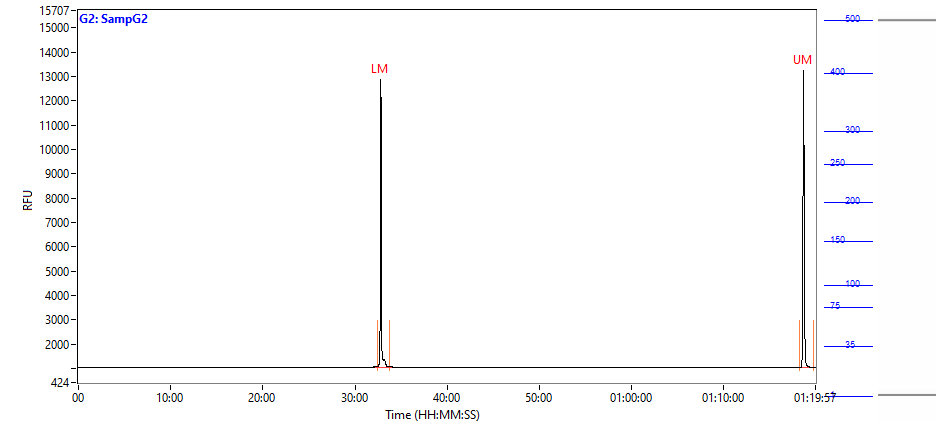

### 2021 10 11 14H 11M G3 SampG3.PNG

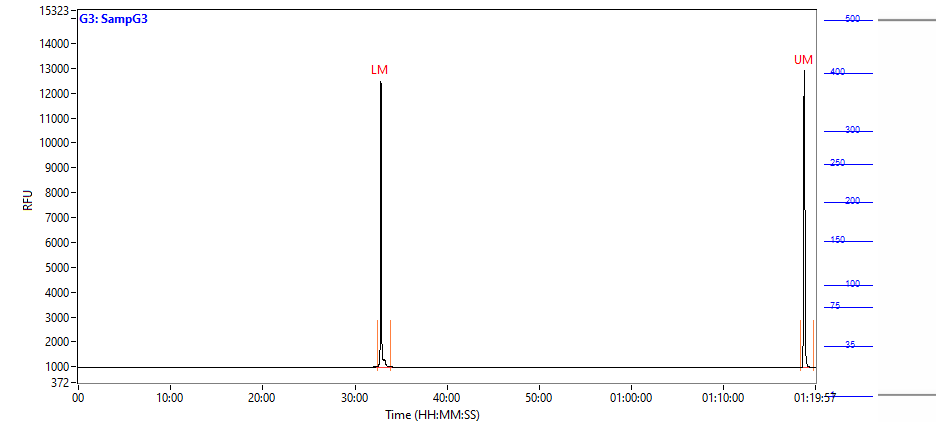

### 2021 10 11 14H 11M G4 SampG4.PNG

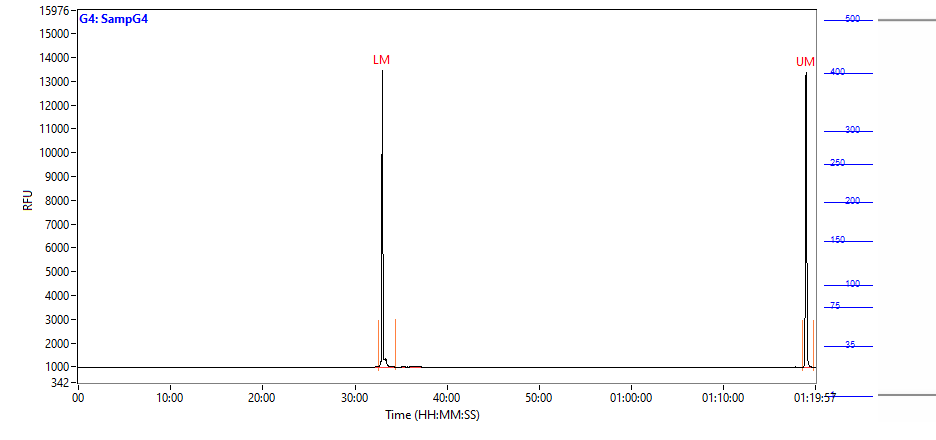

### 2021 10 11 14H 11M G5 SampG5.PNG

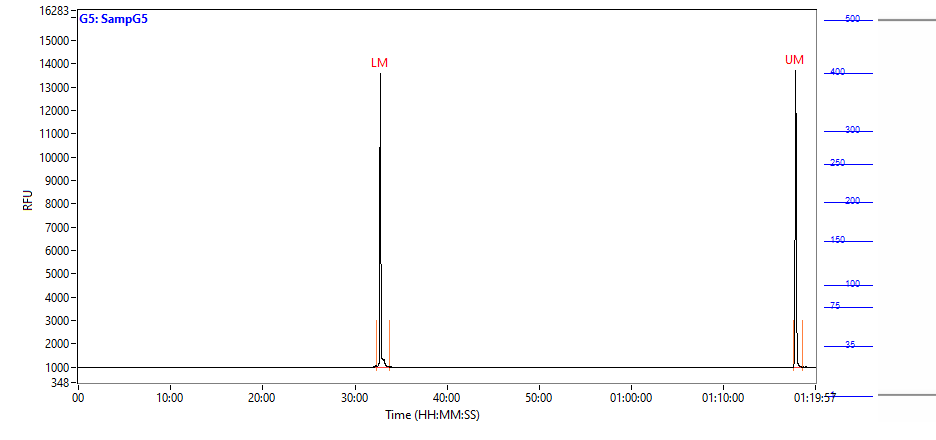

### 2021 10 11 14H 11M G6 SampG6.PNG

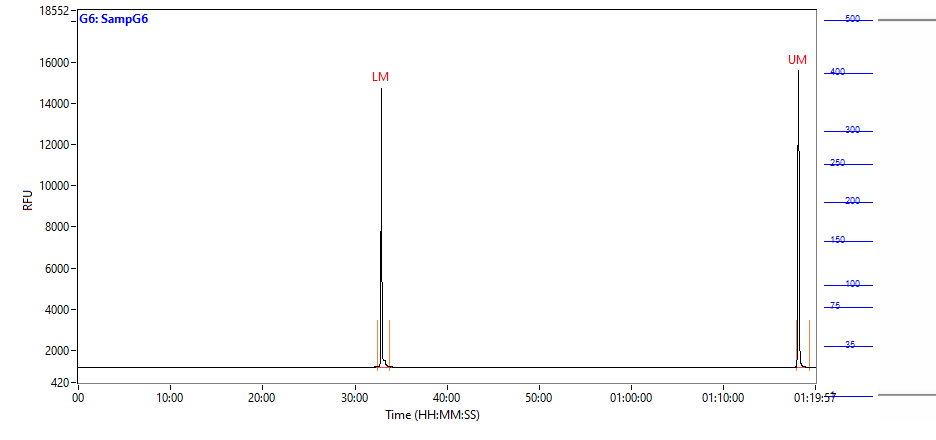

### 2021 10 11 14H 11M G7 SampG7.PNG

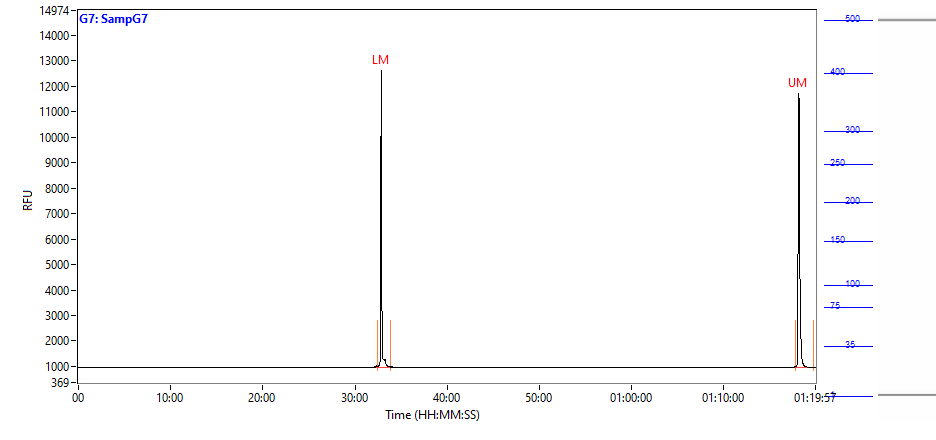

### 2021 10 11 14H 11M G9 SampG9.PNG

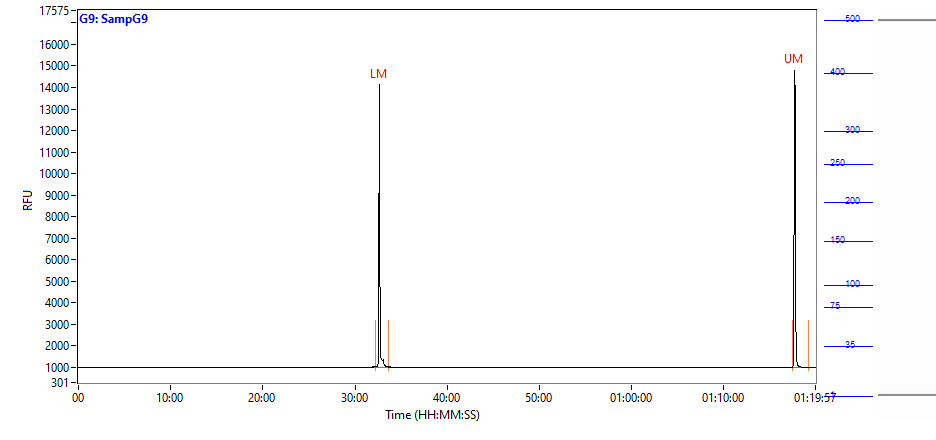

### 2021 10 11 14H 11M G10 SampG10.PNG

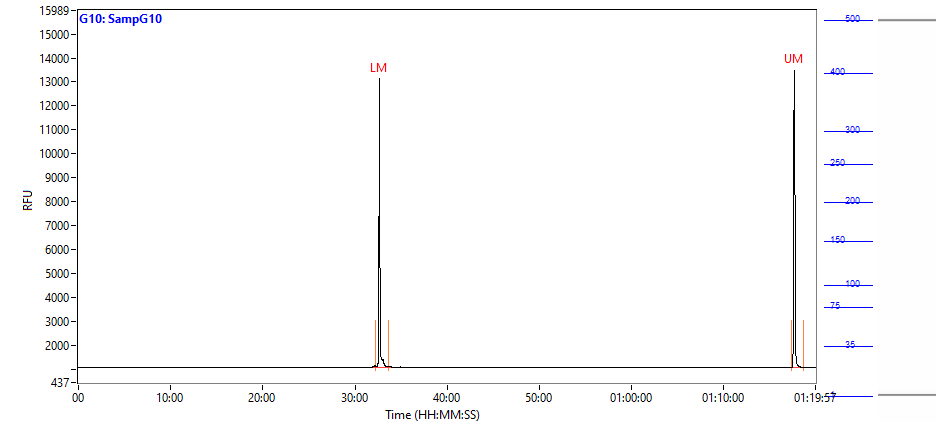

### 2021 10 11 14H 11M G11 SampG11.PNG

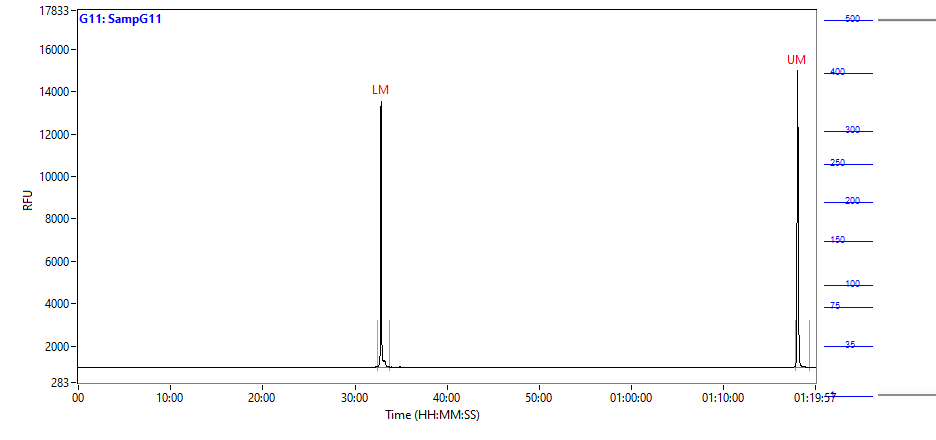

### 2021 10 11 14H 11M G12 SampG12.PNG

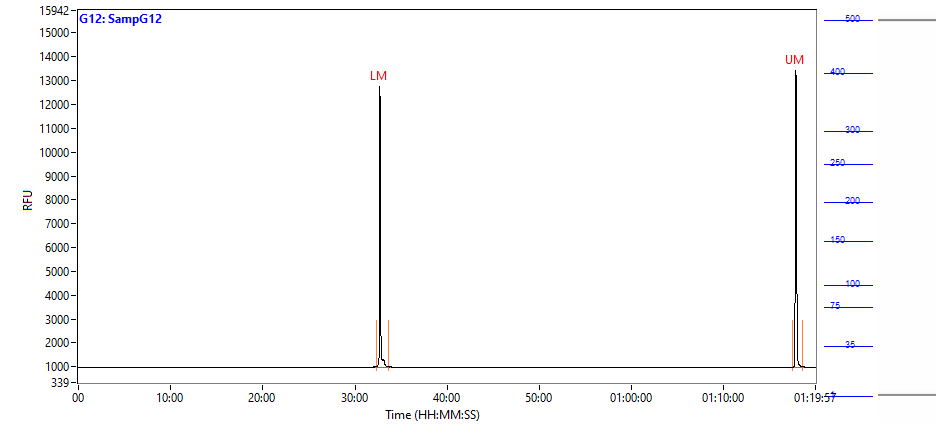

### 2021 10 11 14H 11M Gel.PNG

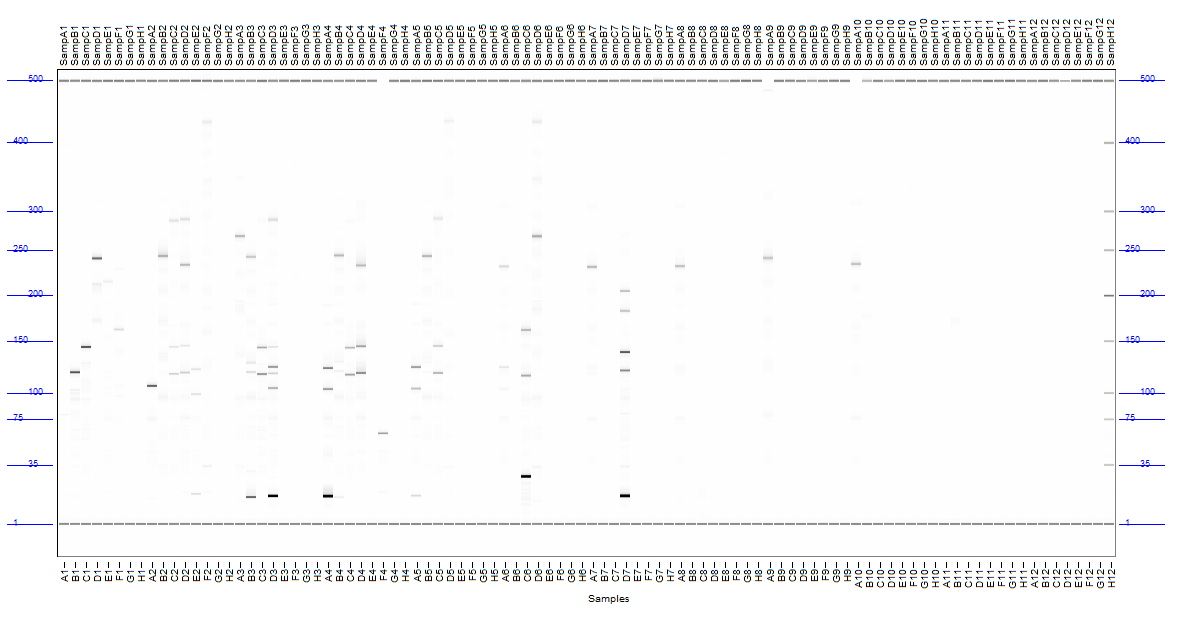

### 2021 10 11 14H 11M H1 SampH1.PNG

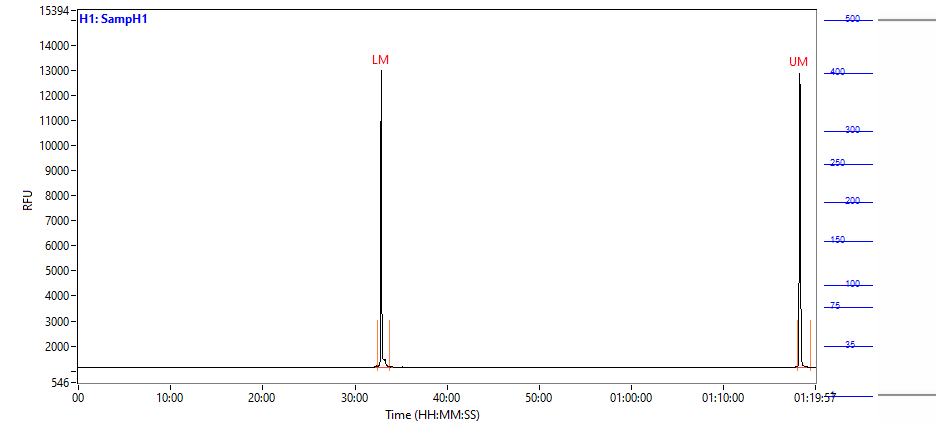

### 2021 10 11 14H 11M H2 SampH2.PNG

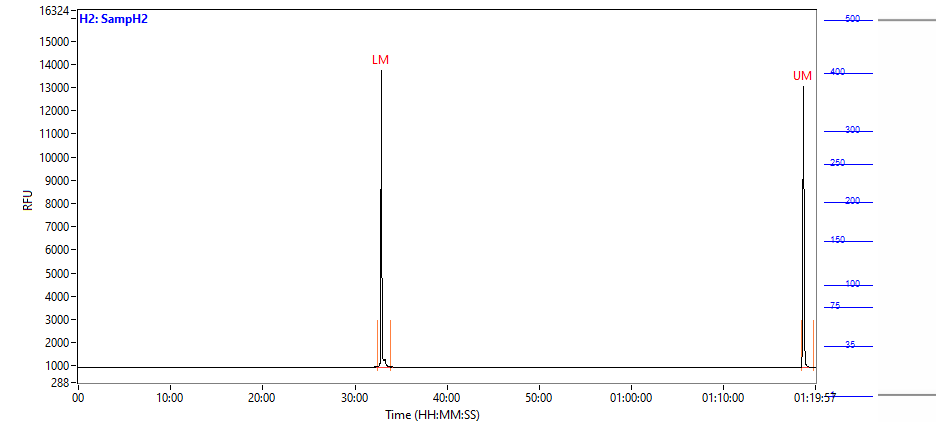

### 2021 10 11 14H 11M H3 SampH3.PNG

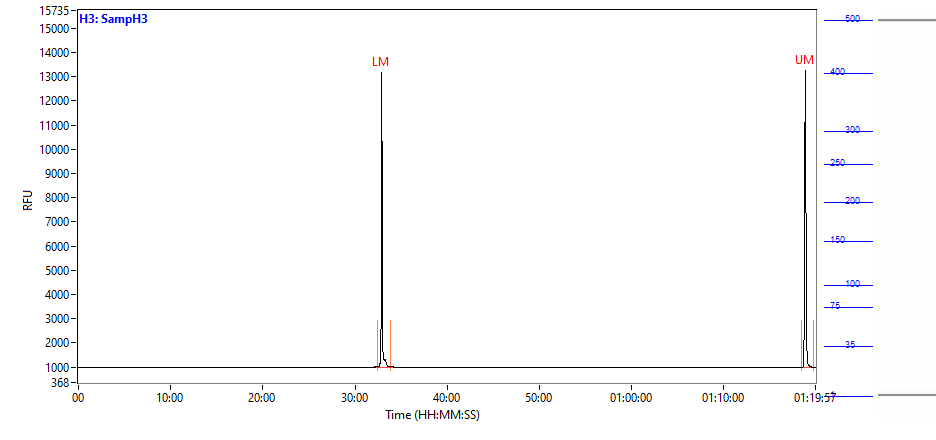

### 2021 10 11 14H 11M H4 SampH4.PNG

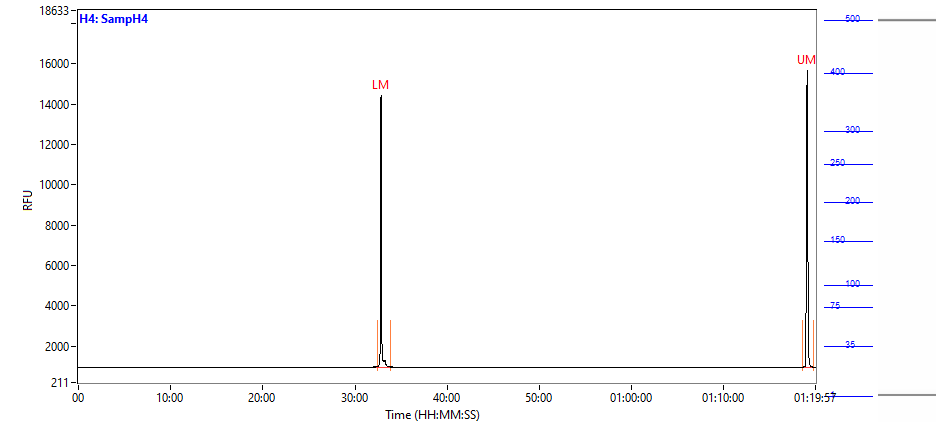

### 2021 10 11 14H 11M H5 SampH5.PNG

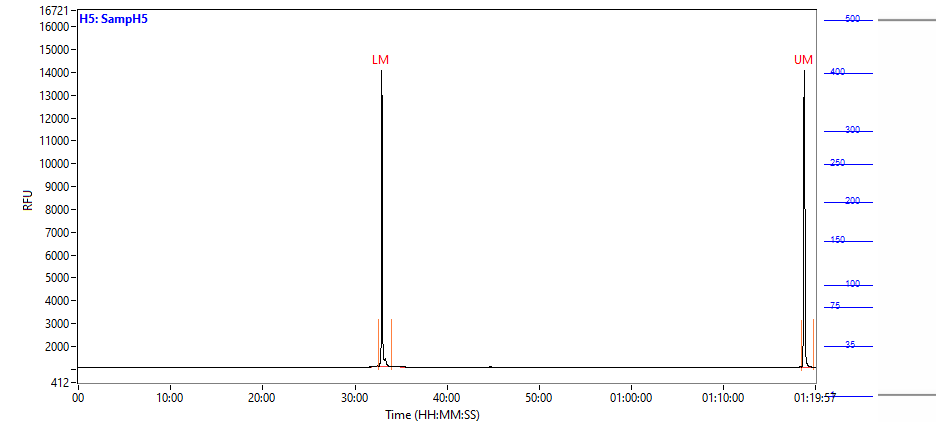

### 2021 10 11 14H 11M H6 SampH6.PNG

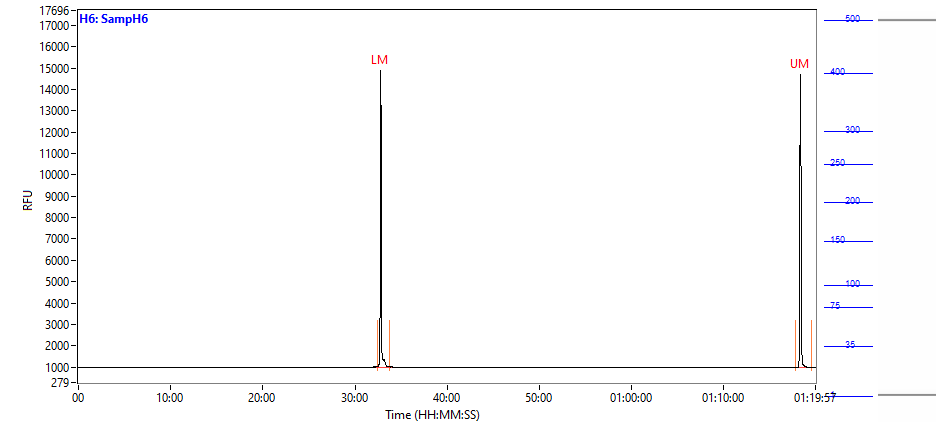

### 2021 10 11 14H 11M H7 SampH7.PNG

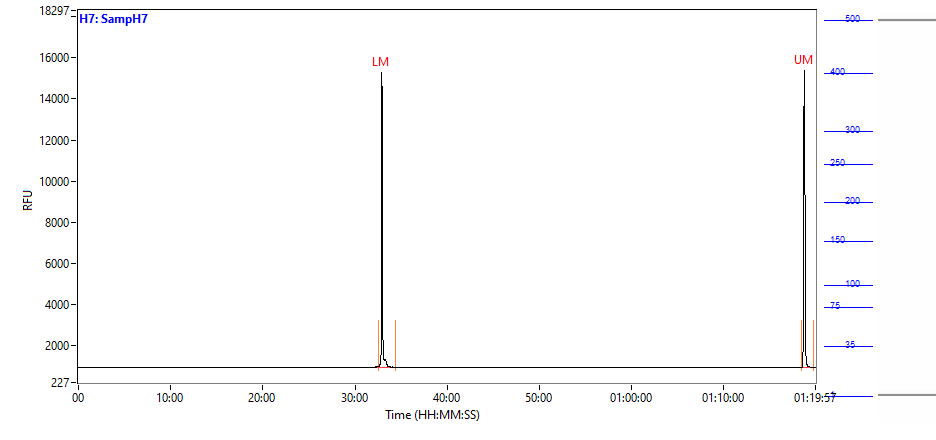

### 2021 10 11 14H 11M H8 SampH8.PNG

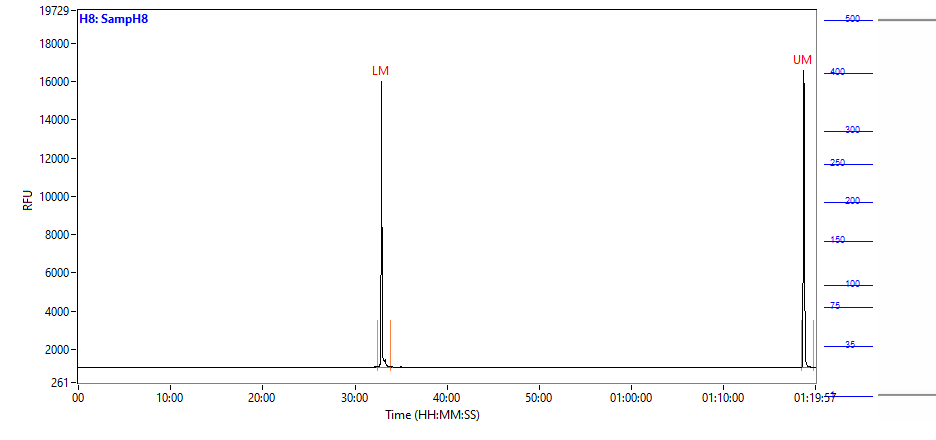

### 2021 10 11 14H 11M H9 SampH9.PNG

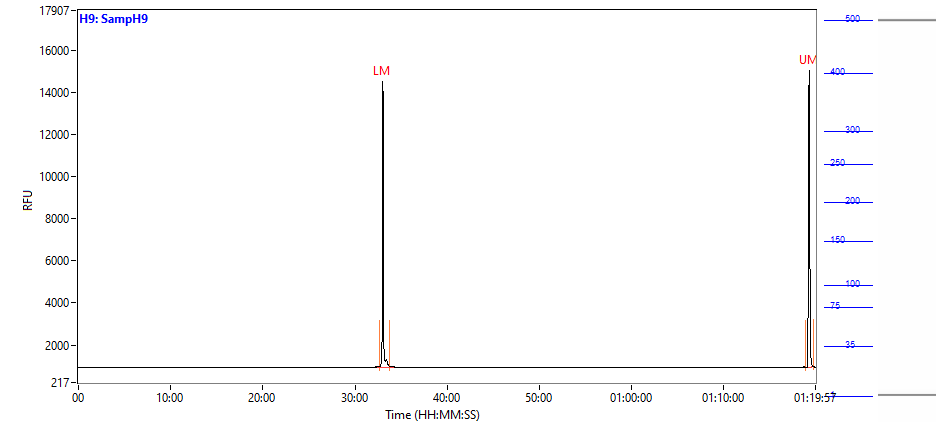

### 2021 10 11 14H 11M H10 SampH10.PNG

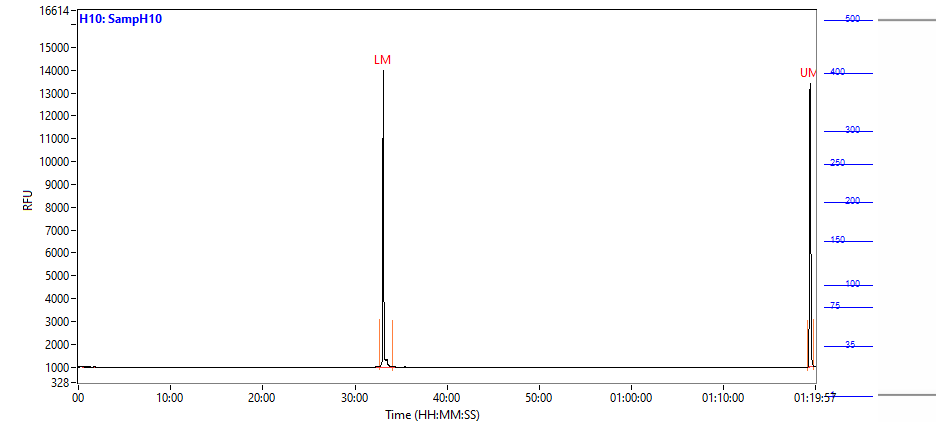

### 2021 10 11 14H 11M H11 SampH11.PNG

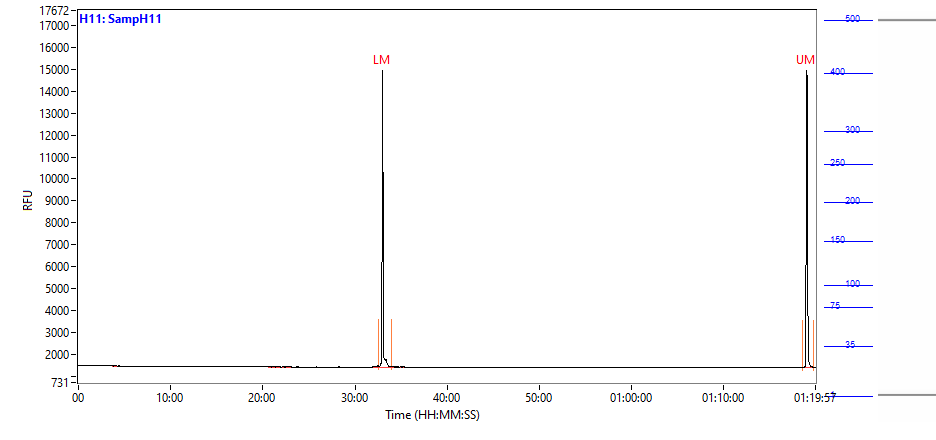

### 2021 10 11 14H 11M H12 SampH12.PNG

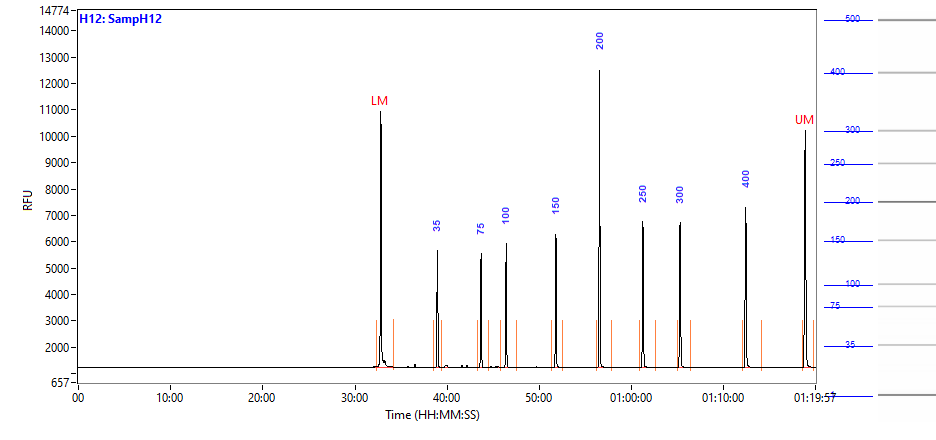

### CameraImage.bmp

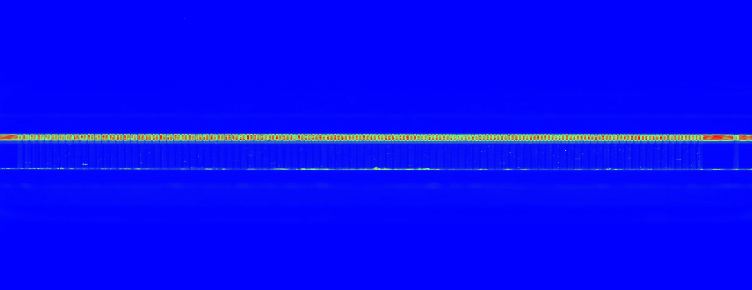
